# Large inversion polymorphisms are widespread in North American songbirds

**DOI:** 10.1101/2025.08.07.669111

**Authors:** Teresa M. Pegan, Benjamin M. Winger

## Abstract

The prevalence and evolutionary importance of inversion polymorphisms in natural populations is poorly known because of limited genome-wide sequence data availability for most species. Inversion studies in wild populations usually target rare cases of major trait polymorphisms or local adaptation whose genetic basis involves inversions, creating a strong impression that inversions in nature are generally maintained by natural selection through links to ecologically relevant phenotypes. By contrast, genome-wide studies in humans and model organisms suggest that inversion polymorphisms are common, subject to highly complex evolutionary processes, and generally difficult to link with clearly observable cases of phenotypic variation. Using a large comparative population genomic dataset generated from 35 codistributed species of birds, we tested the hypothesis that inversions are common even within populations that lack known phenotypic polymorphisms. We leveraged analytical methods suitable for low-coverage whole genome sequencing to reveal evidence for over 170 putative inversion polymorphisms within 28 species. We find that many polymorphisms are large and present at balanced frequencies, and some are shared across species boundaries. Yet, most polymorphisms do not deviate significantly from Hardy-Weinberg Equilibrium, raising the possibility that many of these massive haploblocks could be segregating neutrally. Our results thereby reveal evidence that inversions show a variety of complex yet largely hidden patterns in natural populations, beyond cases where they contribute to known variation in ecologically relevant traits.

**Significance:** Inversions are DNA segments that evolve as tightly linked blocks, predisposing them to contribute to phenotypic variation and local adaptation. Studies of inversions in natural populations of non-model species usually involve rare cases where notable trait polymorphisms are controlled by inversions. But how common are inversion polymorphisms that do not mediate known trait variation? We generated population genomic data from 35 codistributed species and show that large inversions are common in passerine birds, despite apparent absence of phenotypic variation and local adaptation in our study populations. Some inversions show patterns suggesting complex evolutionary scenarios, such as balancing selection and shared polymorphism across species, while others may be neutral. Our study reveals that inversions commonly persist in natural populations even without obvious phenotypic variation.

## Introduction

Inversions are genomic structural variants that can play a significant role in phenotypic evolution, but their prevalence and evolutionary dynamics remain poorly understood in most species (Wellenreuther and Bernatchez 2018; Connallon and Olito 2022). Inversions facilitate special cases of phenotypic evolution because recombination between inverted and non-inverted haplotypes is suppressed, allowing genetic variants with phenotypic effects to evolve together in tightly linked haplotypes (reviewed in (Wellenreuther and Bernatchez 2018; Berdan et al. 2023). Many studies in wild populations have found that inversion polymorphisms contribute to notable phenotypic polymorphisms (Nishikawa et al. 2015; Küpper et al. 2016; Lamichhaney et al. 2016; Tuttle et al. 2016; Jay et al. 2018; Kay et al. 2022), local adaptation (Ayala et al. 2013; Huang et al. 2020; Akopyan et al. 2022; Harringmeyer and Hoekstra 2022; Battlay et al. 2023) and even speciation (Feder et al. 2003; Fuller et al. 2019; Hooper et al. 2019; Knief et al. 2024). However, such cases may not represent the general evolutionary dynamics of inversions in populations (Mérot 2020; Mérot et al. 2020; Hirabayashi and Owens 2023). In model organisms and humans, for which extensive whole genome population sampling has been available for longer than non-model organisms, genome-wide surveys have found that inversion polymorphisms are common but rarely associated with easily-observable phenotypes (Corbett-Detig and Hartl 2012; Catacchio et al. 2018; Giner-Delgado et al. 2019; Kapun and Flatt 2019; Charlesworth 2023). Theoretical work and analyses with model organisms further demonstrate that effects of inversions on fitness can be highly complex and often not linked to discrete phenotypic polymorphisms (Kapun and Flatt 2019; Berdan et al. 2023; Pei et al. 2023). To gain a fuller picture of the role of inversion polymorphisms in the evolution of non-model organisms, it is important to evaluate their prevalence in genome-wide surveys across many focal taxa, rather than focusing only on polymorphisms with known or suspected phenotypic consequences (Harringmeyer and Hoekstra 2022; Hirabayashi and Owens 2023). Yet, only recently has it become feasible to perform whole genome sequencing across many individuals of many species to afford comparative investigation of inversion prevalence in non-model species. Here, we leverage a large multi-species population genomic dataset (1660 individuals) to assess evidence for autosomal inversion polymorphisms within 35 species of North American passerine songbirds and woodpeckers that breed sympatrically across the boreal ecoregion (Fig. 1A, Table 1), without *a priori* evidence of phenotypic polymorphisms.

**Figure 1.**
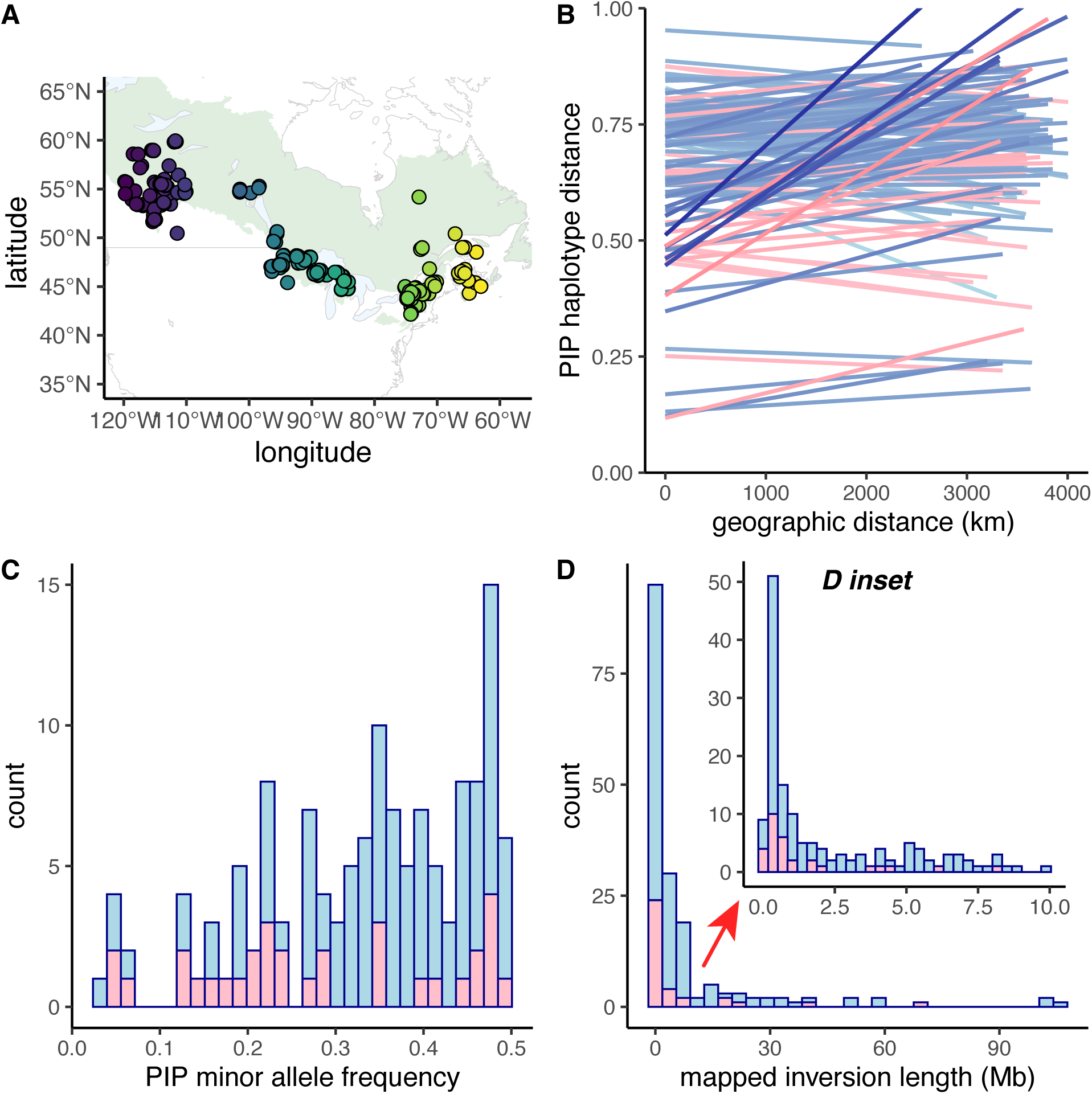
Summary of population genetic sampling and putative inversion polymorphism (PIP) patterns in our dataset. In panels **B** and **C**, colors represent whether each PIP’s status as an inversion was supported by linkage disequilibrium analyses (blue) or whether these results were inconclusive (pink). **A.** Sampling locations colored by longitude for all 35 species in this study from the North American boreal ecoregion (shown in pale green; (Omernik and Griffith 2014). Each species was sampled as broadly and evenly from this region as possible. Fig 1A is adapted from (Pegan et al. 2025). **B.** Slopes of isolation-by-distance (IBD) are shown for each biallelic PIP (for which individuals can be assigned to homozygous or heterozygous genotypes). Lines represent the slope and intercept of linear models of the relationship between PIP haplotype distance and geographic distance among pairs of conspecific individuals. A strongly positive IBD slope represents cases where pairs of individuals close to each other in geographic space tend to have more similar PIP genotypes than those distant from each other, indicating spatial structure in the distribution of PIP haplotypes. The darkness of each line corresponds to the IBD slope to help visually highlight the strongest slopes, and slopes are plotted in order such that the strongest slopes are shown at the front. With a few exceptions (Table 2), most PIPs show relatively flat slopes indicating lack of spatial sorting of PIP haplotypes. **C.** Histogram of minor haplotype (allele) frequency for biallelic PIPs. The distribution is skewed to the right, indicating that most PIP haplotypes exist at relatively balanced frequencies. **D.** Histograms of the mapped length of each PIP in the dataset. The inset shows a more detailed view of the left side of the plot (PIPs < 10 Mb in mapped length). Most identified PIPs are larger than 1 Mb in mapped length, and many are larger than 10 Mb (Dataset S1).

**Table 1.**
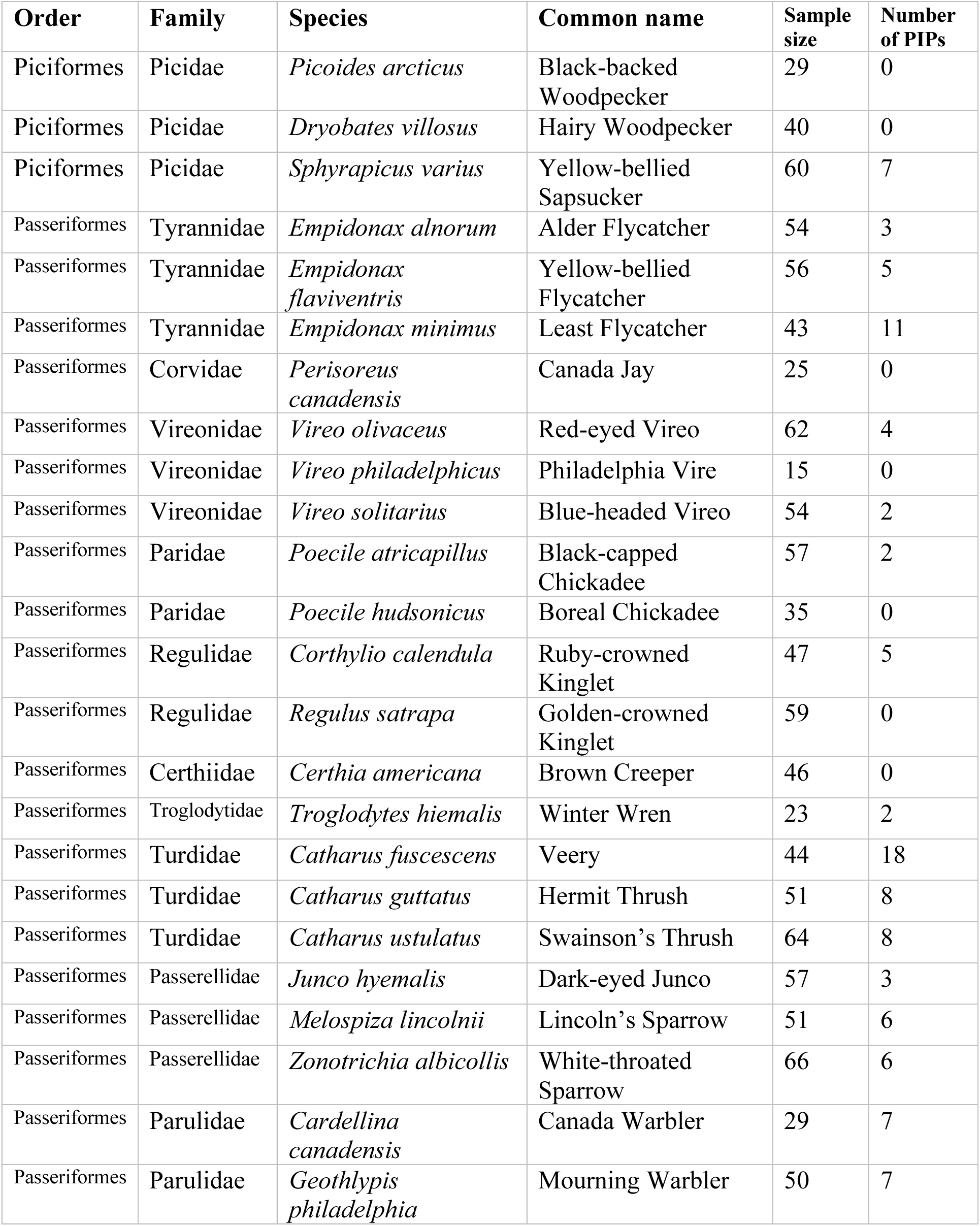
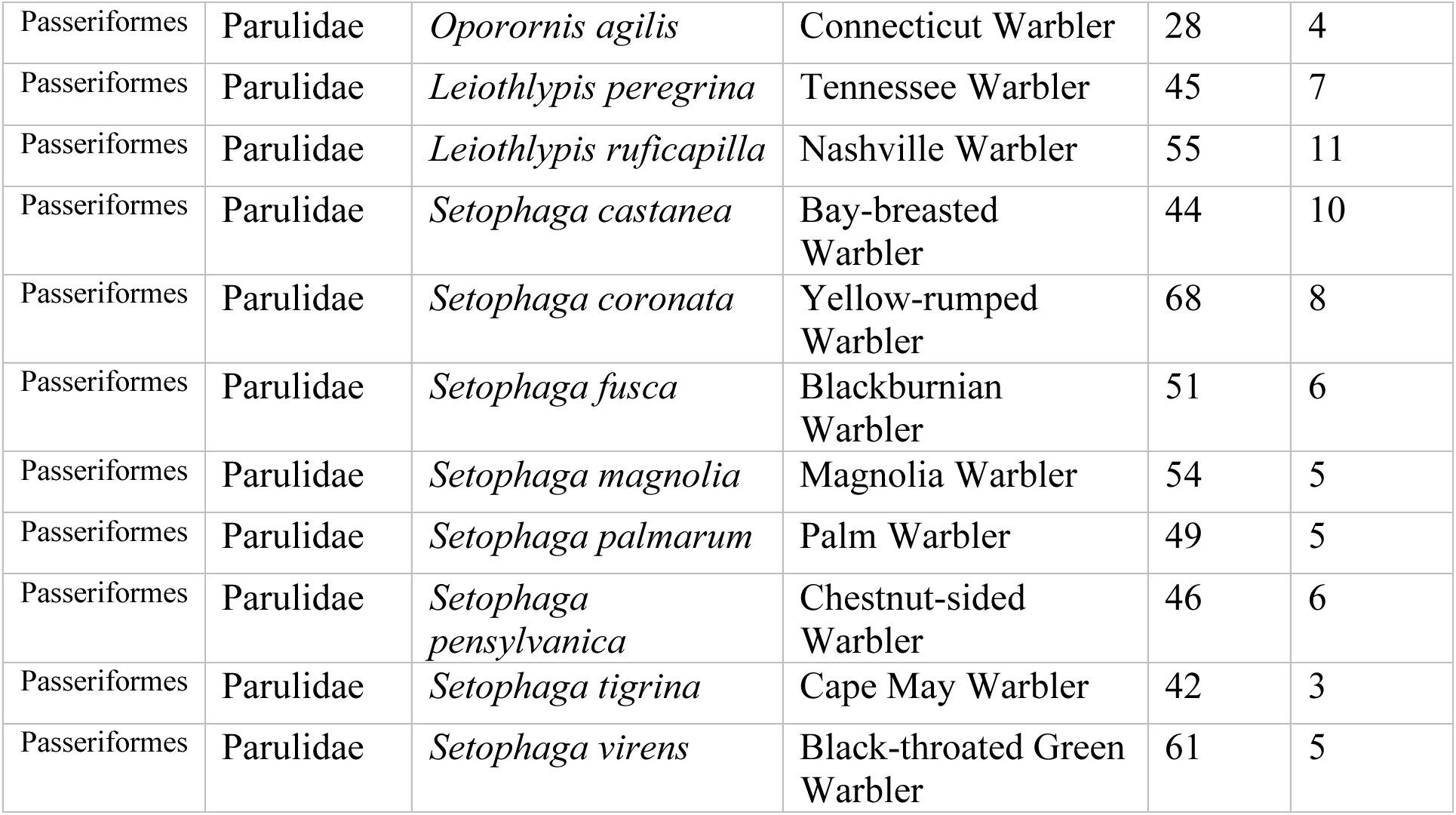
North American songbird and woodpecker species examined in the study, with the sample size for each species and the total number of putative inversion polymorphisms (PIPs) identified. Species are shown in taxonomic order. All individuals were sampled from across the contiguous North American boreal ecoregion as broadly and evenly as possible, as described in *Materials and Methods* and in (Pegan et al. 2025).

**Table 2.** 27 noteworthy polymorphisms in our dataset, including trans-specific polymorphisms spanning multiple species, polymorphisms that deviate significantly from Hardy-Weinberg Equilibrium (HWE), the 10 largest polymorphisms in our dataset (≥ 38 Mb in length), and those showing clear spatial structure. We also include information on the number of genotypes present in our samples (as indicated by PCA clusters) for each PIP. More detailed metadata for each PIP can be found in Table S2, Dataset S1, and Dataset S2.

| Mapped chromosome | Species | Number of genotypes | $\geq 38$ Mb in length | Notable spatial structure | Trans-specific polymorphism | Significantly out of HWE? | HWE D value |
| --- | --- | --- | --- | --- | --- | --- | --- |
| CM020533.1 | *Empidonax flaviventris* | 3 | T; 59 Mb | F | F | F | NA |
| CM020533.1 | *Empidonax minimus* | 3 | T; 40 Mb | F | F | F | NA |
| CM020534.1 | *Empidonax alnorum* | 3 | T; 38 Mb | F | F | F | NA |
| CM020534.1 | *Empidonax flaviventris* | 2 | T; 102 Mb | F | F | F | NA |
| CM020534.1 | *Empidonax minimus* | 6 | T; 50 Mb | F | F | F | NA |
| CM020536.1 | *Empidonax flaviventris* | 3 | T; 57 Mb | F | F | T; $p=0.024$ | -2.84 |
| CM020537.1 | *Empidonax minimus* | 3 | T; 69 Mb | T | F | F | NA |
| CM003717.2 | *Poecile atricapillus* | 3 | F | F | F | T; $p< 0.0001$ | -5.58 |
| CM020341.1 | *Catharus fuscescens* | 3 | F | T | F | F | NA |
| CM020367.1 | *Catharus guttatus*,<br>*Catharus fuscescens*,<br>*Catharus ustulatus* | 3 | F | F | T | F | NA |
| CM020368.1 | *Catharus guttatus*,<br>*Catharus fuscescens*,<br>*Catharus ustulatus* | 3 | F | F | T | F | NA |
| CM020371.1 | *Catharus guttatus*,<br>*Catharus fuscescens*,<br>*Catharus ustulatus* | 3 | F | F | T | F | NA |
| CM042573.1 | *Zonotrichia albicollis*<br>(previously described 3/3<sup>a</sup> inversion) | 3 | T; 101 Mb | F | F | F | NA |
| CM042576.1 | *Zonotrichia albicollis*<br>(previously described 2/2<sup>m</sup><br>inversion) | 2 | T; 106 Mb | F | F | F | NA |
| CM019929.1 | *Oporornis agilis*,<br>*Geothlypis*<br>*philadelphia* | 3 | F | F | T | F | NA |
| CM027507.1 | *Setophaga*<br>*coronata* | 3 | F | T | F | 0.0019 | -4.69 |
| CM027508.1 | *Setophaga*<br>*magnolia* | 3 | F | T | F | 0.037 | -3.11 |
| CM027510.1 | *Cardellina*<br>*canadensis* | 3 | T; 53<br>Mb | F | F | F | NA |
| CM027511.1 | *Leiothlypis*<br>*peregrina* | 3 | F | F | F | T; p=0.0096 | -2.33 |
| CM027513.1 | *Leiothlypis*<br>*ruficapilla*,<br>*Setophaga*<br>*castanea*,<br>*Setophaga*<br>*palmarum* | 2-3 | F | F | T | F | NA |
| CM027528.1 | *Leiothlypis*<br>*peregrina* | 3 | F | F | F | T; p< 0.0001 | -8.95 |
| CM027530.1 | 11 warbler species<br>aligned to<br>GCA_001746935.2 | 3 | F | F | T | F | NA |
| CM027532.1 | 11 warbler species<br>aligned to<br>GCA_001746935.2 | 3 | F | F | T | F | NA |
| CM027532.1 | *Setophaga*<br>*pensylvanica* | 3 | F | F | T | T; p=0.036 | -3.43 |
| CM027532.1 | *Setophaga tigrina* | 3 | F | F | T | T; p=0.0039 | -4.85 |
| CM027534.1 | 9 warbler species<br>aligned to<br>GCA_001746935.2 | 3 | F | F | T | F | NA |
| CM027536.1 | *Setophaga*<br>*pensylvanica* | 3 | F | F | F | T; p< 0.0001 | -6.24 |

Several species of birds have served as focal systems for understanding phenotypic effects of inversion polymorphisms in wild populations. These include the Ruff (*Calidris pugnax*) and the White-throated Sparrow (*Zonotrichia albicollis*), both of which display linked plumage and behavioral polymorphisms that are associated with inversions (Küpper et al. 2016; Lamichhaney et al. 2016; Tuttle et al. 2016). In other bird species, inversions have been identified as the basis of polymorphisms in plumage (Funk et al. 2021; Sanchez-Donoso et al. 2022), sperm morphology, (Kim et al. 2017; Knief et al. 2017), and migratory behavior (Sanchez-Donoso et al. 2022; Lundberg et al. 2023). Typically, such inversions have been discovered by searching for genetic variants that correlate with known phenotypic variation of interest such as plumage coloration or migratory behavior. By contrast, broader surveys for inversion polymorphisms in wild birds have been scarce, but evidence suggests that inversion polymorphisms are typically present in bird populations (Hooper and Price 2015, 2017; Knief et al. 2016; Ishigohoka et al. 2021; Edwards et al. 2025). The prevalence of inversion polymorphisms in wild bird populations, and the extent to which they tend to be linked to clear cases of local adaptation or phenotypic polymorphism, is not well known.

We hypothesized that North American bird species, like the model organisms that have been studied, display a variety of inversion polymorphisms beyond those that can be easily linked to discrete phenotypic variation. Our study species provide an opportunity to test this hypothesis by surveying for inversions within large, broadly distributed populations of wild birds that lack both phylogeographic structure (Pegan et al. 2025) and the type of phenotypic variation that is commonly the focus of studies on avian inversion evolution (Küpper et al. 2016; Lamichhaney et al. 2016; Tuttle et al. 2016; Sanchez-Donoso et al. 2022). Among the 35 species we analyzed, there are no described intraspecific phenotypic polymorphisms within the boreal ecoregion we sampled, with one exception: the sparrow *Zonotrichia albicollis* displays a species-wide plumage and behavioral polymorphism associated with a well-studied inversion (the *2/2^m^* polymorphism; an additional inversion polymorphism in this species called *3/3^a^* may also influence phenotype (Thorneycroft 1975; Tuttle et al. 2016)). Because North American birds receive substantial attention from researchers and birdwatchers, it is likely that any undescribed phenotypic polymorphisms in our study species are subtle or not associated with easily observable behavior or signaling traits such as plumage or song. This does not mean that no phenotypic polymorphisms exist—many populations exhibit polymorphisms in important traits that are difficult to detect without dedicated study, such as sensitivity to environmental stressors (Hoffmann et al. 2004; Tepolt and Palumbi 2020; An et al. 2024). Even so, our study provides a distinct perspective on the prevalence and role of inversions in wild bird populations by surveying for inversions without relying on hypotheses generated from phenotypic variation.

We focus specifically on surveying for large inversion polymorphisms present at relatively balanced frequencies in their respective populations. Large, balanced polymorphisms are particularly unlikely to be neutral with respect to fitness (Connallon and Olito 2022; Berdan et al. 2023), and their presence within populations suggests that they may have complex and consequential roles in the evolutionary dynamics of these populations, irrespective of their connection to discrete phenotypes. We leverage indirect inversion polymorphism detection methods (Li and Ralph 2019; Huang et al. 2020; Mérot 2020); Fig S1), which are well-suited to detect polymorphisms that are large, old, and present at relatively balanced frequencies in the population (Mérot 2020; Berdan et al. 2023). Indirect inversion detection relies on evaluation of the genetic consequences of recombination suppression, as opposed to direct evidence of physical inversion. These indirect methods therefore carry the caveat that they cannot confirm that polymorphic haploblocks are caused by physical inversion as opposed to other sources of recombination suppression (e.g., (Bascón-Cardozo et al. 2022; Shipilina et al. 2023; Irwin et al. 2025). Nonetheless, indirect methods are a valuable step in quantifying inversion diversity of non-model organisms (Mérot 2020), and many of the consequential evolutionary dynamics that are most frequently discussed in the context of inversions also apply to other kinds of haploblocks that experience reduced recombination (Shipilina et al. 2023; Dallaire et al. 2025; Irwin et al. 2025). Using PCA-based methods available for low-coverage whole genomes (Meisner and Albrechtsen 2018; Simon 2023), we apply the indirect inversion detection protocol described by (Huang et al. 2020; Mérot 2020) to low-coverage whole genome datasets, which were feasible for us to generate among many individuals in our non-model focal taxa.

Upon identifying putative inversion polymorphisms in our study populations, we describe characteristics of each polymorphism including its mapped length and the relative frequency of its alternative haplotypes, and we test three additional hypotheses. First, we test whether polymorphisms deviate significantly from Hardy-Weinberg Equilibrium (HWE), as expected if they are under strong selection. Second, we hypothesize that some inversion polymorphisms play a role in local adaptation across the >3500 km expanse of our sampling region, as has been described in other wild species including birds (Harringmeyer and Hoekstra 2022; Battlay et al. 2023; Knief et al. 2024; Wooldridge et al. 2024). We predict that polymorphisms involved in local adaptation will show spatial structure in the sampling region, reflecting differential selection across environmental gradients. Third, we hypothesize that polymorphisms which appear to be shared by multiple related species may have a common origin—i.e., that they do not represent independent inversion events. Such “trans-specific” polymorphisms can arise when ancient polymorphisms are maintained in daughter species after speciation or when inversion haplotypes are shared across species through introgression (Fontaine et al. 2015). We evaluate evidence for trans-specific polymorphism by examining genetic distance-based phylogenetic trees of putatively inverted regions. As all our study taxa are well-defined species, there is a strong general expectation that conspecific individuals should cluster more closely with each other in a phylogenetic tree than with heterospecifics. We predict that trees made from trans-specific polymorphisms will violate this expectation, reflecting the distinct evolutionary history of these regions compared with the rest of the genome (Fontaine et al. 2015; Knief et al. 2024).

## Results

### Most species show evidence of autosomal inversion polymorphisms

Out of the 35 species we analyzed, 28 species (80%) showed evidence of at least one autosomal inversion polymorphism (Table 1). Polymorphisms were particularly prevalent in flycatchers (Tyrannidae), thrushes (Turdidae), warblers (Parulidae), and sparrows (Passerellidae). By contrast, we detected no polymorphisms in two out of the three woodpecker species we examined. We identified approximately 170 previously undescribed putative inversion polymorphisms (which we refer to as PIPs) across 28 species (Dataset S1). We also detected polymorphisms that have been described previously in sparrows, including the two well-known inversions in *Zonotrichia albicollis*—the *2/2^m^*inversion and the *3/3^a^* inversion (Thorneycroft 1975)—and two polymorphisms in *Junco hyemalis* which may be those described in past cytological work (Shields 1973). We verified the *Z. albicollis 2/2^m^* inversion using 22 samples with associated museum voucher specimens to confirm the association between the inversion genotype our methods assigned to these individuals and the plumage phenotype it is known to control (Table S1, Supplementary Information). Of the 174 total PIPs in our dataset, 139 polymorphisms produced patterns consistent with inversion in both PCA and linkage disequilibrium analyses (Fig. 2C,G, Dataset S2; see *Materials and Methods*). The remaining 35 polymorphisms had clear evidence for inversion in PCA (Fig. 2C, Dataset S2) but ambiguous results from linkage analyses. Among species with evidence of inversion polymorphism, the mean number of LD-supported PIPs identified per species was 5 (sd = 3), ranging from 1 to 14 per species. We did not evaluate evidence for inversions on sex chromosomes due to the difficulties of accurately mapping short reads to sex chromosomes, especially in cases where the corresponding reference genome lacks the W sex chromosome.

**Figure 2.**
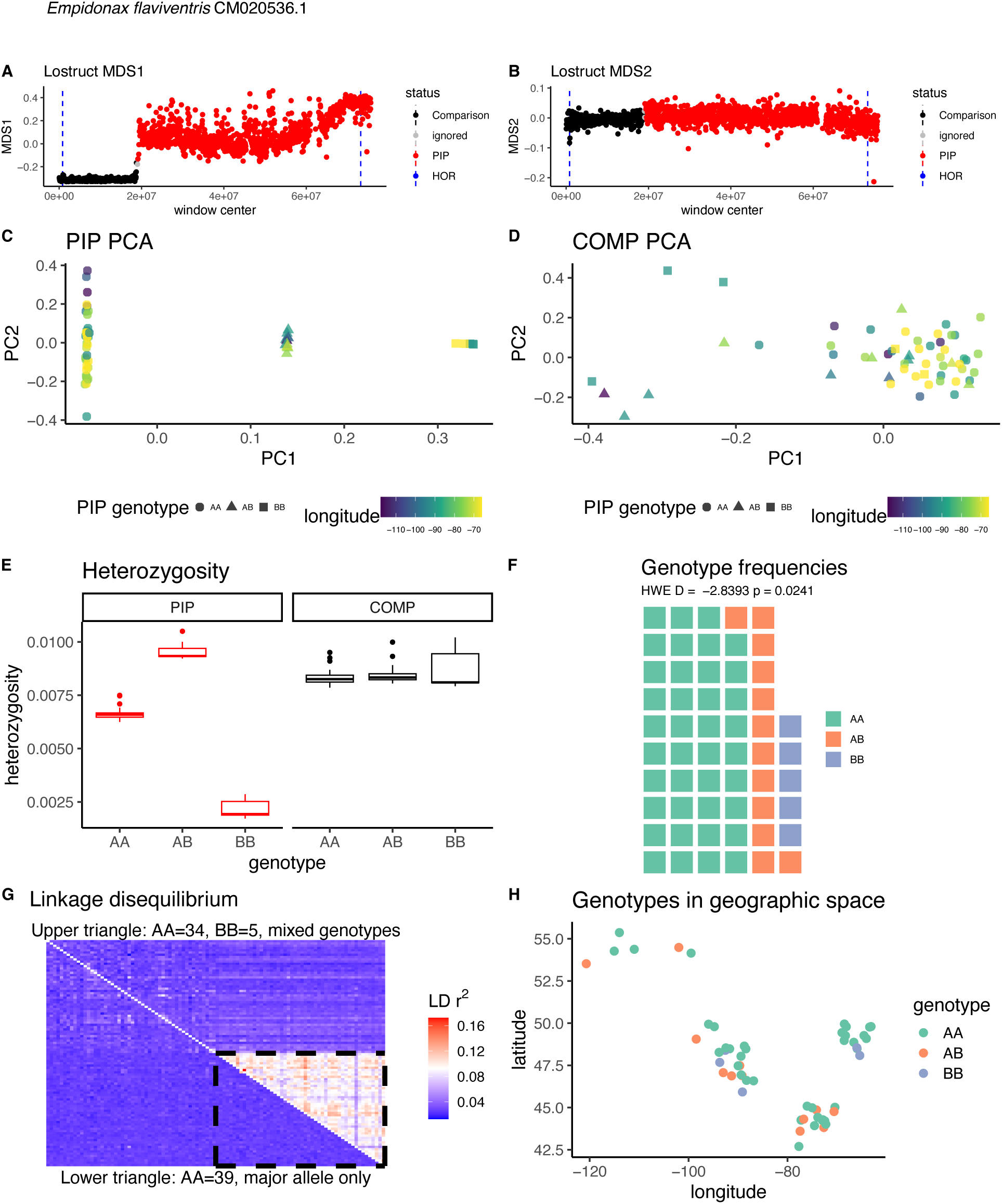
Summary plots from a single example putative inversion polymorphism (PIP) in *Empidonax flaviventris,* mapped to the chromosome CM020536.1. Plots for all PIPs can be found in Dataset S2. **A.** Analysis with lostruct (see *Materials and Methods*) reveals a long segment of the chromosome with distinct population structure identified by multidimensional scaling (MDS). The pattern is present on MDS 1 but not MDS 2 (**B**). The highlighted red region is identified as a PIP (see *Materials and Methods*). Blue dashed lines in panels **A** and **B** represent annotations of high-order repeats (HORs)—repetitive regions associated with centromeres and sometimes inversions—on the reference chromosome. **C.** PCA reveals a 3-cluster pattern in the PIP region, consistent with the presence of an inversion polymorphism. Individuals in the central cluster are assumed to be heterozygotes for the polymorphism, whereas those in the right and left clusters are assumed to be homozygotes (Mérot 2020a). **D.** The 3-cluster pattern is not present in the comparison (COMP) region of the chromosome, where no inversion polymorphisms are thought to be present. Points in panels **C** and **D** are colored by the longitude of individuals’ sampling location, as in Fig. 1A, and show no evident geographic structure. **E.** Putatively heterozygous individuals show elevated heterozygosity within the PIP region compared to putatively homozygous individuals, consistent with the presence of an inversion polymorphism (Mérot 2020). This pattern does not exist in the COMP region. **F.** PIP genotype frequencies of the sampled population. Haplotypes are named A and B arbitrarily based on their frequency, where A is the more common haplotype. **G.** Linkage disequilibrium (LD) analysis shows a pattern consistent with the presence of an inversion polymorphism, where LD in a sample of individuals with both classes of homozygous genotypes (upper triangle) is elevated in the PIP region compared with the COMP region. No such pattern exists in a sample of individuals that all share the same genotype (lower triangle), which is expected in cases of inversion because recombination suppression should create linkage disequilibrium only between chromosomes that have alternative haplotypes (Mérot 2020). **H.** Individuals are shown in geographic space, colored by their PIP genotype, to illustrate the spatial distributions of the haplotypes. Individual locations are jittered to allow all points to be seen across the range (actual sampling locations can be seen in Fig. 1A). In this case, both PIP haplotypes are present throughout the range, indicating low spatial structure in the PIP.

### Detected polymorphisms tend to be large and some are near centromeric repeats

The mean mapped length of LD-supported PIPs in our dataset was 8.59 Mb (sd = 18.16 Mb). PIPs lacking LD support showed a tendency to be smaller, though many were still quite large (mean = 6.04 Mb, sd = 13.80 Mb). Across all PIPs, mapped length ranged from 50 Kb to 106.0 Mb (Fig. 1D). The largest PIP we detected is the well-studied *2/2^m^* inversion polymorphism in *Zonotrichia albicollis*. Flycatchers in the genus *Empidonax* showed a strong tendency to have large PIPs: seven out of the ten largest PIPs in our dataset, all greater than 38 Mb in length, belonged to the three *Empidonax* species that we analyzed (Table 2). PIP length is necessarily bounded by the length of the reference chromosomes to which we mapped our data, which affects the distribution of lengths. In Fig. S2, we show the distribution of PIP lengths as a proportion of their reference chromosomes, demonstrating that a substantial proportion of PIPs occupy 10% or more of the reference chromosome they are mapped to. PCA-based methods are particularly well-suited to detecting large polymorphisms (Mérot 2020; Berdan et al. 2023), and there may be many small inversion polymorphisms in our datasets (especially those < 50 Kb in length) that we were not able to detect.

Large inversions (>1 Mb in length) have been shown to be associated with repetitive regions such as centromeric satellites, which may promote their formation (Gozashti et al. 2025). Variation in reference genome quality prevented us from systematically evaluating the repeat landscape in our dataset or inferring whether PIPs are paracentric or pericentric (see Supplementary Information), but we found that many PIPs have breakpoints that were mapped close to high-order repeats (HORs) (Dataset S3, Supplementary Information), which are repetitive satellite regions associated with centromeres (Qi et al. 2025).

### Detected polymorphisms tend to be present at balanced frequencies and in Hardy-Weinberg Equilibrium

The PIPs we identified tend to show relatively balanced haplotype frequencies (Fig. 1C). The majority of PIPs (76%, *n* = 133) produce 3 clusters on a PCA, corresponding to all three possible genotype states for a locus with two alleles (Mérot 2020) (Dataset S1-S3). We also identified 34 PIPs that produced only two PCA clusters, which may represent cases where one of the possible genotypes is rare or lethal. This category includes the *2/2^m^* inversion polymorphism in *Zonotrichia albicollis*, for which one of the two classes of homozygotes is extremely rare and not present in our dataset (Tuttle et al. 2016). Finally, we identified 7 PIPs that produced 4, 5 or 6 clusters on PCA, which may represent cases where the polymorphism comprises more than two alleles (Fig. S3) (Dallaire et al. 2025). We hereafter refer to PIPs that produce 3 PCA clusters as “biallelic PIPs.” PIPs that produce 2 clusters may be biallelic as well, but we are not able to confirm allele frequencies in these cases because we cannot infer which clusters represent heterozygotes versus homozygotes with the available data (Huang et al. 2020; Mérot 2020); see *Materials and Methods*).

In the subset of 101 LD-supported biallelic PIPs, we found that the mean minor allele frequency across all species was 0.35 (sd = 0.11). The 32 biallelic PIPs that lacked LD support tended to show lower minor allele frequencies (mean = 0.28, sd = 0.14), raising the possibility that low-coverage analyses of LD (*see Materials and Methods*) may work better with polymorphisms at more balanced frequencies.

The vast majority (92.5%) of the 133 biallelic PIPs show genotype frequencies that do not deviate from expectations under Hardy-Weinberg Equilibrium (HWE; Dataset S1). Among the 10 biallelic PIPs that did deviate significantly from HWE, the D value was negative, indicating an excess of heterozygotes compared to expectations (Table 2). We did not observe a relationship between the length of a PIP and its likelihood of being in HWE (Fig. S4). PIPs that produce only 2 clusters in PCA may be missing an entire class of genotypes and may therefore be more likely to deviate significantly from HWE than those that produce 3 clusters, but it is not possible to evaluate HWE in these cases because we cannot be certain whether the missing genotype class involves homozygotes or heterozygotes.

### Putative heterozygotes tend to have elevated heterozygosity within PIP regions

Individuals with a heterozygous inversion genotype are generally expected to show higher heterozygosity in the inverted region than homozygous individuals (Mérot 2020) (Fig. 2E, Dataset S2). This is because inversion haplotypes do not recombine, which means that the haplotype-specific variation that they capture upon formation, and later accumulate through mutation, is necessarily heterozygous when the inversion haplotype is heterozygous. We observed this expected elevation of heterozygosity within the heterozygous inversion genotype in 76 out of the 133 biallelic PIPs. Among the remaining biallelic PIPs, 41 showed a pattern where heterozygosity was significantly lower in one class of homozygotes than in the other two genotypes, which did not differ significantly from each other. The remaining 16 PIPs in this category showed neither pattern. Among the 34 PIPs with 2 observed genotypes, we observed a significant difference in heterozygosity between the genotypes in 19 cases (56%).

### Most polymorphisms do not show spatial structure within the sampled region

We assessed the potential role of PIPs in local adaptation by evaluating spatial structure in the distribution of PIP haplotypes. Inversions can facilitate local adaptation across discrete habitat types (Huang et al. 2020; Harringmeyer and Hoekstra 2022) and environmental gradients (Akopyan et al. 2022; Knief et al. 2024), both of which produce spatial patterns in the distribution of inversion haplotypes. Although our samples come from within consistent habitat types across the sampling region (boreal or hemiboreal forest), subtler environmental gradients across the >3500 km breadth of the sampling region could create local selective pressures that lead to spatial gradients in PIP haplotype frequency. We visually assessed PCA and spatial sampling plots for each PIP to evaluate qualitative evidence for spatial structure (Fig. 2C,H, Dataset S2). Further, we estimated a slope of isolation by distance (IBD) for each biallelic PIP (Fig. 1B). We observed very little notable spatial variation in genotype or haplotype frequencies. Across all biallelic PIPs, mean IBD slope ranged from -0.00008 to 0.00019. Heterozygotes tend to exist across the entire breadth of the sampling region, even in most of the PIPs with the strongest IBD slopes. We observed four notable exceptions to the general lack of spatial structure wherein one portion of the range was strongly dominated by a particular haplotype (Table 2; Dataset S2).

### Several polymorphisms are shared across species

We identified 26 cases where PIPs were identified on the same chromosome in multiple species that had been mapped to the same reference genome (Table S2). In 12 cases (Table 2), a genetic distance neighbor-joining tree of the overlapping region showed some individuals clustering by PIP genotype instead of by species (Fig. 3, Dataset S3), which is consistent with trans-specific polymorphism (Fontaine et al. 2015; Knief et al. 2024). That is, within the PIP region, some individuals homozygous for the A haplotype were found to be more closely related to a heterospecific than to a conspecific homozygous for the alternative B haplotype. These patterns suggest that the corresponding polymorphism did not have independent origins in each species, because if this were the case, we would expect haplotypes from conspecifics to be more closely related to each other than to haplotypes from heterospecifics—the general expectation for the rest of the genome (e.g., Fig. 3A).

**Figure 3.**
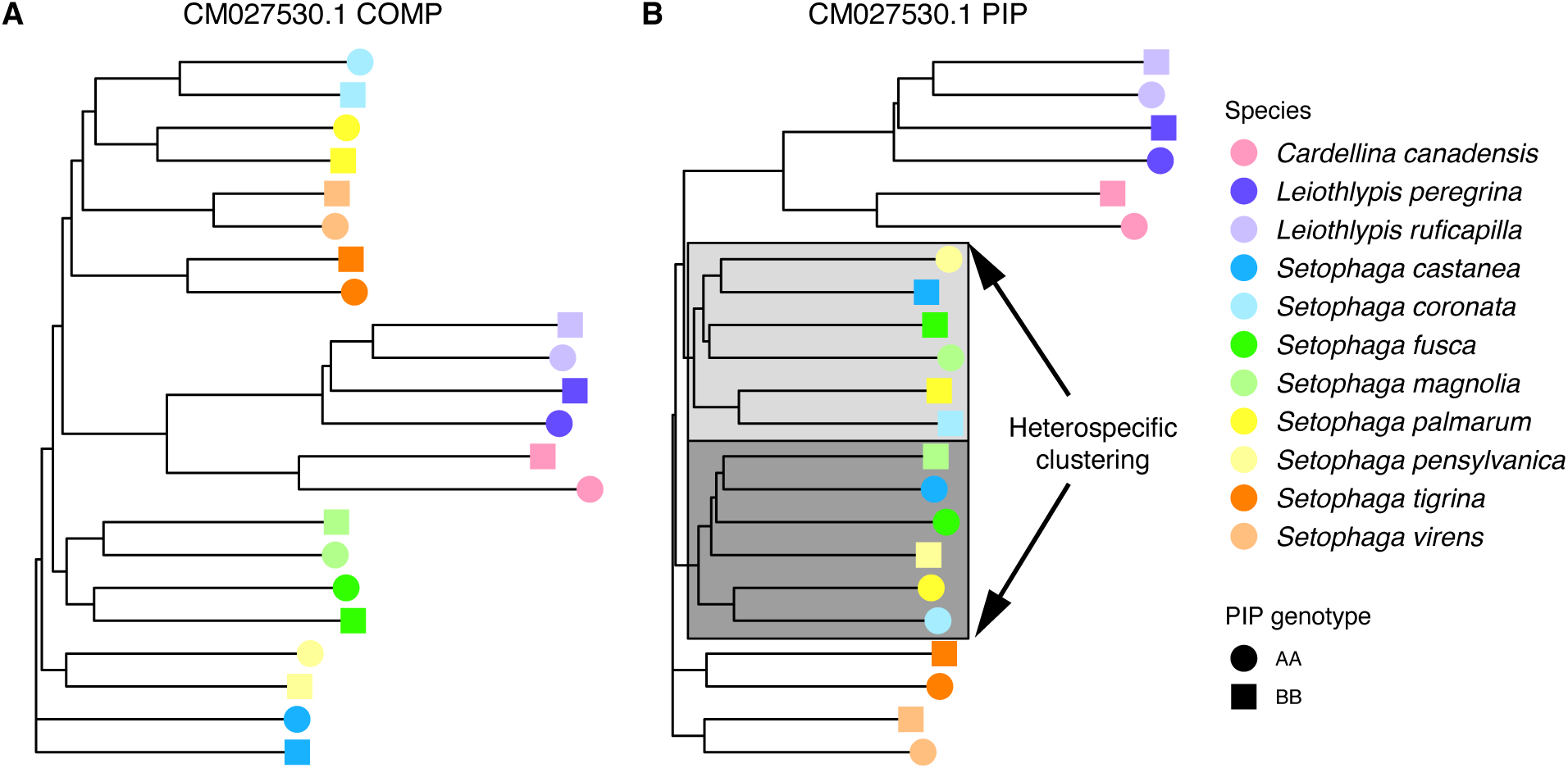
Neighbor-joining genetic distance trees illustrate different patterns of relatedness within a region of putative inversion polymorphism (PIP) versus a comparison (COMP) region, showing evidence that this PIP may be a trans-specific polymorphism. Plots for all trees can be found in Dataset S3. This example illustrates trees from two regions on the chromosome CM027530.1, which is part of the reference genome to which we mapped the 11 species of warblers (Parulidae) shown in the legend. All species show a PIP mapped to the same place on this chromosome. Each tree includes a randomly chosen representative of each of the two alternative homozygous PIP genotypes for each species. **A.** In the comparison (COMP) region, where no inversions are thought to be present, individuals cluster with their conspecifics. By contrast, in the PIP region (**B**), relationships among individuals of six *Setophaga* species (those in shades of blue, green, and yellow) do not follow this expected pattern. Instead, pairs of conspecifics are divided into two heterospecific clusters of six (highlighted in light and dark gray), where each cluster represents homozygotes for one of the PIP haplotypes. Note that genotypes are named arbitrarily based on haplotype frequencies within their respective species. This plot suggests that the B haplotypes of *S. magnolia* and *S. pensylvanica* may be homologous with the A haplotype of the other four species involved in these clusters, and vice versa.

## Discussion

Our study uses PCA-based methods (Li and Ralph 2019; Huang et al. 2020; Mérot 2020) to provide evidence that large inversion polymorphisms may be widespread in passerine bird populations. Whereas the majority of research on inversions in non-model organisms relates to effects of inversions on phenotypic appearance and behavior, our results indicate that large inversions can be common in populations even when they cannot be linked to easily observable phenotypes, as is well known in model systems (Berdan et al. 2023). Many studies in wild, non-model organisms have relied on analysis of *F_st_* to identify inversions, which requires *a priori* identification of populations or phenotypic categories to compare. This approach is similar to the way that early molecular geneticists approached genetic variation in general—by identifying a genetic variant through its effect on an observable phenotype (Charlesworth et al. 2016). Hubby and Lewontin’s (1966) breakthrough in measuring genetic variation directly, without relying on phenotypic hypotheses, revealed that the evolutionary dynamics influencing genetic variation are much more complex than could be appreciated with a phenotype-driven approach (Hubby and Lewontin 1966; Lewontin and Hubby 1966; Charlesworth et al. 2016). Analogously, this and other genome-wide surveys for inversion polymorphisms are revealing that structural variants are pervasive in wild populations, beyond cases where they can be linked to discrete phenotypic polymorphisms (Hooper and Price 2015, 2017; Ishigohoka et al. 2021; Knief et al. 2024; Wooldridge et al. 2024; Edwards et al. 2025). Some of these newly detected polymorphisms may be associated with phenotypes that are important for fitness but difficult to observe, such as immune function, fecundity, or environmental tolerance (Hoffmann et al. 2004; Tepolt and Palumbi 2020; Minias et al. 2022; An et al. 2024). More generally, the increasing availability of empirical data on inversion polymorphisms is revealing patterns which emphasize that there are many fundamental unanswered questions about the evolutionary processes maintaining diversity of large polymorphic haploblocks in populations, mirroring questions about genetic diversity maintenance that have been studied for decades using single nucleotide polymorphisms (SNPs) (Charlesworth et al. 2016; Buffalo 2021; Johnson et al. 2023).

Most of the polymorphisms we identified (*n =* 123) are present in genotype frequencies that do not significantly deviate from Hardy-Weinberg Equilibrium (HWE), which means that we cannot reject the hypothesis that they could be segregating neutrally. The presence of so many large polymorphisms in apparent HWE is surprising given the theoretical expectation that large inversions (or other large haploblocks caused by recombination suppression) are particularly unlikely to be selectively neutral. This expectation is based on the hypothesis that large haploblocks are unlikely to capture combinations of internal genetic variants, such as SNPs, with a net neutral effect on fitness—more likely, the specific combination of SNPs in a haploblock will be at least slightly beneficial or deleterious, exposing the haploblock to natural selection for fixation or loss (Connallon and Olito 2022; Berdan et al. 2023). As such, maintenance of large haploblocks at intermediate frequencies suggests a role for balancing selection or other complex evolutionary forces acting to prevent fixation or loss (Connallon and Olito 2022; Berdan et al. 2023). Yet, our empirical dataset suggests that large polymorphisms with genotype frequencies indicative of selective pressure are the exception rather than the rule. Despite their large physical size, many of these polymorphisms may have fitness effects weak enough to subject them to nearly-neutral dynamics, where the efficacy of selection to remove deleterious alleles depends on effective population size (Ohta 1992). Recent pangenome analyses in scrub-jay (*Aphelocoma*) species with varying effective population size suggests that most structural variants, including inversions, are weakly deleterious and conform to predictions from nearly neutral theory (Edwards et al. 2025).

Among the 10 polymorphisms that do significantly deviate from HWE, we observe an excess of heterozygotes, as expected under forms of balancing selection such as heterozygote advantage (Wellenreuther and Bernatchez 2018; Berdan et al. 2023). We also observed 41 polymorphisms that we could not evaluate for HWE because they produce more or fewer than 3 clusters on a PCA, and many of these could experiencing complex selective dynamics as well— for example, the *2/2^m^* polymorphism of *Zonotrichia albicollis* produces only two PCA clusters and deviates strongly from HWE due to the extreme rarity of one of the classes of homozygotes (Tuttle et al. 2016). We do not know of ecological variation that may explain why some polymorphisms appear to be in balancing selection, but they could be associated with difficult to observe phenotypes, such as aspects of the immune system (Minias et al. 2022).

The polymorphisms we identified show an intriguing combination of characteristics— some patterns are consistent with described characteristics of inversion polymorphisms, while others deserve further attention. For example, we found that haploblock breakpoints often correspond with centromeric satellites, which is consistent with the observation that large inversions are often located near centromeric satellites and that these satellites may even promote inversion formation (Gozashti et al. 2025). However, centromeres themselves also experience recombination suppression and can produce large haploblock polymorphisms that might look like inversions on PCA, raising the possibility that some PIPs represent polymorphic centromeres rather than inversions (Logsdon et al. 2024; Irwin et al. 2025). Using low-coverage short-read sequencing data, we are unable to directly evaluate the presence of structural inversions in the genome, so some of these polymorphisms could be large haploblocks that exist for other reasons (Shipilina et al. 2023; Dallaire et al. 2025; Irwin et al. 2025). Most haploblocks in our data showed patterns of linkage disequilibrium expected for inversions, but some (20%) deviated from these expectations. However, LD calculations are computationally intensive and the reduced datasets we used for them may be prone to issues caused by the limitations of low-coverage inference methods.

A substantial minority of biallelic polymorphisms (37%) deviate from expectations about average heterozygosity of single nucleotide polymorphisms (SNPs) along the span of an inversion (Huang et al. 2020; Mérot 2020). Individuals heterozygous for an inversion haplotype are expected to have higher average SNP heterozygosity in the affected region than those with homozygous haplotypes because recombination suppression promotes divergence between the haplotypes. Yet, it is not clear how general these heterozygosity expectations should be for inversions, given that genetic differentiation between inversion haplotypes arises through complex processes and changes with the age of the polymorphism (Charlesworth 2023). For example, very young inversions have not had much time to develop haplotype-specific variants and may show strongly depressed SNP heterozygosity when homozygous, without demonstrating elevated SNP heterozygosity when in a heterozygous state. This pattern— depressed SNP heterozygosity in one class of homozygotes compared to the other two genotypes—accounts for most of the “unexpected” heterozygosity patterns we observed. As more empirical data on inversions become available across a variety of systems, there will be new opportunities to investigate how the evolutionary dynamics of inversions shape patterns of heterozygosity within affected genome regions.

Direct evaluation of the genomic structure underlying PIPs will require population-level long-read sequencing, such as a pangenome approach (Secomandi et al. 2025), which is not yet practical for variant discovery in the large number of samples we analyze here (*n* = 1660 of 35 species). Another approach for confirmation could be to use cytology, for example with fluorescence in-situ hybridization (FISH) to link sequence data with chromosomal architecture visible in a karyotype (Shearer et al. 2014; Artemov et al. 2017; Deakin et al. 2019). A few of our study species have been involved in cytological studies from past decades describing inversion differences between species or populations (Shields 1973; Rising and Shields 1980; Shields et al. 1987; Hooper and Price 2017), but it is not possible to be certain of how the physical chromosomes photographed in older karyotypes correspond to the scaffolded, sequence-based “chromosomes” inferred by our reference-mapped analyses (Lewin et al. 2019; Iannucci et al. 2021). Regardless of whether all of the massive, well-differentiated polymorphisms we detected are caused by genomic inversions or other processes with similar effects, their maintenance in populations and their consequences for evolution warrant further investigation.

Many studies have identified inversion polymorphisms that show spatial gradients corresponding to environmental variation, which can be interpreted as signs of local adaptation (Huang et al. 2020; Harringmeyer and Hoekstra 2022; Knief et al. 2024; Wooldridge et al. 2024). By contrast, we find that most PIPs in boreal bird populations are geographically well mixed. Not all inversion polymorphisms necessarily capture locally adaptive combinations of alleles. Moreover, the populations sampled in our boreal belt study system may be less likely to demonstrate inversion-associated local adaptation than populations that span stabler or more heterogeneous habitats. The environment across our sampling region is relatively homogenous and lacks barriers to gene flow (van Els et al. 2012; Ralston et al. 2021; Pegan et al. 2025), which can erode local adaptation. Further, the spatial distributions of species in this region have undergone major disruptions throughout Pleistocene glacial cycles (Kimmitt et al. 2023), so contemporary patterns of local adaptation have not had a long time to accrue. Among PIPs that do show notable spatial structure (Table 2), further investigation is necessary to evaluate whether spatial structure is driven by local adaptation versus non-adaptive phylogeographic processes.

For example, a haplotype may have become fixed in an historical glacial refugium by chance, and then subsequently become the predominant haplotype in parts of the contemporary range closest to that refugium. With increased sampling and careful study design, these possibilities could be investigated using genotype-environment analysis.

Our data, alongside results from other studies, suggest that maintenance of inversion polymorphisms across species boundaries may be relatively common in closely related bird species (Tuttle et al. 2016; Hooper and Price 2017; Knief et al. 2024). Polymorphisms may be present in multiple species because they originated prior to species divergence or because they were shared across species through introgression, and their maintenance across species boundaries suggests they may be maintained as polymorphisms by selection (Fontaine et al. 2015; Brelsford et al. 2020; Jamie and Meier 2020; Yan et al. 2020; Knief et al. 2024). Incomplete lineage sorting can also maintain polymorphisms for long periods and may be an explanation for trans-specific polymorphism in recently diverged lineages. The groups of species displaying trans-specific polymorphisms in our dataset have divergence times ranging from about 0.5-4 million years ago, although divergence times are not well resolved in all cases (Barker et al. 2015; Everson et al. 2019; Harvey et al. 2020). Phylogenetic analyses based on higher-coverage data (as opposed to the distance-based trees we present here) is necessary to provide further insights into how the haplotypes involved in trans-specific polymorphism are related to each other and when they diverged (Fontaine et al. 2015).

### Conclusions: Inversions show under-appreciated patterns in diverse species

Our approach facilitated the discovery of polymorphisms in wild bird species that may have important, yet hidden, implications about the evolution of their corresponding populations. Within populations that appear phenotypically homogenous, we observed many large polymorphisms, including several that seem to be involved in complex evolutionary dynamics— as evidenced by signs of balancing selection, spatial structure, and maintenance across species boundaries (Table 2). Yet we found that the majority of large polymorphisms do not show such patterns and are present at genotype frequencies consistent with Hardy-Weinberg Equilibrium. It is not clear what evolutionary processes have resulted in the maintenance of these polymorphisms, setting up questions about the forces shaping diversity in large haploblocks that parallel similar questions about genetic diversity in general (Hubby and Lewontin 1966; Lewontin and Hubby 1966). It remains challenging to disentangle the evolutionary processes affecting inversions among non-model species that are not amenable to captive breeding or genetic manipulation, but we demonstrate that methods of indirect inversion detection can provide a feasible way to identify interesting cases as opportunities for further investigation, even in low-coverage sequencing data. Although indirect methods such as those employed here cannot confirm the presence of a physical inversion underlying each of the large polymorphic haploblocks we identify, our analyses demonstrate that most of these regions show patterns of linkage disequilibrium and heterozygosity consistent with inversion. Finally, our dataset adds to the scarce but proliferating availability of empirical data on inversion polymorphisms in wild populations, which will be useful for testing predictions informed by the long history of theoretical work on inversion evolution (Wellenreuther et al. 2019; Connallon and Olito 2022).

## Materials and Methods

### Species and sampling

We generated population genomic data from 35 species of North American boreal birds, as described in (Pegan et al. 2025). These codistributed species represent a substantial subset of the avian community in the forested North American boreal ecoregion. Three species are woodpeckers (Piciformes), and the remaining are from 17 genera and 10 families of songbirds (Passeriformes). We sampled species broadly and evenly along the longitudinal axis of the boreal ecoregion (Fig. 1A), intentionally excluding related populations in other ecoregions (e.g., Rocky Mountains) with known phenotypic or phylogeographic differences from boreal populations (Pegan et al. 2025). Most (∼88%) of the samples were ethanol-preserved or flash-frozen specimen-vouchered tissues provided by contributing natural history collections, while the remaining ∼12% came from unvouchered blood samples. After the filtering steps described below, mean sample size per species was = 47 ± 13 (range = 15–68 samples per species, for a total of 1660 samples; Dataset S4). Individuals from all species were sampled from across the 3500+ km longitudinal breadth of the eastern boreal belt (Fig 1A). This contiguous ecoregion lacks major geographic barriers, and consequently bird species typically do not show deep phylogeographic structure within this region even though it spans most of the longitudinal breadth of North America (Pegan et al. 2025).

### Sequencing

DNA extraction and low-coverage sequencing techniques are described in Pegan et al 2025, but in brief, we extracted DNA using DNeasy Blood and Tissue Kits (Qiagen Sciences, Germantown, MD, USA) or phenol-chloroform extraction and prepared libraries using a modified Illumina Nextera library preparation protocol (Therkildsen and Palumbi 2017; Schweizer et al. 2021). We generated paired-end sequence data with Illumina platforms (HiSeq, NovaSeq 6000, NovaSeq X). Demultiplexed data were processed using AdapterRemoval v2.3.1(Schubert et al. 2016) and fastp v0.23.2 (Chen et al. 2018) as described in (Pegan et al. 2025). Average genome-wide depth of alignment coverage, including sites with missing data, was approximately 2.8x.

### Alignment to reference genomes, filtering, and identification of SNPs to inform detection of possible inversions

We aligned samples to a reference genome from a related species published on GenBank (Zhang et al. 2014; Laine et al. 2016; Toews et al. 2016; Ruegg et al. 2018; Feng et al. 2020; Manthey et al. 2021; Friis et al. 2022; Sly et al. 2022) (Dataset S1) using bwa mem (Li 2013). Species were aligned to the most closely related species as possible with a chromosome-assembled reference genome available of GenBank at the time of analysis. We conducted additional supporting analyses to evaluate whether genetic distance between each aligned species and its reference genome biases PIP detection, finding no evidence for biases (Fig. S5). As described in detail in (Pegan et al. 2025), we filtered data to remove duplicates and read overlap, and we removed individuals from the dataset if they showed evidence of species misidentification or sample contamination. We used ANGSD v0.941 (Korneliussen et al. 2014) to calculate genotype likelihoods for all sites inferred to be SNPs at an alpha-level 0.05. We applied ngsParalog v1.3.2 (Linderoth 2018) to filter out SNPs with a high likelihood of occurring within a mis-mapped or paralogous region. We excluded all sex chromosomes from our analysis.

### Identification of putative inversion polymorphisms (PIPs)

We identified putative inversion polymorphisms primarily based on the presence of a chromosome region whose SNP data produces PCA plots with 2 or 3 distinct clusters on PC1 (Huang et al. 2020; Mérot 2020); Fig. S1; Dataset S2). In 7 cases, we flagged regions as putative inversion polymorphisms when they showed 4-6 clusters distributed across PC1 and PC2, which may reflect patterns caused by polymorphisms with more than two alleles (e.g. Fig. S3, Dataset S1, Dataset S2). We also assessed linkage disequilibrium (LD) at regions of interest to identify cases where LD is elevated in a sample of individuals from multiple PCA clusters (representing individuals showing a mix of inversion haplotypes) compared with a sample of individuals from within a cluster (representing individuals homozygous for one of the haplotypes). This pattern is expected of inversions, but not regions experiencing recombination suppression for other reasons, because chromosomes involved in inversions only experience recombination suppression when paired in meiosis with a chromosome that has the opposite inversion orientation haplotype (Huang et al. 2020). Most regions we identified using PCA also produced LD patterns consistent with inversion polymorphism (*n =*139 regions, Dataset S2). Evaluating linkage disequilibrium is computationally demanding and challenging using low-coverage sequence data. For this reason, we retained putative inversion polymorphisms in our downstream analyses if they produced 3 PCA clusters, even if LD analysis provided unclear evidence that the region was an inversion (*n* = 35 regions, Dataset S2). In the Results, we clearly distinguish between regions that satisfy both identification criteria (“LD-supported” PIPs) versus those that show PCA evidence only. We discarded regions from downstream analyses if they only produced 2 clusters on PCA and lacked evidence for inversion in LD analysis.

Figure S1 provides a detailed summary of how we identified putative inversion polymorphisms (PIPs) across the genome through a series of analysis steps (*A1-A4*) and decisions (*D1-D6*) based on analysis results. To identify chromosome windows showing evidence of inversion polymorphism, we used the program lostruct (Li and Ralph 2019) in a genotype likelihood framework with the program PCAngsd (Meisner and Albrechtsen 2018) using custom scripts available at https://github.com/alxsimon/local_pcangsd (Simon 2023) (Fig. S1, *A1*). With the SNP genotype likelihoods we generated in ANGSD, we applied lostruct to 50-kb genome windows with a minimum variant number of 500 loci per window. We also used lostruct’s “corners” function to sort chromosome windows based on whether they display one of the three most extreme patterns on the chromosome, as identified by lostruct, and to create a PCA for windows in each “corner” separately. This provides an automated, albeit coarse, way to inspect PCA patterns representing potential inversions. We specified that each corner should contain 30% of the windows on the chromosome. We visually examined lostruct’s output plots and we flagged regions for further analysis (Fig. S1, *D1*) when they met two conditions: 1) lostruct indicated that the region comprises a contiguous series of 50-kb windows with distinct local structure, as indicated by highly diverged values of MDS1 (i.e. a visually distinct “spike” in MDS1 in either the positive or negative direction), and 2) PCA of the “corner” windows associated with the spike produced a pattern of 2-6 distinct clusters in PC1 and PC2. We proceeded with analysis of regions that failed to meet the second condition in some cases when the spike on MDS1 was very clear (Fig. S1, *D2*) because the method of using “corners” to sort windows only coarsely captures the putative inversion region (i.e. when the inversion is small, many windows from beyond the region will be included in each corner).

To identify the specific location of each PIP, we used changepoint analysis on MDS1 values produced by lostruct (Fig. S1, *A2*, Dataset S2). This analysis does not identify specific inversion breakpoint loci, but rather identifies the 50-kb windows at which the distinct local structure pattern first appears and then disappears along the chromosome. We assigned the start and end of each PIP as the center of the 50-kb windows identified by the changepoint analysis.

In most chromosomes, we also identified “comparison” (COMP) regions that apparently do not involve inversions, which we used as a point of comparison with the corresponding PIP. We included a buffer of 5 ignored windows (∼250,000 kb) on either side of each PIP in a chromosome. In a few cases (*n* = 6), we were unable to identify an appropriate COMP region because the putative inversion polymorphisms spanned too great a proportion of the chromosome or was mapped onto the chromosome in a complex way. We used PCAngsd to create a PCA of each PIP and each COMP (Fig. S1, *A3*). If the PIP PCA contained at least two clearly definable clusters on PC1 (Fig. S1, *D3,* Dataset S2) that were not related to geographic structure or sex, we proceeded with further analysis.

Finally, we used ngsLD (Fox et al. 2019) on ANGSD-generated genotype likelihoods to calculate linkage disequilibrium across each chromosome with a PIP (Fig. S1, *A4,* Dataset S2). Inversions are expected to create a pattern where elevated LD can be detected in analyses involving individuals with a mix of genotypes, but not in analyses involving a subset of individuals that all share the same homozygous genotype (Huang et al. 2020; Mérot 2020; Harringmeyer and Hoekstra 2022). That is, we expect LD estimates from an inversion polymorphism to show relative differences depending on whether LD is calculated with a set of individuals showing mixed inversion genotypes compared to calculations from a subset of homozygotes that all share the same inversion haplotype. Calculating LD across an entire chromosome is extremely computationally intensive, so it is necessary to down-sample SNPs before performing these calculations. The program ngsLD is designed for use with low-coverage datasets, but in our experience, random down-sampling of SNPs to a computationally-feasible level (e.g., 3000 SNPs, which requires 9,000,000 LD calculations) can produce datasets that are too noisy to produce meaningful results, even in cases where there is very strong evidence of inversion polymorphism from other analyses. We maximized our ability to detect elevated LD in a PIP, should it exist, by selecting non-random subsets of SNPs using the “SNP weights” output of PCAngsd, which reflects the contribution of each SNP to the genetic covariance matrix underlying the PCA. For each chromosome, we identified the 3000 SNPs with highest absolute value SNP weights from separate PCA of the PIP and COMP regions, and we applied ngsLD to this non-random subset and plotted the results. Selecting SNPs with high PCAngsd weights certainly biases the estimates of LD we obtain; however, doing so should not create bias in the *relative* LD displayed by putatively homozygous samples compared with mixed-genotype samples, which is the main aim of this analysis. After generating LD estimates, we plotted resulting LD matrices in R as rasters after aggregating them by a factor of 20 with the function ‘aggregate()’ from the R package ‘terra’ (Hijmans 2024). NA values in LD matrices were ignored during aggregation. We visually assessed LD plots to determine whether patterns of recombination suppression support the hypothesis that PCA clustering patterns associated with the PIP were caused by an inversion (Fig. S1, *D6,* Dataset S2).

In some cases, plots of ngsLD results indicated the existence of a potential inversion within the COMP region (Fig. S1, *D4*). When this occurred, we re-examined lostruct results to identify whether the newly identified region corresponded with any spikes on MDS2, 3, or 4 (Fig. S1, *D5*). If so, we added the newly identified region to our set of PIPs and analyzed it as described above (starting from Fig. S1, *A2*).

### Assigning PIP genotypes to individuals

We used PCAngsd plots to assign putative genotypes to individuals based on PCA clusters (Dataset S4). For regions showing 3 clusters on PCA plots (the majority of identified PIPs, *n* = 133), we assigned individuals to the middle cluster as heterozygotes (“AB”) and the individuals in the right and left clusters as homozygotes (“AA” or “BB”, where “AA” is the more frequent genotype) (Huang et al. 2020; Mérot 2020). It is less clear how to assign specific genotypes within regions that show 2 clusters or 4-6 clusters (*n* = 41 PIPs). Regions that cluster into 2 groups may represent polymorphisms where one of the genotypes is lethal or exceedingly rare, as is the case in the well-studied *Z. albicollis* inversion (Tuttle et al. 2016). The presence >3 clusters along PC1 and PC2 may indicate the involvement of more than two inversion alleles. Since we cannot unambiguously identify which clusters represent homozygotes vs heterozygotes in these cases, we arbitrarily designate their genotypes using the labels “C” and “D” for 2-cluster regions and “C” through “H” for regions with up to 6 clusters.

### Characteristics of identified polymorphic regions

We evaluated the length and average heterozygosity of each PIP region and also estimated average heterozygosity from COMP regions for comparison. We estimated the mapped length of the polymorphism by calculating the distance between the start and end of the PIP that we identified using changepoint analysis on lostruct output (described above). To estimate heterozygosity, we used ANGSD to generate site allele frequencies for all loci (including invariant loci) within each PIP or COMP region separately. We then used winsfs v0.7.0 (Rasmussen et al. 2022) to estimate a 1d site frequency spectrum (SFS) for each individual at each PIP and COMP, where heterozygosity is the number of sites in the second SFS bin divided by the total number of sites in the SFS.

We used boxplots to visualize how average heterozygosity varies across genotypes for each PIP and its corresponding COMP (Dataset S2). For PIPs with 3 clusters, we evaluated whether heterozygosity was significantly elevated in putative heterozygotes over both classes of putative homozygotes, or whether heterozygosity was significantly suppressed in one class of homozygotes compared with the other two genotypes (Mérot 2020). To make these comparisons, we used the R package ‘emmeans’ (Lenth 2023) to apply post-hoc significance testing to linear models where genotype was the predictor and heterozygosity the response variable. Finally, we also used emmeans to test for significant differences in heterozygosity between the “C” and “D” genotypes in PIPs with 2 clusters.

### Population-level patterns of putative inversion polymorphism

We describe genotype frequencies in all PIPs, and 3 additional patterns based on haplotype frequencies in PIPs presumed to be biallelic—those with 3 observed PCA clusters (assigned to genotypes AA, AB, BB). We are not able to confidently infer haplotype frequencies in PIPs with 2 or more than 3 PCA clusters. For each PIP, we calculated the frequency of each genotype and we visualized how genotypes vary in continuous space by plotting samples colored by their genotype across the sampling region (Dataset S2). Using biallelic PIPs, we calculated minor haplotype frequency, tested whether the PIP deviated from Hardy-Weinberg Equilibrium, and estimated a slope of isolation by distance in geographic space. Minor haplotype frequency is analogous to minor allele frequency, where individuals are assigned a minor haplotype count of 2 (for individuals in the less-common class of homozygotes), 1 (for heterozygotes), or 0 (for the more-common class of homozygotes), and the corresponding sum is divided by the total number of haplotypes in the sample (i.e., 2*n* for these diploid organisms) to calculate the frequency. We tested for deviation from Hardy-Weinberg Equilibrium using the function ‘HWExact’ in the R package ‘HardyWeinberg’ v1.7.8 (Graffelman 2015) with an alpha-level of 0.05. Finally, to evaluate spatial structure, we visually assessed plots of genotype frequencies in space (Dataset S2) and we calculated isolation by distance by estimating the linear slope of pairwise PIP haplotype distance (the number of haplotypes shared by a pair of samples, which can be 0, 1, or 2) and pairwise geographic distance between individuals.

### Comparative analysis of putative inversion polymorphisms across multiple species

While analyzing many species at once, we noted many instances where different species mapped to the same reference genome showed evidence of inversion polymorphism on the same chromosomes (*n =* 24 chromosomes where PIPs occur in at least 2 species; Dataset S1). To further evaluate these PIPs, we created genetic distance-based neighbor-joining trees for each potentially homologous PIP and, when possible, an associated comparison (“COMP”) region (Dataset S3). Each tree comprised all species mapped to the corresponding reference genome, regardless of whether they showed a PIP on the corresponding chromosome, and included conspecific representatives from multiple PIP genotypes. We expected that trees made using independently-formed PIPs should show branching pattern consistent with species-level relationships shown by the rest of the genome (as represented by the tree from the COMP region). In other words, because our study species are relatively well diverged, the general expectation is that conspecifics should be more closely related to each other than they are to heterospecifics. PIP trees showing alternative branching patterns, whereby individuals are grouped by PIP genotype instead of by species, are consistent with scenarios where the polymorphism predates speciation or has been shared across species through introgression (Fontaine et al. 2015; Knief et al. 2024).

Out of 24 chromosomes with PIPs in multiple species, 4 contained PIPs that mapped to completely non-overlapping positions across different species, which presumably represent totally independent polymorphisms. Each of the remaining 20 chromosomes contained one or more PIPs that at least partially overlapped across multiple species (*n* = 26 overlapping PIPs). We identified the largest possible region that overlapped all identified PIPs, which we term a “shared PIP” region. In many cases the width of PIPs varied somewhat across species, such that the length of the shared PIP regions used in this comparative genetic distance analysis was often smaller than that of the species-level PIPs. We also selected new COMP regions for this analysis by choosing an arbitrary locus in the chromosome that was the same length as the shared PIP and overlapped with no PIPs in any species.

After identifying shared PIP and shared COMP regions, we created gene trees using representative individuals selected from each species (Dataset S5). For species that did not show evidence of a PIP on the chromosome, we selected one individual randomly. For species involved in the shared PIP, we randomly selected one individual from each identified genotype, except that we excluded putatively heterozygous (“AB”) individuals.

We then calculated pairwise genetic distance among the selected individuals for each shared PIP and shared COMP using a custom script adapted from https://github.com/marqueda/PopGenCode/tree/master (Marques et al. 2018). These calculations estimate genetic distance as the expected number of pairwise nucleotide differences between individuals, inferred in a genotype likelihood framework. We first used ANGSD to estimate site allele frequencies for all loci within a given region, for each pair of individuals in the analysis. We then used winsfs on these pairwise site allele frequency files to estimate a 2d SFS for each pair of individuals. Whereas the custom script from (Marques et al. 2018) estimates genetic distance across genome windows based windowed estimates of the 2d SFS, we used an adapted version of the script to generate a single estimate of genetic distance for the 2d SFS that represents the entire region of interest. Finally, we used a matrix of pairwise genetic distances with the R function ‘nj()’ from the package ‘ape’ v5.7-1 (Paradis and Schleip 2019) to plot a neighbor-joining tree.

## Acknowledgements

For helpful suggestions and insights, we thank the Winger and Bradburd labs at the University of Michigan, as well as Peter Ralph, Jun Ishigohoka, Tim Sackton, Emma Berdan, and Brian Charlesworth. Nicole Adams shared custom code for estimating genetic distance. For providing samples, we thank the curators, collections staff, and field collectors from the following institutions: American Museum of Natural History, Cleveland Museum of Natural History, Cornell University Museum of Vertebrates, New York State Museum, Royal Alberta Museum, Royal Ontario Museum, University of Alaska Museum of the North, University of Minnesota Museum of Natural History, University of California, Berkeley Museum of Vertebrate Zoology, University of Michigan Museum of Zoology, University of Washington Burke Museum. For additional samples, we thank J. Tremblay (Environment and Climate Change Canada). Next-generation sequencing for this project was partially carried out in the Advanced Genomics Core at the University of Michigan. This research was also supported in part through computational resources and services provided by Advanced Research Computing (ARC), a division of Information and Technology Services (ITS) at the University of Michigan, Ann Arbor.

## Funding

National Science Foundation grant 2146950 (BMW)

National Science Foundation Graduate Research Fellowship DGE 1256260, Fellow ID 2018240490 (TMP)

National Science Foundation Postdoctoral Research Fellowships in Biology Program Grant No. 2410085 (TMP)

University of Michigan Rackham Graduate Student Research Grant (TMP)

## Data Availability Statement

The sequence data underlying this article are available on the NCBI Sequence Read Archive and can be accessed with BioProject numbers PRJNA1043688 and PRJNA1130443. Other relevant data will available in the supplementary materials upon publication.

## Notes

### Competing Interest Statement

The authors have declared no competing interest.

