## Supplementary material for "Large inversion polymorphisms are widespread in North American songbirds": Dataset S2

**Dataset S2**. For each putative inversion polymorphism (PIP), we show 8 plots generated by the analyses we used to identify PIPs (Fig. S1) and to evaluate some of their characteristics. The species and name of the PIP is given at the top of each 8-plot page. A. Analysis with lostruct (Fig. S1, *A1*) indicates which segments of chromosomes have distinct population structure, as identified by multidimensional scaling (MDS) (Li & Ralph 2019). The highlighted red region is identified as a PIP (Fig. S1, *A2*). Panel A shows MDS1 values along each chromosome. B. MDS2 values from lostruct along each chromosome. In some cases, PIPs may be highlighted on MDS2 and not MDS1, particularly when more than one PIP is present on a chromosome. C. PCA plots from each PIP region (Fig. S1, *A3)* show distinct clustering patterns consistent with inversion (Mérot 2020). Genotypes are assigned to individuals based on these clusters. In cases where PCA produces 3 clusters, individuals in the central cluster are assumed to be heterozygotes for the polymorphism (AB genotype), while those in the right and left clusters are assumed to be homozygotes (AA and BB genotypes; (Mérot 2020)). Otherwise, individuals are assigned arbitrary genotype codes (C-H, depending on the number of clusters). D. The cluster pattern is not present in the comparison (COMP) region of the chromosome, where no inversion polymorphisms are thought to be present. E. Boxplots show individual heterozygosity values from inside the PIP region and inside a comparison (COMP) region in the same chromosome. F. A waffle plot shows PIP genotype frequencies of the sampled population. Genotypes are assigned based on PCA (see panel C). G. Linkage disequilibrium (LD) is plotted from a subsample of SNPs in the PIP and COMP regions (Fig. S1, *A4*). Inversions are expected to produce a pattern where LD in a sample of individuals with both classes of homozygous genotypes (upper triangle) is elevated in the PIP region compared with the COMP region (Mérot 2020). This pattern should not exist in sample of individuals that all share the same genotype (lower triangle). PIPs that meet these expectations are considered "LD-confirmed" PIPs. H. Individuals are shown in geographic space, colored by their PIP genotype, illustrating the spatial distributions of the genotypes. Individual locations are jittered to allow all points to be seen across the range (actual sampling locations can be seen in Fig. 1A). Note that in some cases, we were unable to identify a suitable COMP region from the same chromosome as a PIP due to the size and/or complexity of the PIP, and COMP data are therefore not shown in panels D and E in these cases. Chromosomes that have more than one PIP on them have only a single COMP region (if any).

*Sphyrapicus varius* CM026025.1

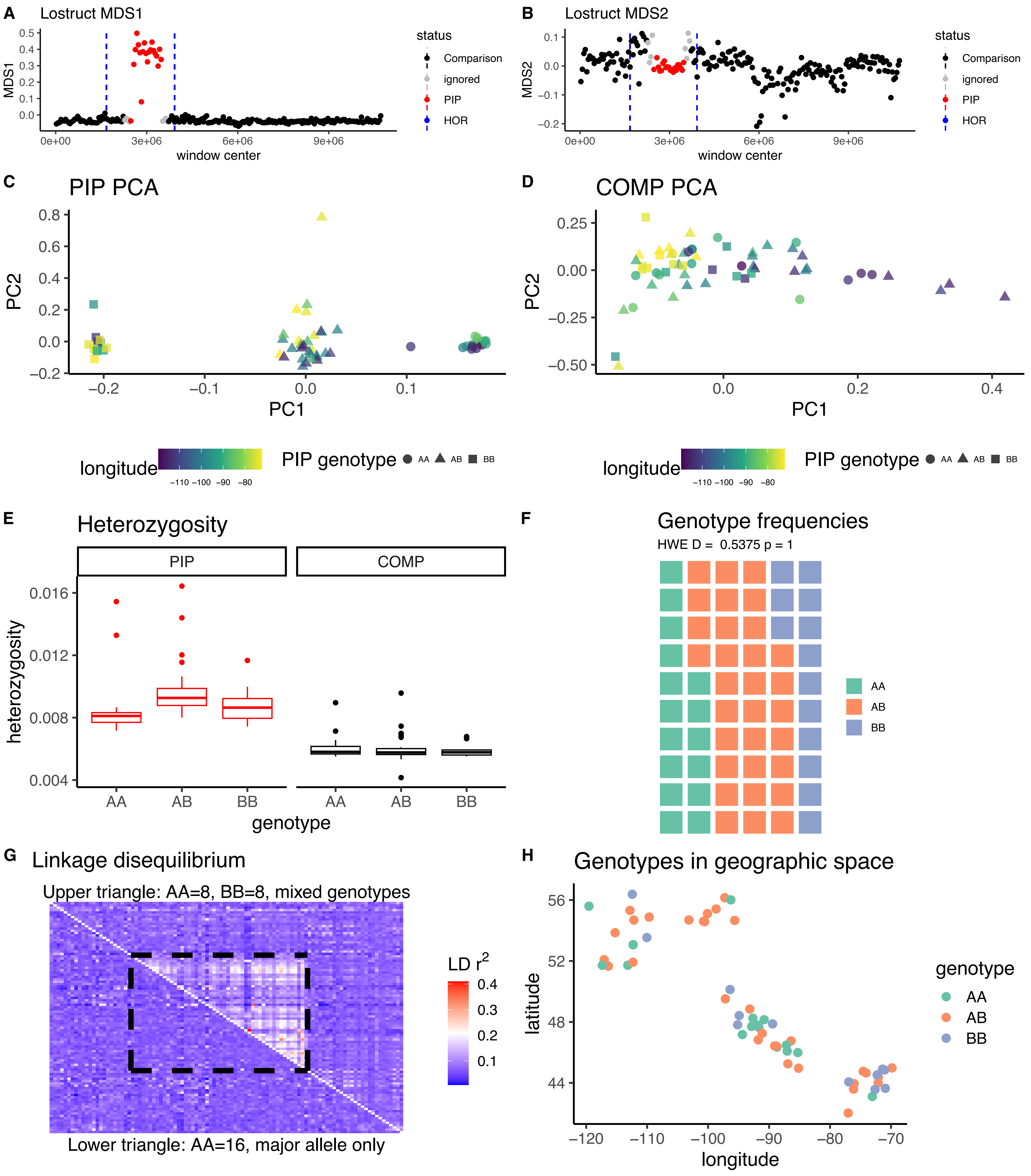

*Sphyrapicus varius* CM026013.1

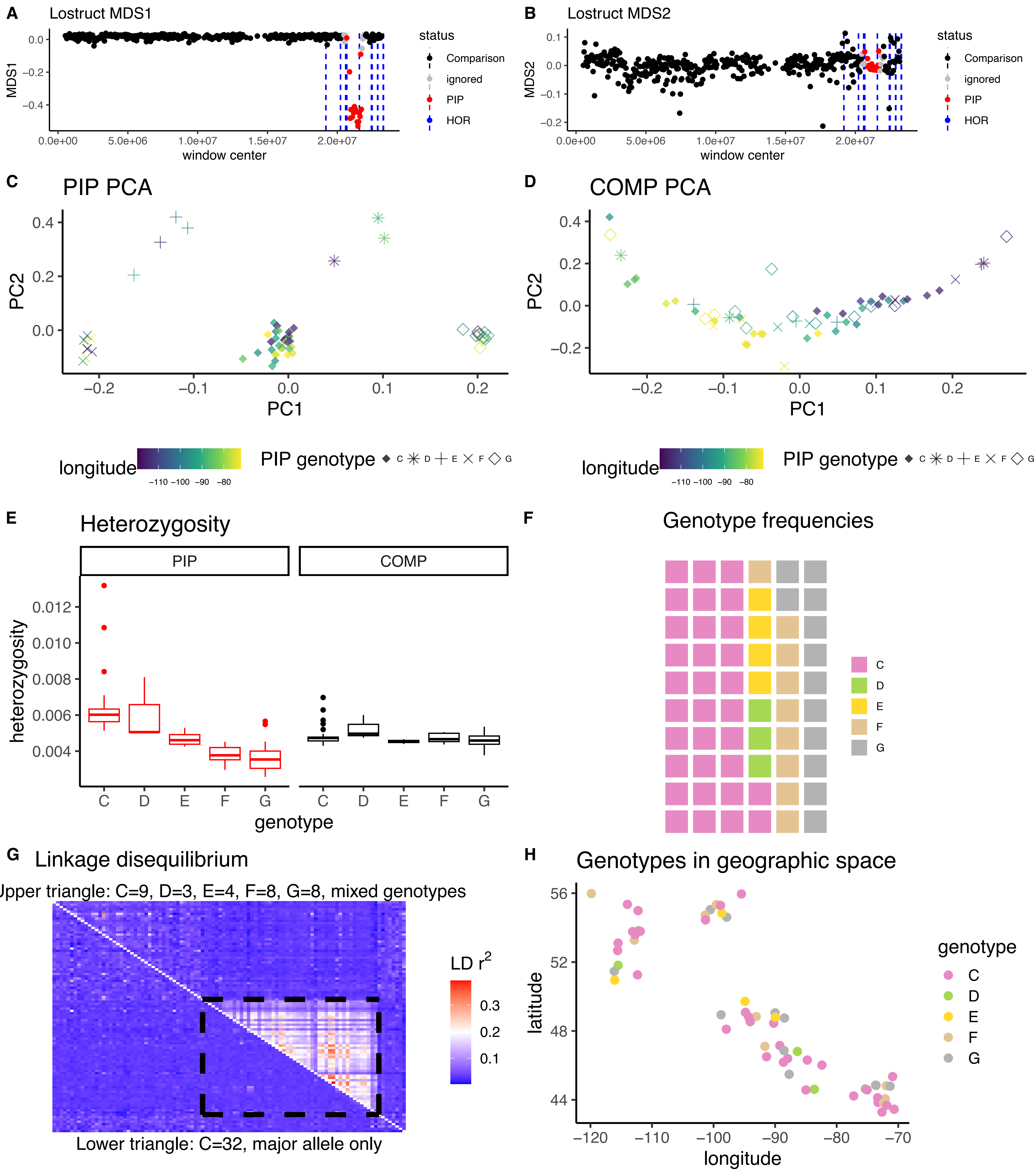

*Sphyrapicus varius* CM026000.1A

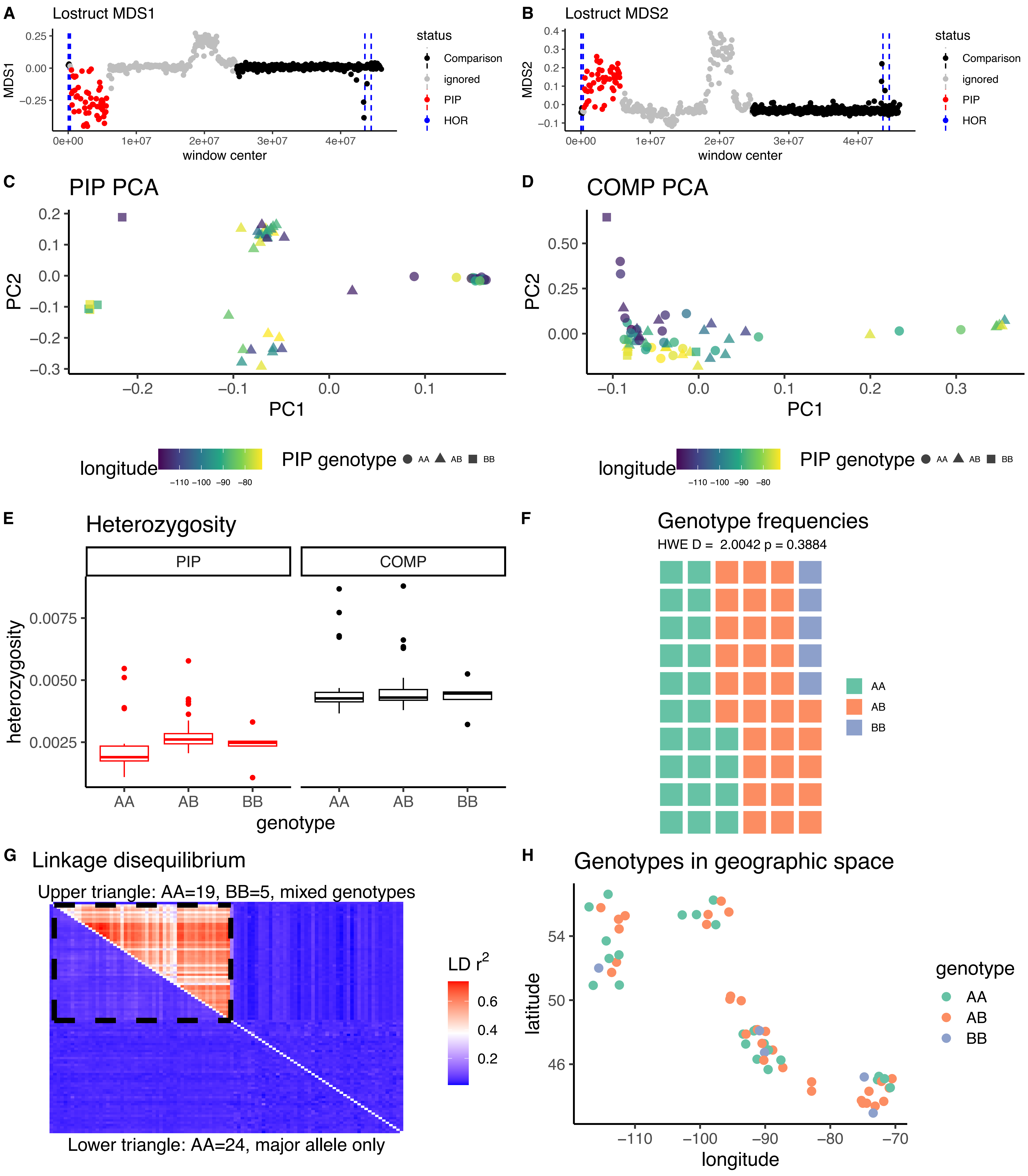

*Sphyrapicus varius* CM026000.1B

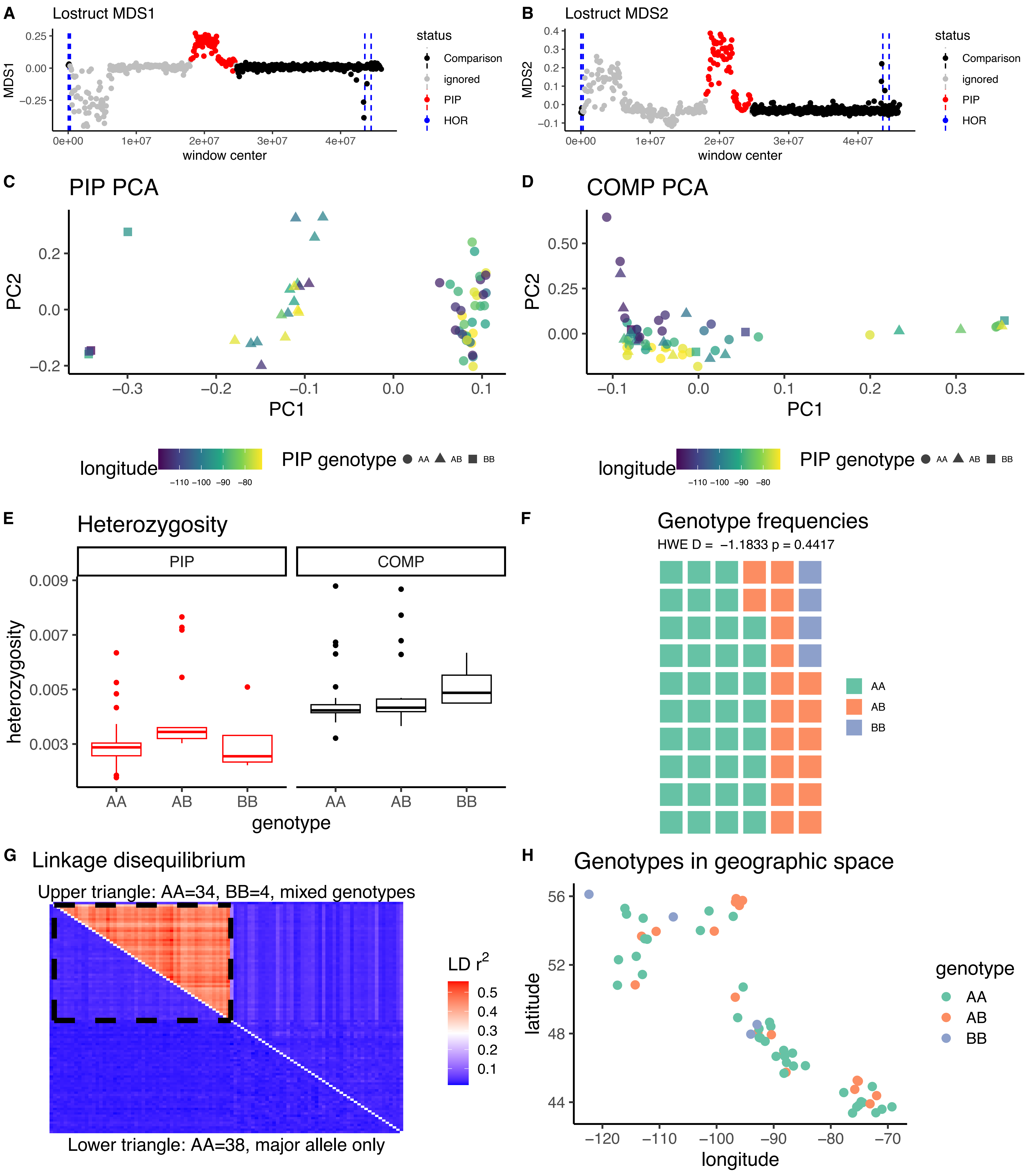

*Sphyrapicus varius* CM025996.1A

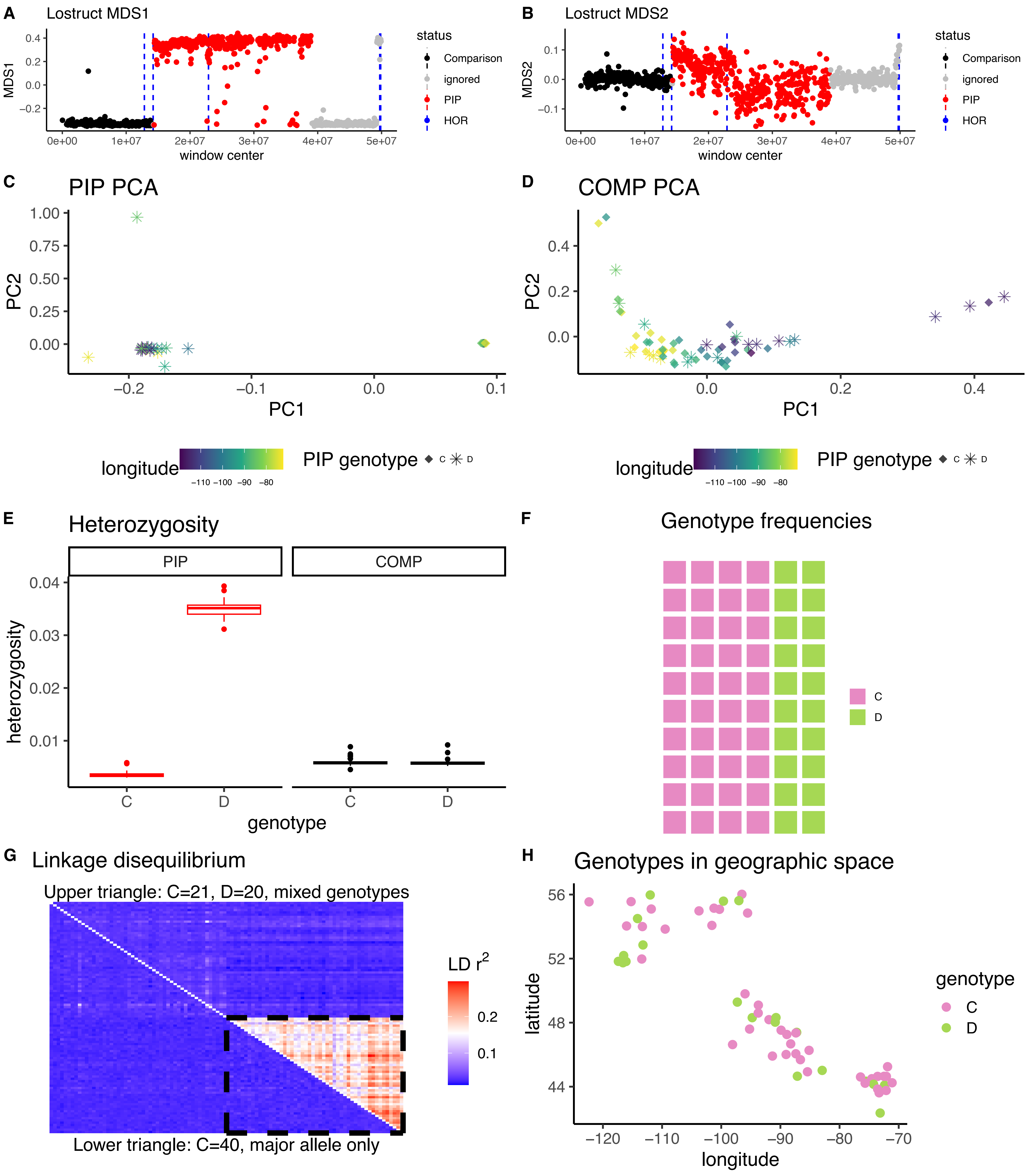

*Sphyrapicus varius* CM025996.1B

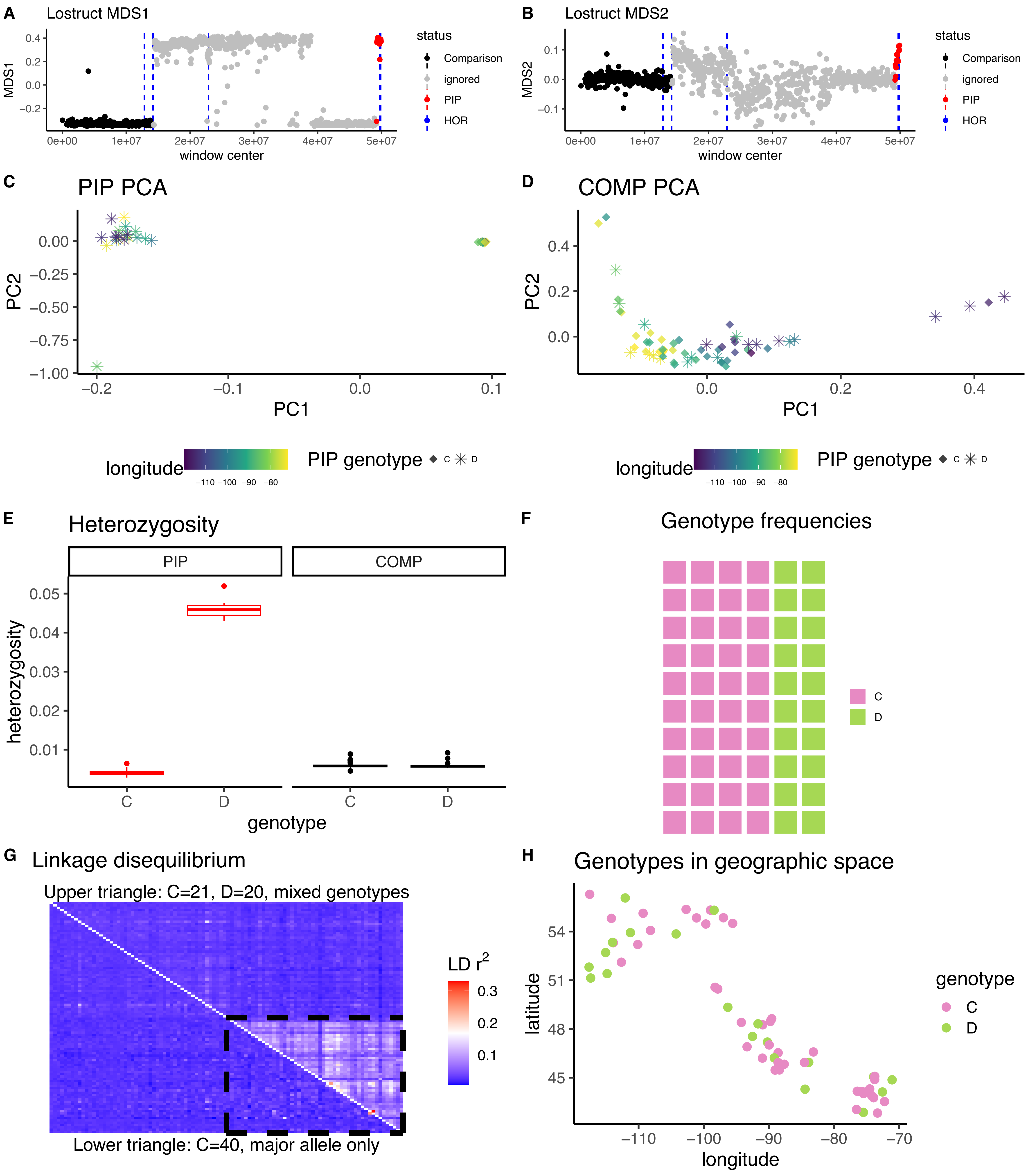

*Sphyrapicus varius* CM025994.1

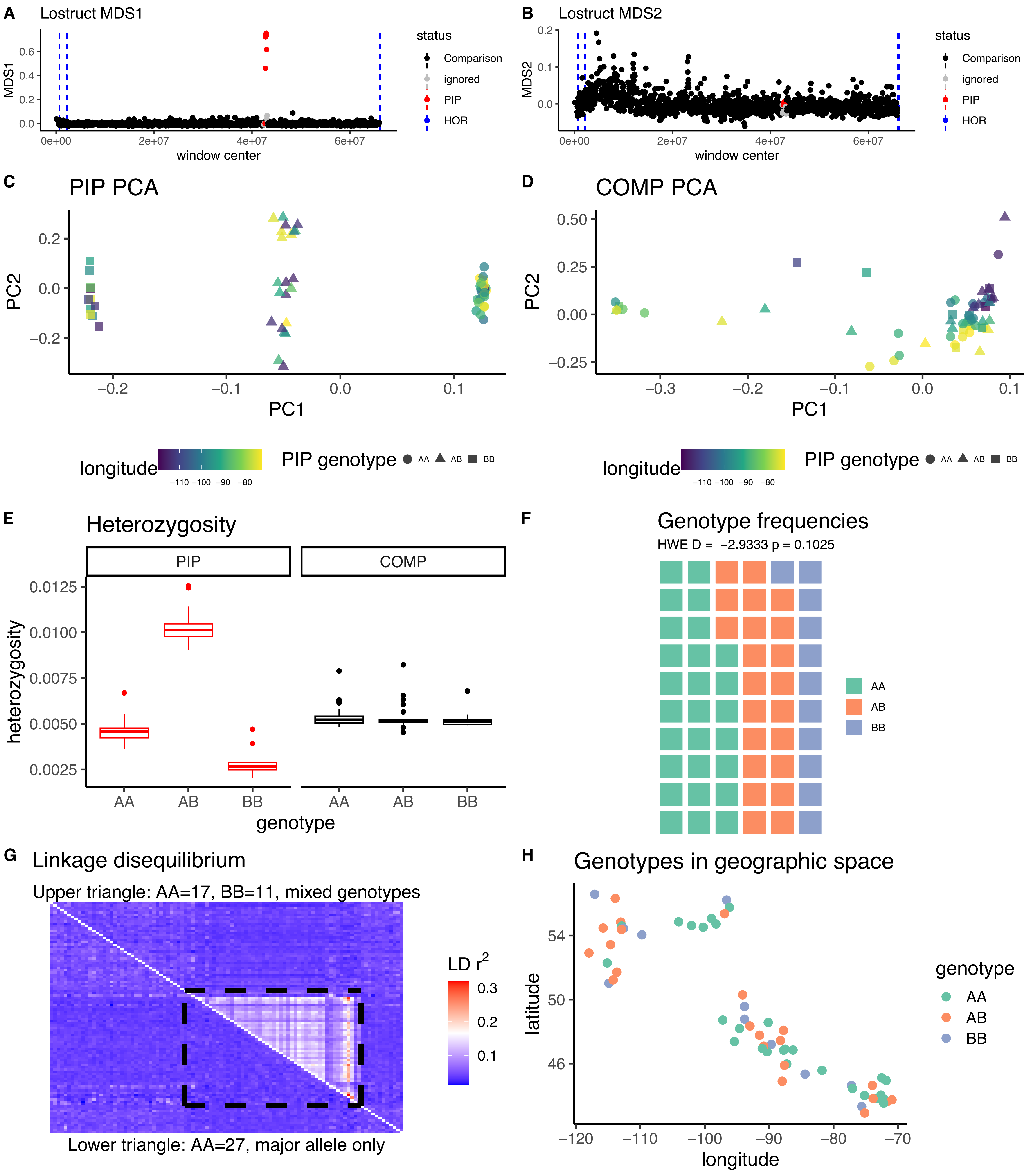

*Empidonax alnorum* CM020534.1

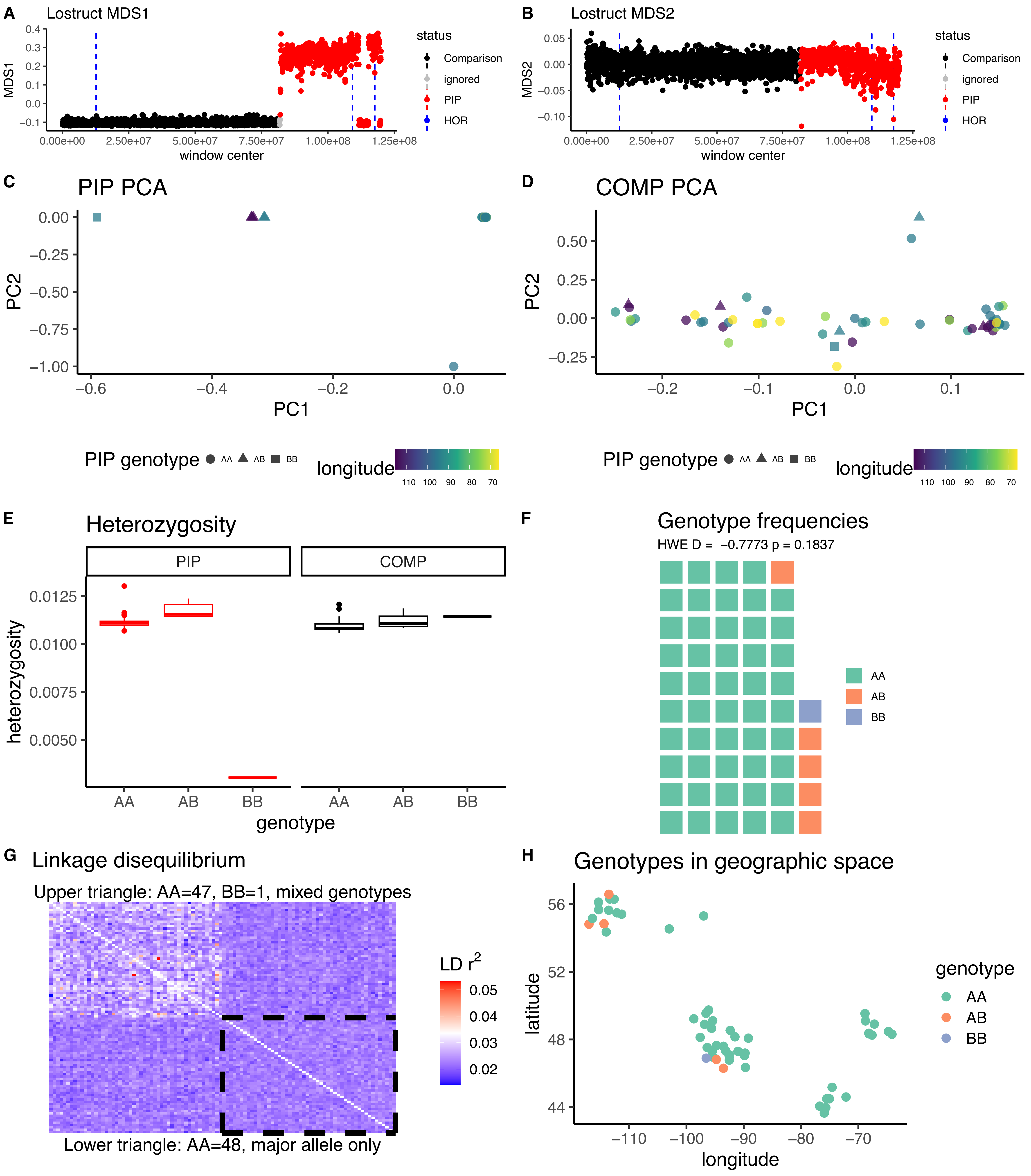

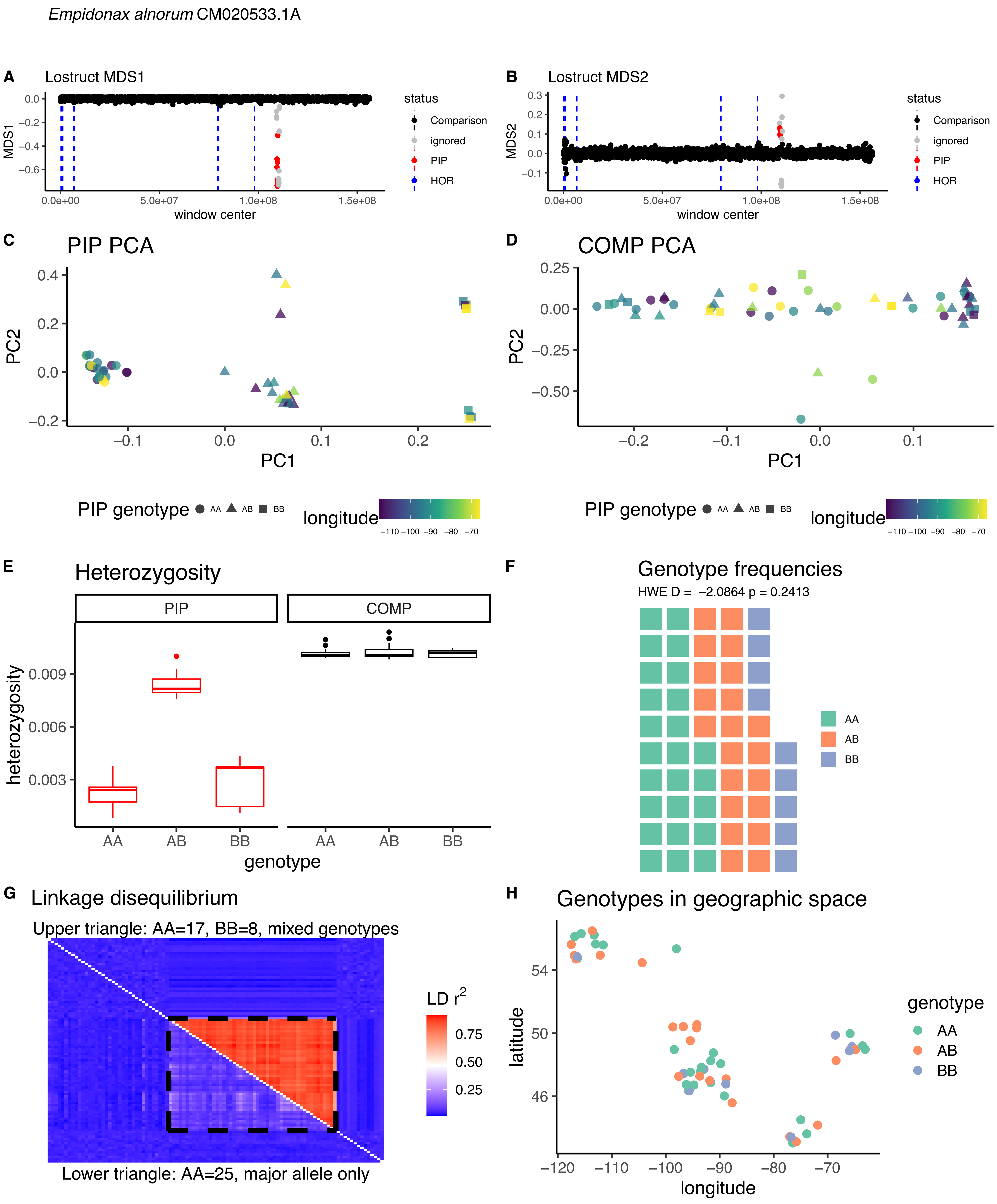

*Empidonax alnorum* CM020533.1A
A
Lostruct MDS1
MDS1
window center
status
Comparison
ignored
PIP
HOR
B
Lostruct MDS2
MDS2
C
PIP PCA
PC1
PC2
D
COMP PCA
PIP genotype AA AB BB
longitude
E
Heterozygosity
PIP
COMP
heterozygosity
genotype
F
Genotype frequencies
HWE D = −2.0864 p = 0.2413
AA
AB
BB
G
Linkage disequilibrium
Upper triangle: AA=17, BB=8, mixed genotypes
Lower triangle: AA=25, major allele only
LD r2
H
Genotypes in geographic space
latitude
longitude
genotype

*Empidonax alnorum* CM020533.1B

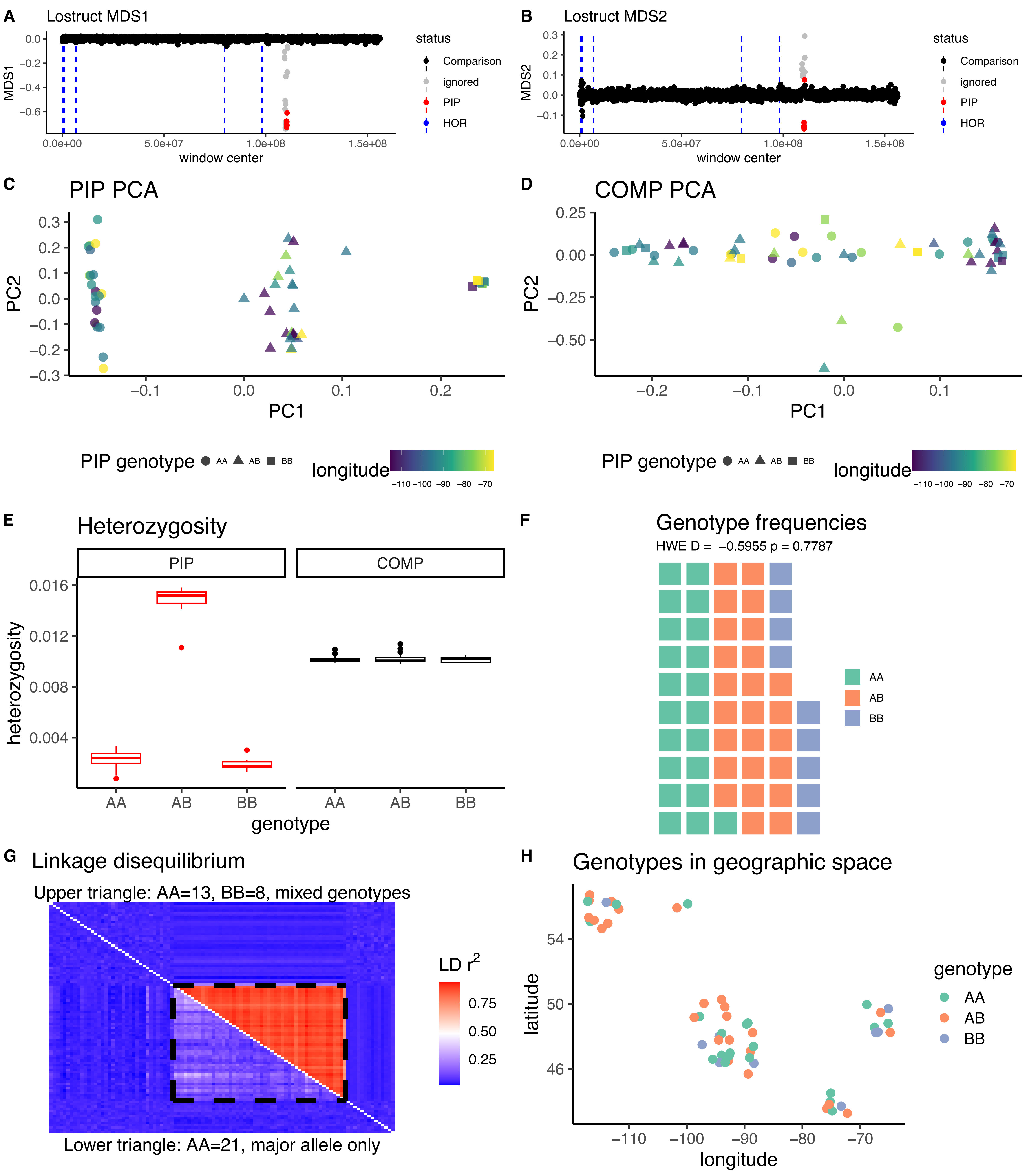

*Empidonax flaviventris* CM020543.1A

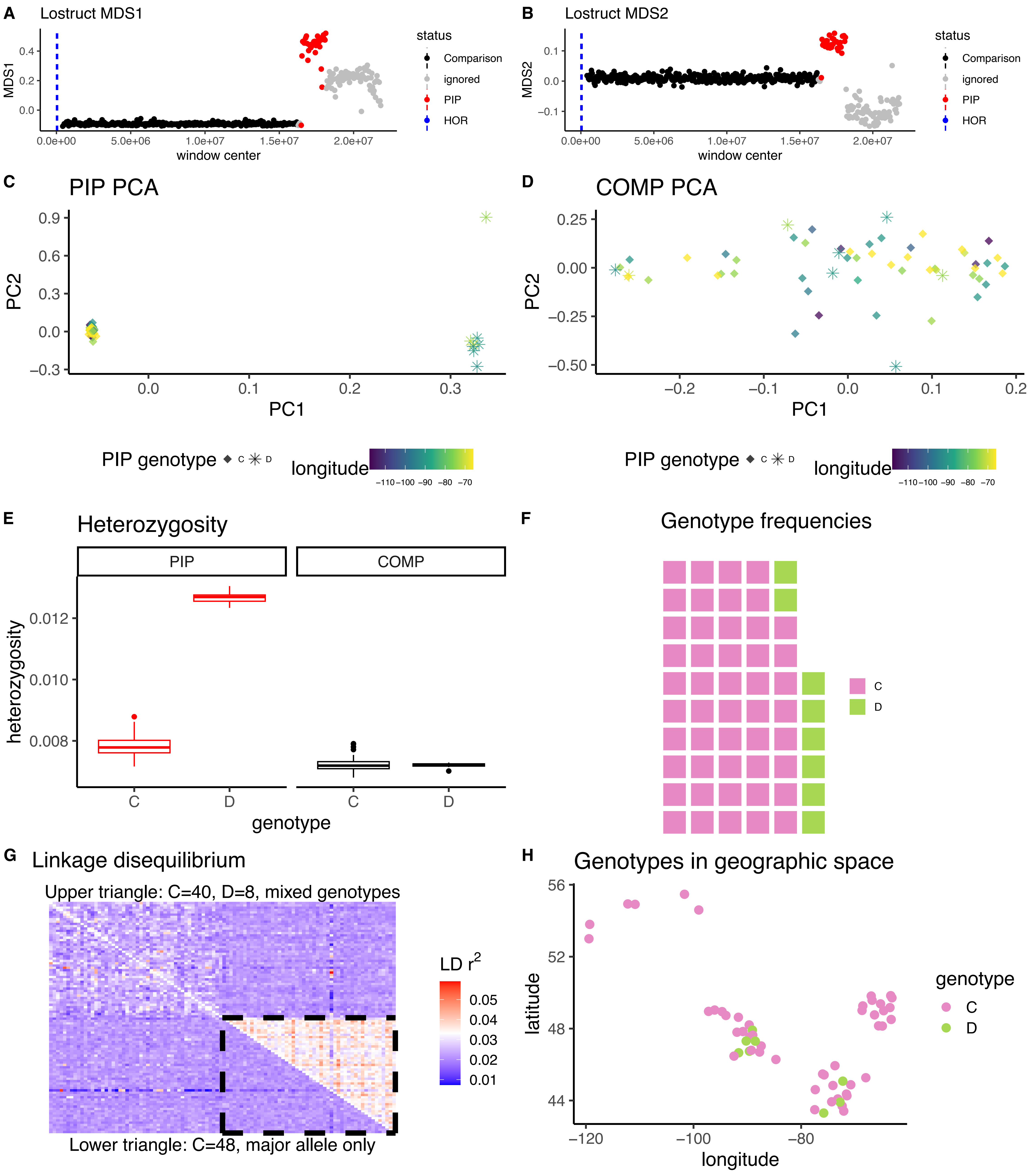

*Empidonax flaviventris* CM020543.1B

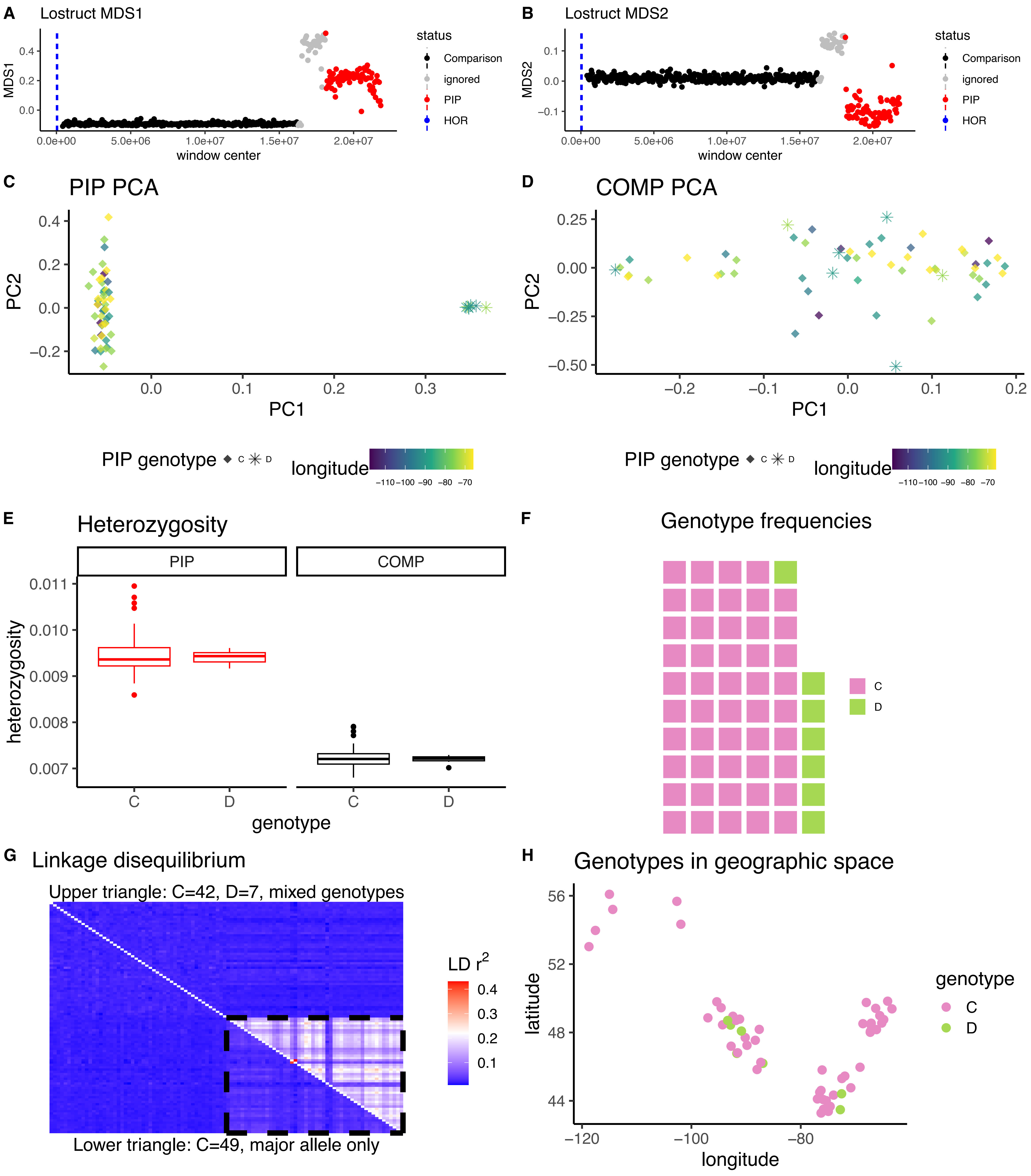

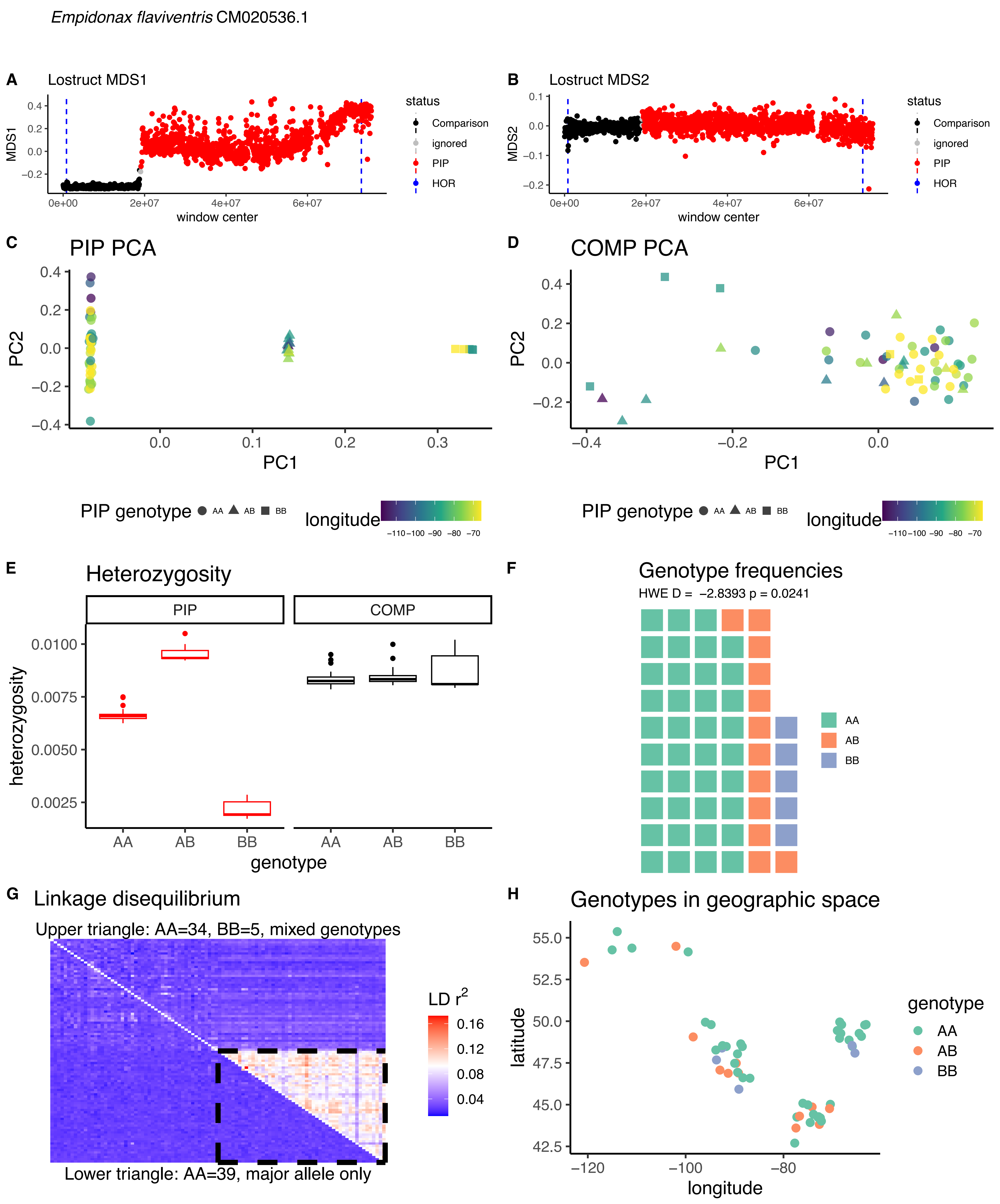

*Empidonax flaviventris* CM020536.1
A
Lostruct MDS1
MDS1
window center
status
Comparison
ignored
PIP
HOR
B
Lostruct MDS2
MDS2
window center
C
PIP PCA
PC1
PC2
D
COMP PCA
PC1
PC2
PIP genotype AA AB BB
longitude
E
Heterozygosity
PIP
COMP
heterozygosity
genotype
F
Genotype frequencies
HWE D = -2.8393 p = 0.0241
AA
AB
BB
G
Linkage disequilibrium
Upper triangle: AA=34, BB=5, mixed genotypes
LD r$^2$
Lower triangle: AA=39, major allele only
H
Genotypes in geographic space
latitude
longitude
genotype
AA
AB
BB

*Empidonax flaviventris* CM020534.1

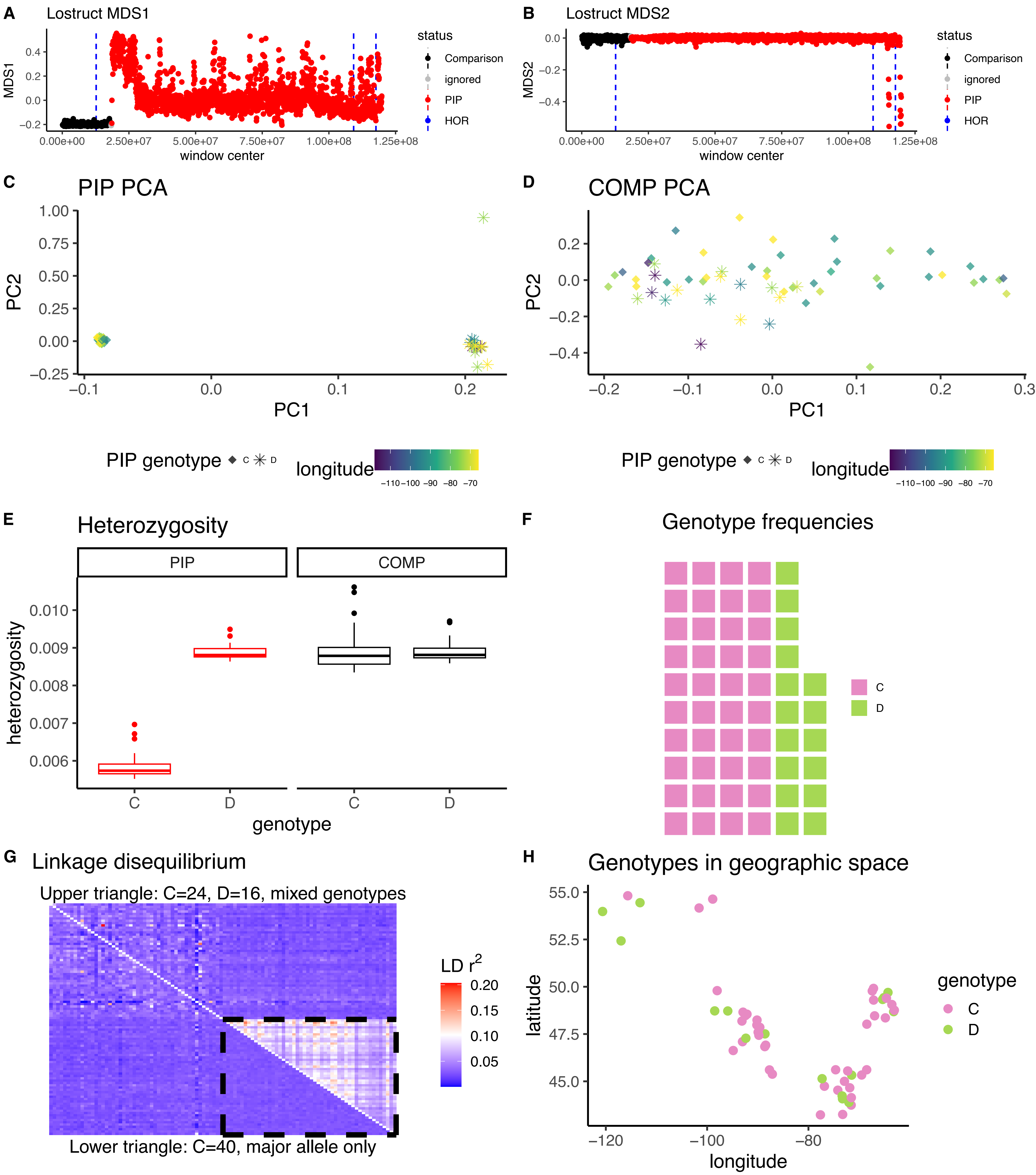

*Empidonax flaviventris* CM020533.1

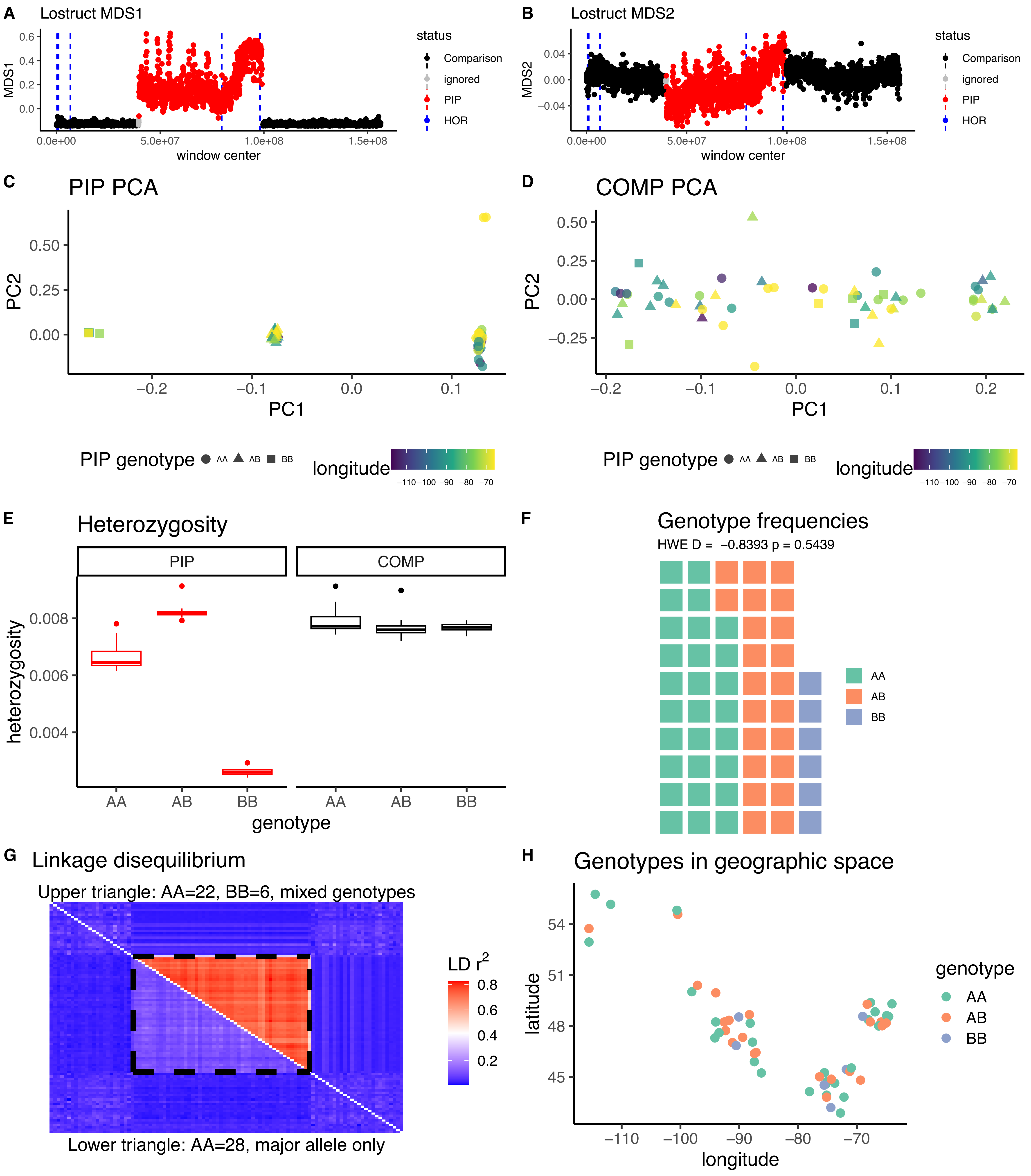

*Empidonax minimus* CM020547.1

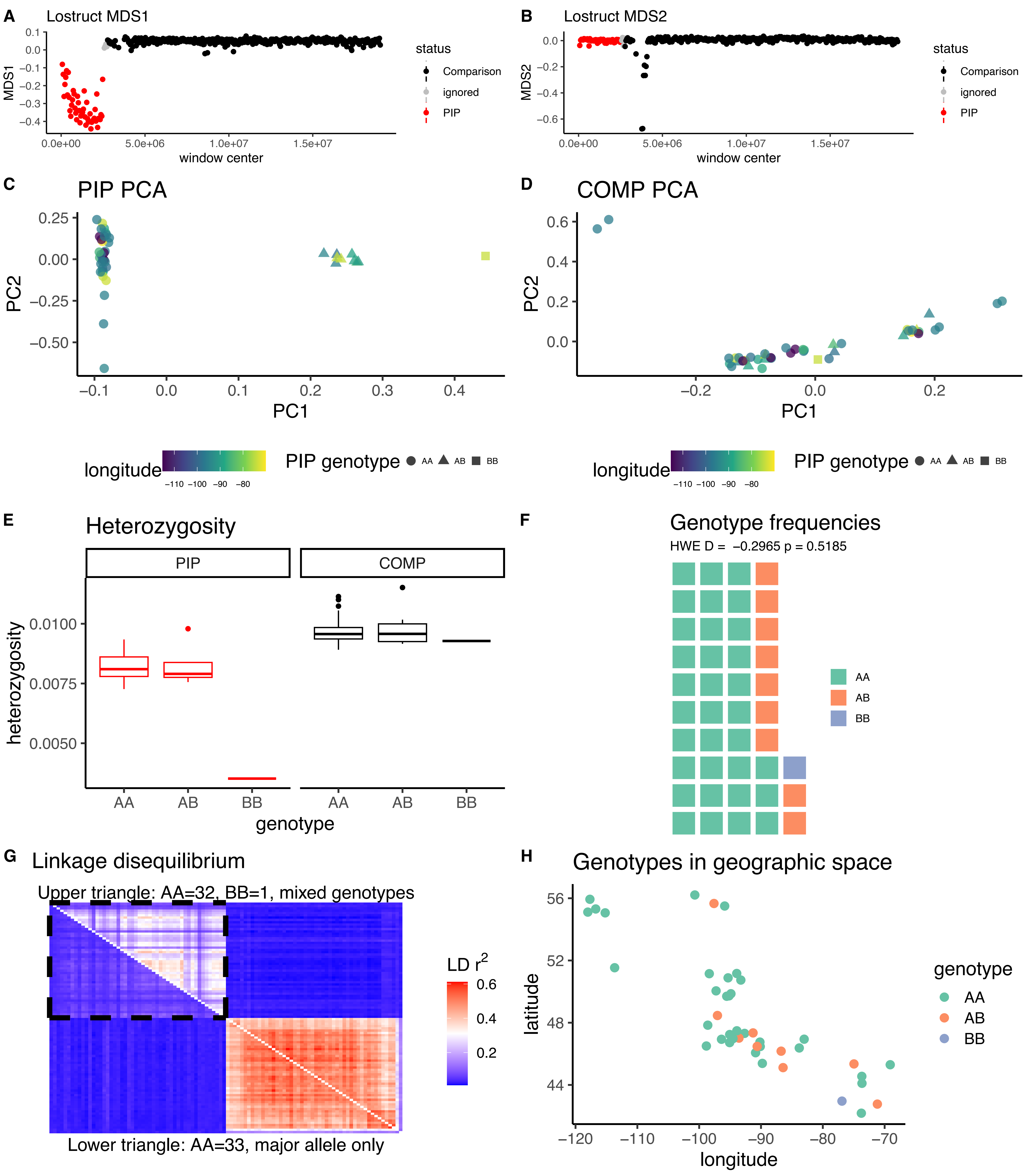

*Empidonax minimus* CM020545.1

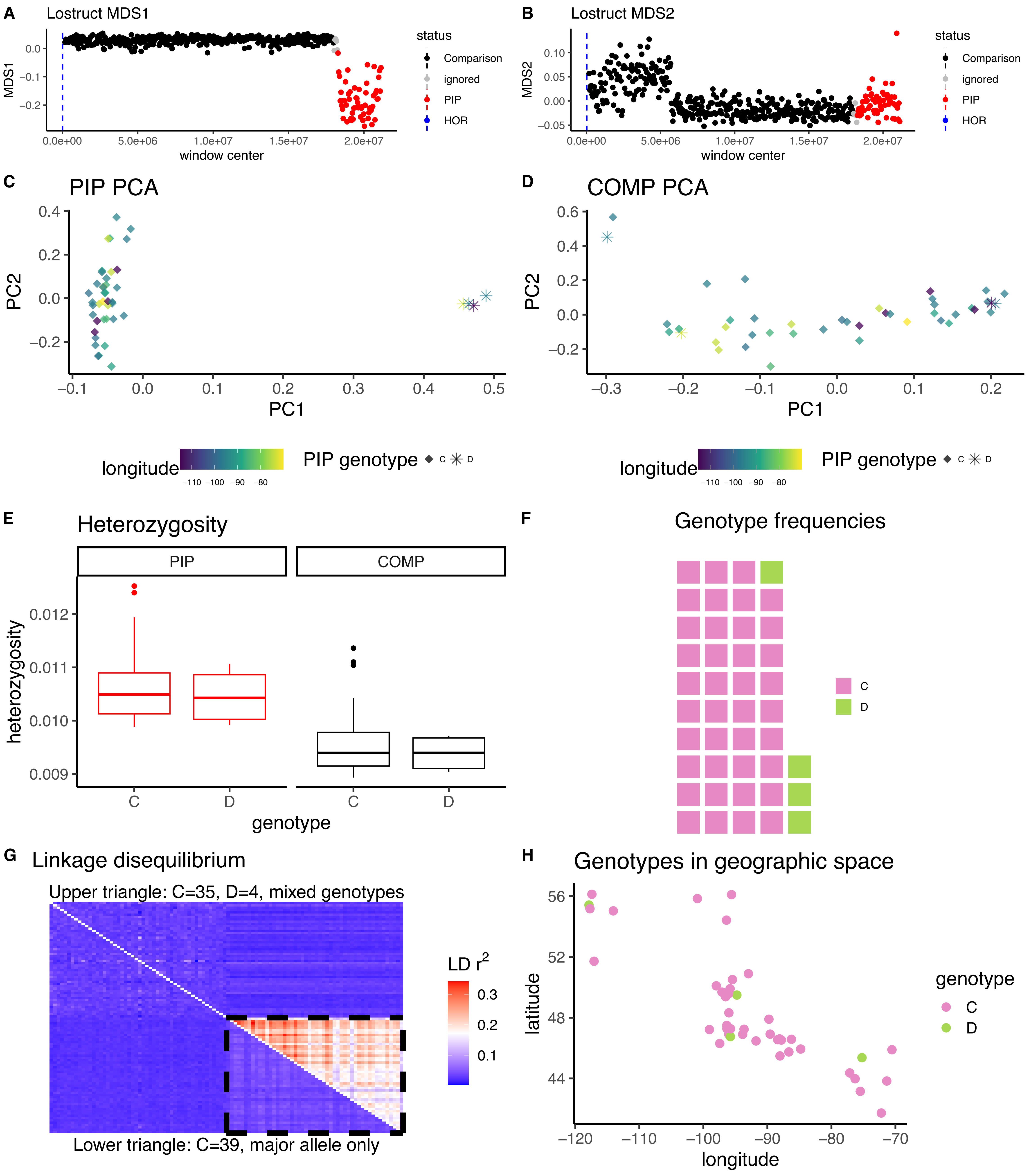

*Empidonax minimus* CM020543.1B

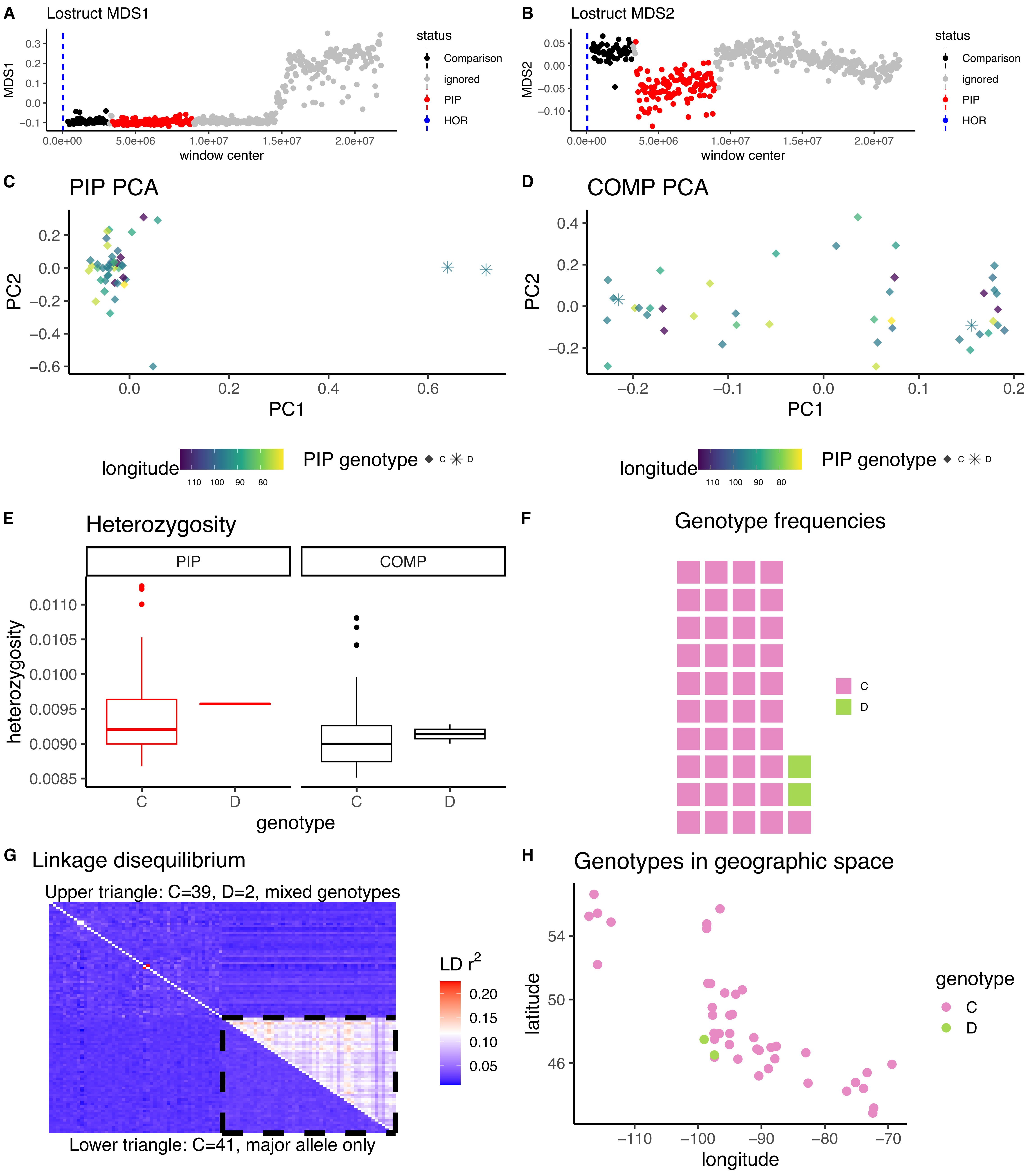

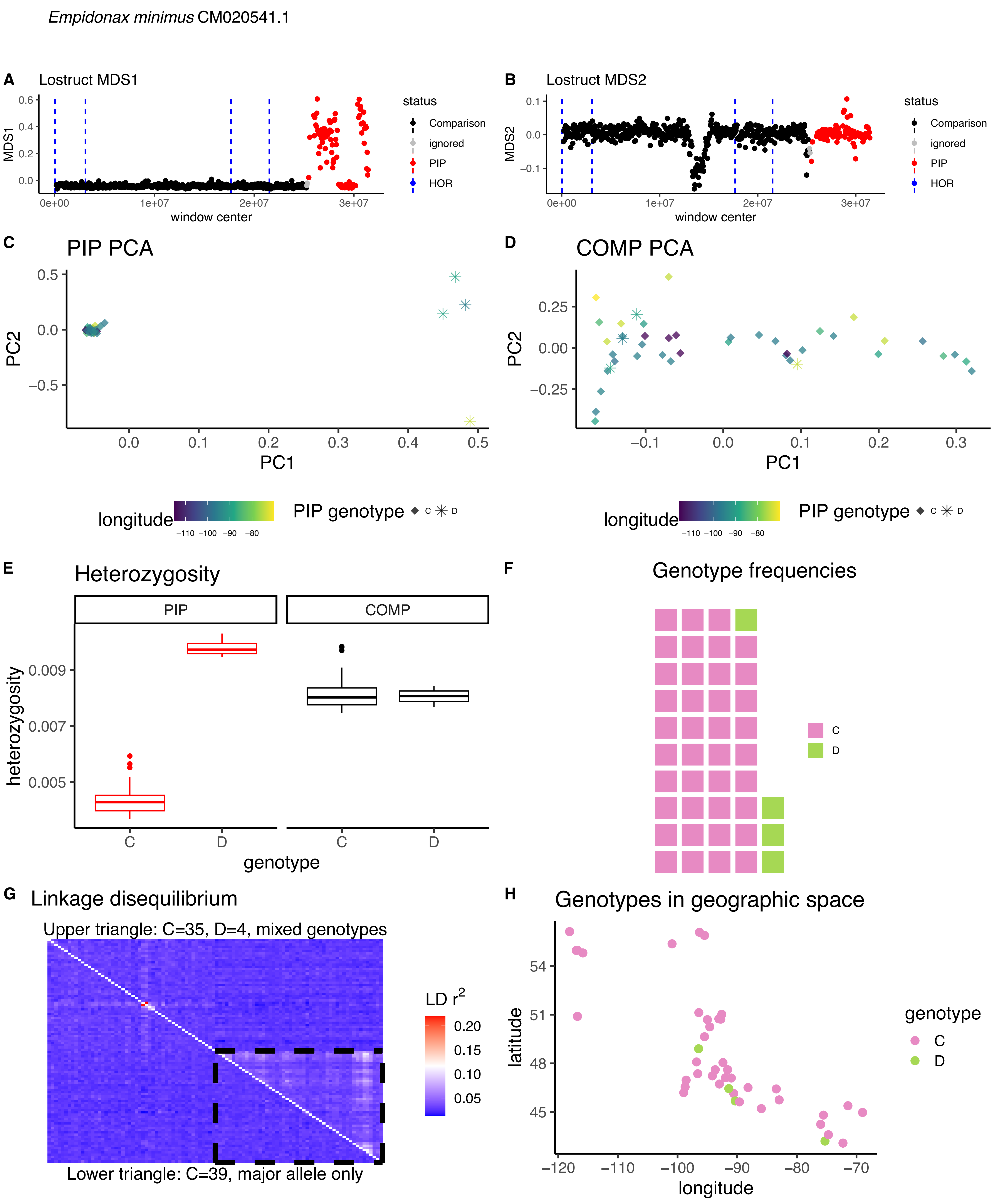

Empidonax minimus CM020541.1
A
Lostruct MDS1
MDS1
window center
status
Comparison
ignored
PIP
HOR
B
Lostruct MDS2
MDS2
C
PIP PCA
PC1
PC2
D
COMP PCA
longitude
PIP genotype
E
Heterozygosity
PIP
COMP
heterozygosity
genotype
F
Genotype frequencies
C
D
G
Linkage disequilibrium
Upper triangle: C=35, D=4, mixed genotypes
Lower triangle: C=39, major allele only
LD r2
H
Genotypes in geographic space
latitude
longitude
genotype

*Empidonax minimus* CM020539.1A

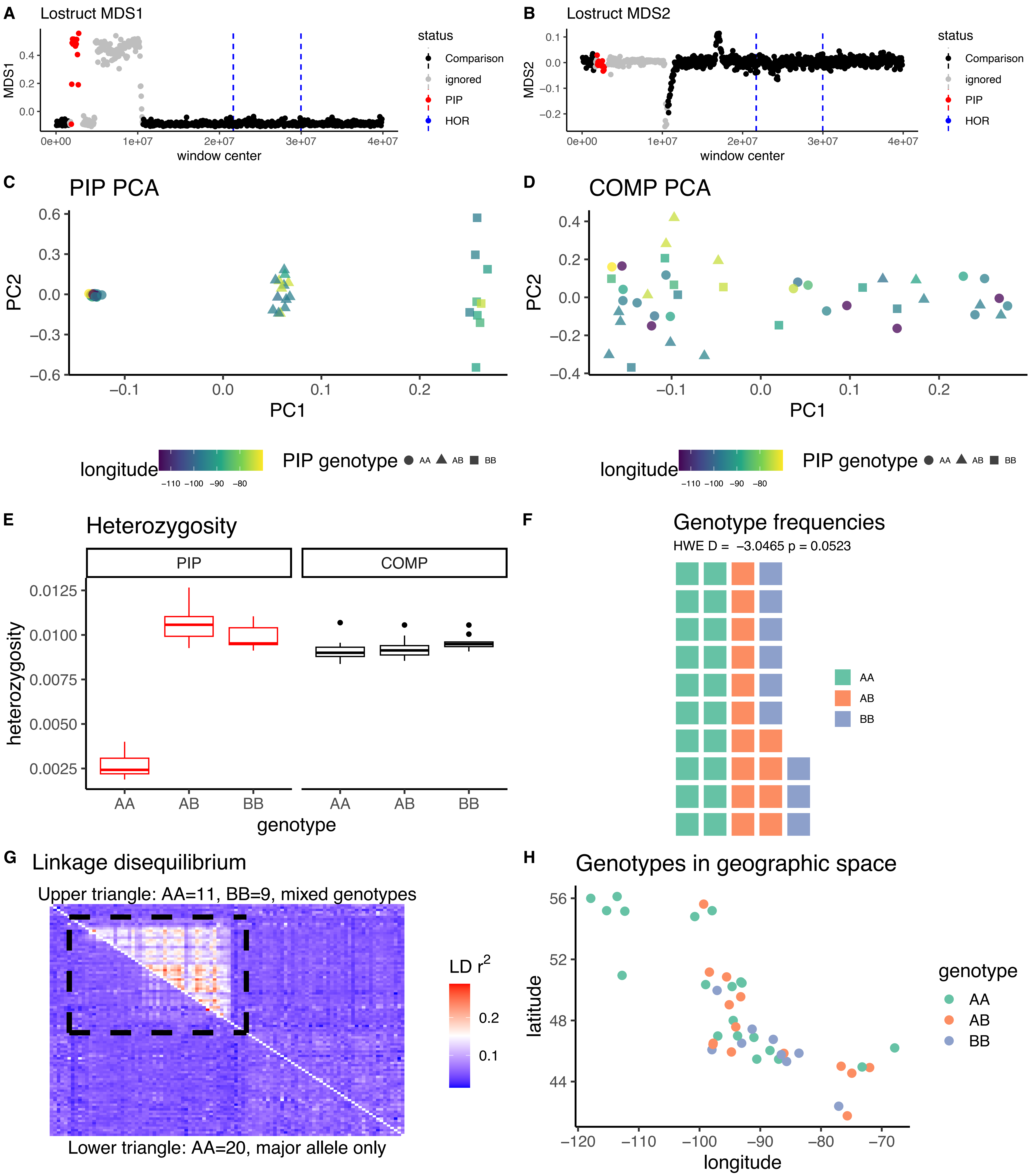

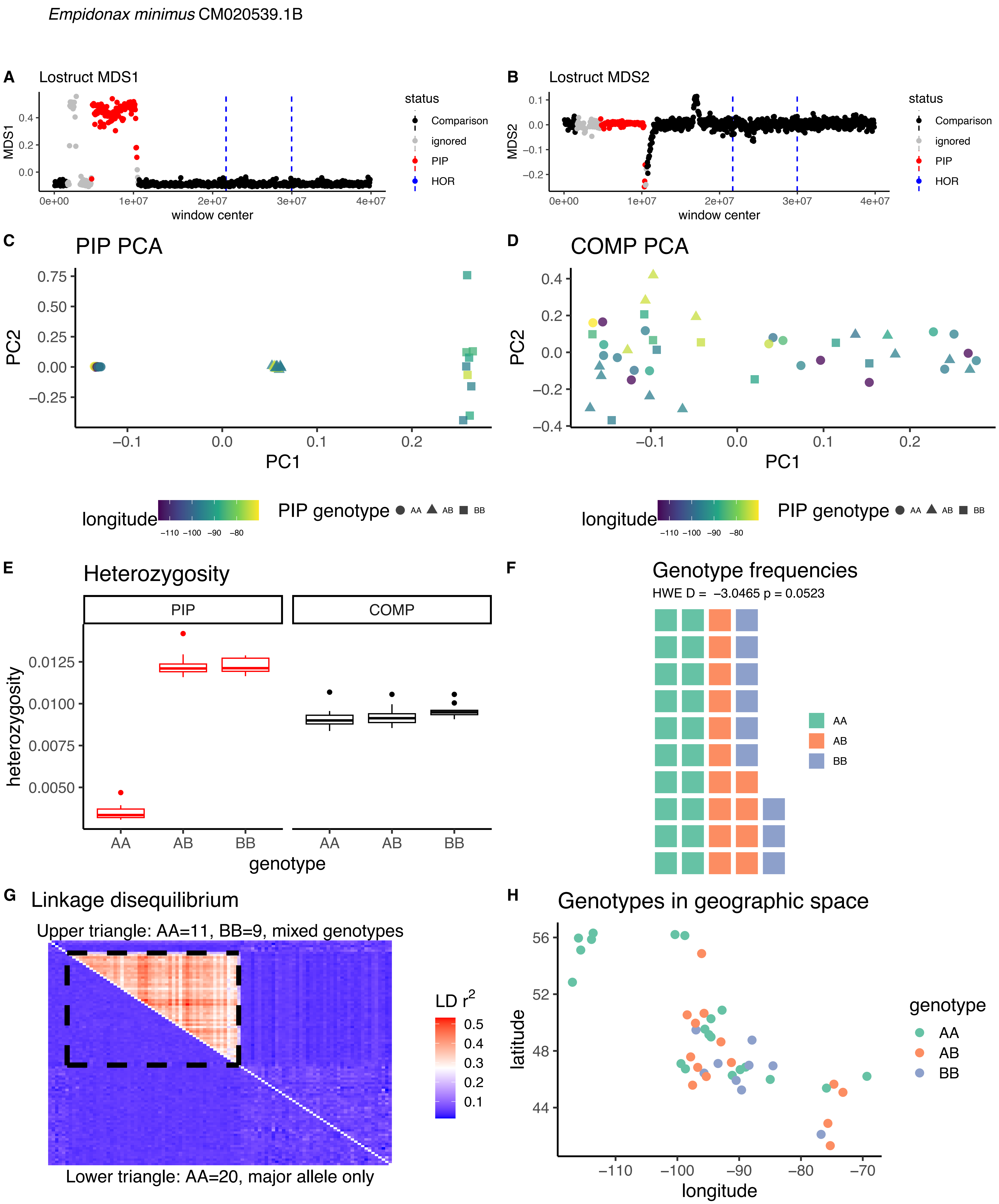

Empidonax minimus CM020539.1B
A Lostruct MDS1
MDS1
window center
status
Comparison
ignored
PIP
HOR
B Lostruct MDS2
MDS2
window center
C PIP PCA
PC2
PC1
longitude
PIP genotype AA AB BB
D COMP PCA
E Heterozygosity
PIP
COMP
heterozygosity
genotype
F Genotype frequencies
HWE D = −3.0465 p = 0.0523
AA
AB
BB
G Linkage disequilibrium
Upper triangle: AA=11, BB=9, mixed genotypes
LD r2
Lower triangle: AA=20, major allele only
H Genotypes in geographic space
latitude
longitude
genotype

*Empidonax minimus* CM020537.1

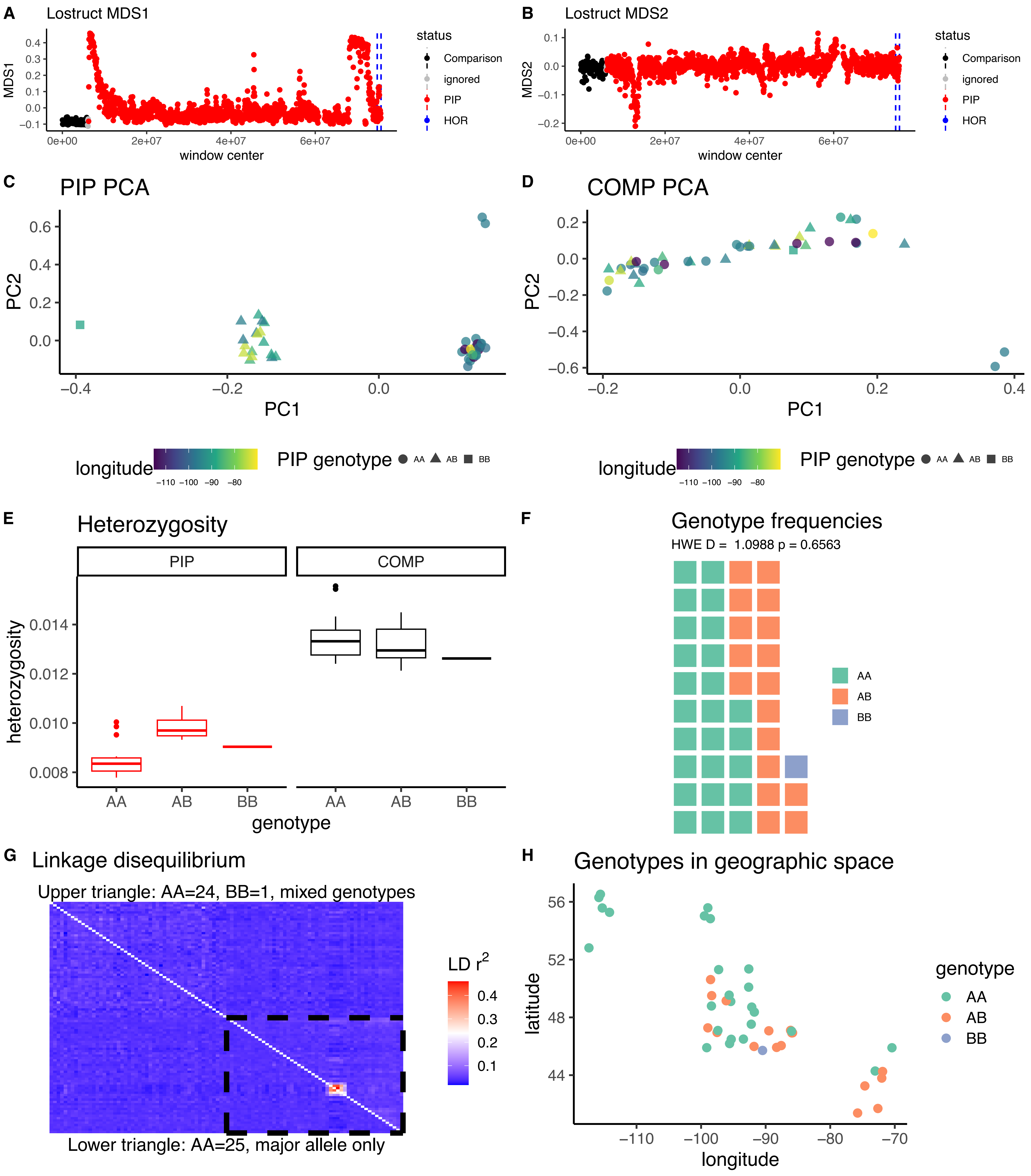

*Empidonax minimus* CM020535.1

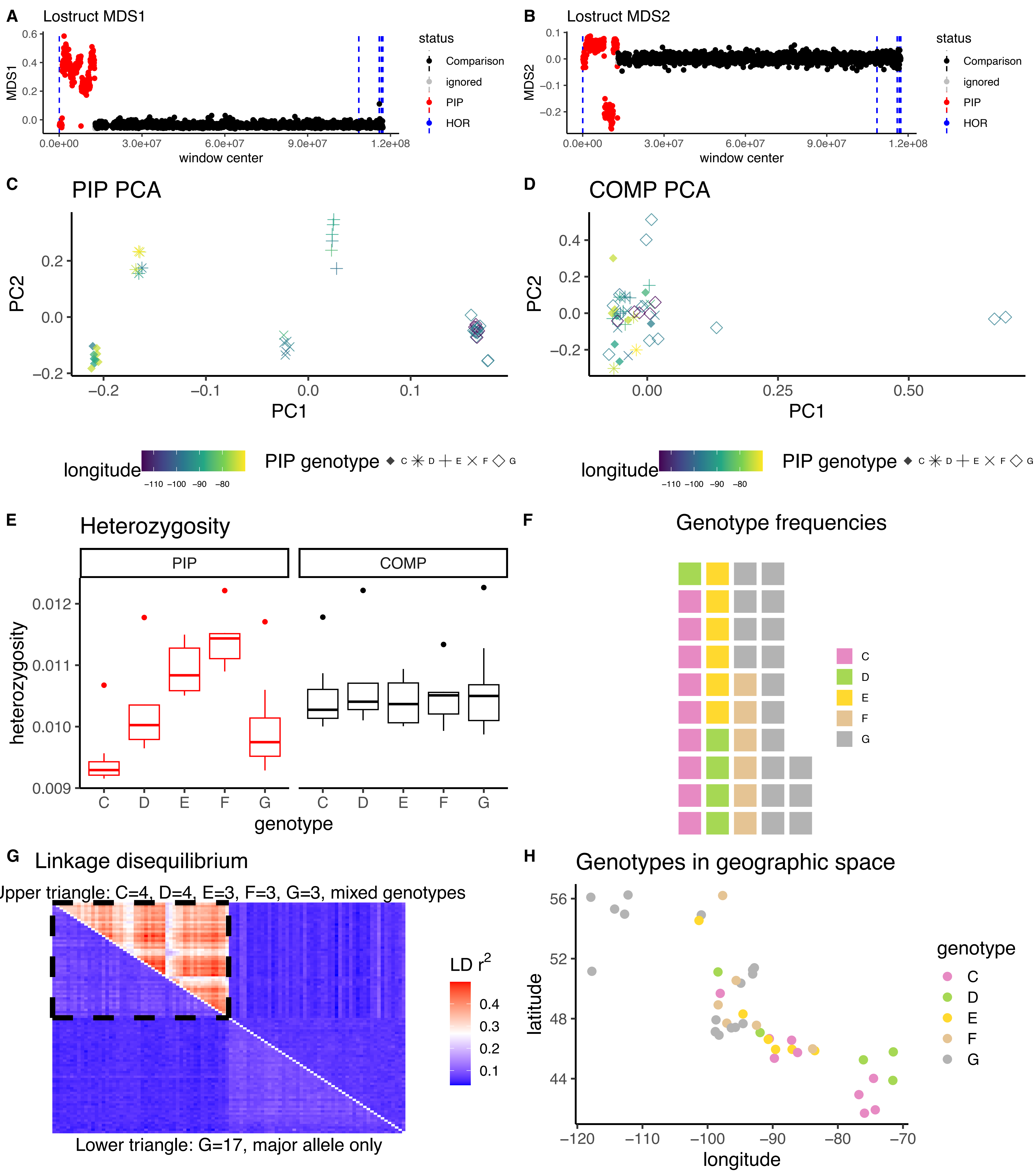

*Empidonax minimus* CM020534.1A

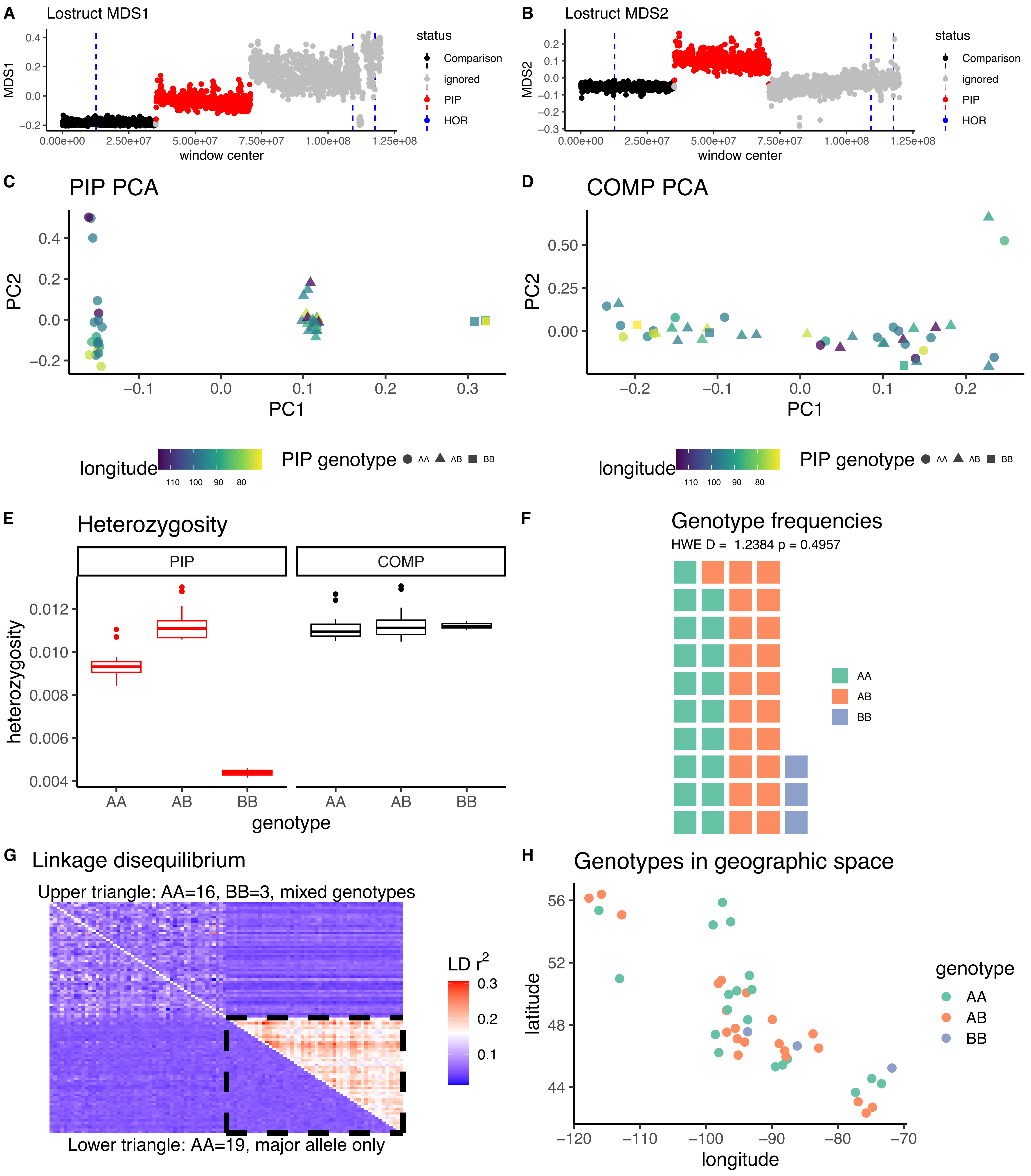

*Empidonax minimus* CM020534.1B

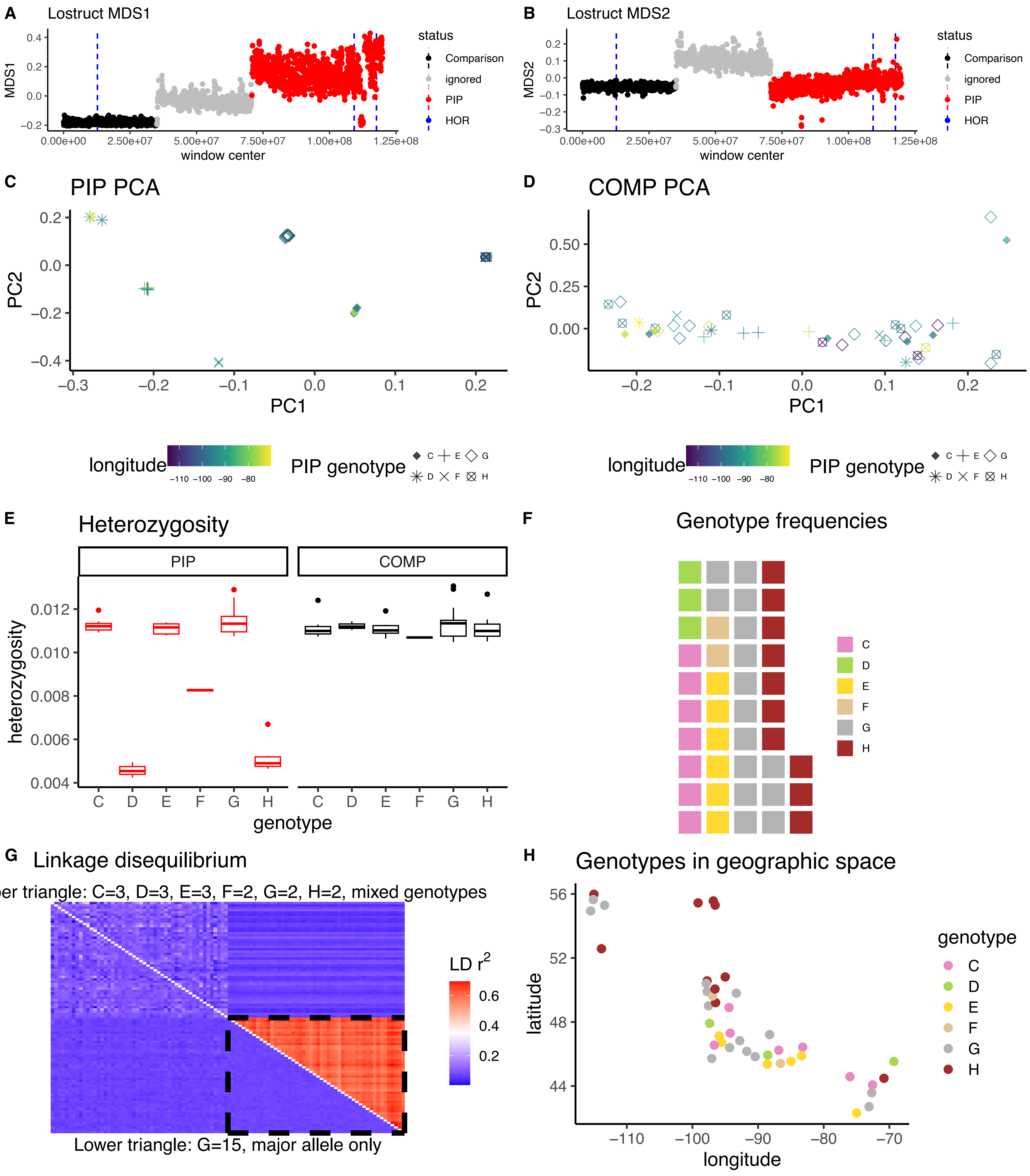

*Empidonax minimus* CM020533.1

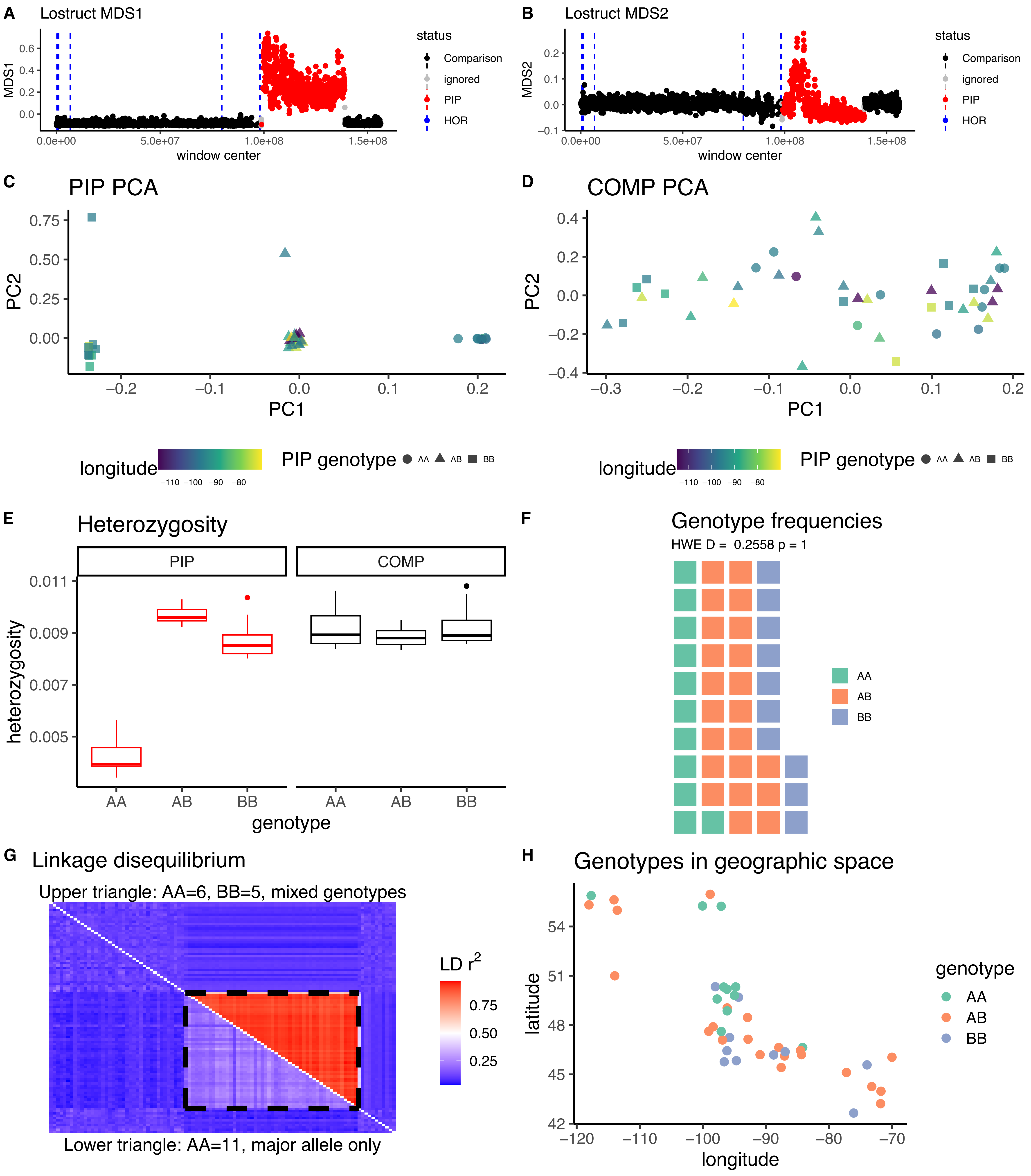

*Vireo olivaceus* CM036381.1

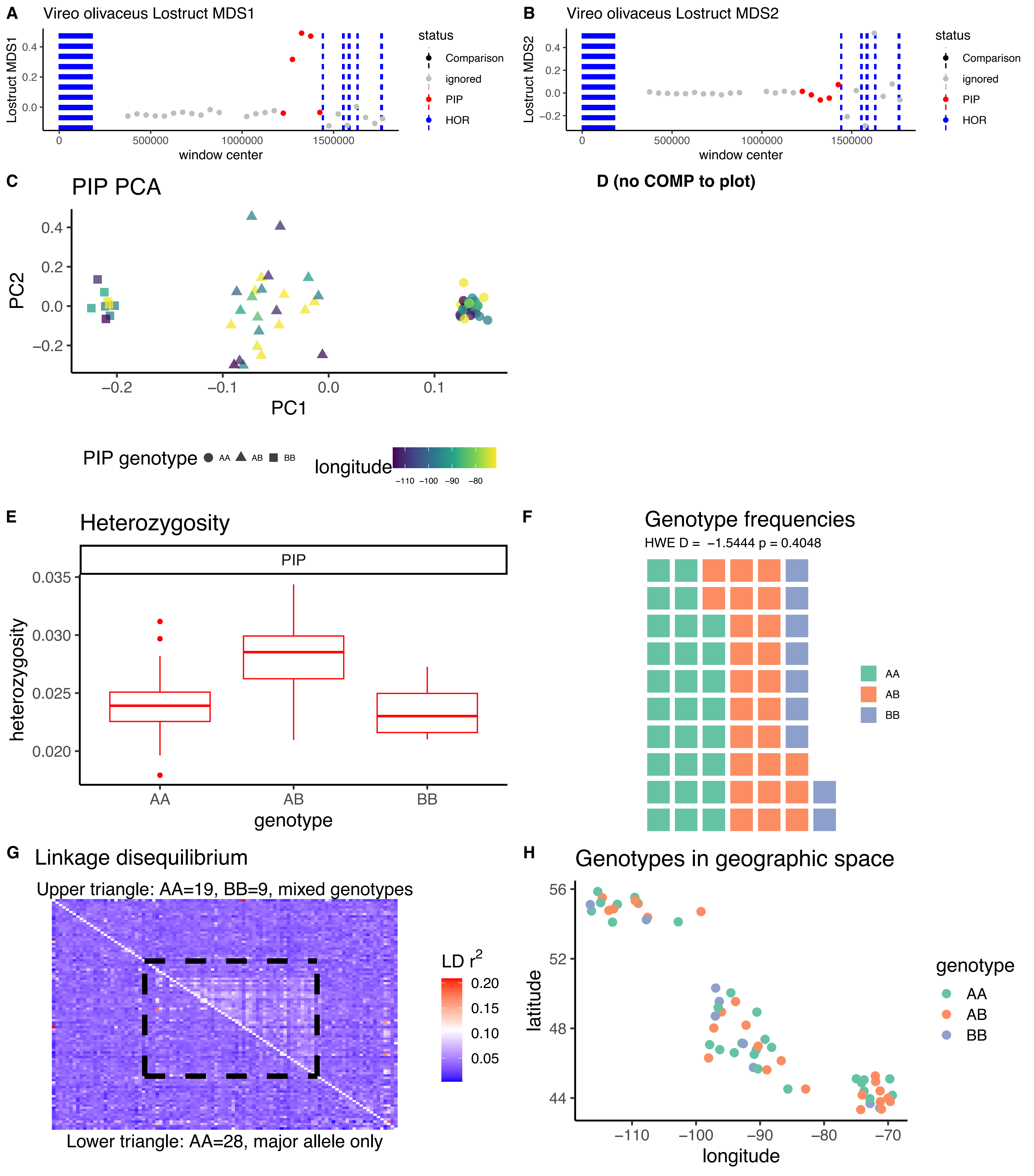

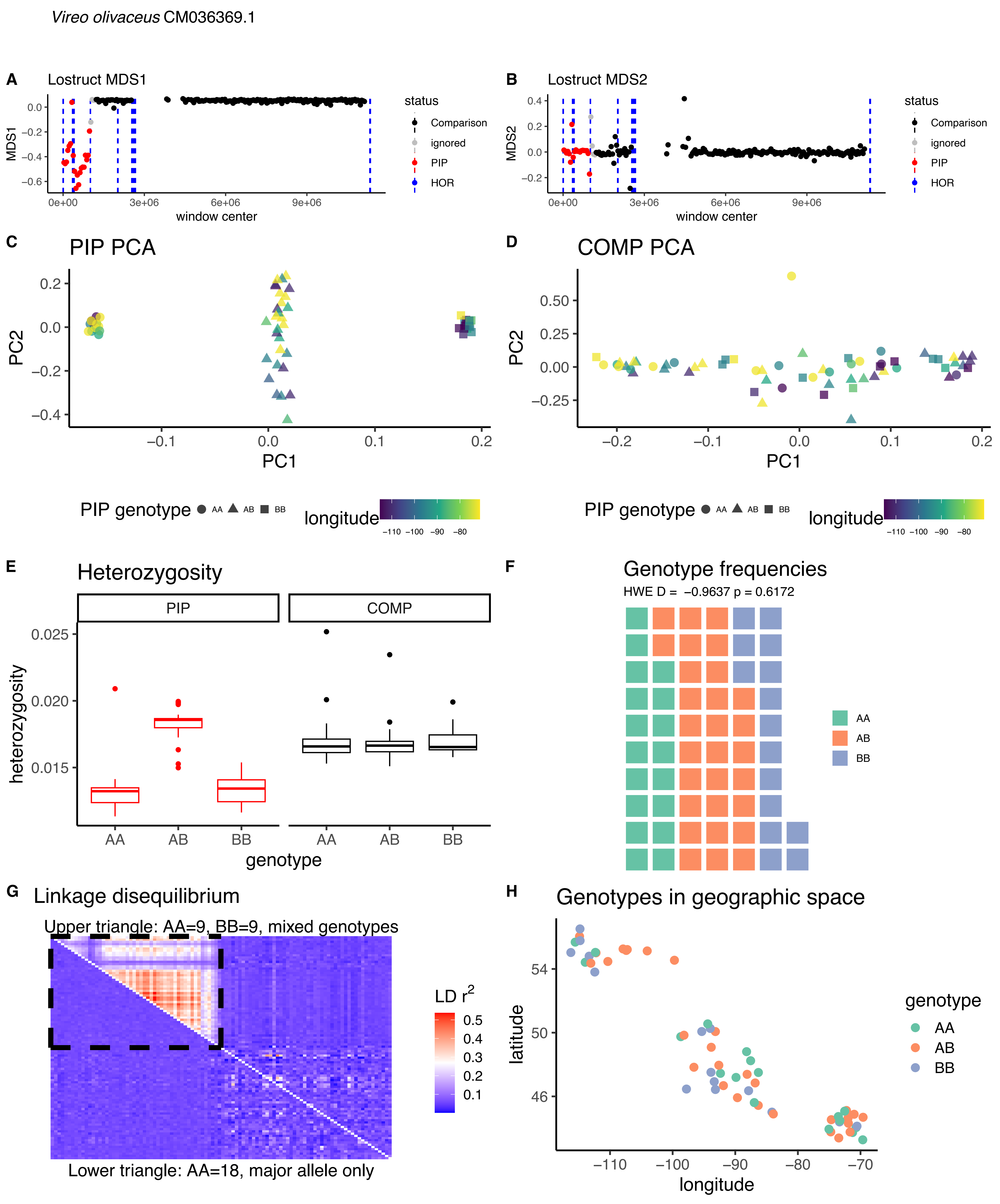
Vireo olivaceus CM036369.1
A
Lostruct MDS1
MDS1
window center
status
Comparison
ignored
PIP
HOR
B
Lostruct MDS2
MDS2
window center
C
PIP PCA
PC1
PC2
D
COMP PCA
PC1
PC2
PIP genotype AA AB BB
longitude
E
Heterozygosity
PIP
COMP
heterozygosity
genotype
F
Genotype frequencies
HWE D = −0.9637 p = 0.6172
AA
AB
BB
G
Linkage disequilibrium
Upper triangle: AA=9, BB=9, mixed genotypes
LD r²
Lower triangle: AA=18, major allele only
H
Genotypes in geographic space
latitude
longitude
genotype
AA
AB
BB

*Vireo olivaceus* CM036357.1

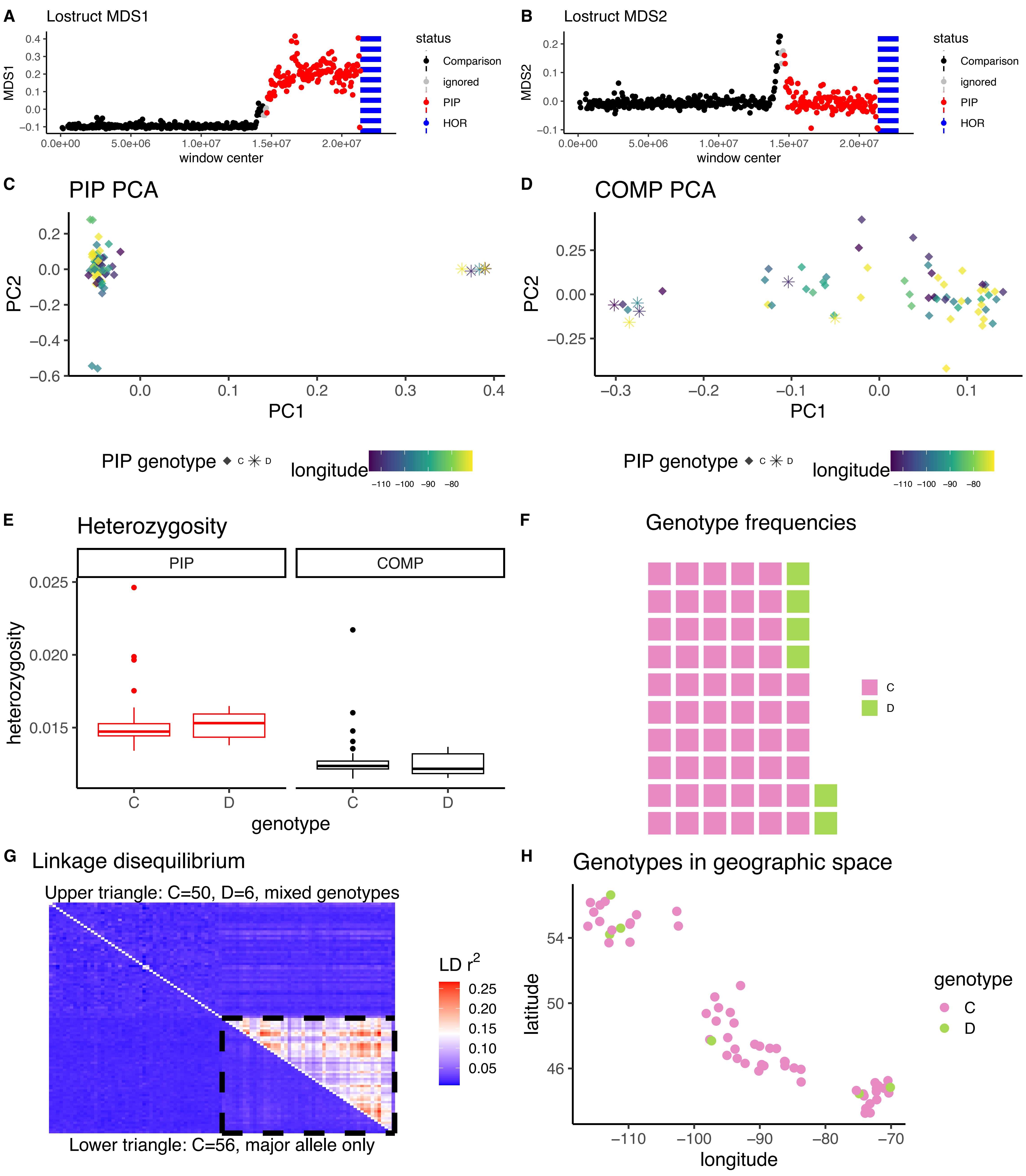

*Vireo olivaceus* CM036353.1

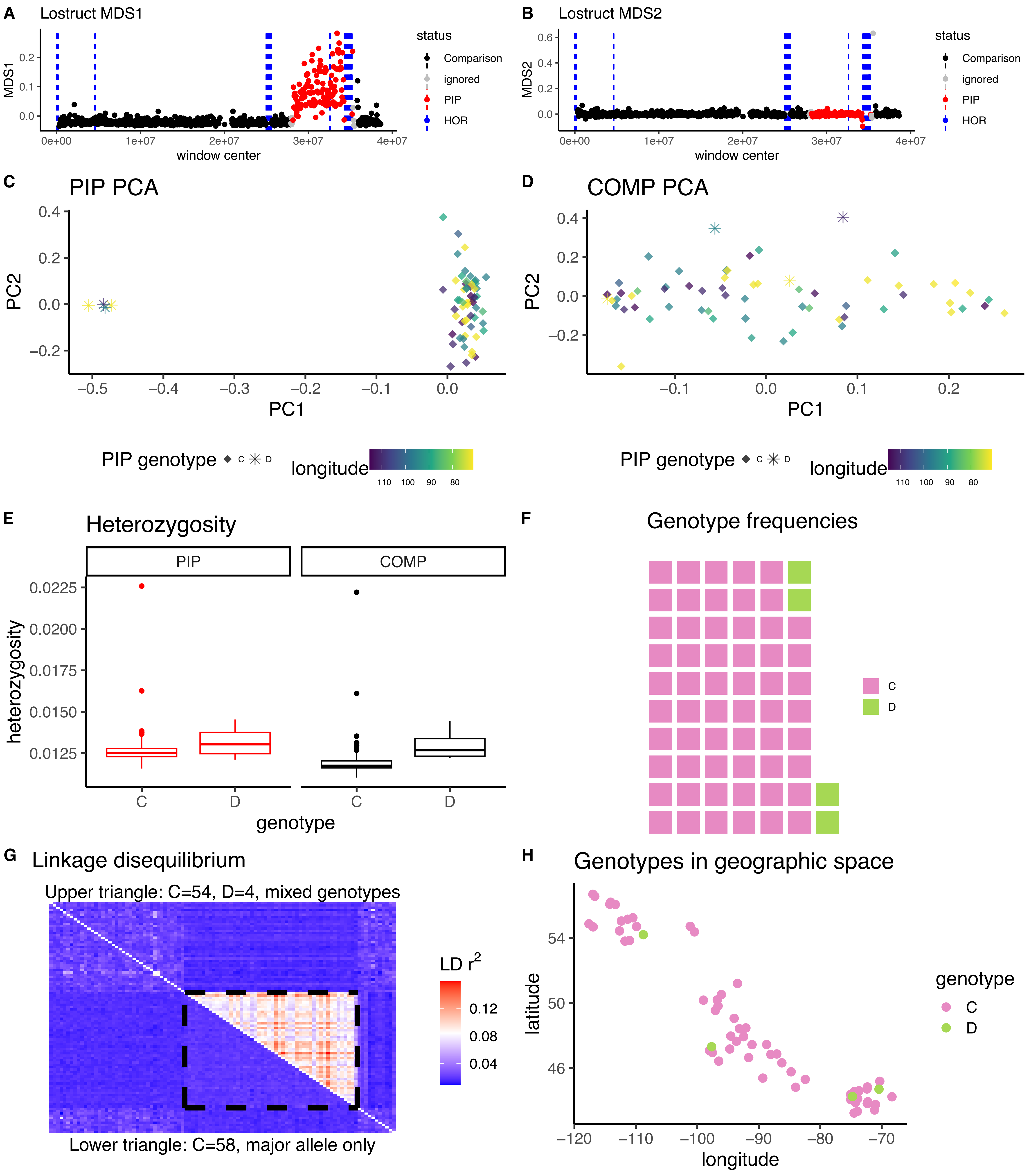

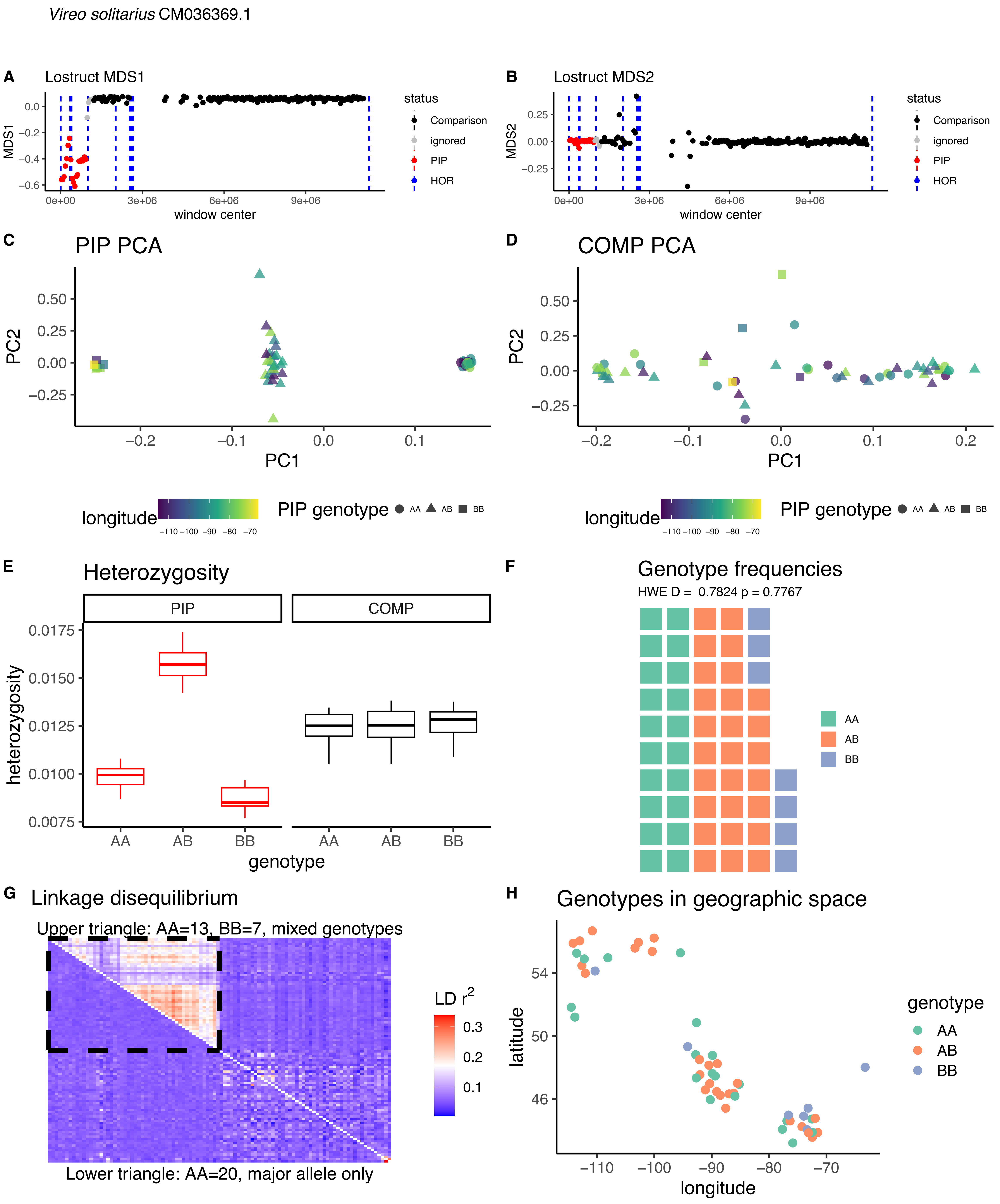

Vireo solitarius CM036369.1
A
Lostruct MDS1
MDS1
window center
status
Comparison
ignored
PIP
HOR
B
Lostruct MDS2
MDS2
window center
C
PIP PCA
PC1
PC2
longitude
PIP genotype AA AB BB
D
COMP PCA
PC1
PC2
longitude
PIP genotype AA AB BB
E
Heterozygosity
PIP
COMP
heterozygosity
genotype
F
Genotype frequencies
HWE D = 0.7824 p = 0.7767
AA
AB
BB
G
Linkage disequilibrium
Upper triangle: AA=13, BB=7, mixed genotypes
LD r²
Lower triangle: AA=20, major allele only
H
Genotypes in geographic space
latitude
longitude
genotype
AA
AB
BB

*Vireo solitarius* CM036353.1

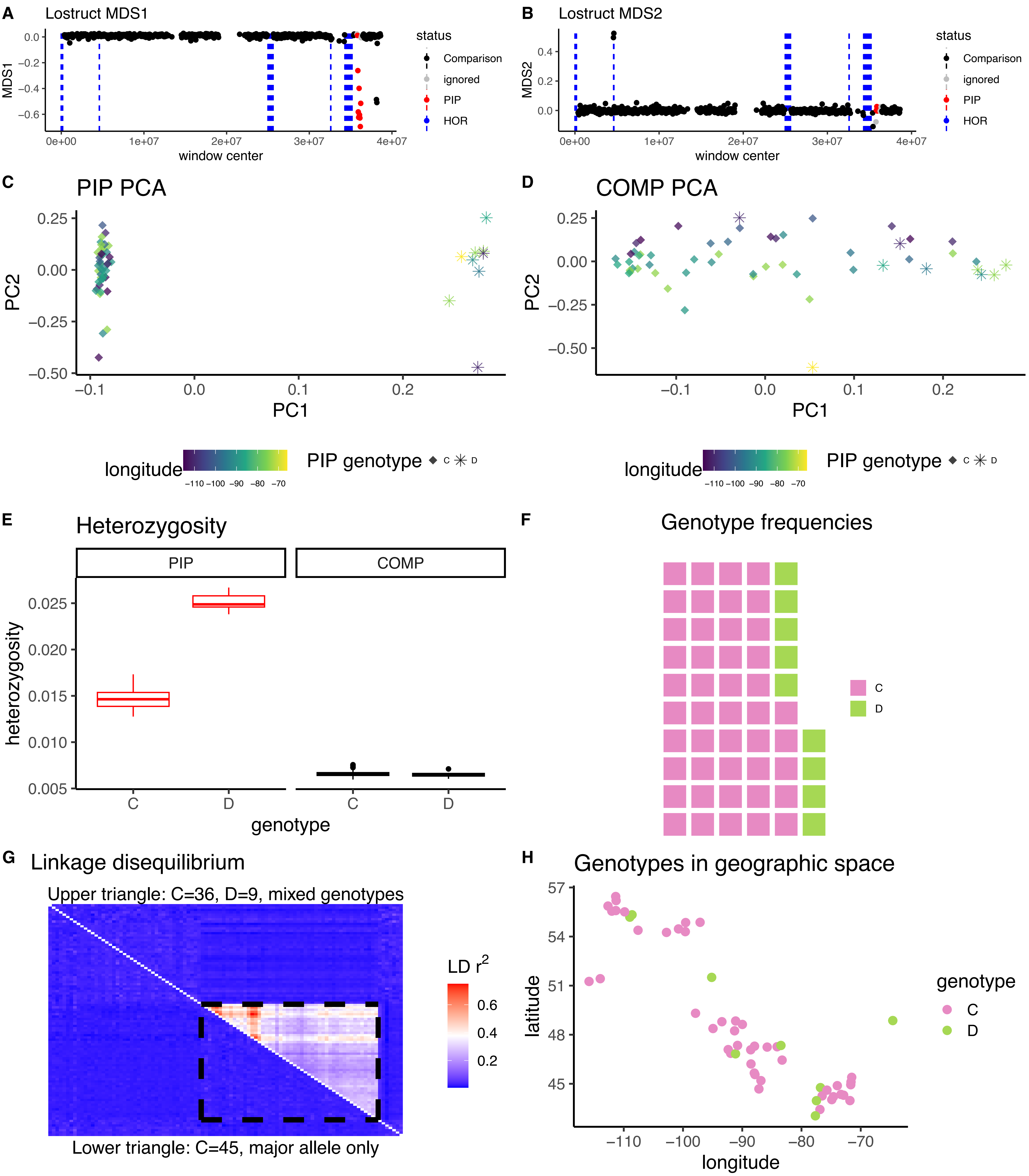

*Poecile atricapillus* CM003722.1

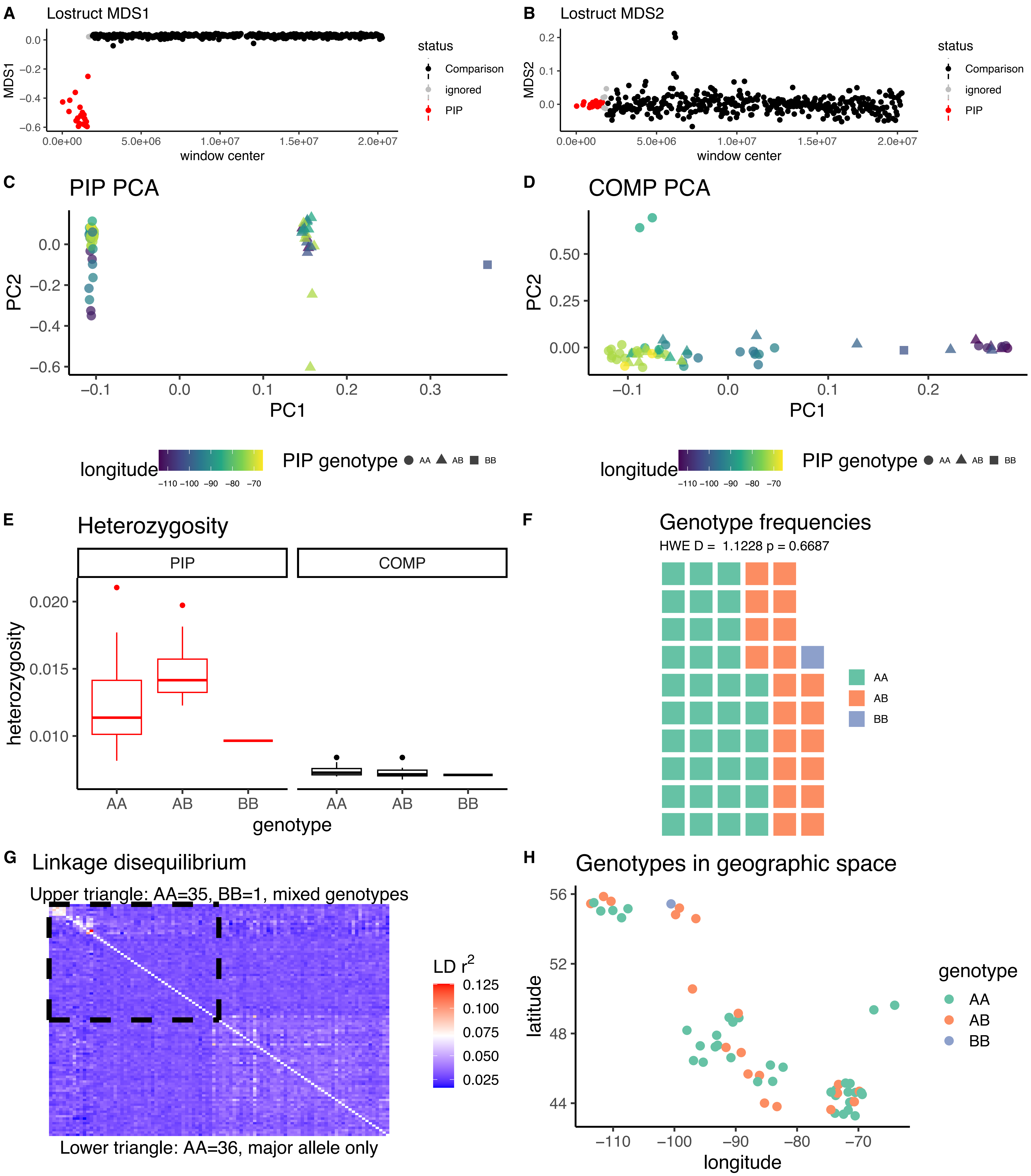

*Poecile atricapillus* CM003717.2

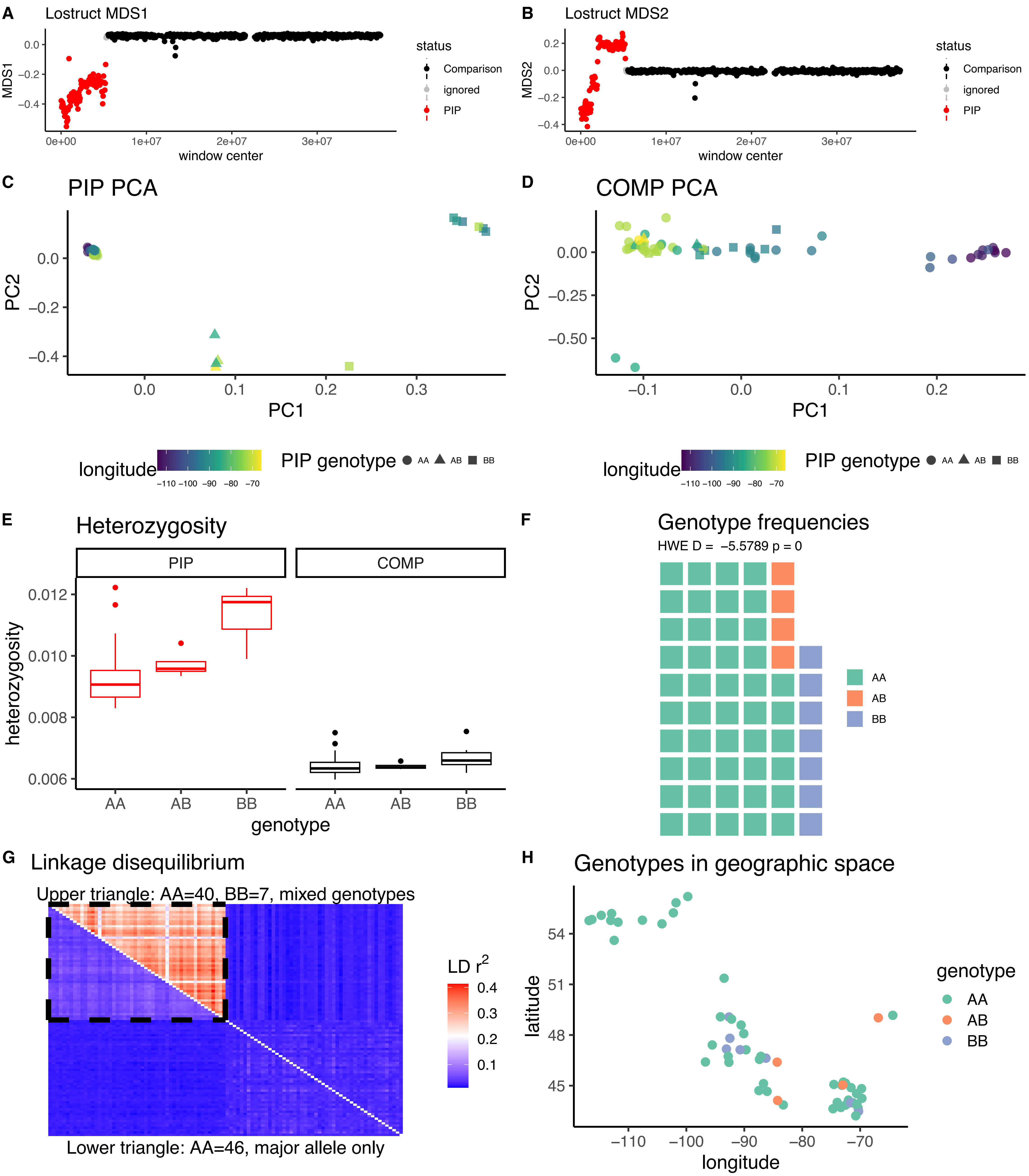

*Corthylio calendula* CM031877.1

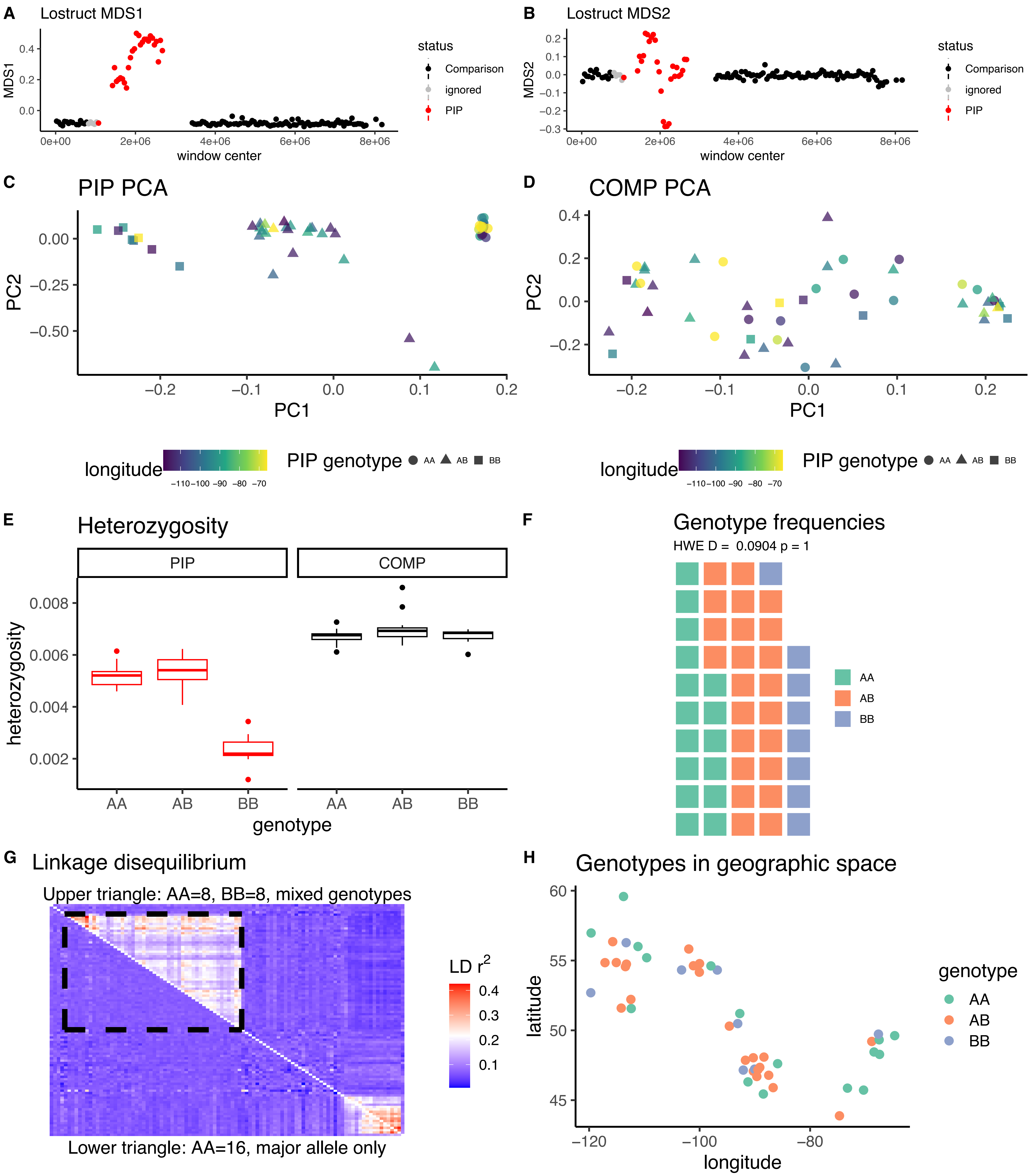

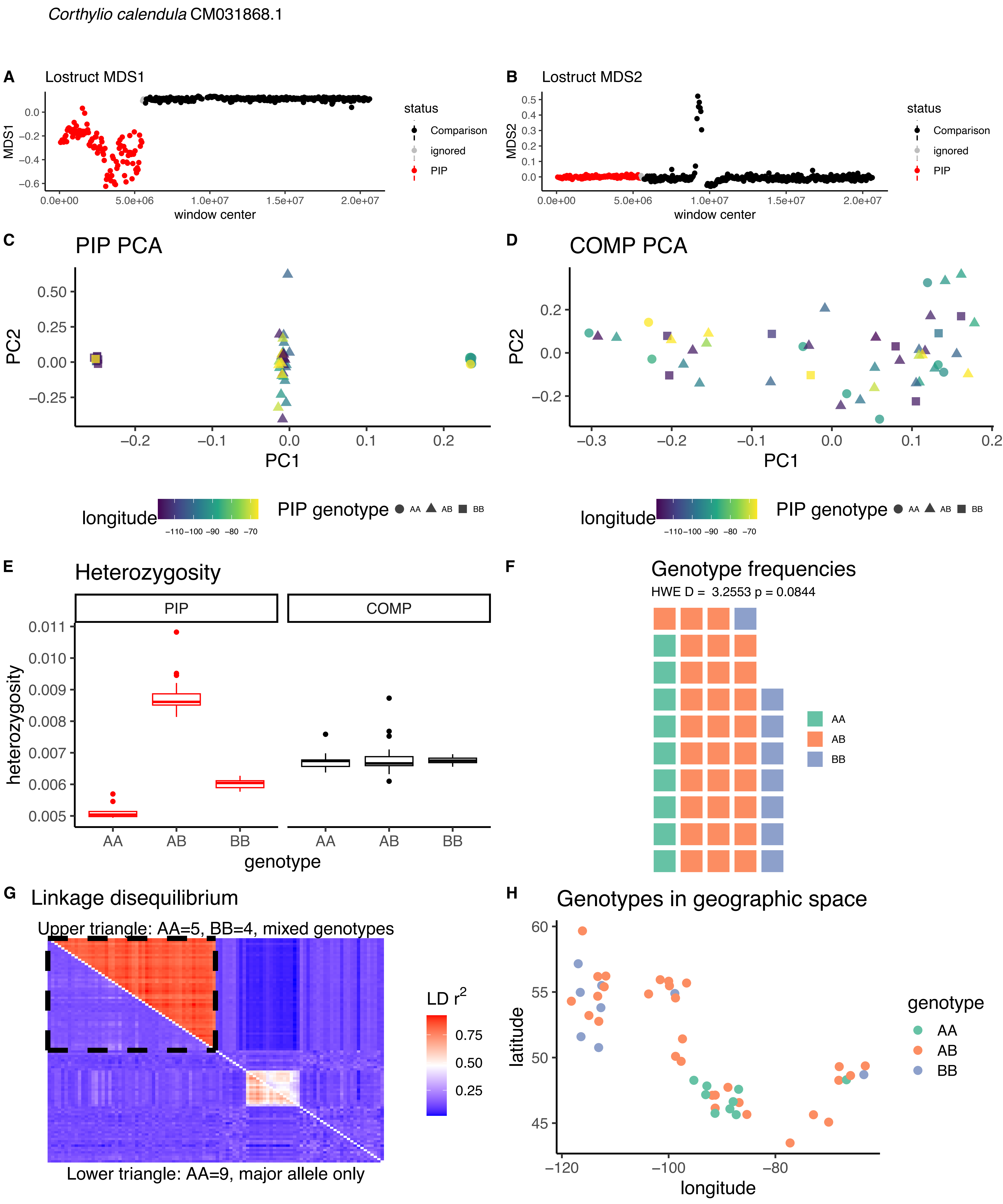
Corthylio calendula CM031868.1
A
Lostruct MDS1
MDS1
window center
status
Comparison
ignored
PIP
B
Lostruct MDS2
MDS2
window center
C
PIP PCA
PC1
PC2
D
COMP PCA
longitude
PIP genotype
AA
AB
BB
E
Heterozygosity
PIP
COMP
heterozygosity
genotype
F
Genotype frequencies
HWE D = 3.2553 p = 0.0844
G
Linkage disequilibrium
Upper triangle: AA=5, BB=4, mixed genotypes
Lower triangle: AA=9, major allele only
LD r2
H
Genotypes in geographic space
latitude
longitude
genotype

*Corthylio calendula* CM031863.1

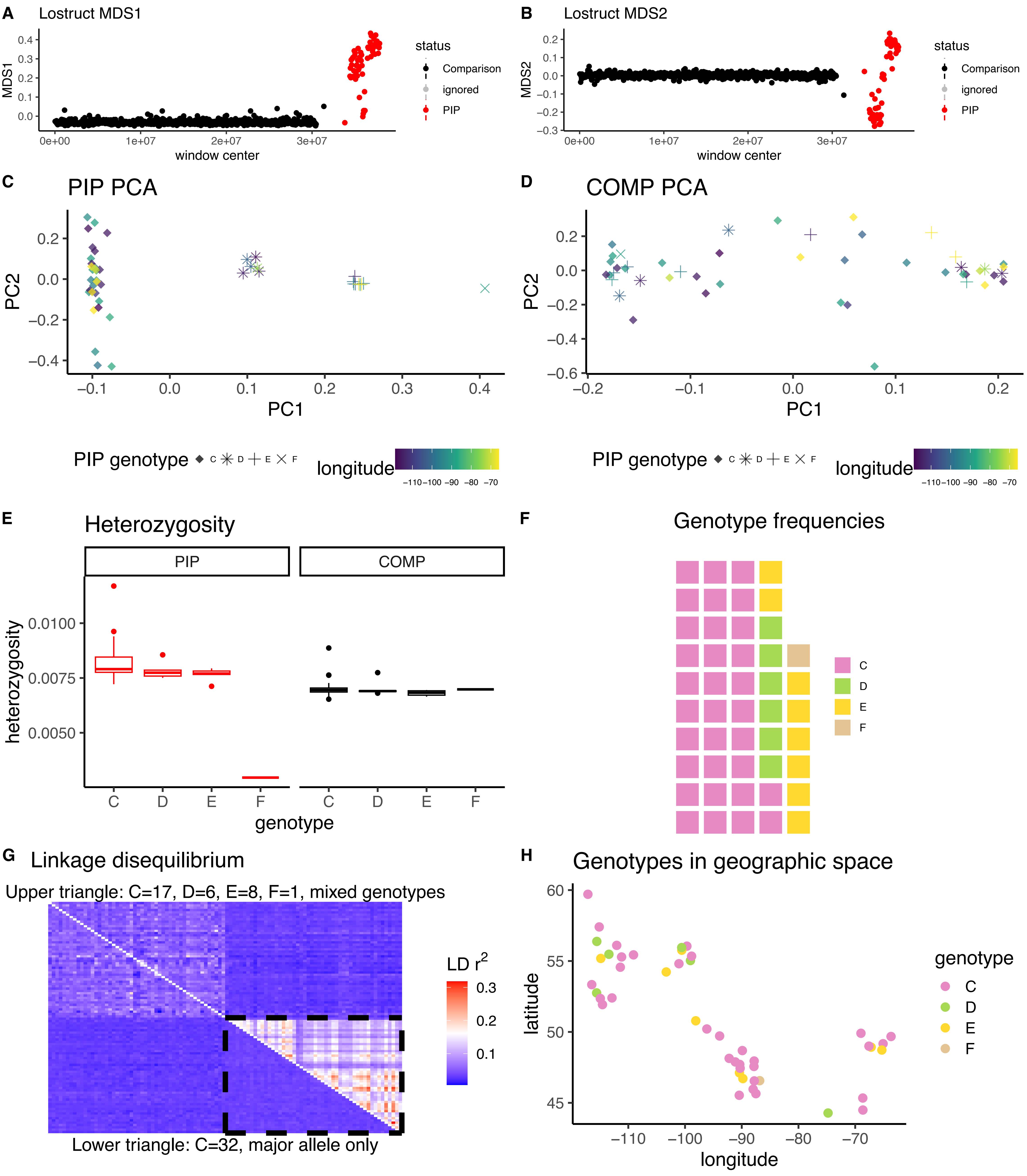

*Corthylio calendula* CM031861.1

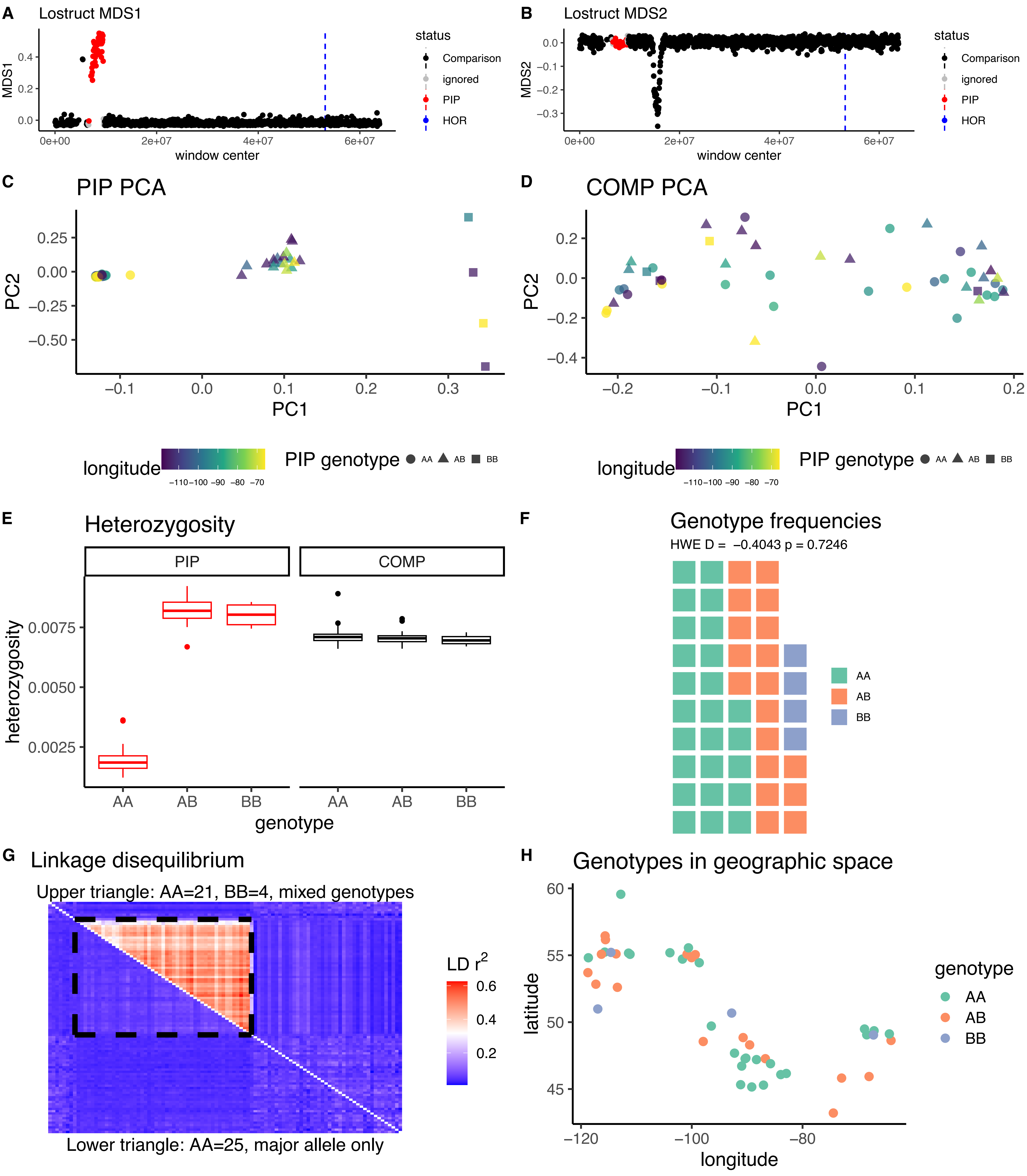

*Corthylio calendula* CM031859.1

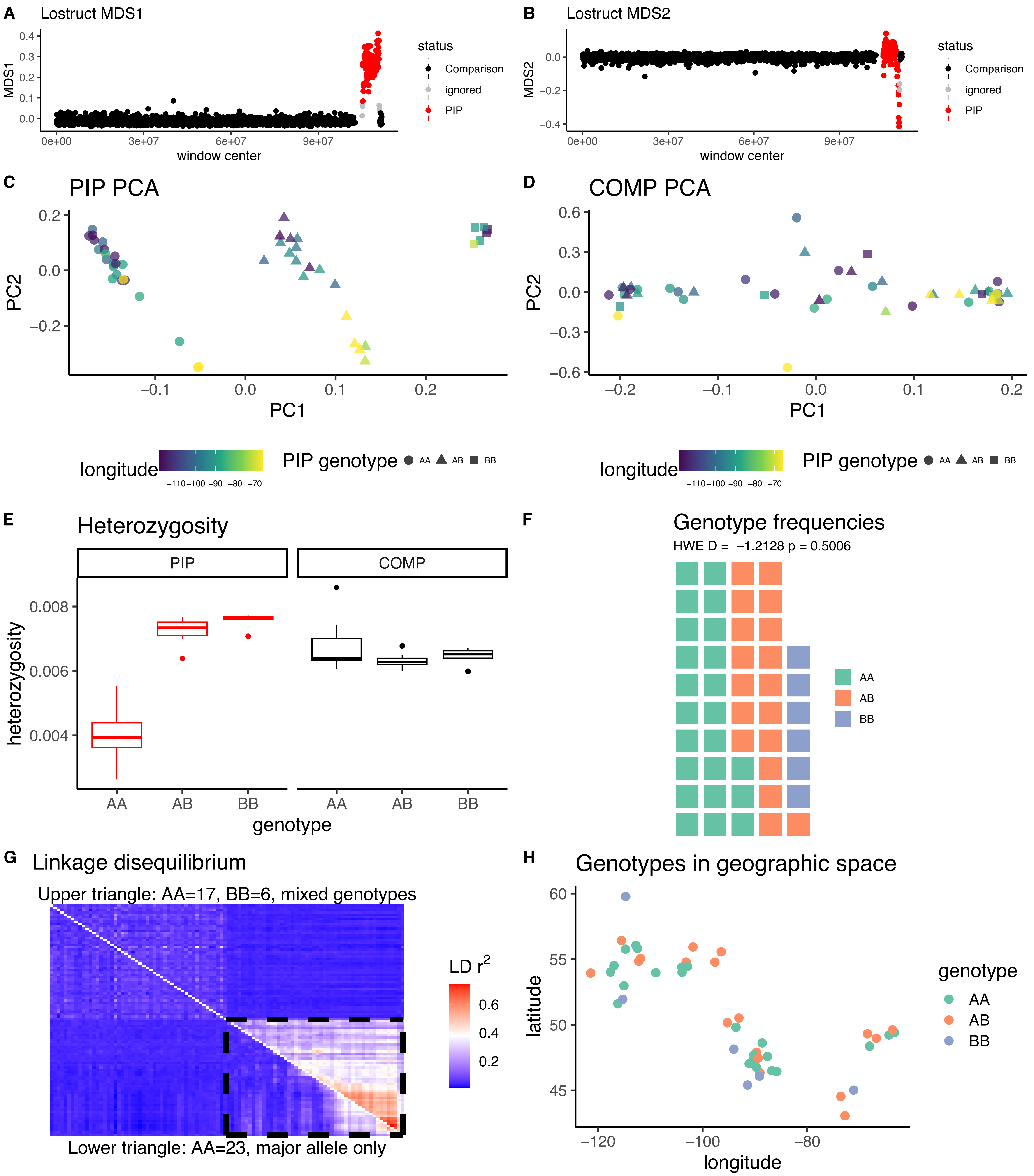

*Troglodytes hiemalis* CM031868.1

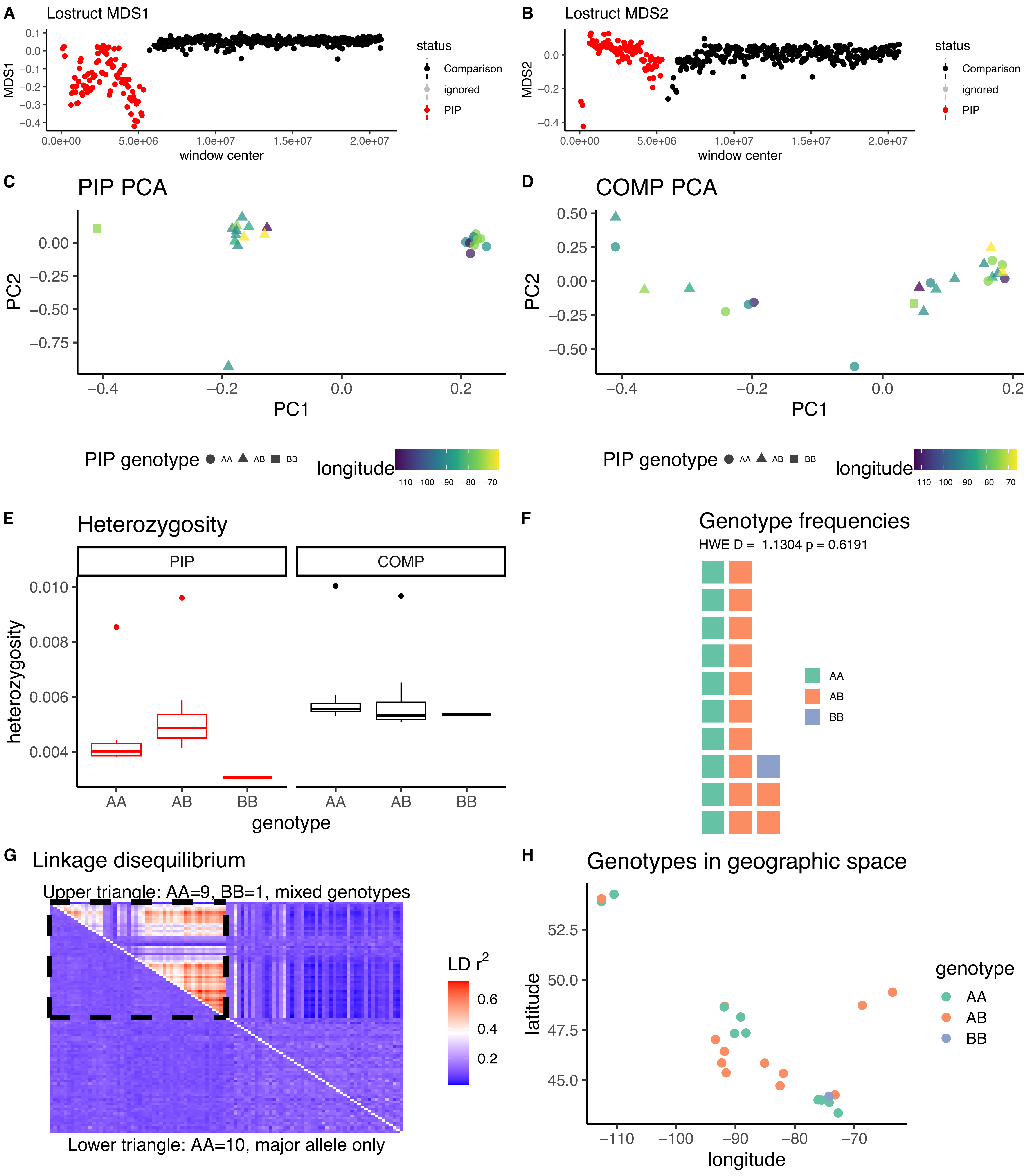

*Troglodytes hiemalis* CM031864.1

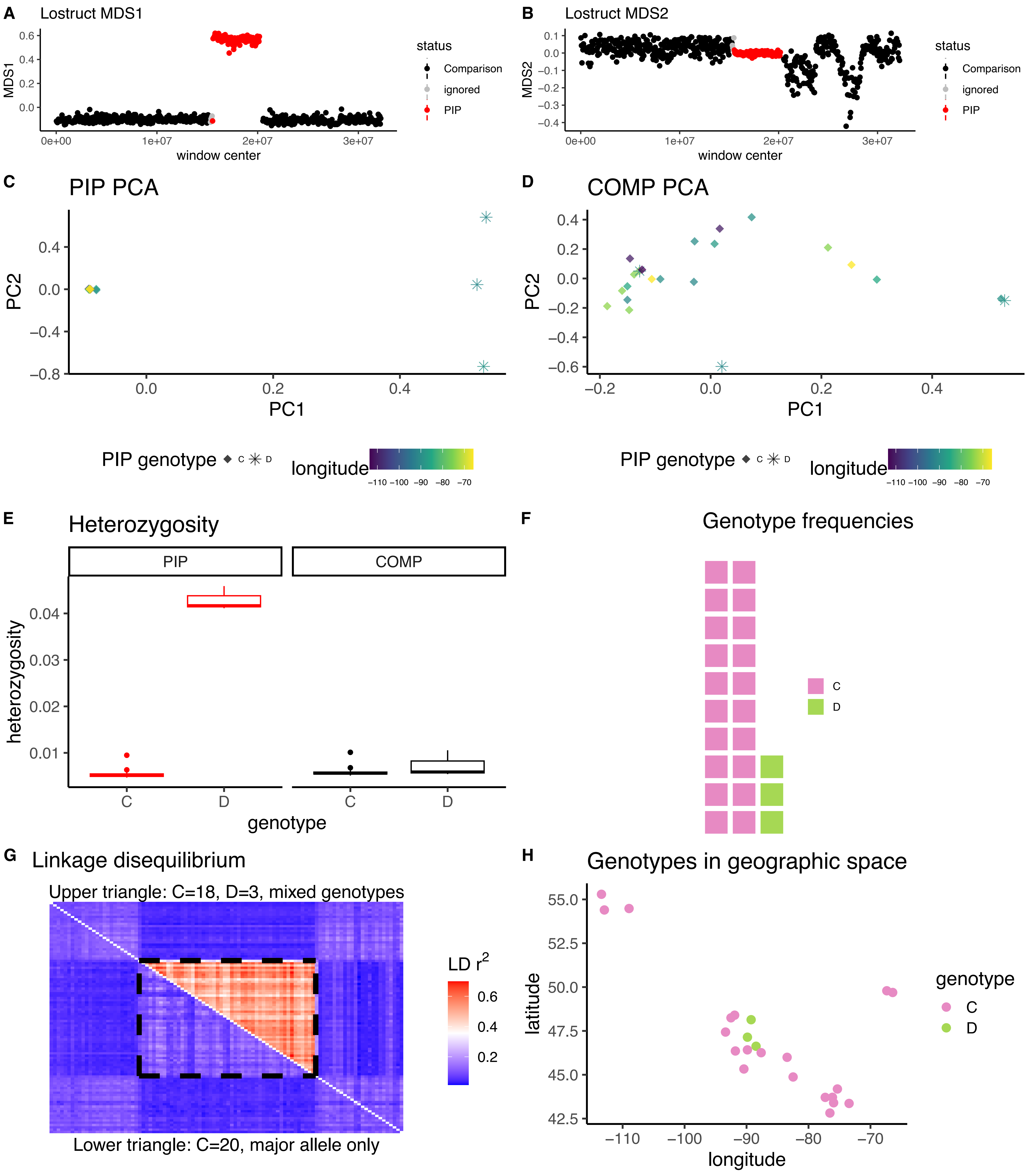

*Catharus fuscescens* CM020372.1

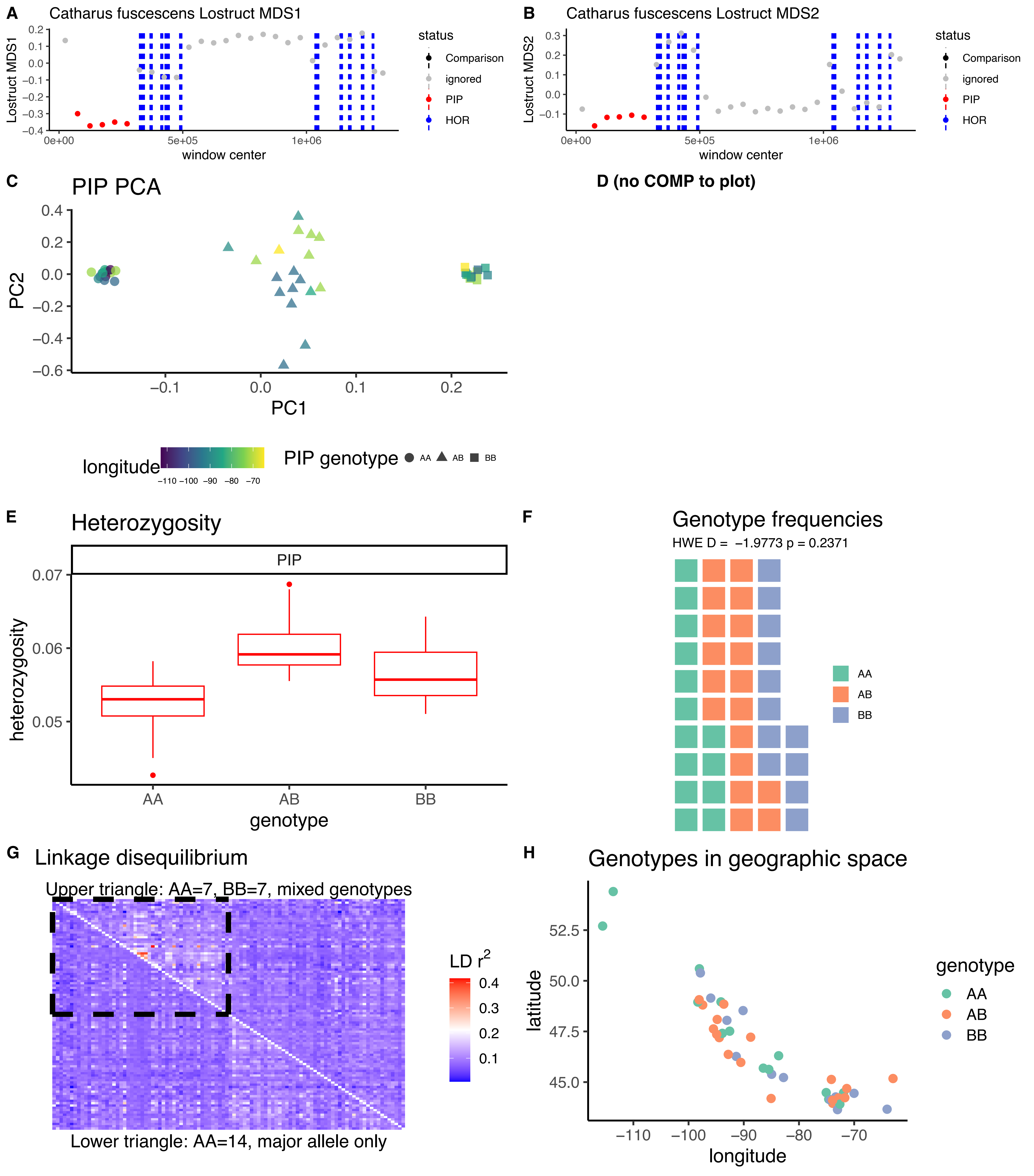

*Catharus fuscescens* CM020371.1A

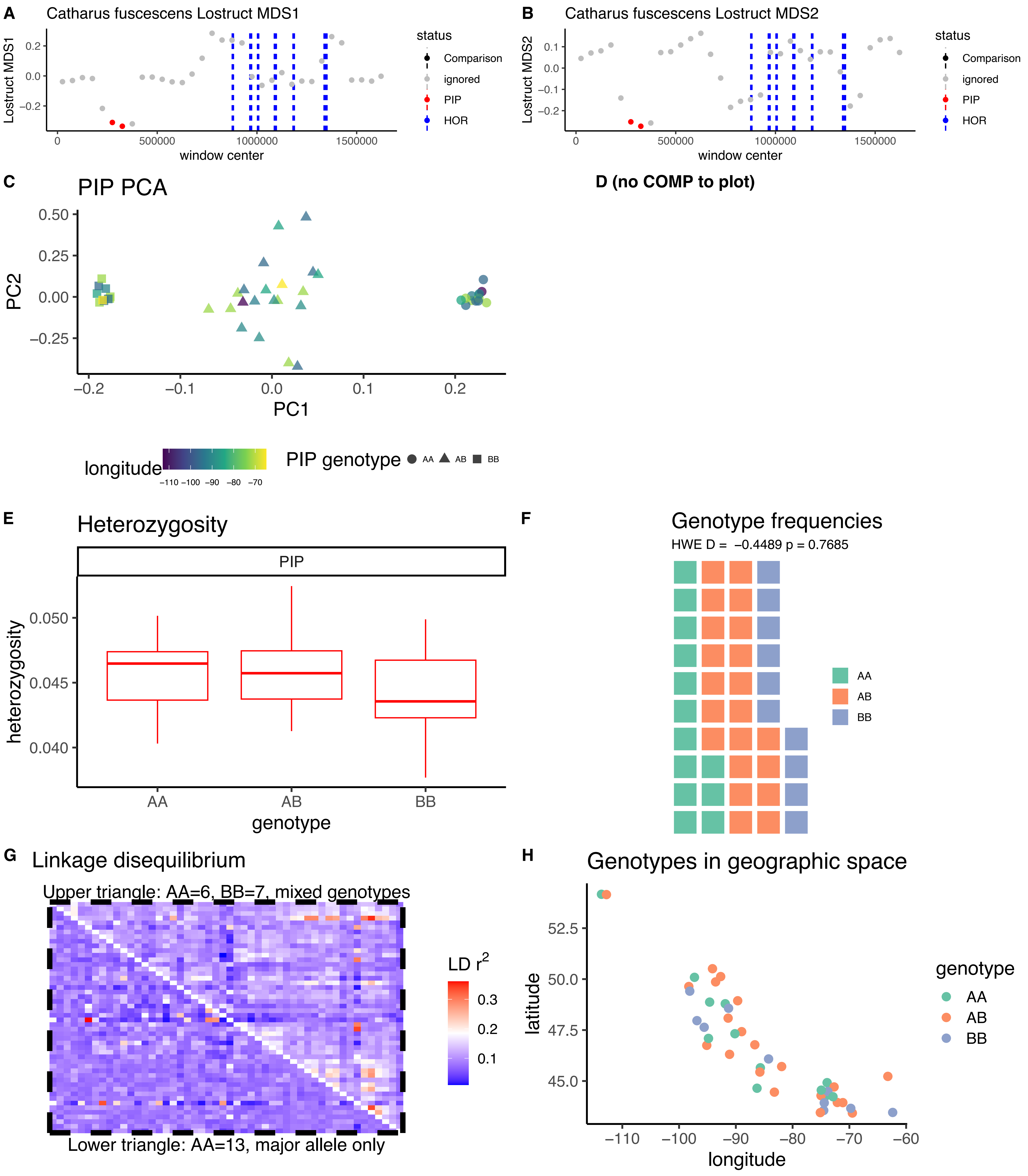

*Catharus fuscescens* CM020371.1B

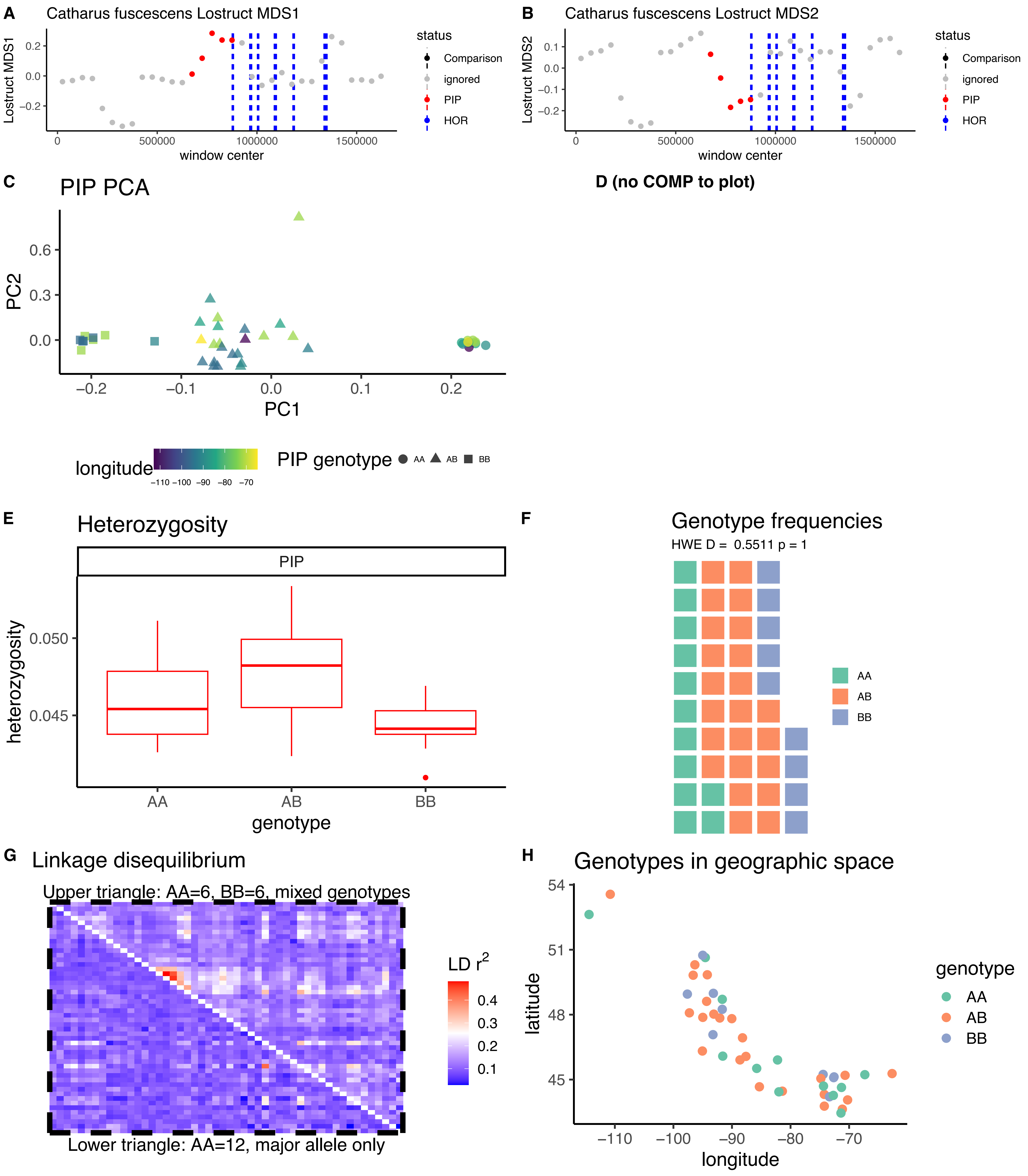

*Catharus fuscescens* CM020369.1

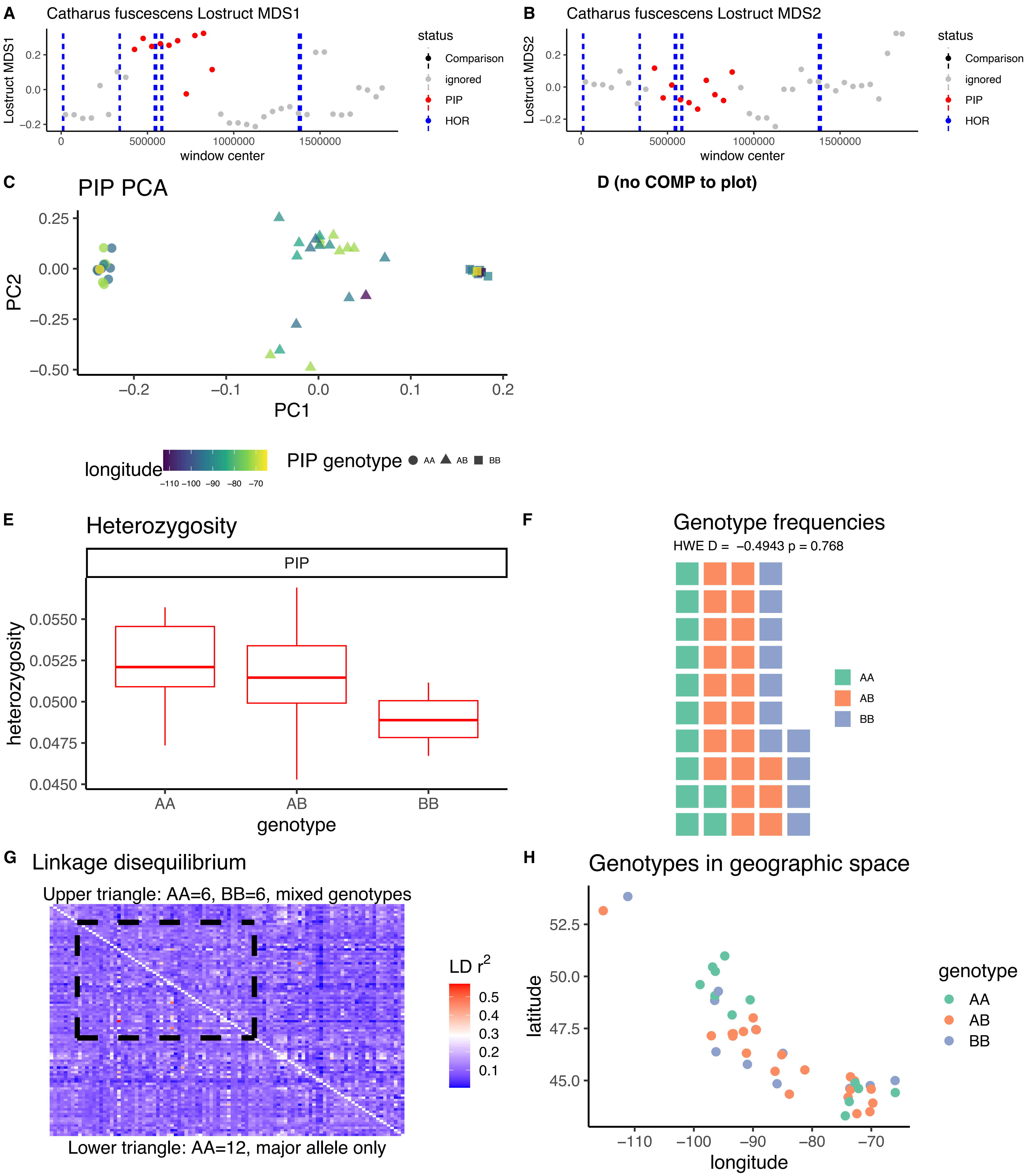

*Catharus fuscescens* CM020368.1

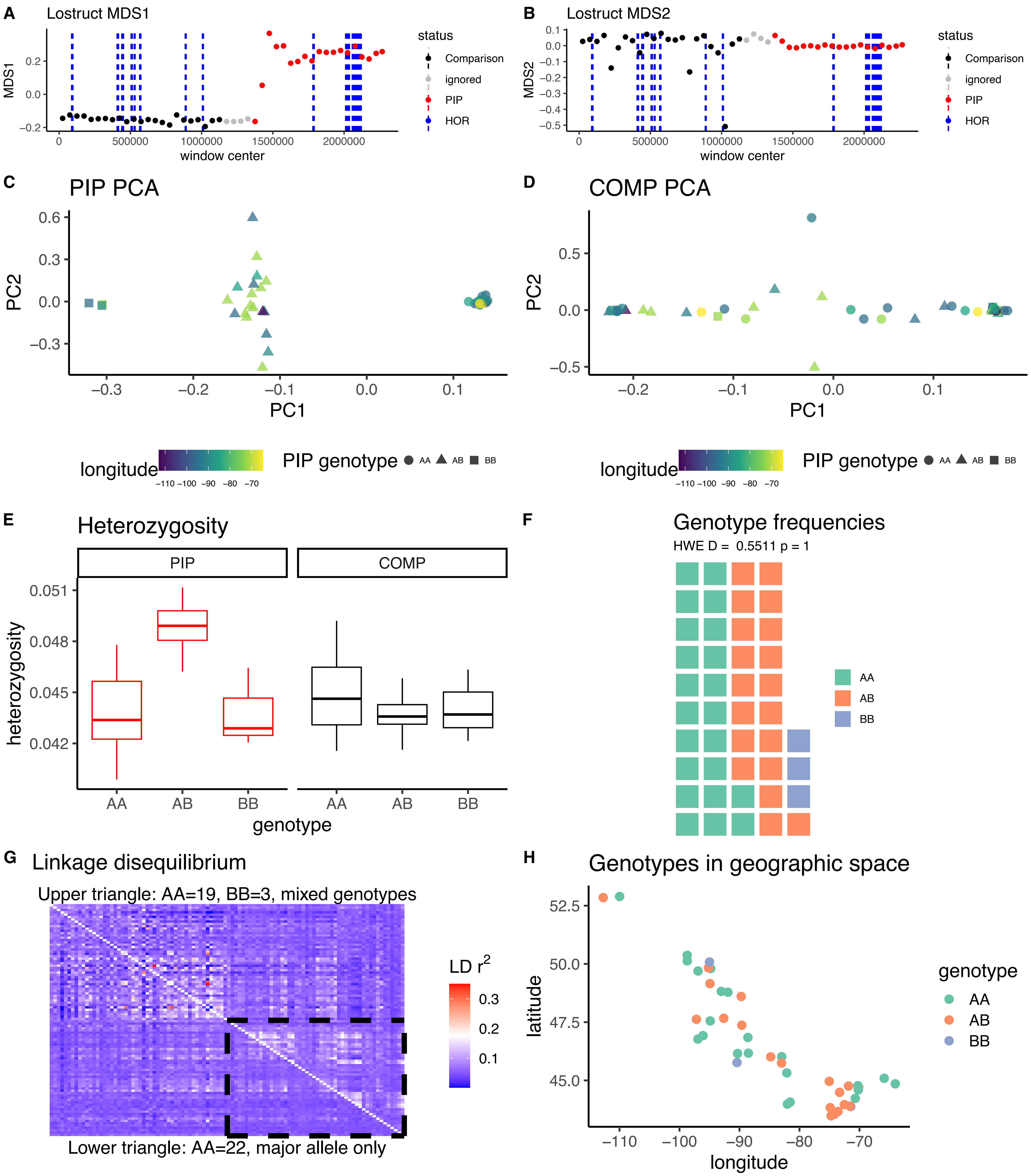

*Catharus fuscescens* CM020367.1A

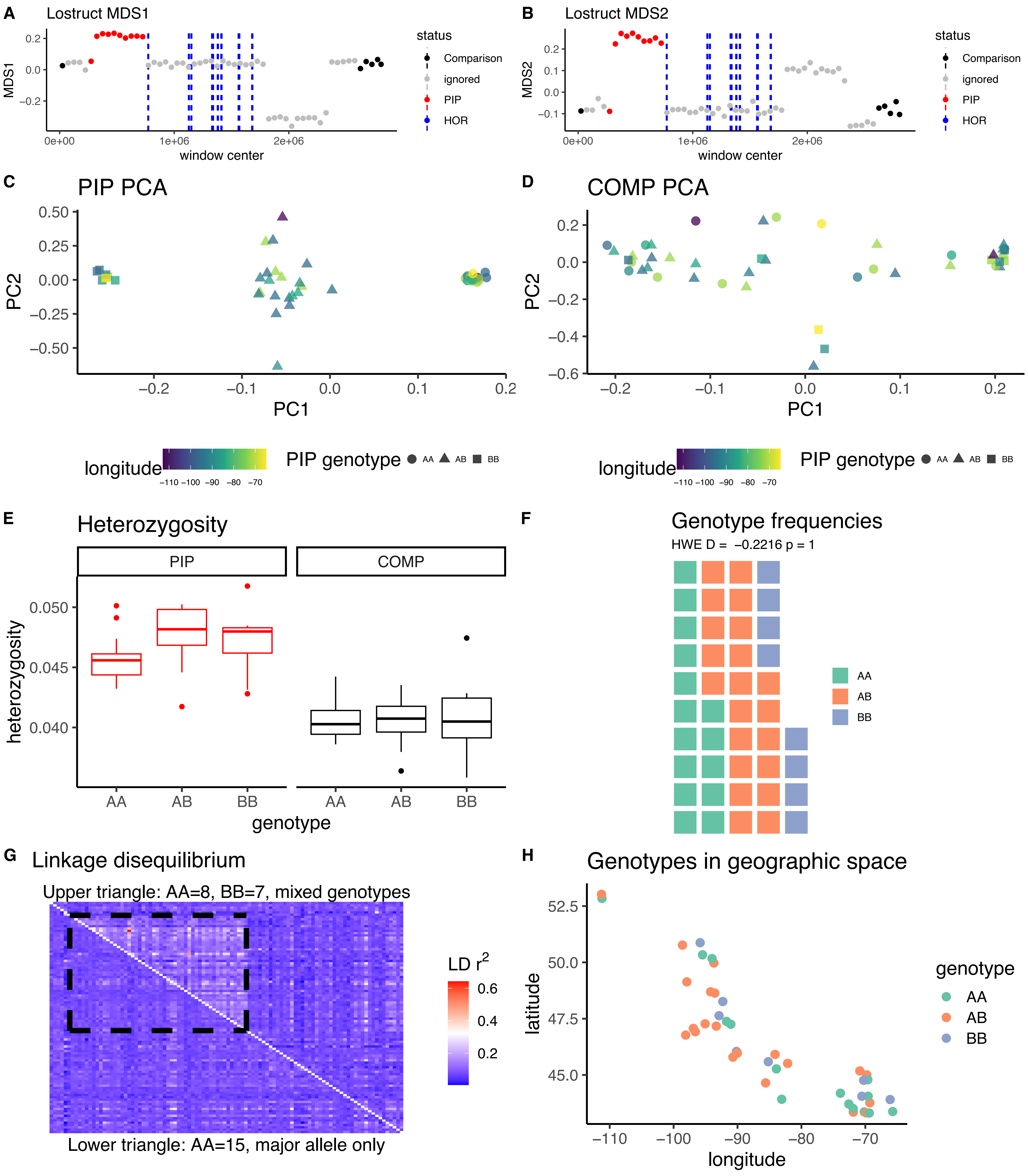

*Catharus fuscescens* CM020367.1B

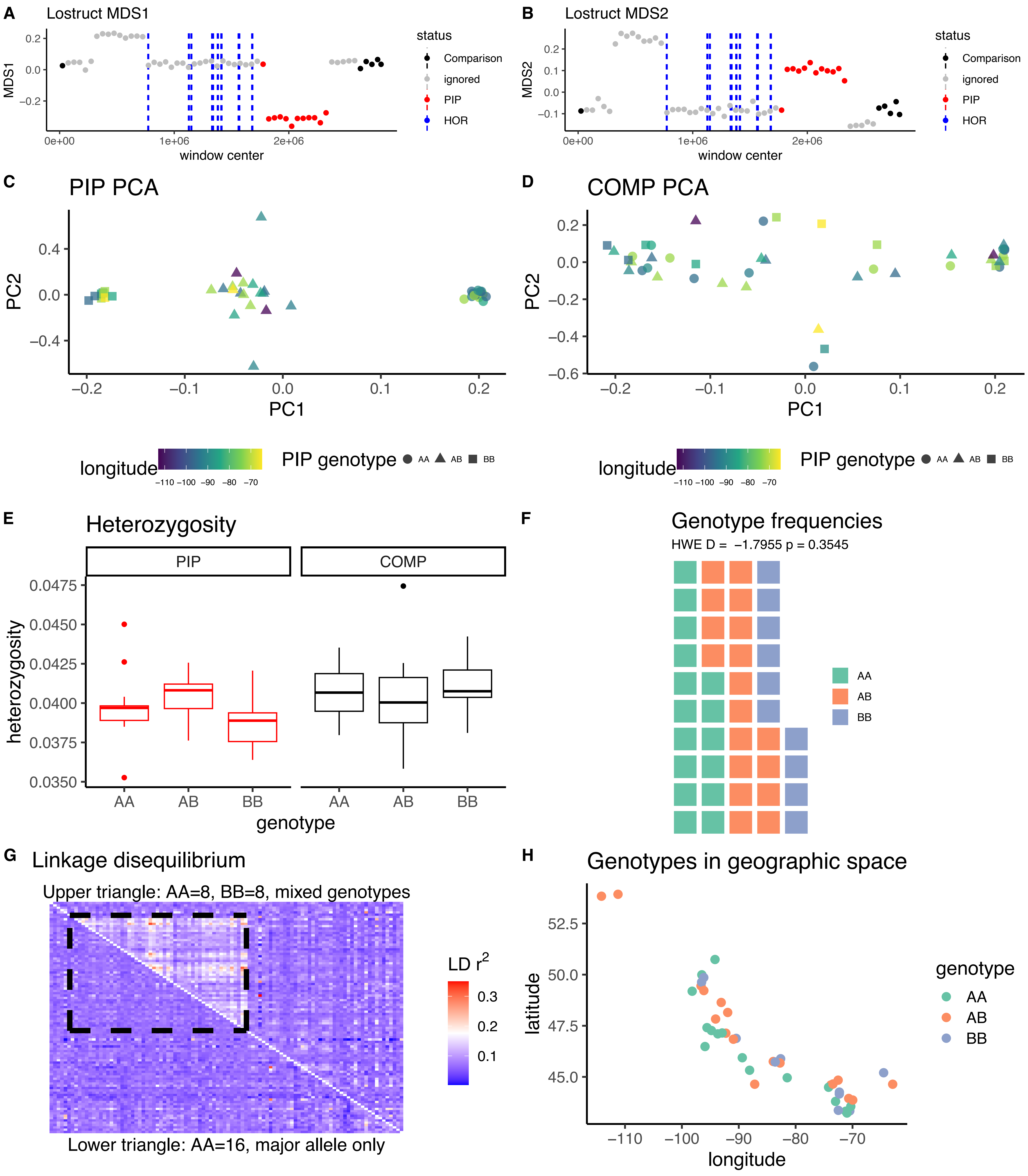

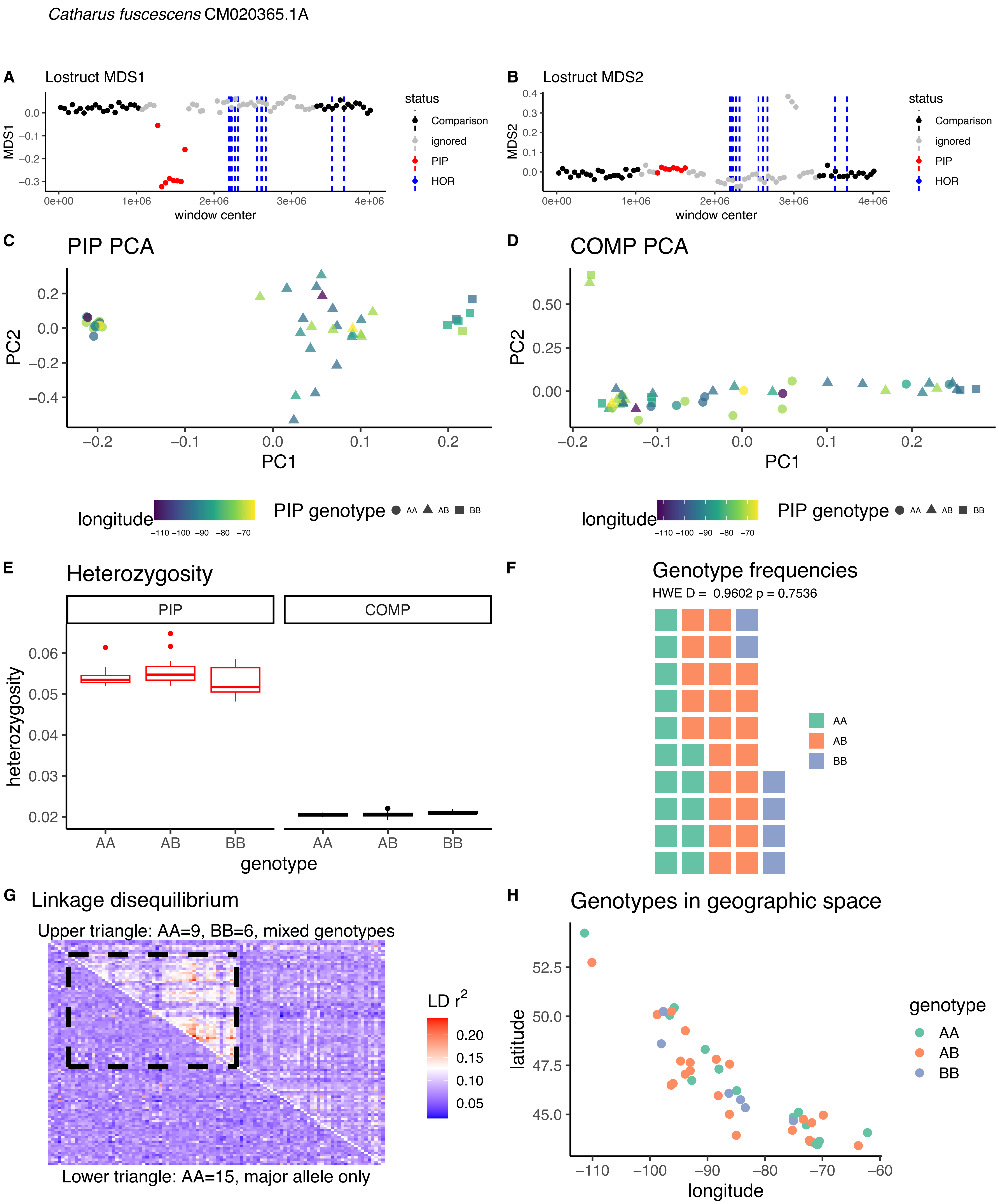

Catharus fuscescens CM020365.1A
A
Lostruct MDS1
status
Comparison
ignored
PIP
HOR
MDS1
window center
B
Lostruct MDS2
MDS2
window center
C
PIP PCA
PC2
PC1
D
COMP PCA
PC2
PC1
longitude
PIP genotype
AA
AB
BB
E
Heterozygosity
PIP
COMP
heterozygosity
genotype
F
Genotype frequencies
HWE D = 0.9602 p = 0.7536
G
Linkage disequilibrium
Upper triangle: AA=9, BB=6, mixed genotypes
LD r2
Lower triangle: AA=15, major allele only
H
Genotypes in geographic space
latitude
longitude
genotype

*Catharus fuscescens* CM020365.1B

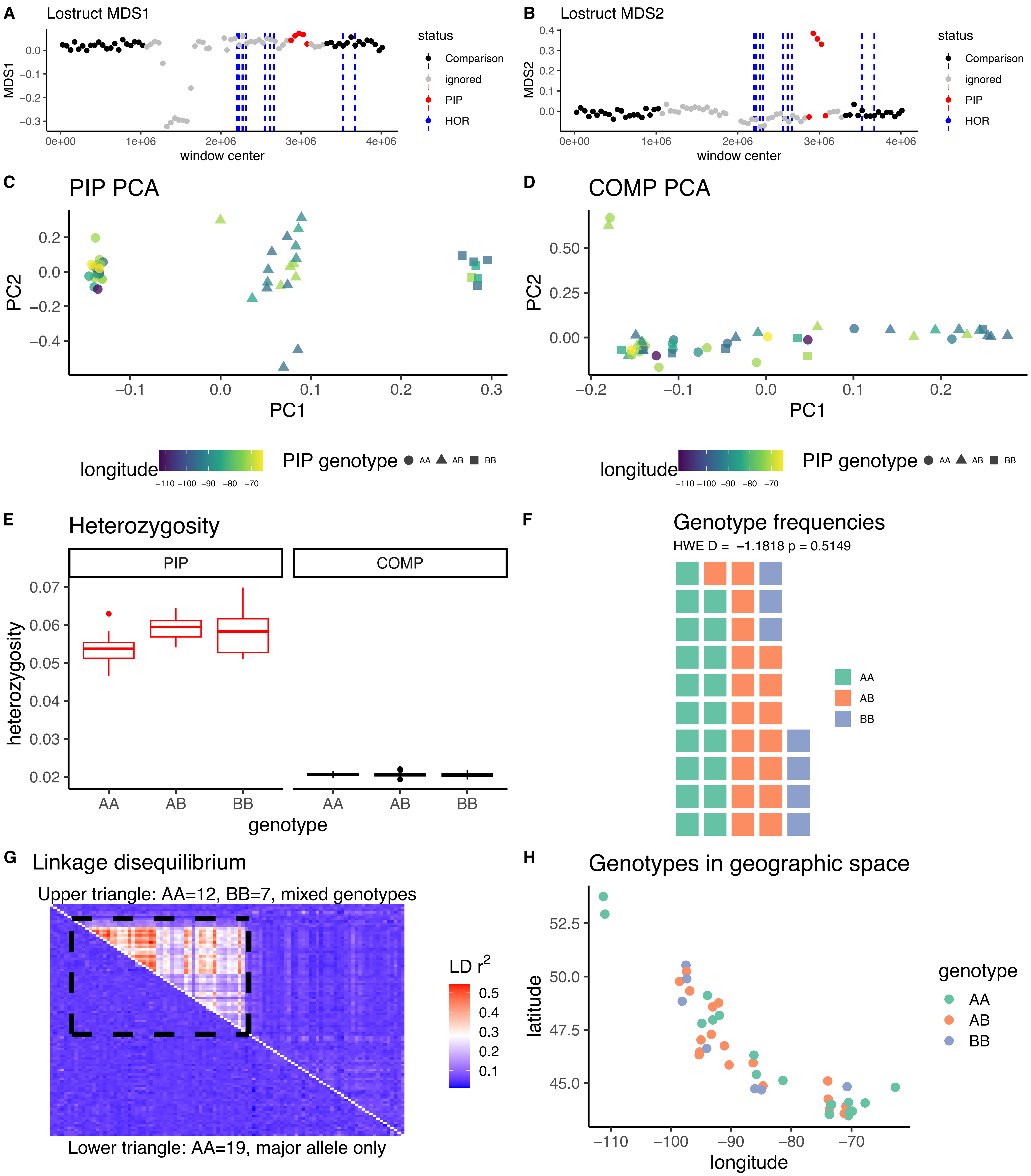

*Catharus fuscescens* CM020358.1

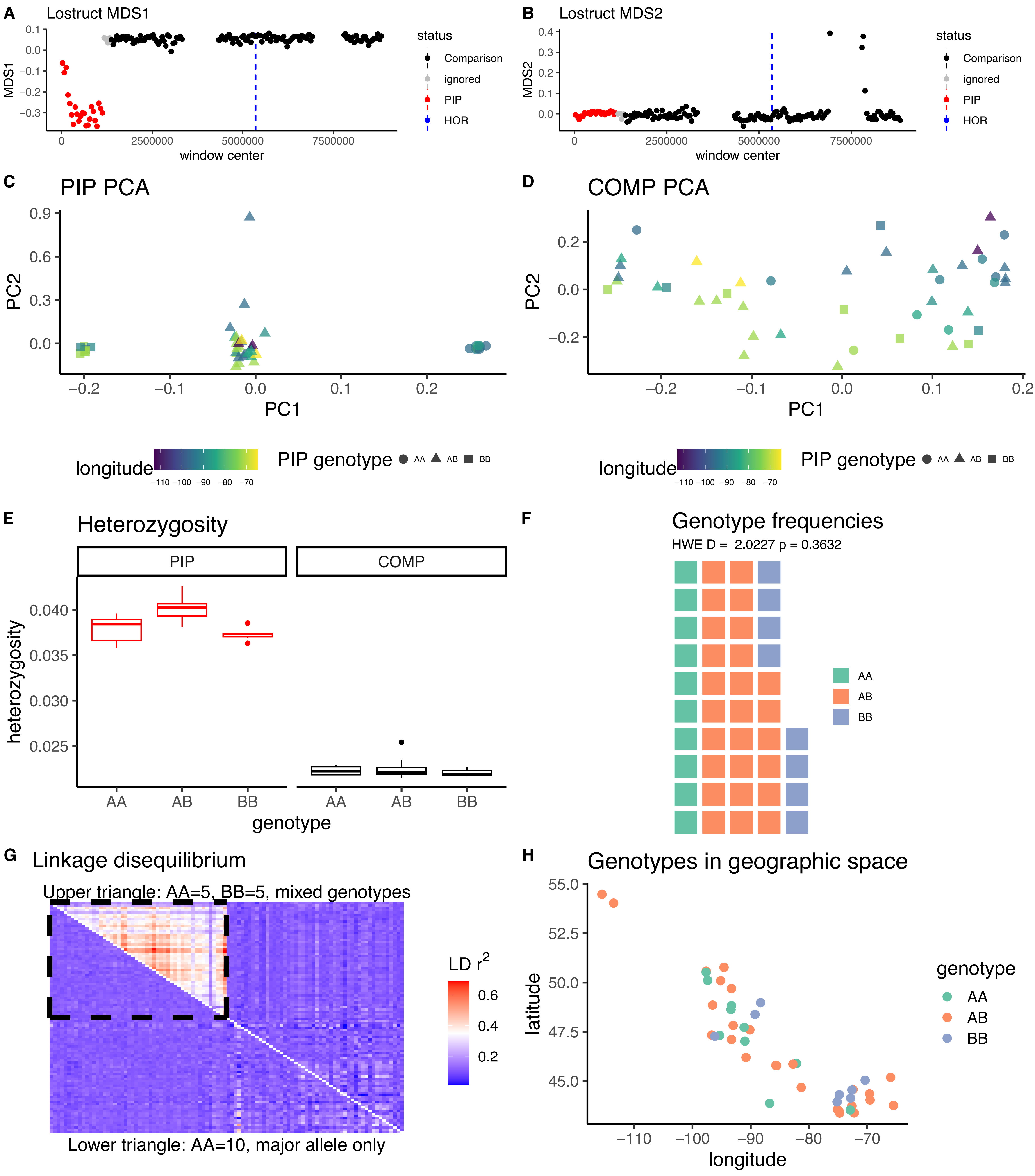

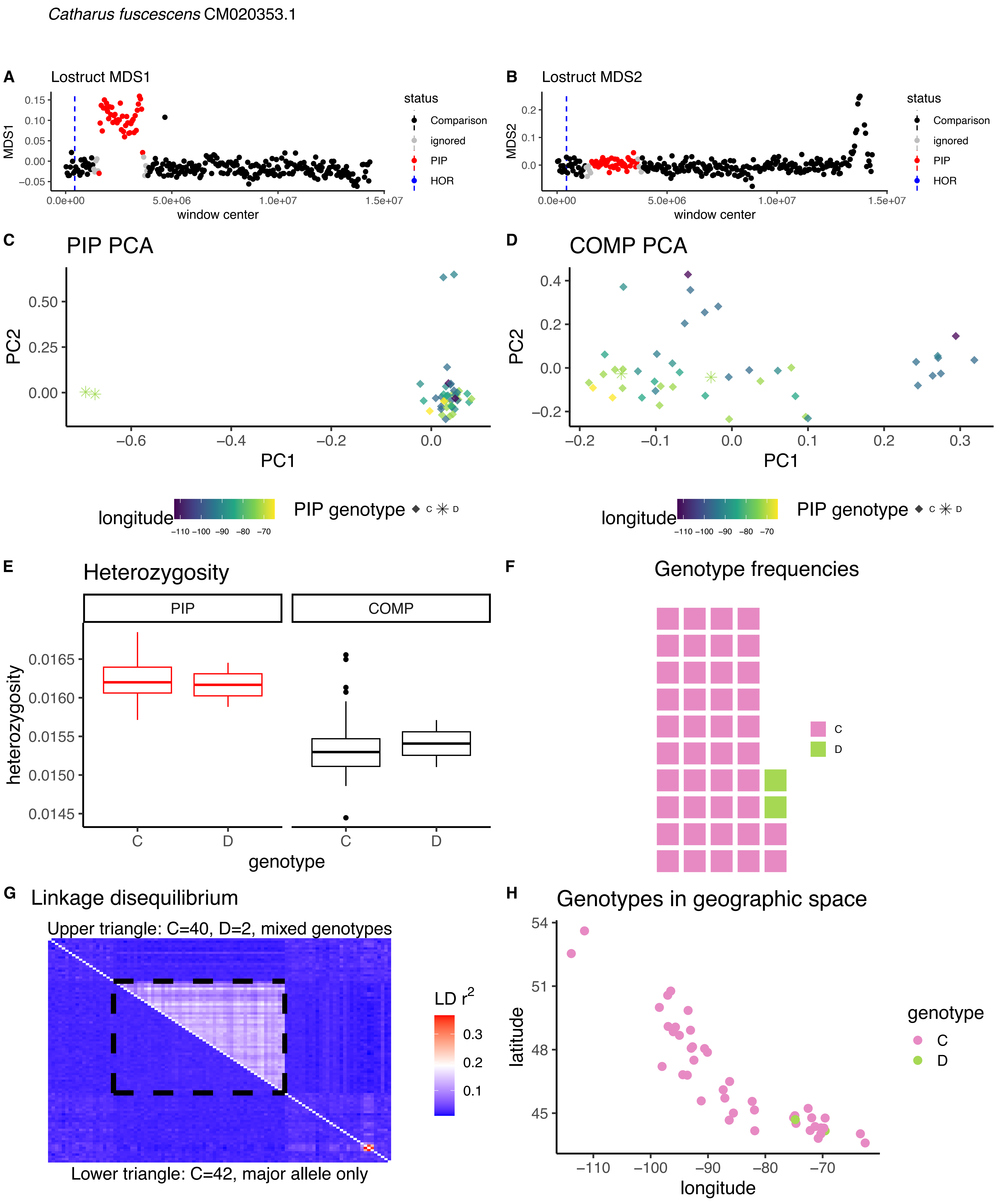

Catharus fuscescens CM020353.1
A
Lostruct MDS1
MDS1
window center
status
Comparison
ignored
PIP
HOR
B
Lostruct MDS2
MDS2
C
PIP PCA
PC1
PC2
D
COMP PCA
longitude
PIP genotype
E
Heterozygosity
PIP
COMP
heterozygosity
genotype
F
Genotype frequencies
C
D
G
Linkage disequilibrium
Upper triangle: C=40, D=2, mixed genotypes
Lower triangle: C=42, major allele only
LD r2
H
Genotypes in geographic space
latitude
longitude
genotype

*Catharus fuscescens* CM020349.1

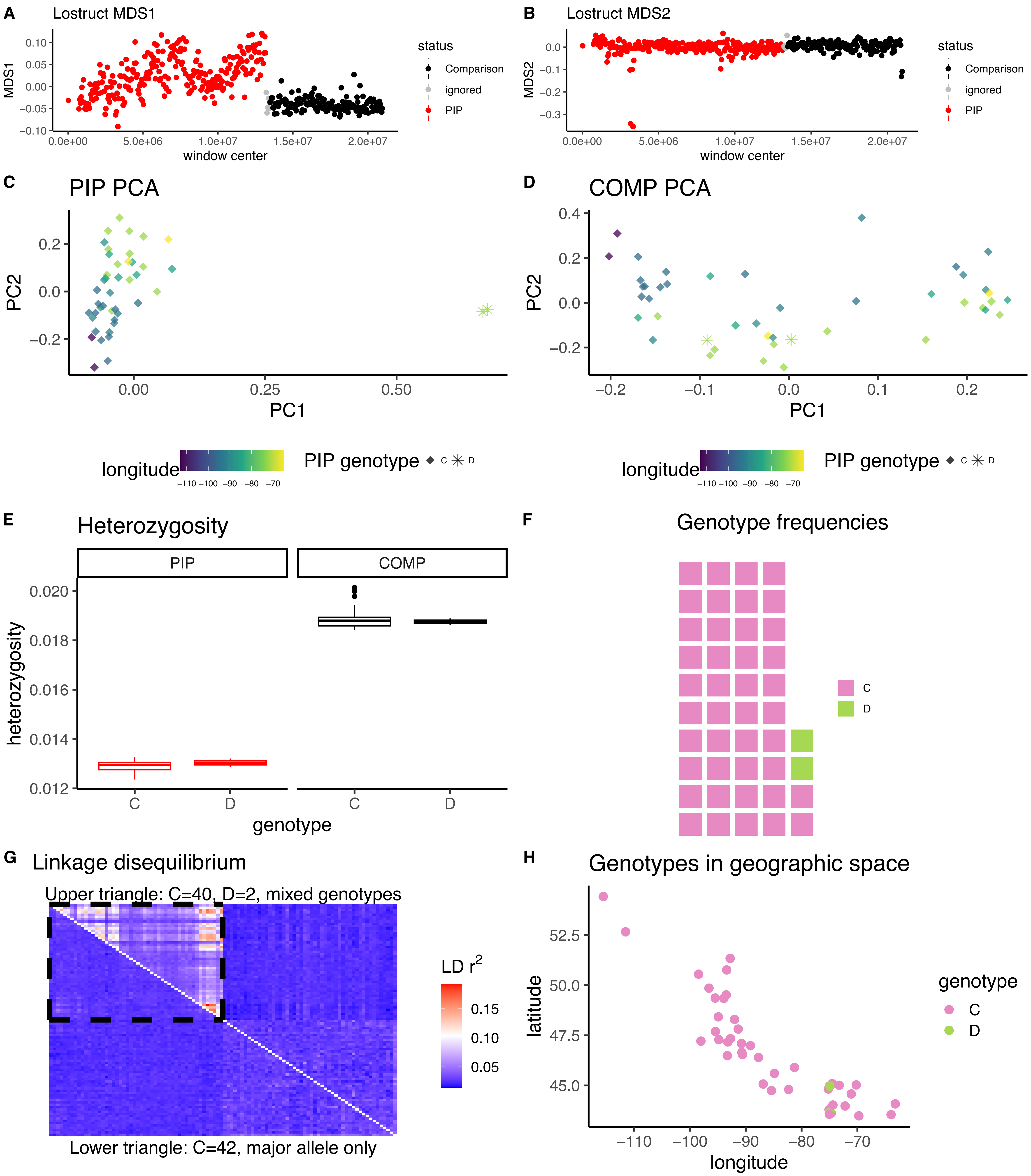

*Catharus fuscescens* CM020348.1

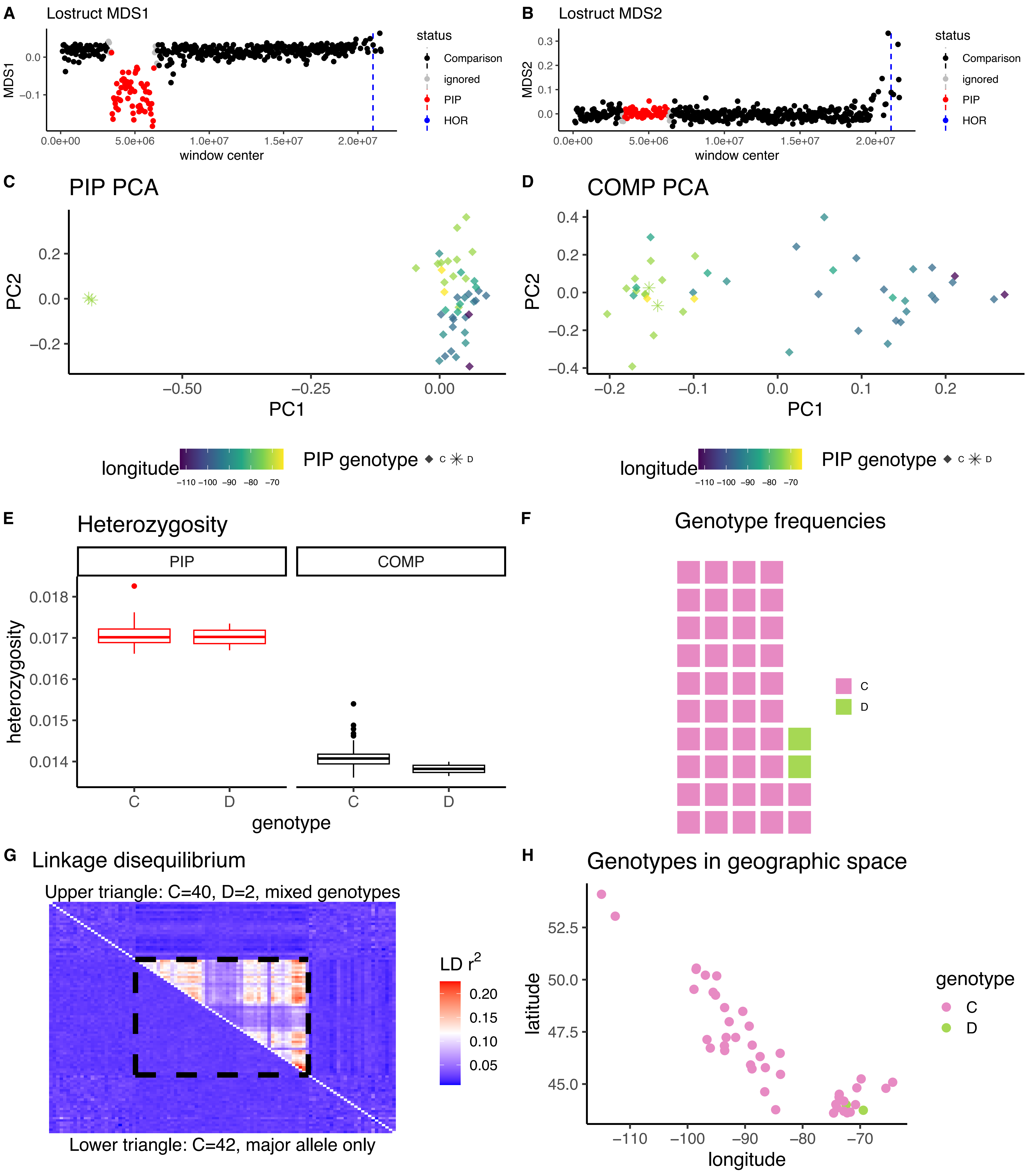

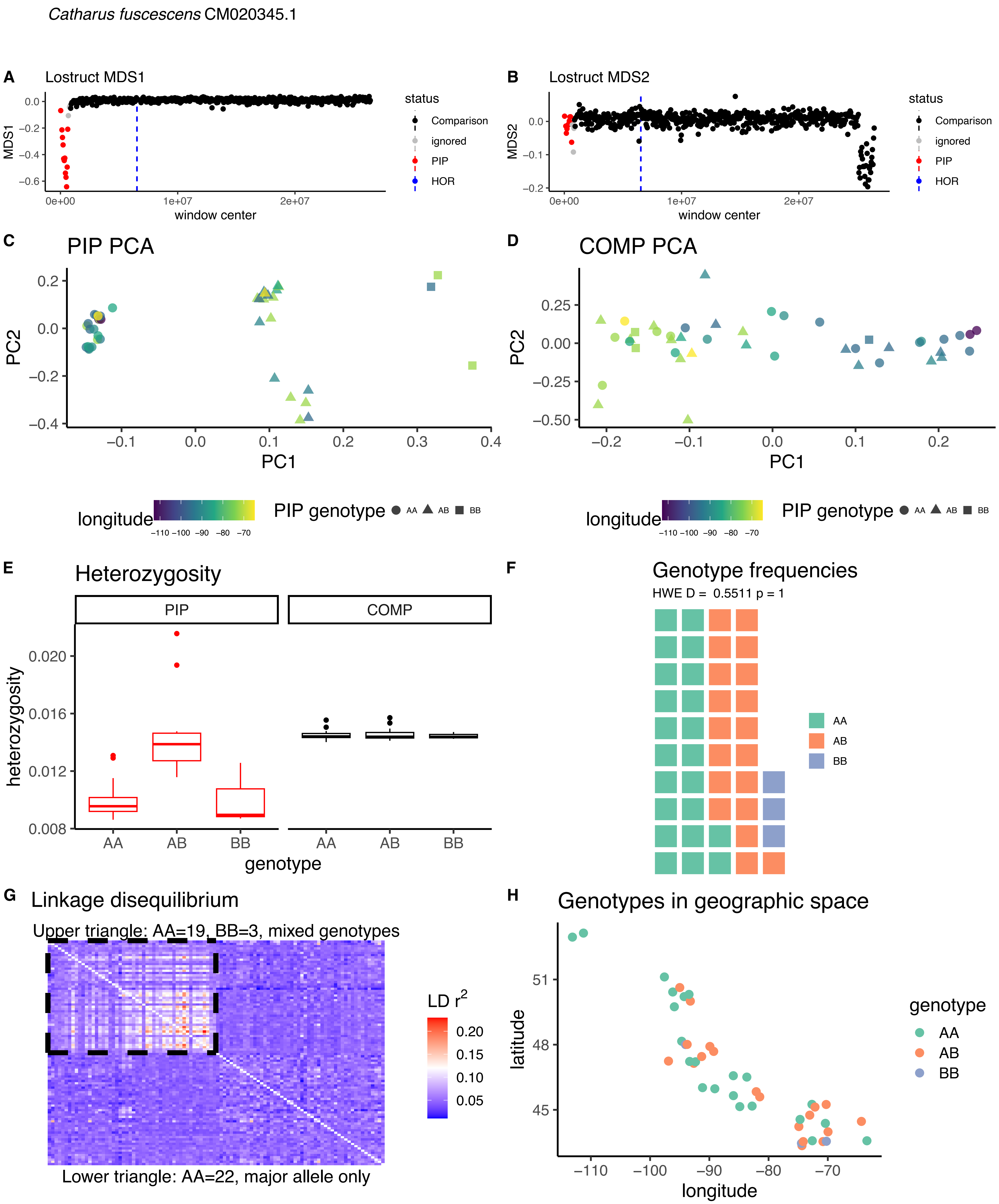
Catharus fuscescens CM020345.1
A
Lostruct MDS1
MDS1
window center
status
Comparison
ignored
PIP
HOR
B
Lostruct MDS2
MDS2
C
PIP PCA
PC1
PC2
D
COMP PCA
longitude
PIP genotype
AA
AB
BB
E
Heterozygosity
PIP
COMP
heterozygosity
genotype
F
Genotype frequencies
HWE D = 0.5511 p = 1
G
Linkage disequilibrium
Upper triangle: AA=19, BB=3, mixed genotypes
LD r2
Lower triangle: AA=22, major allele only
H
Genotypes in geographic space
latitude

*Catharus fuscescens* CM020341.1A

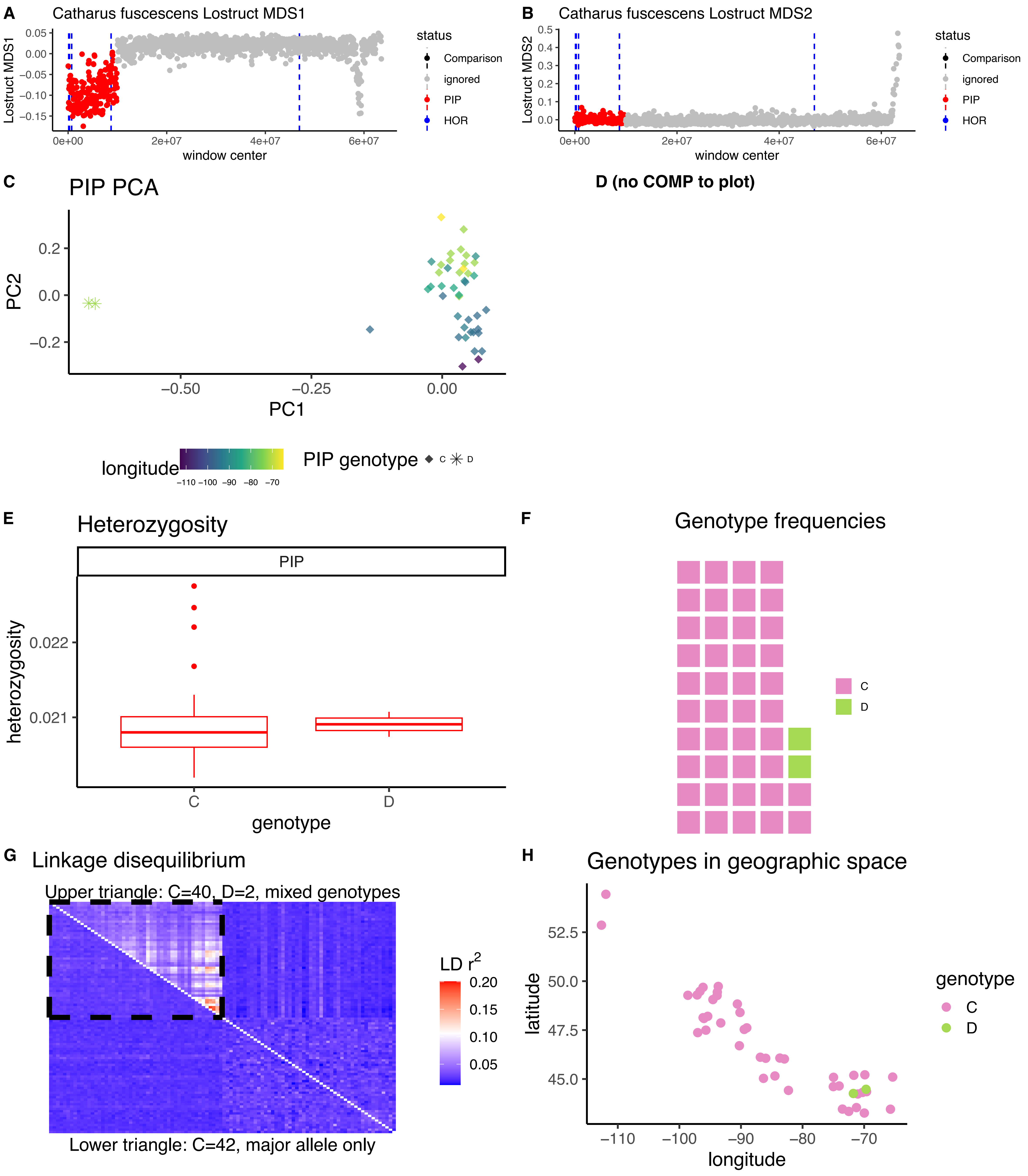

*Catharus fuscescens* CM020341.1B

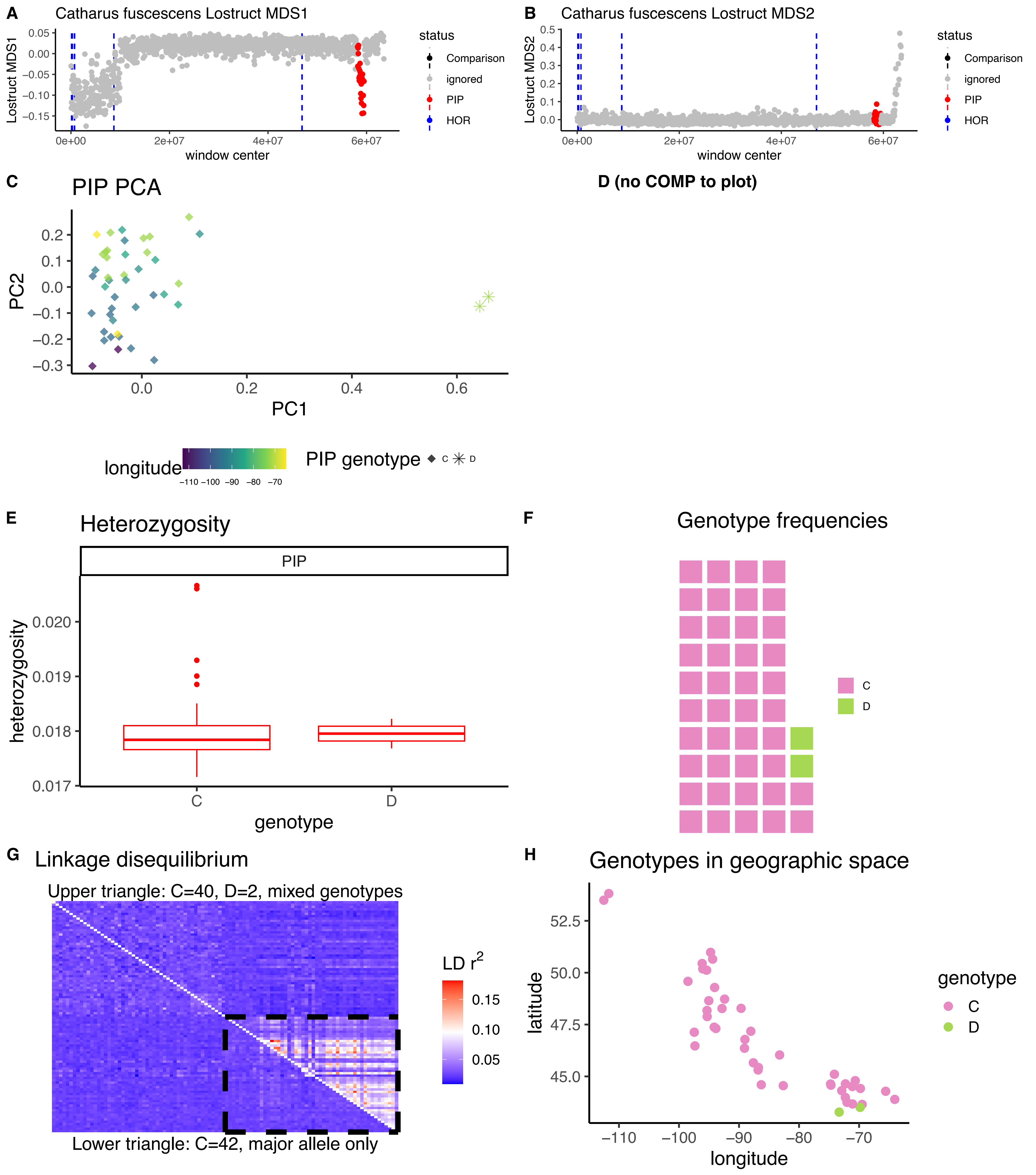

*Catharus fuscescens* CM020341.1C

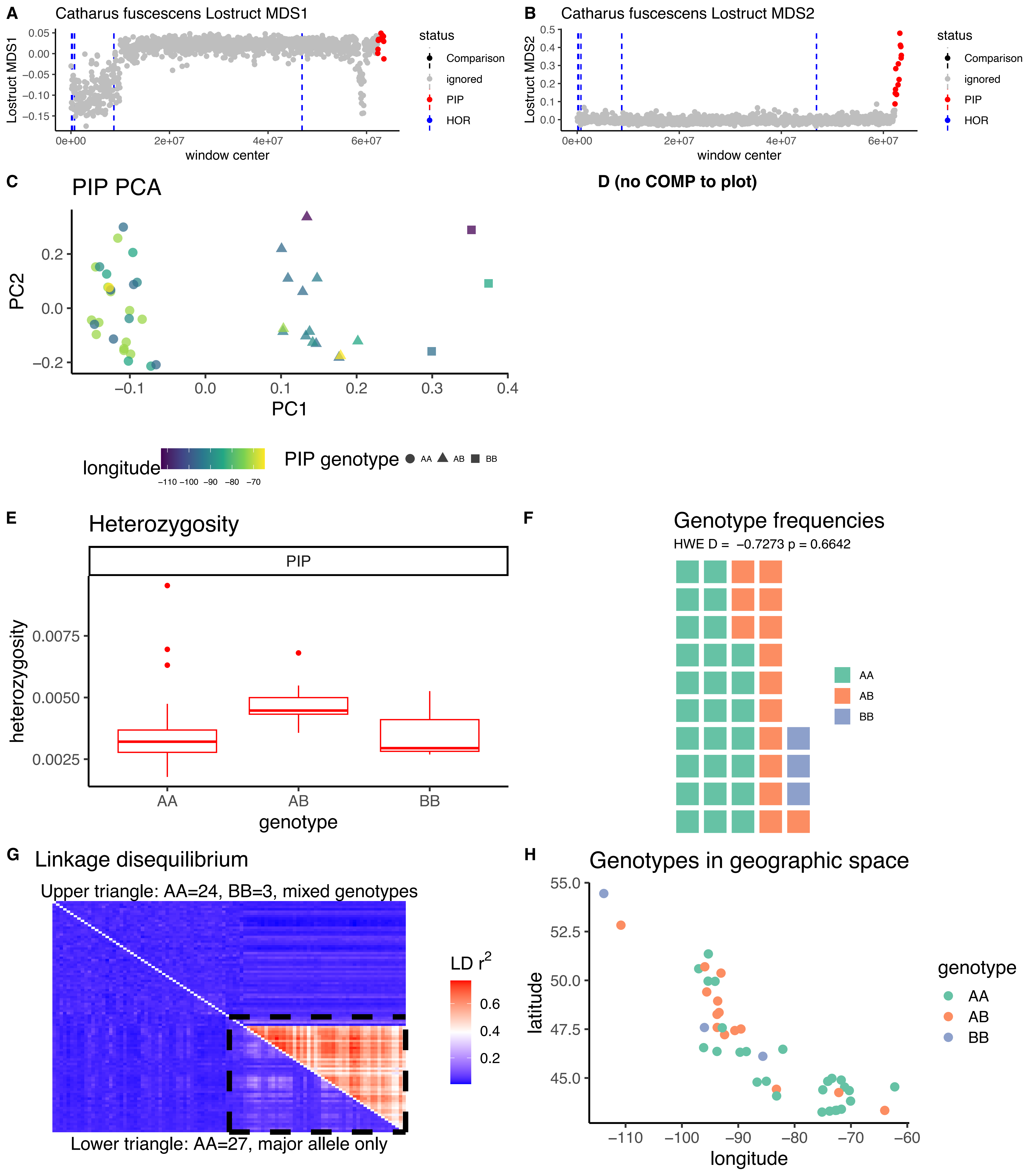

*Catharus fuscescens* CM020338.1

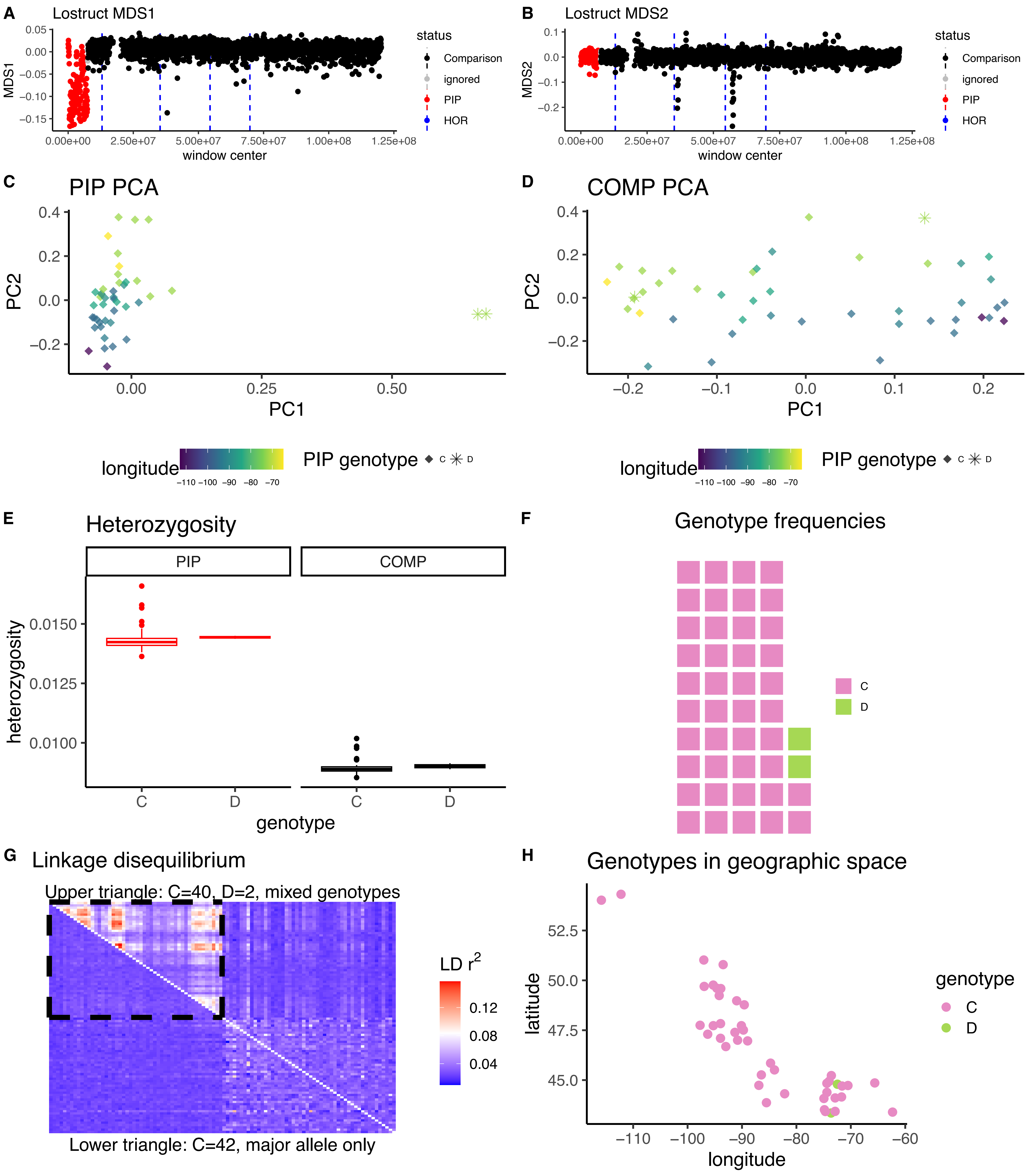

*Catharus guttatus* CM020371.1A

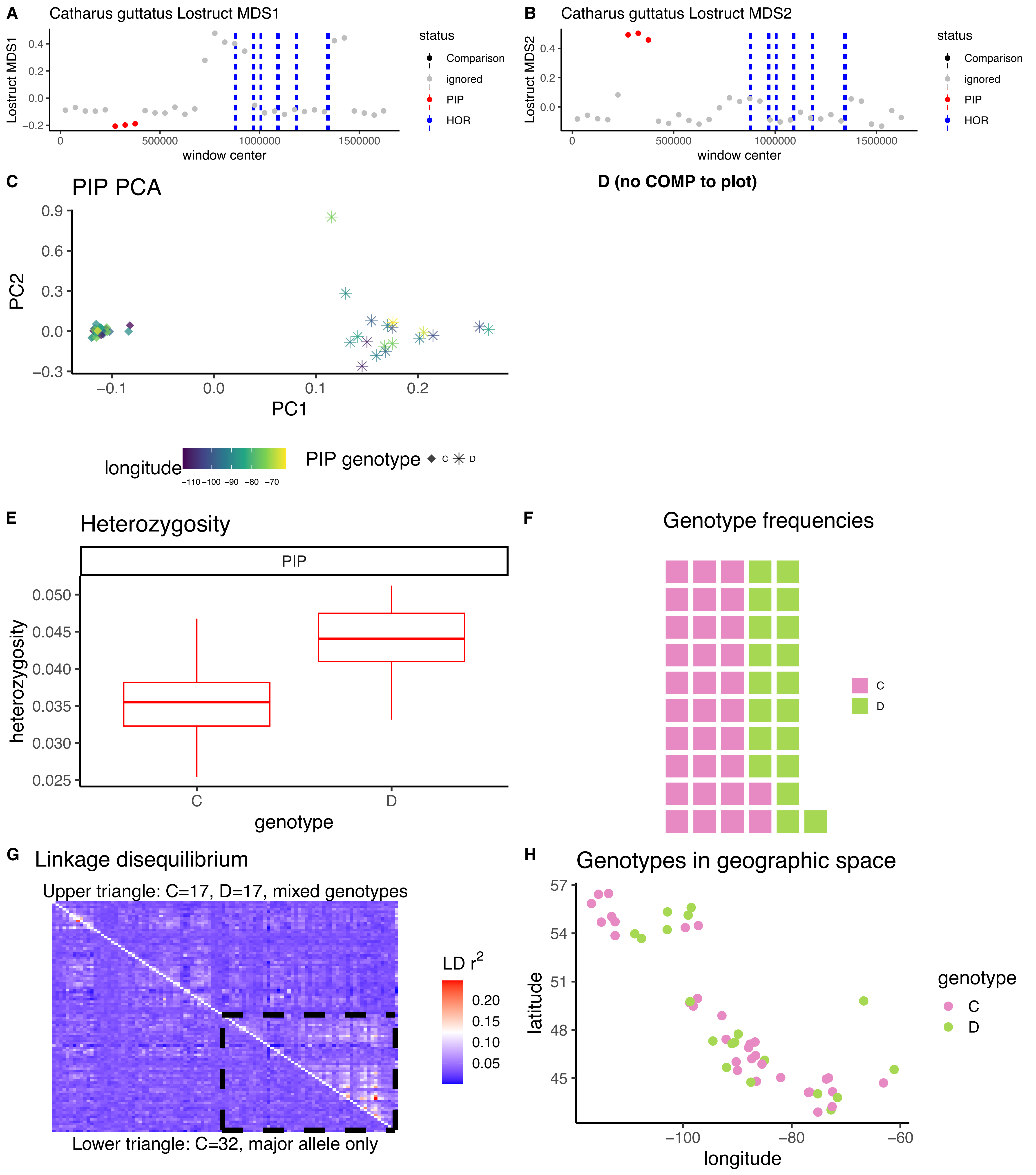

*Catharus guttatus* CM020371.1B

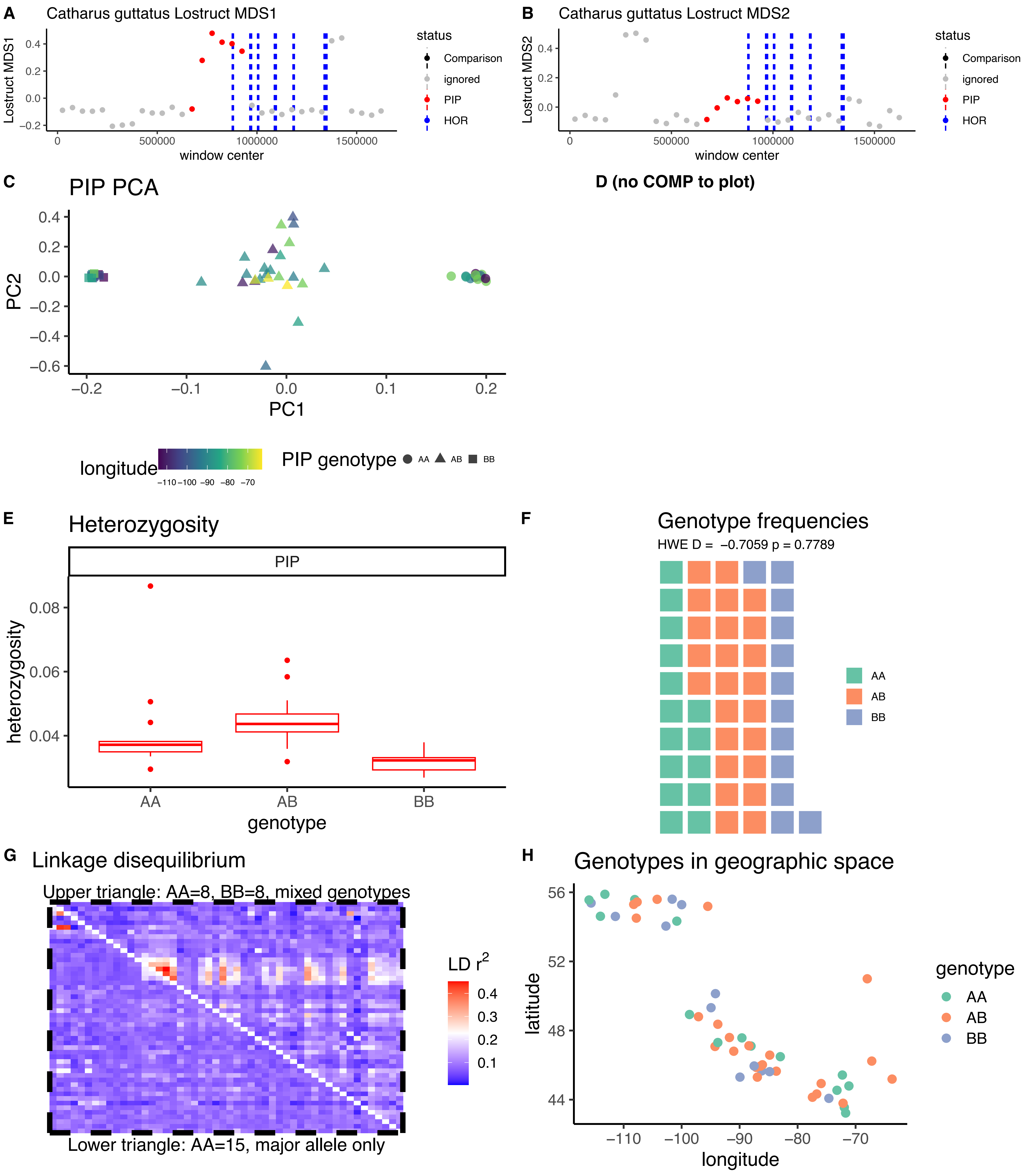

*Catharus guttatus* CM020371.1C

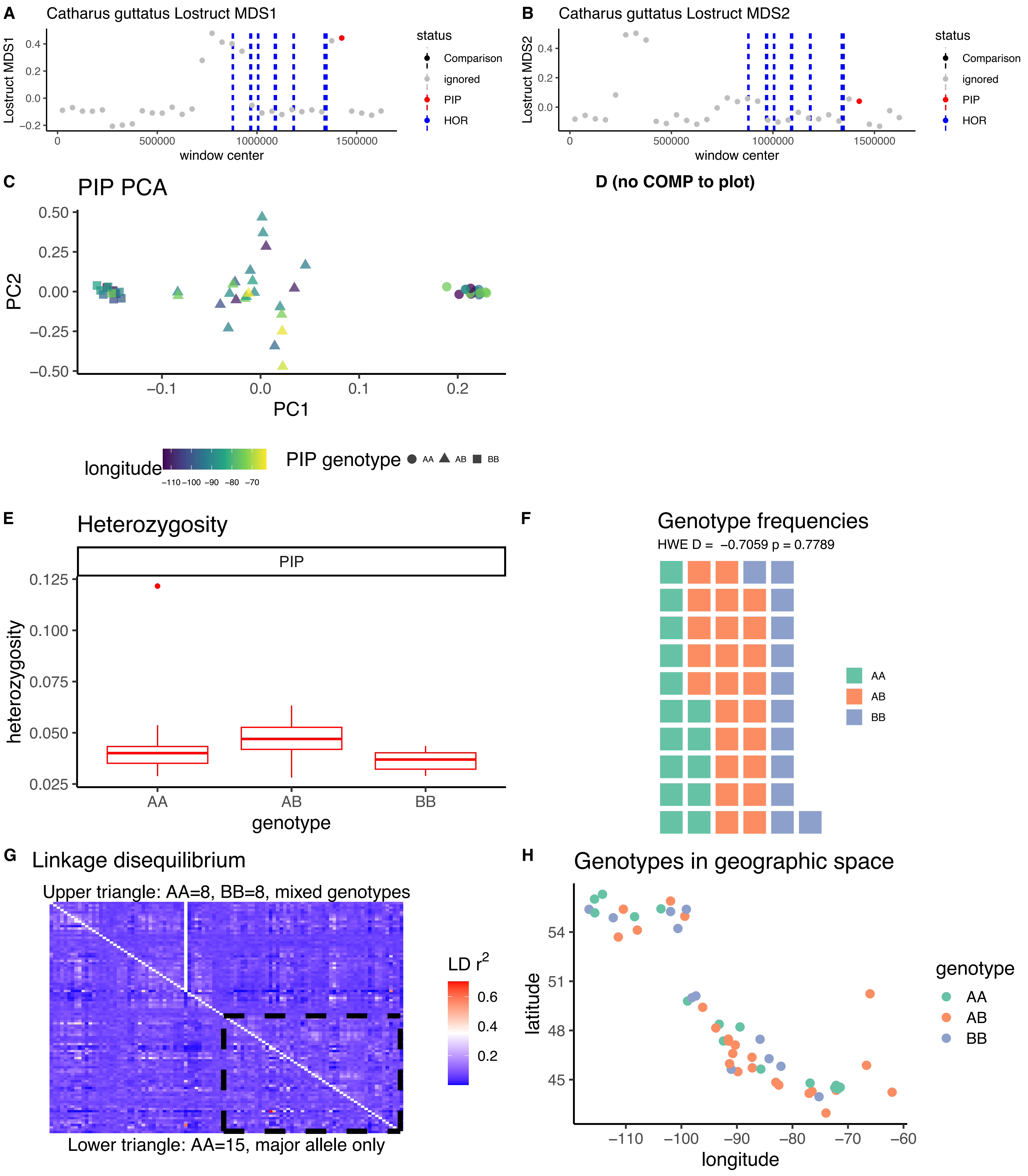

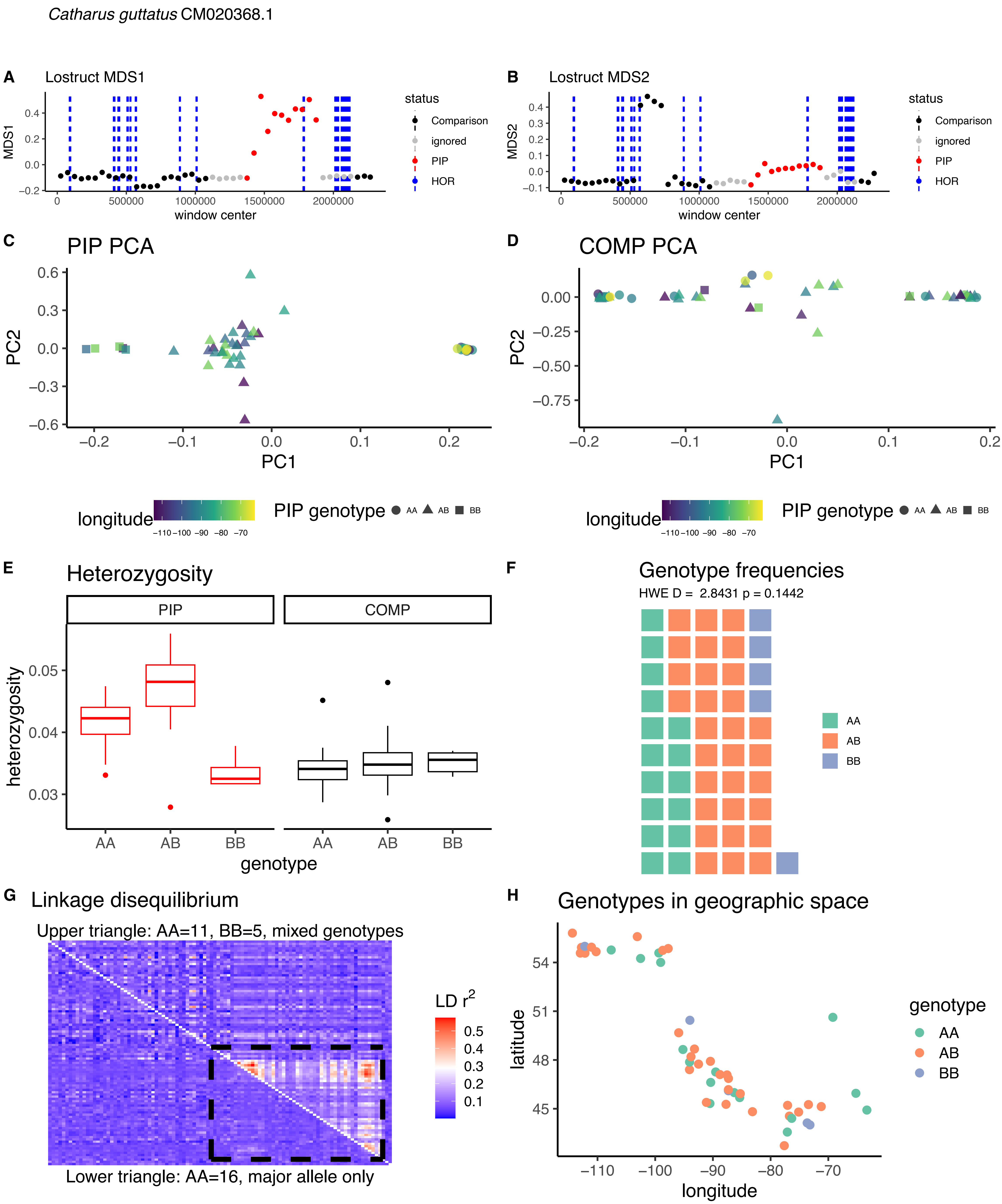

Catharus guttatus CM020368.1
A
Lostruct MDS1
MDS1
window center
status
Comparison
ignored
PIP
HOR
B
Lostruct MDS2
MDS2
window center
C
PIP PCA
PC1
PC2
D
COMP PCA
longitude
PIP genotype
AA
AB
BB
E
Heterozygosity
PIP
COMP
heterozygosity
genotype
F
Genotype frequencies
HWE D = 2.8431 p = 0.1442
G
Linkage disequilibrium
Upper triangle: AA=11, BB=5, mixed genotypes
LD r2
Lower triangle: AA=16, major allele only
H
Genotypes in geographic space
latitude
longitude
genotype

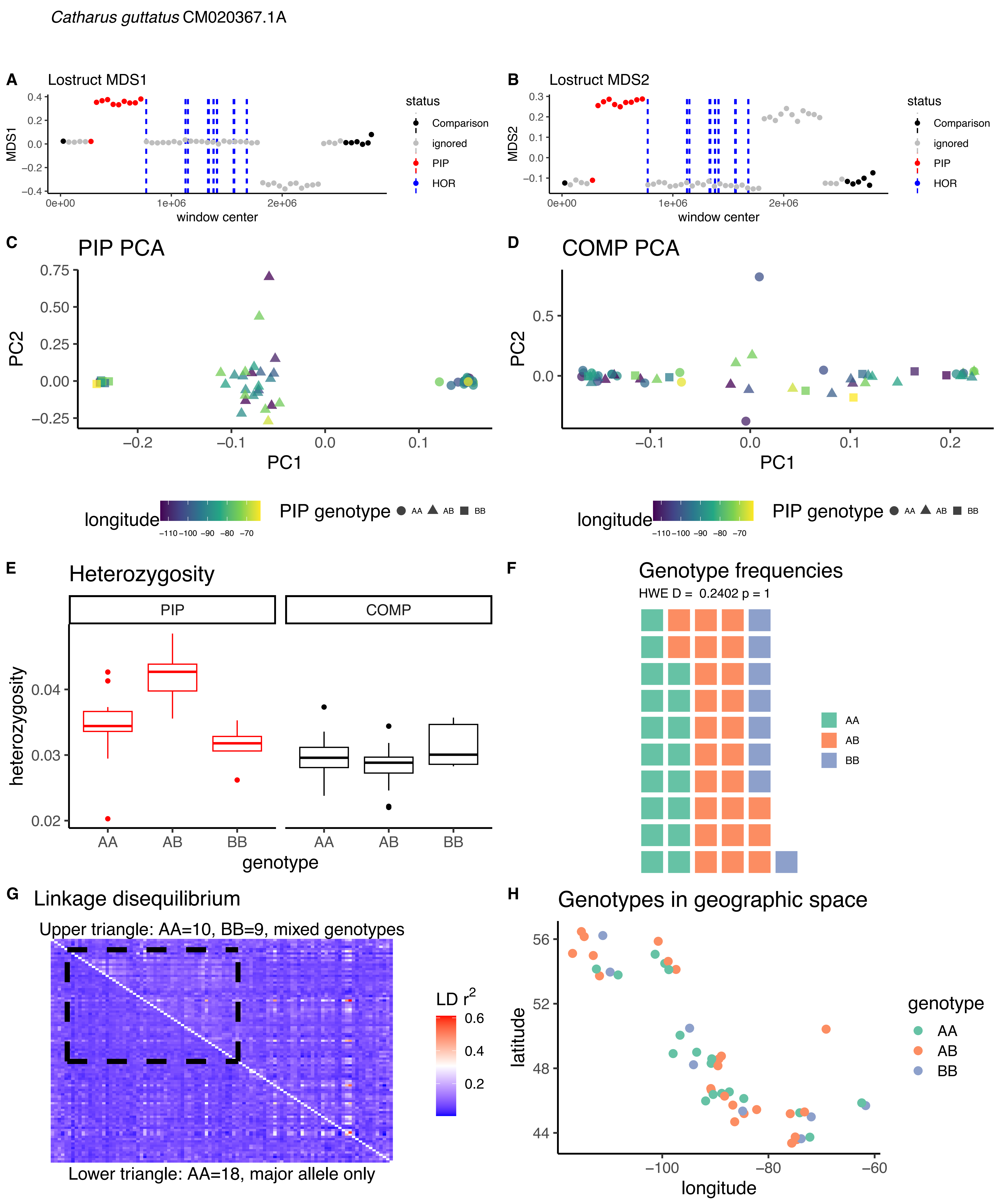

Catharus guttatus CM020367.1A
A
Lostruct MDS1
MDS1
window center
status
Comparison
ignored
PIP
HOR
B
Lostruct MDS2
MDS2
C
PIP PCA
PC2
PC1
D
COMP PCA
longitude
PIP genotype
AA
AB
BB
E
Heterozygosity
PIP
COMP
heterozygosity
genotype
F
Genotype frequencies
HWE D = 0.2402 p = 1
G
Linkage disequilibrium
Upper triangle: AA=10, BB=9, mixed genotypes
LD r2
Lower triangle: AA=18, major allele only
H
Genotypes in geographic space
latitude
longitude
genotype

*Catharus guttatus* CM020367.1B

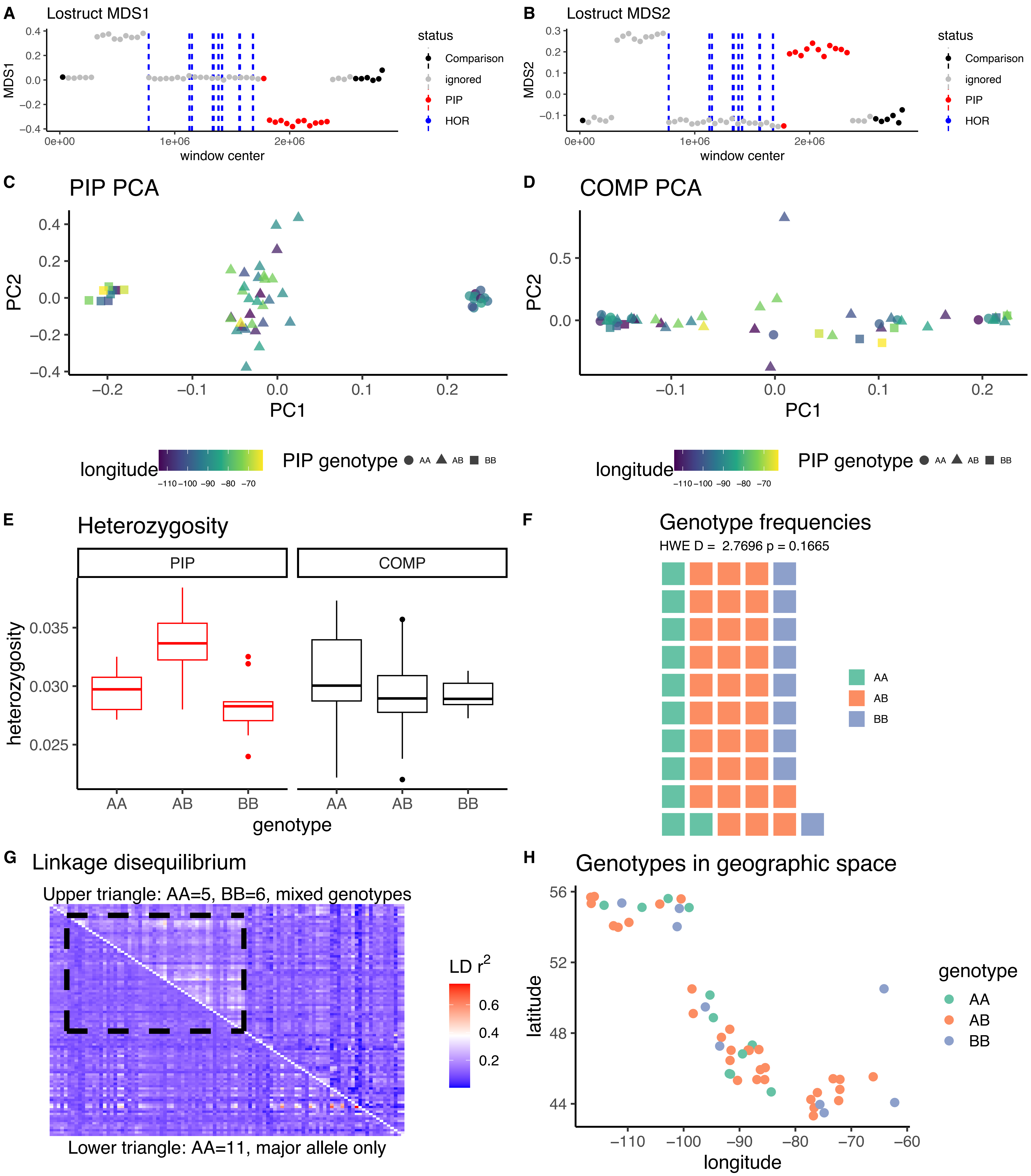

*Catharus guttatus* CM020365.1A

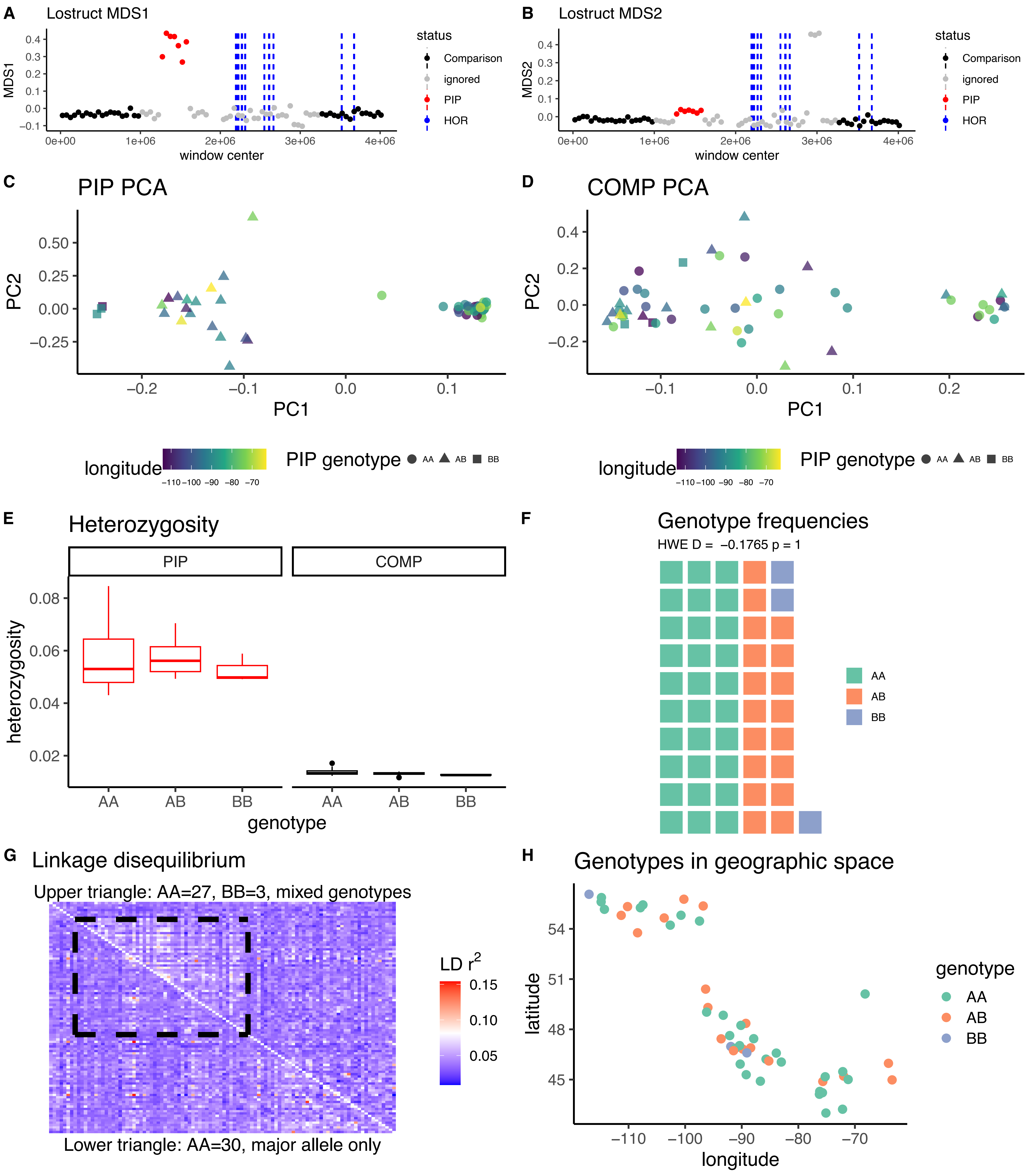

*Catharus guttatus* CM020365.1B

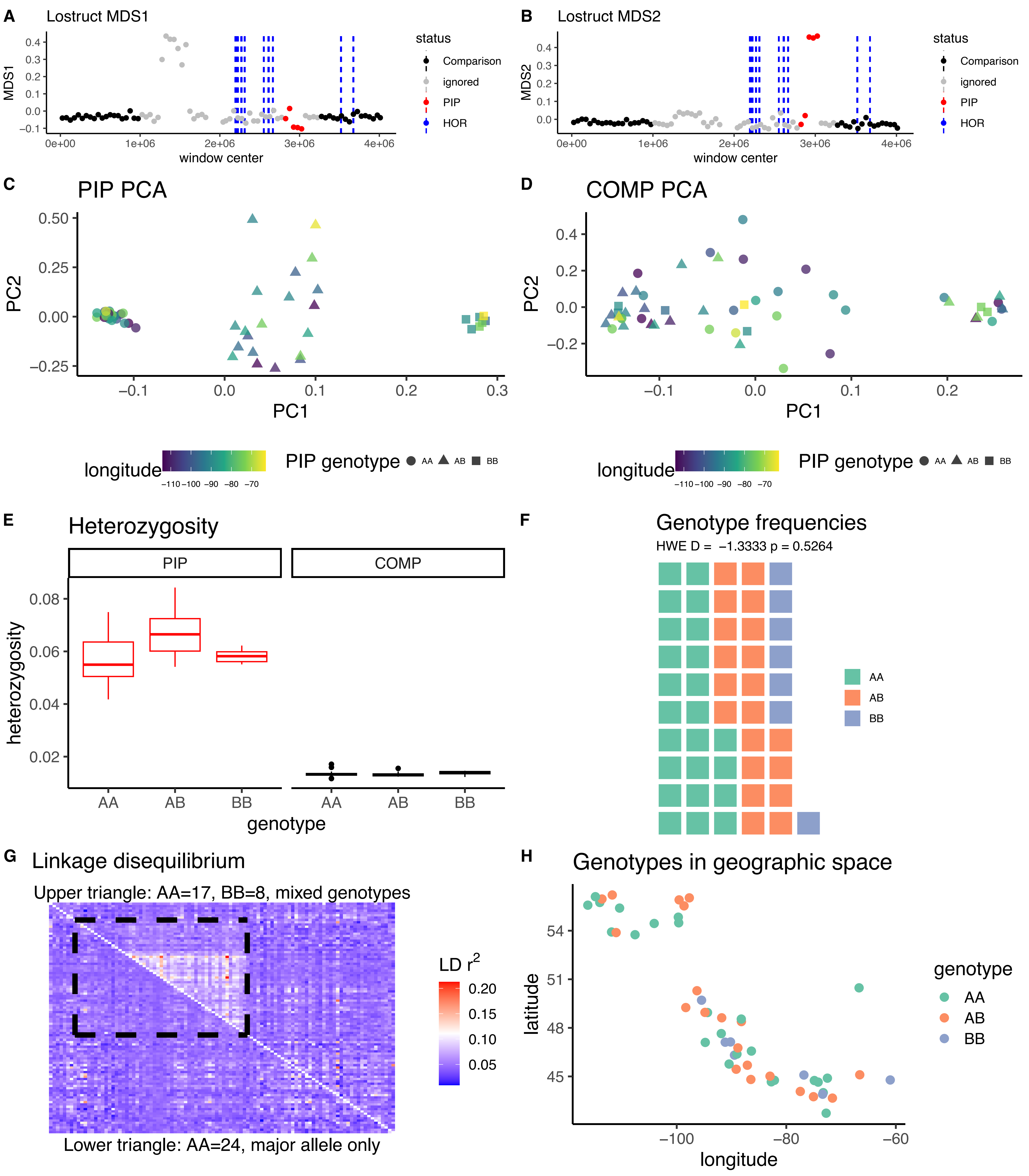

*Catharus ustulatus* CM020371.1A

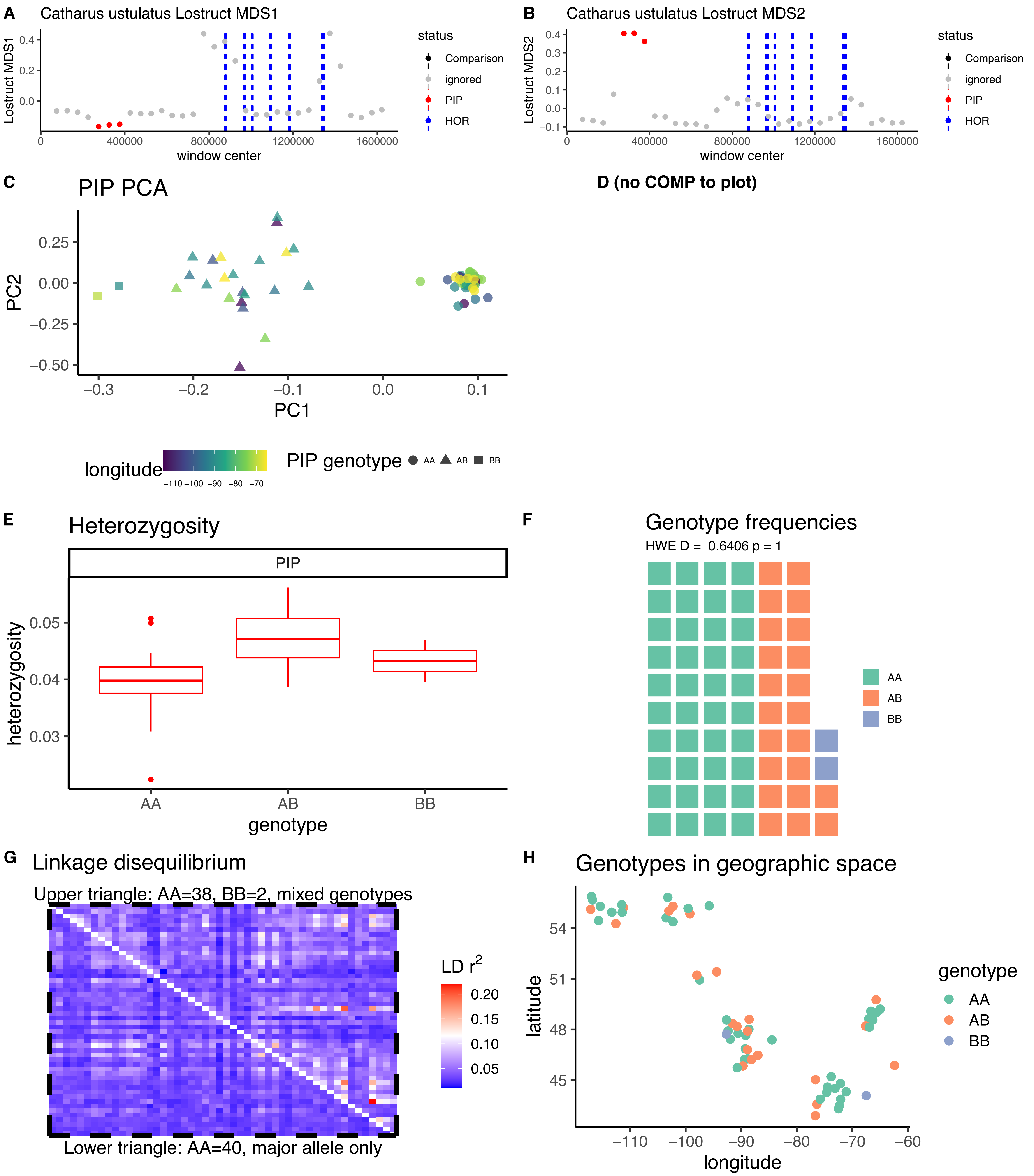

*Catharus ustulatus* CM020371.1B

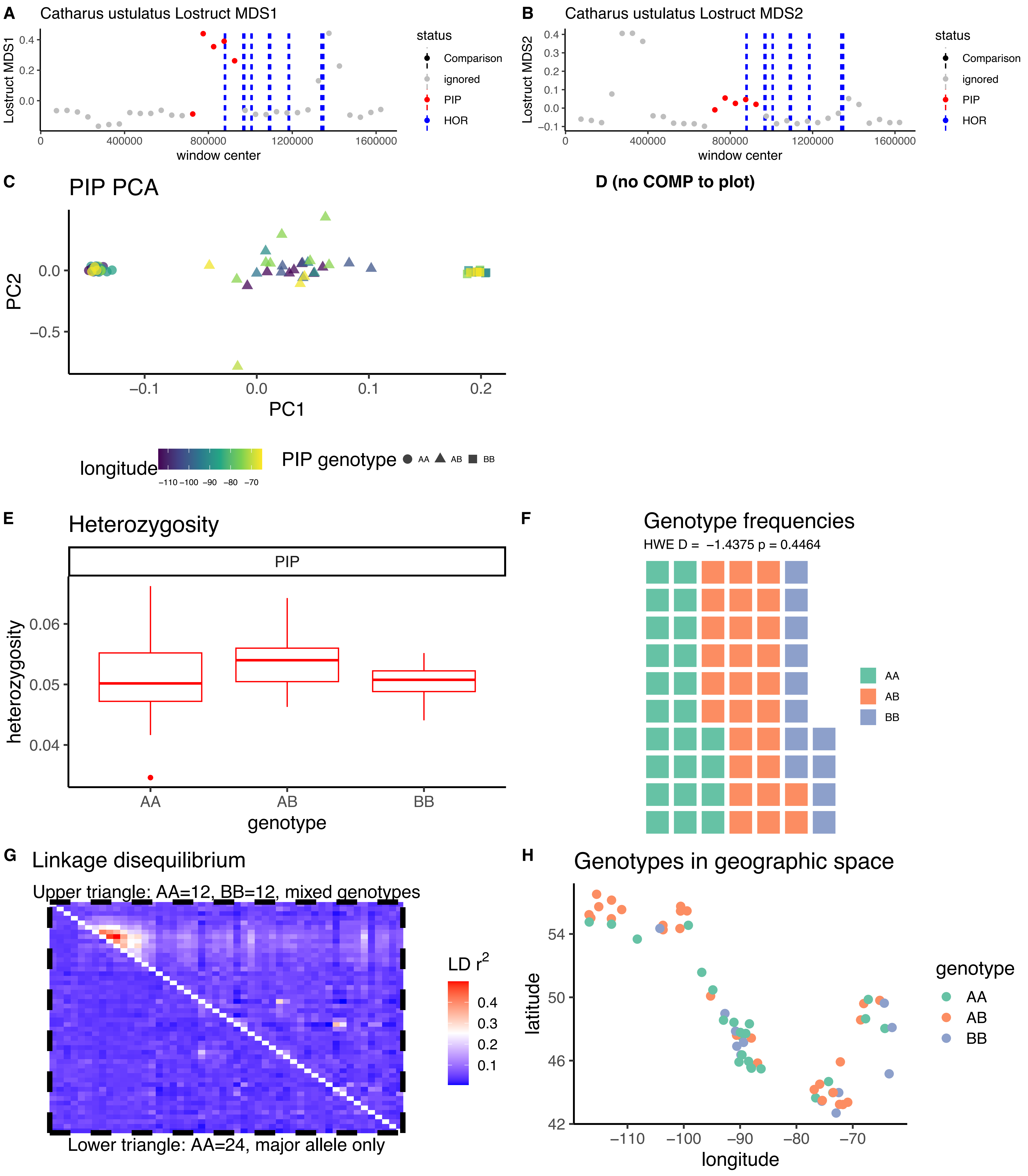

*Catharus ustulatus* CM020371.1C

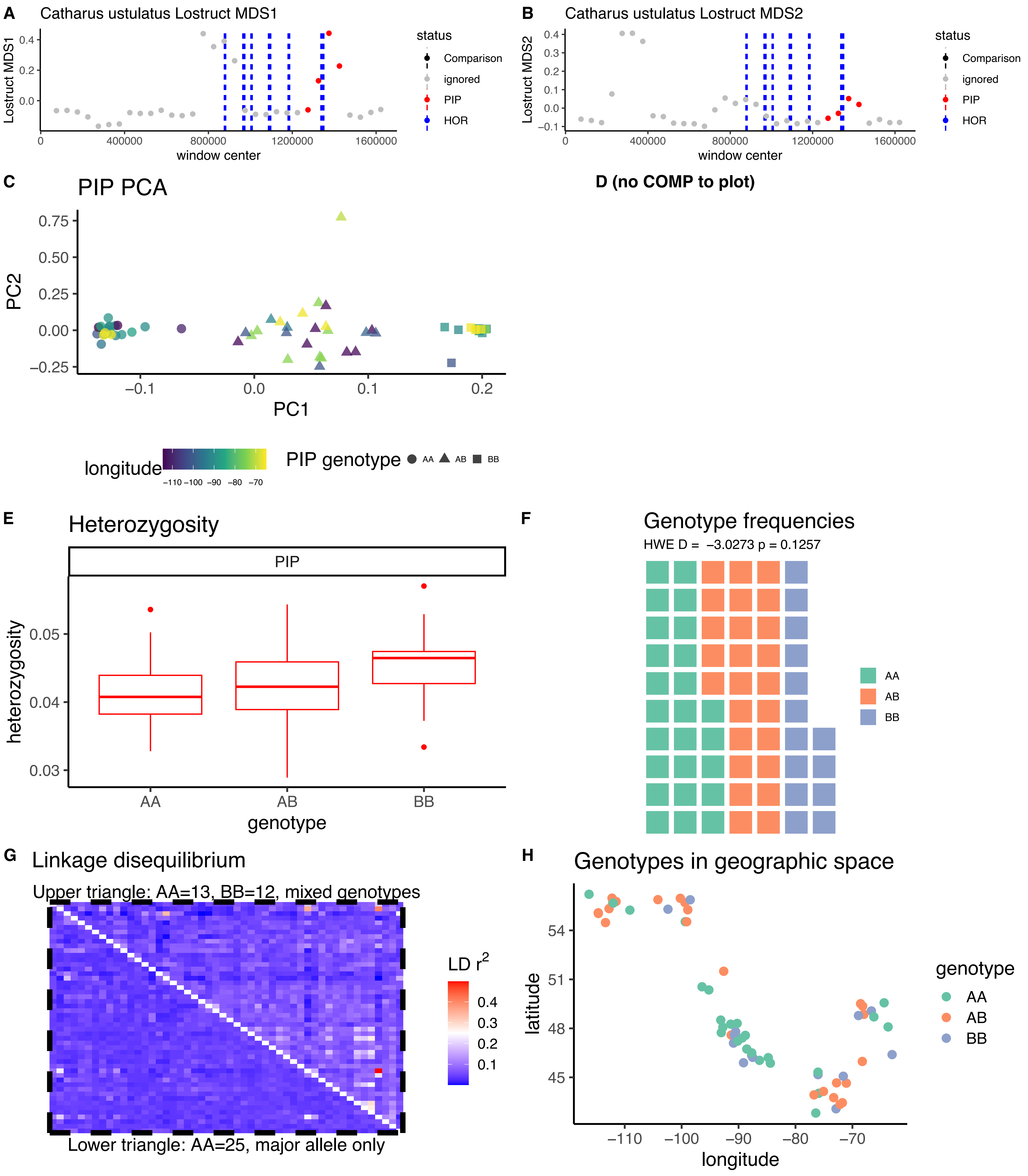

*Catharus ustulatus* CM020368.1

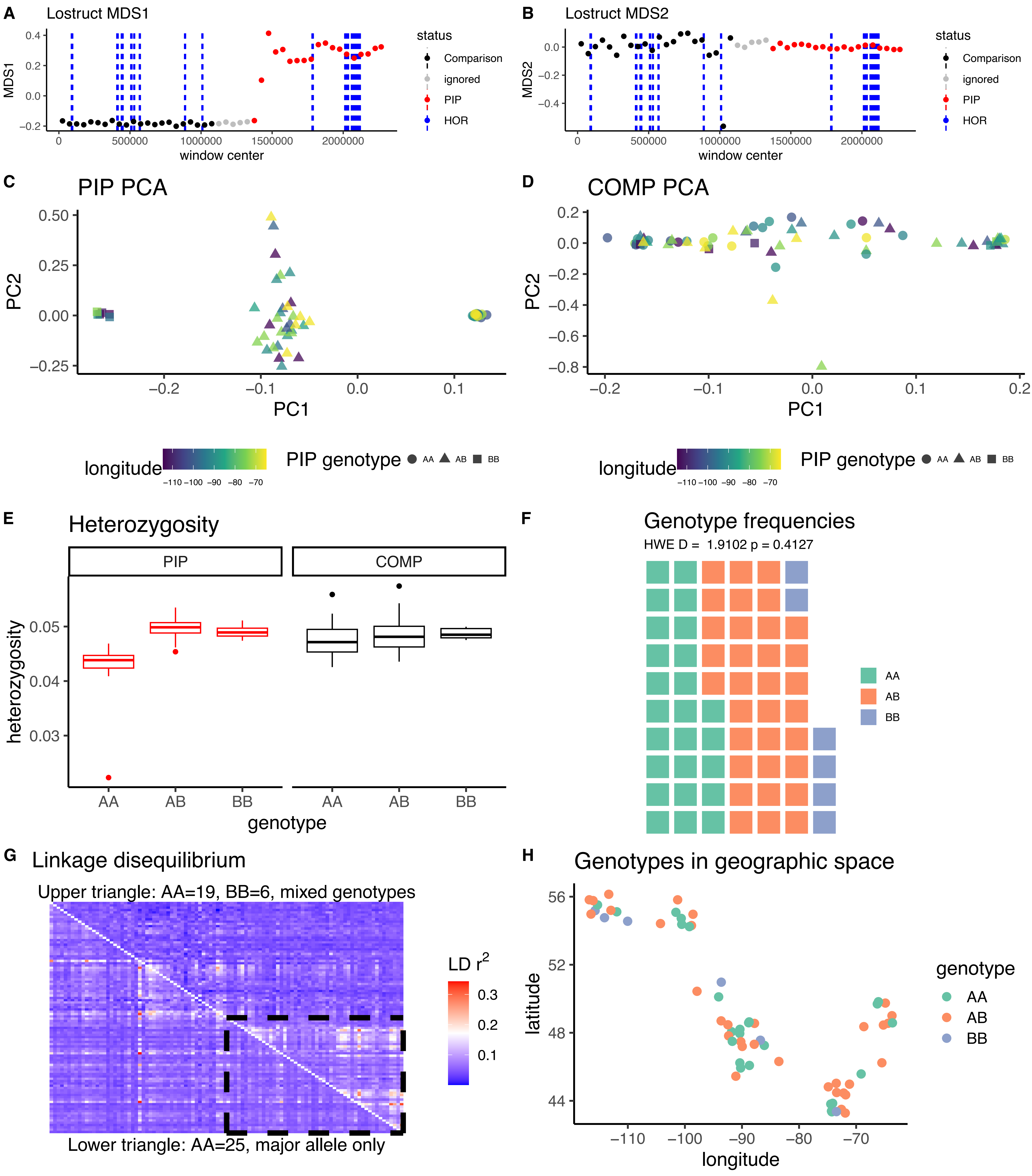

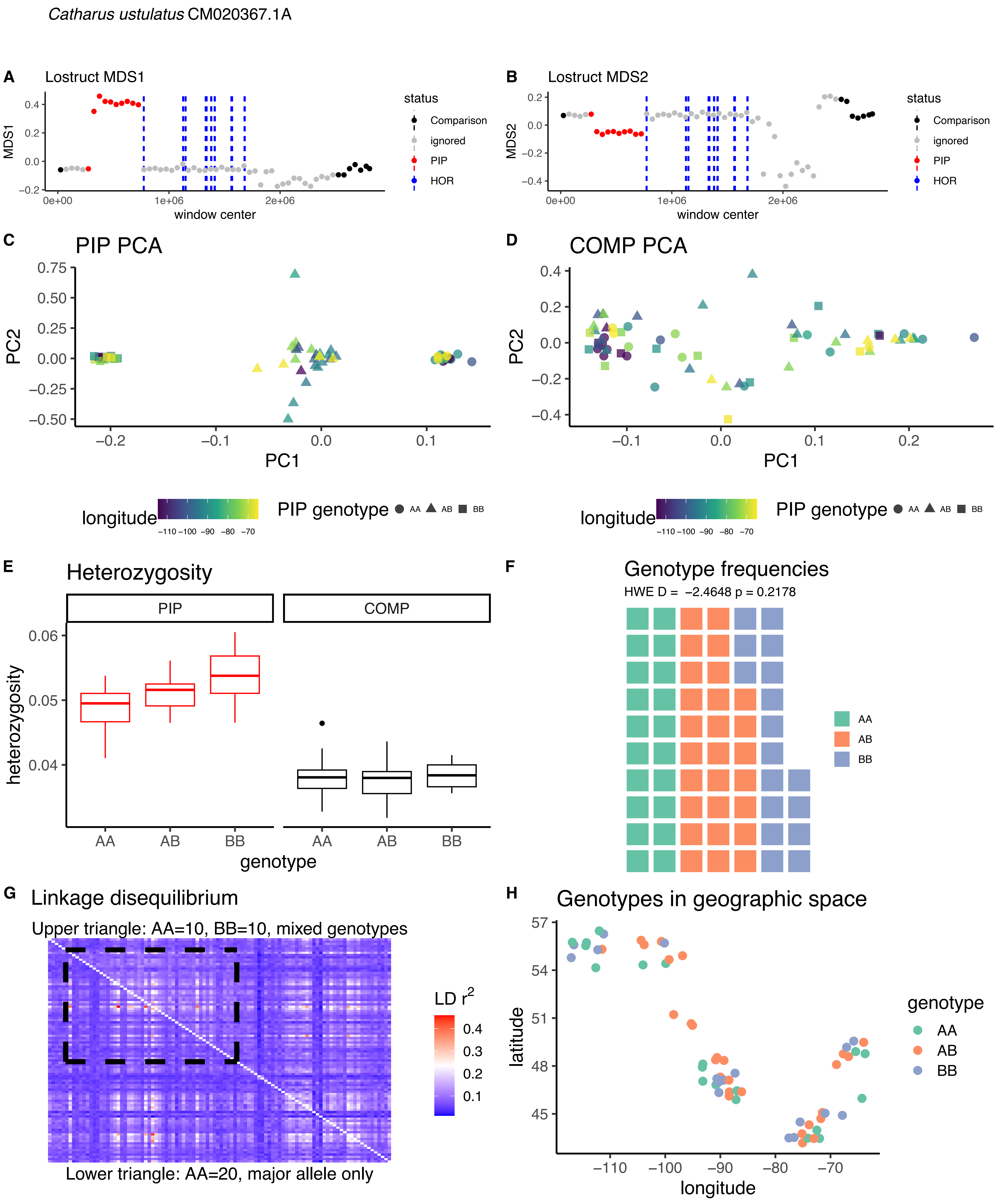
Catharus ustulatus CM020367.1A
A
Lostruct MDS1
MDS1
window center
status
Comparison
ignored
PIP
HOR
B
Lostruct MDS2
MDS2
window center
C
PIP PCA
PC2
PC1
D
COMP PCA
longitude
PIP genotype
AA
AB
BB
E
Heterozygosity
PIP
COMP
heterozygosity
genotype
F
Genotype frequencies
HWE D = −2.4648 p = 0.2178
G
Linkage disequilibrium
Upper triangle: AA=10, BB=10, mixed genotypes
Lower triangle: AA=20, major allele only
LD r2
H
Genotypes in geographic space
latitude
longitude
genotype

*Catharus ustulatus* CM020367.1B

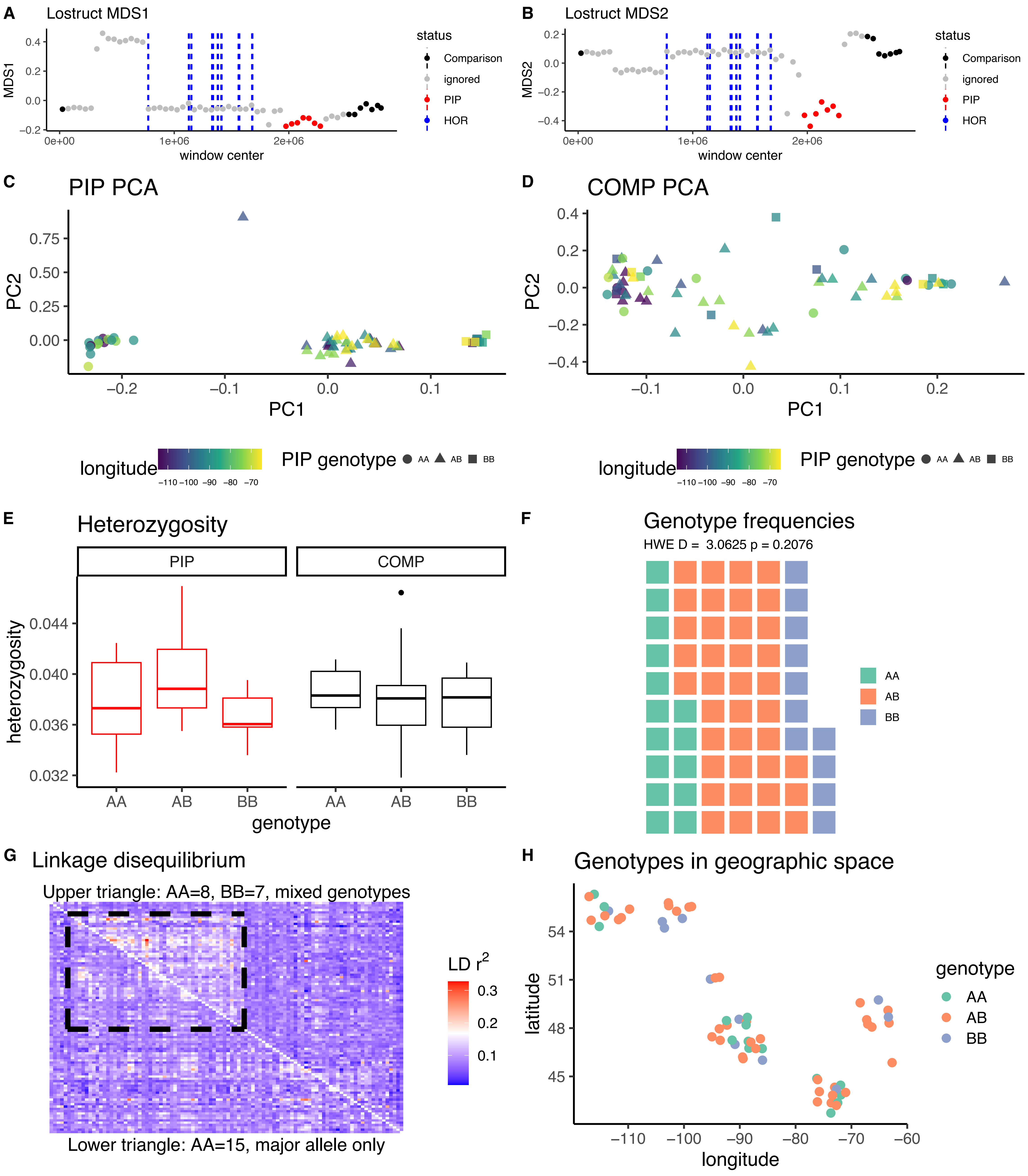

*Catharus ustulatus* CM020365.1

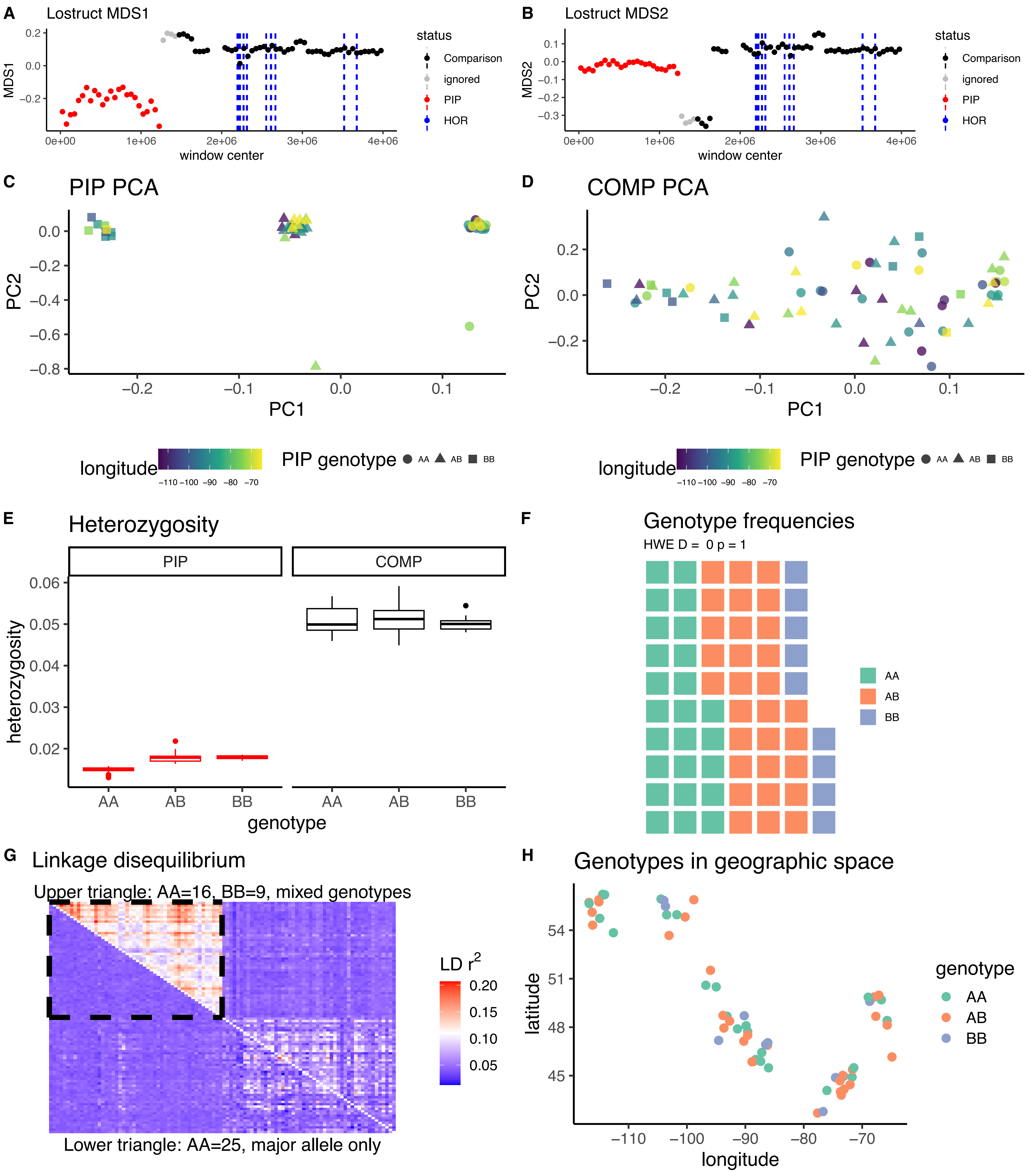

*Catharus ustulatus* CM020344.1

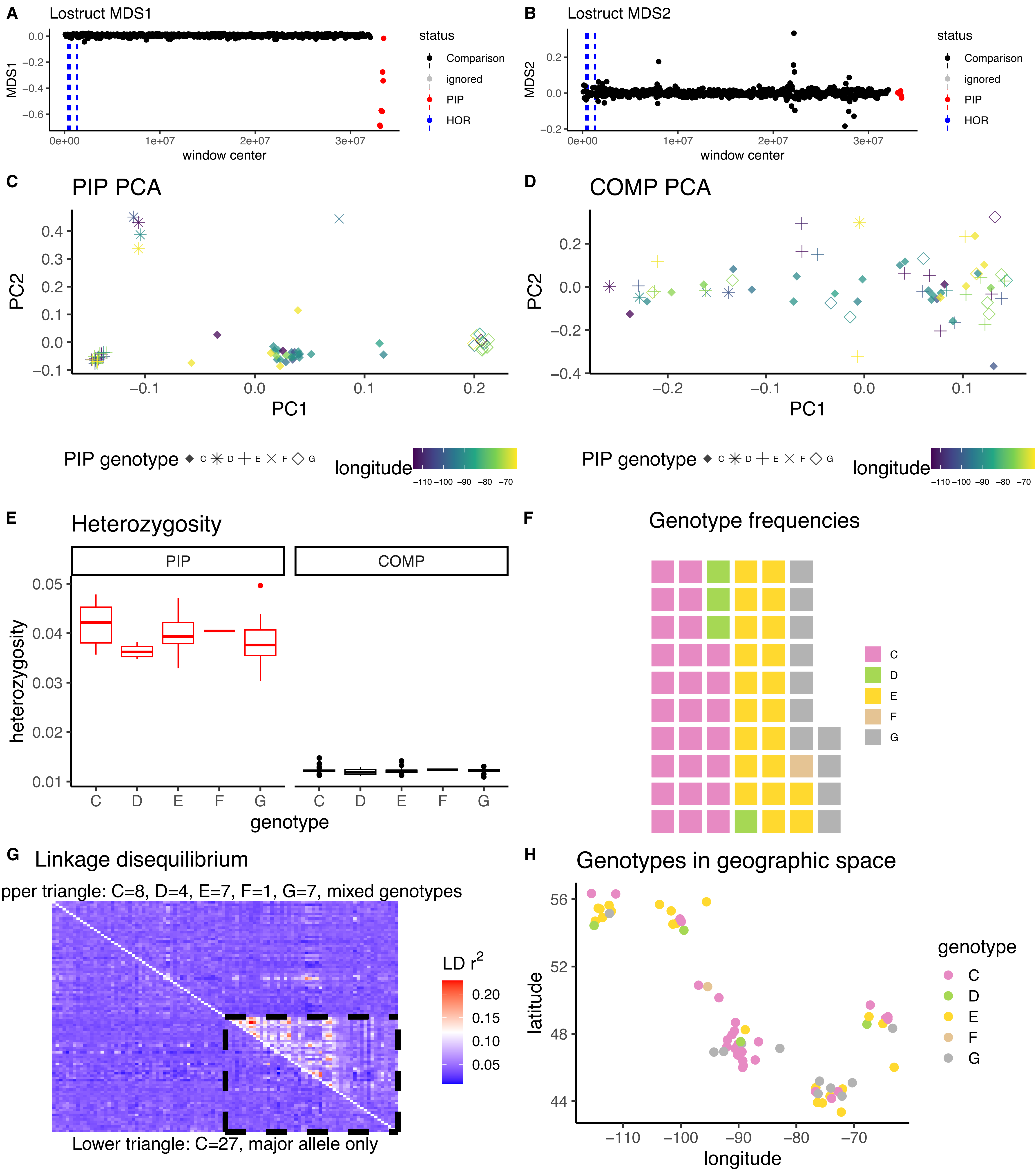

*Junco hyemalis* CM042598.1

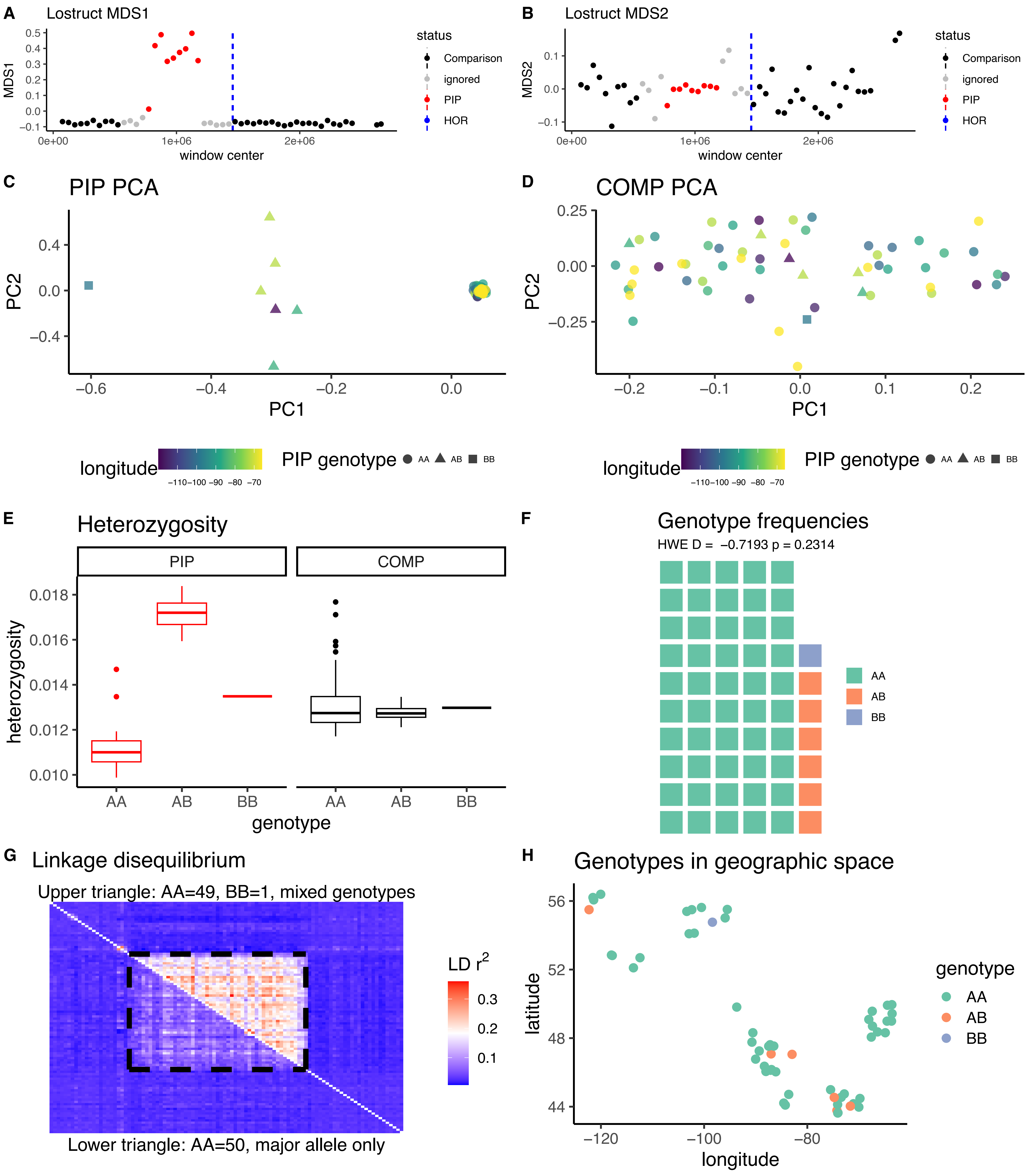

*Junco hyemalis* CM042579.1

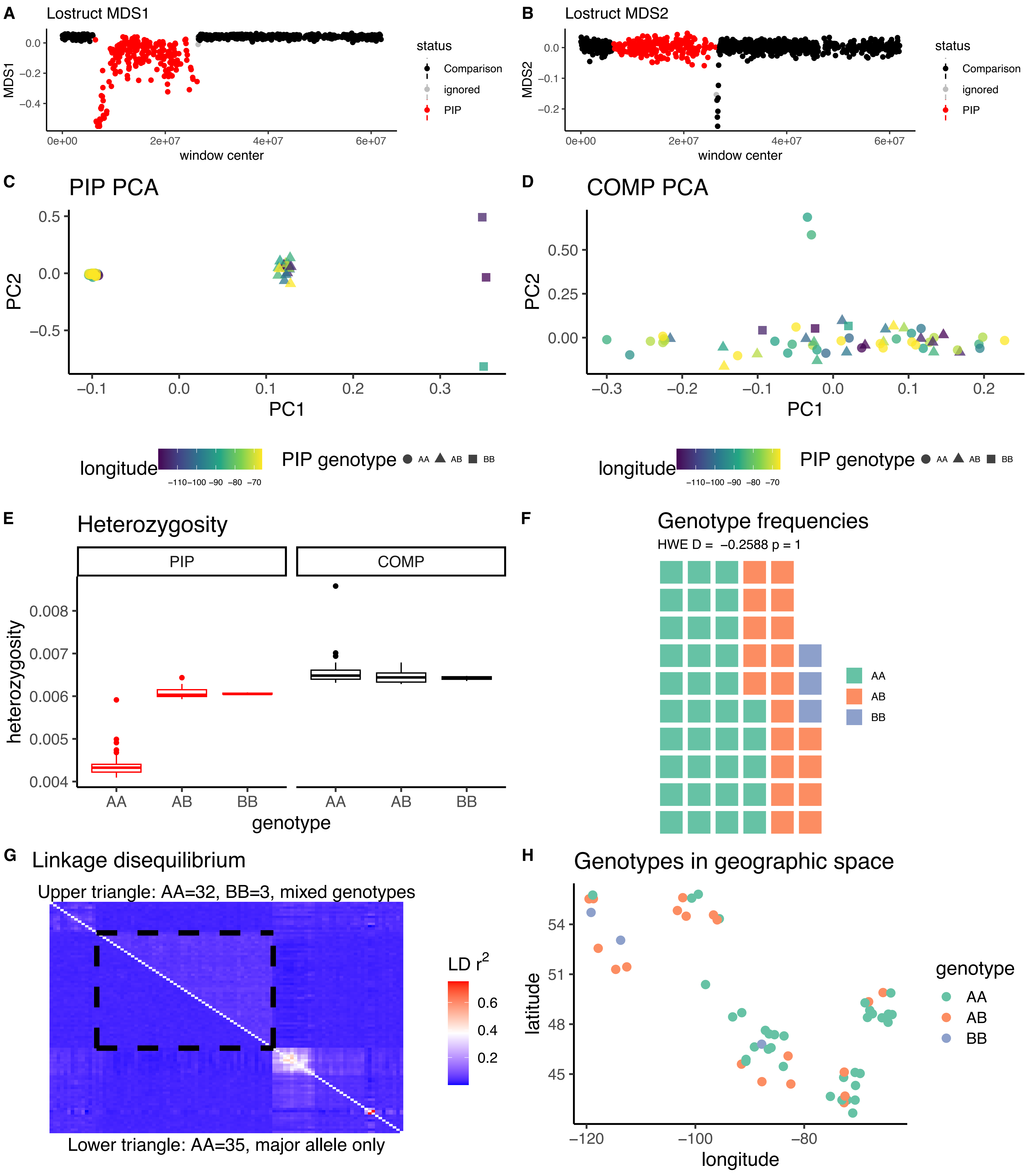

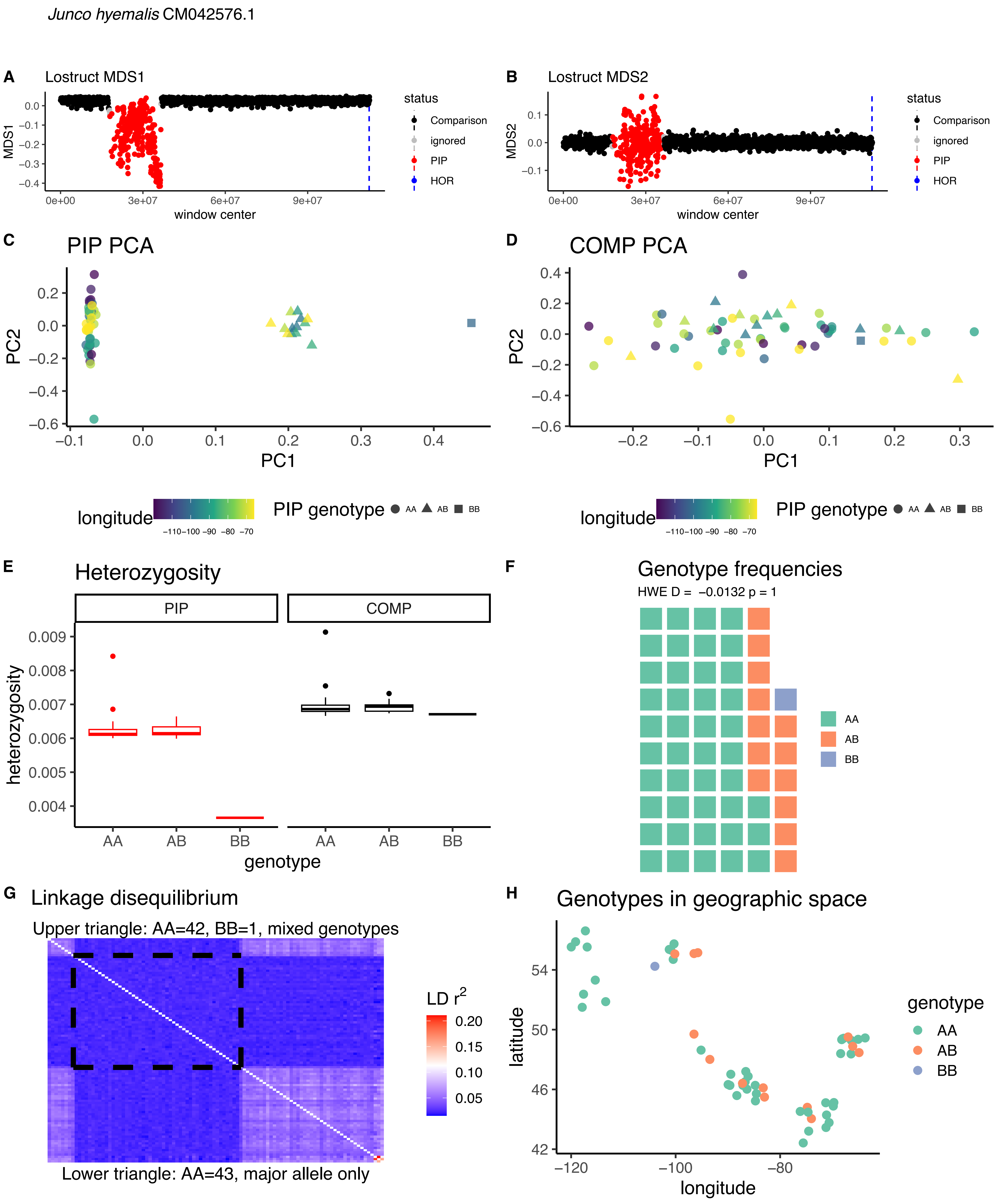
Junco hyemalis CM042576.1
A
Lostruct MDS1
MDS1
window center
status
Comparison
ignored
PIP
HOR
B
Lostruct MDS2
MDS2
C
PIP PCA
PC1
PC2
D
COMP PCA
longitude
PIP genotype
AA
AB
BB
E
Heterozygosity
PIP
COMP
heterozygosity
genotype
F
Genotype frequencies
HWE D = −0.0132 p = 1
G
Linkage disequilibrium
Upper triangle: AA=42, BB=1, mixed genotypes
Lower triangle: AA=43, major allele only
LD r²
H
Genotypes in geographic space
latitude
longitude
genotype

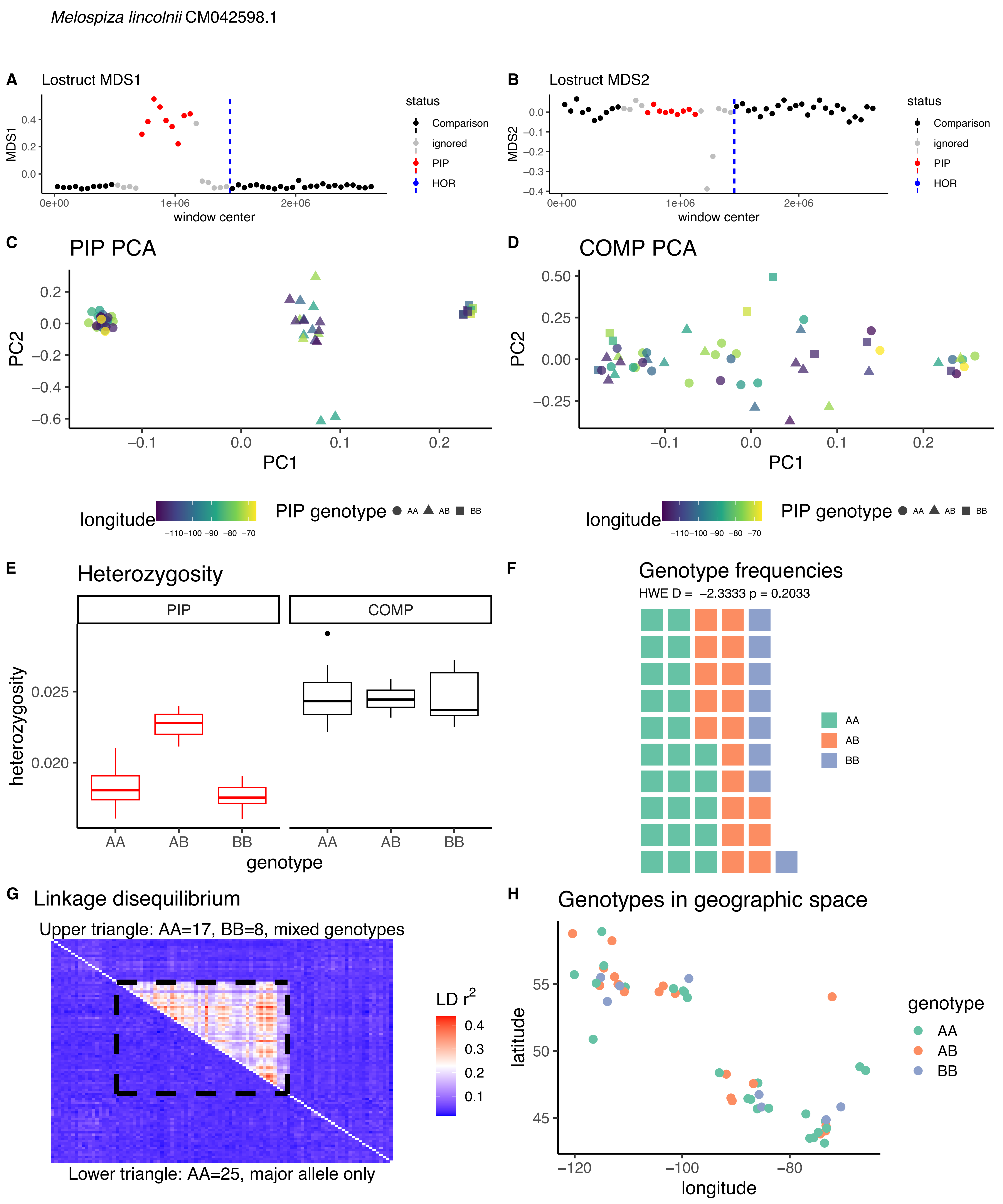

Melospiza lincolnii CM042598.1
A
Lostruct MDS1
MDS1
window center
status
Comparison
ignored
PIP
HOR
B
Lostruct MDS2
MDS2
window center
C
PIP PCA
PC1
PC2
D
COMP PCA
PC1
PC2
longitude
PIP genotype
AA
AB
BB
E
Heterozygosity
PIP
COMP
heterozygosity
genotype
F
Genotype frequencies
HWE D = -2.3333 p = 0.2033
AA
AB
BB
G
Linkage disequilibrium
Upper triangle: AA=17, BB=8, mixed genotypes
LD $r^2$
Lower triangle: AA=25, major allele only
H
Genotypes in geographic space
latitude
longitude
genotype
AA
AB
BB

*Melospiza lincolnii* CM042581.1A

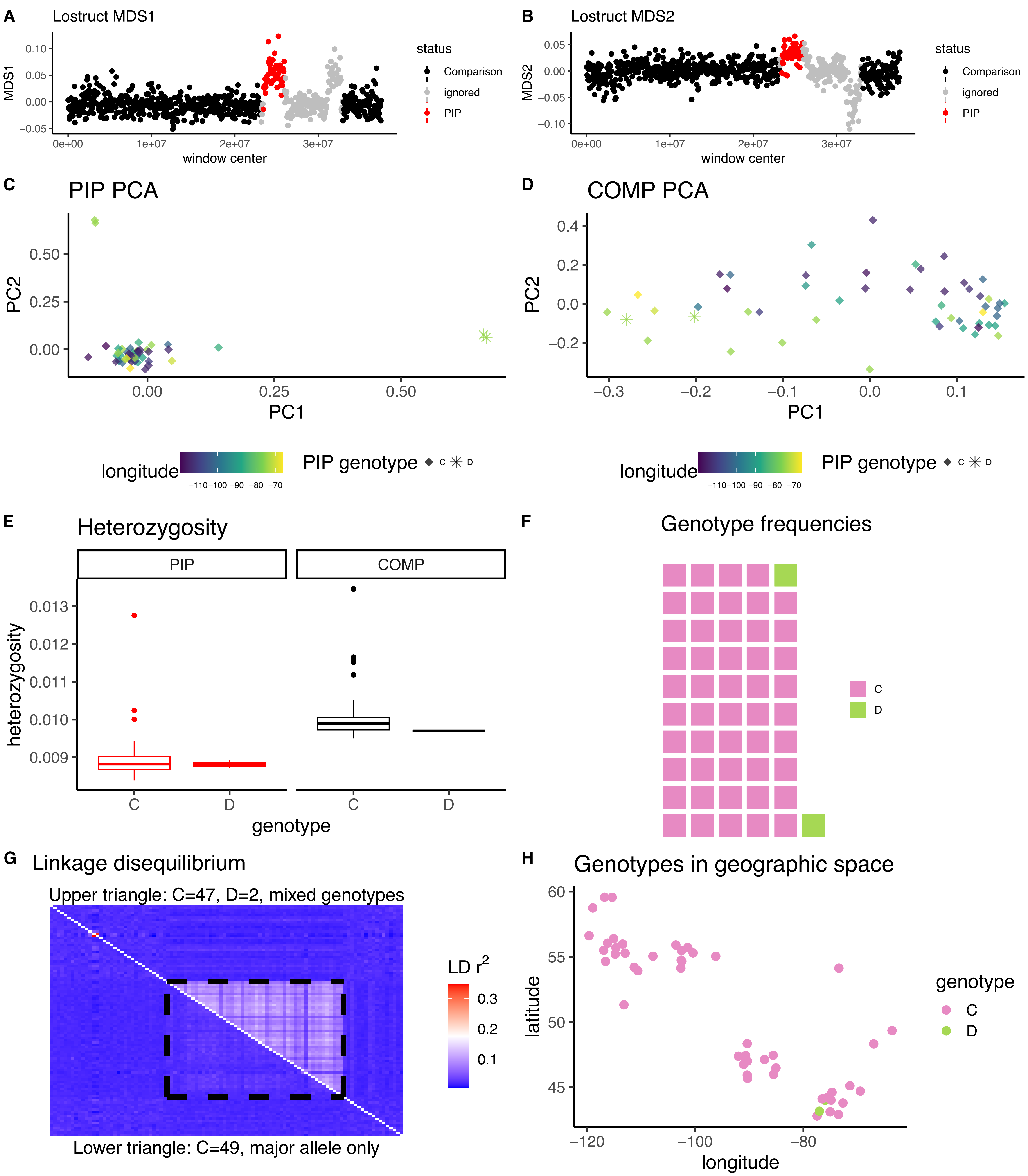

*Melospiza lincolnii* CM042581.1B

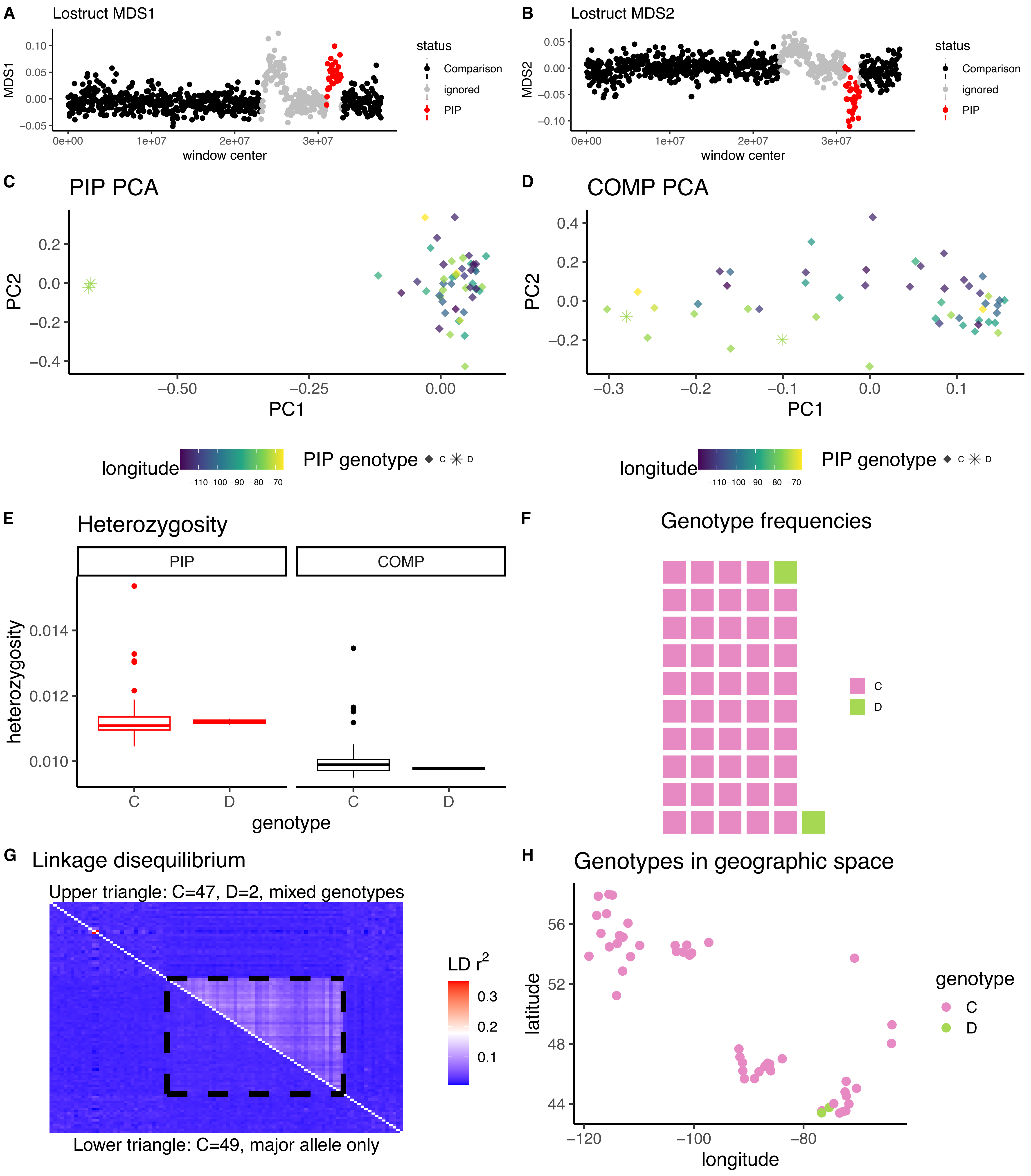

*Melospiza lincolnii* CM042579.1A

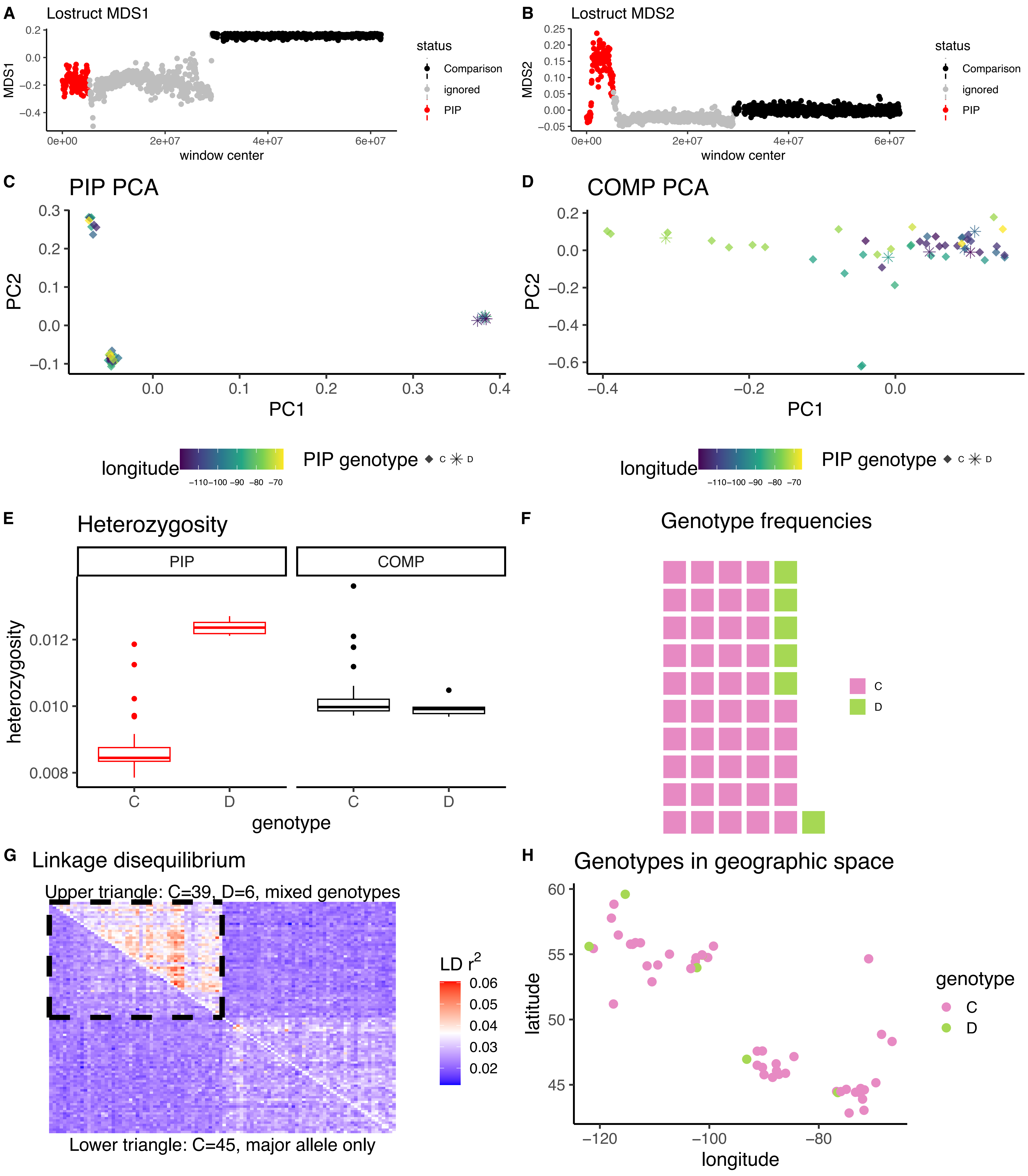

*Melospiza lincolnii* CM042579.1B

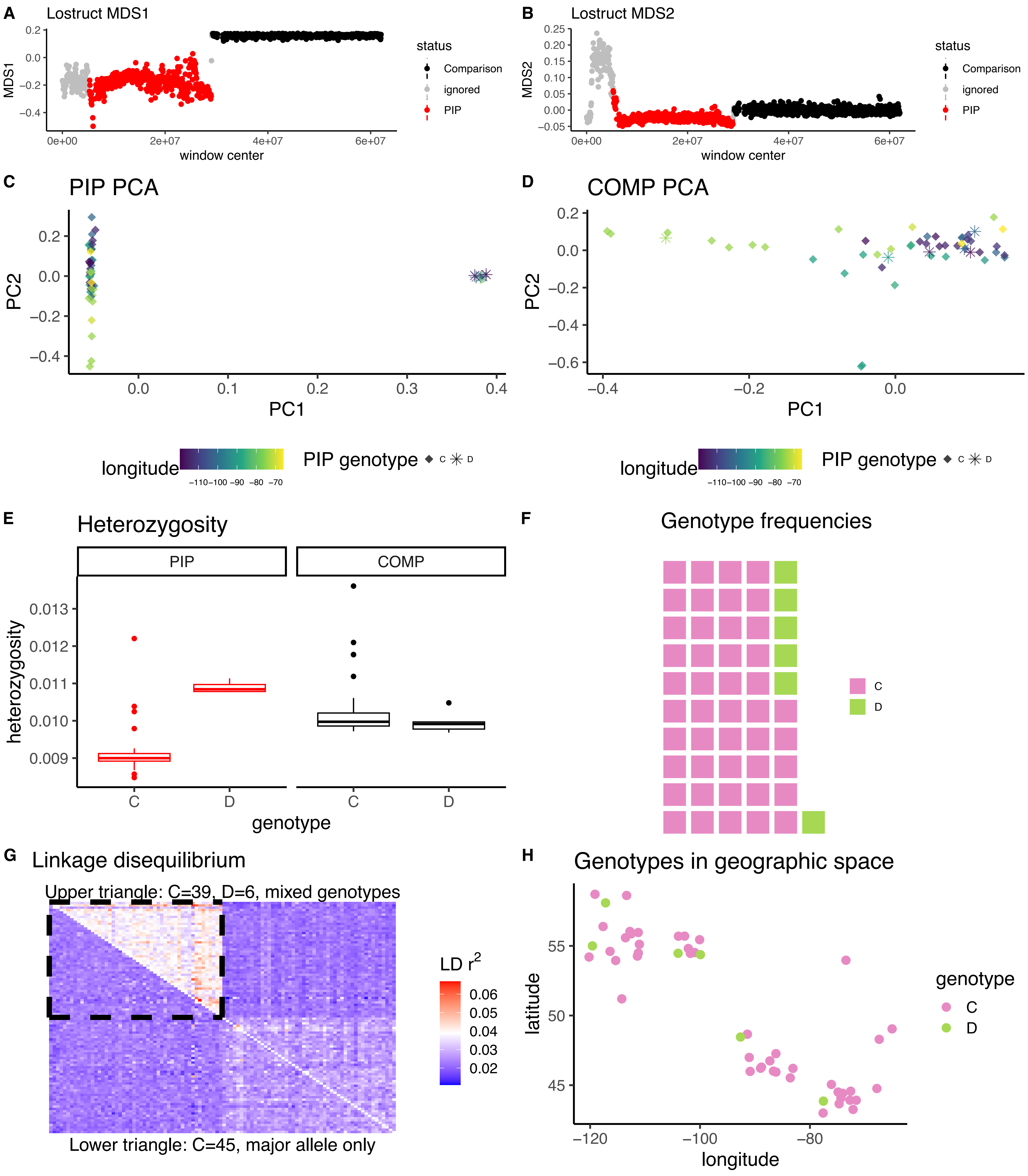

*Melospiza lincolnii* CM042577.1

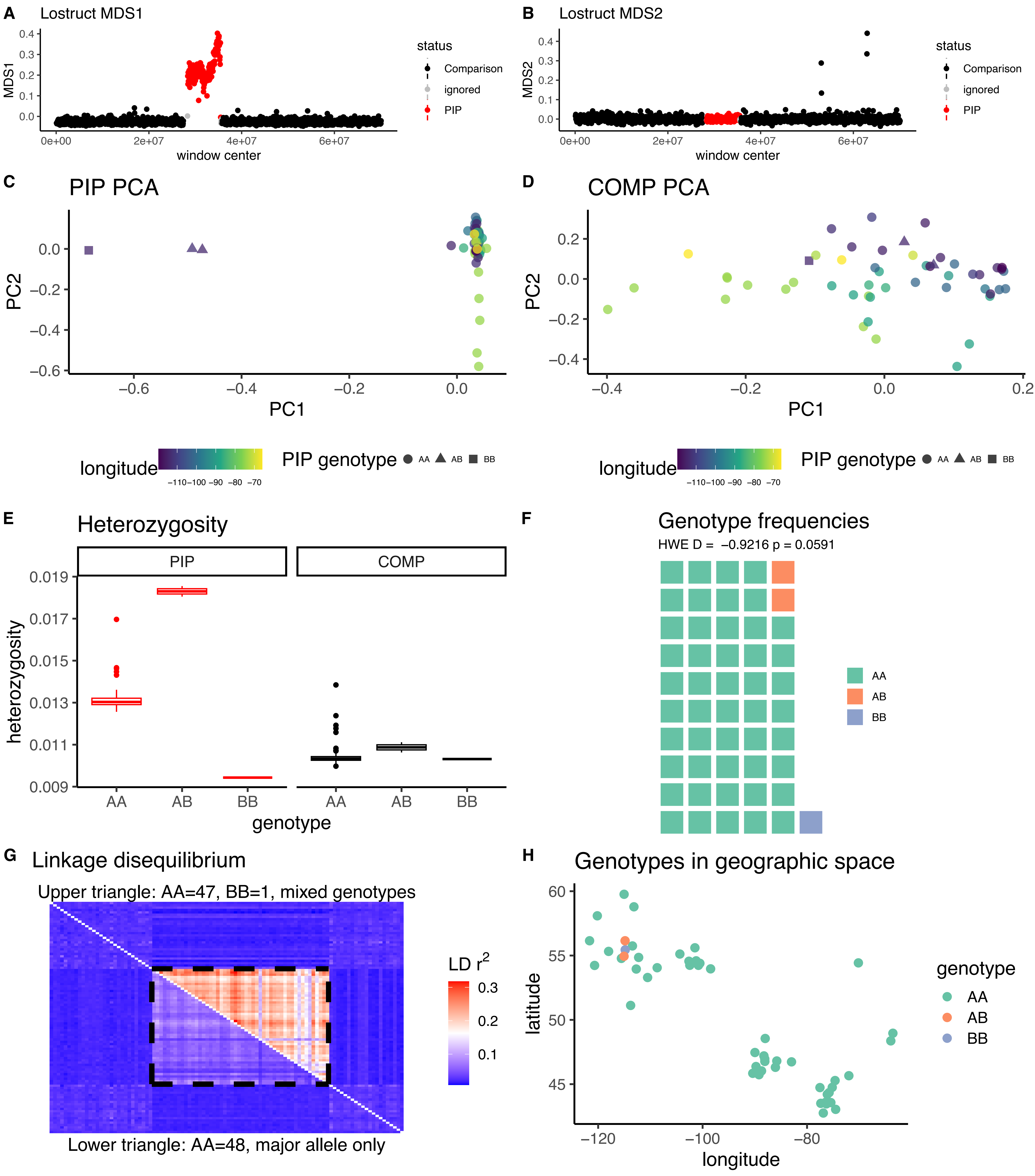

*Zonotrichia albicollis* CM042600.1

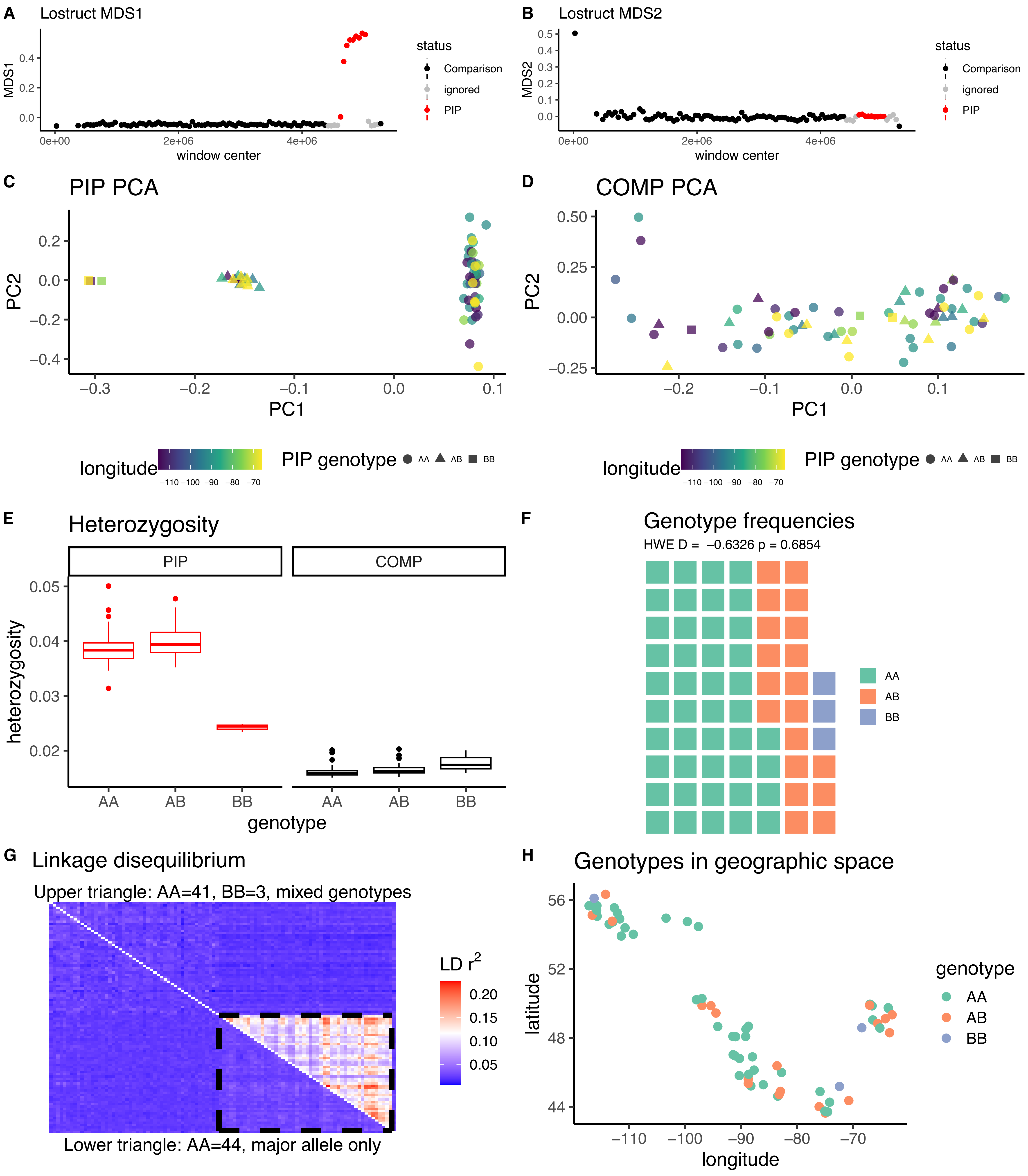

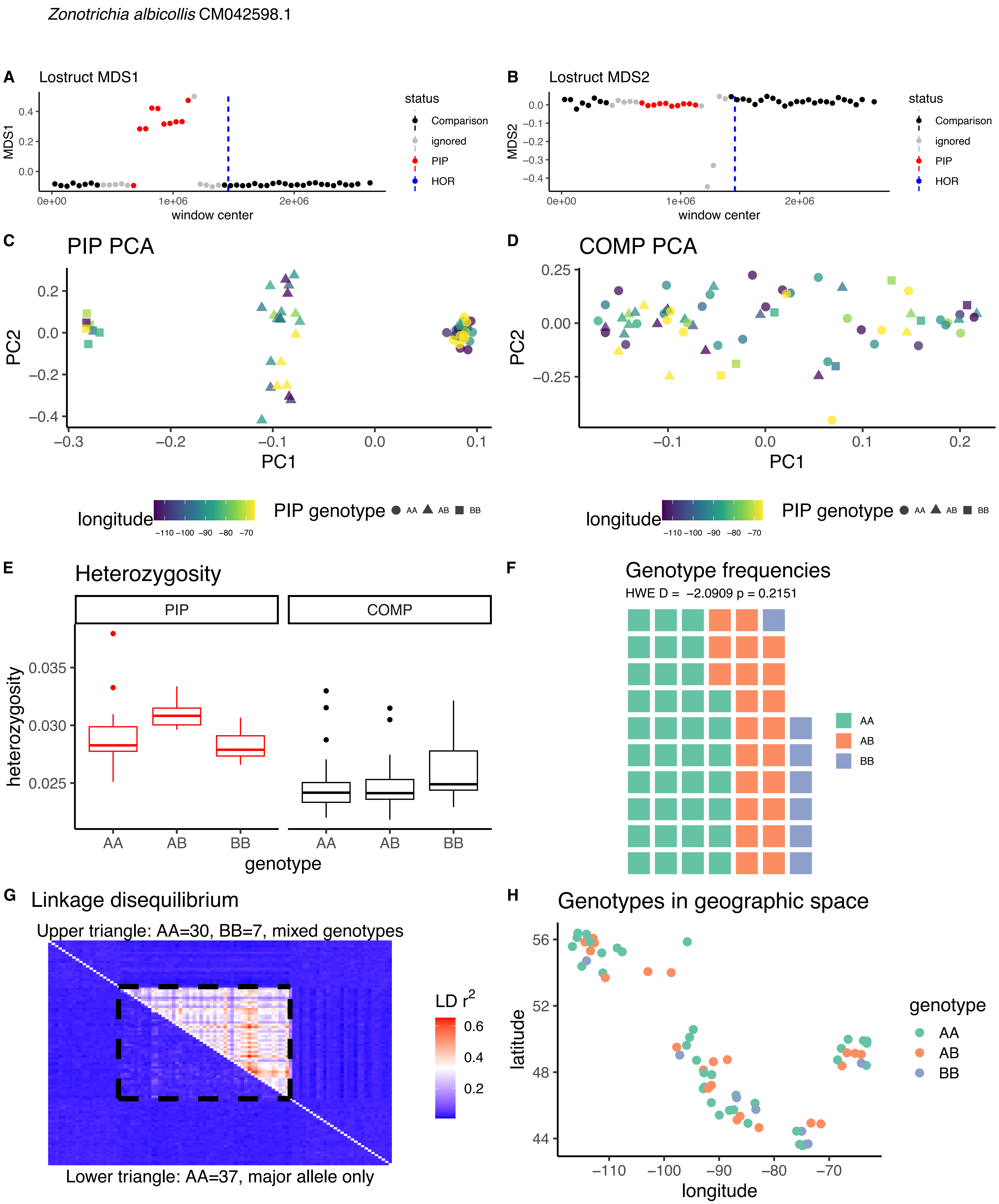

Zonotrichia albicollis CM042598.1
A
Lostruct MDS1
MDS1
window center
status
Comparison
ignored
PIP
HOR
B
Lostruct MDS2
MDS2
C
PIP PCA
PC1
PC2
D
COMP PCA
longitude
PIP genotype
AA
AB
BB
E
Heterozygosity
PIP
COMP
heterozygosity
genotype
F
Genotype frequencies
HWE D = −2.0909 p = 0.2151
G
Linkage disequilibrium
Upper triangle: AA=30, BB=7, mixed genotypes
LD r2
Lower triangle: AA=37, major allele only
H
Genotypes in geographic space
latitude
longitude
genotype

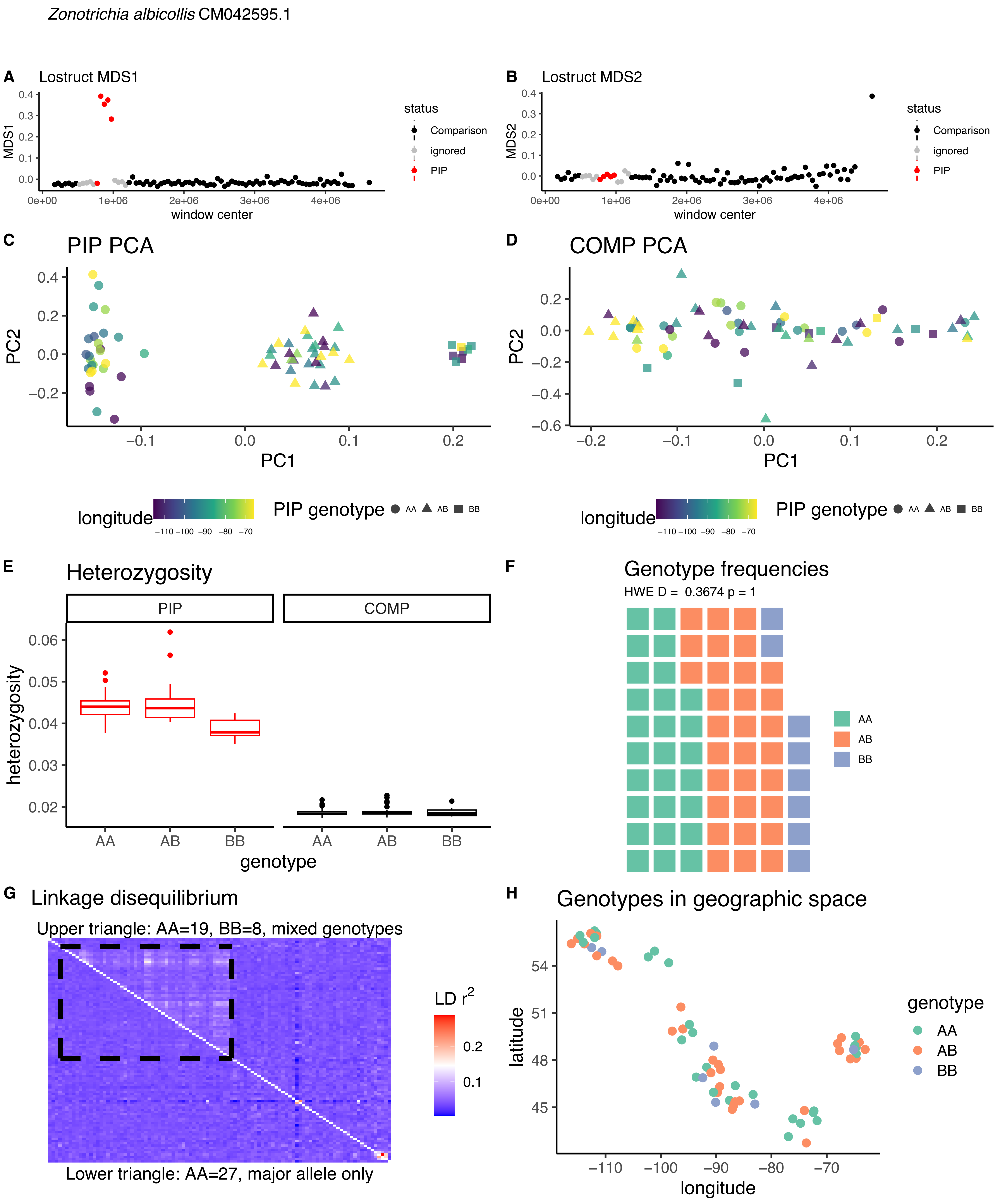

Zonotrichia albicollis CM042595.1
A Lostruct MDS1
B Lostruct MDS2
status
Comparison
ignored
PIP
window center
C PIP PCA
D COMP PCA
longitude
PIP genotype AA AB BB
E Heterozygosity
PIP
COMP
heterozygosity
genotype
F Genotype frequencies
HWE D = 0.3674 p = 1
G Linkage disequilibrium
Upper triangle: AA=19, BB=8, mixed genotypes
Lower triangle: AA=27, major allele only
LD r2
H Genotypes in geographic space
latitude
longitude
genotype

*Zonotrichia albicollis* CM042576.1

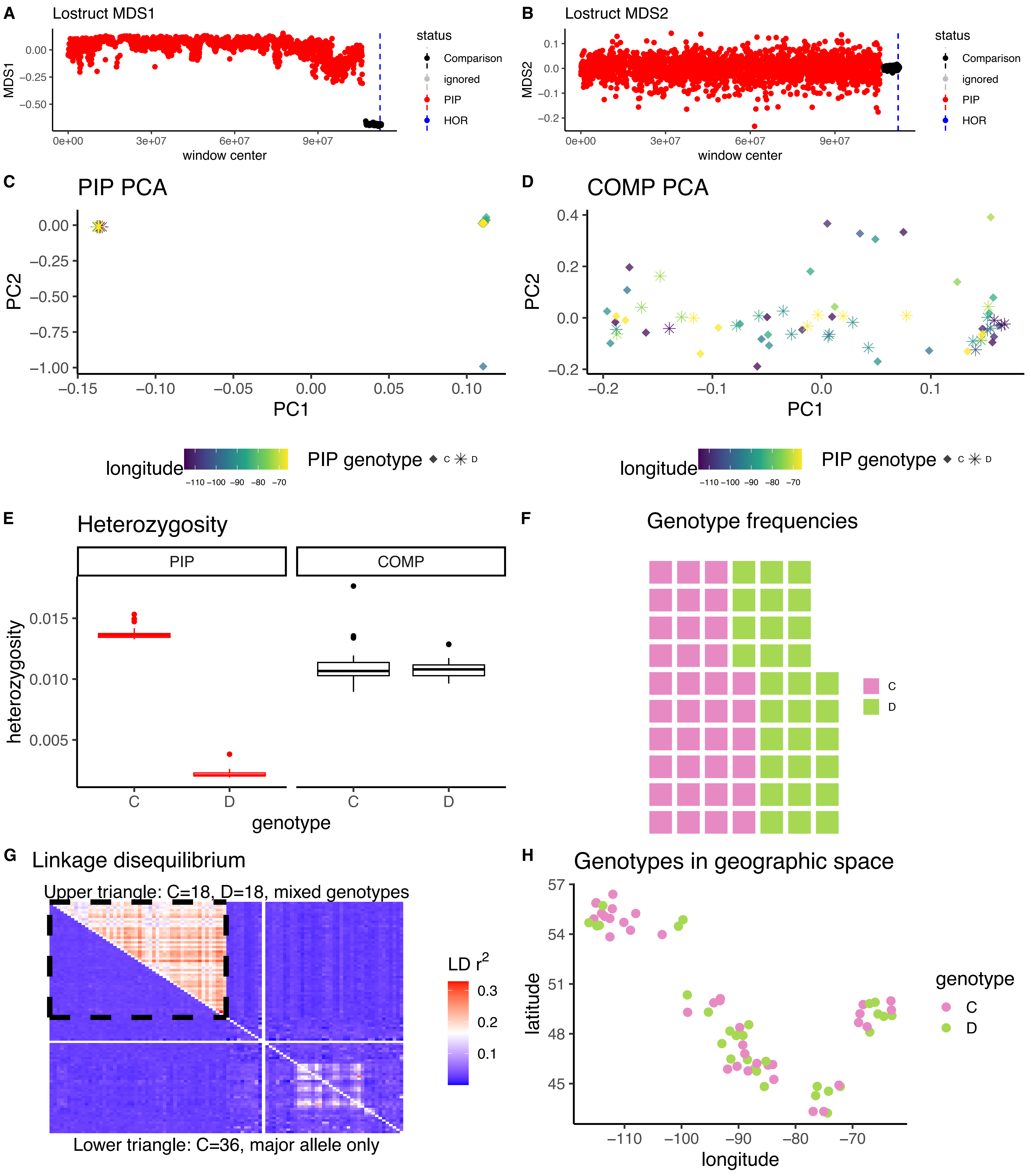

*Zonotrichia albicollis* CM042573.1A

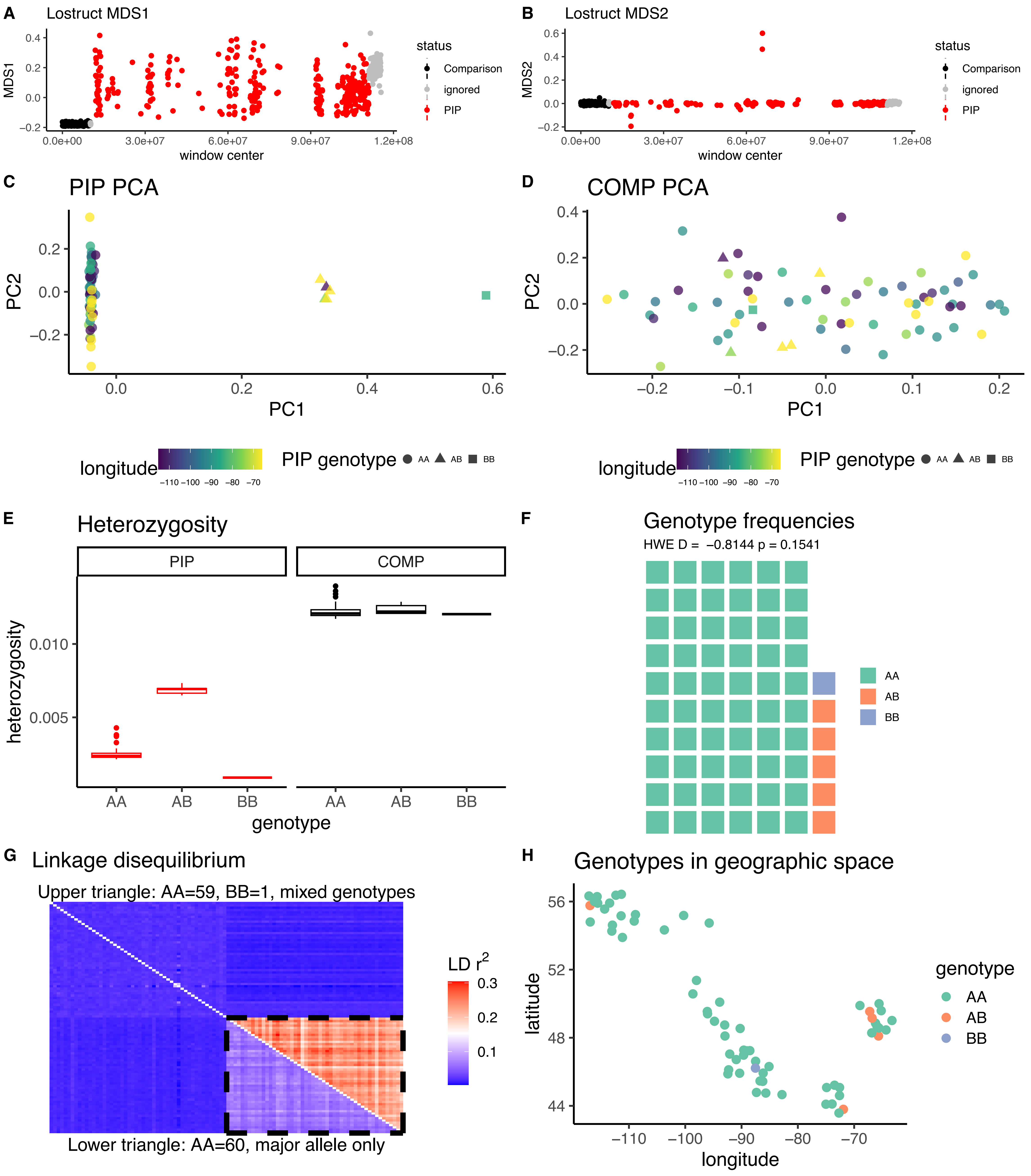

*Zonotrichia albicollis* CM042573.1B

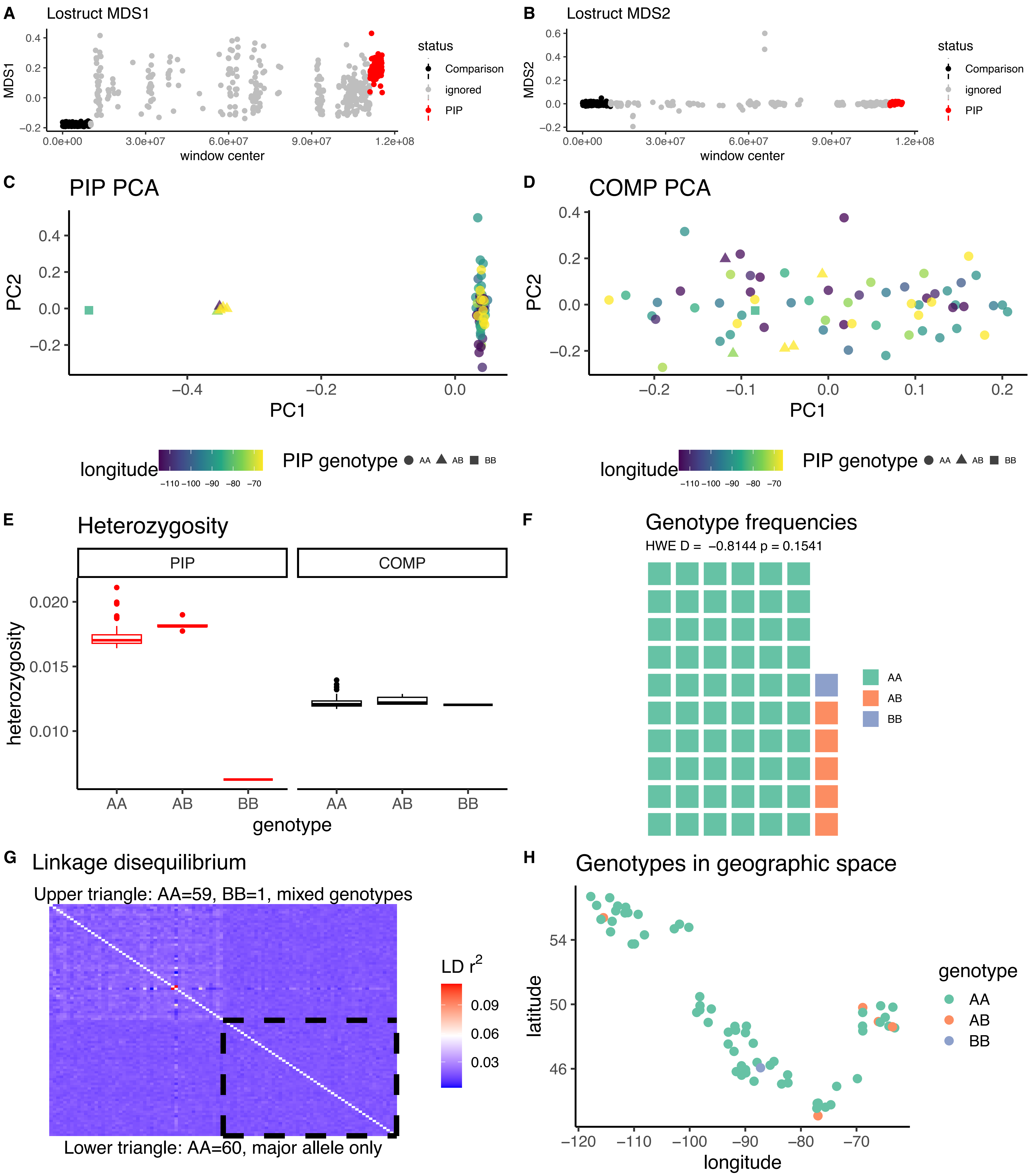

*Cardellina canadensis* CM027534.1

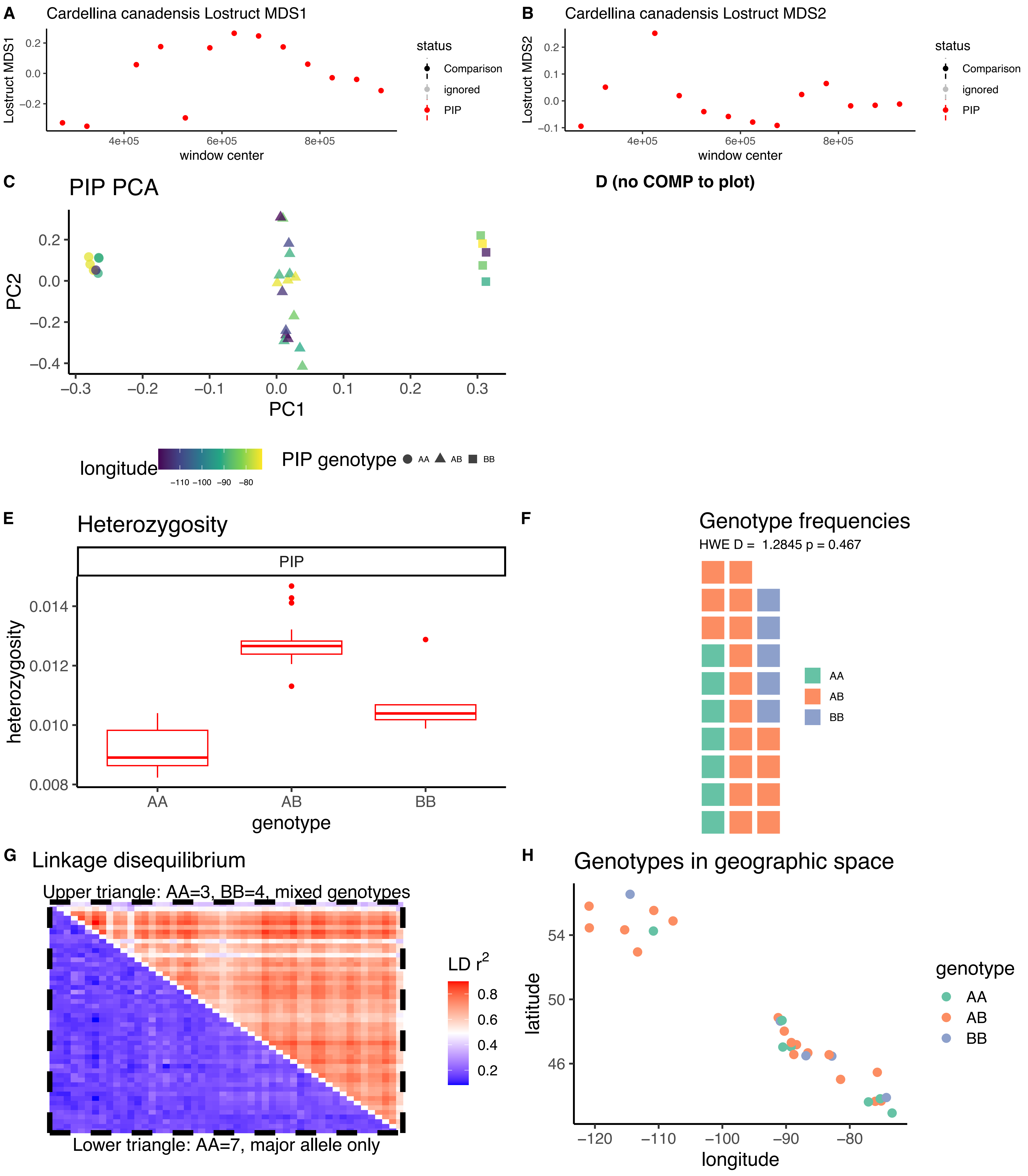

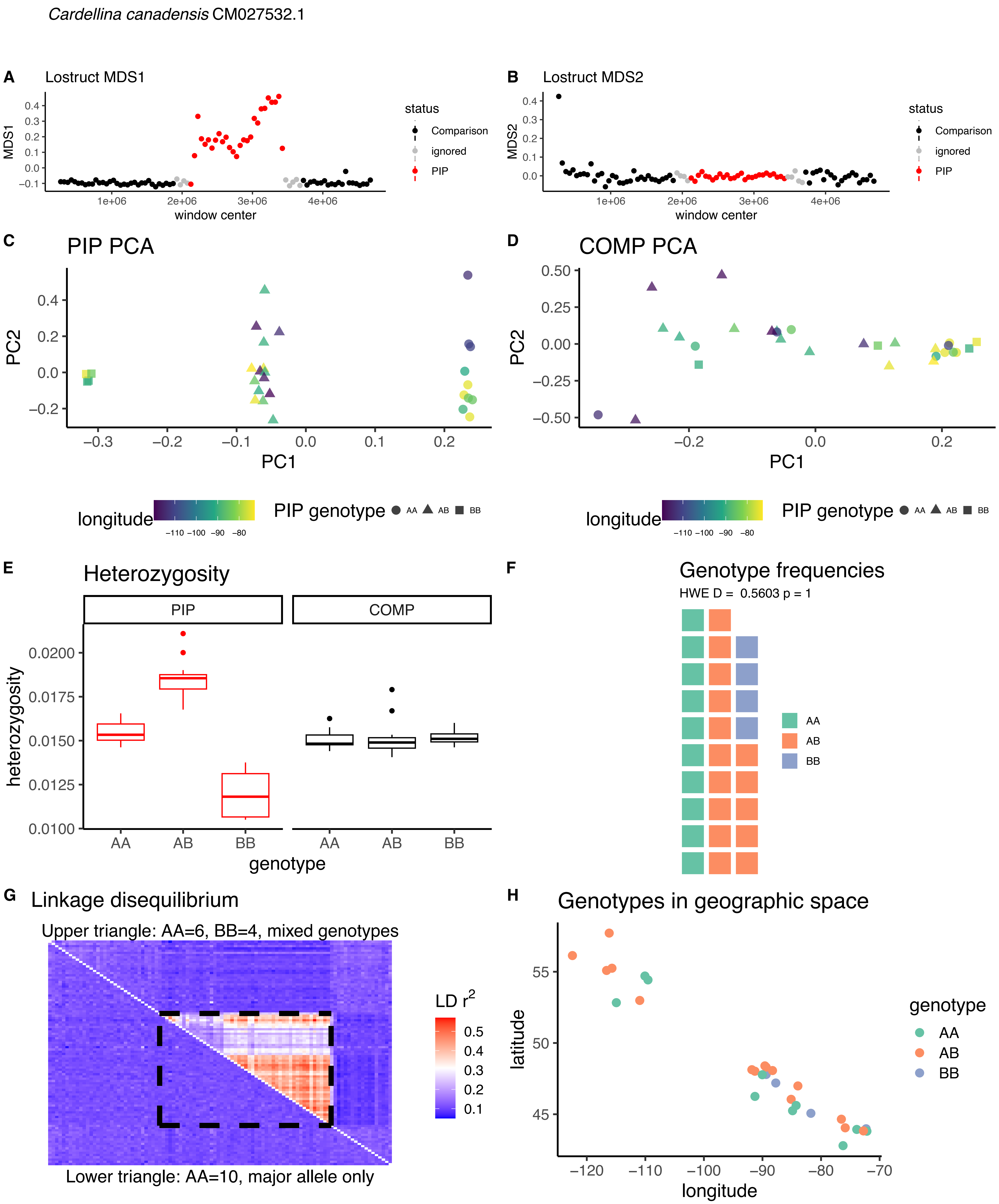
*Cardellina canadensis* CM027532.1
A
Lostruct MDS1
MDS1
window center
status
Comparison
ignored
PIP
B
Lostruct MDS2
MDS2
C
PIP PCA
PC1
PC2
D
COMP PCA
longitude
PIP genotype
AA
AB
BB
E
Heterozygosity
PIP
COMP
heterozygosity
genotype
F
Genotype frequencies
HWE D = 0.5603 p = 1
G
Linkage disequilibrium
Upper triangle: AA=6, BB=4, mixed genotypes
Lower triangle: AA=10, major allele only
LD $r^2$
H
Genotypes in geographic space
latitude
longitude
genotype

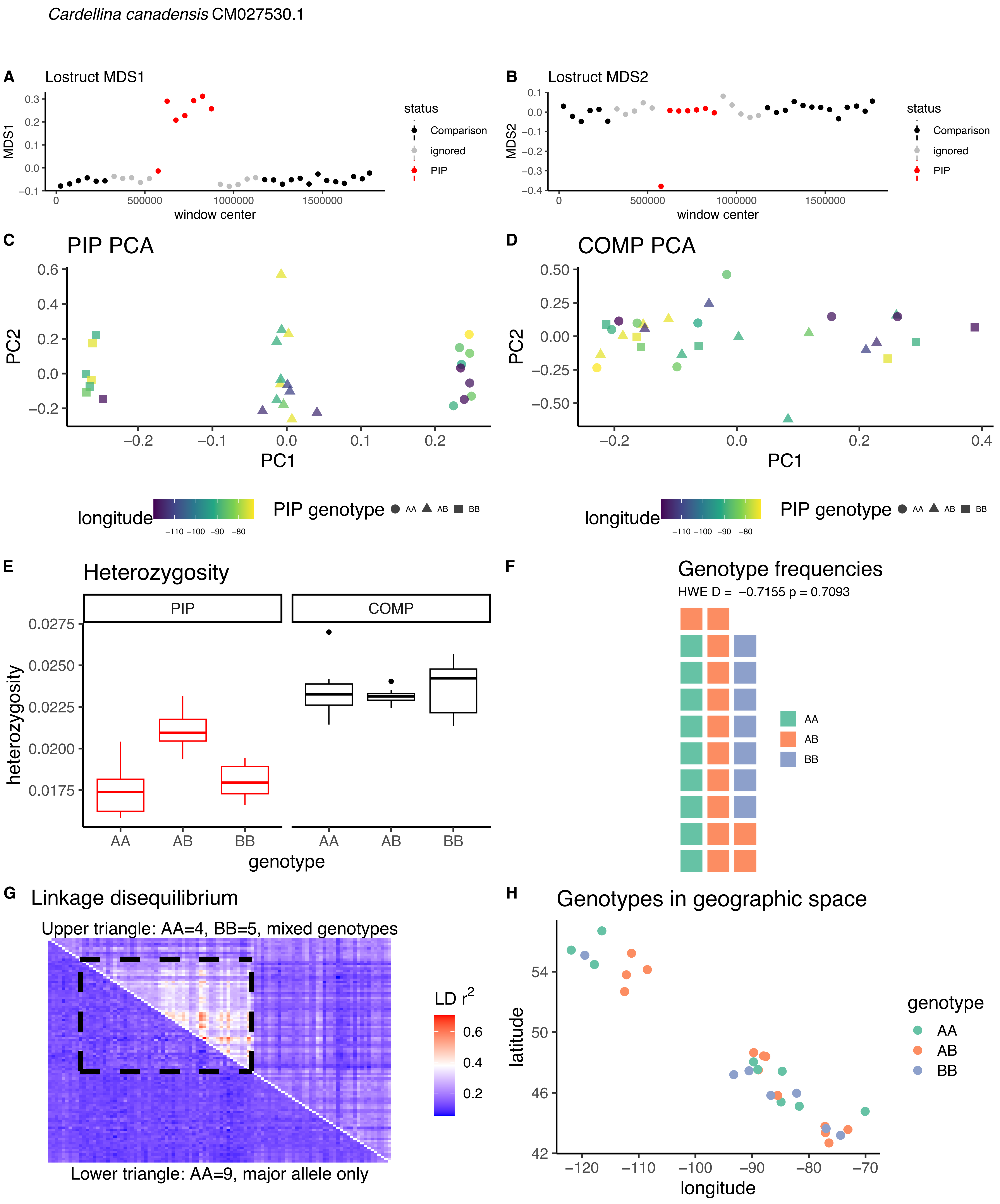
Cardellina canadensis CM027530.1
A
Lostruct MDS1
MDS1
window center
status
Comparison
ignored
PIP
B
Lostruct MDS2
MDS2
window center
status
Comparison
ignored
PIP
C
PIP PCA
PC2
PC1
D
COMP PCA
PC2
PC1
longitude
PIP genotype AA AB BB
longitude
PIP genotype AA AB BB
E
Heterozygosity
PIP
COMP
heterozygosity
genotype
F
Genotype frequencies
HWE D = −0.7155 p = 0.7093
AA
AB
BB
G
Linkage disequilibrium
Upper triangle: AA=4, BB=5, mixed genotypes
LD r2
Lower triangle: AA=9, major allele only
H
Genotypes in geographic space
latitude
longitude
genotype
AA
AB
BB

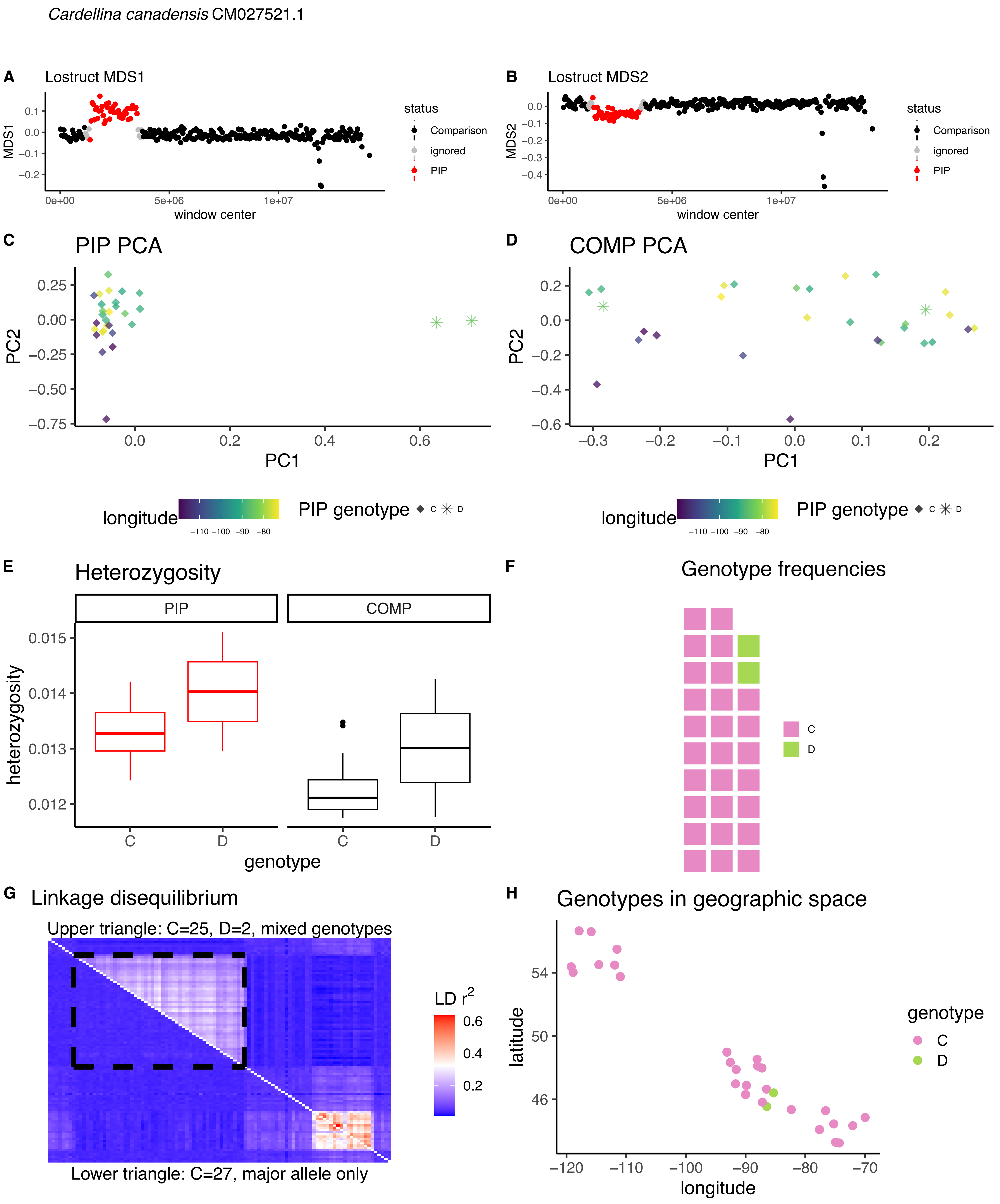

*Cardellina canadensis* CM027521.1
A
Lostruct MDS1
MDS1
window center
status
Comparison
ignored
PIP
B
Lostruct MDS2
MDS2
C
PIP PCA
PC2
PC1
D
COMP PCA
longitude
PIP genotype
C
D
E
Heterozygosity
PIP
COMP
heterozygosity
genotype
F
Genotype frequencies
G
Linkage disequilibrium
Upper triangle: C=25, D=2, mixed genotypes
Lower triangle: C=27, major allele only
LD $r^2$
H
Genotypes in geographic space
latitude

*Cardellina canadensis* CM027511.1A

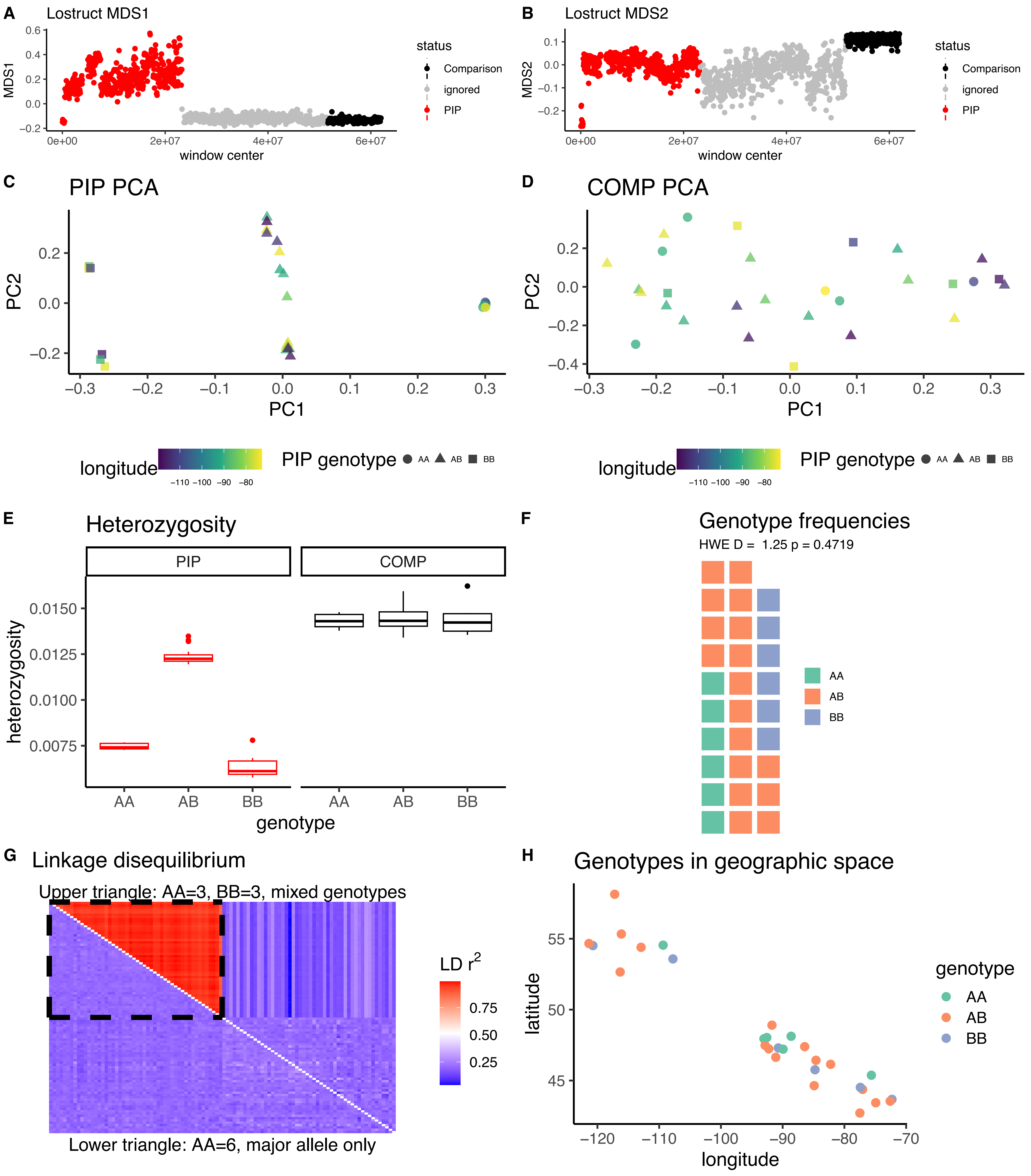

*Cardellina canadensis* CM027511.1B

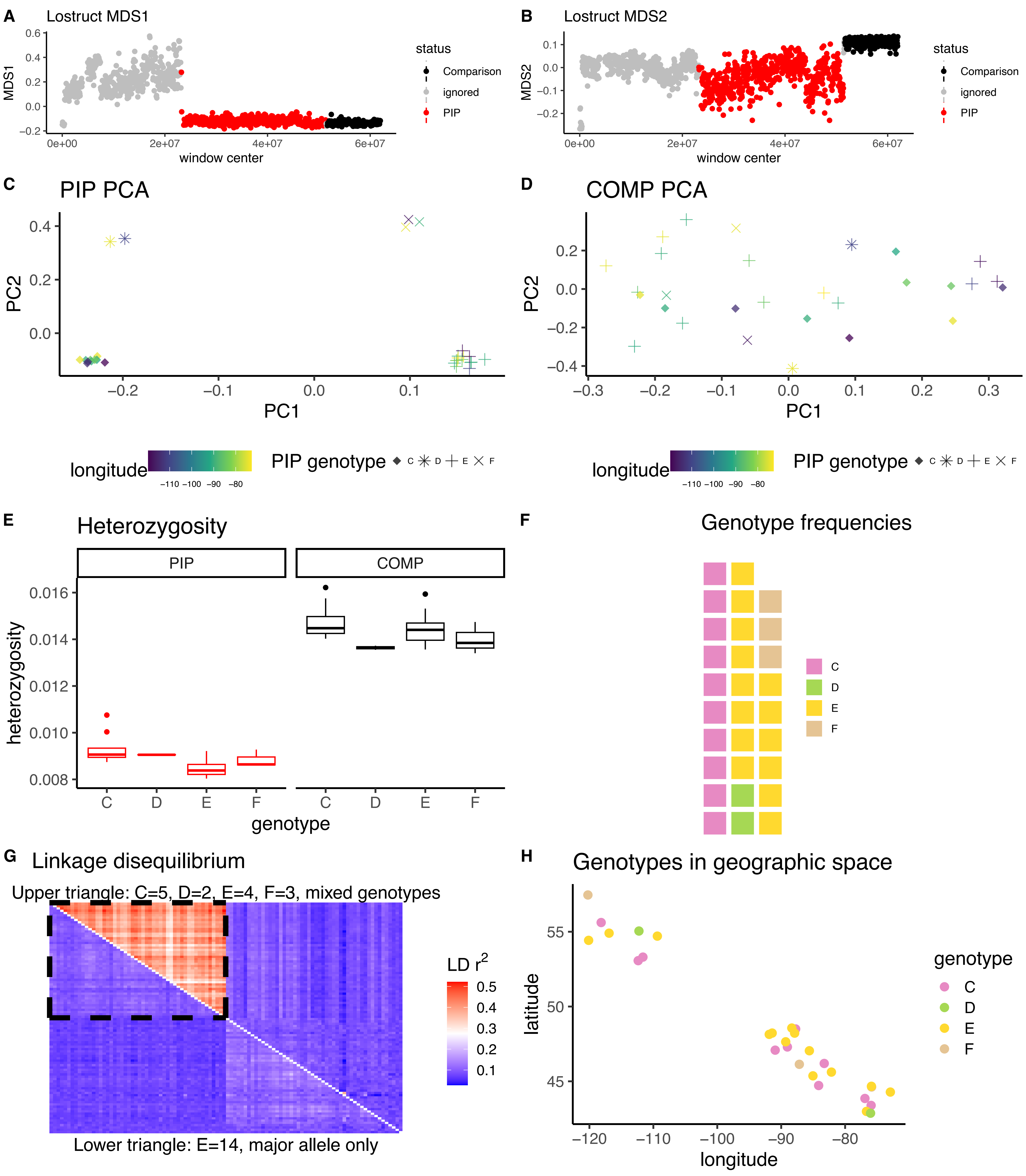

*Cardellina canadensis* CM027510.1

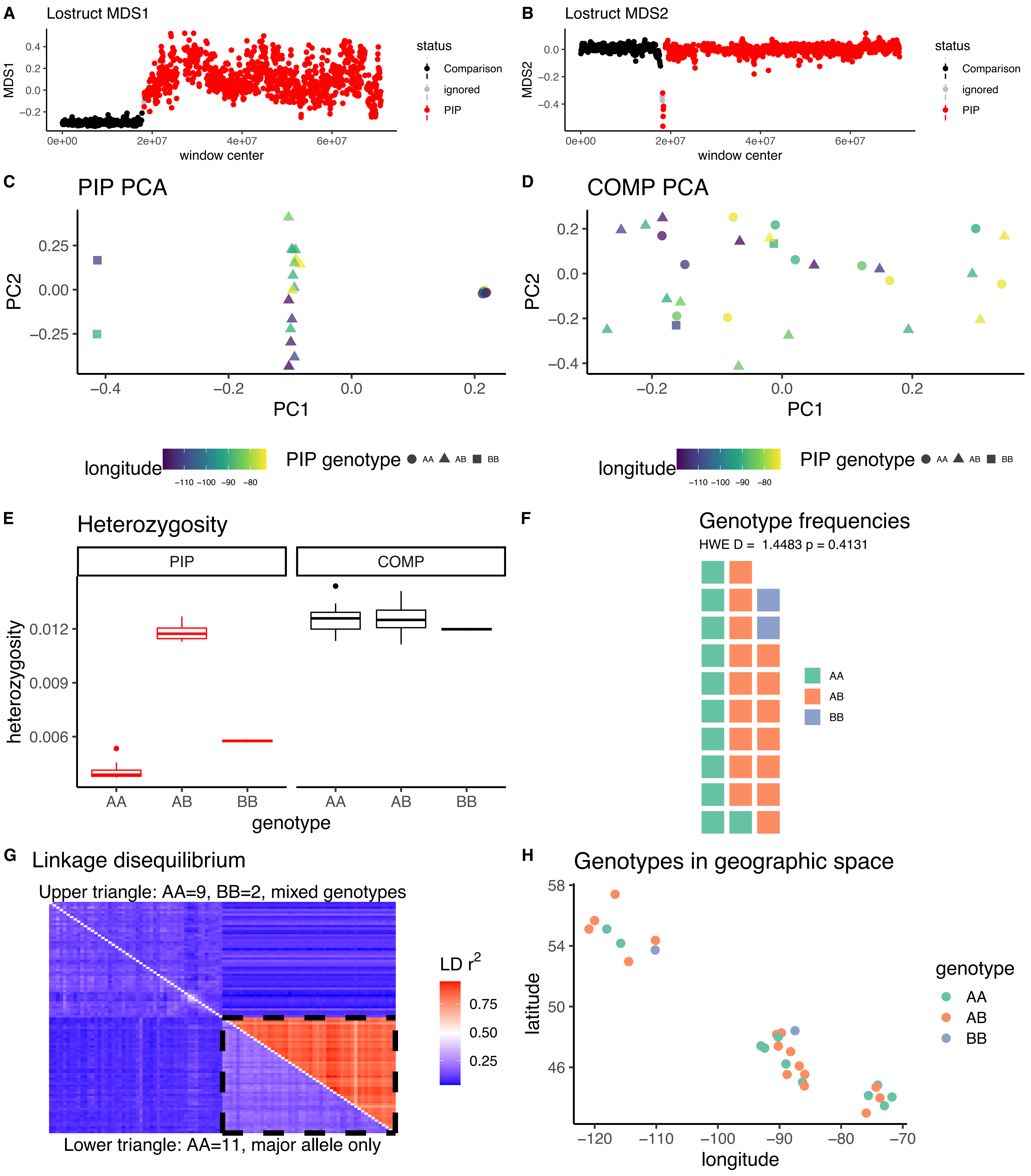

*Geothlypis philadelphia* CM019932.1

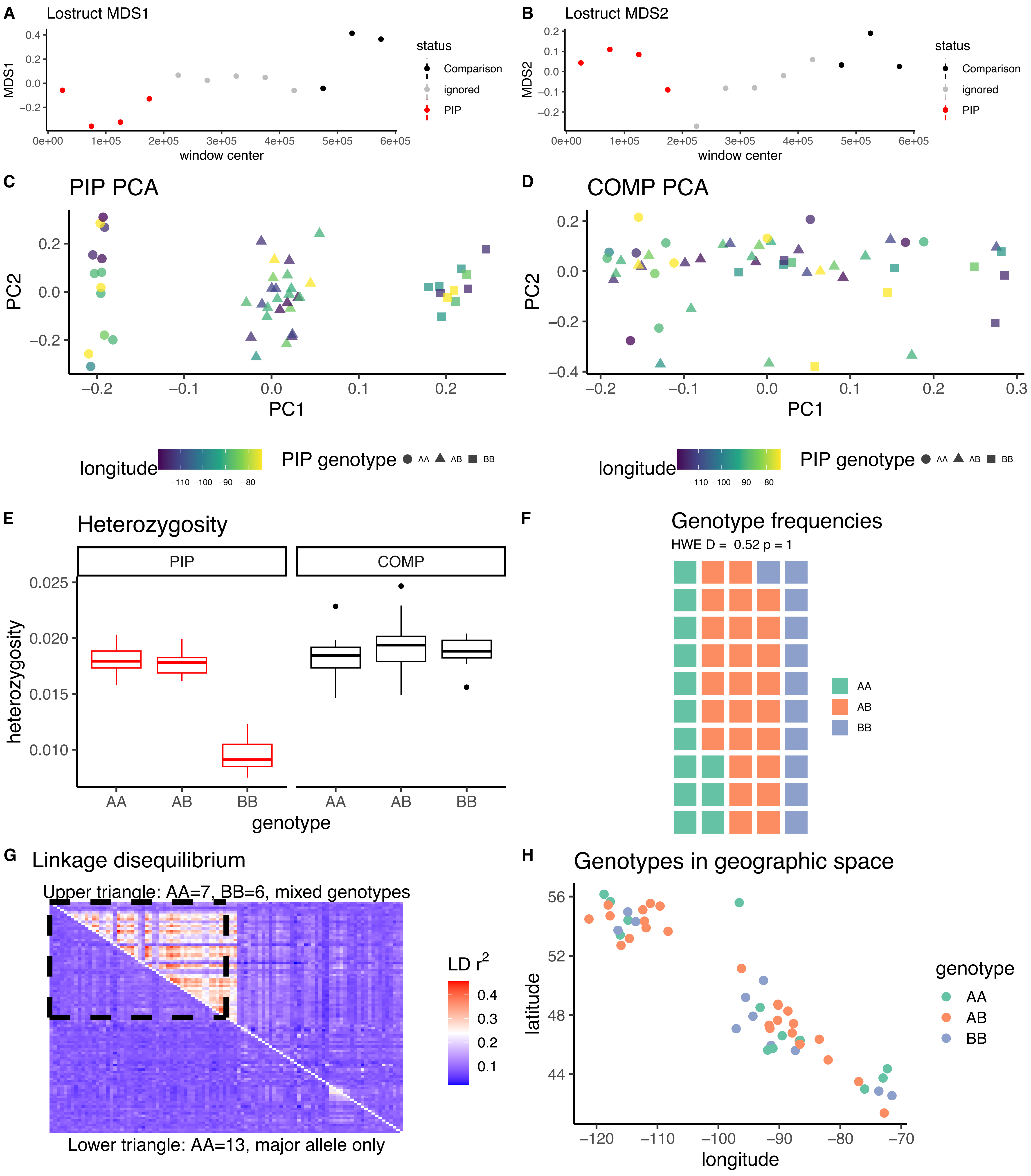

*Geothlypis philadelphia* CM019929.1

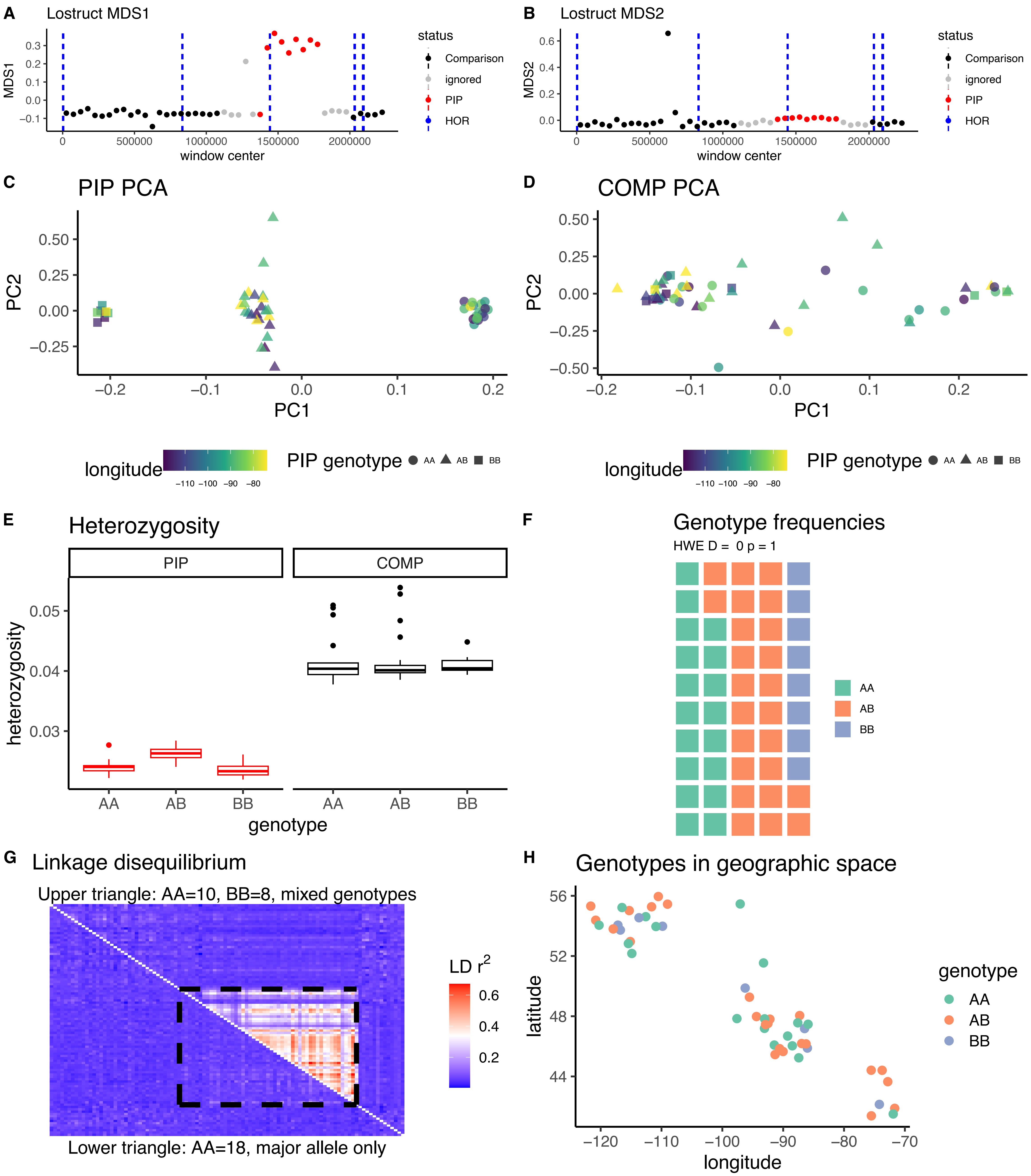

*Geothlypis philadelphia* CM019926.1

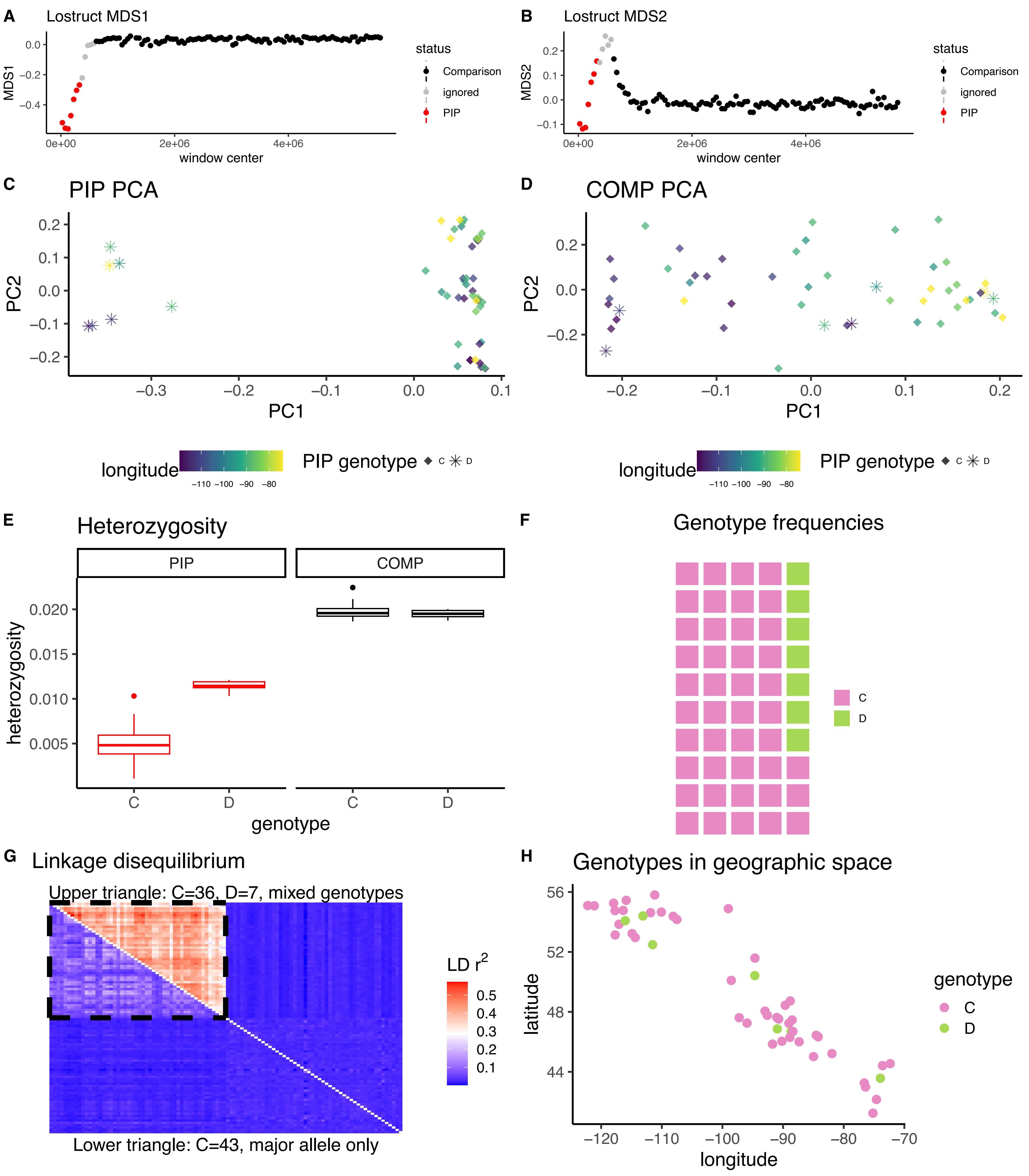

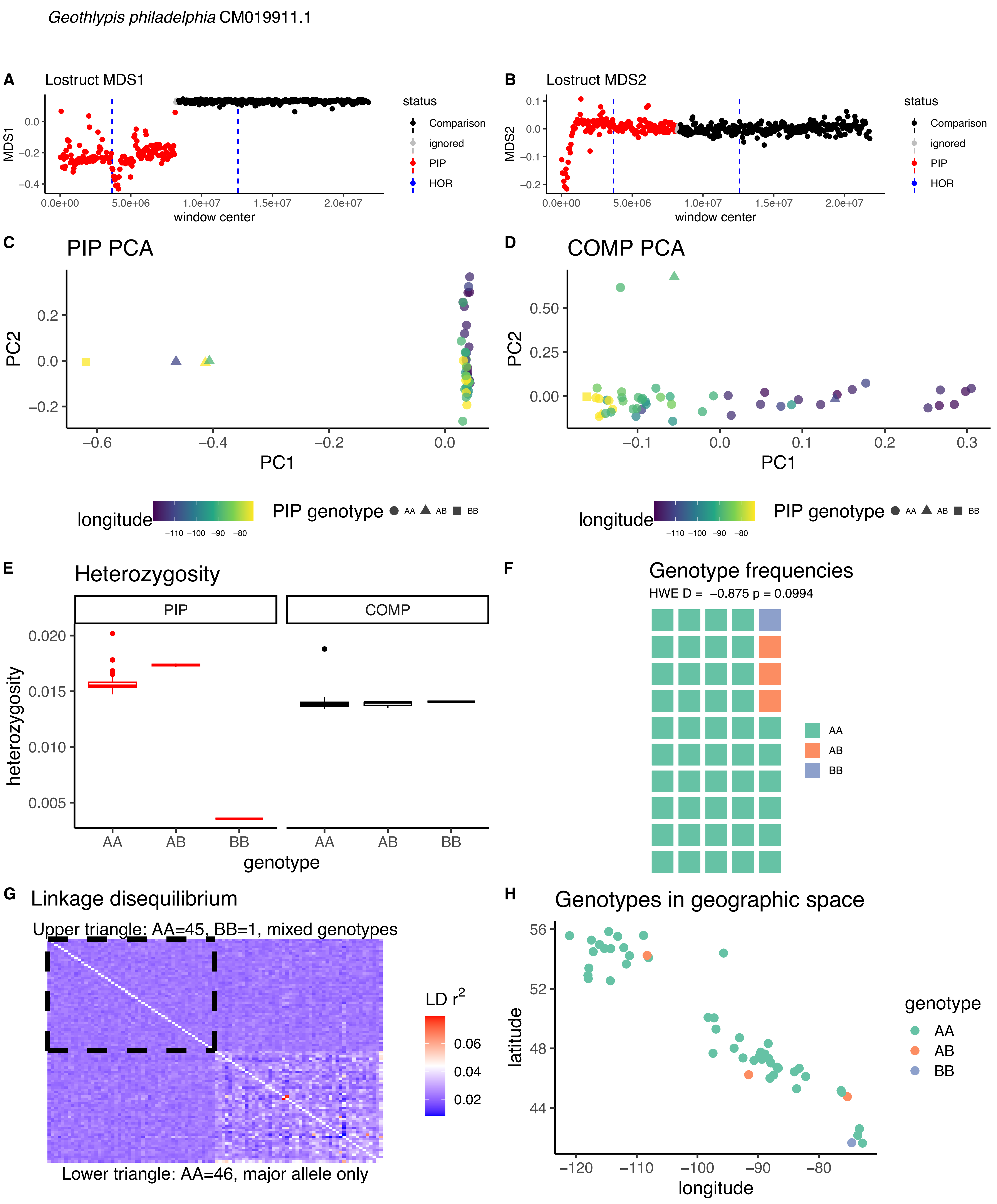

Geothlypis philadelphia CM019911.1
A
Lostruct MDS1
MDS1
window center
status
Comparison
ignored
PIP
HOR
B
Lostruct MDS2
MDS2
window center
C
PIP PCA
PC1
PC2
longitude
PIP genotype
AA
AB
BB
D
COMP PCA
E
Heterozygosity
PIP
COMP
heterozygosity
genotype
F
Genotype frequencies
HWE D = −0.875 p = 0.0994
G
Linkage disequilibrium
Upper triangle: AA=45, BB=1, mixed genotypes
LD r2
Lower triangle: AA=46, major allele only
H
Genotypes in geographic space
latitude
longitude
genotype

*Geothlypis philadelphia* CM019906.1

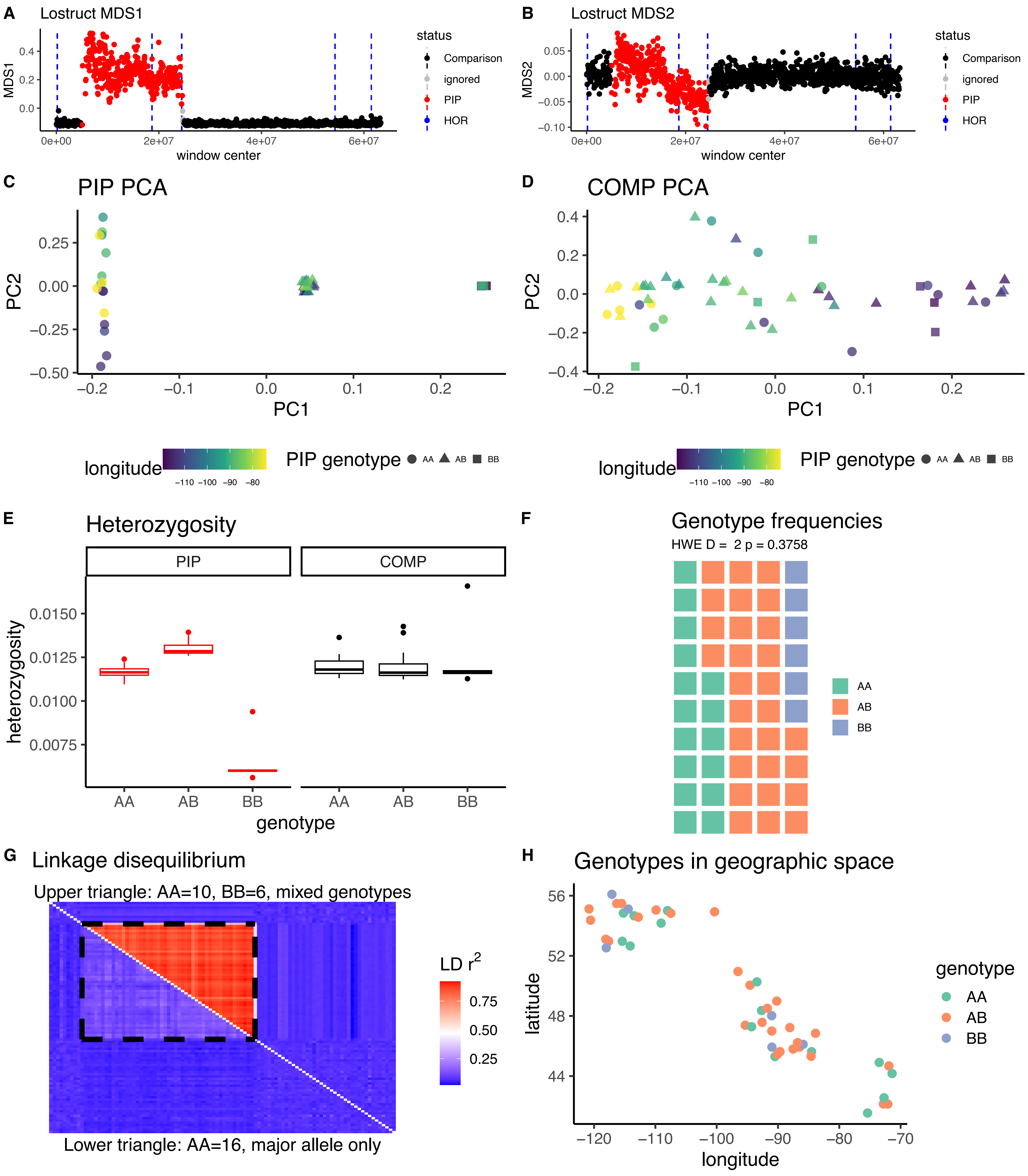

*Geothlypis philadelphia* CM019903.1

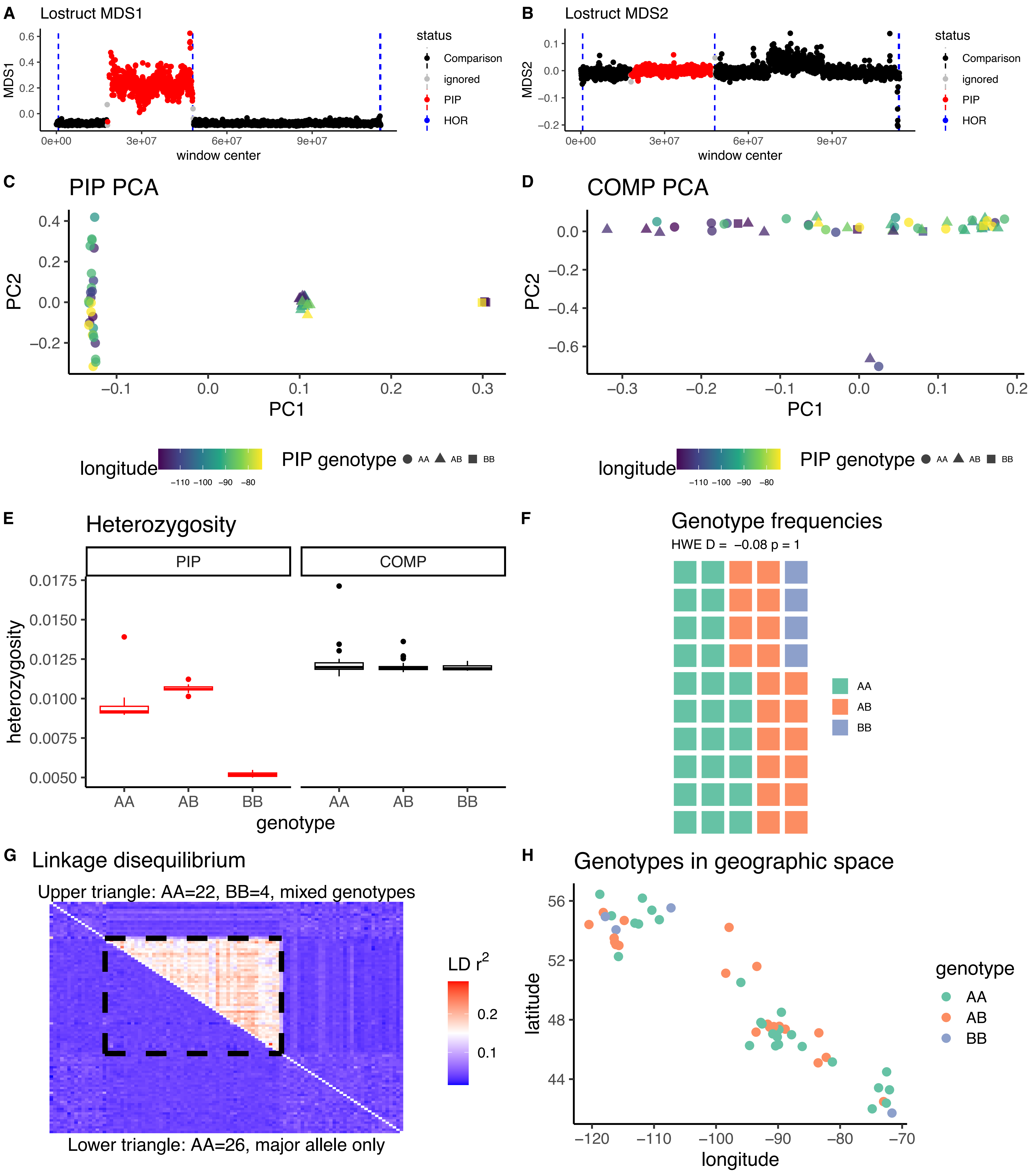

*Geothlypis philadelphia* CM019902.1

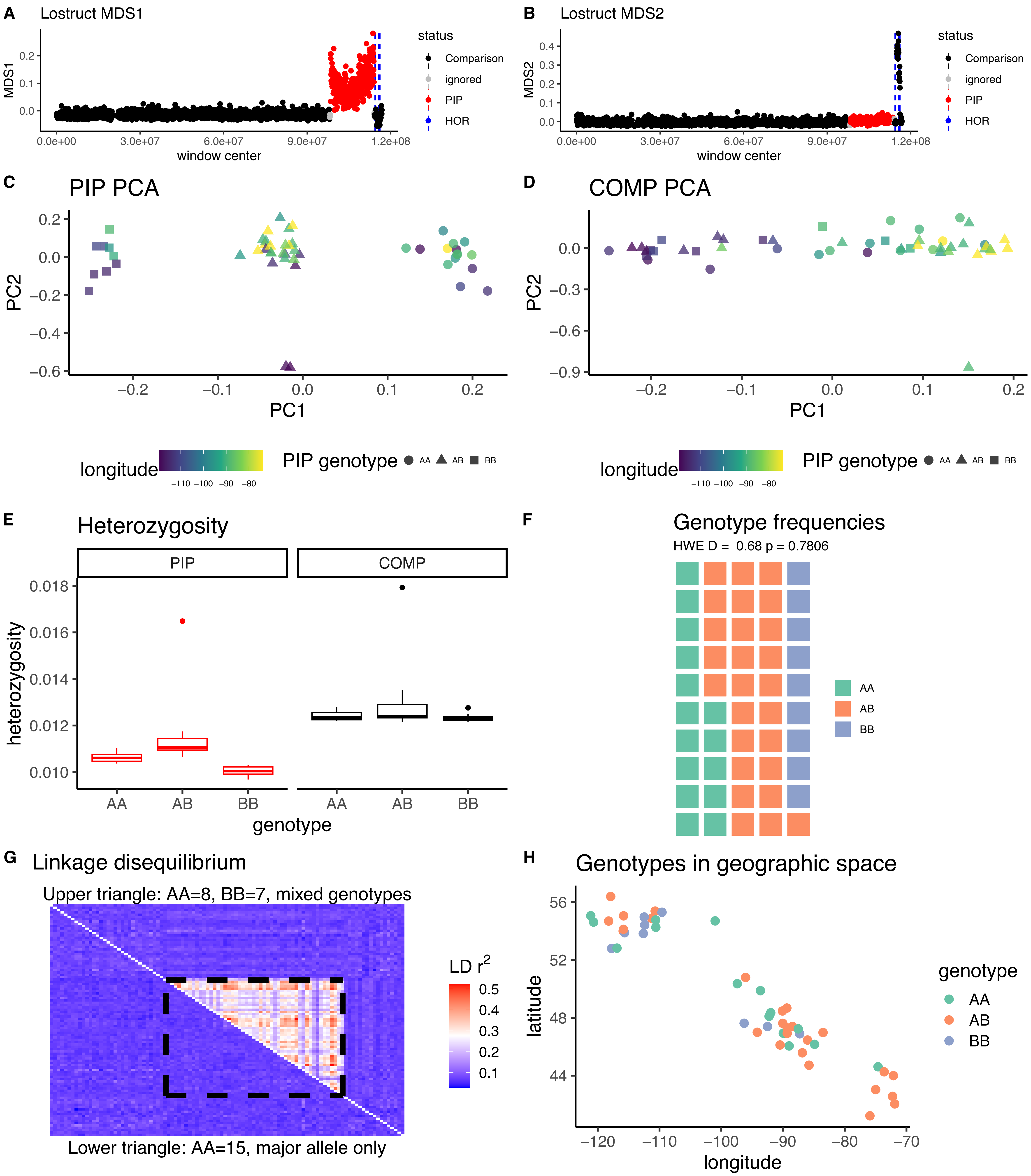

*Leiothlypis peregrina* CM027534.1

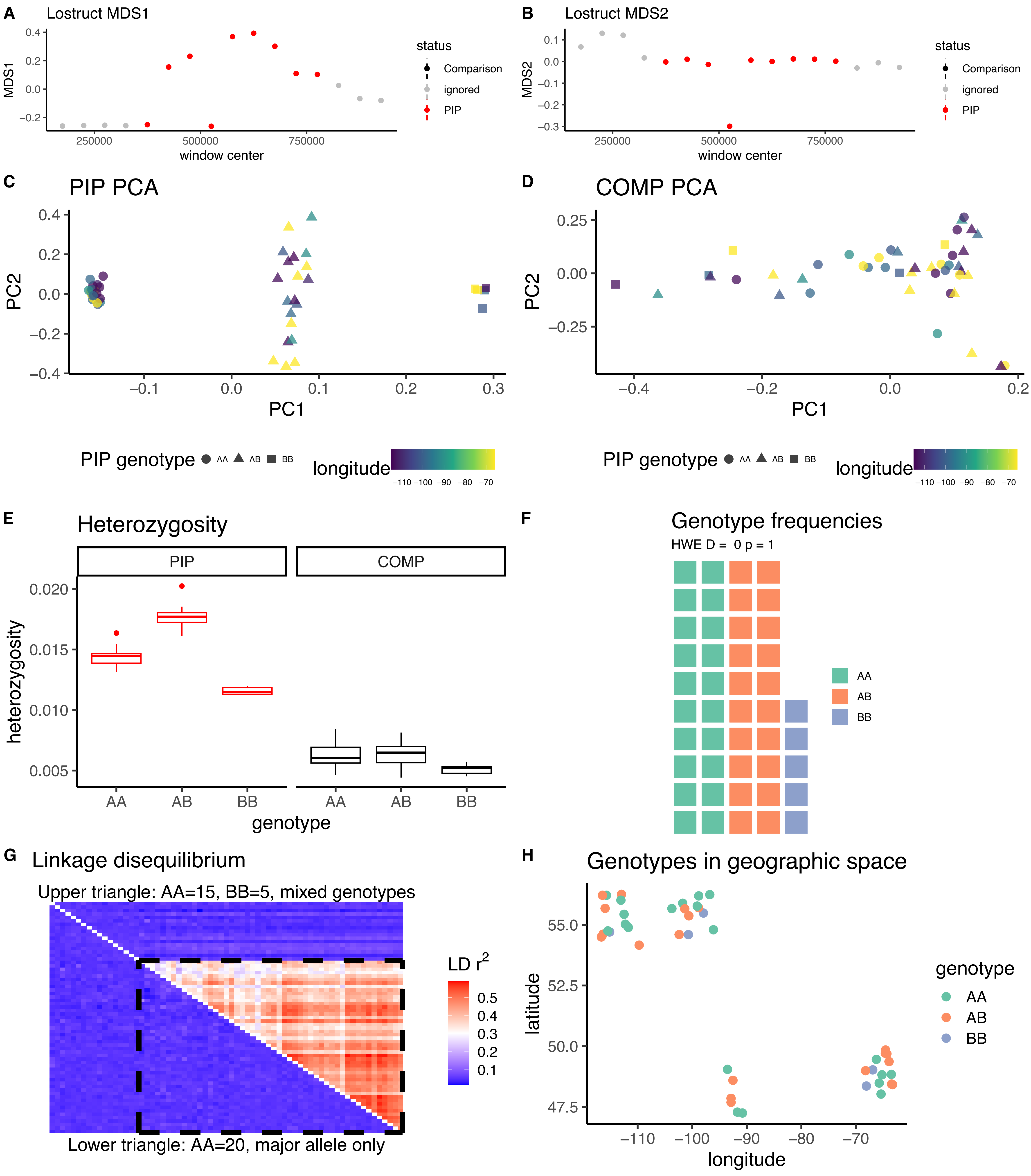

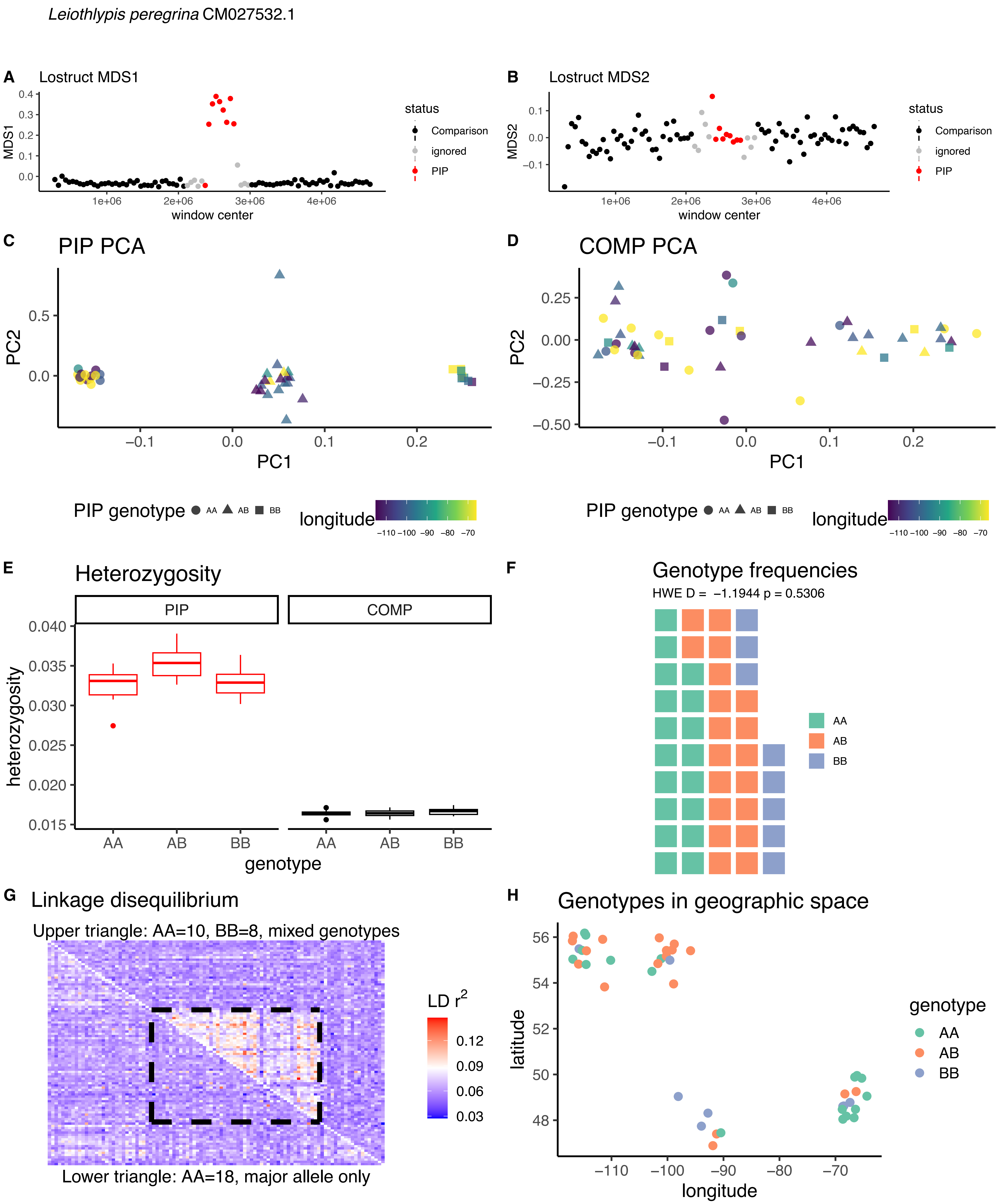

Leiothlypis peregrina CM027532.1
A
Lostruct MDS1
MDS1
window center
status
Comparison
ignored
PIP
B
Lostruct MDS2
MDS2
window center
C
PIP PCA
PC1
PC2
D
COMP PCA
PIP genotype
AA
AB
BB
longitude
E
Heterozygosity
PIP
COMP
heterozygosity
genotype
F
Genotype frequencies
HWE D = −1.1944 p = 0.5306
G
Linkage disequilibrium
Upper triangle: AA=10, BB=8, mixed genotypes
LD r2
Lower triangle: AA=18, major allele only
H
Genotypes in geographic space
latitude
longitude

*Leiothlypis peregrina* CM027530.1

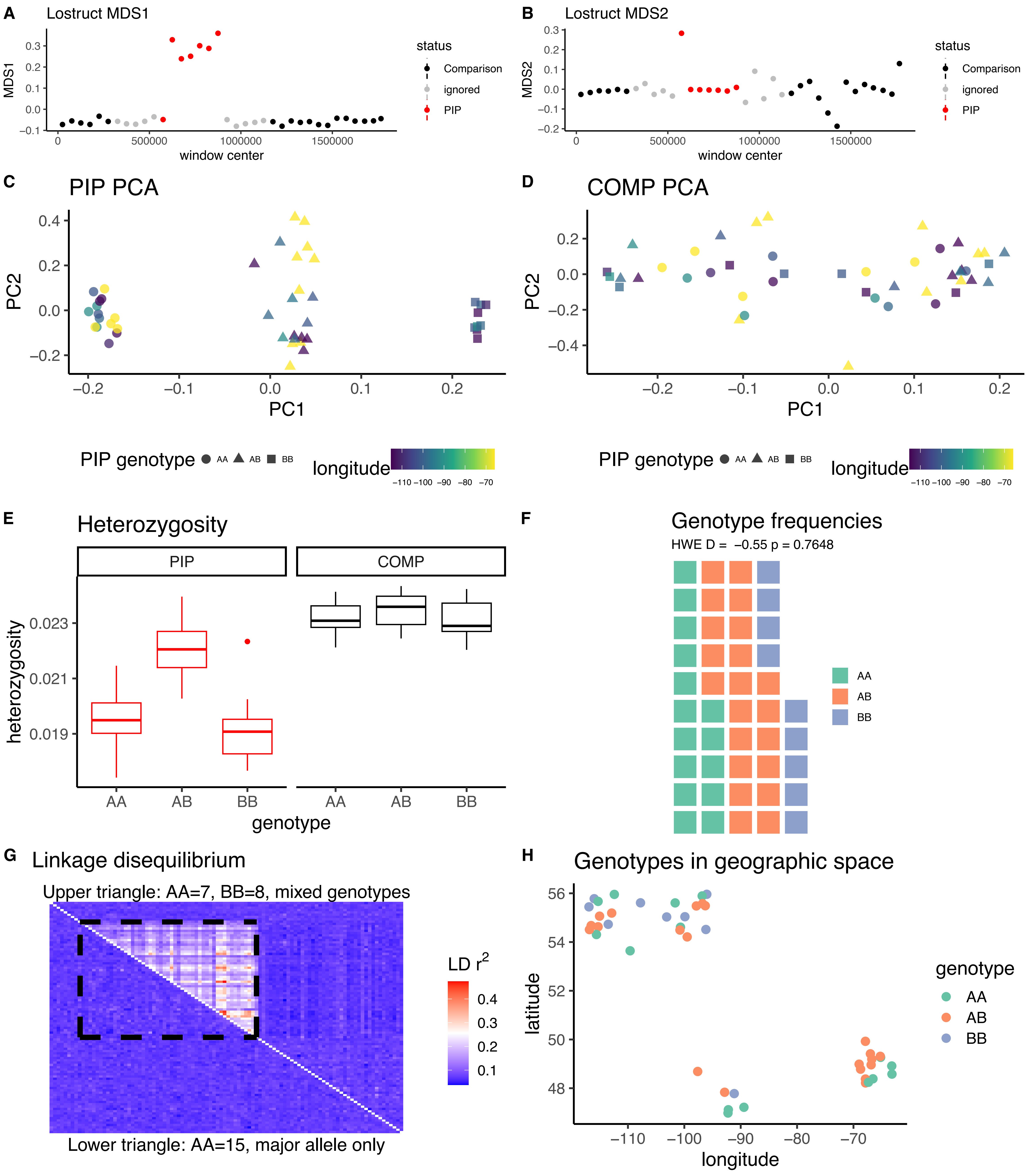

*Leiothlypis peregrina* CM027528.1

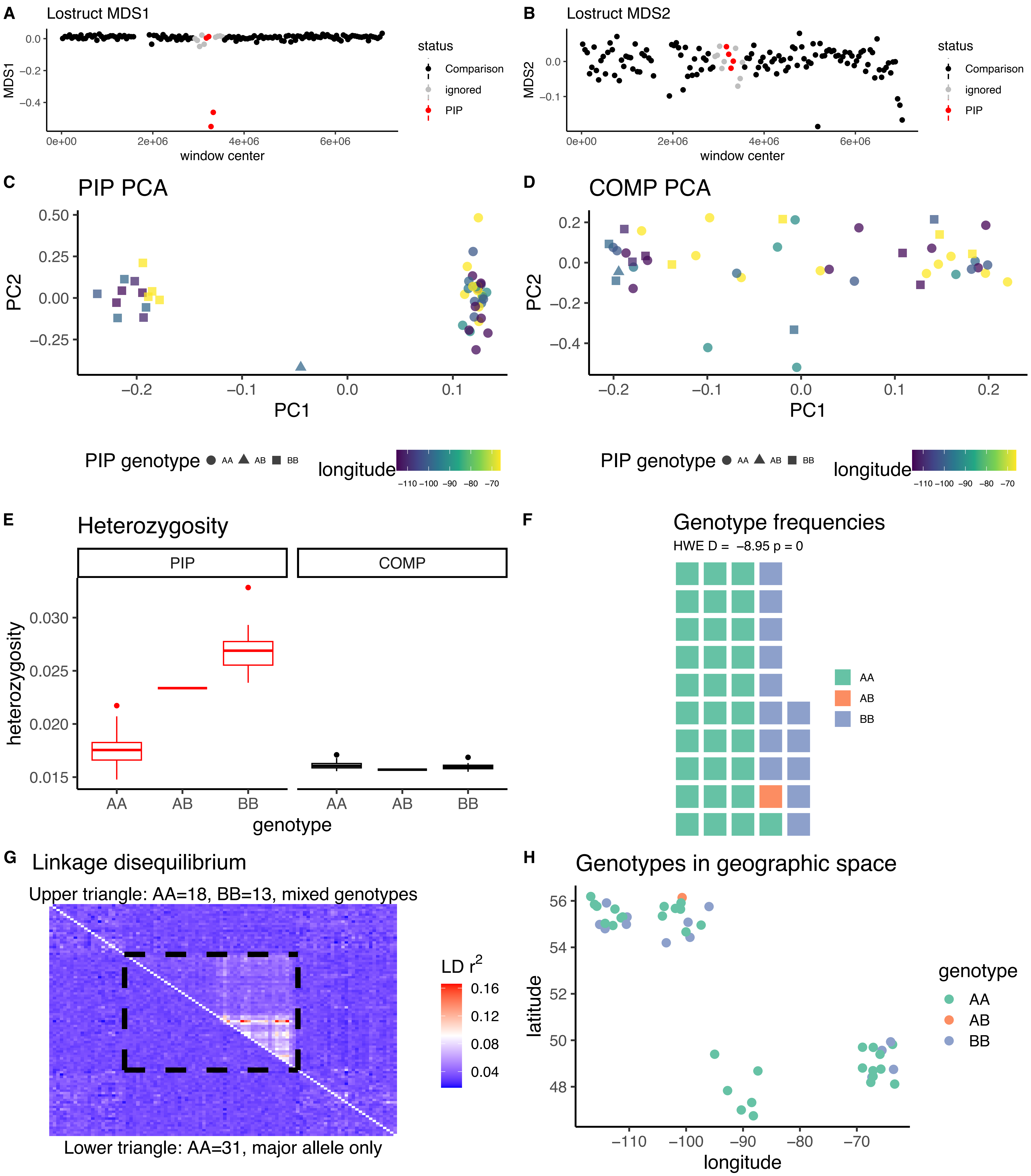

*Leiothlypis peregrina* CM027512.1

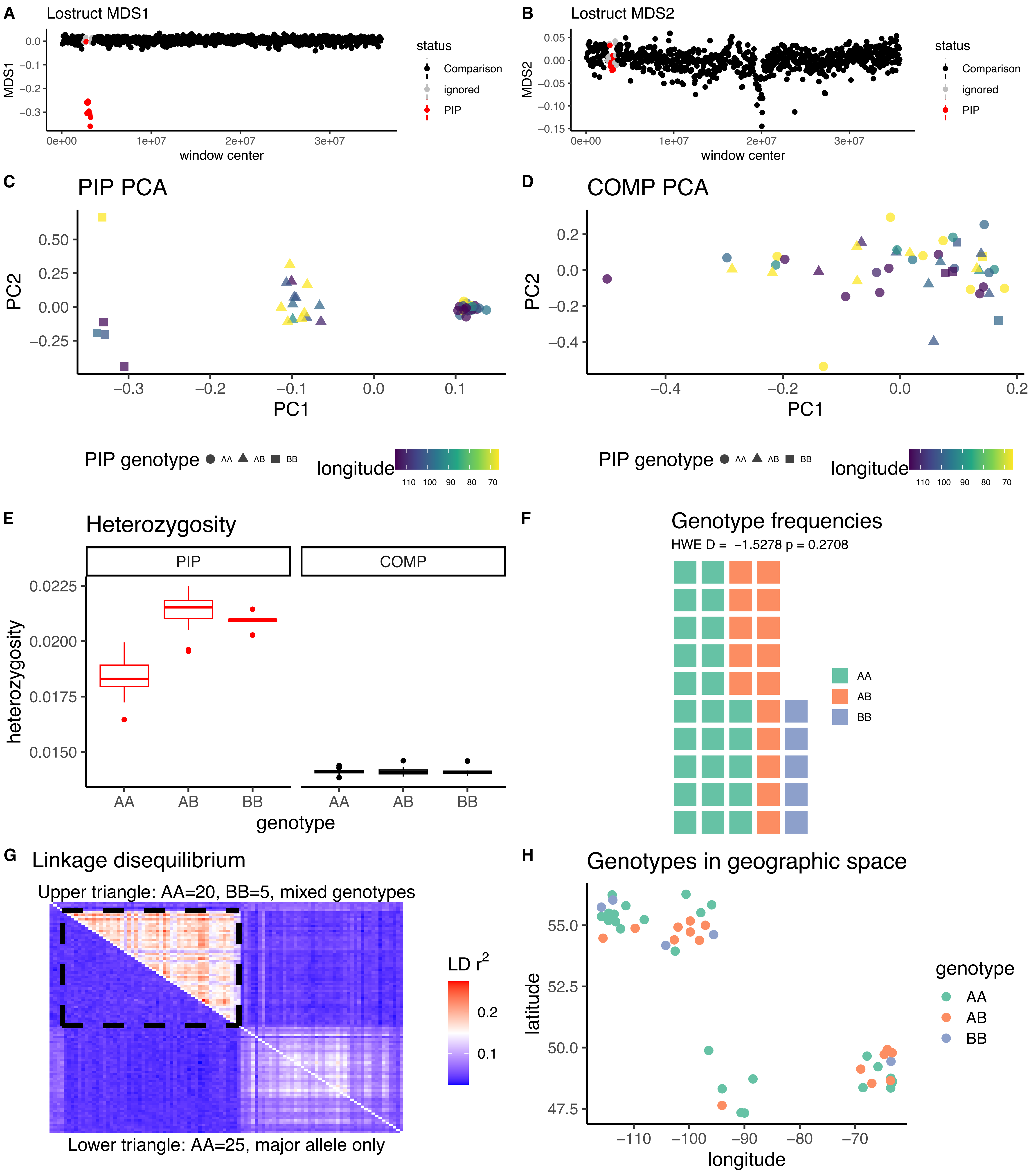

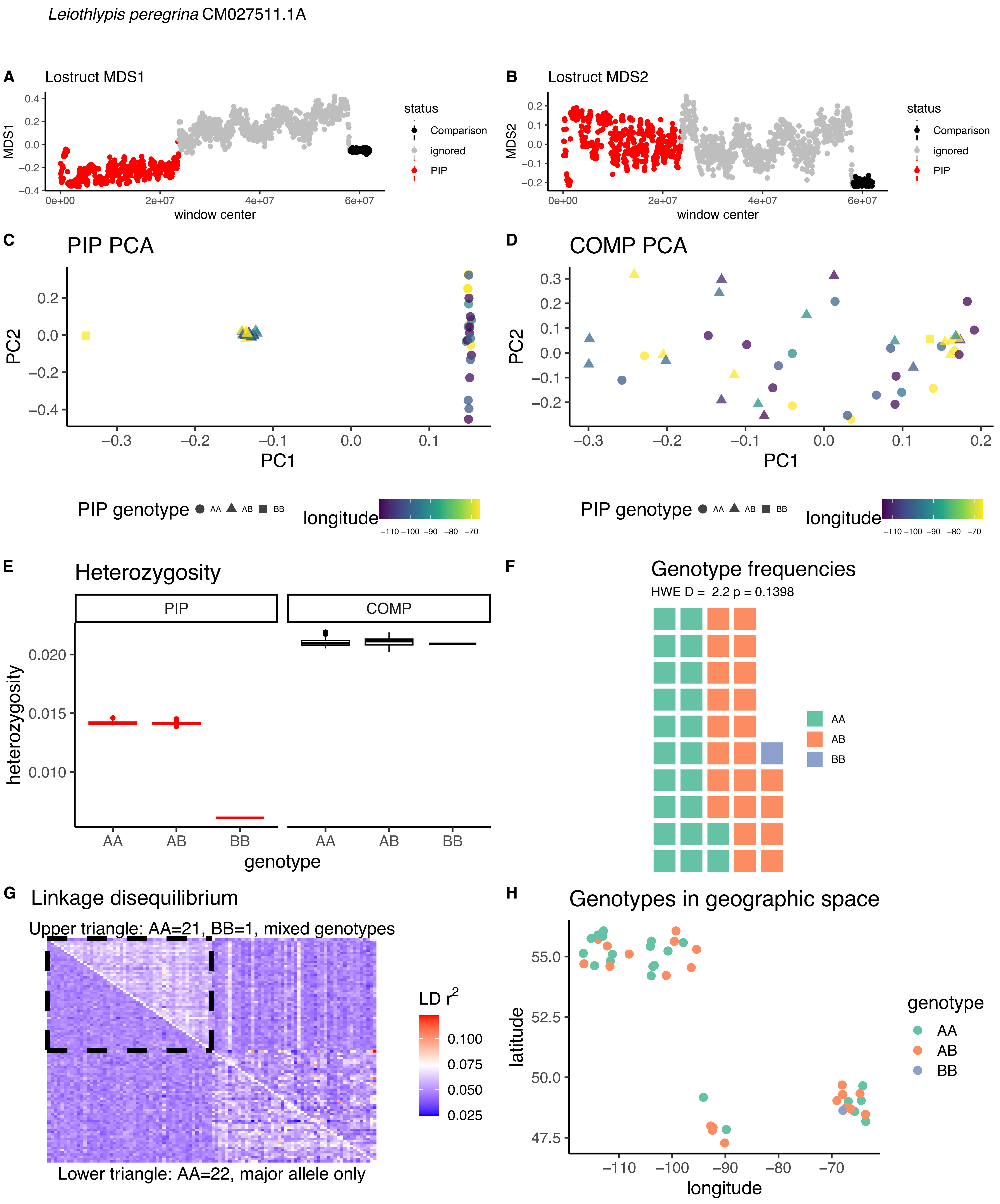

*Leiothlypis peregrina* CM027511.1A
A Lostruct MDS1
B Lostruct MDS2
MDS1
MDS2
window center
status
Comparison
ignored
PIP
C PIP PCA
D COMP PCA
PC1
PC2
PIP genotype AA AB BB
longitude
E Heterozygosity
PIP
COMP
heterozygosity
genotype
F Genotype frequencies
HWE D = 2.2 p = 0.1398
AA
AB
BB
G Linkage disequilibrium
Upper triangle: AA=21, BB=1, mixed genotypes
Lower triangle: AA=22, major allele only
LD $r^2$
H Genotypes in geographic space
latitude
longitude
genotype
AA
AB
BB

*Leiothlypis peregrina* CM027511.1B

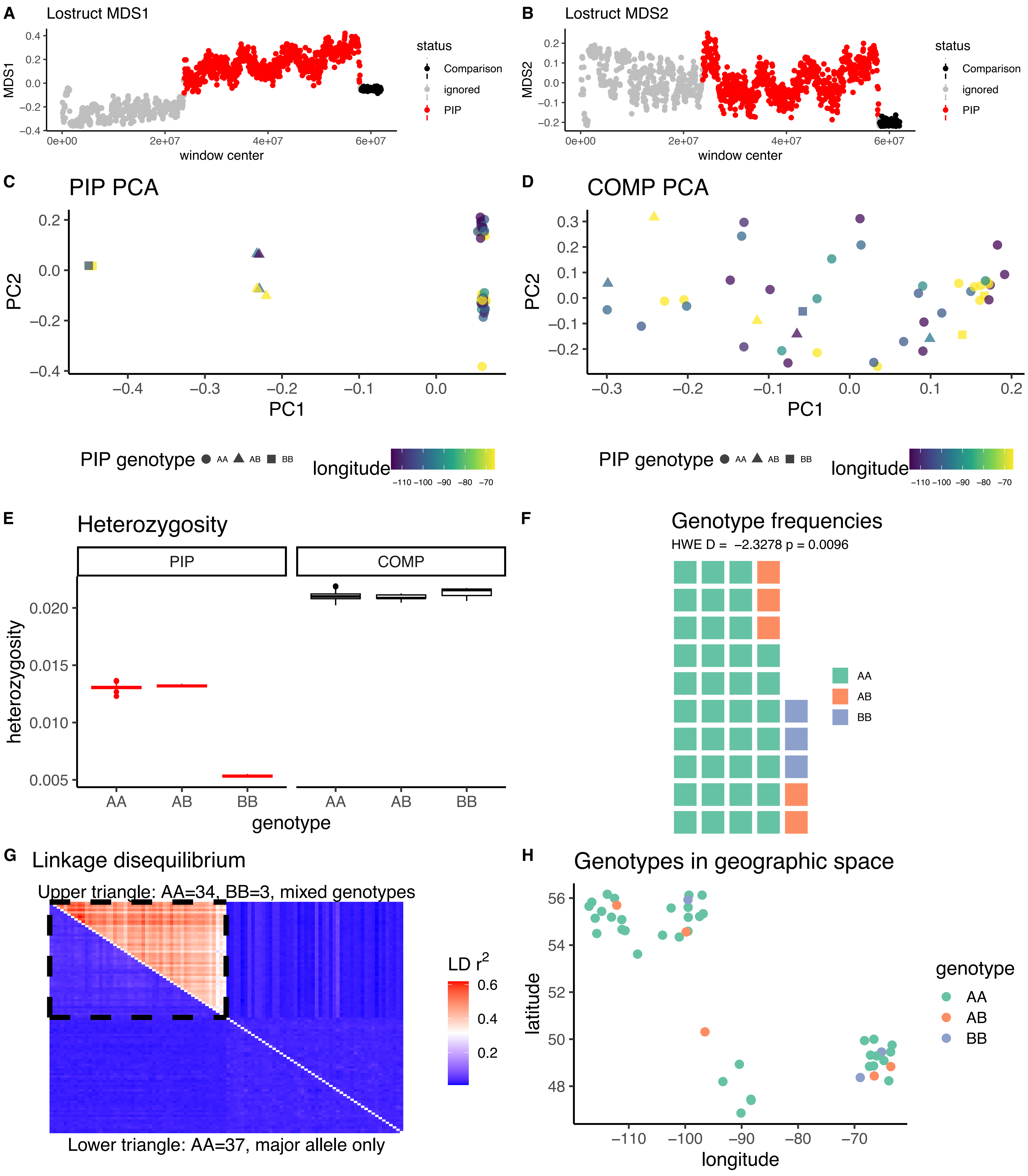

*Leiothlypis ruficapilla* CM027534.1

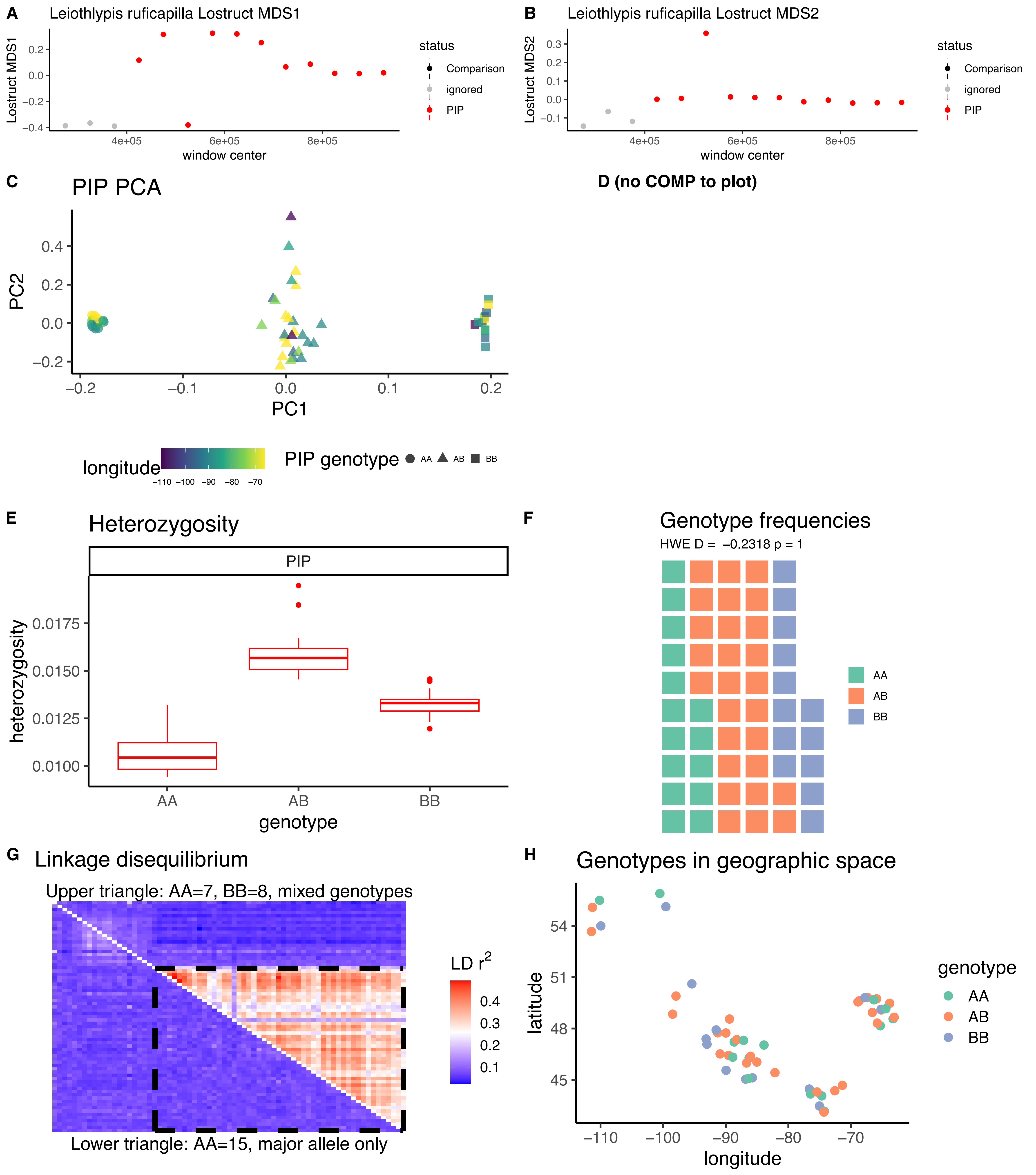

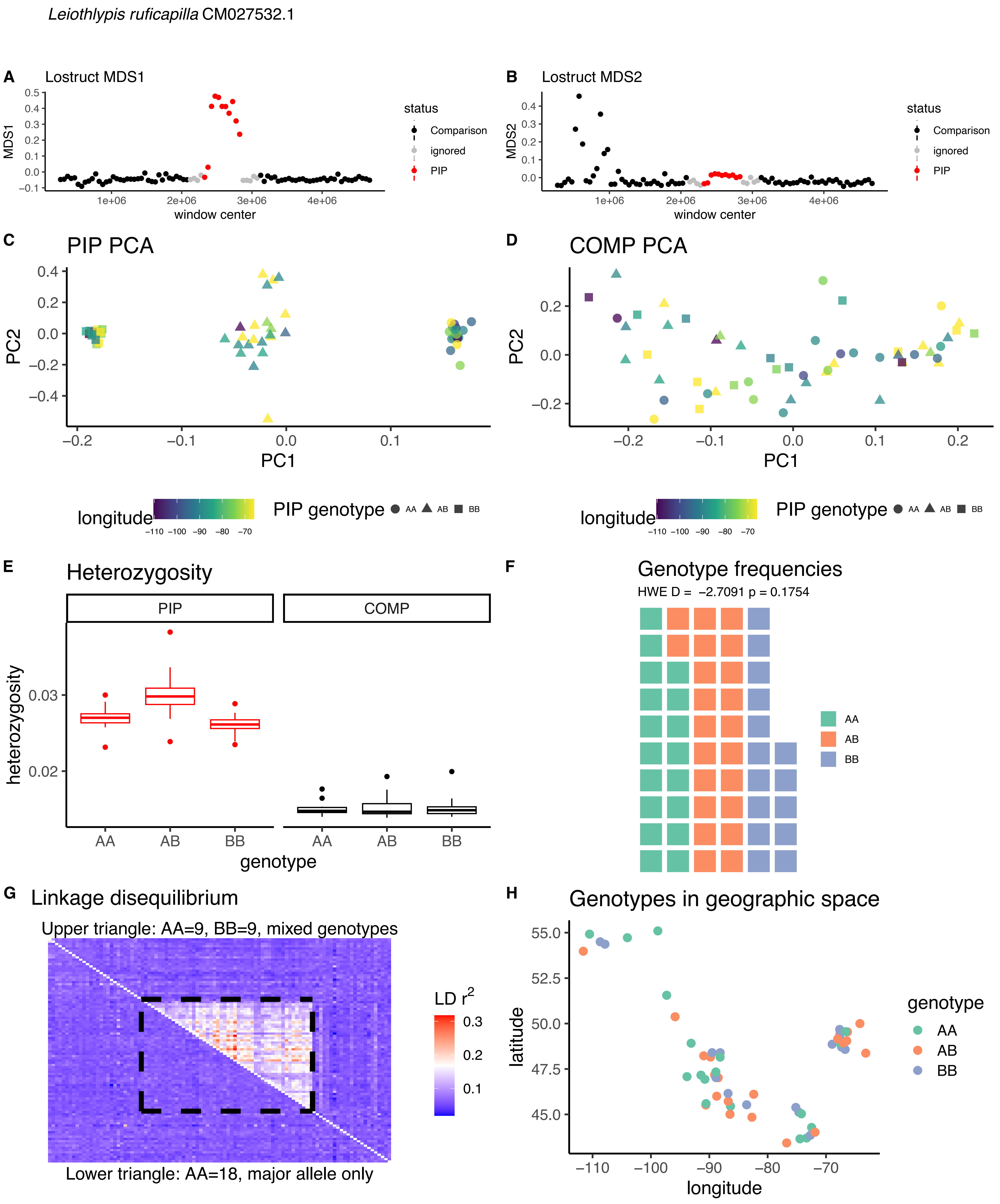

Leiothlypis ruficapilla CM027532.1
A
Lostruct MDS1
MDS1
window center
status
Comparison
ignored
PIP
B
Lostruct MDS2
MDS2
window center
C
PIP PCA
PC1
PC2
D
COMP PCA
longitude
PIP genotype
AA
AB
BB
E
Heterozygosity
PIP
COMP
heterozygosity
genotype
F
Genotype frequencies
HWE D = -2.7091 p = 0.1754
G
Linkage disequilibrium
Upper triangle: AA=9, BB=9, mixed genotypes
Lower triangle: AA=18, major allele only
LD r2
H
Genotypes in geographic space
latitude
longitude
genotype

*Leiothlypis ruficapilla* CM027531.1

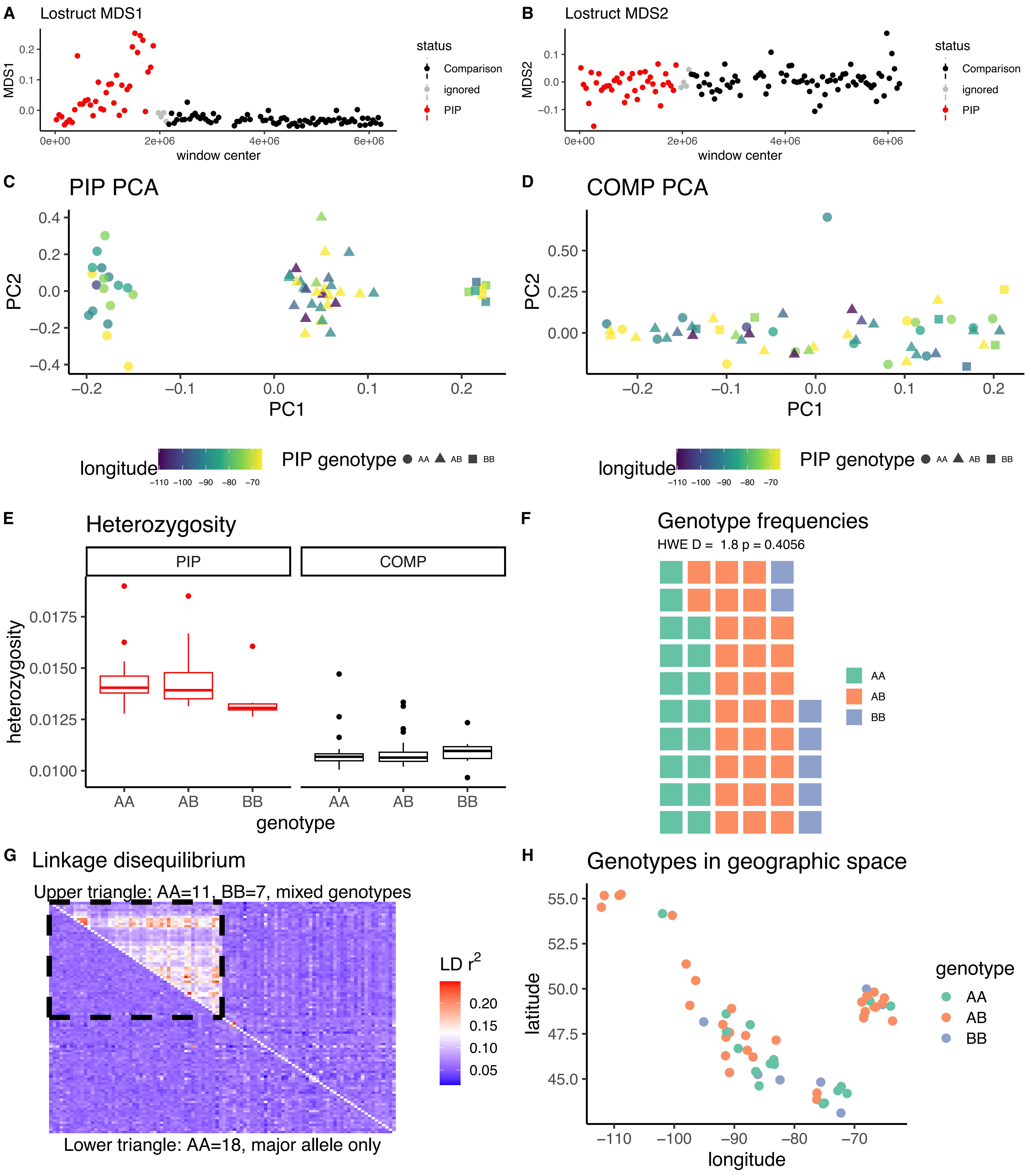

*Leiothlypis ruficapilla* CM027530.1

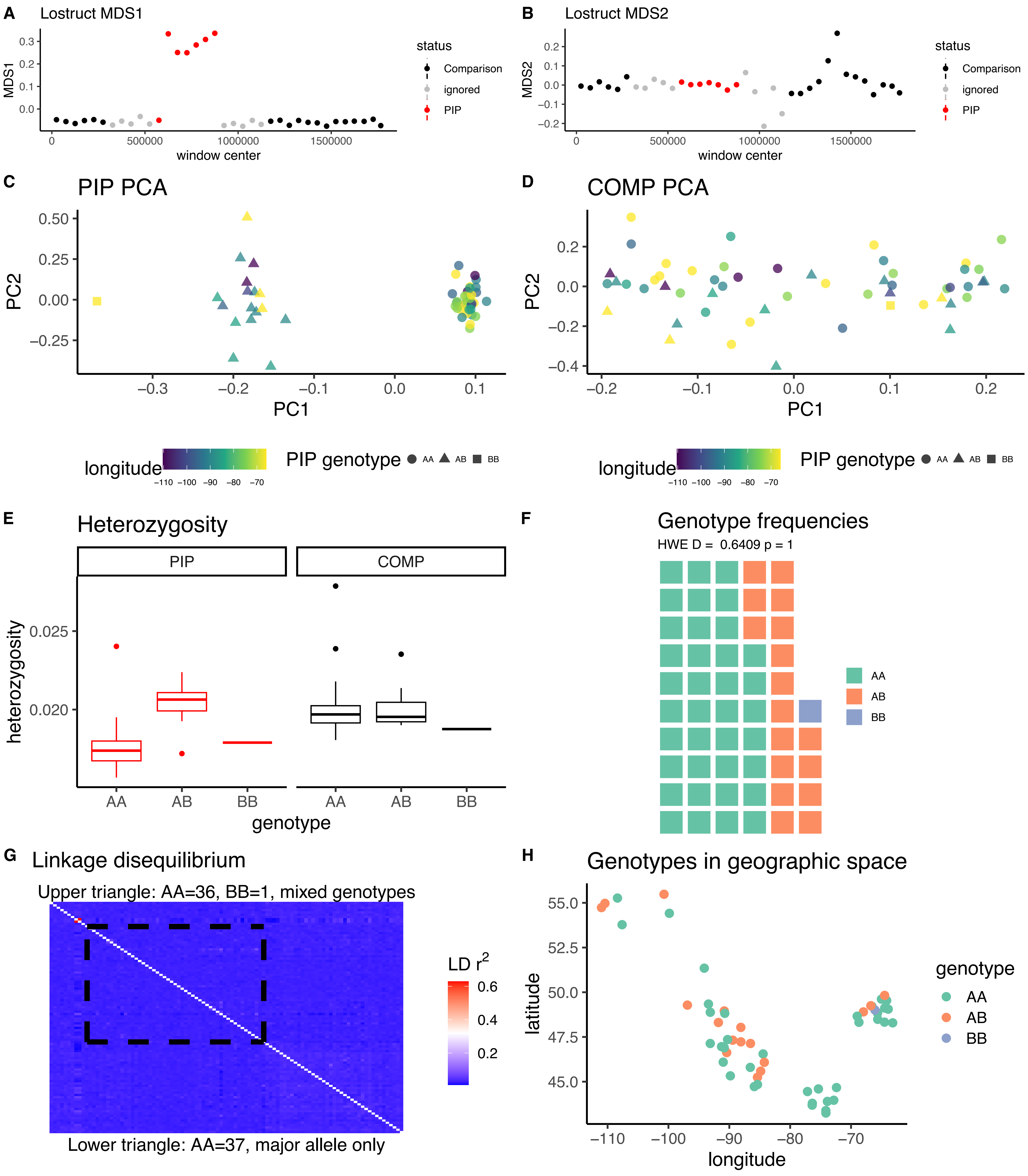

*Leiothlypis ruficapilla* CM027518.1A

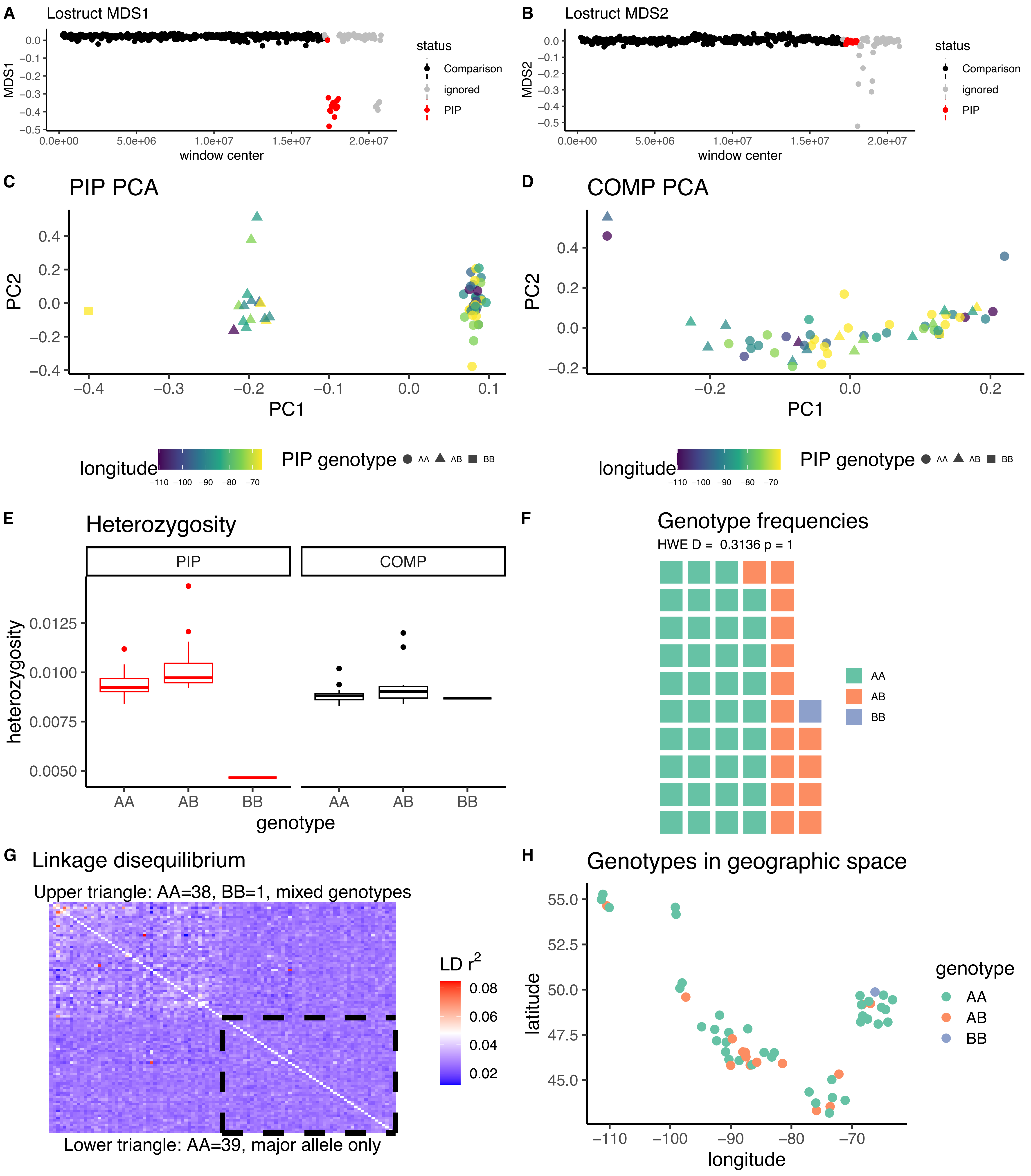

*Leiothlypis ruficapilla* CM027518.1B

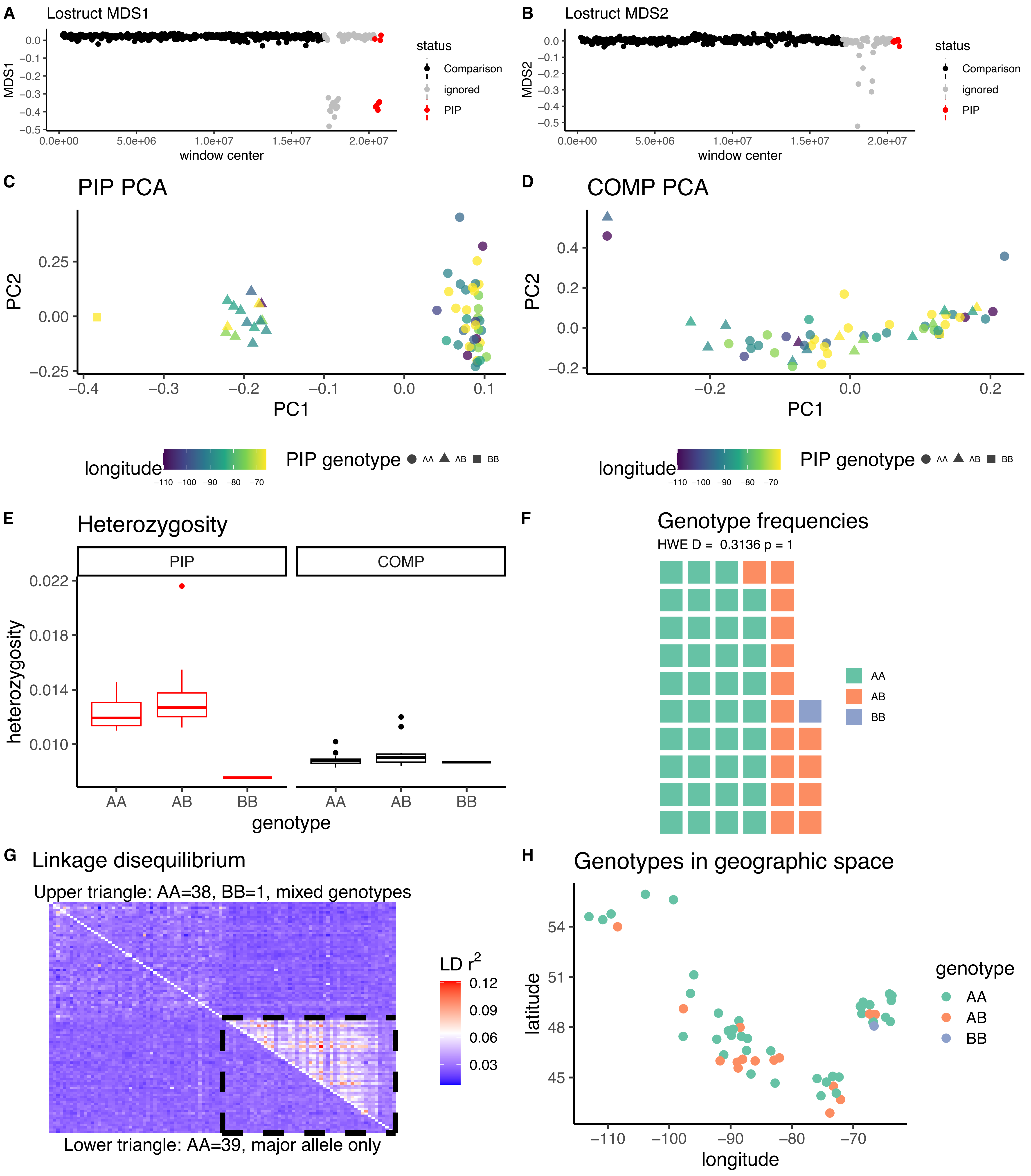

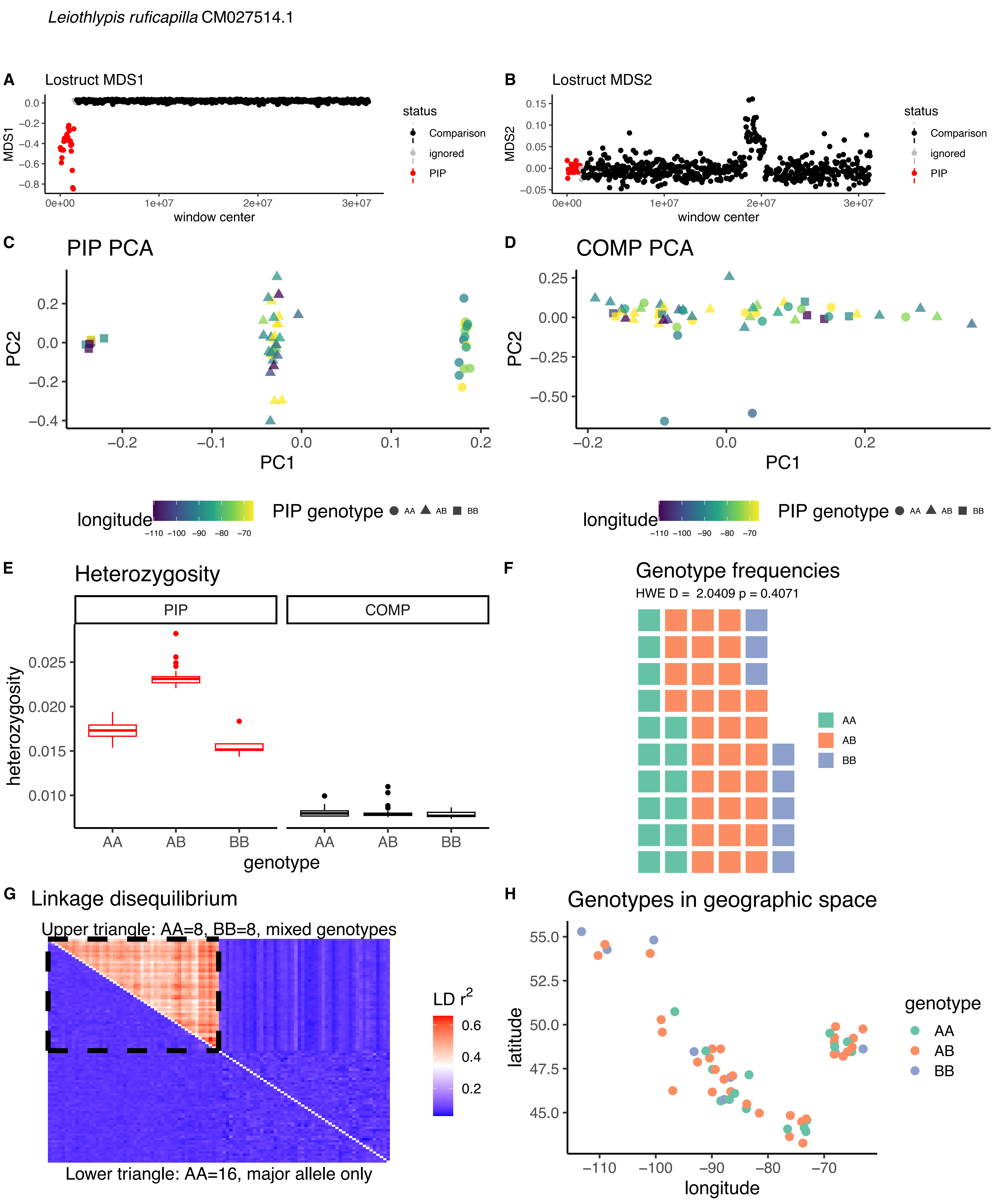
Leiothlypis ruficapilla CM027514.1
A
Lostruct MDS1
MDS1
window center
status
Comparison
ignored
PIP
B
Lostruct MDS2
MDS2
C
PIP PCA
PC1
PC2
D
COMP PCA
longitude
PIP genotype
AA
AB
BB
E
Heterozygosity
PIP
COMP
heterozygosity
genotype
F
Genotype frequencies
HWE D = 2.0409 p = 0.4071
G
Linkage disequilibrium
Upper triangle: AA=8, BB=8, mixed genotypes
Lower triangle: AA=16, major allele only
LD r2
H
Genotypes in geographic space
latitude
longitude

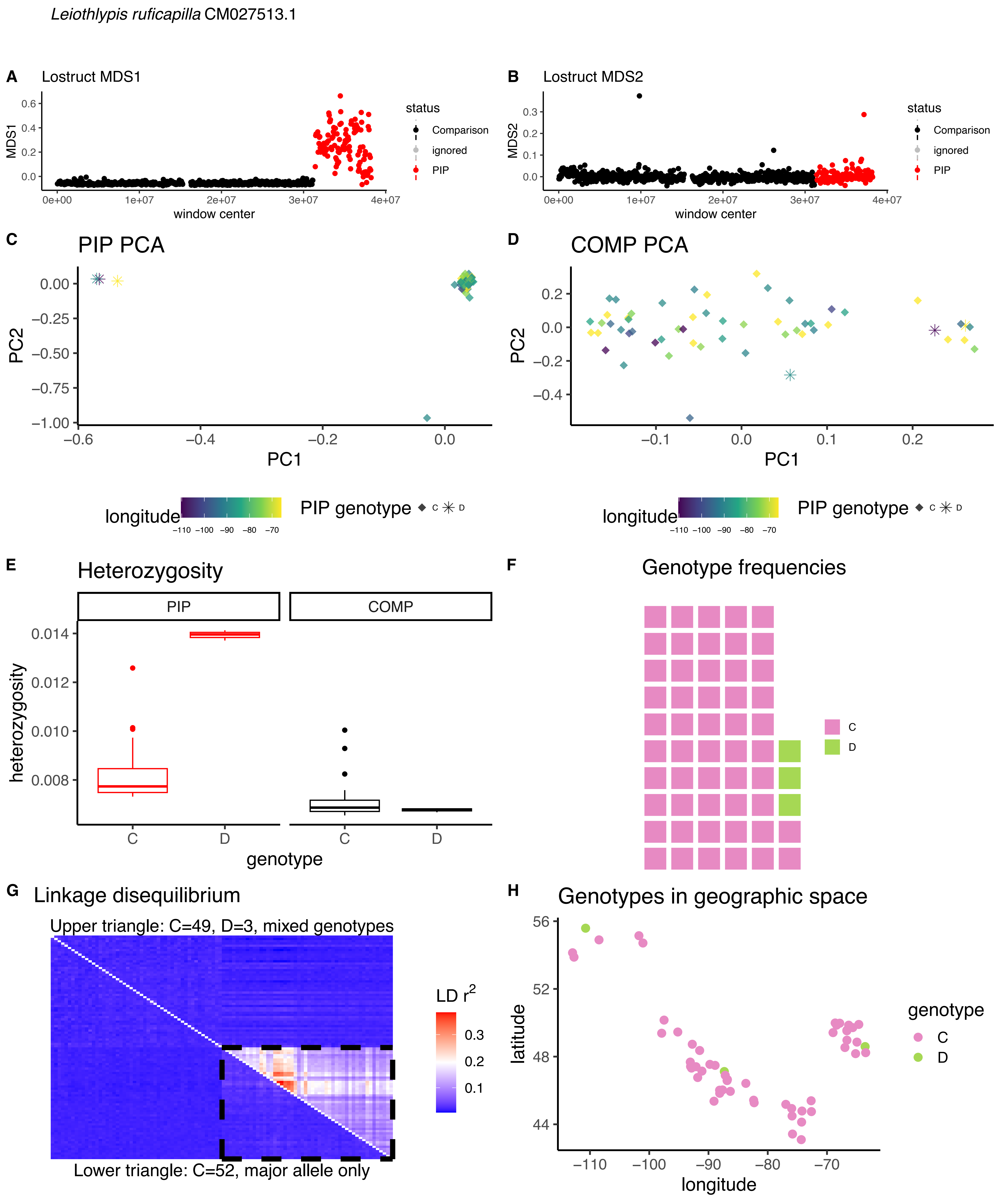

Leiothlypis ruficapilla CM027513.1
A
Lostruct MDS1
MDS1
window center
status
Comparison
ignored
PIP
B
Lostruct MDS2
MDS2
window center
C
PIP PCA
PC1
PC2
D
COMP PCA
longitude
PIP genotype
C
D
E
Heterozygosity
PIP
COMP
heterozygosity
genotype
F
Genotype frequencies
G
Linkage disequilibrium
Upper triangle: C=49, D=3, mixed genotypes
Lower triangle: C=52, major allele only
LD $r^2$
H
Genotypes in geographic space
latitude
longitude
genotype

*Leiothlypis ruficapilla* CM027512.1

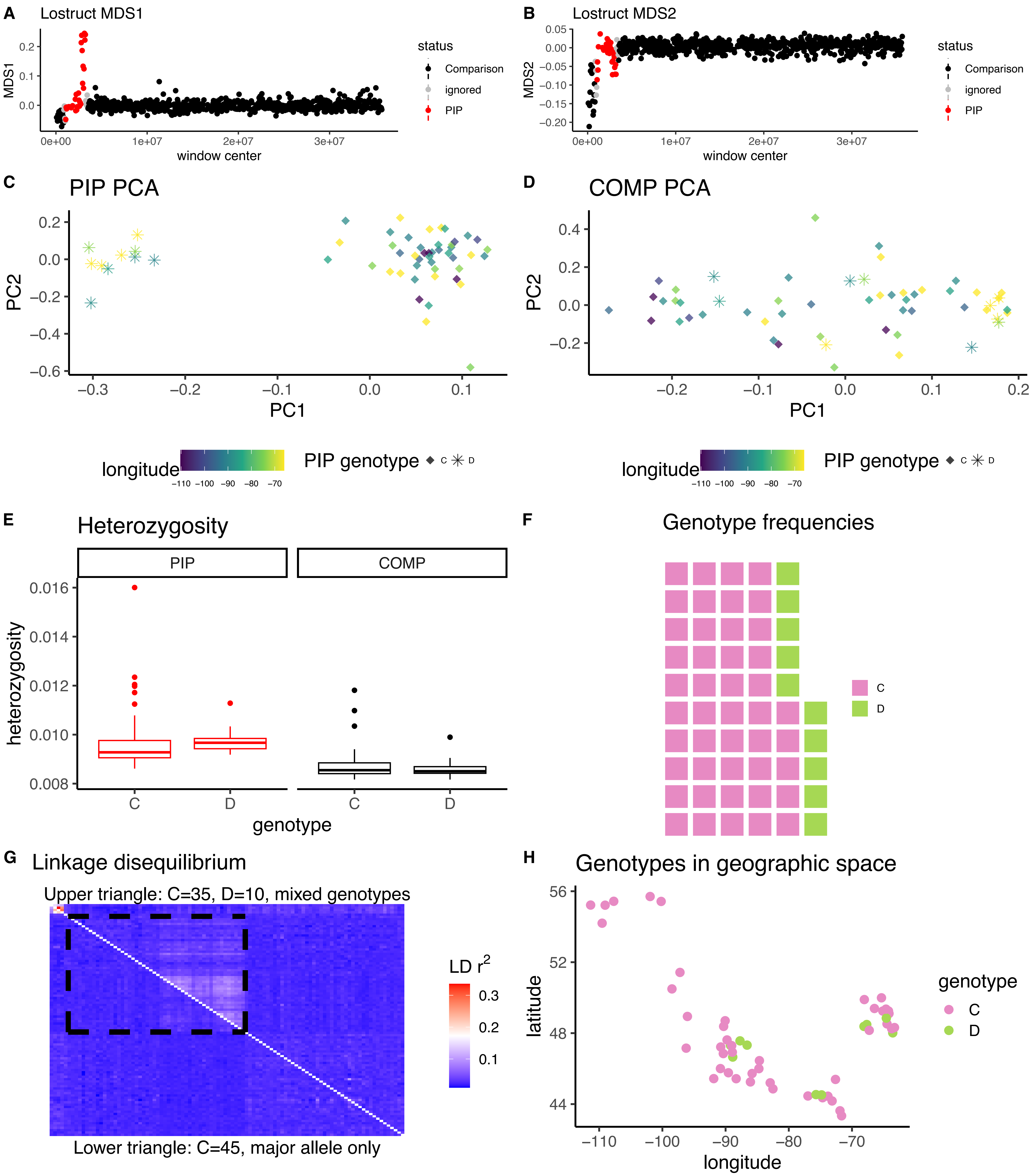

*Leiothlypis ruficapilla* CM027510.1

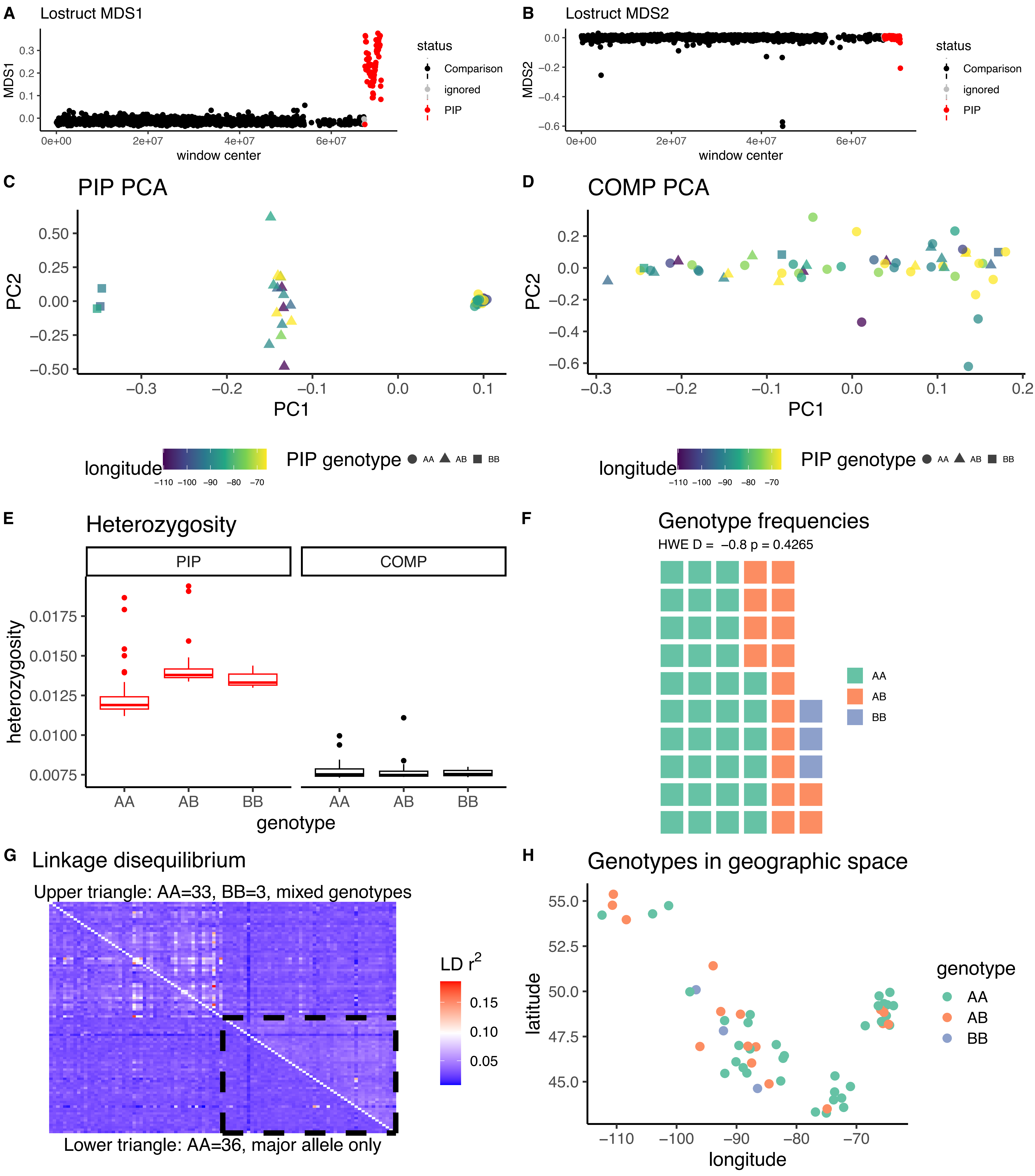

*Leiothlypis ruficapilla* CM027508.1

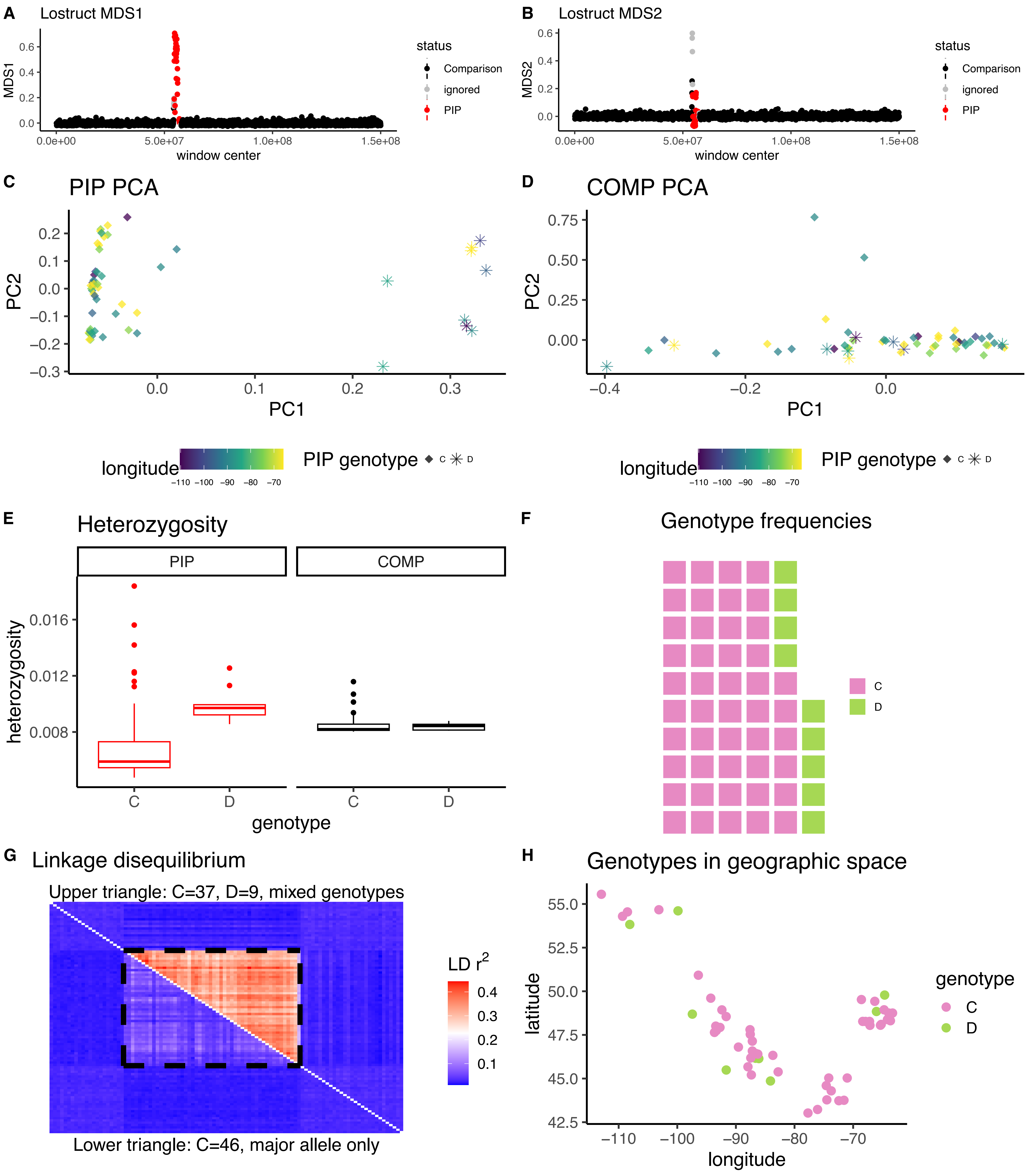

*Oporornis agilis* CM019929.1

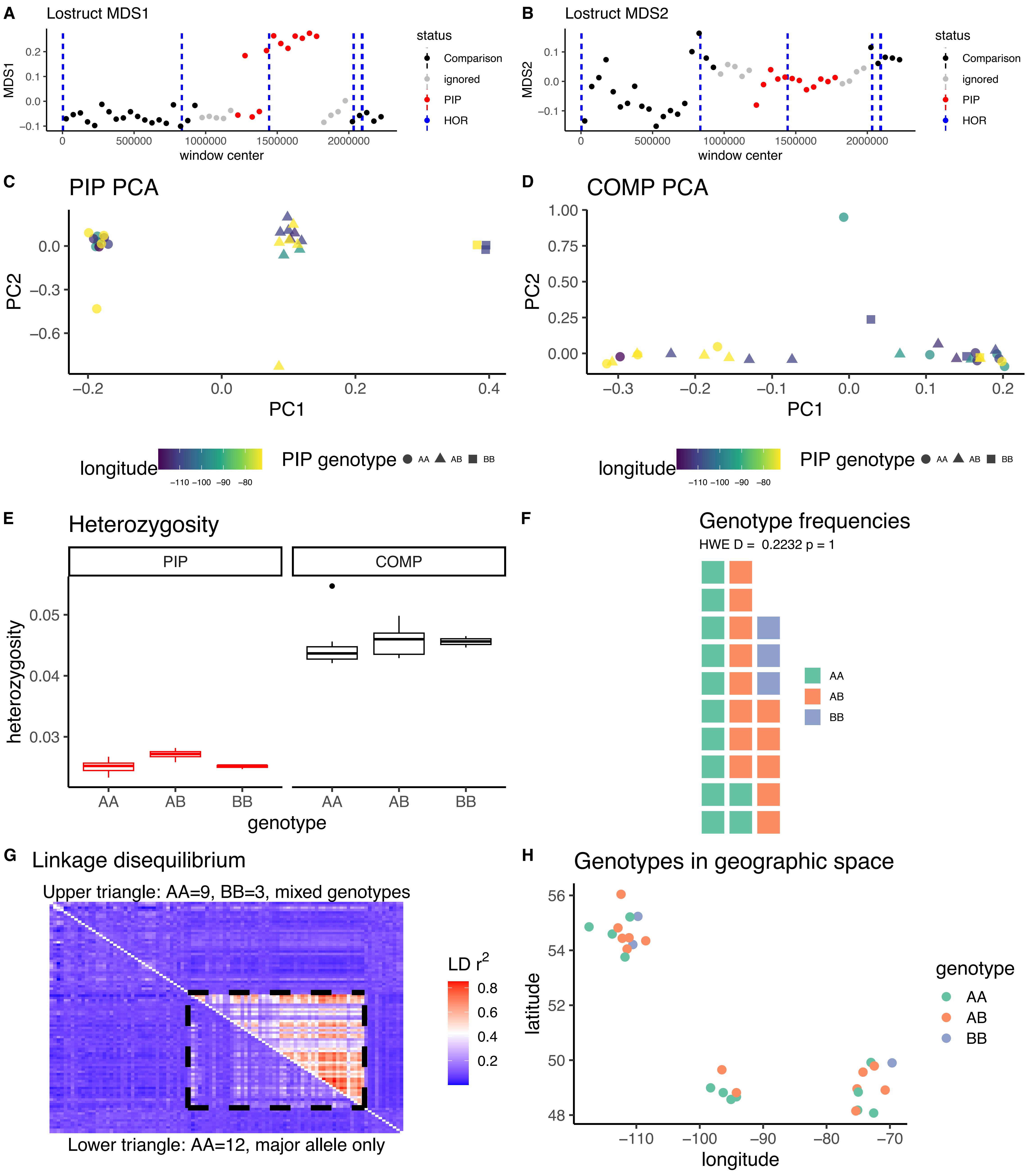

*Oporornis agilis* CM019921.1

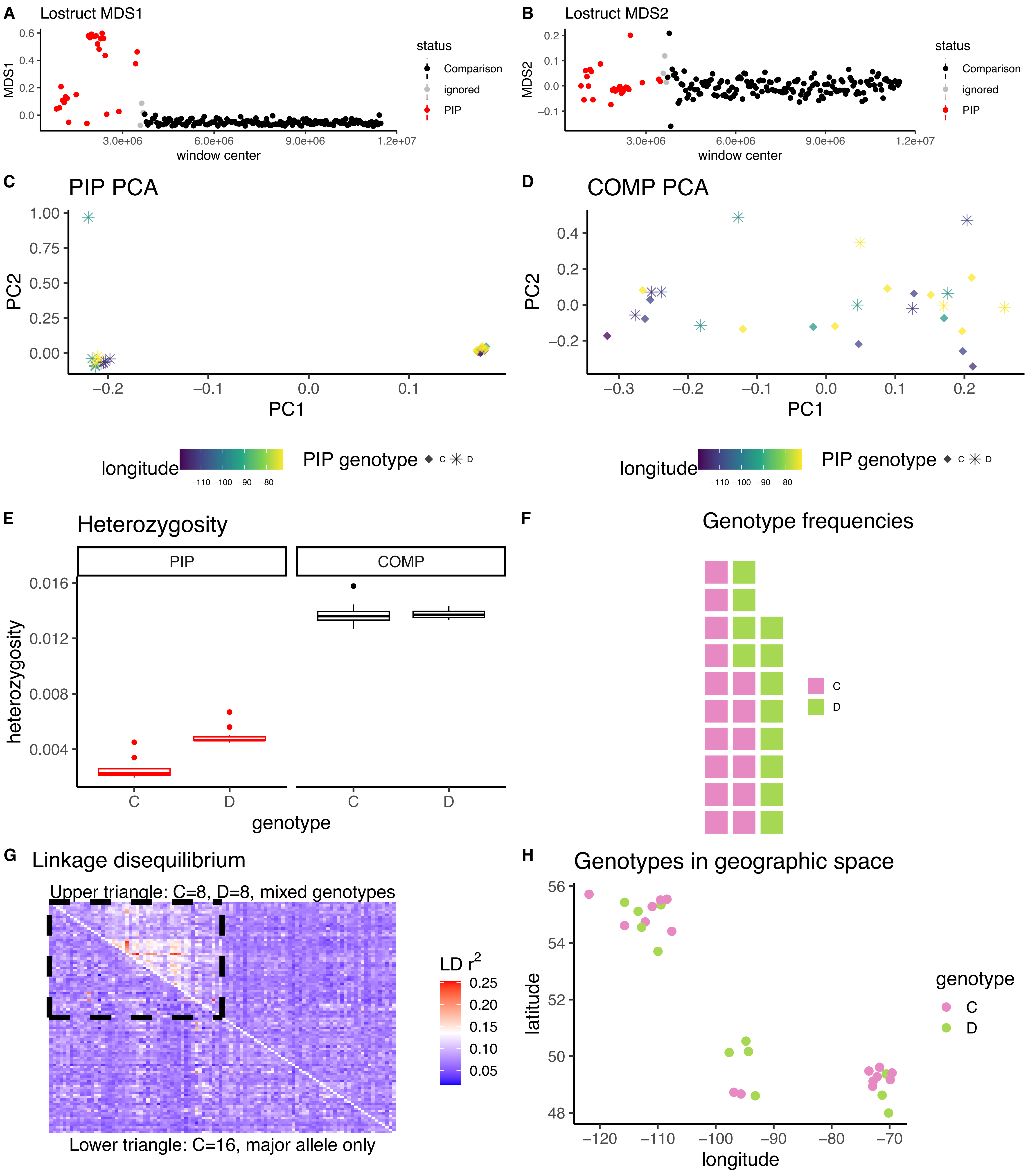

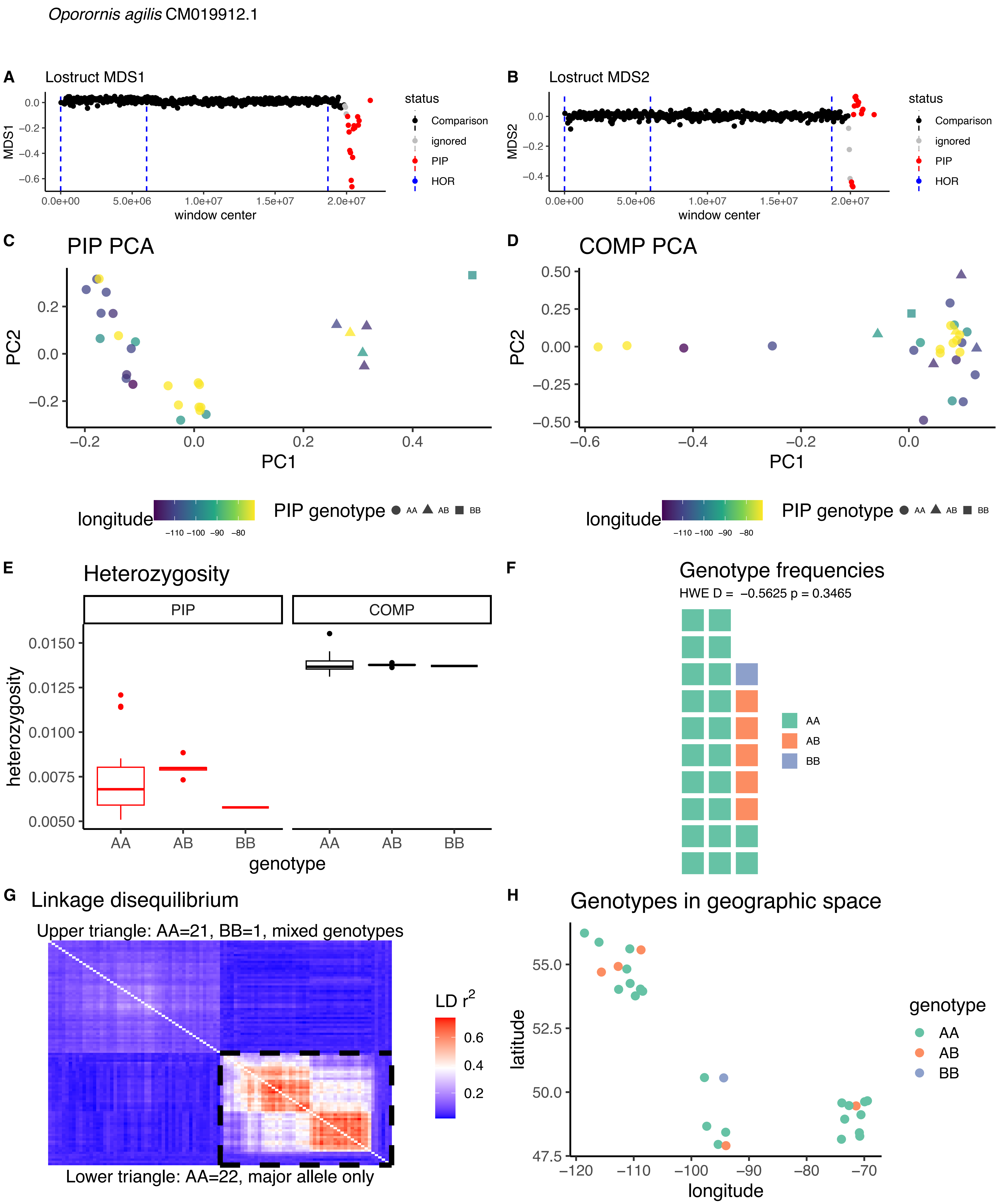

Oporornis agilis CM019912.1
A Lostruct MDS1
B Lostruct MDS2
MDS1
MDS2
window center
status
Comparison
ignored
PIP
HOR
C PIP PCA
D COMP PCA
PC1
PC2
longitude
PIP genotype AA AB BB
E Heterozygosity
PIP
COMP
heterozygosity
genotype
F Genotype frequencies
HWE D = −0.5625 p = 0.3465
AA
AB
BB
G Linkage disequilibrium
Upper triangle: AA=21, BB=1, mixed genotypes
Lower triangle: AA=22, major allele only
LD r2
H Genotypes in geographic space
latitude
longitude
genotype

*Oporornis agilis* CM019902.1

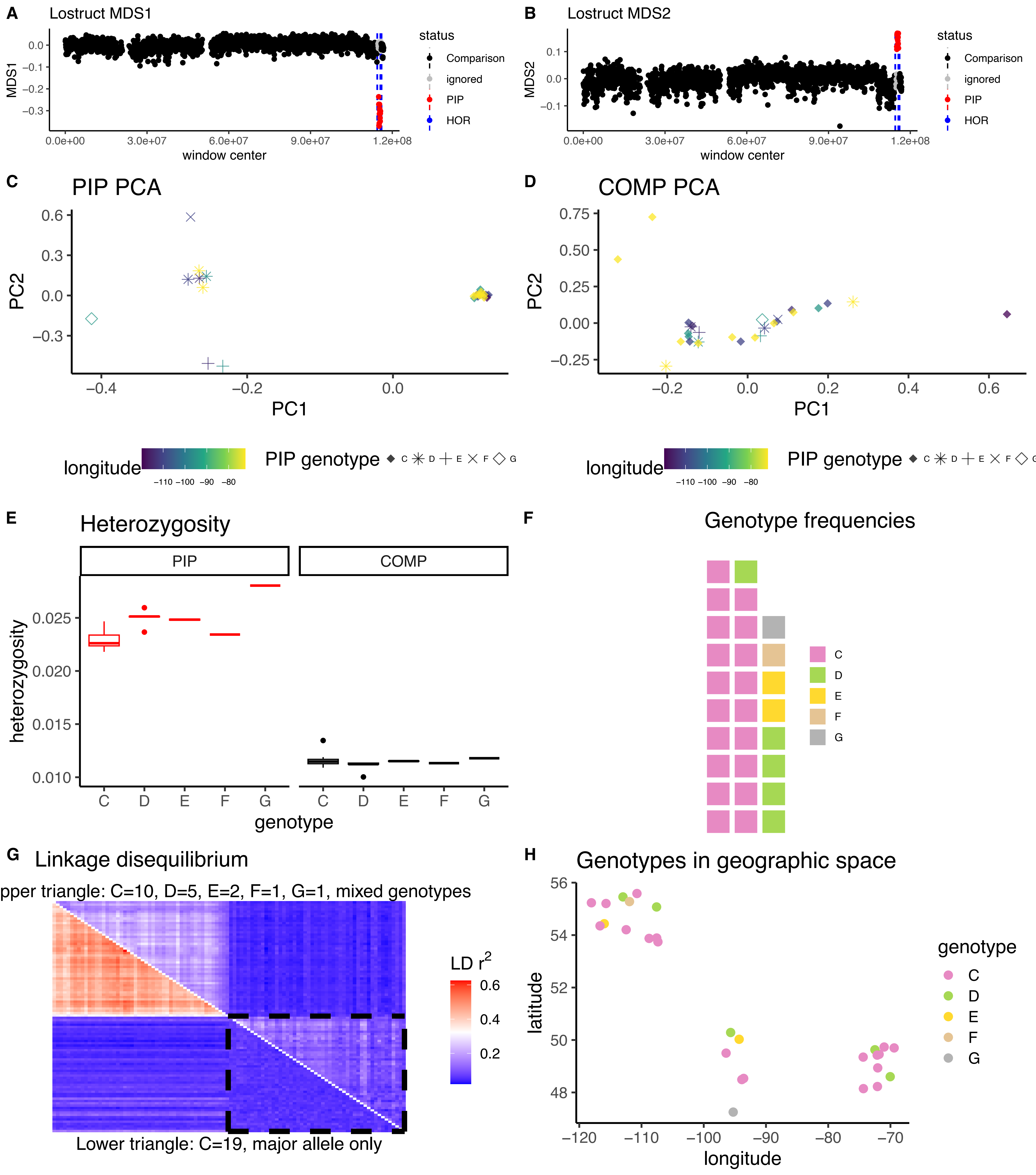

*Setophaga castanea* CM027534.1

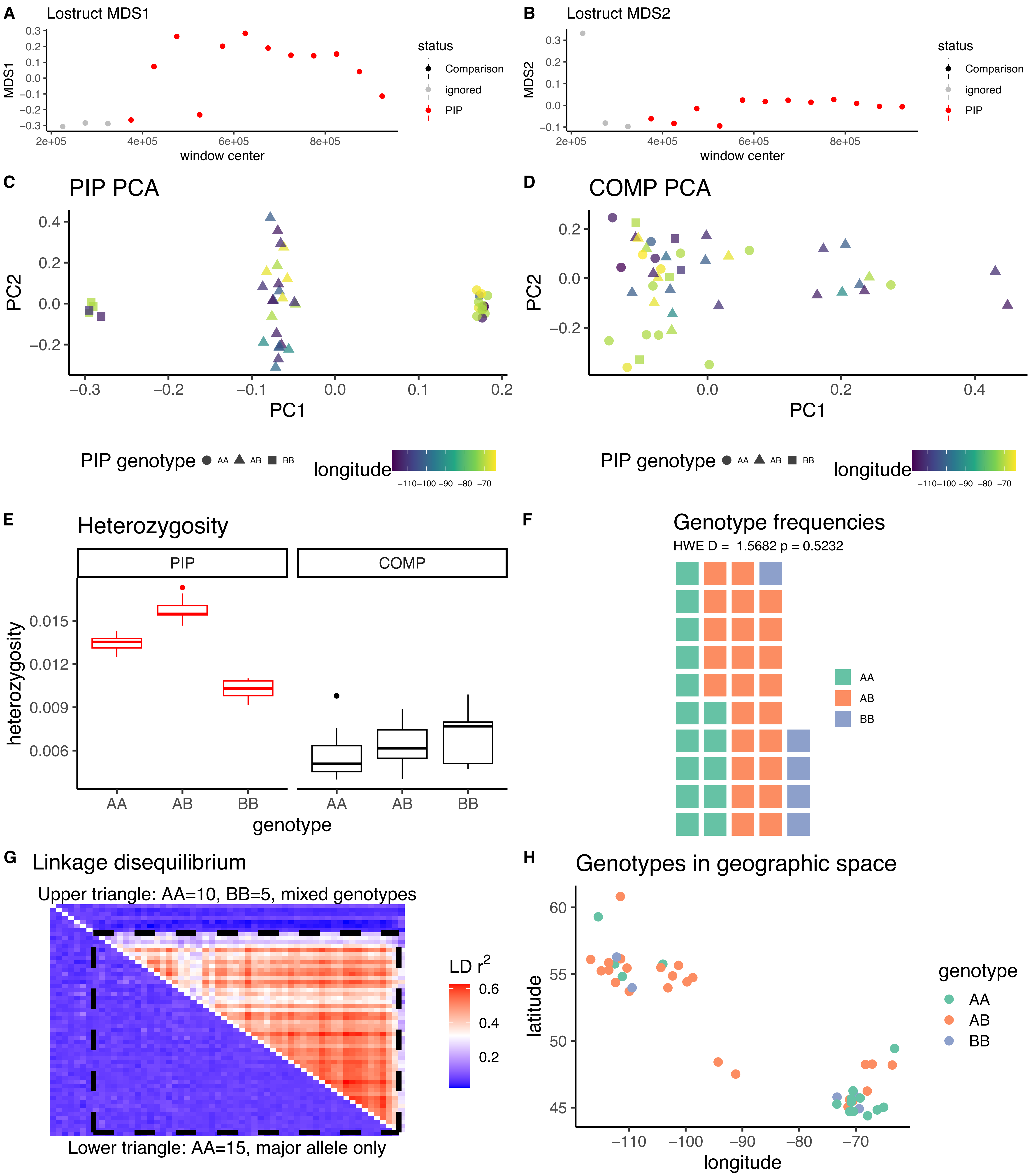

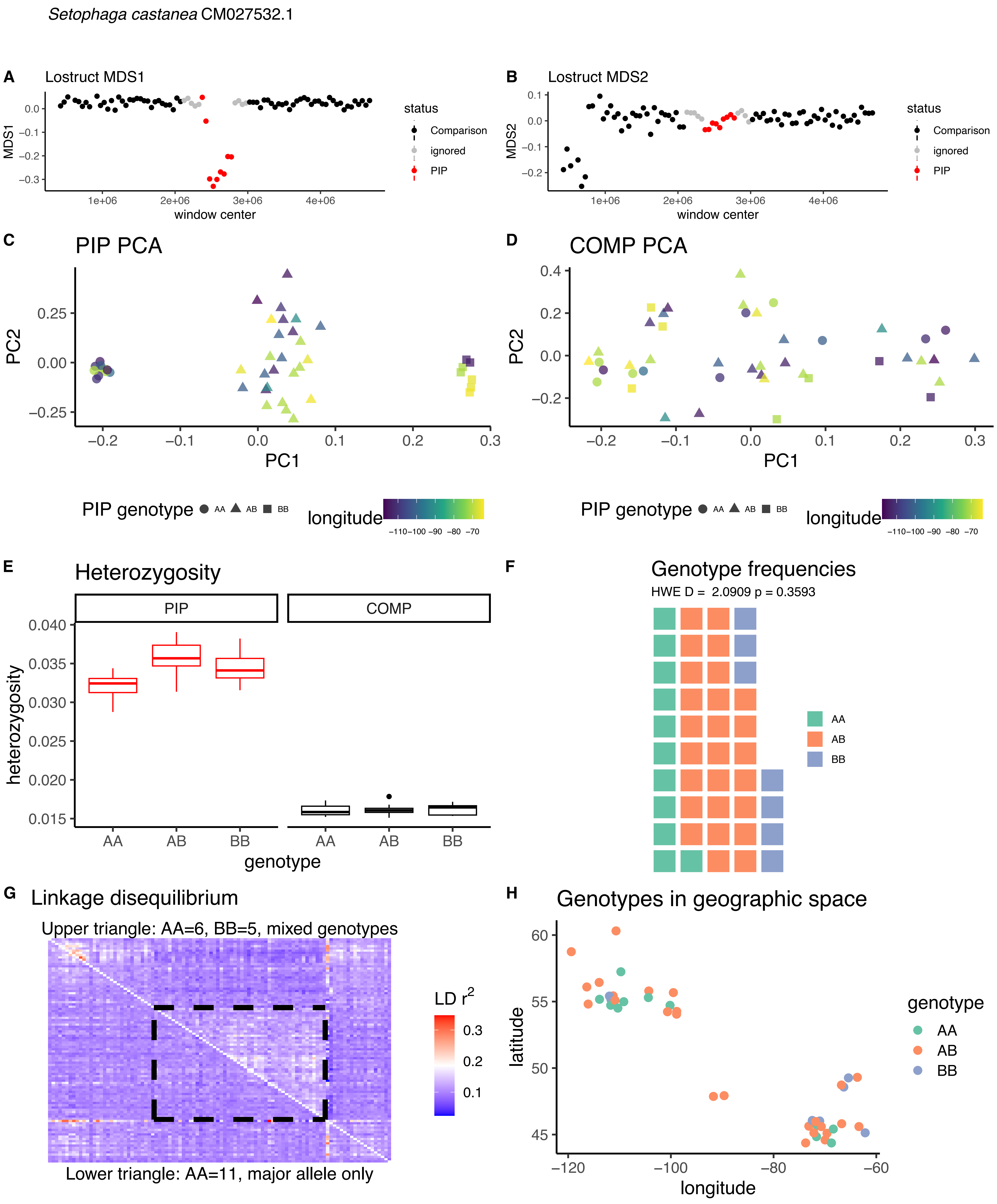

*Setophaga castanea* CM027532.1
A Lostruct MDS1
MDS1
window center
status
Comparison
ignored
PIP
B Lostruct MDS2
MDS2
C PIP PCA
PC1
PC2
D COMP PCA
PIP genotype AA AB BB
longitude
E Heterozygosity
PIP
COMP
heterozygosity
genotype
F Genotype frequencies
HWE D = 2.0909 p = 0.3593
AA
AB
BB
G Linkage disequilibrium
Upper triangle: AA=6, BB=5, mixed genotypes
LD r2
Lower triangle: AA=11, major allele only
H Genotypes in geographic space
latitude
longitude
genotype

*Setophaga castanea* CM027530.1

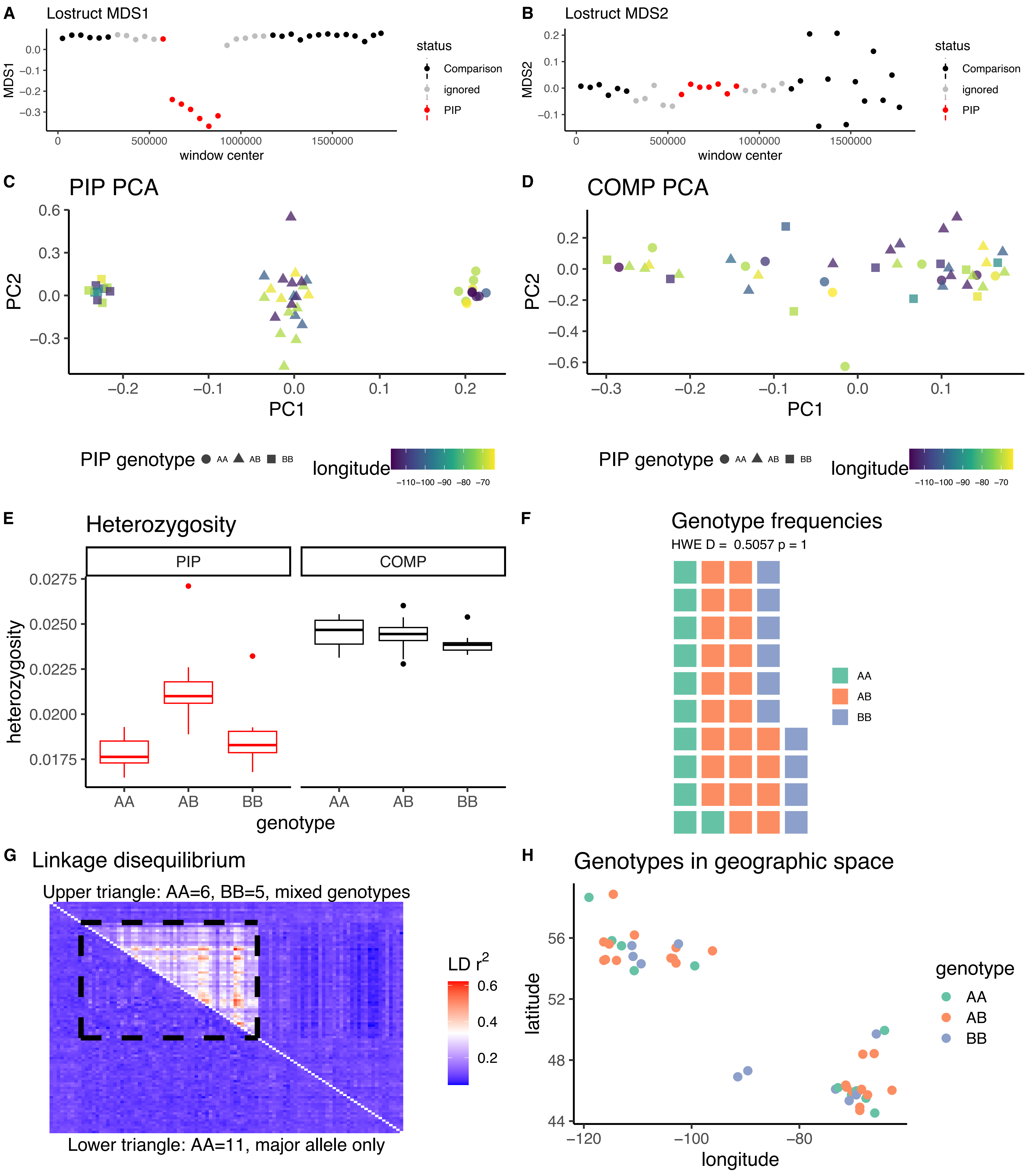

*Setophaga castanea* CM027518.1

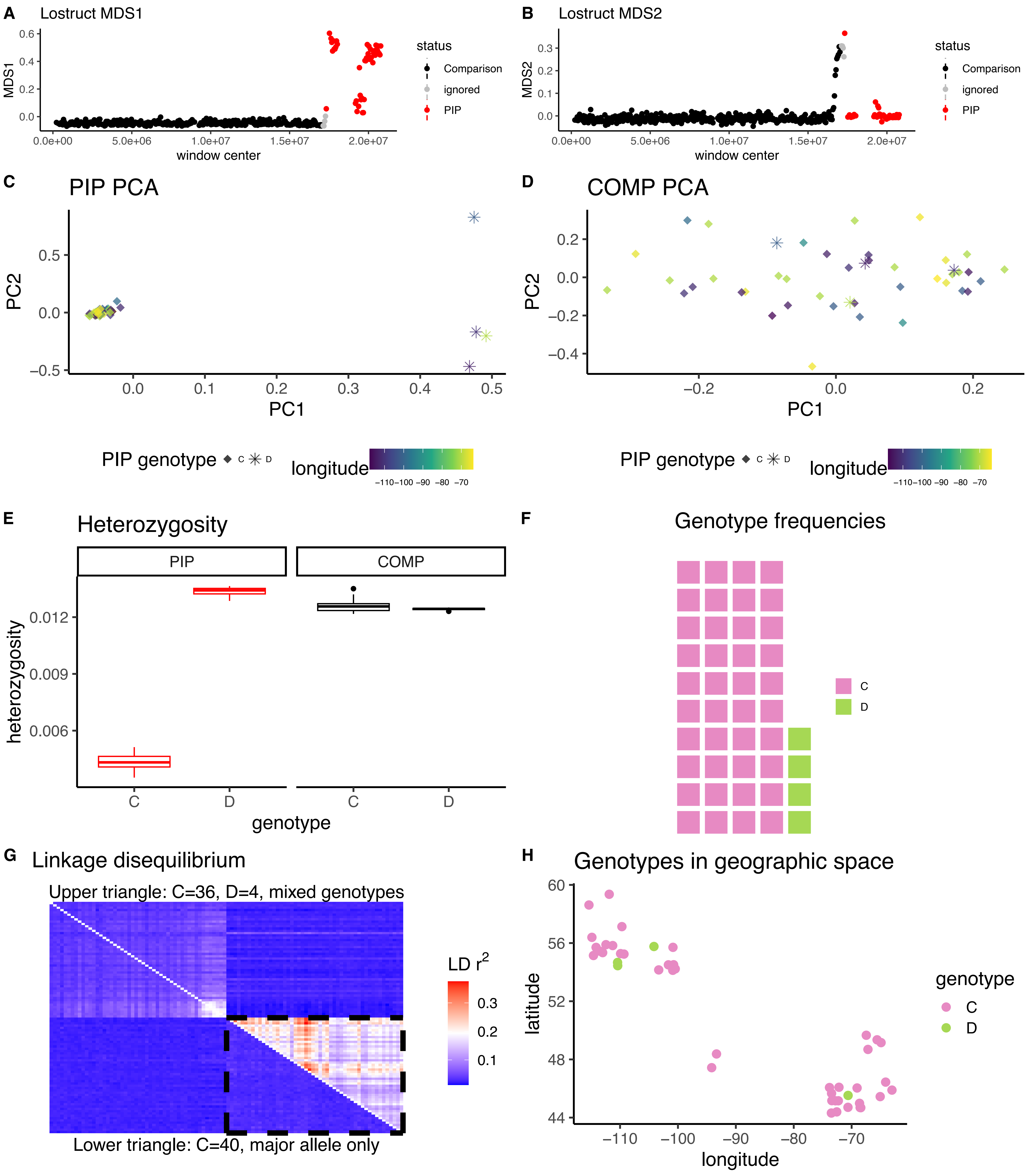

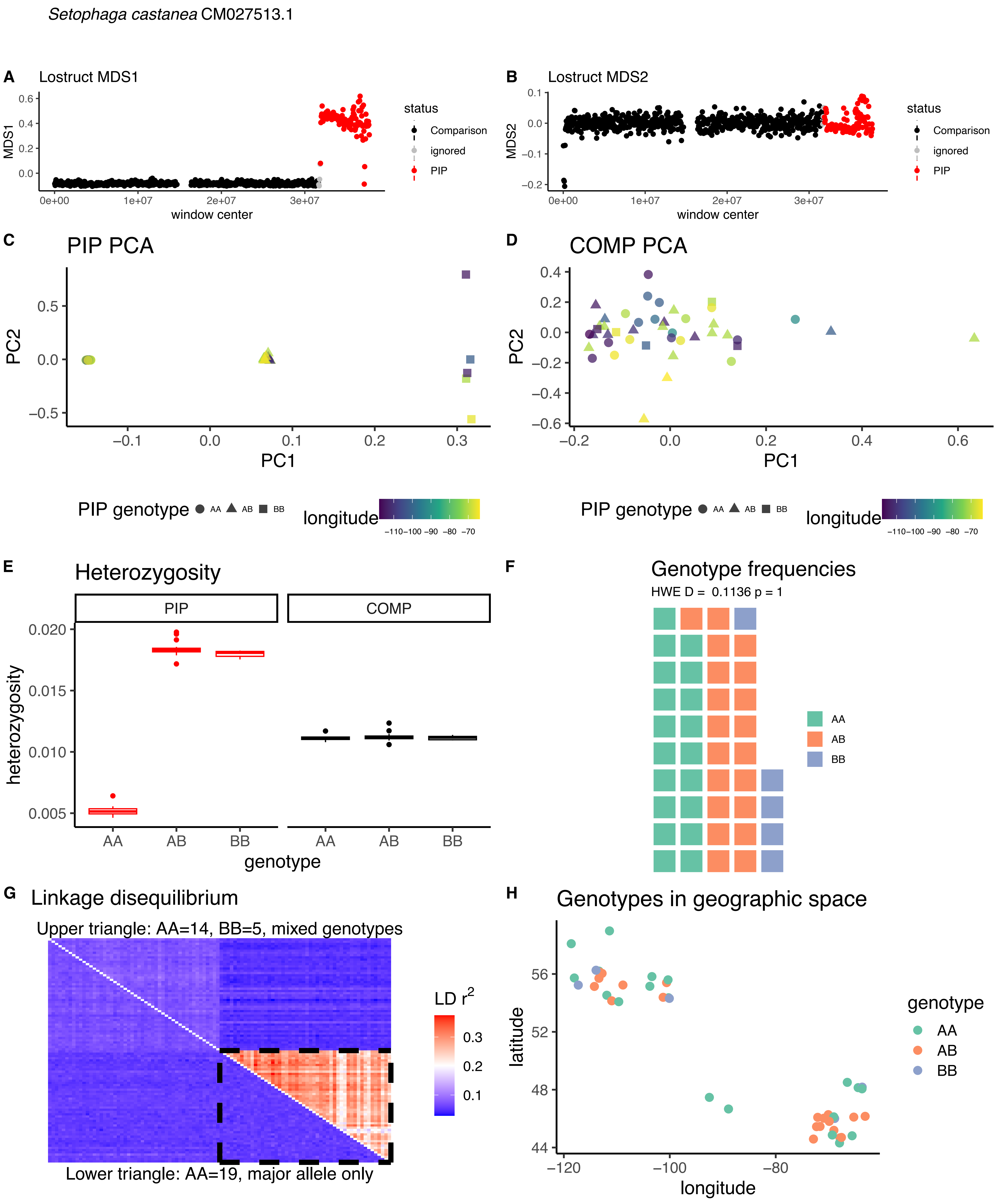

Setophaga castanea CM027513.1
A
Lostruct MDS1
MDS1
window center
status
Comparison
ignored
PIP
B
Lostruct MDS2
MDS2
window center
C
PIP PCA
PC1
PC2
D
COMP PCA
PIP genotype AA AB BB
longitude
E
Heterozygosity
PIP
COMP
heterozygosity
genotype
F
Genotype frequencies
HWE D = 0.1136 p = 1
AA
AB
BB
G
Linkage disequilibrium
Upper triangle: AA=14, BB=5, mixed genotypes
Lower triangle: AA=19, major allele only
LD r2
H
Genotypes in geographic space
latitude
longitude
genotype

*Setophaga castanea* CM027512.1

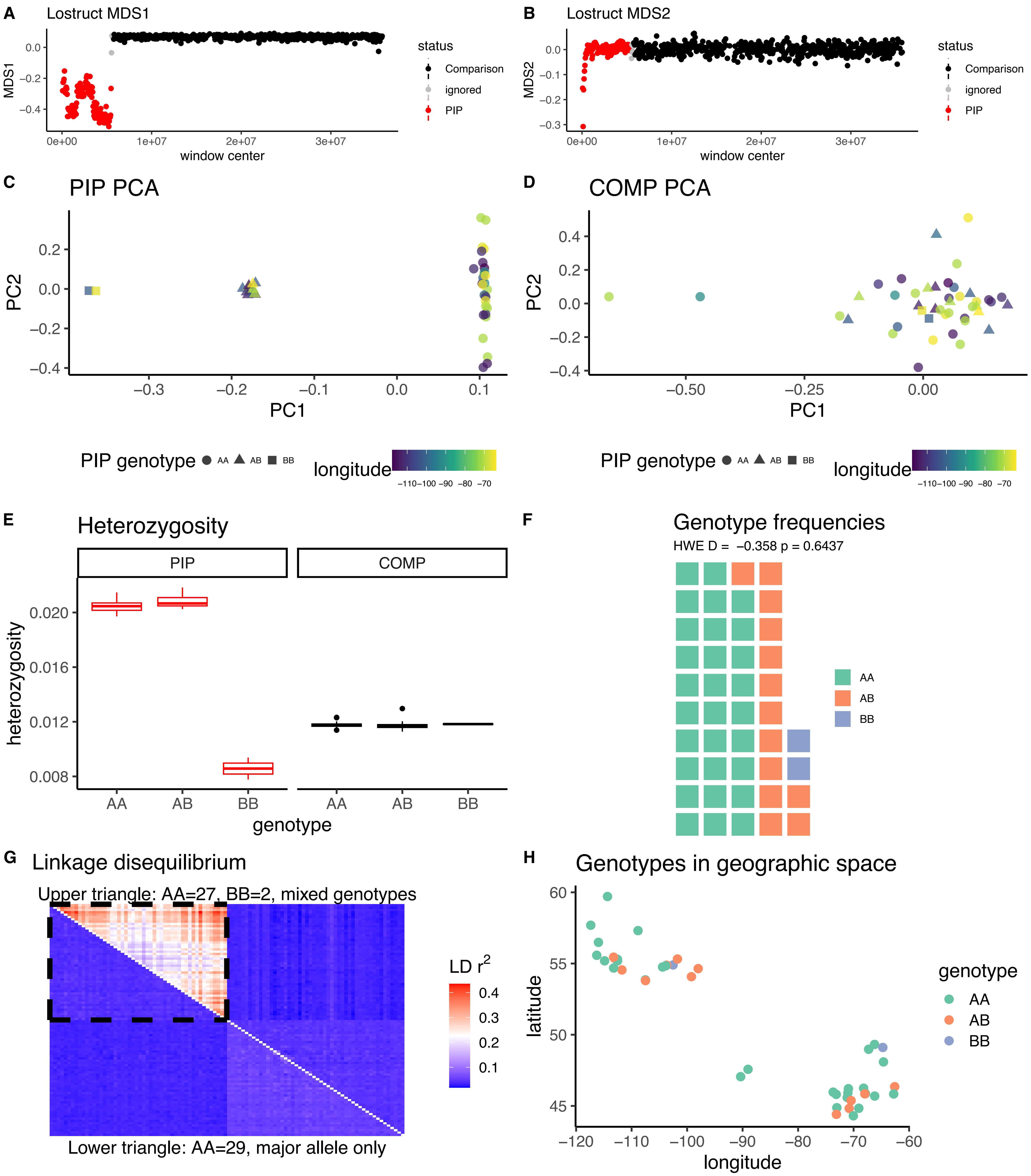

*Setophaga castanea* CM027511.1A

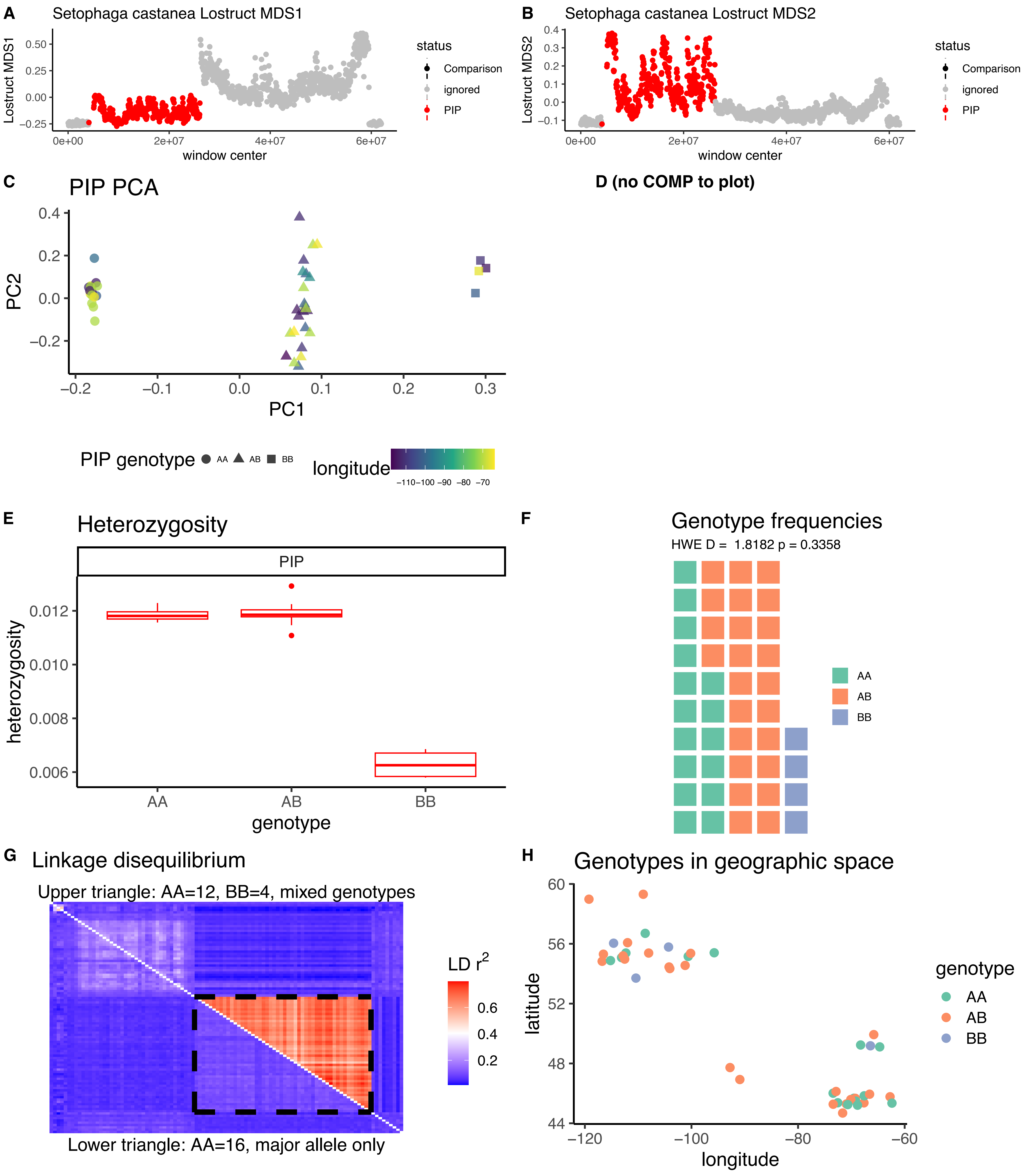

*Setophaga castanea* CM027511.1B

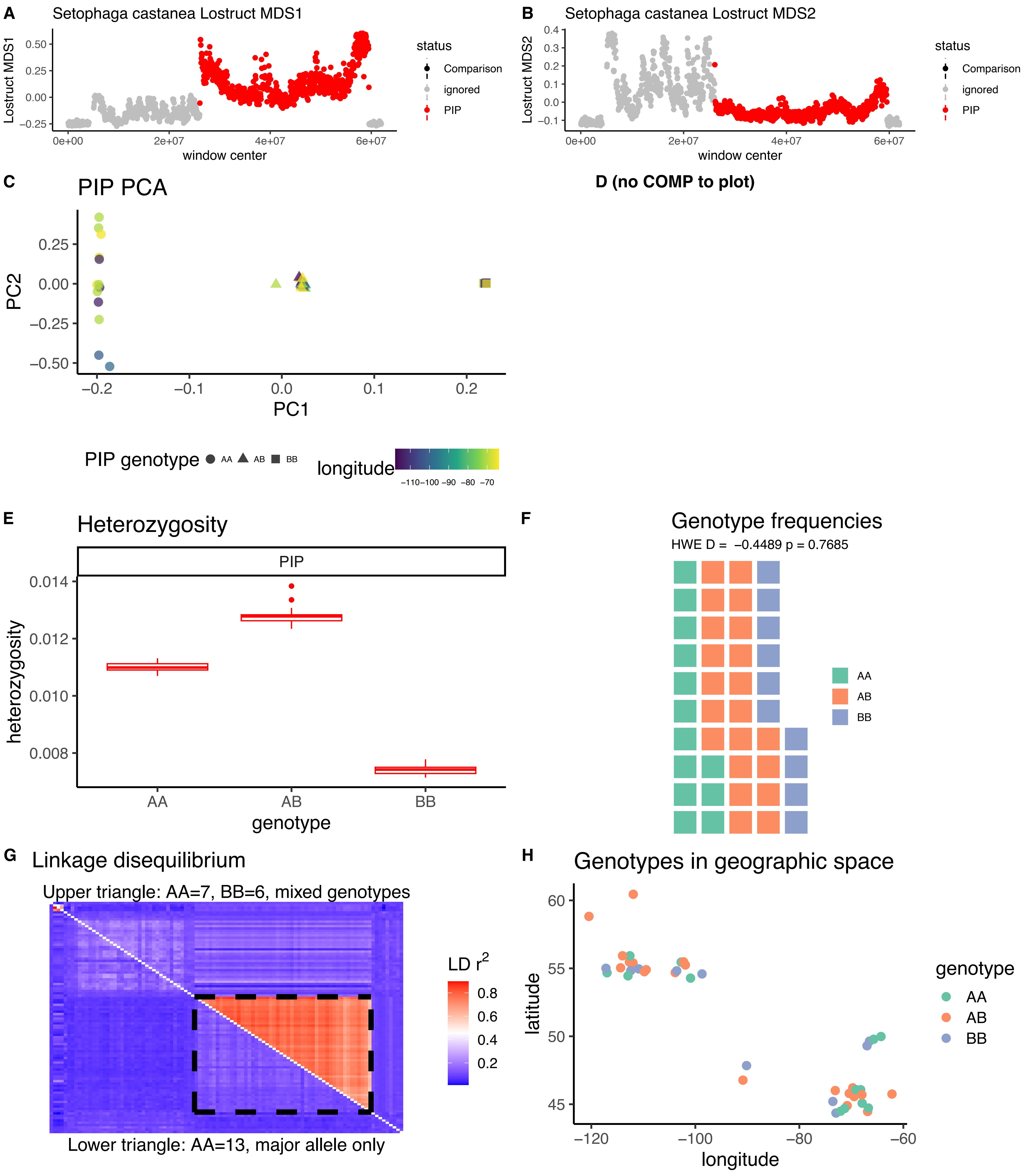

*Setophaga castanea* CM027510.1

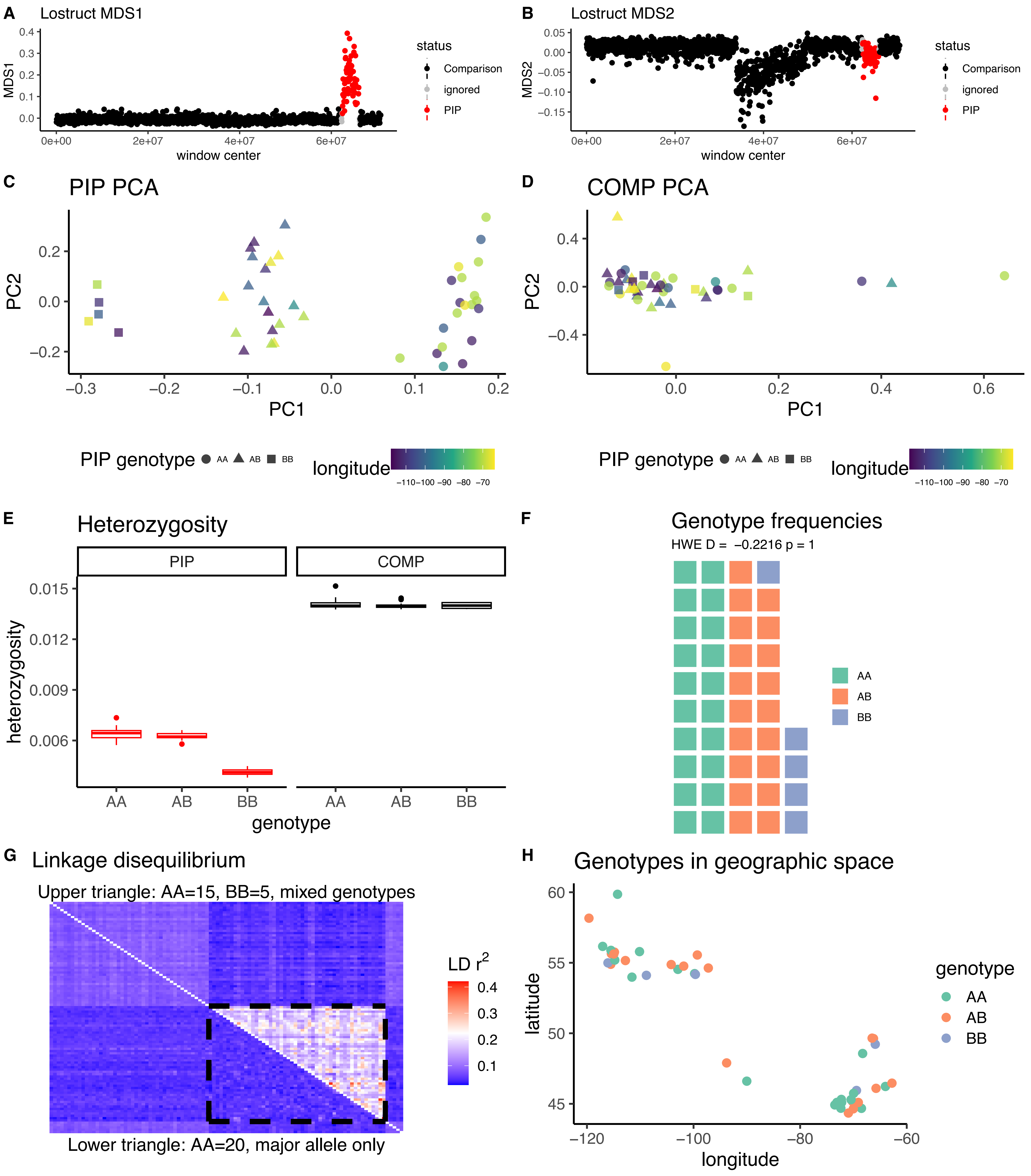

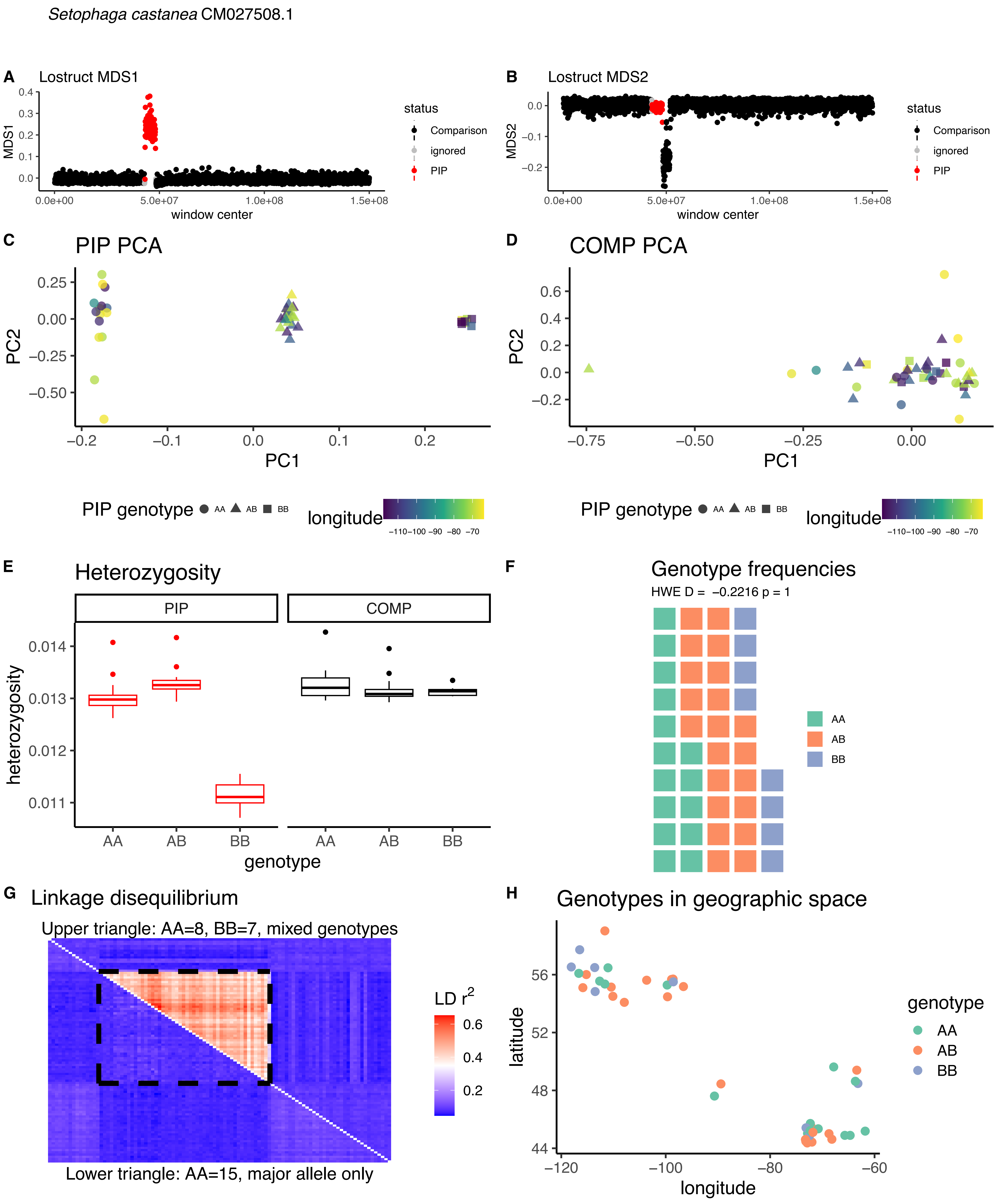
*Setophaga castanea* CM027508.1
A
Lostruct MDS1
MDS1
window center
status
Comparison
ignored
PIP
B
Lostruct MDS2
MDS2
window center
C
PIP PCA
PC1
PC2
D
COMP PCA
PIP genotype
AA
AB
BB
longitude
E
Heterozygosity
PIP
COMP
heterozygosity
genotype
F
Genotype frequencies
HWE D = −0.2216 p = 1
G
Linkage disequilibrium
Upper triangle: AA=8, BB=7, mixed genotypes
Lower triangle: AA=15, major allele only
LD r2
H
Genotypes in geographic space
latitude
genotype

*Setophaga coronata* CM027532.1

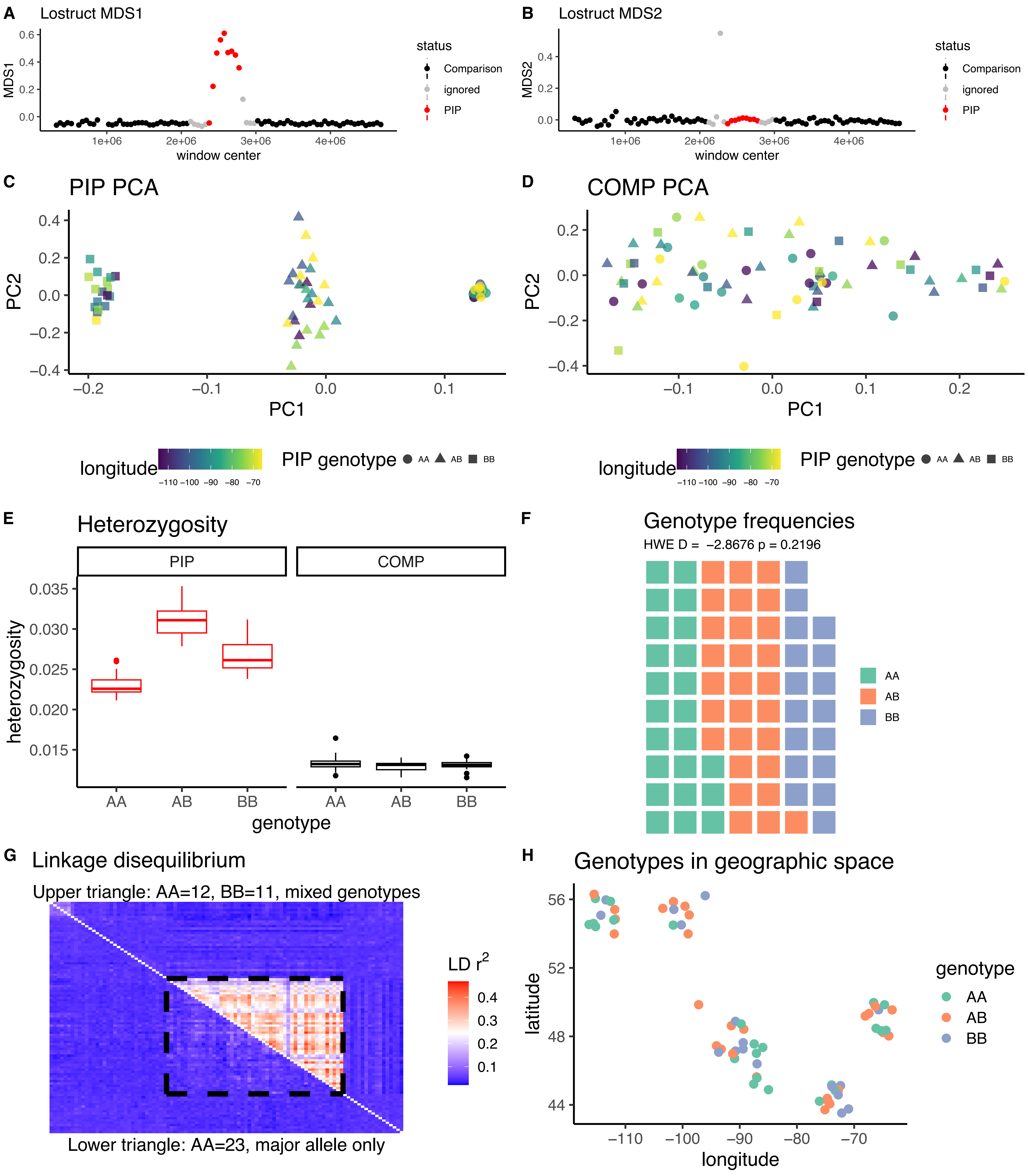

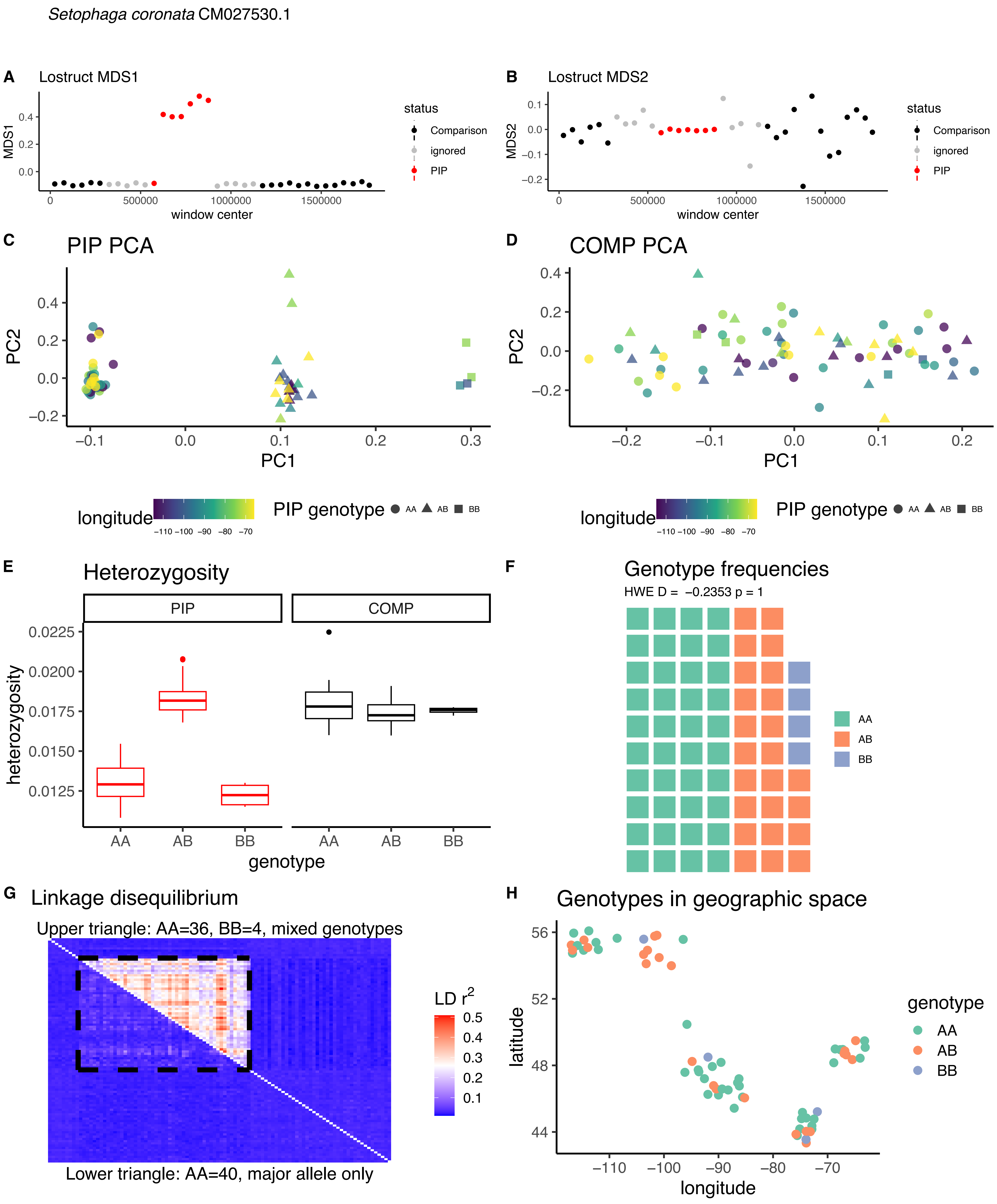

Setophaga coronata CM027530.1
A Lostruct MDS1
B Lostruct MDS2
status
Comparison
ignored
PIP
window center
MDS1
MDS2
C PIP PCA
D COMP PCA
PC1
PC2
longitude
PIP genotype AA AB BB
E Heterozygosity
PIP
COMP
heterozygosity
genotype
F Genotype frequencies
HWE D = −0.2353 p = 1
AA
AB
BB
G Linkage disequilibrium
Upper triangle: AA=36, BB=4, mixed genotypes
Lower triangle: AA=40, major allele only
LD r2
H Genotypes in geographic space
latitude
longitude
genotype

*Setophaga coronata* CM027510.1A

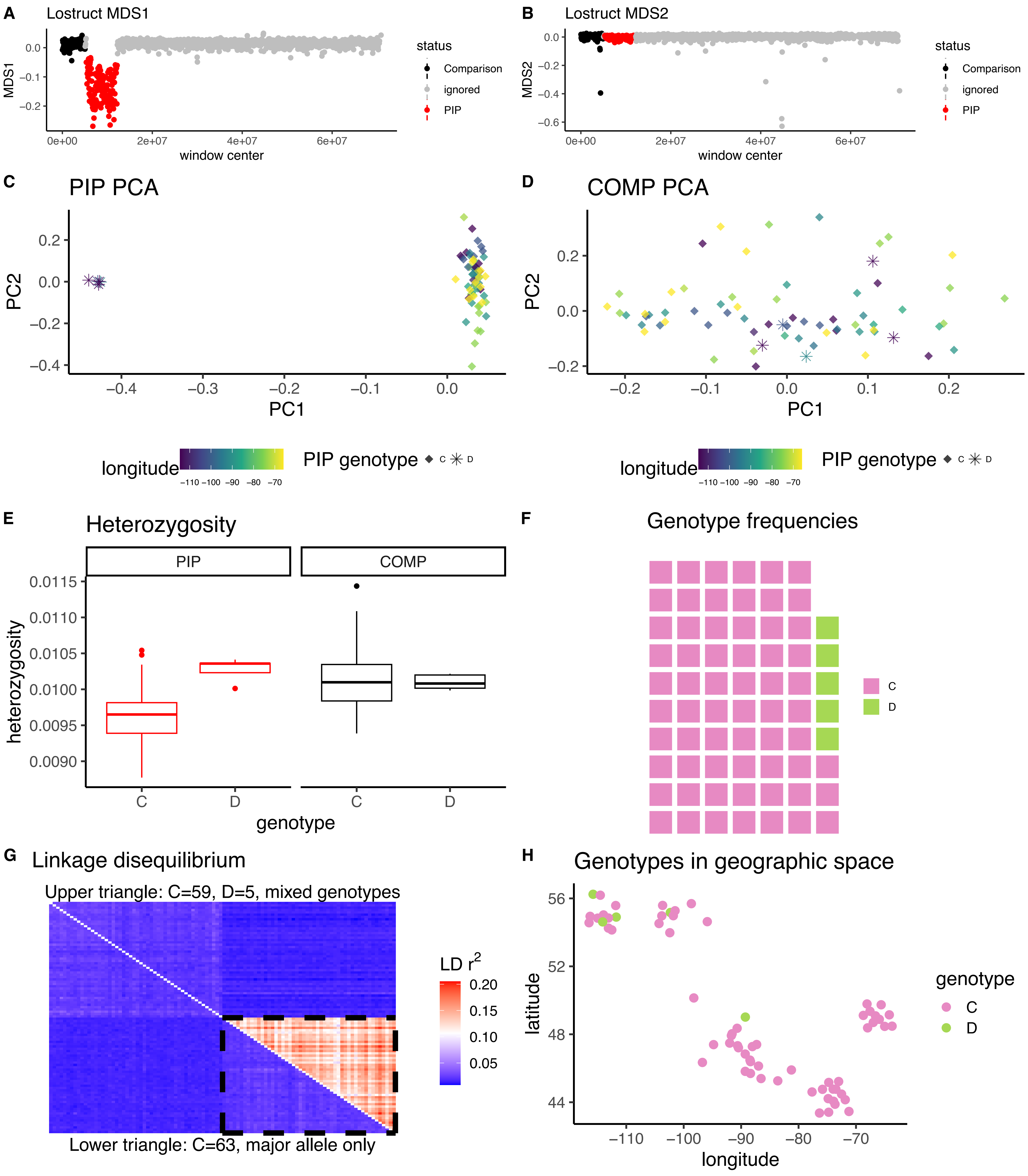

*Setophaga coronata* CM027510.1B

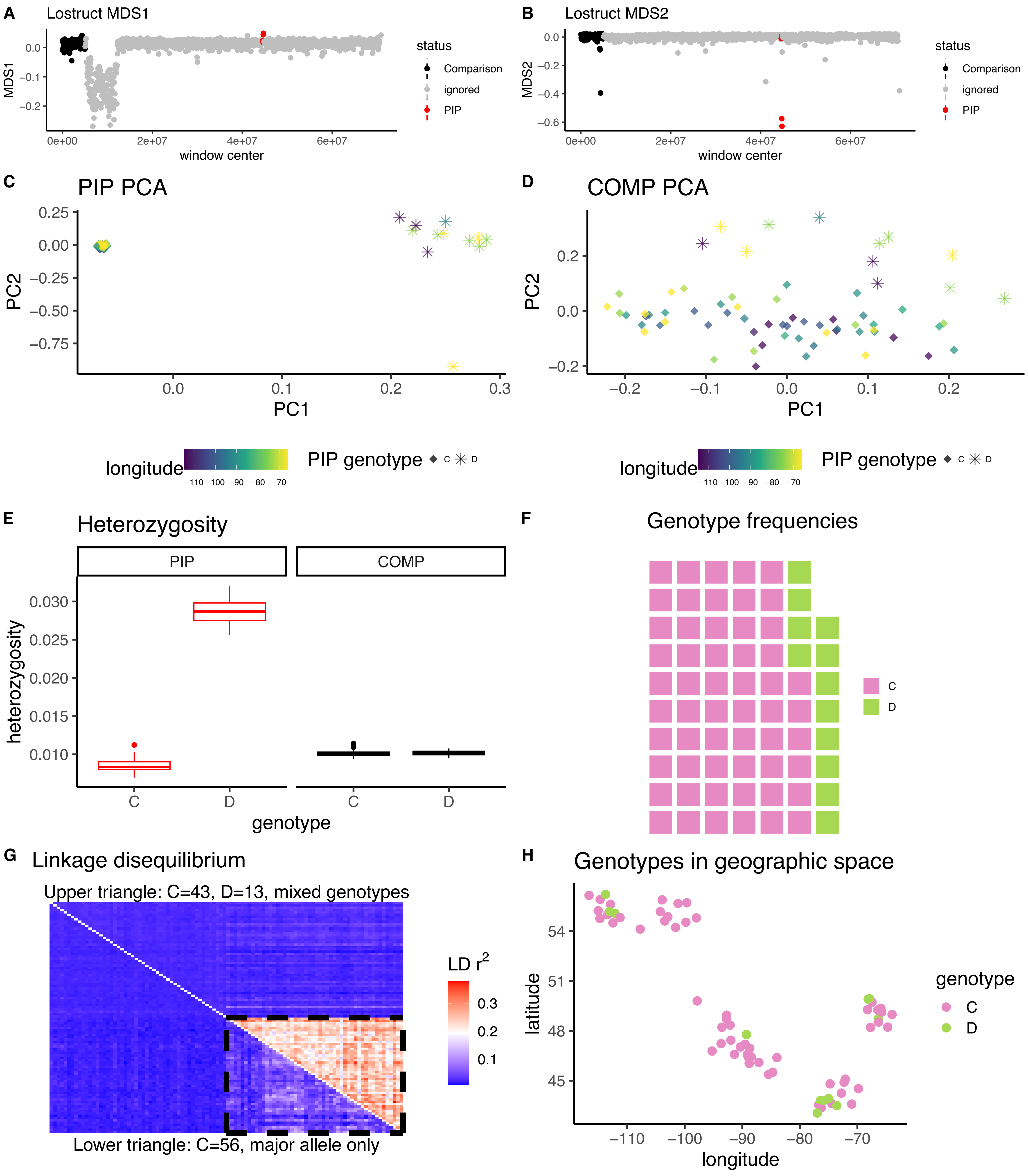

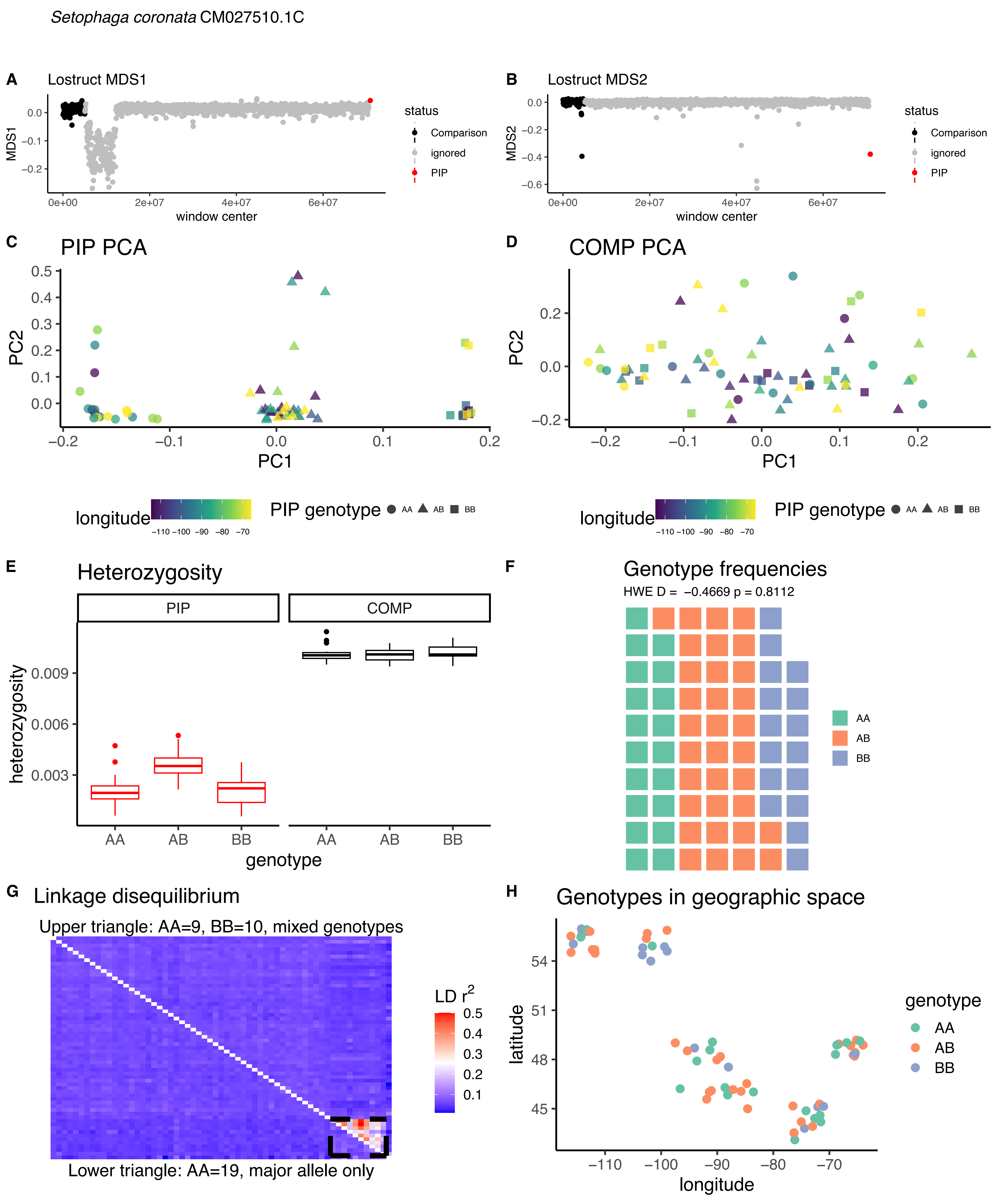

Setophaga coronata CM027510.1C
A
Lostruct MDS1
MDS1
window center
status
Comparison
ignored
PIP
B
Lostruct MDS2
MDS2
C
PIP PCA
PC1
PC2
D
COMP PCA
longitude
PIP genotype
AA
AB
BB
E
Heterozygosity
PIP
COMP
heterozygosity
genotype
F
Genotype frequencies
HWE D = −0.4669 p = 0.8112
G
Linkage disequilibrium
Upper triangle: AA=9, BB=10, mixed genotypes
LD r2
Lower triangle: AA=19, major allele only
H
Genotypes in geographic space
latitude
longitude
genotype

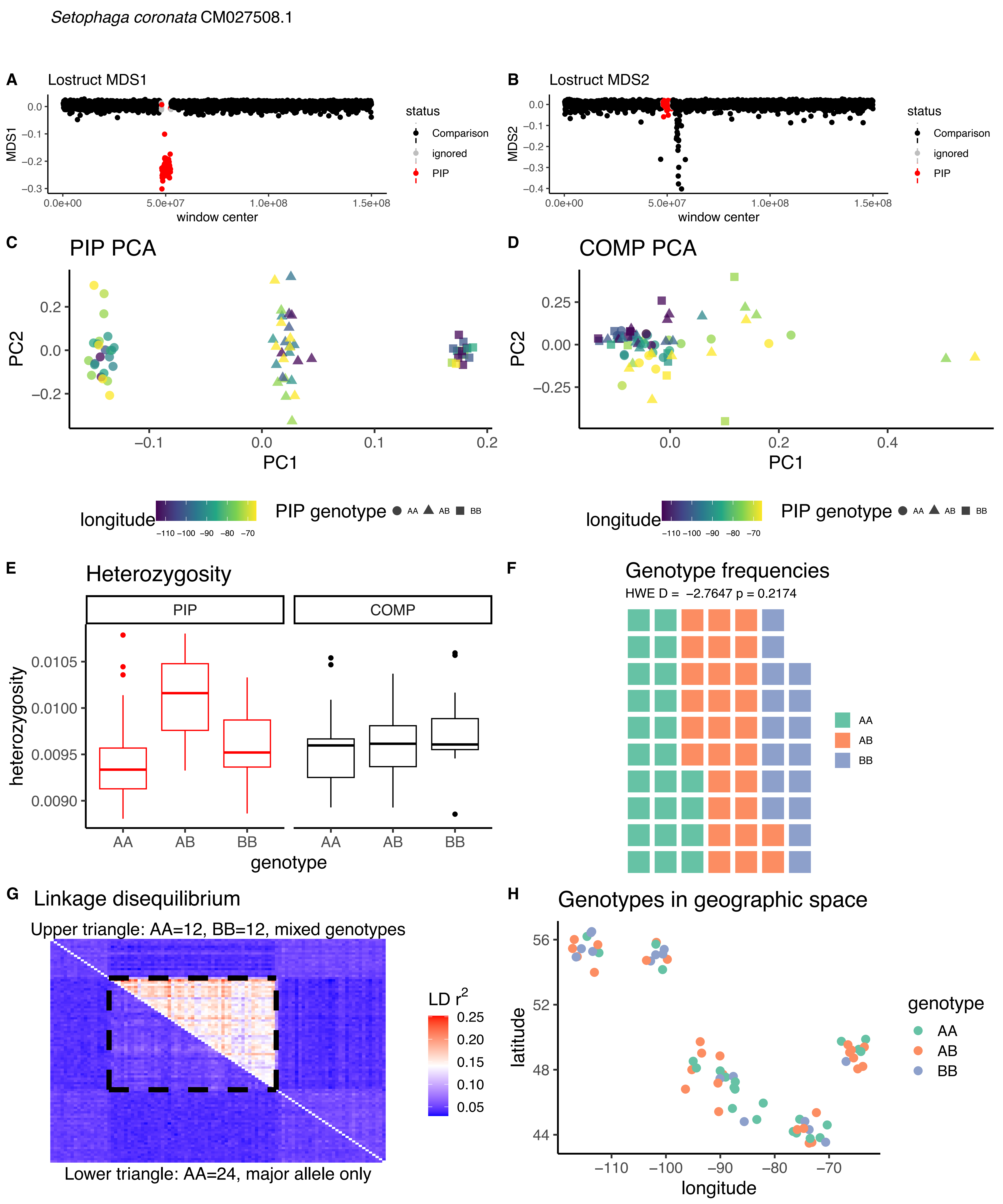

*Setophaga coronata* CM027508.1
A
Lostruct MDS1
MDS1
window center
status
Comparison
ignored
PIP
B
Lostruct MDS2
MDS2
C
PIP PCA
PC1
PC2
D
COMP PCA
longitude
PIP genotype
AA
AB
BB
E
Heterozygosity
PIP
COMP
heterozygosity
genotype
F
Genotype frequencies
HWE D = −2.7647 p = 0.2174
G
Linkage disequilibrium
Upper triangle: AA=12, BB=12, mixed genotypes
LD $r^2$
Lower triangle: AA=24, major allele only
H
Genotypes in geographic space
latitude
longitude
genotype

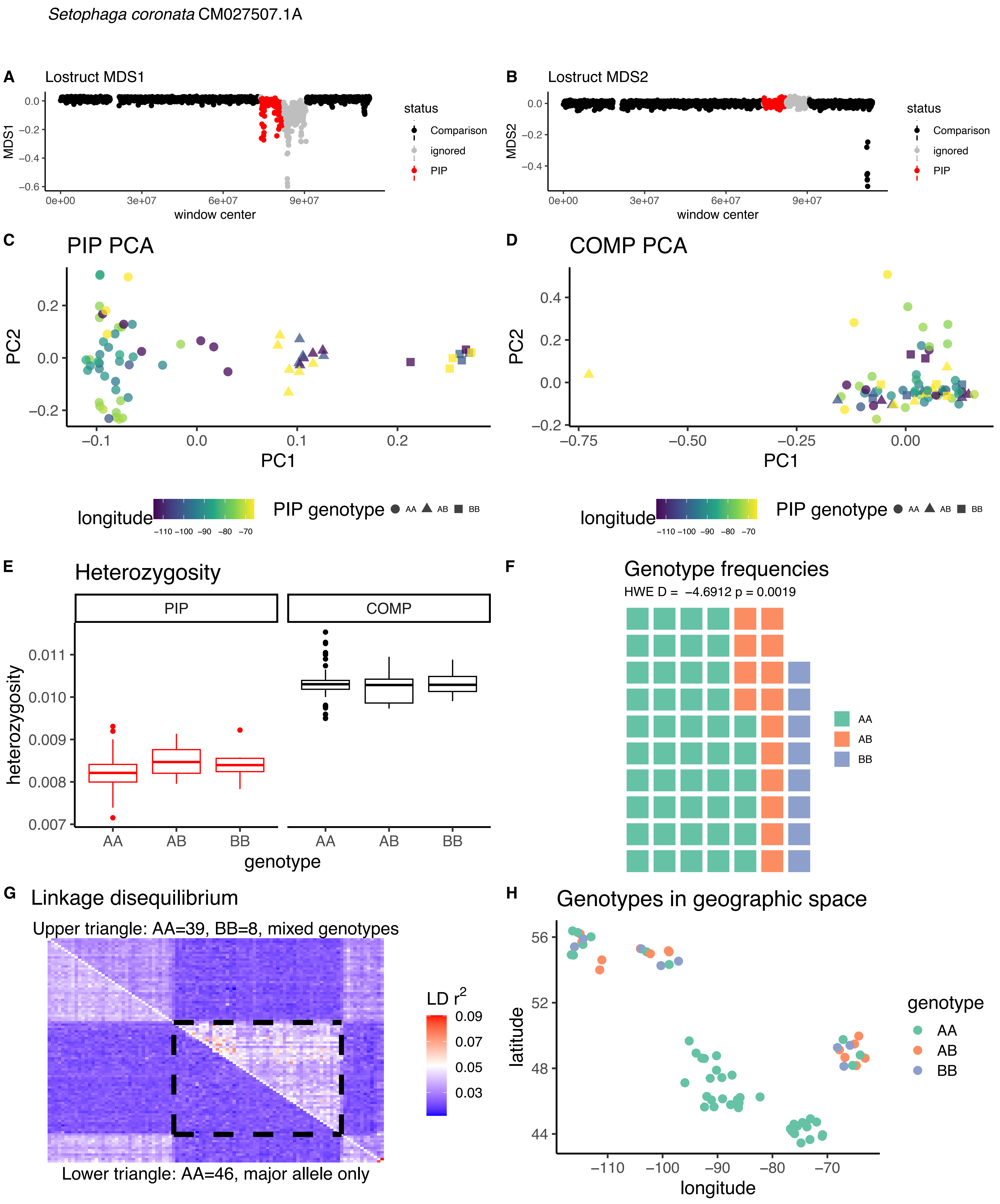

*Setophaga coronata* CM027507.1A
A
Lostruct MDS1
MDS1
window center
status
Comparison
ignored
PIP
B
Lostruct MDS2
MDS2
C
PIP PCA
PC1
PC2
D
COMP PCA
longitude
PIP genotype
AA
AB
BB
E
Heterozygosity
PIP
COMP
heterozygosity
genotype
F
Genotype frequencies
HWE D = −4.6912 p = 0.0019
G
Linkage disequilibrium
Upper triangle: AA=39, BB=8, mixed genotypes
Lower triangle: AA=46, major allele only
LD r²
H
Genotypes in geographic space
latitude
genotype

*Setophaga coronata* CM027507.1B

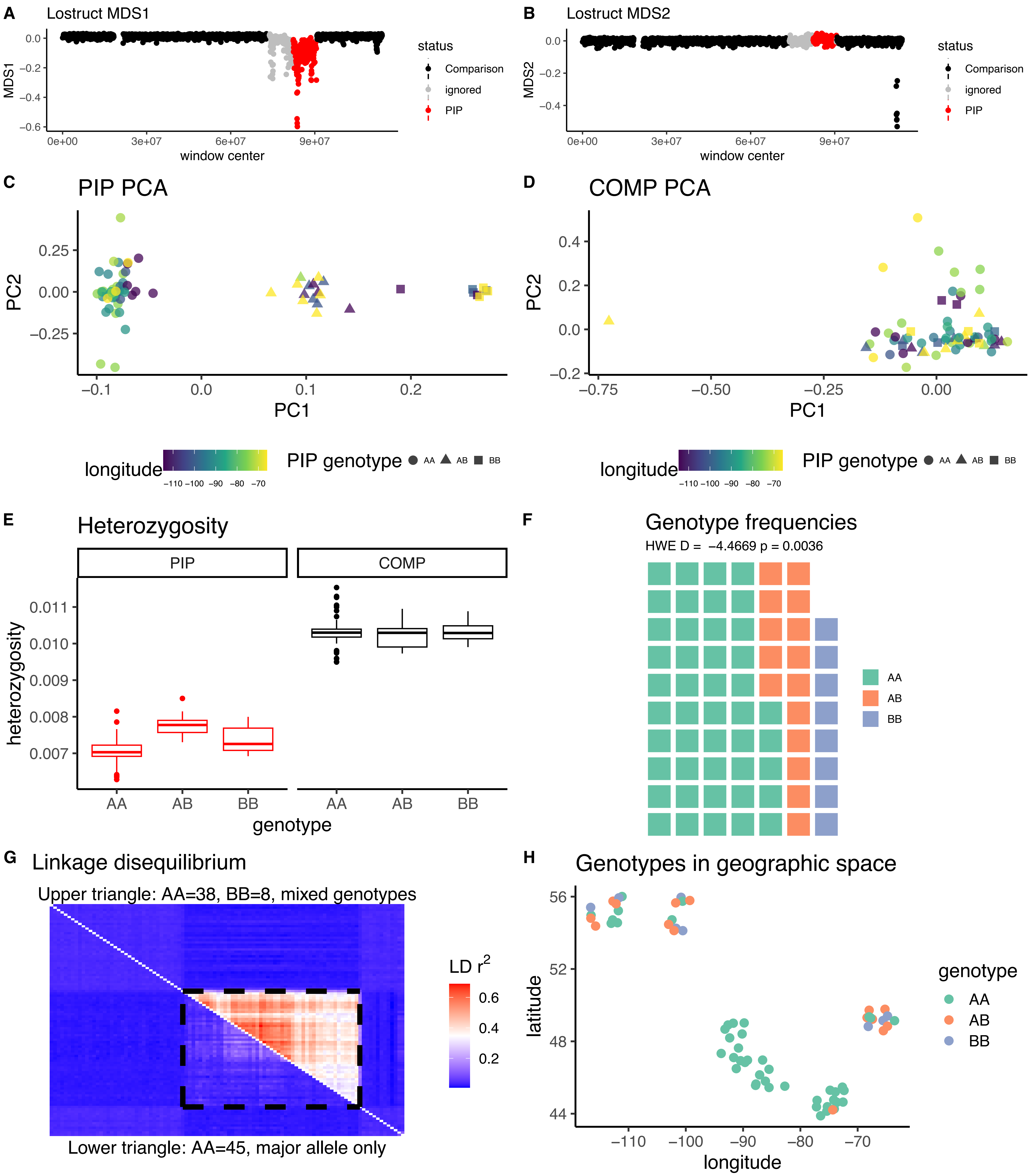

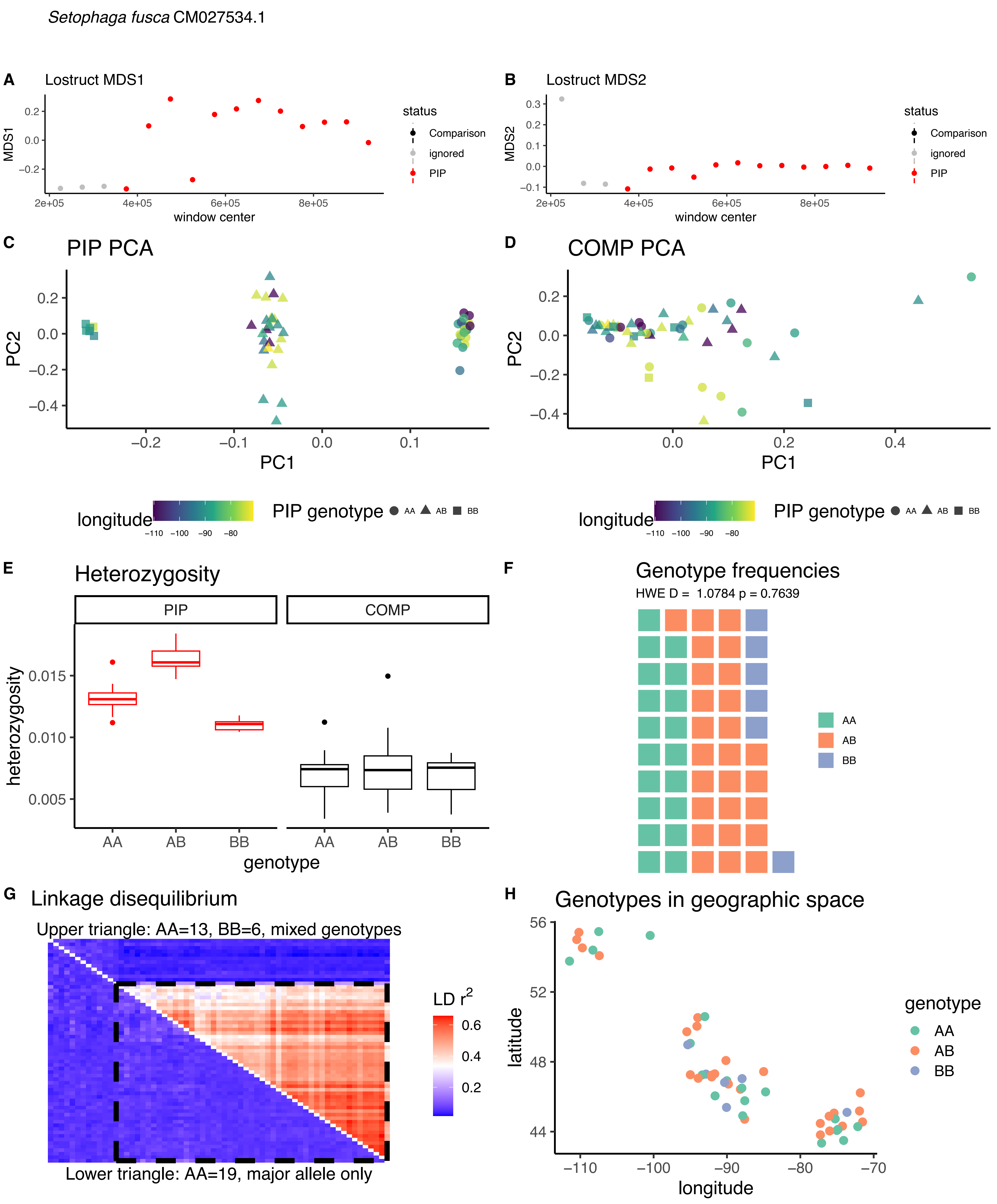

*Setophaga fusca* CM027534.1
A Lostruct MDS1
MDS1
window center
status
Comparison
ignored
PIP
B Lostruct MDS2
MDS2
window center
C PIP PCA
PC1
PC2
D COMP PCA
longitude
PIP genotype AA AB BB
E Heterozygosity
PIP
COMP
heterozygosity
genotype
F Genotype frequencies
HWE D = 1.0784 p = 0.7639
AA
AB
BB
G Linkage disequilibrium
Upper triangle: AA=13, BB=6, mixed genotypes
Lower triangle: AA=19, major allele only
LD r²
H Genotypes in geographic space
latitude
longitude
genotype

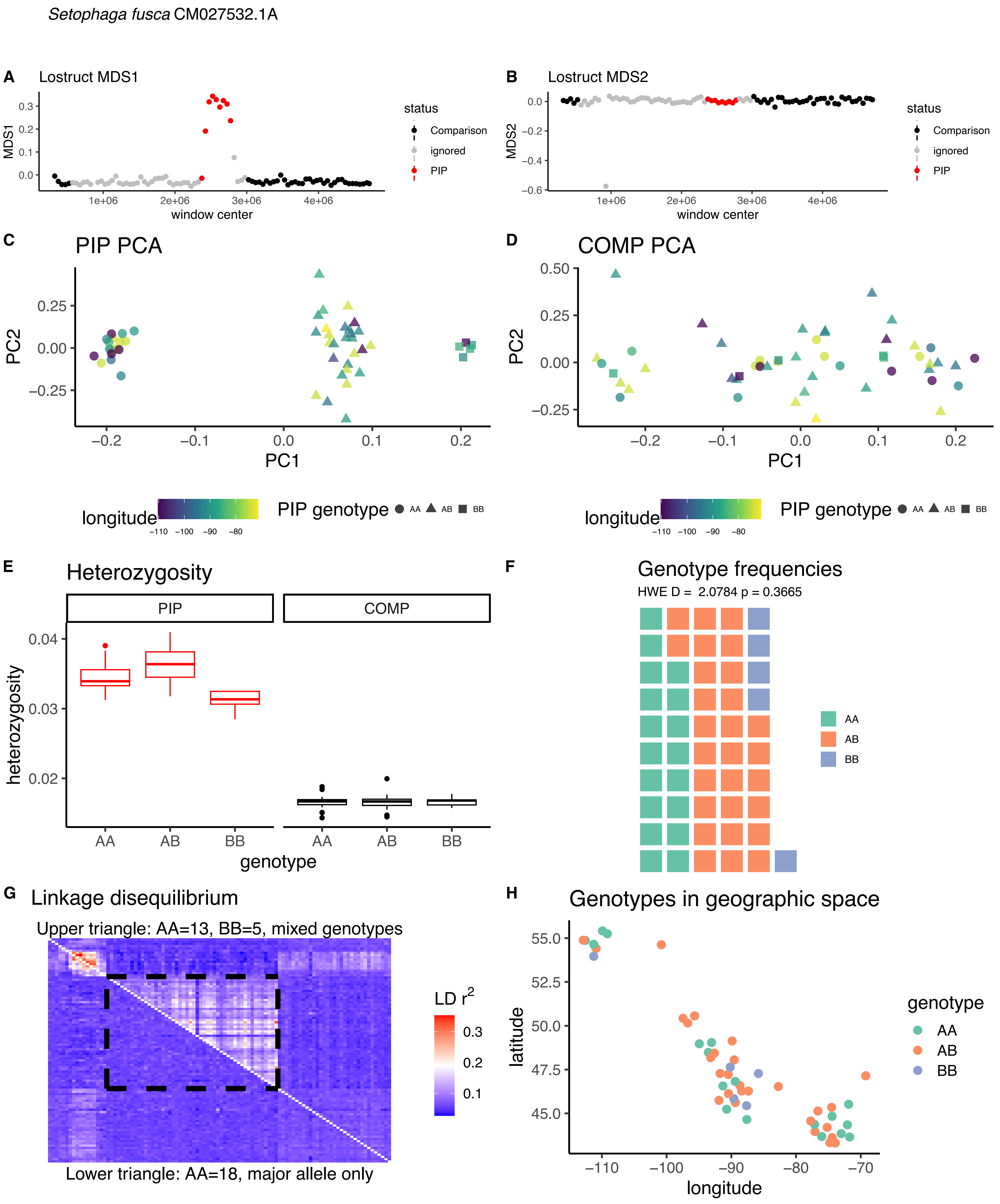

Setophaga fusca CM027532.1A
A Lostruct MDS1
MDS1
window center
status
Comparison
ignored
PIP
B Lostruct MDS2
MDS2
window center
C PIP PCA
PC1
PC2
D COMP PCA
longitude
PIP genotype AA AB BB
E Heterozygosity
PIP
COMP
heterozygosity
genotype
F Genotype frequencies
HWE D = 2.0784 p = 0.3665
AA
AB
BB
G Linkage disequilibrium
Upper triangle: AA=13, BB=5, mixed genotypes
LD r2
Lower triangle: AA=18, major allele only
H Genotypes in geographic space
latitude
longitude
genotype

*Setophaga fusca* CM027532.1B

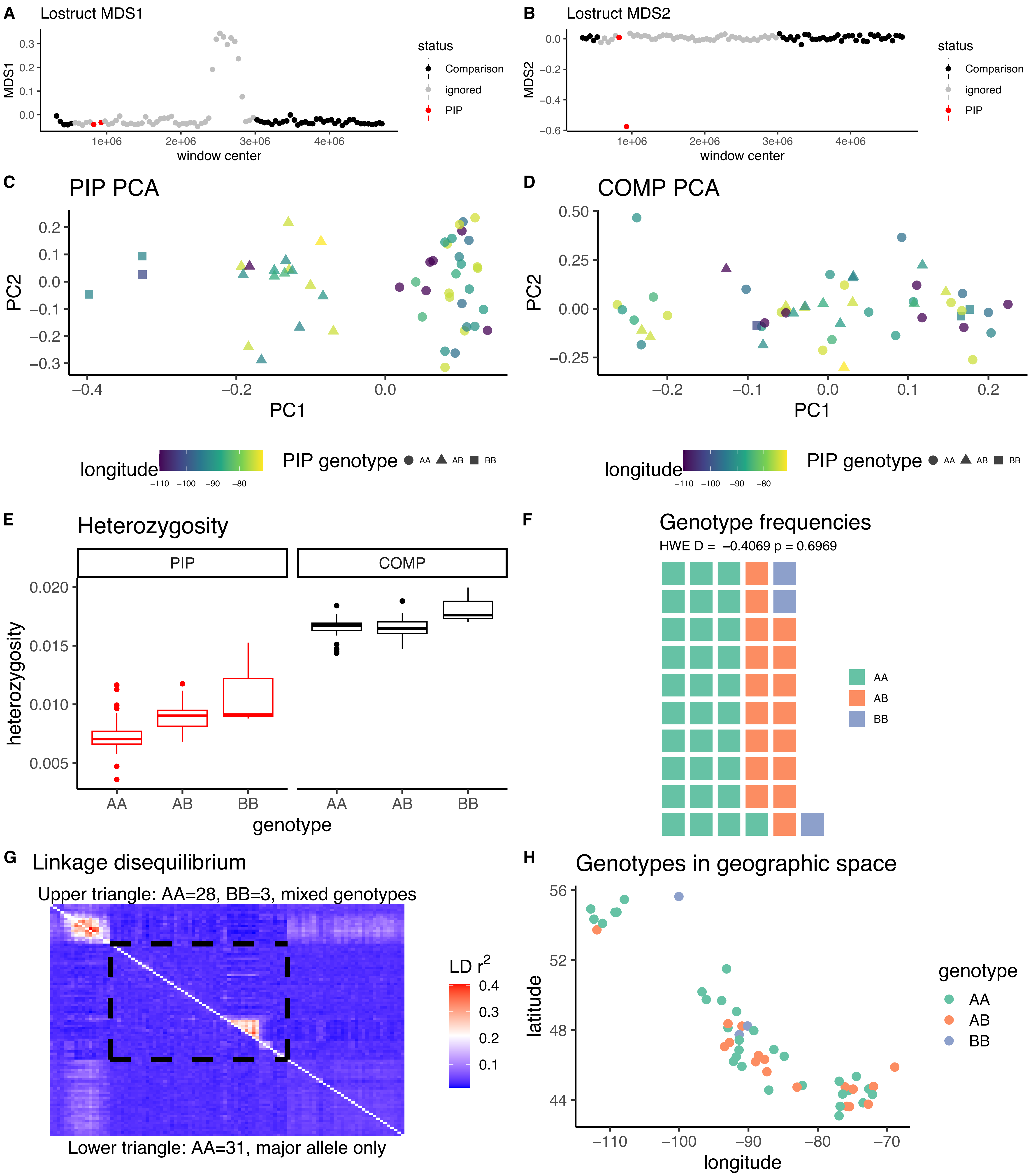

*Setophaga fusca* CM027530.1

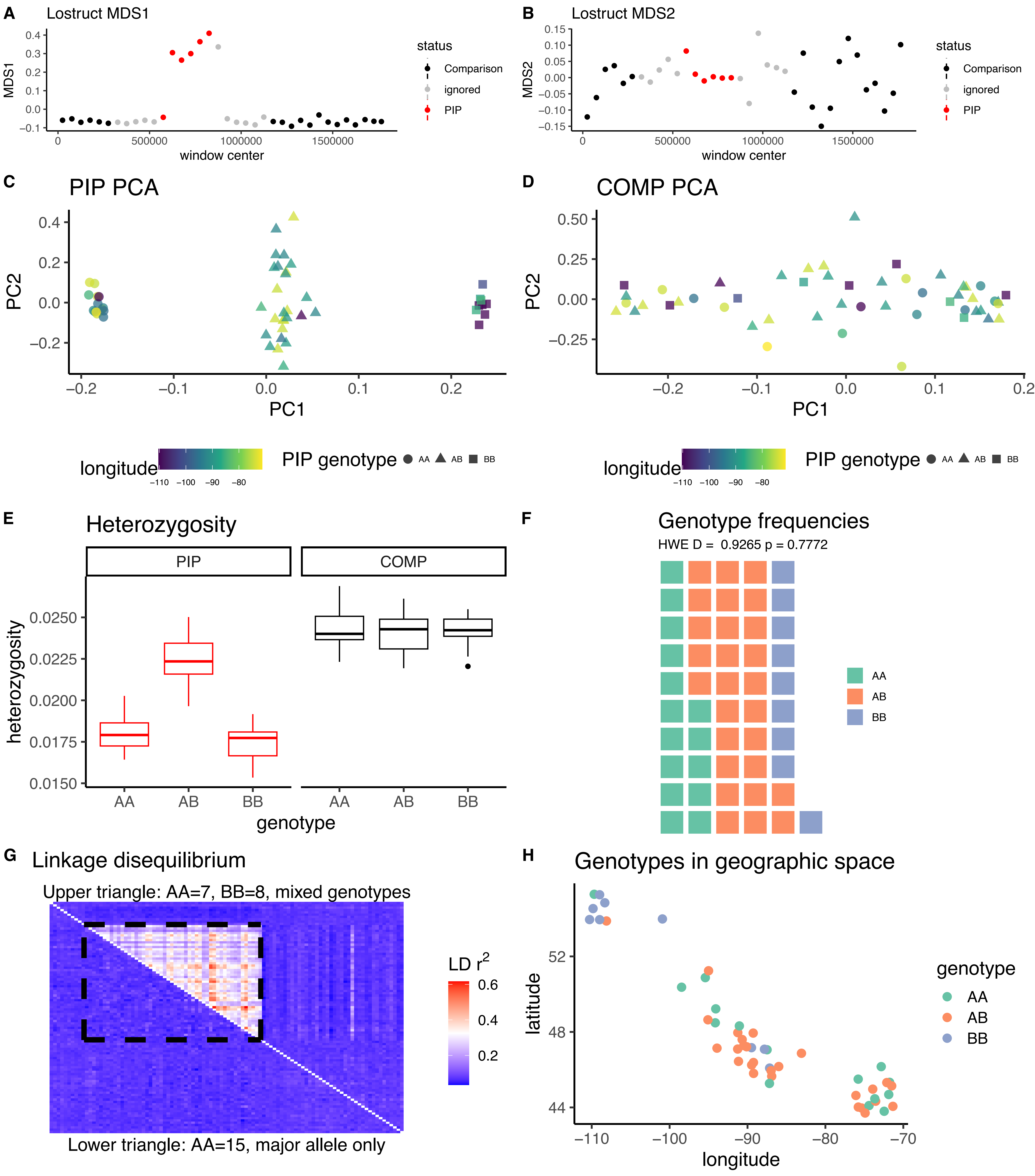

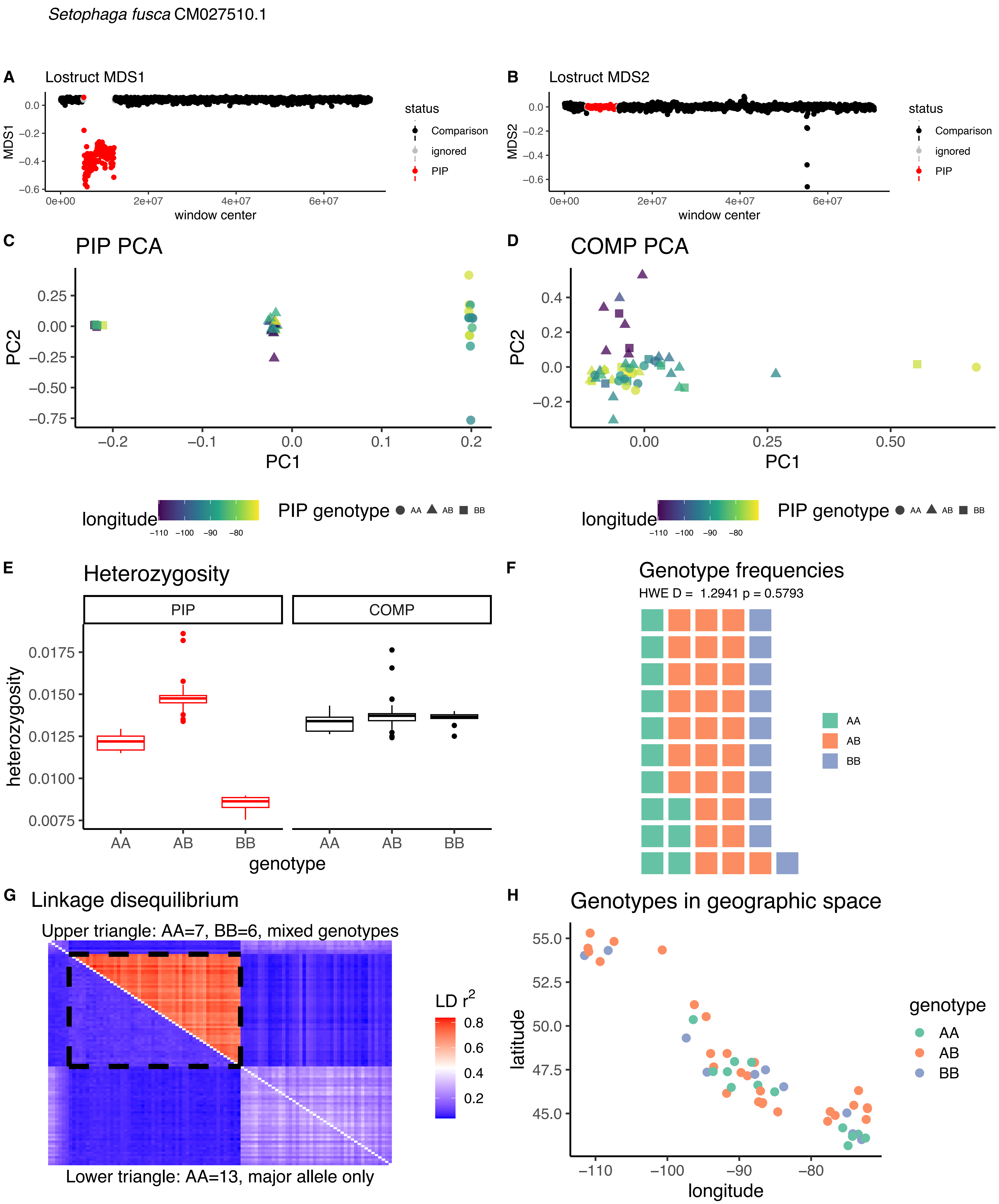

*Setophaga fusca* CM027510.1
A
Lostruct MDS1
MDS1
window center
status
Comparison
ignored
PIP
B
Lostruct MDS2
MDS2
window center
C
PIP PCA
PC2
PC1
D
COMP PCA
longitude
PIP genotype
AA
AB
BB
E
Heterozygosity
PIP
COMP
heterozygosity
genotype
F
Genotype frequencies
HWE D = 1.2941 p = 0.5793
G
Linkage disequilibrium
Upper triangle: AA=7, BB=6, mixed genotypes
Lower triangle: AA=13, major allele only
LD r²
H
Genotypes in geographic space
latitude
longitude
genotype

*Setophaga fusca* CM027508.1

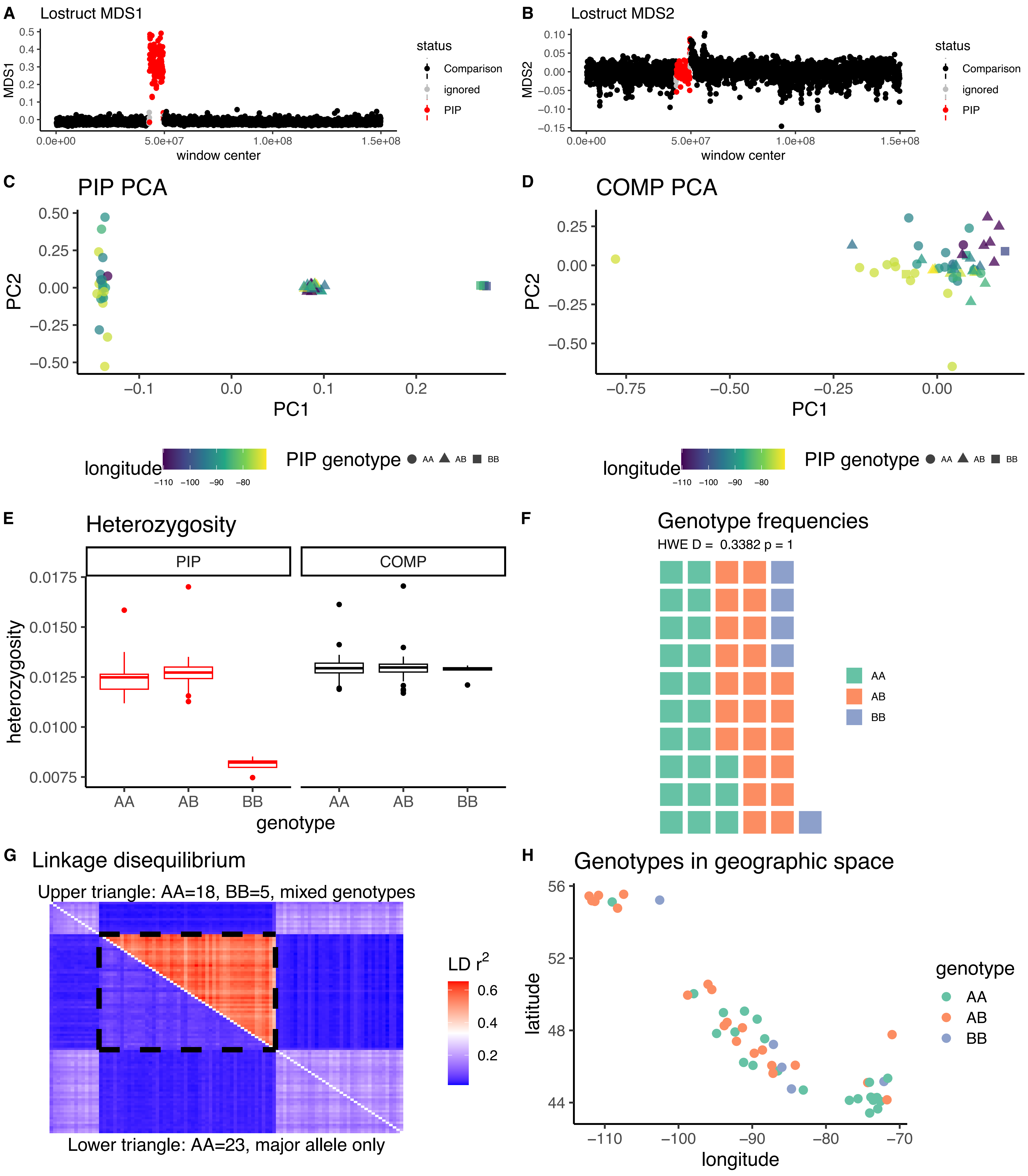

*Setophaga magnolia* CM027534.1

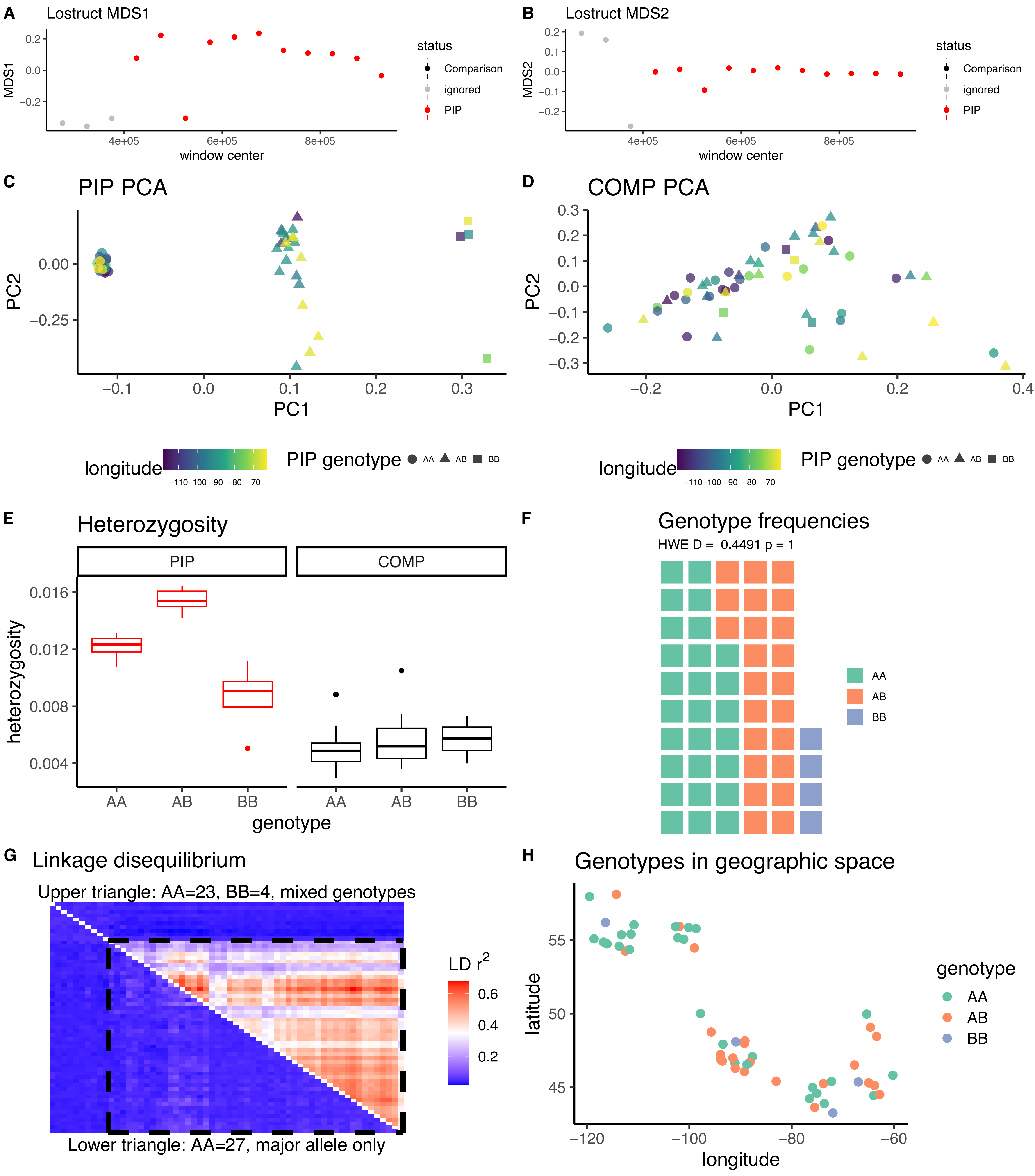

*Setophaga magnolia* CM027532.1

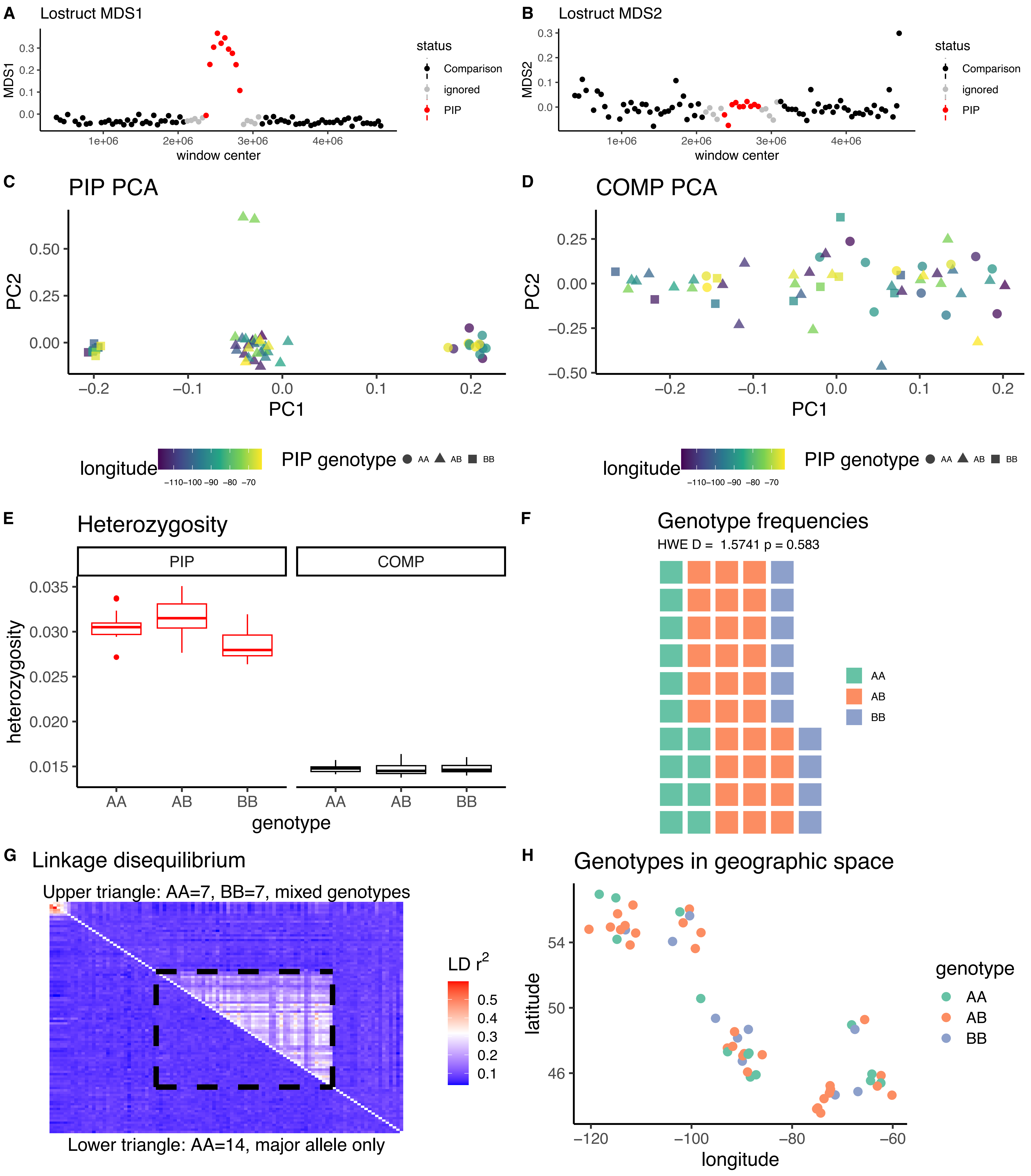

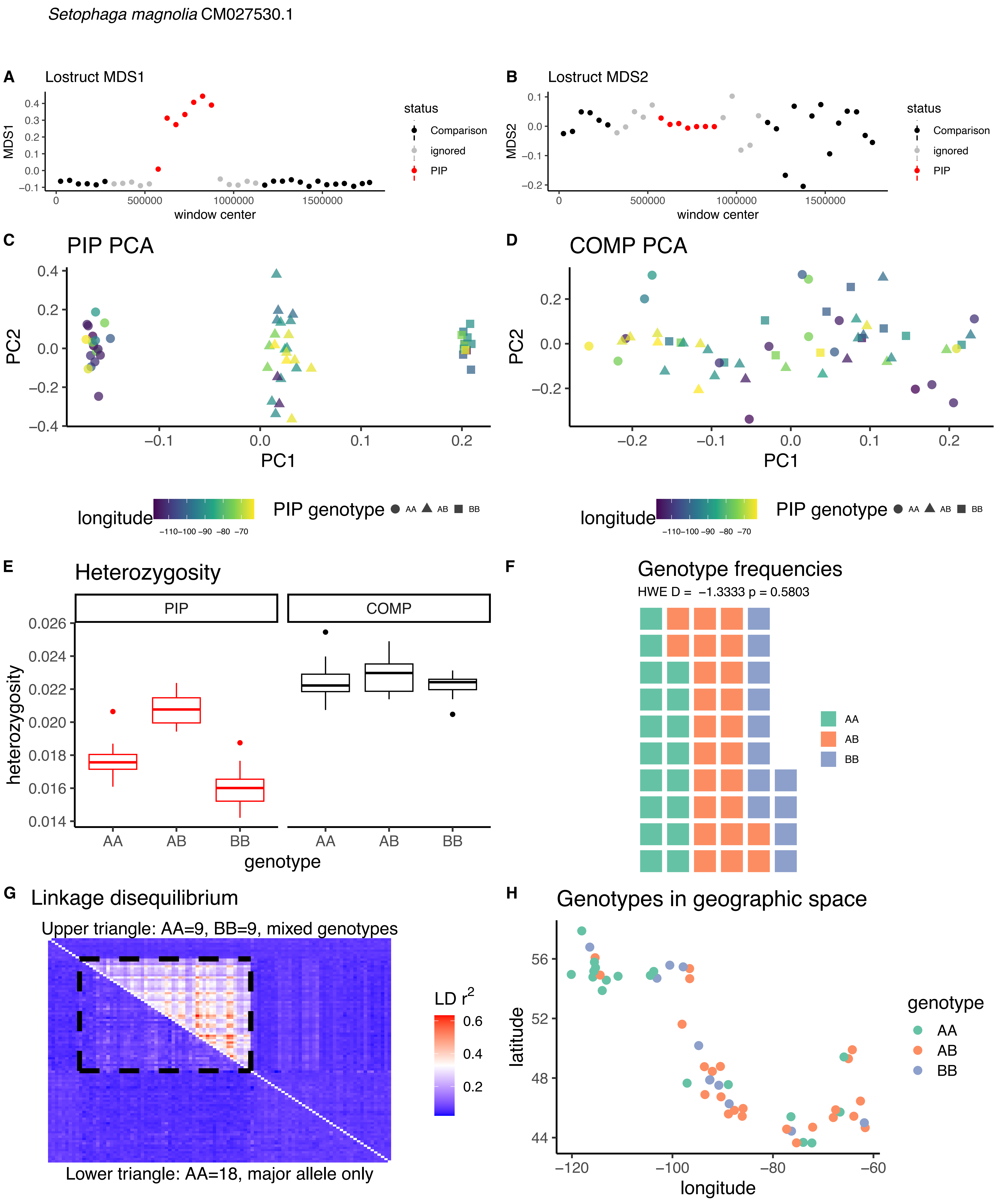

Setophaga magnolia CM027530.1
A
Lostruct MDS1
MDS1
window center
status
Comparison
ignored
PIP
B
Lostruct MDS2
MDS2
window center
status
Comparison
ignored
PIP
C
PIP PCA
PC1
PC2
D
COMP PCA
PC1
PC2
longitude
PIP genotype
AA
AB
BB
E
Heterozygosity
PIP
COMP
heterozygosity
genotype
F
Genotype frequencies
HWE D = -1.3333 p = 0.5803
AA
AB
BB
G
Linkage disequilibrium
Upper triangle: AA=9, BB=9, mixed genotypes
Lower triangle: AA=18, major allele only
LD r2
H
Genotypes in geographic space
latitude
longitude
genotype
AA
AB
BB

*Setophaga magnolia* CM027508.1A

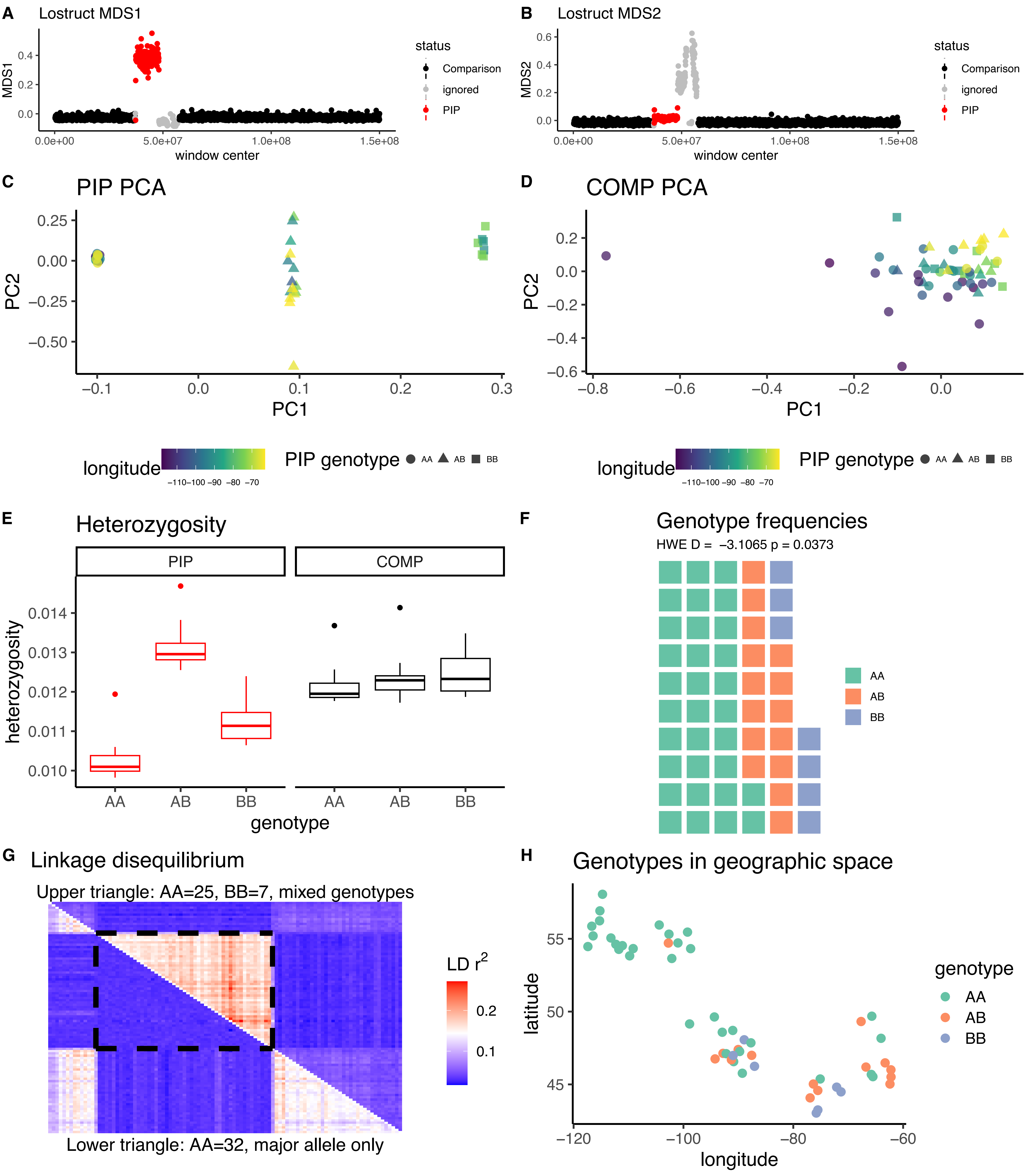

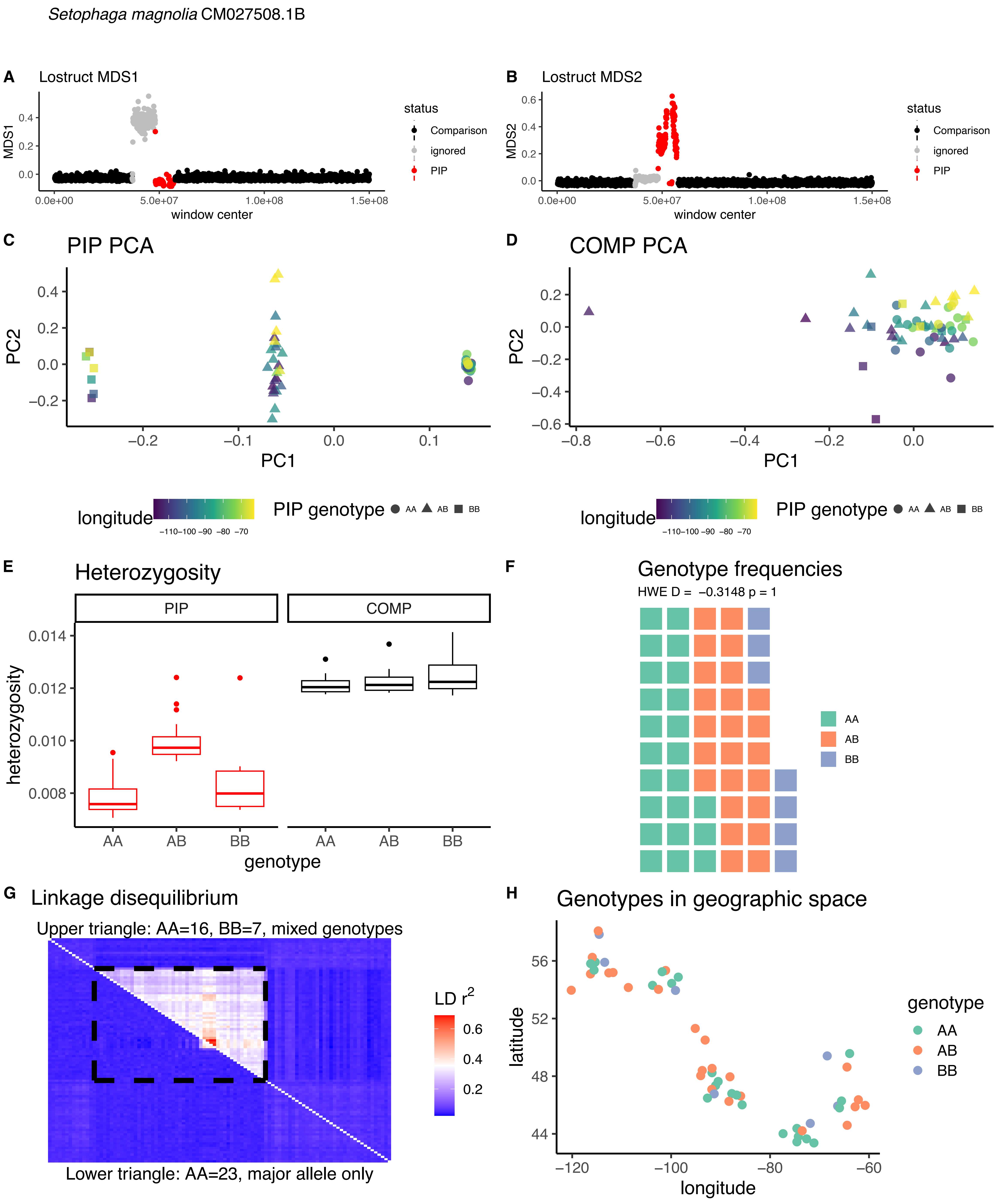

Setophaga magnolia CM027508.1B
A Lostruct MDS1
MDS1
window center
status
Comparison
ignored
PIP
B Lostruct MDS2
MDS2
C PIP PCA
PC1
PC2
D COMP PCA
longitude
PIP genotype AA AB BB
E Heterozygosity
PIP
COMP
heterozygosity
genotype
F Genotype frequencies
HWE D = −0.3148 p = 1
AA
AB
BB
G Linkage disequilibrium
Upper triangle: AA=16, BB=7, mixed genotypes
LD r2
Lower triangle: AA=23, major allele only
H Genotypes in geographic space
latitude
longitude
genotype

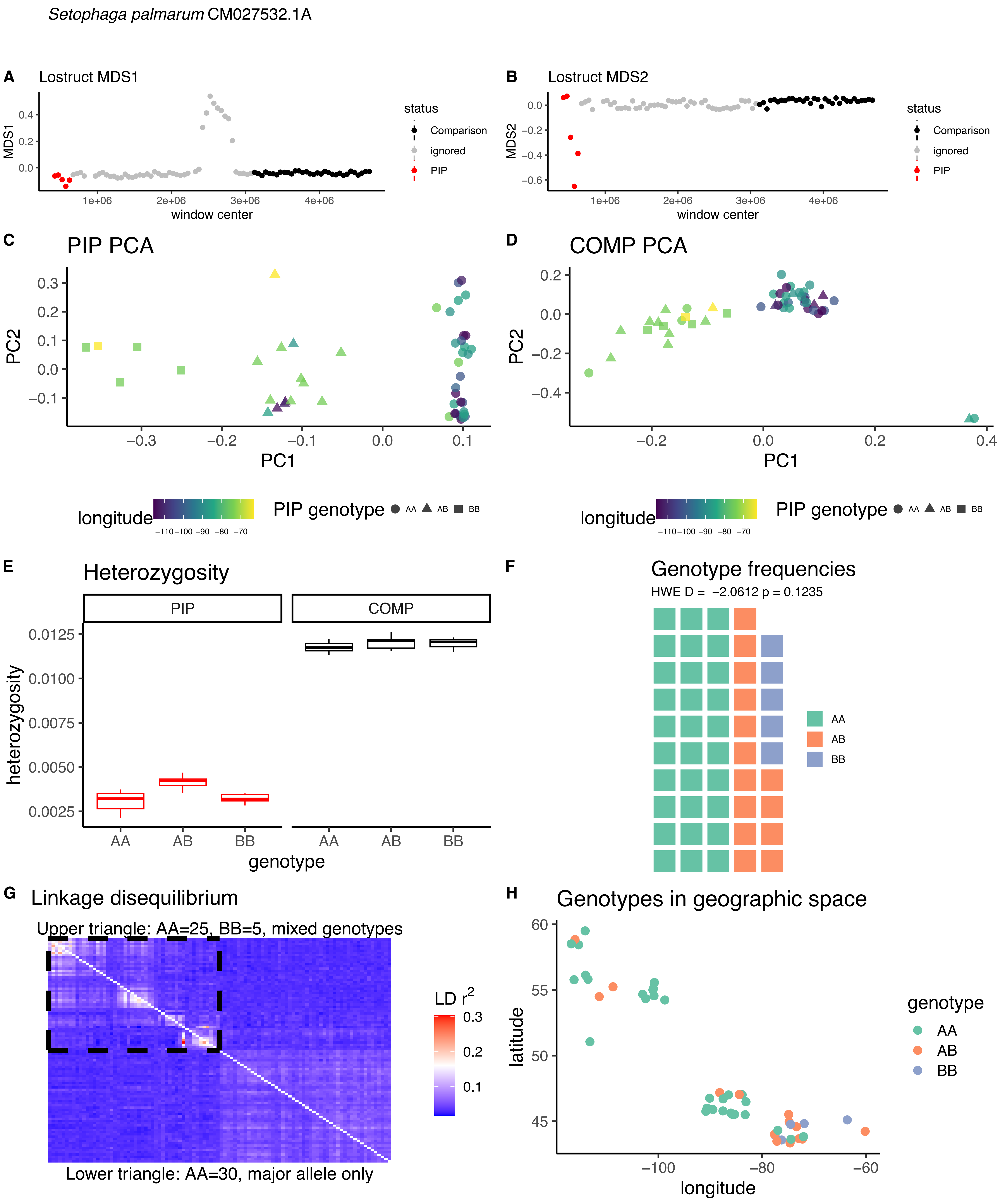

Setophaga palmarum CM027532.1A
A
Lostruct MDS1
MDS1
0.4
0.2
0.0
1e+06
2e+06
3e+06
4e+06
window center
status
Comparison
ignored
PIP
B
Lostruct MDS2
MDS2
0.0
−0.2
−0.4
−0.6
1e+06
2e+06
3e+06
4e+06
window center
status
Comparison
ignored
PIP
C
PIP PCA
PC2
0.3
0.2
0.1
0.0
−0.1
−0.3
−0.2
−0.1
0.0
0.1
PC1
D
COMP PCA
PC2
0.2
0.0
−0.2
−0.4
−0.2
0.0
0.2
0.4
PC1
longitude
−110 −100 −90 −80 −70
PIP genotype
AA
AB
BB
longitude
−110 −100 −90 −80 −70
PIP genotype
AA
AB
BB
E
Heterozygosity
PIP
COMP
heterozygosity
0.0125
0.0100
0.0075
0.0050
0.0025
AA
AB
BB
AA
AB
BB
genotype
F
Genotype frequencies
HWE D = −2.0612 p = 0.1235
AA
AB
BB
G
Linkage disequilibrium
Upper triangle: AA=25, BB=5, mixed genotypes
LD r²
0.3
0.2
0.1
Lower triangle: AA=30, major allele only
H
Genotypes in geographic space
latitude
60
55
50
45
−100
−80
−60
longitude
genotype
AA
AB
BB

*Setophaga palmarum* CM027532.1B

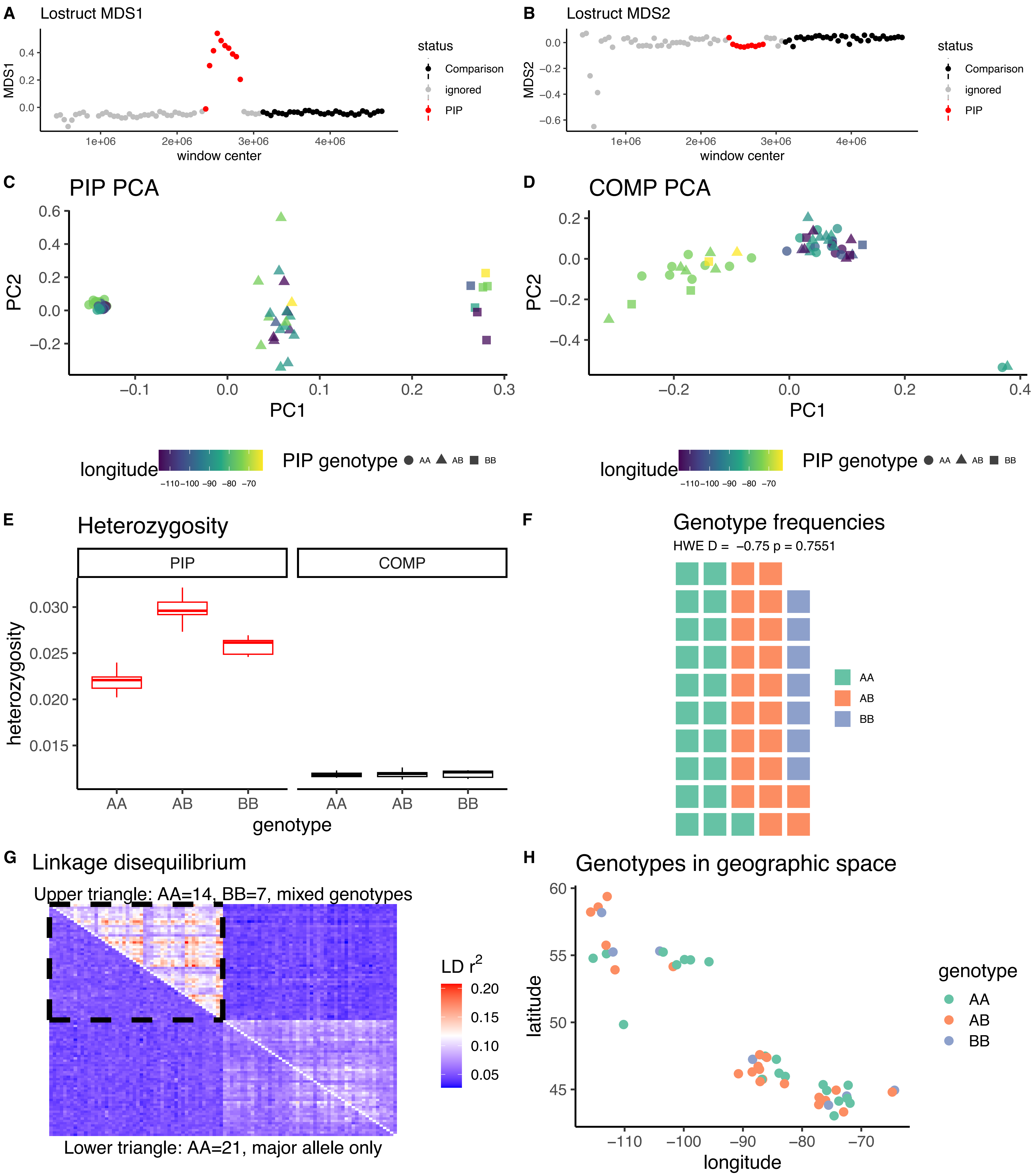

*Setophaga palmarum* CM027530.1

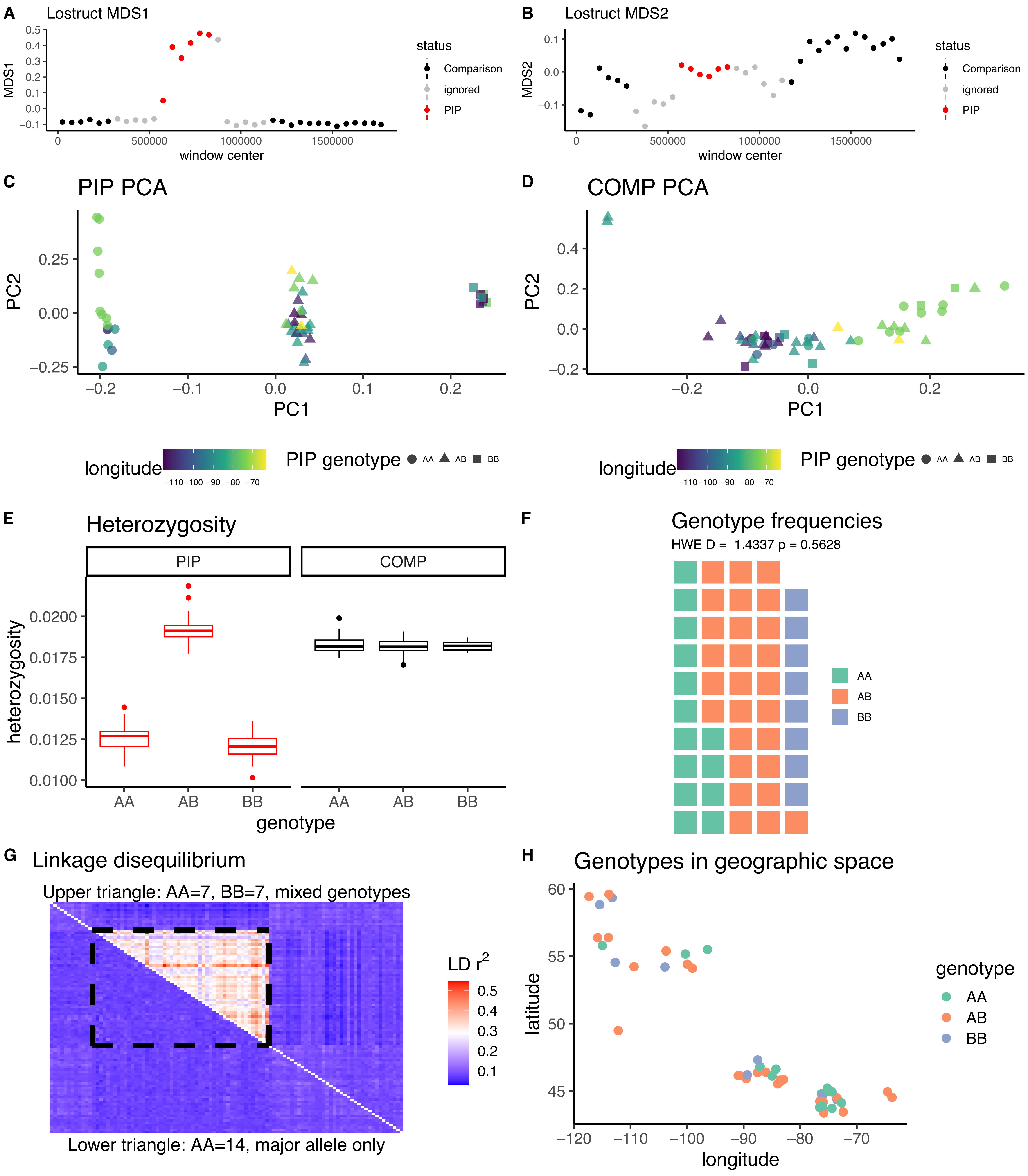

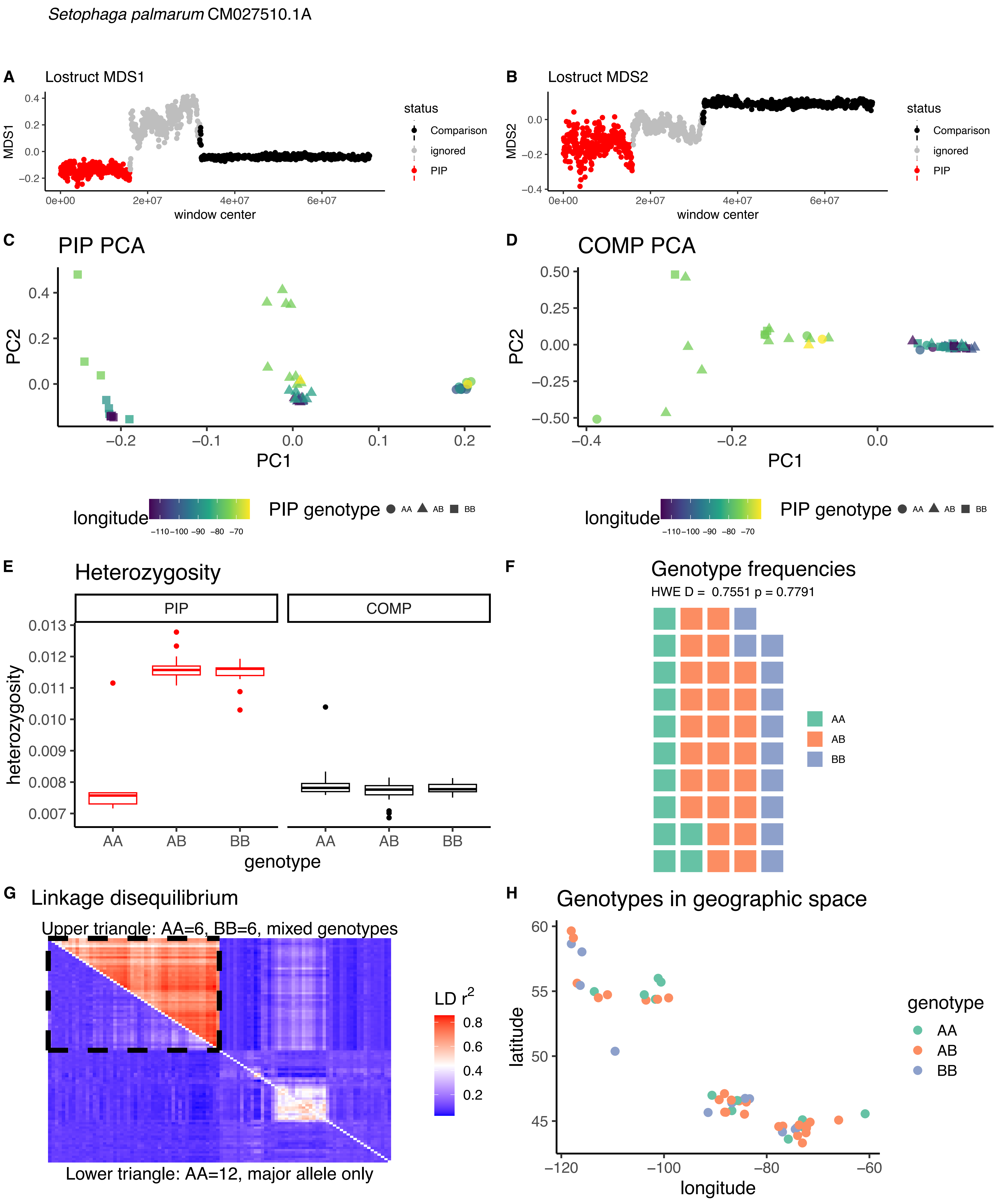

Setophaga palmarum CM027510.1A
A
Lostruct MDS1
MDS1
window center
status
Comparison
ignored
PIP
B
Lostruct MDS2
MDS2
window center
C
PIP PCA
PC1
PC2
D
COMP PCA
longitude
PIP genotype
AA
AB
BB
E
Heterozygosity
PIP
COMP
heterozygosity
genotype
F
Genotype frequencies
HWE D = 0.7551 p = 0.7791
G
Linkage disequilibrium
Upper triangle: AA=6, BB=6, mixed genotypes
LD r^2
Lower triangle: AA=12, major allele only
H
Genotypes in geographic space
latitude
longitude
genotype

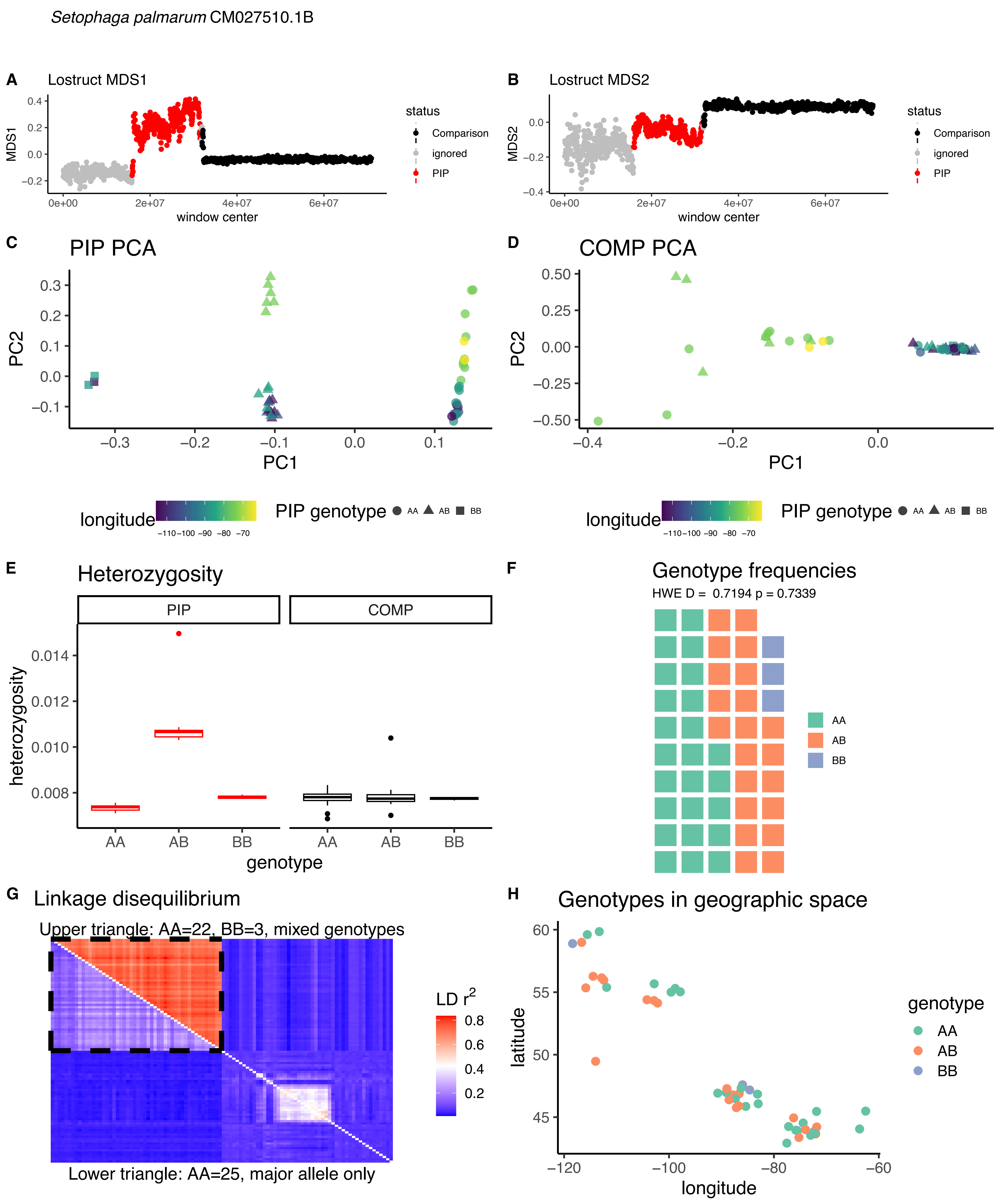

Setophaga palmarum CM027510.1B
A
Lostruct MDS1
MDS1
window center
status
Comparison
ignored
PIP
B
Lostruct MDS2
MDS2
window center
C
PIP PCA
PC1
PC2
D
COMP PCA
longitude
PIP genotype
AA
AB
BB
E
Heterozygosity
PIP
COMP
heterozygosity
genotype
F
Genotype frequencies
HWE D = 0.7194 p = 0.7339
G
Linkage disequilibrium
Upper triangle: AA=22, BB=3, mixed genotypes
Lower triangle: AA=25, major allele only
LD r2
H
Genotypes in geographic space
latitude
genotype

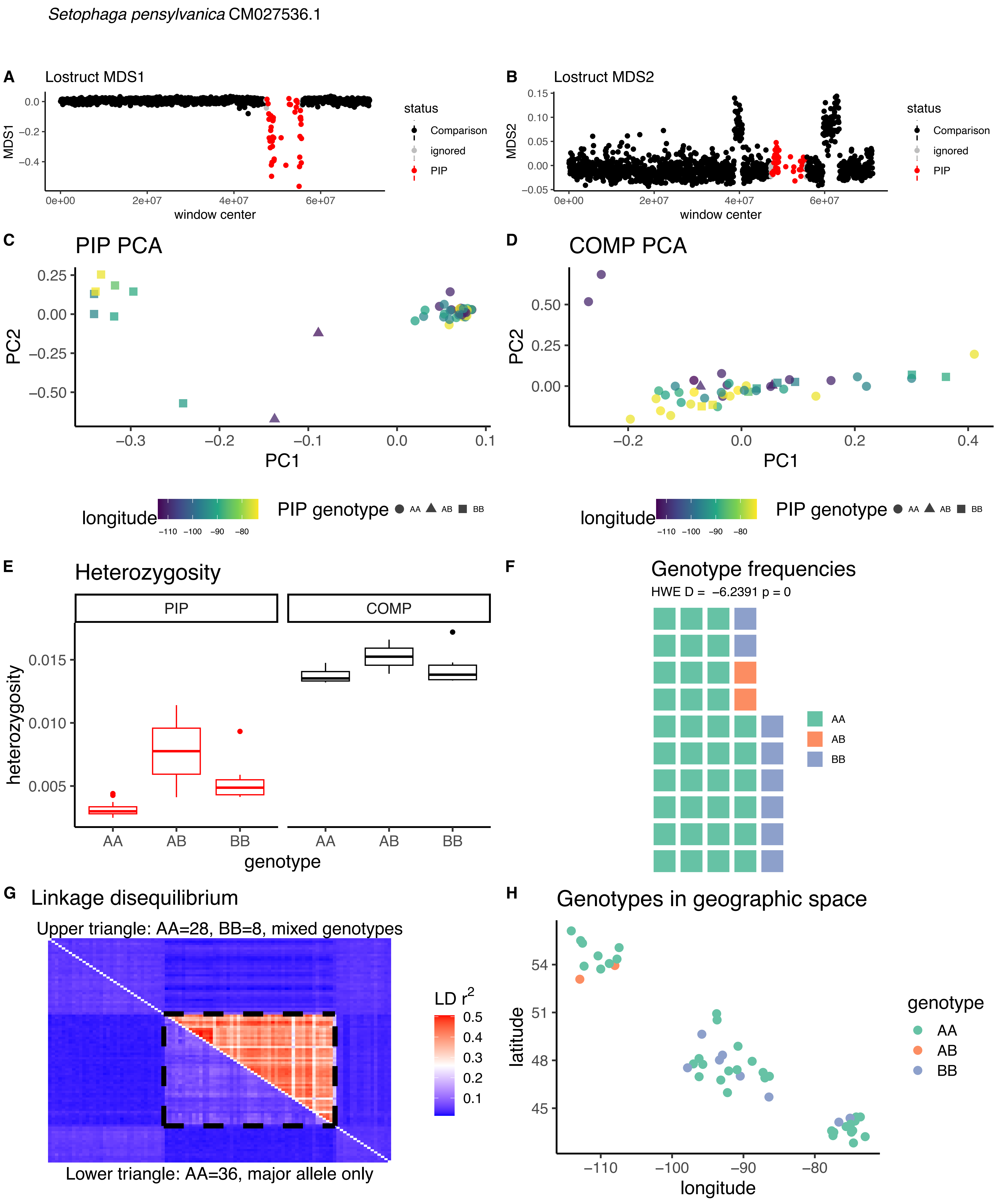

Setophaga pensylvanica CM027536.1
A
Lostruct MDS1
MDS1
window center
status
Comparison
ignored
PIP
B
Lostruct MDS2
MDS2
C
PIP PCA
PC2
PC1
D
COMP PCA
longitude
PIP genotype
AA
AB
BB
E
Heterozygosity
PIP
COMP
heterozygosity
genotype
F
Genotype frequencies
HWE D = −6.2391 p = 0
G
Linkage disequilibrium
Upper triangle: AA=28, BB=8, mixed genotypes
Lower triangle: AA=36, major allele only
LD r²
H
Genotypes in geographic space
latitude
genotype

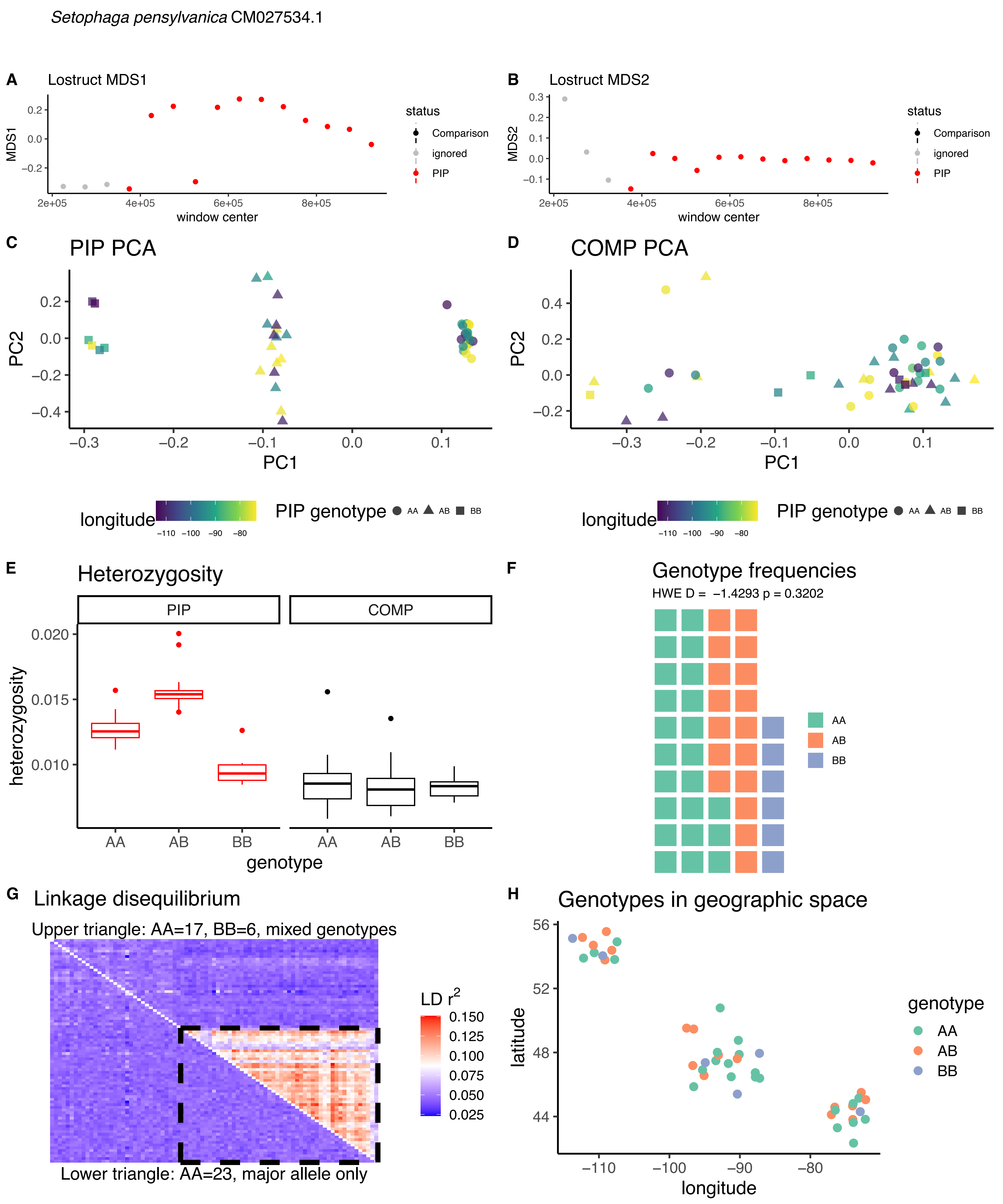
*Setophaga pensylvanica* CM027534.1
A
Lostruct MDS1
MDS1
window center
status
Comparison
ignored
PIP
B
Lostruct MDS2
MDS2
window center
status
Comparison
ignored
PIP
C
PIP PCA
PC1
PC2
D
COMP PCA
PC1
PC2
longitude
PIP genotype AA AB BB
longitude
PIP genotype AA AB BB
E
Heterozygosity
PIP
COMP
heterozygosity
genotype
AA AB BB AA AB BB
F
Genotype frequencies
HWE D = −1.4293 p = 0.3202
AA
AB
BB
G
Linkage disequilibrium
Upper triangle: AA=17, BB=6, mixed genotypes
LD r^2
Lower triangle: AA=23, major allele only
H
Genotypes in geographic space
latitude
longitude
genotype
AA
AB
BB

*Setophaga pensylvanica* CM027532.1

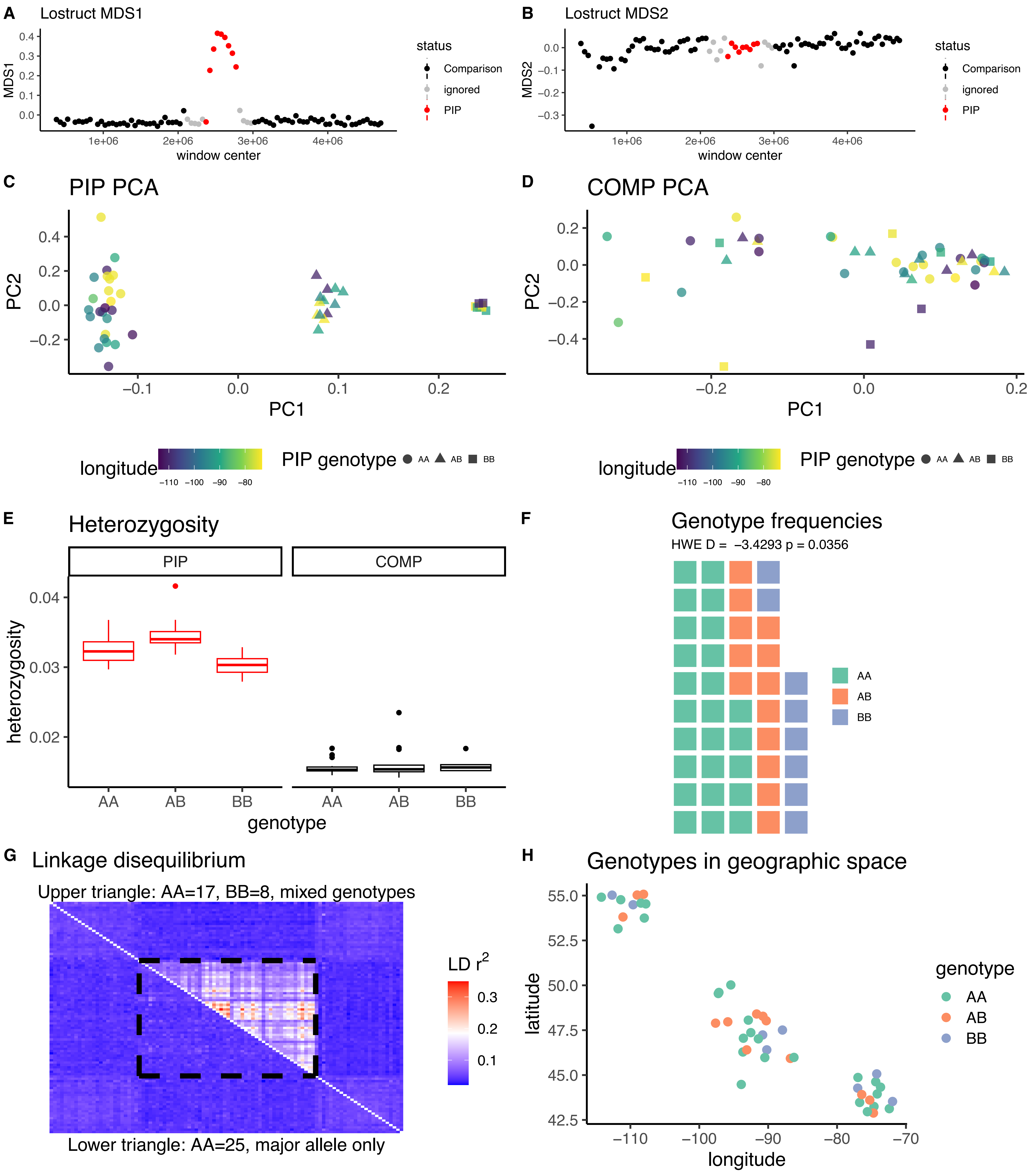

*Setophaga pensylvanica* CM027530.1

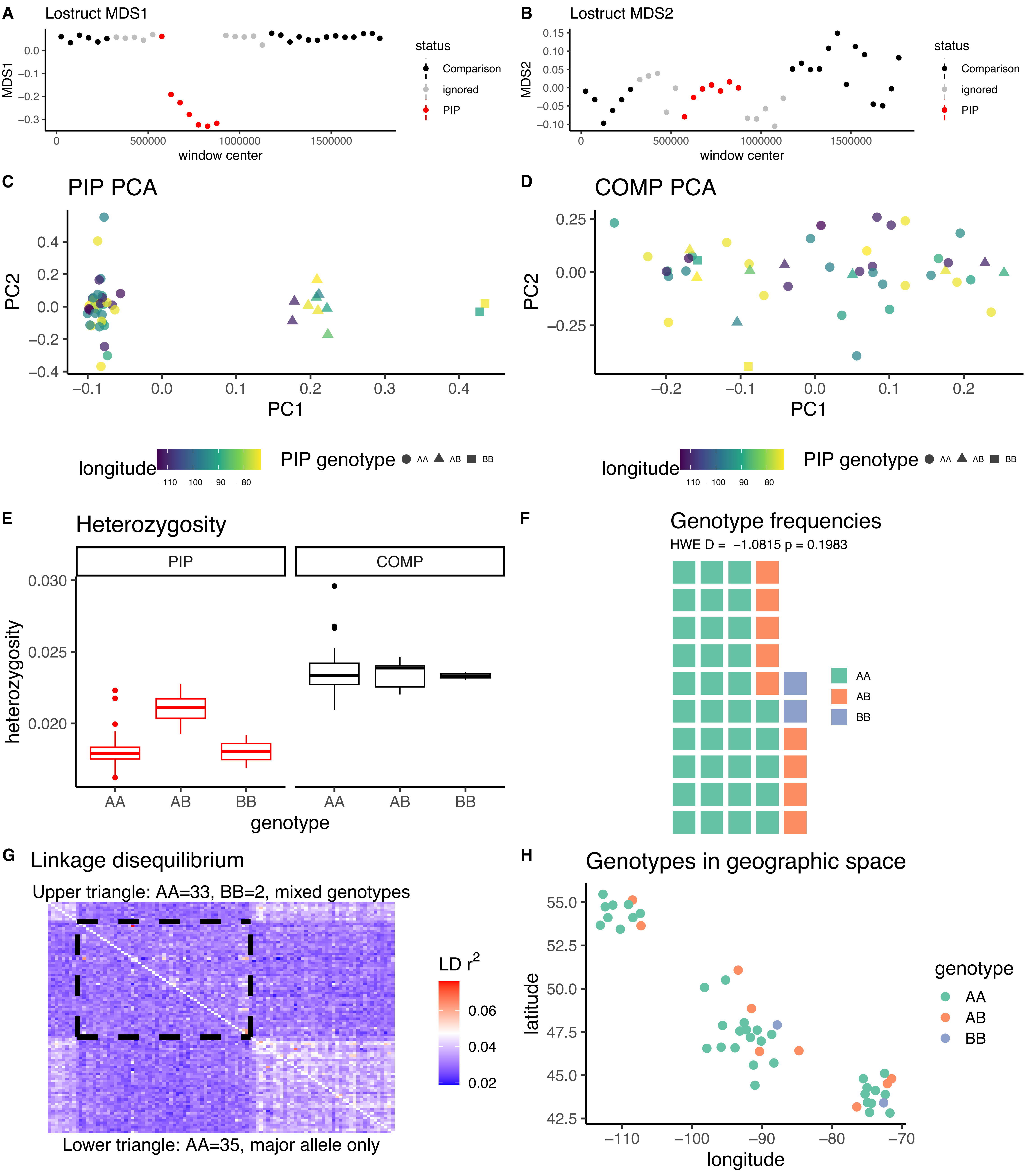

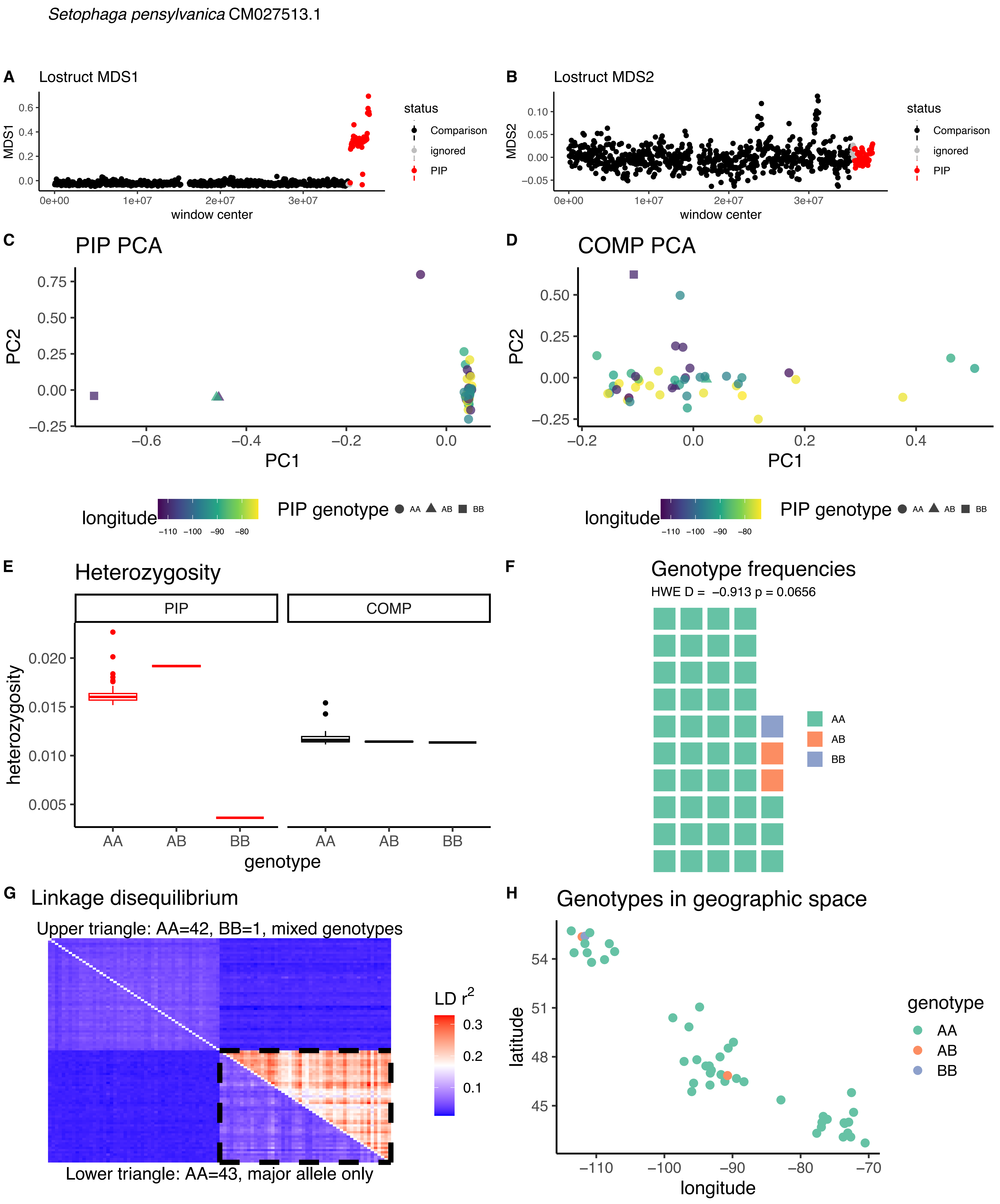
Setophaga pensylvanica CM027513.1
A Lostruct MDS1
MDS1
window center
status
Comparison
ignored
PIP
B Lostruct MDS2
MDS2
window center
C PIP PCA
PC1
PC2
D COMP PCA
longitude
PIP genotype AA AB BB
E Heterozygosity
PIP
COMP
heterozygosity
genotype
F Genotype frequencies
HWE D = −0.913 p = 0.0656
AA
AB
BB
G Linkage disequilibrium
Upper triangle: AA=42, BB=1, mixed genotypes
Lower triangle: AA=43, major allele only
LD r2
H Genotypes in geographic space
latitude
longitude
genotype

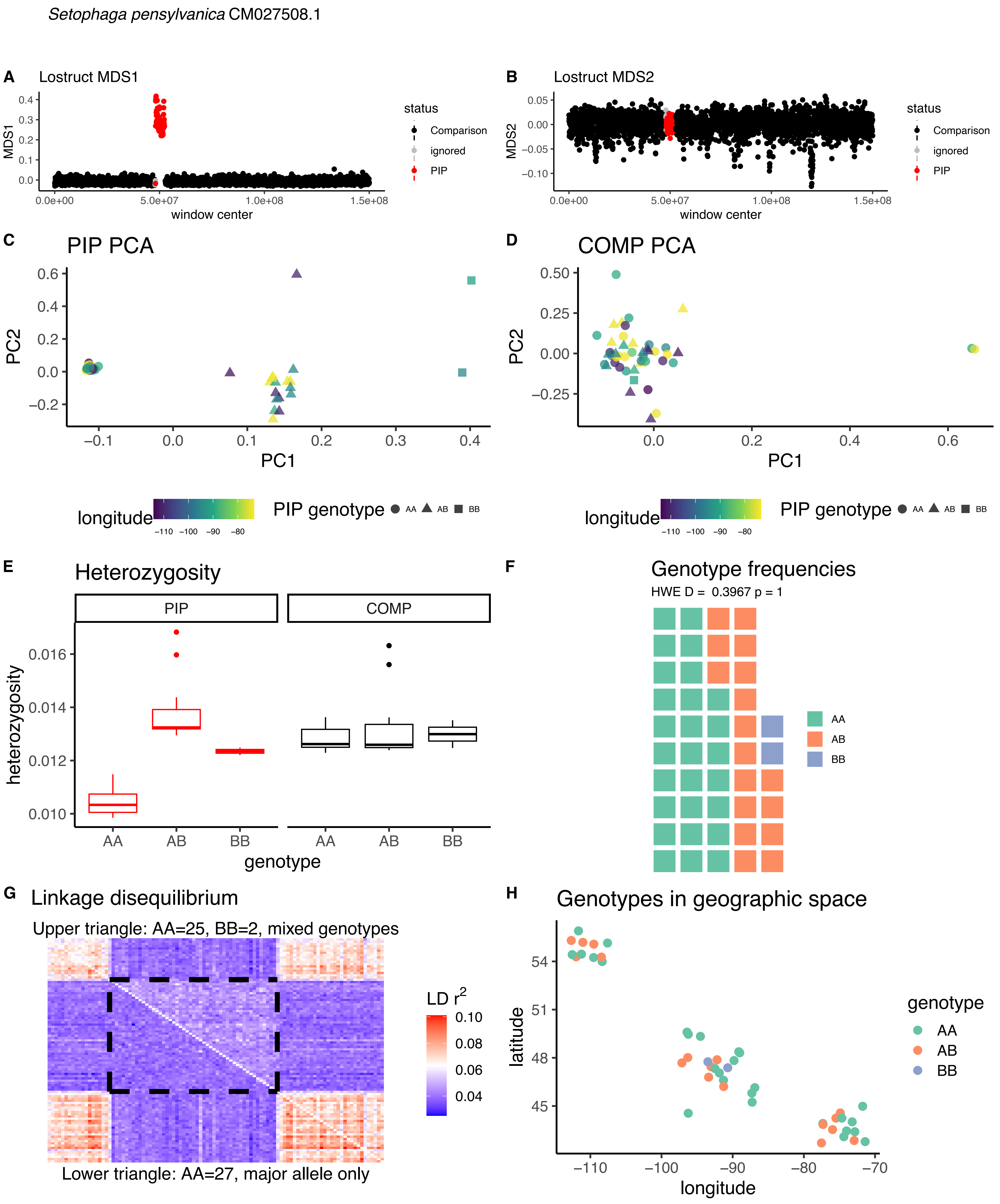

*Setophaga pensylvanica* CM027508.1
A
Lostruct MDS1
MDS1
window center
status
Comparison
ignored
PIP
B
Lostruct MDS2
MDS2
C
PIP PCA
PC1
PC2
D
COMP PCA
longitude
PIP genotype
AA
AB
BB
E
Heterozygosity
PIP
COMP
heterozygosity
genotype
F
Genotype frequencies
HWE D = 0.3967 p = 1
G
Linkage disequilibrium
Upper triangle: AA=25, BB=2, mixed genotypes
LD $r^2$
Lower triangle: AA=27, major allele only
H
Genotypes in geographic space
latitude
longitude
genotype

*Setophaga tigrina* CM027534.1

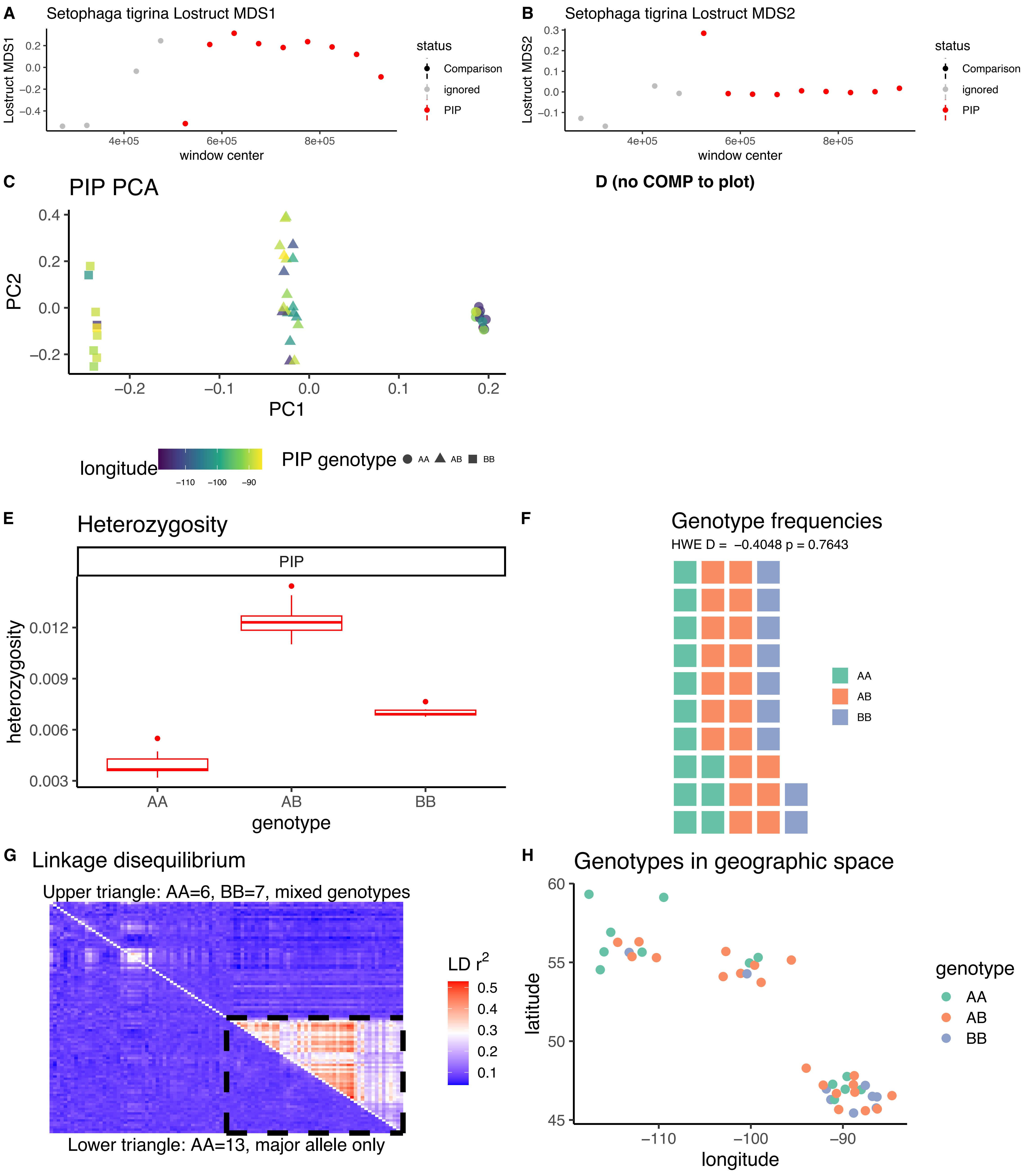

*Setophaga tigrina* CM027532.1

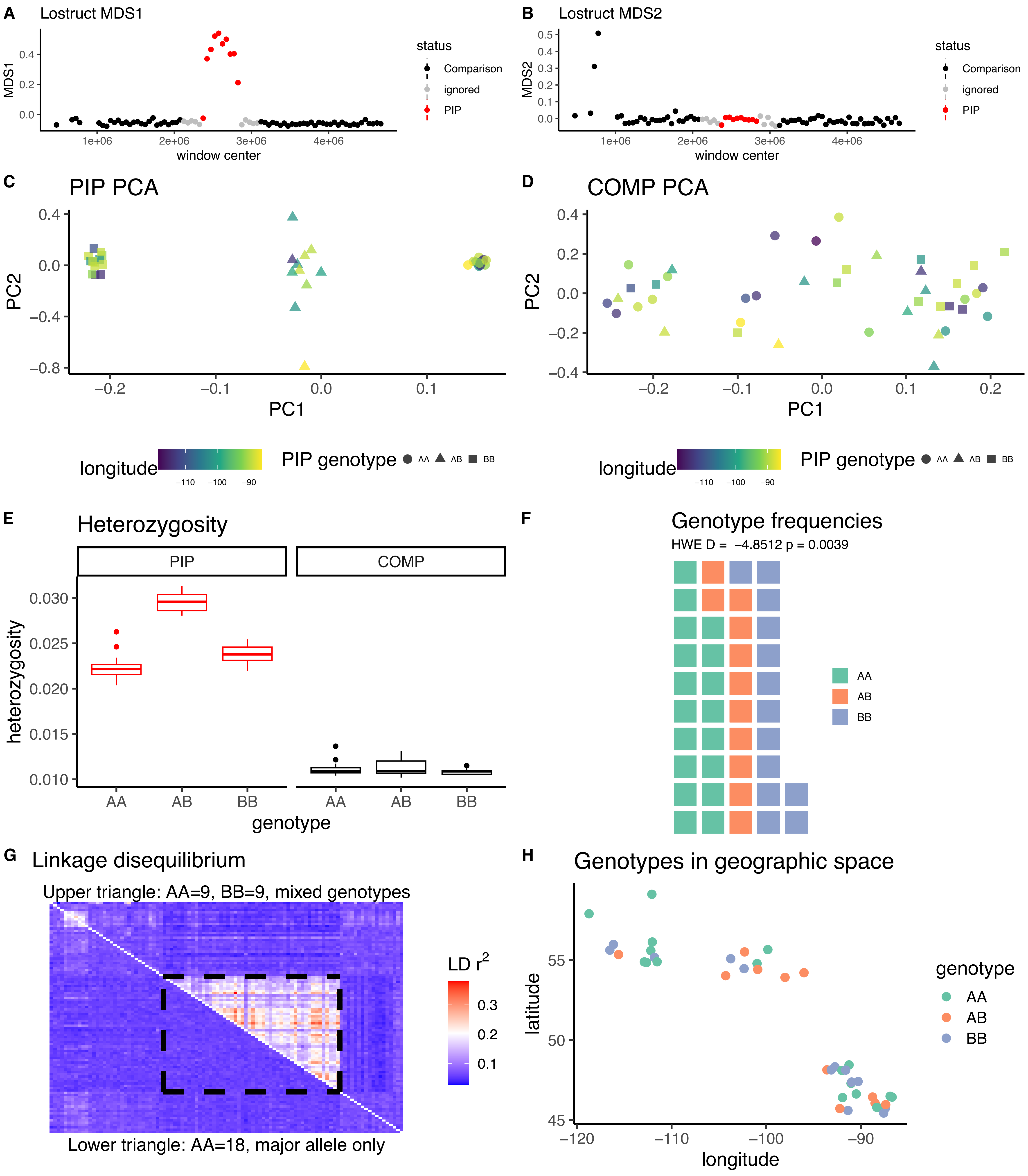

*Setophaga tigrina* CM027530.1

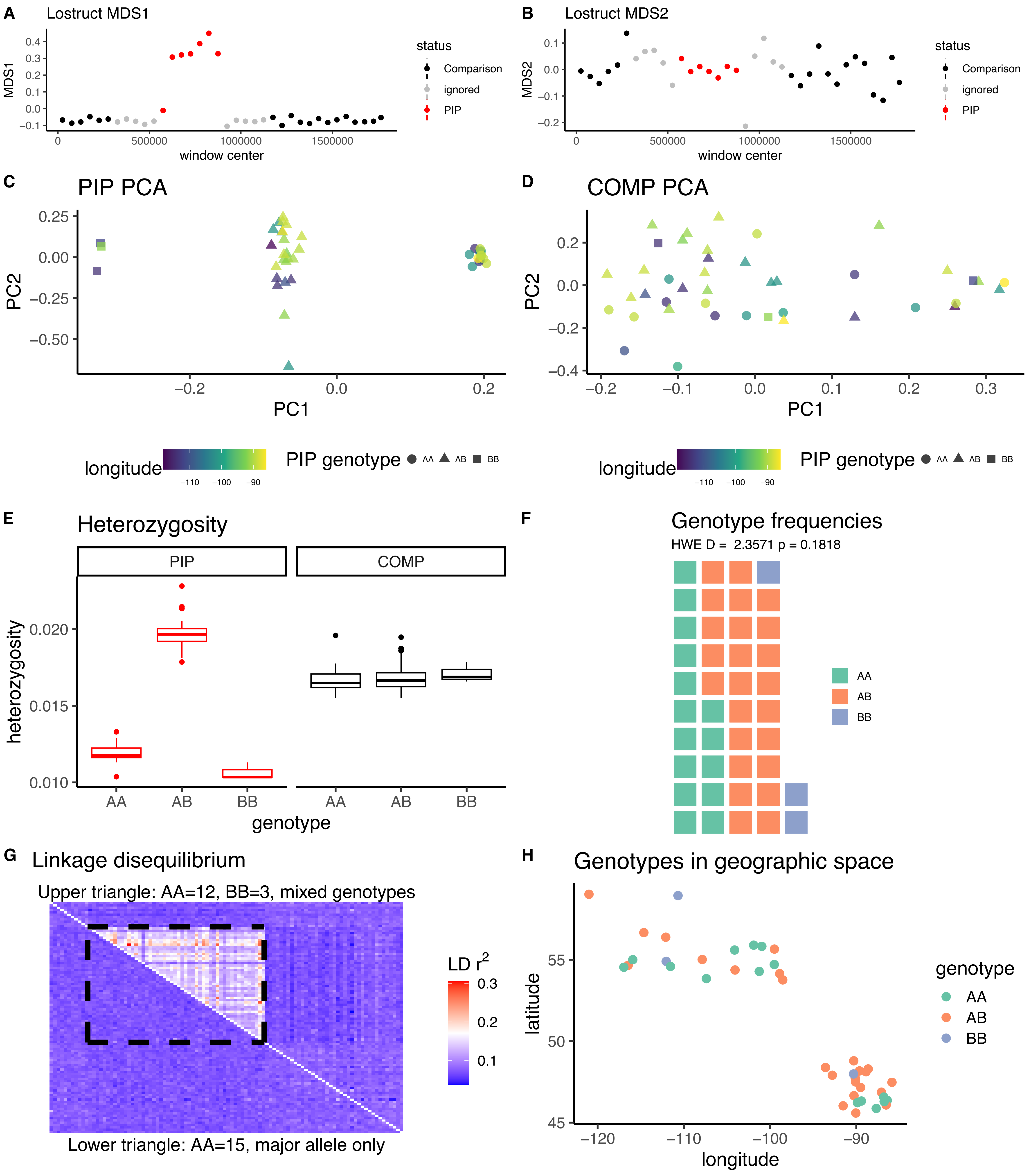

*Setophaga virens* CM027534.1

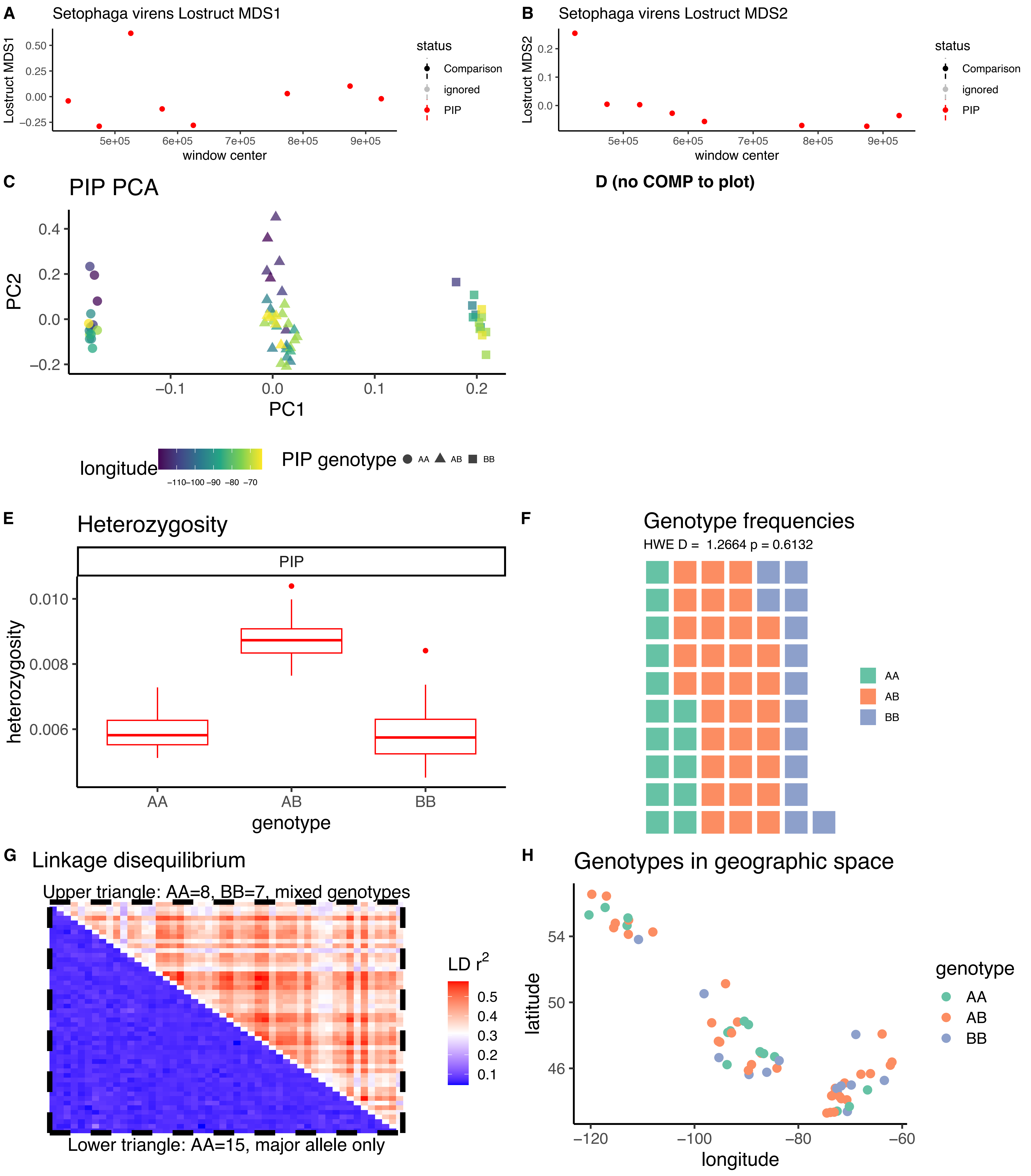

*Setophaga virens* CM027532.1

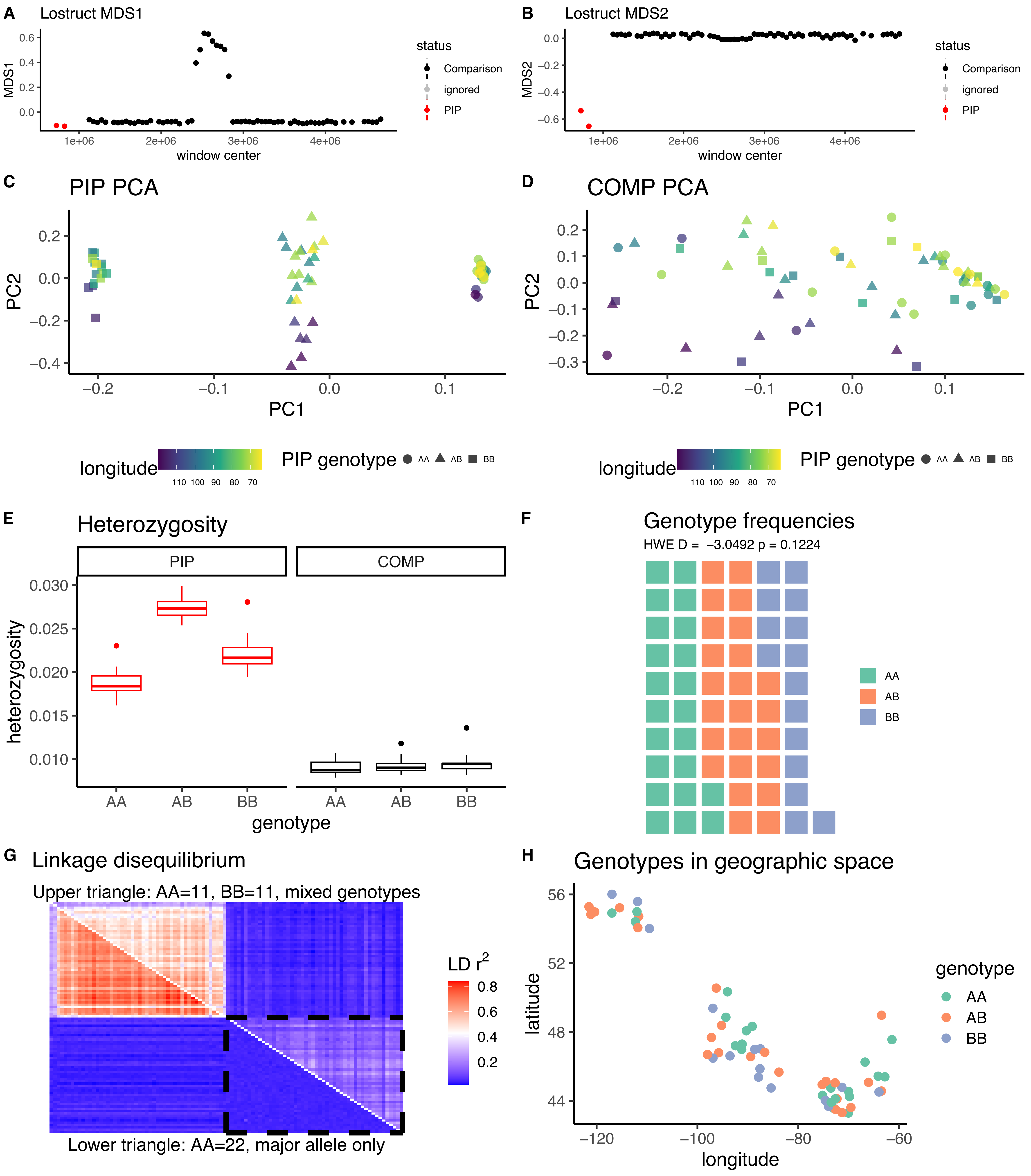

*Setophaga virens* CM027530.1

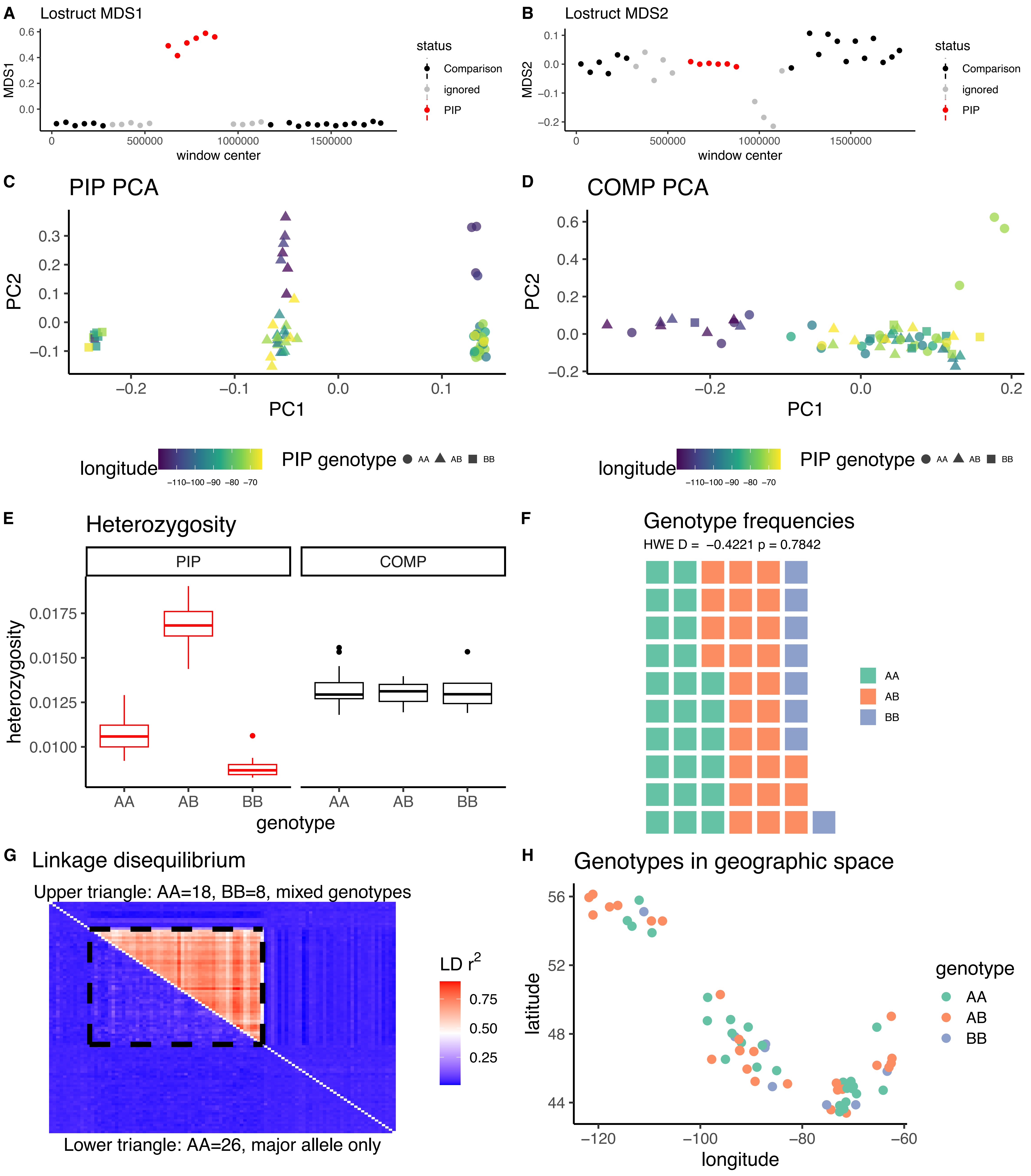

*Setophaga virens* CM027508.1A

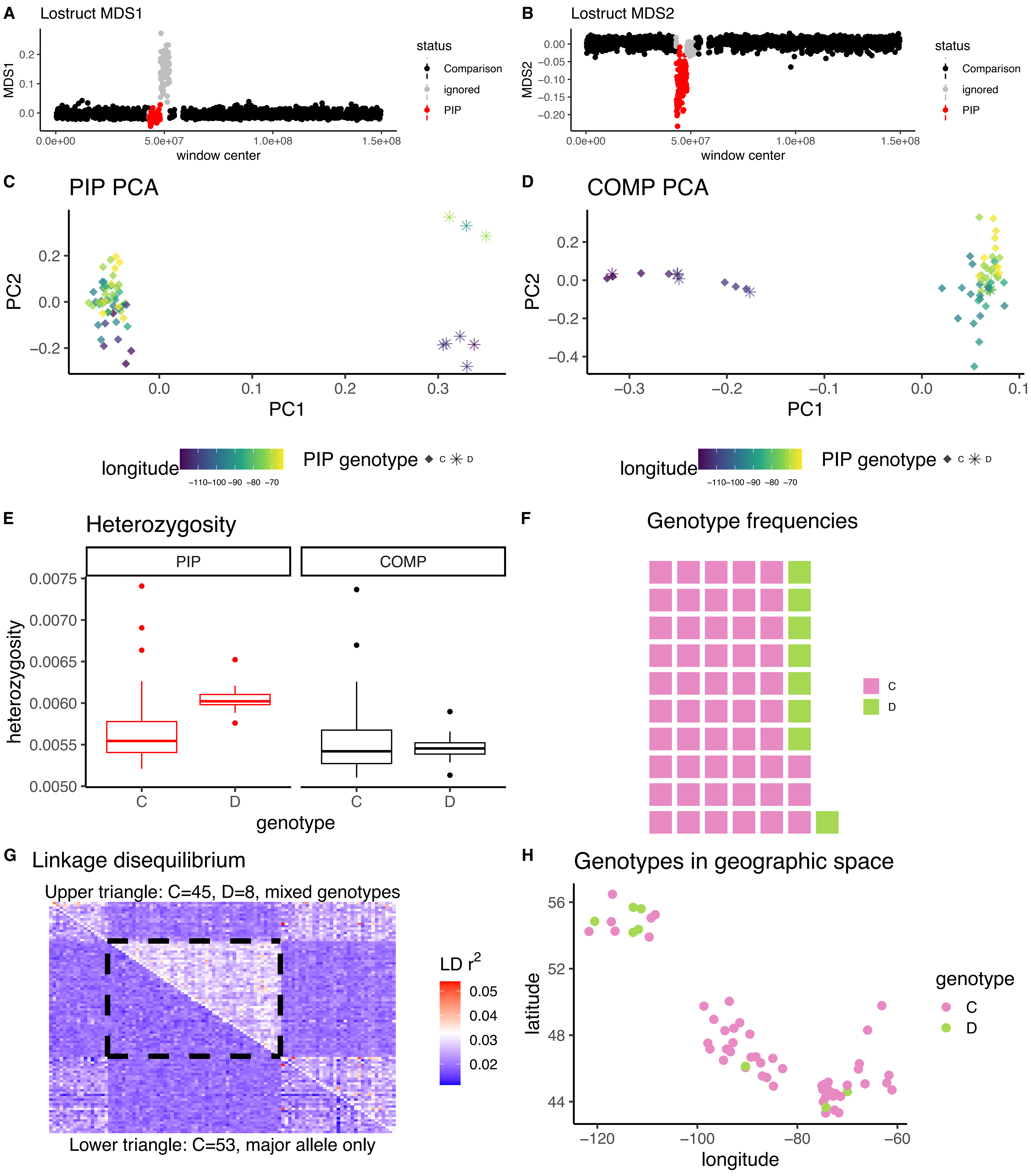

*Setophaga virens* CM027508.1B

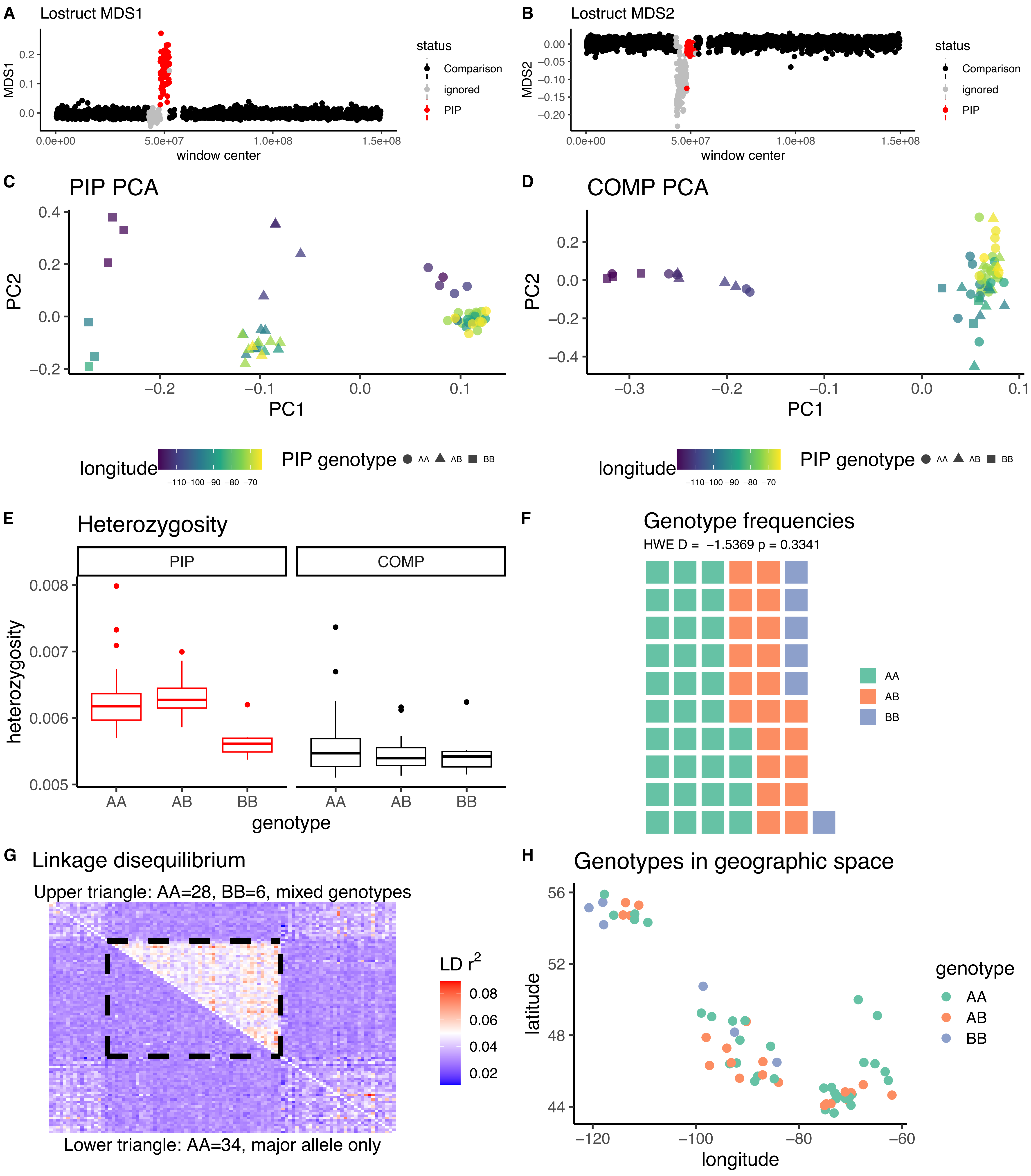
