## Supplementary material for "Large inversion polymorphisms are widespread in North American songbirds": Dataset S3

**Dataset S3**. In 26 cases, PIPs were identified in multiple species that were mapped to the same reference genome. Here we provide neighbor-joining genetic distance trees for each shared PIP, along with trees from comparison (COMP) regions of the same chromosome. Note that it was not always possible to identify an appropriate COMP region, and that PIPs from the same chromosome share a single COMP region. Each tree includes all species mapped to a given reference chromosome, including one randomly-chosen representative of each identified PIP genotype for each species. We excluded putative heterozygotes (genotype=AB) from this analysis. In cases where species did not have an identified PIP on the relevant chromosome, we chose one individual randomly. The shape at each tip on the tree indicates the PIP genotype of that individual, while color represents species. In some cases, branching patterns show individuals clustering by inversion genotype rather than by species, suggesting the existence of trans-specific polymorphisms. Note that AA and BB genotypes are named arbitrarily based on haplotype frequencies within their respective species, not homology—i.e., in cases of trans-specific polymorphisms, the A haplotype of one species may be apparently homologous with the B haplotype of another species.

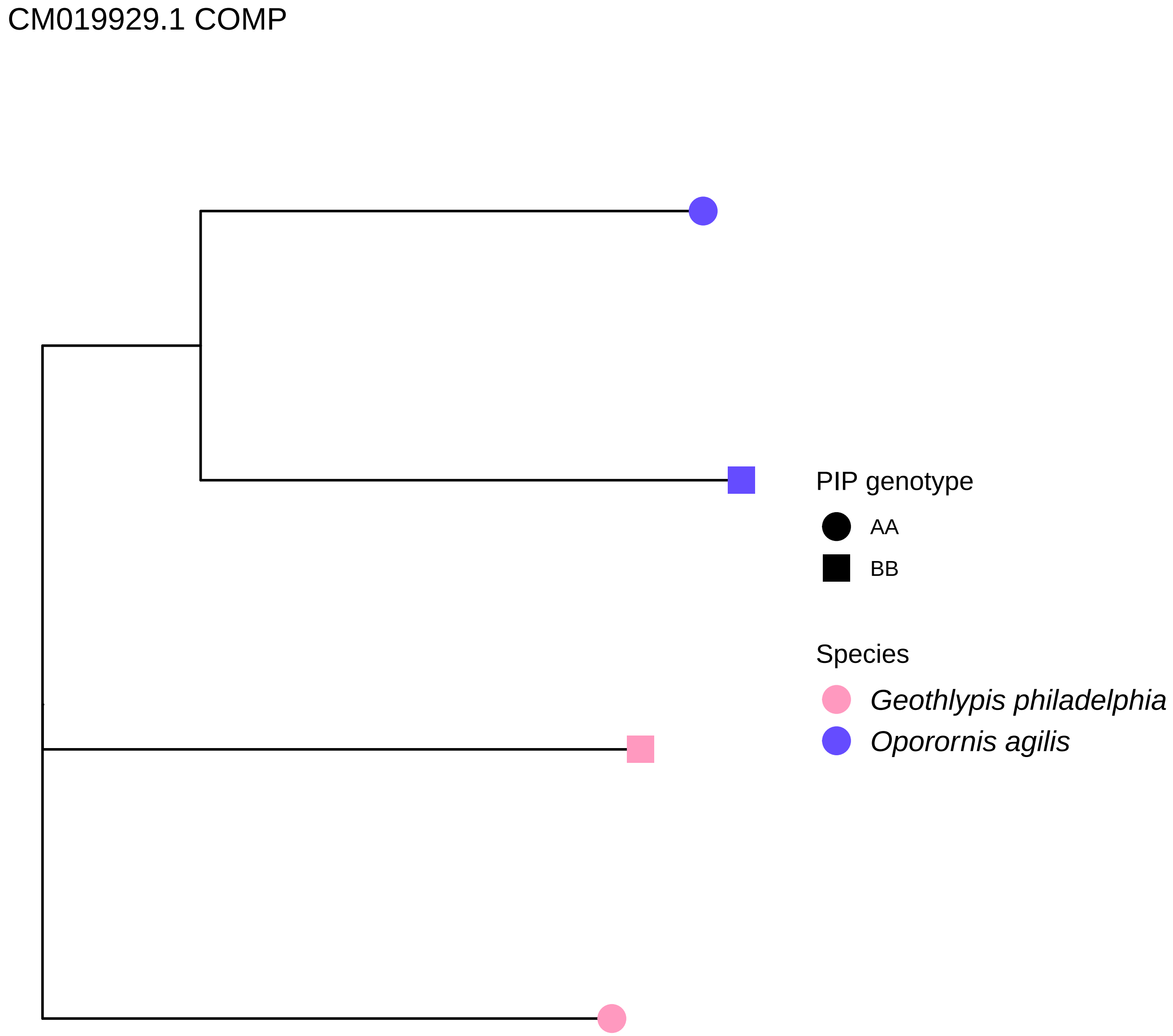
CM019929.1 COMP
PIP genotype
AA
BB
Species
Geothlypis philadelphia
Oporornis agilis

CM019929.1 PIP

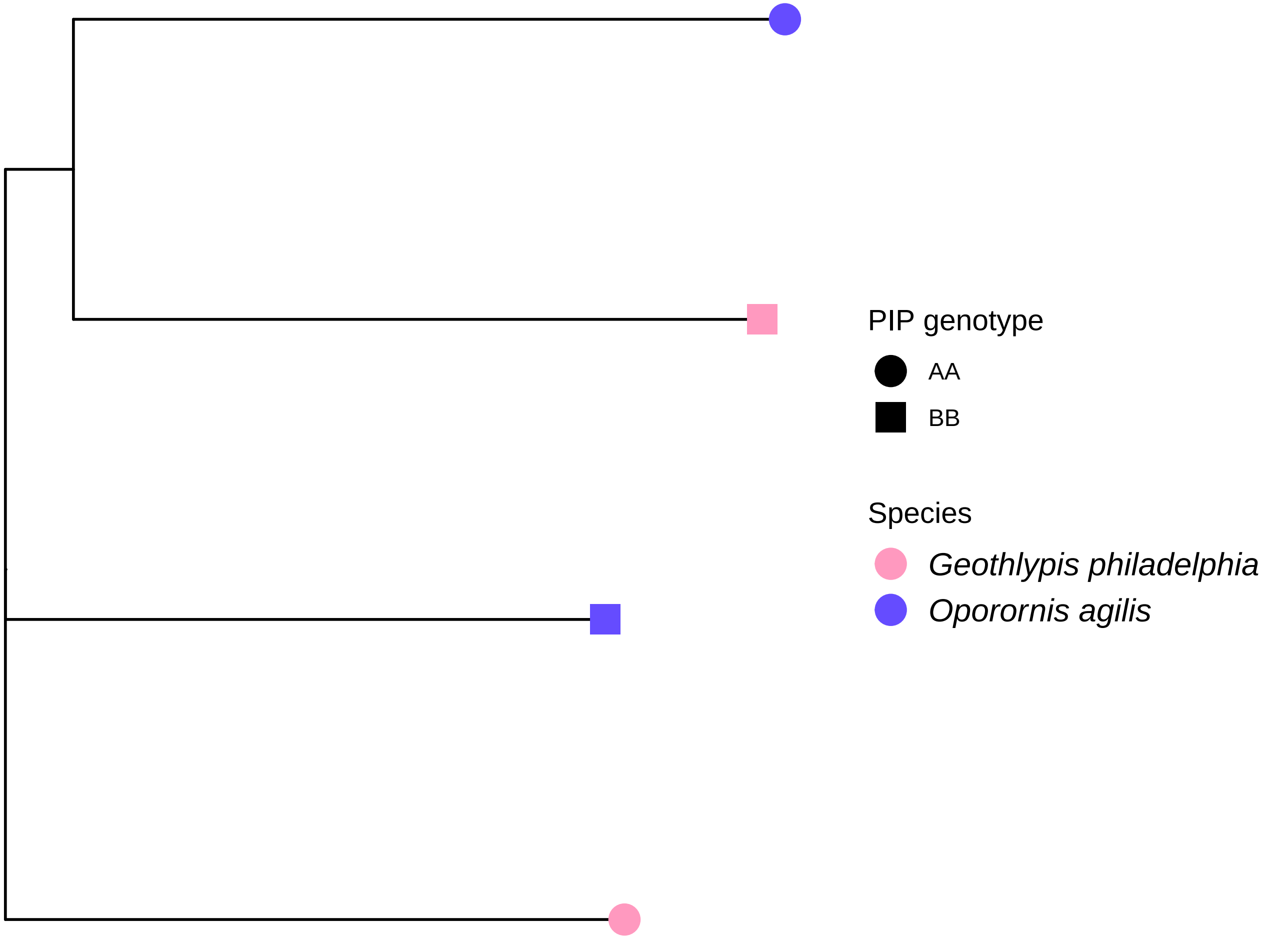

CM020365.1 COMP

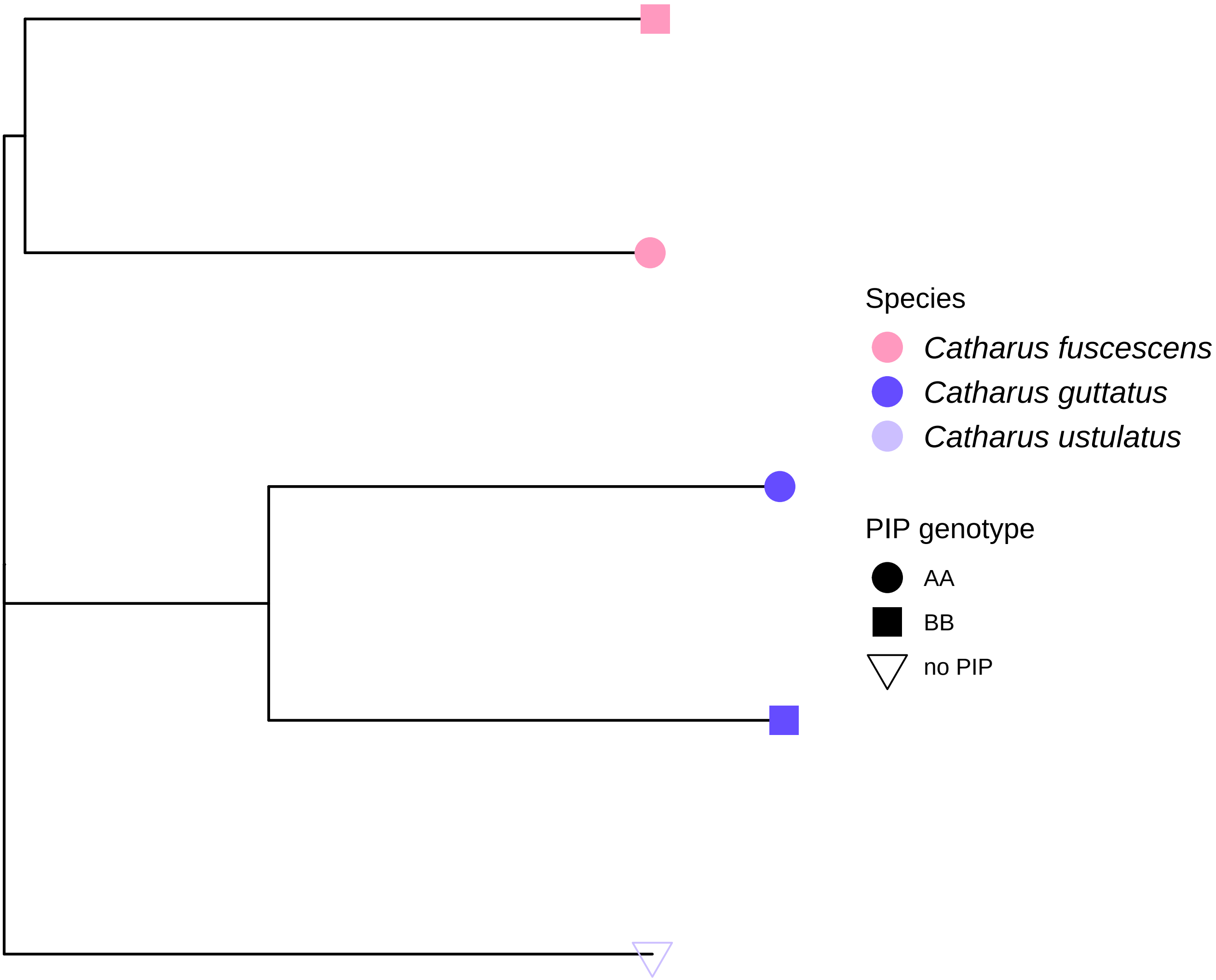

CM020365.1 PIP

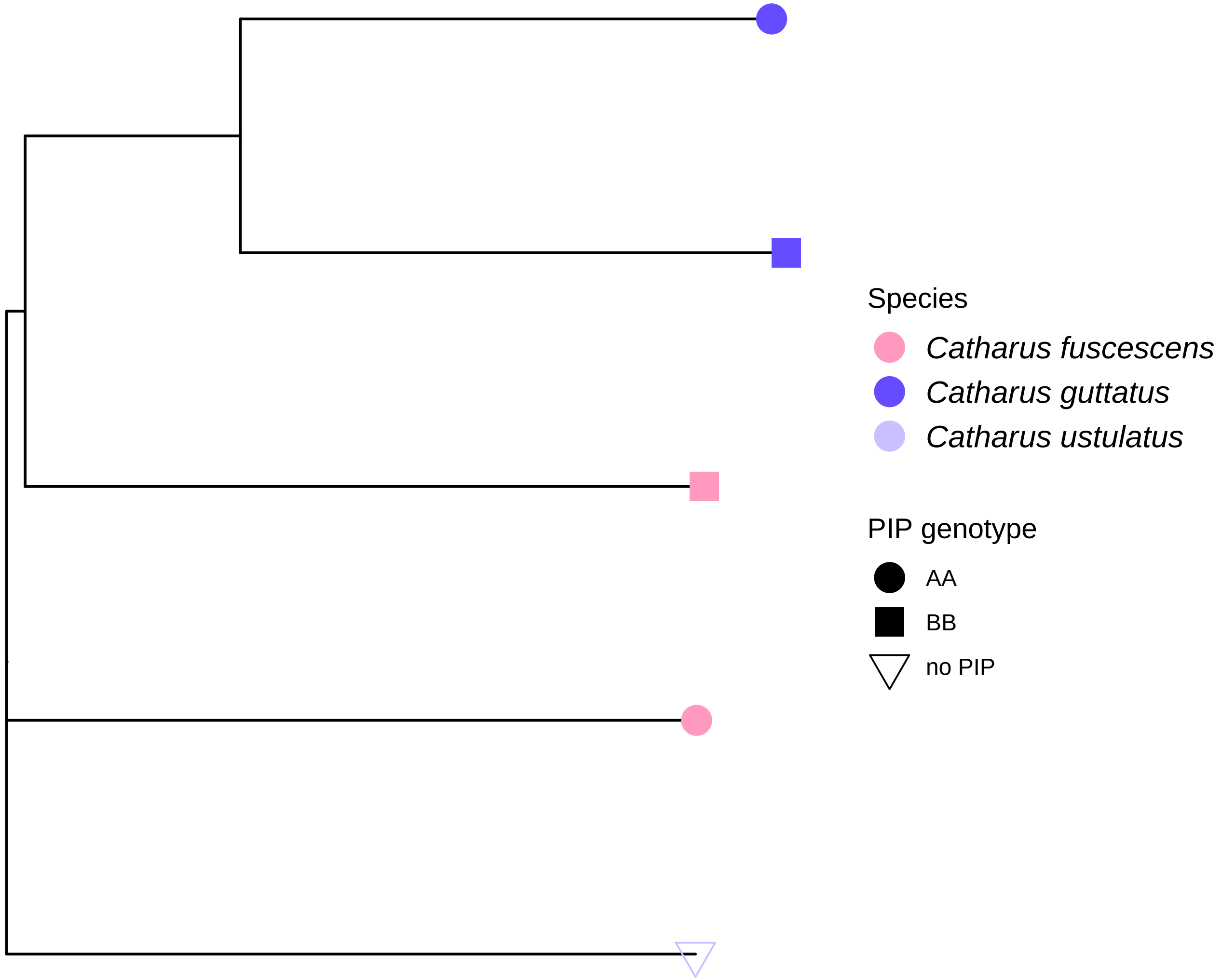

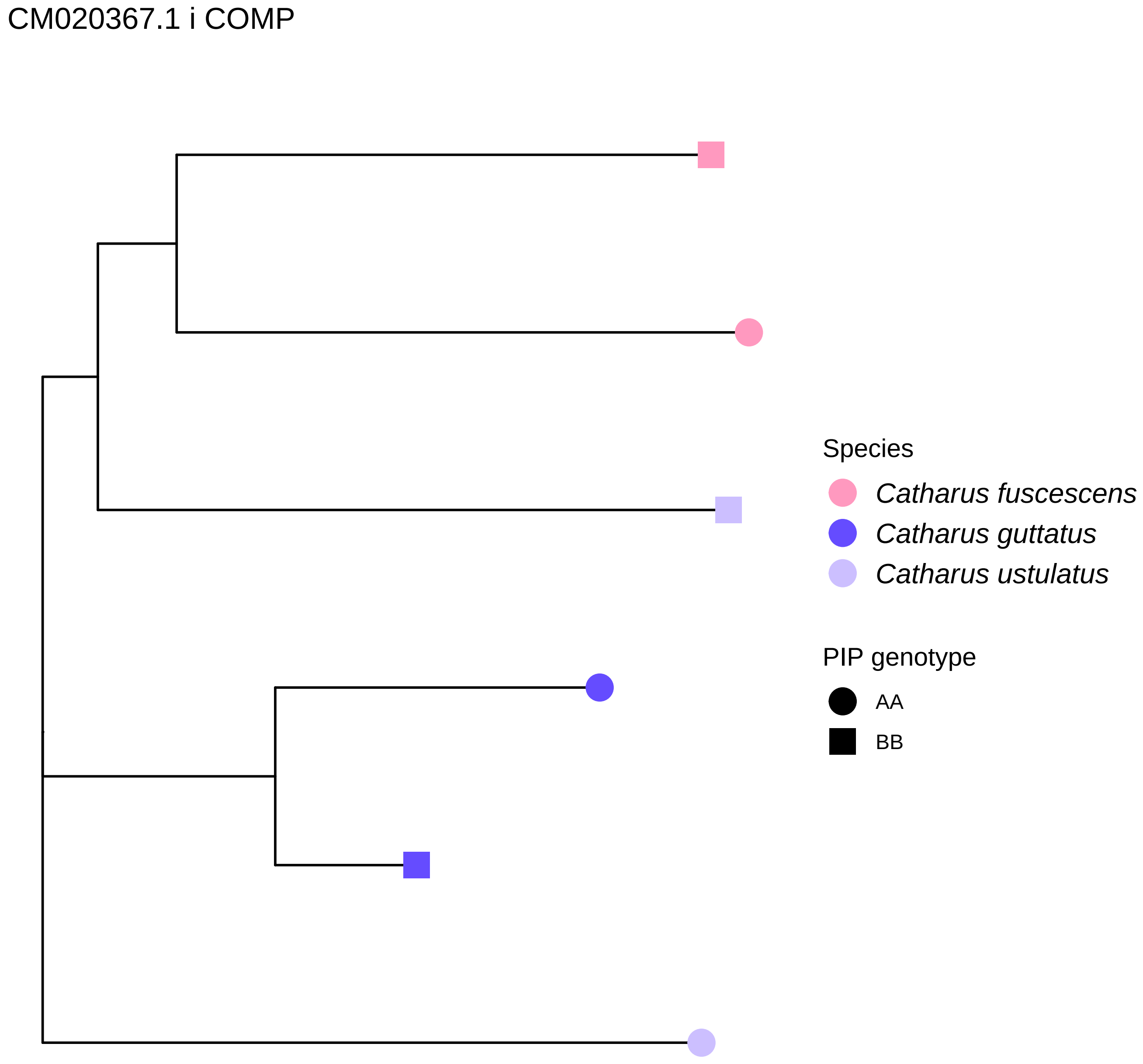
CM020367.1 i COMP
Species
Catharus fuscescens
Catharus guttatus
Catharus ustulatus
PIP genotype
AA
BB

CM020367.1 i PIP

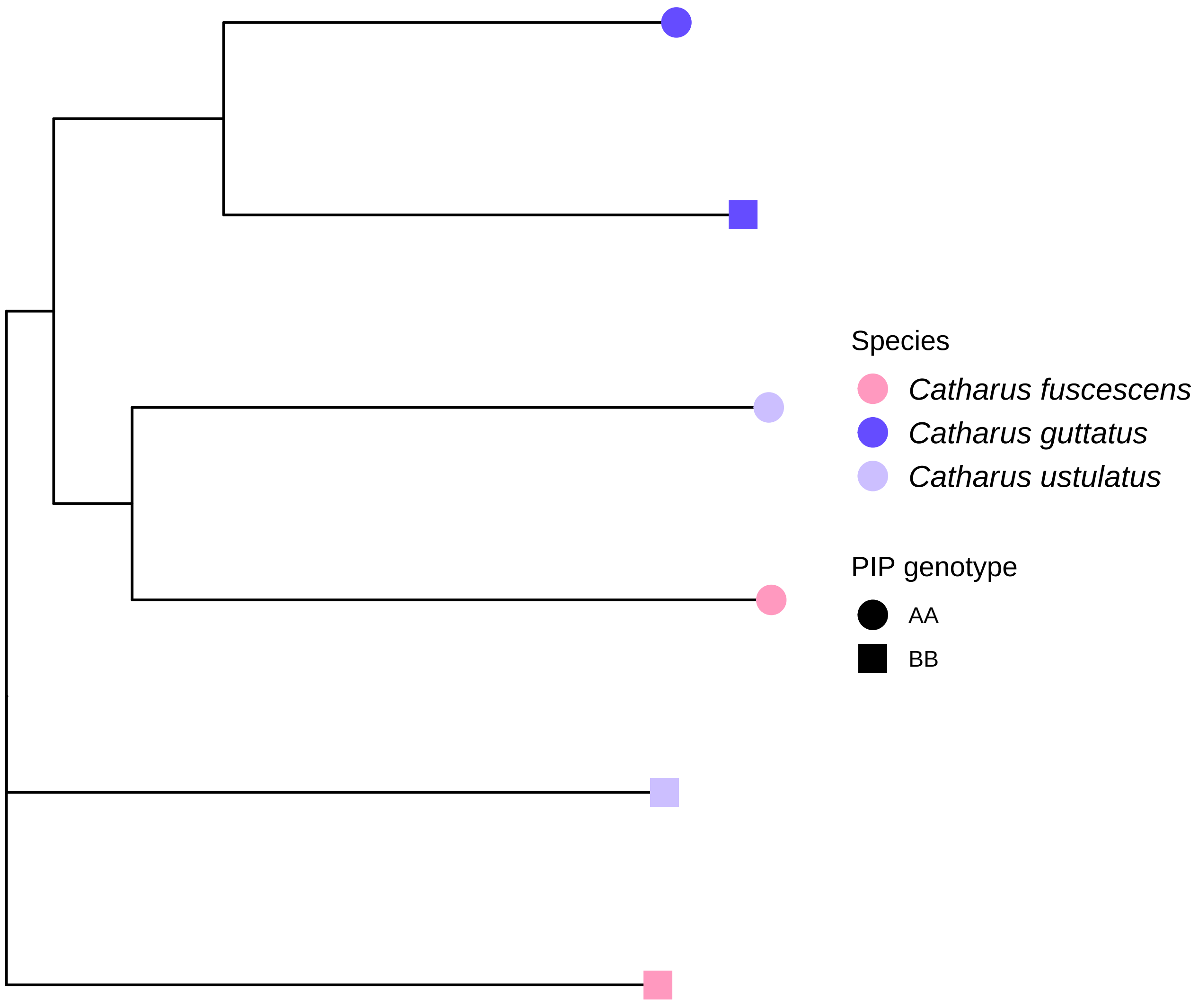

CM020367.1 ii PIP

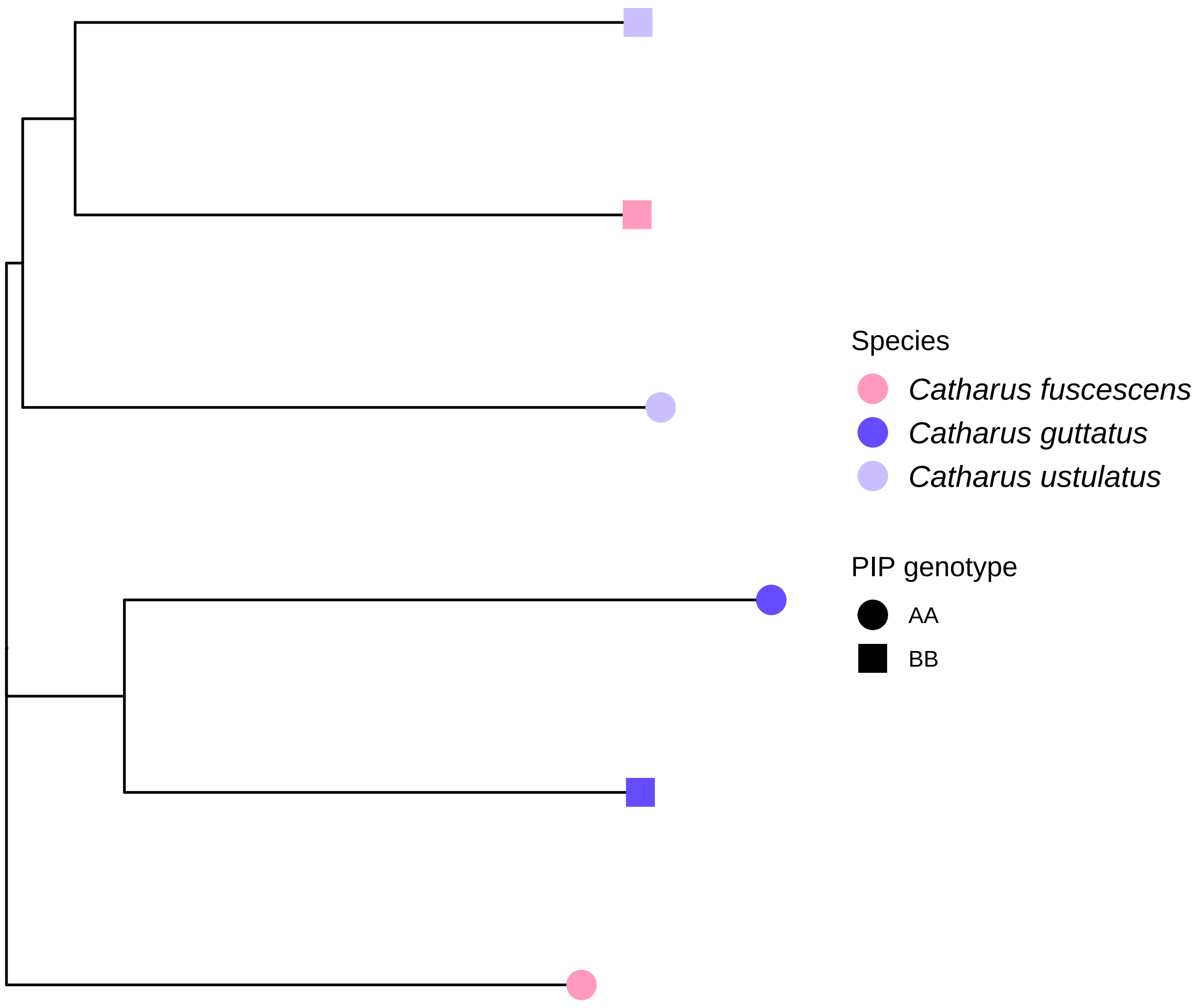

CM020368.1 COMP

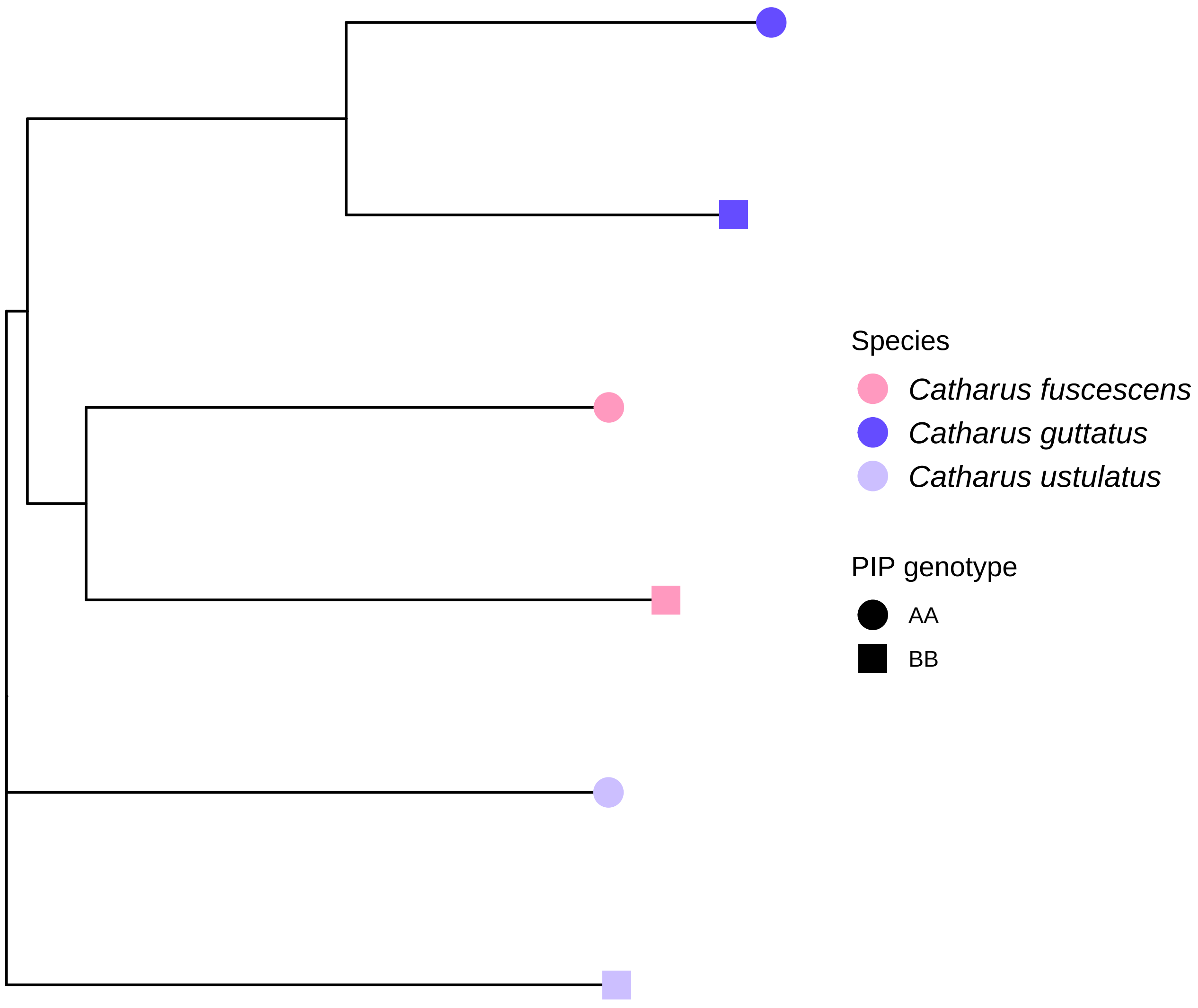

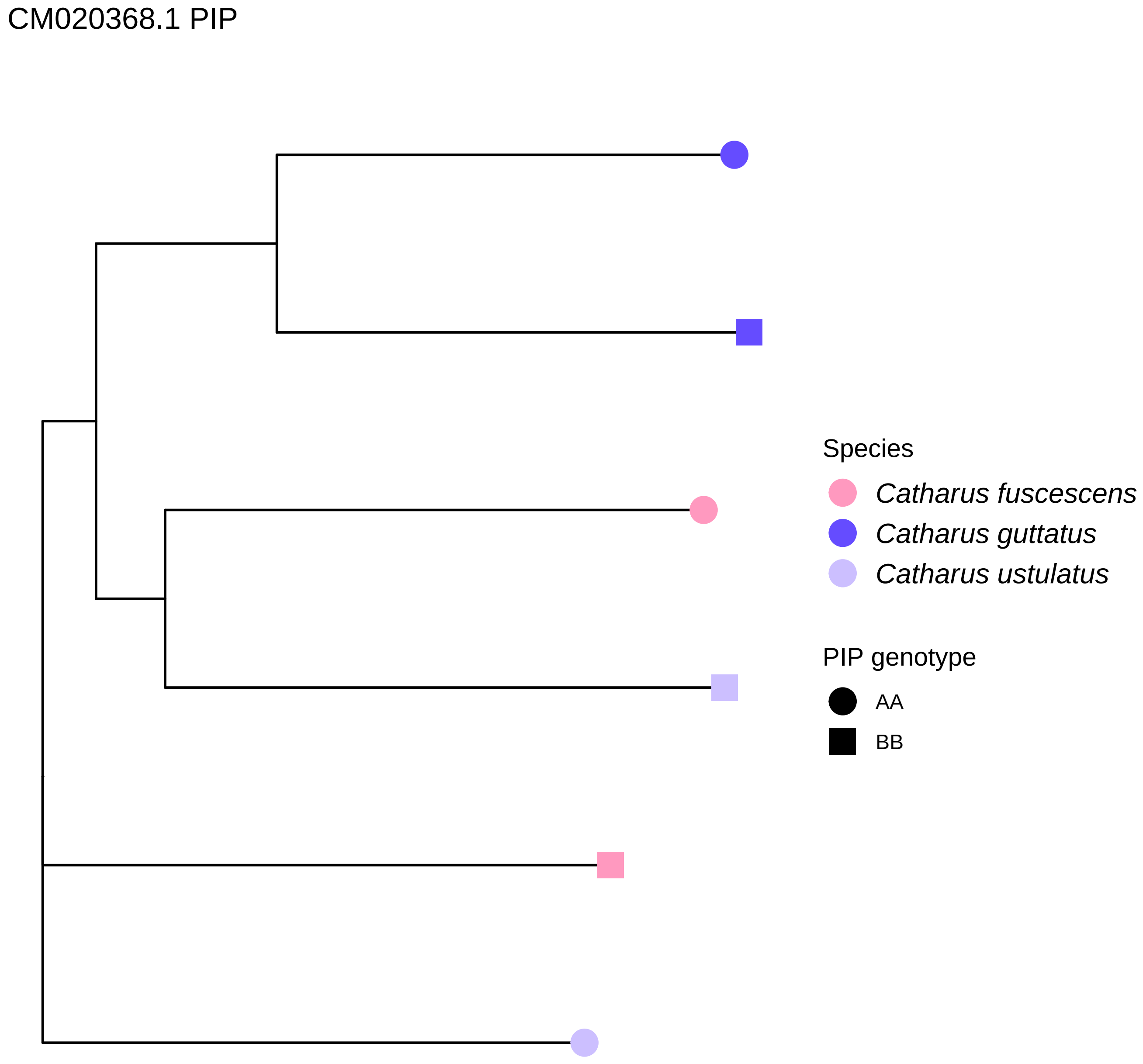
CM020368.1 PIP
Species
Catharus fuscescens
Catharus guttatus
Catharus ustulatus
PIP genotype
AA
BB

### CM020371.1 i PIP

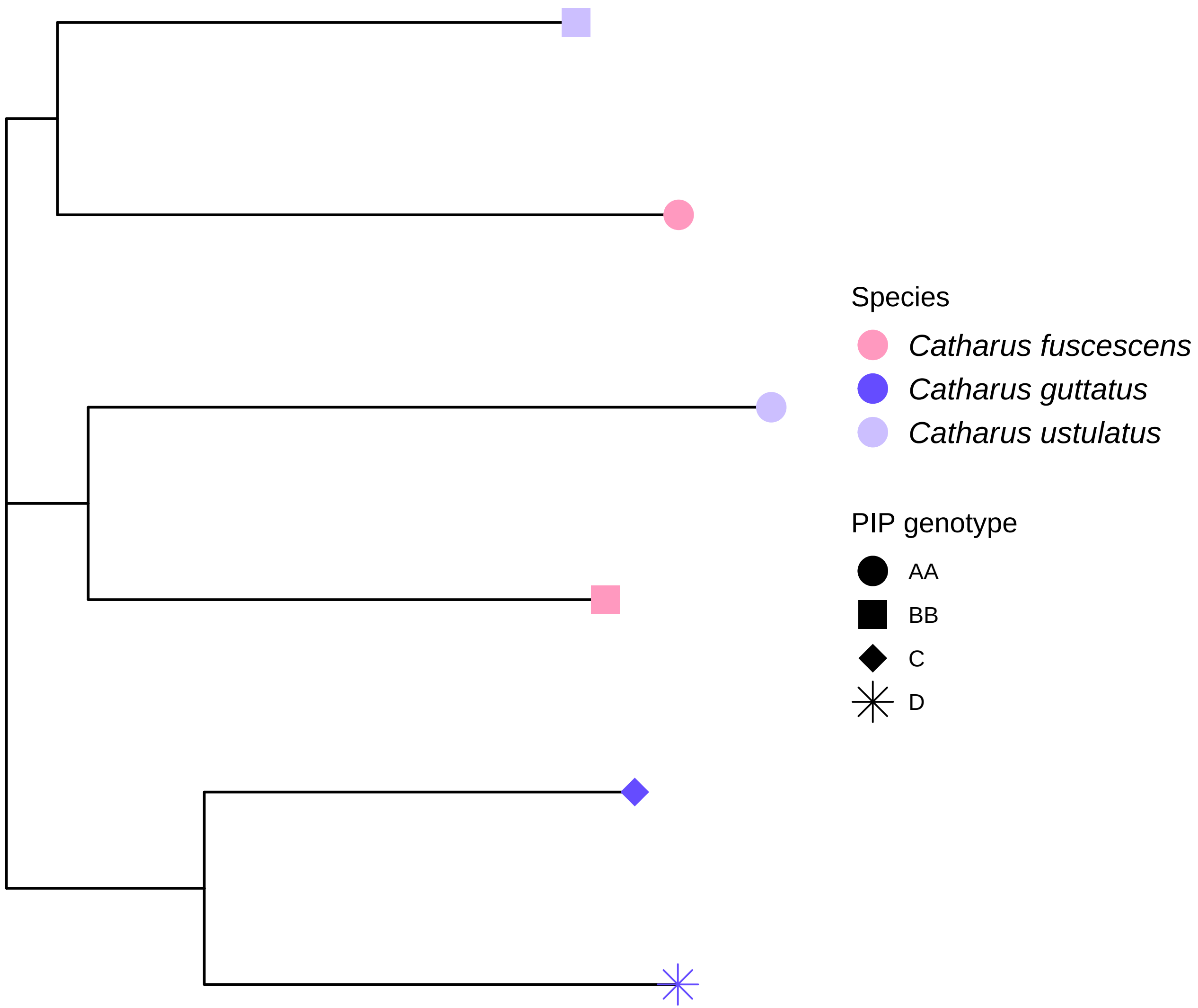

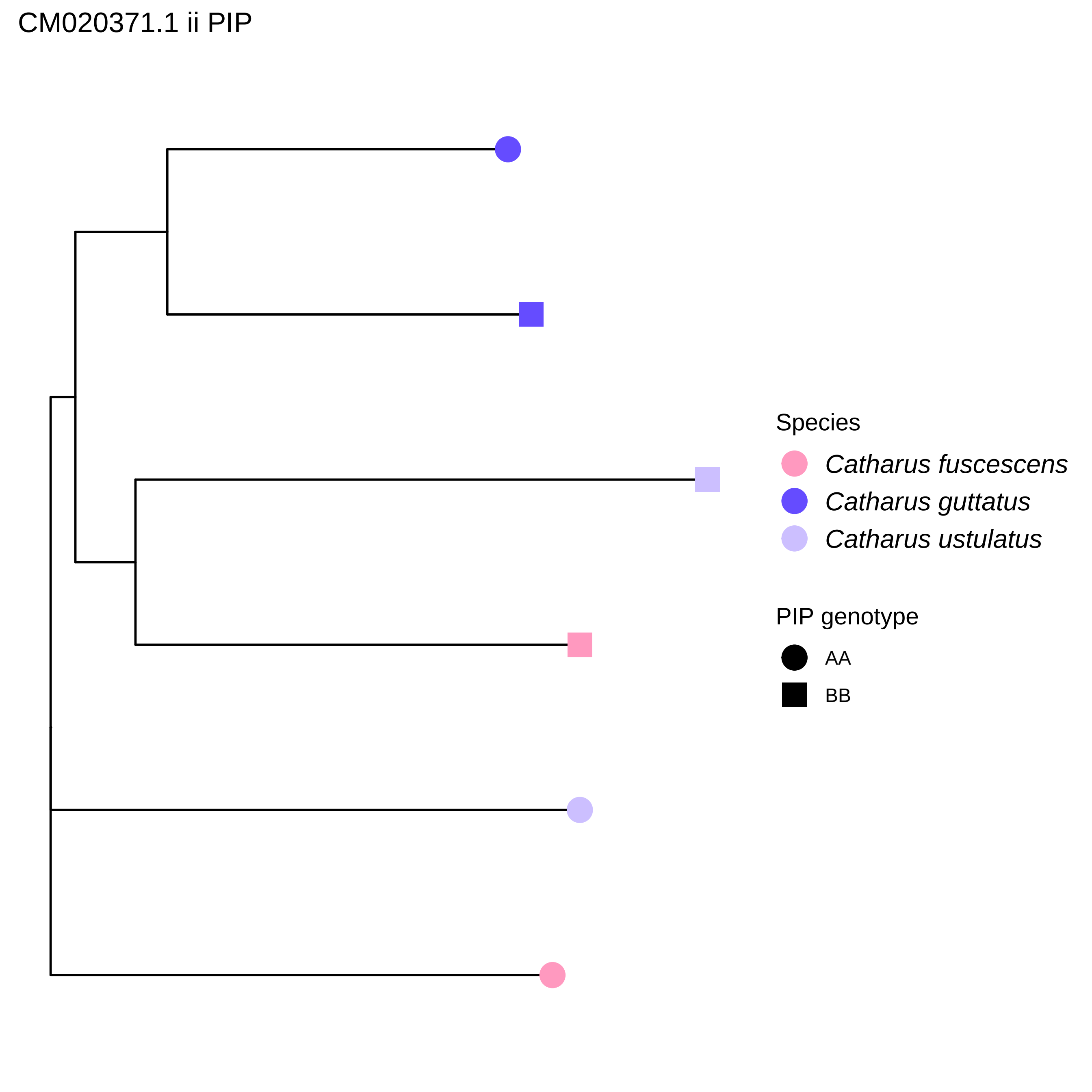
CM020371.1 ii PIP
Species
Catharus fuscescens
Catharus guttatus
Catharus ustulatus
PIP genotype
AA
BB

CM020371.1 iii PIP

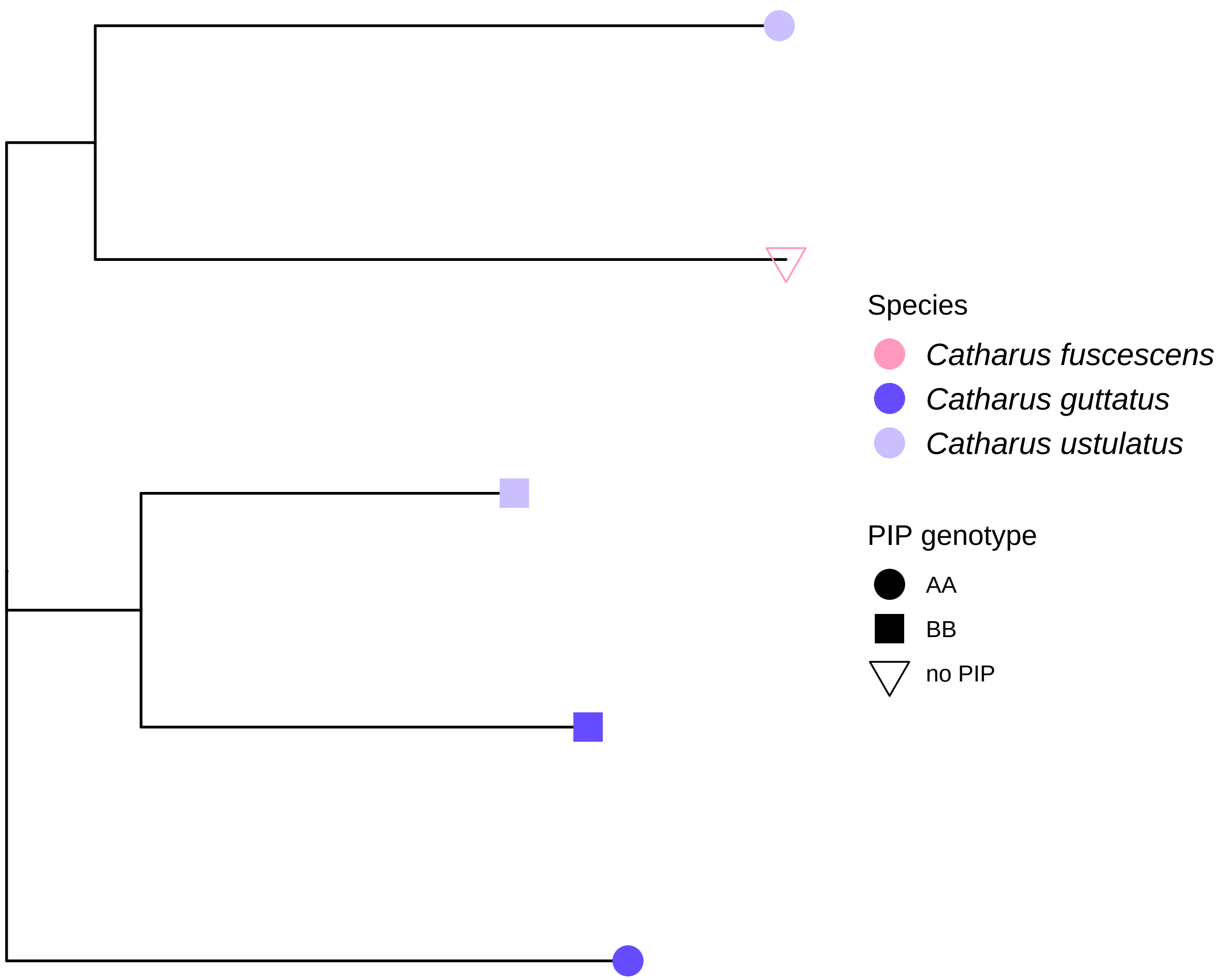

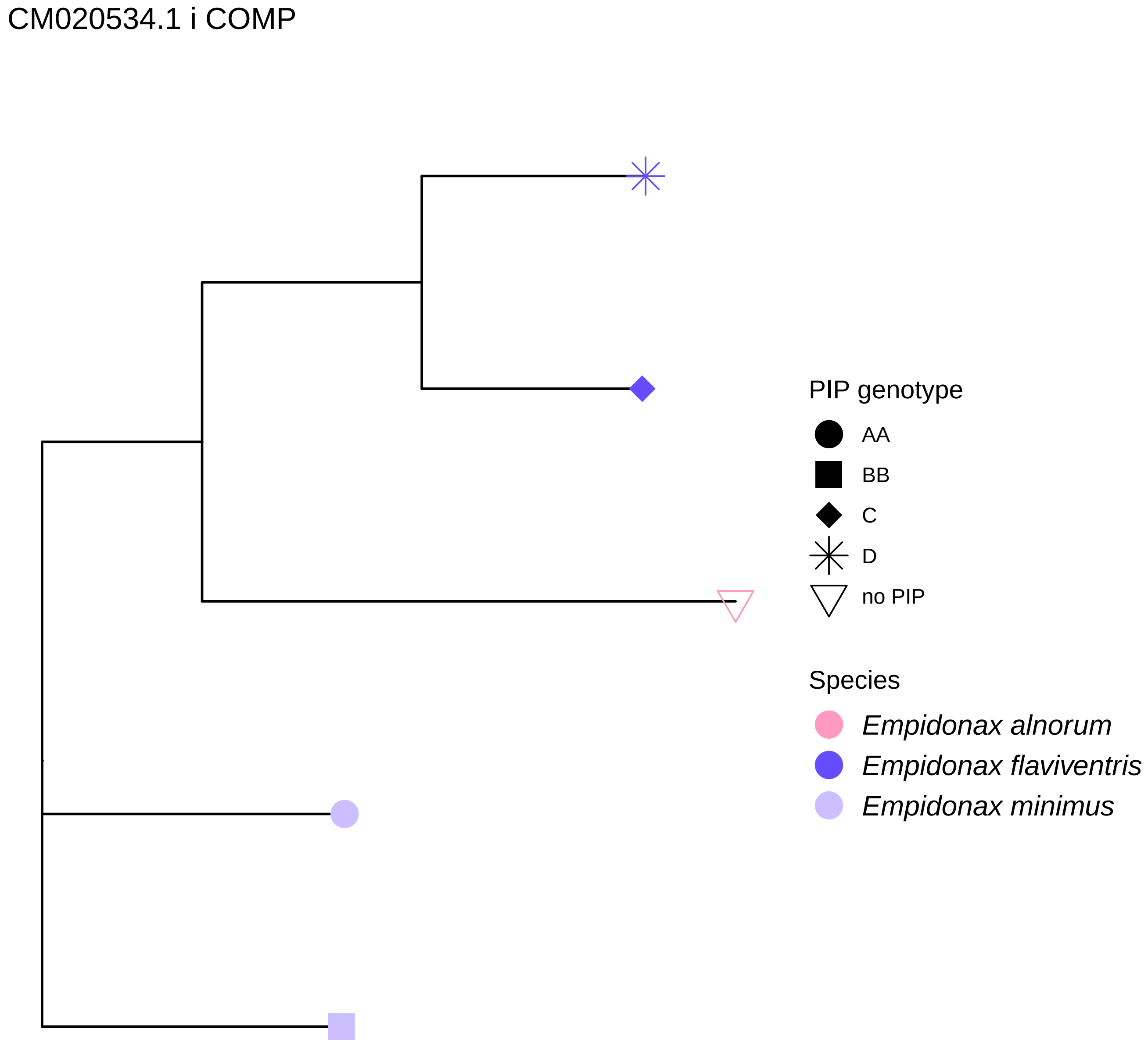
CM020534.1 i COMP
PIP genotype
AA
BB
C
D
no PIP
Species
Empidonax alnorum
Empidonax flaviventris
Empidonax minimus

CM020534.1 i PIP

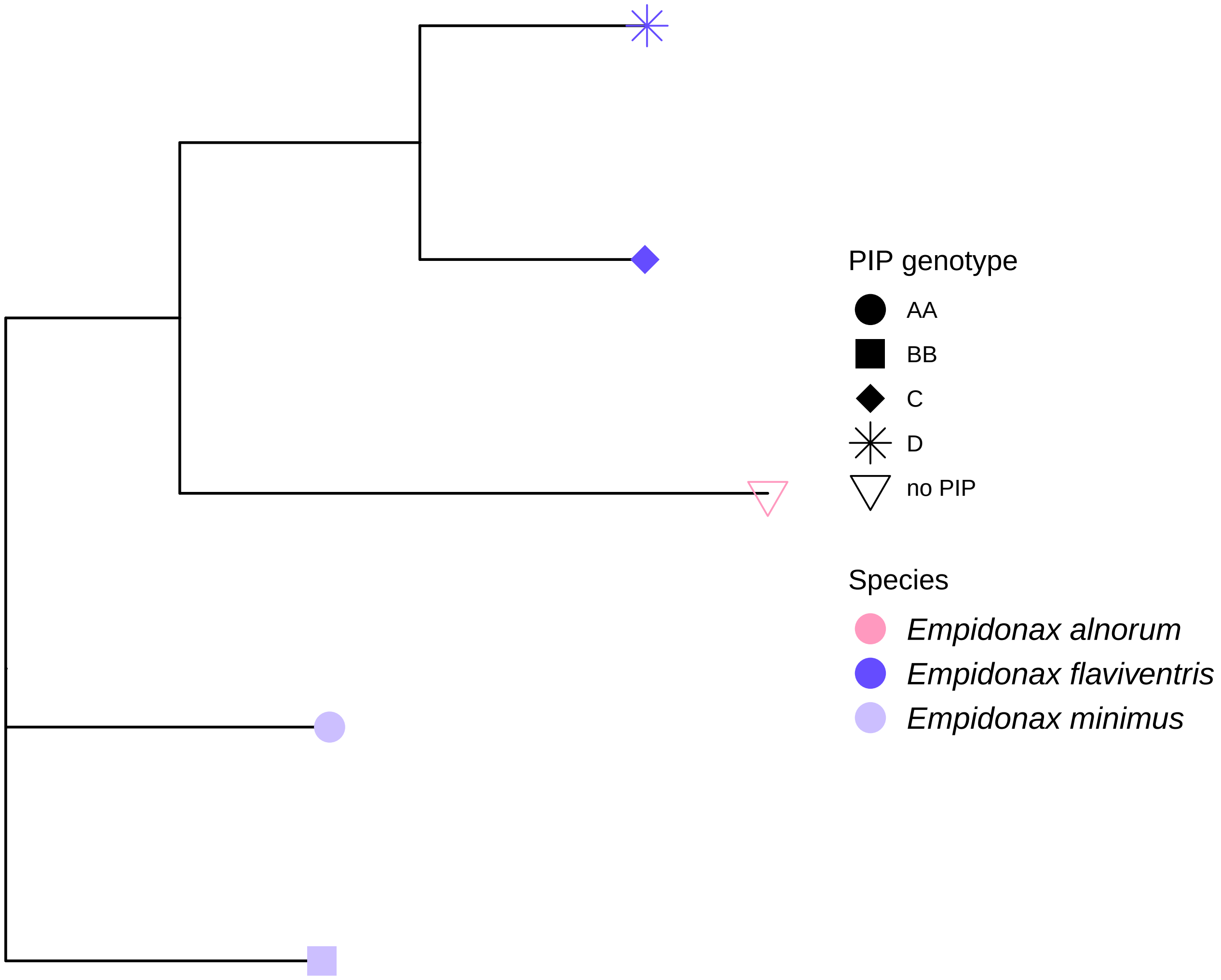

CM020534.1 ii PIP

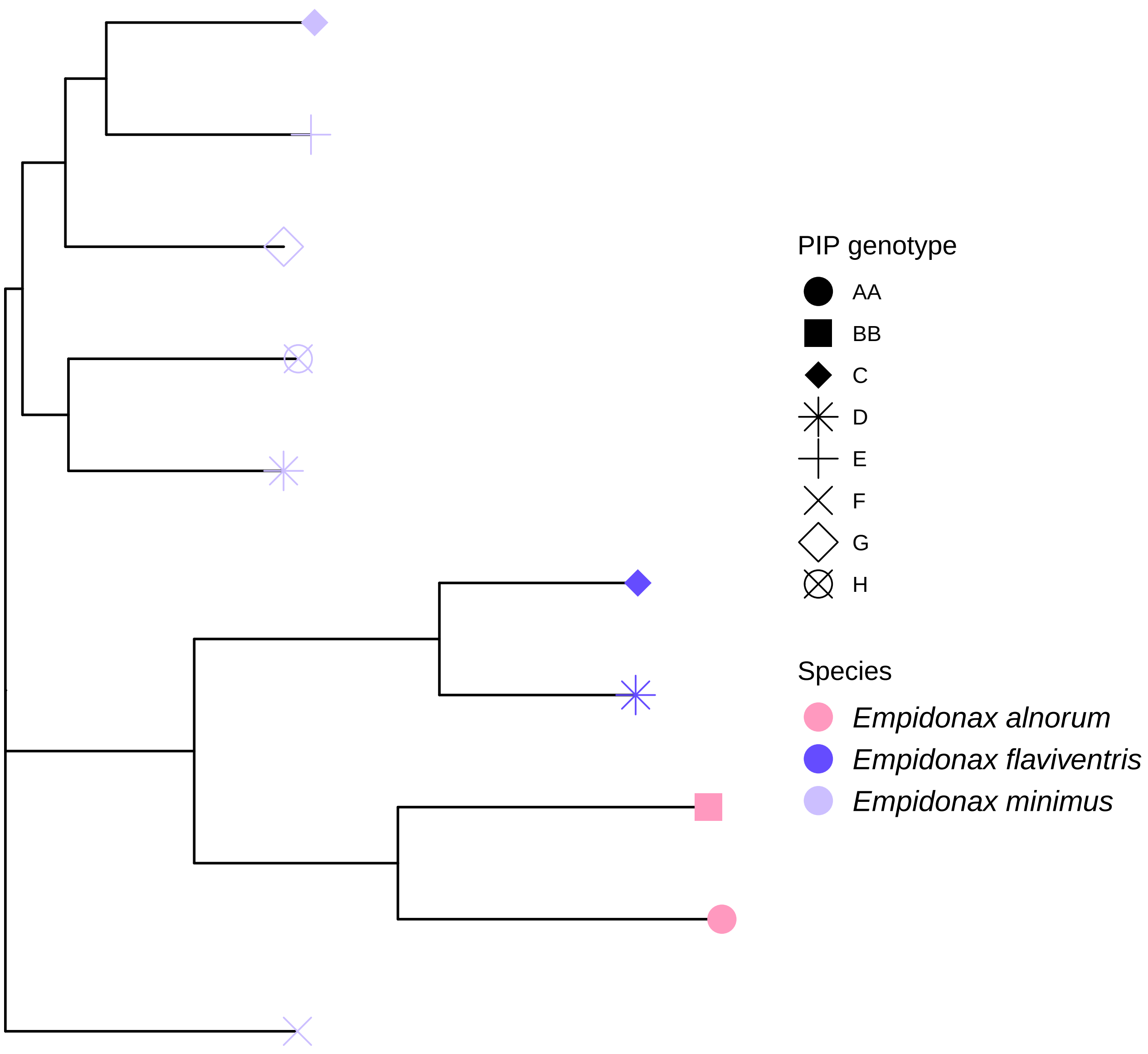

CM020543.1 COMP

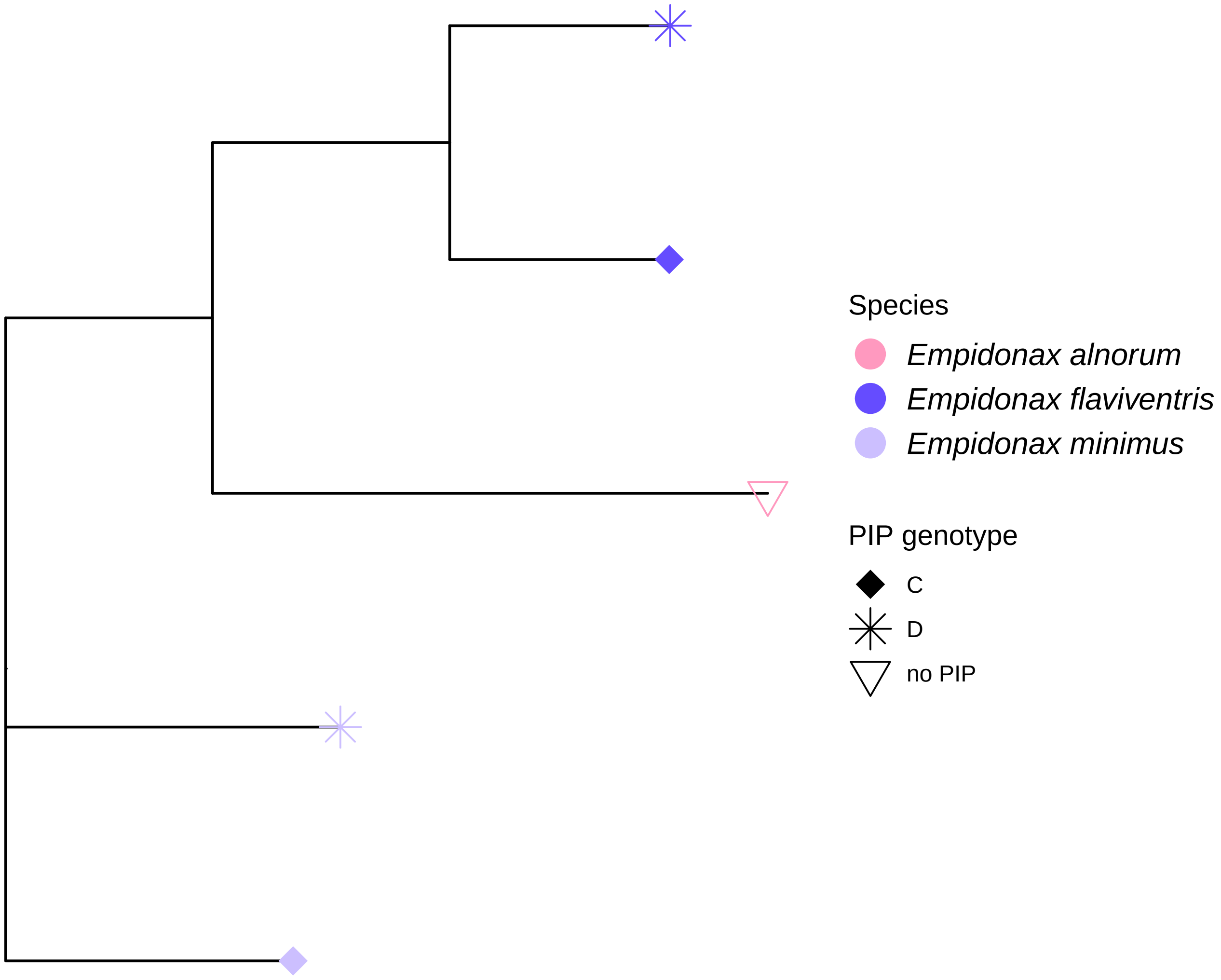

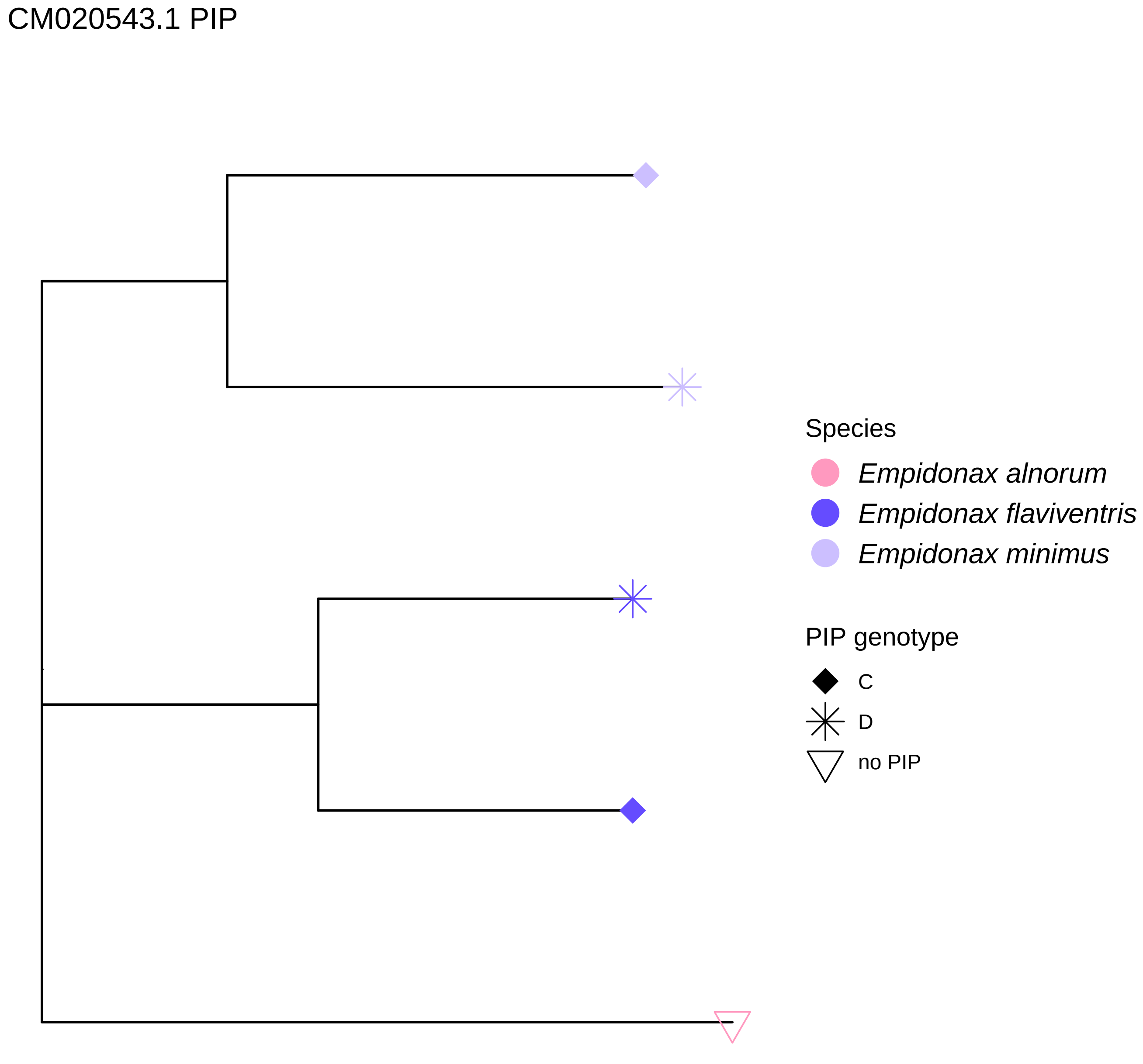
CM020543.1 PIP
Species
Empidonax alnorum
Empidonax flaviventris
Empidonax minimus
PIP genotype
C
D
no PIP

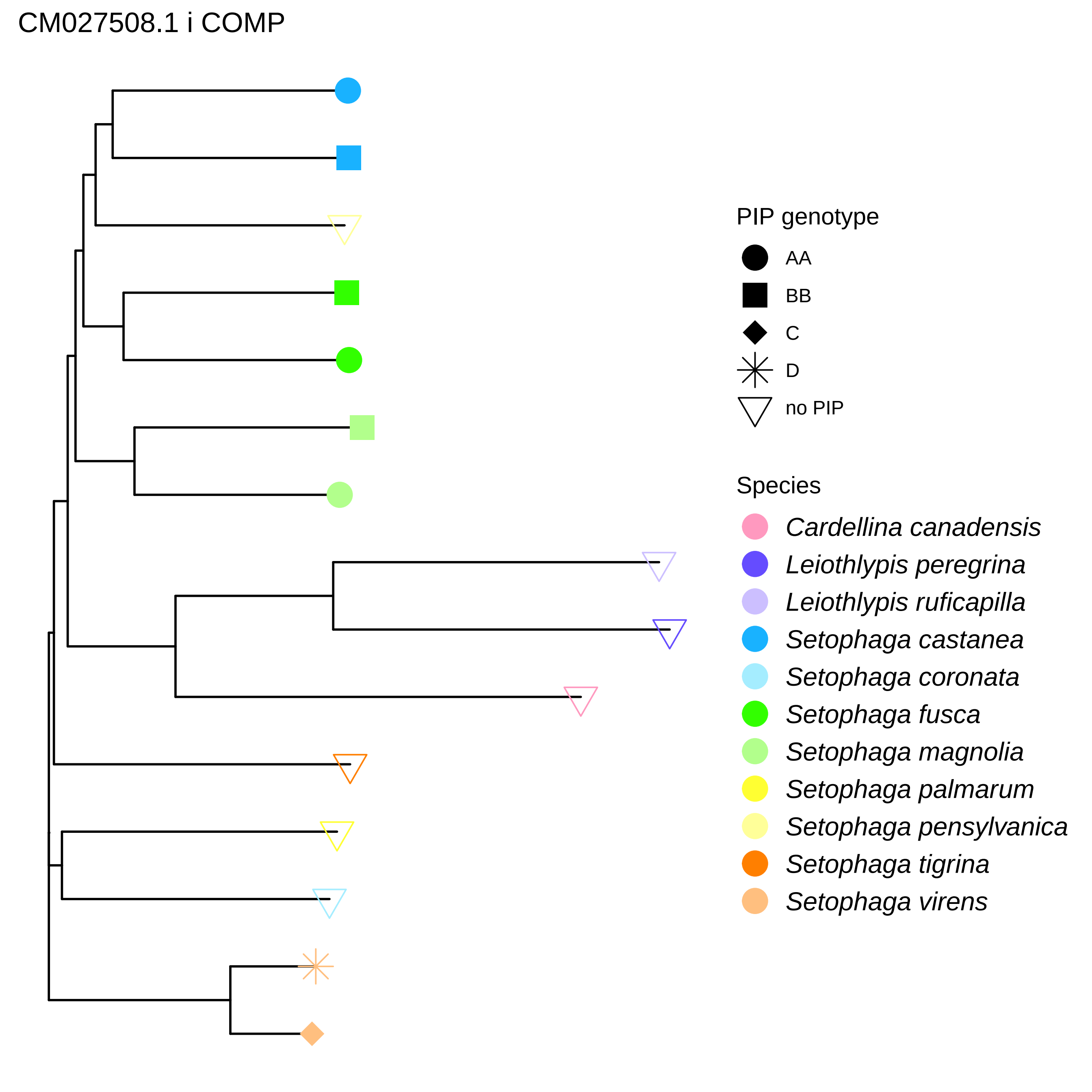
CM027508.1 i COMP
PIP genotype
AA
BB
C
D
no PIP
Species
Cardellina canadensis
Leiothlypis peregrina
Leiothlypis ruficapilla
Setophaga castanea
Setophaga coronata
Setophaga fusca
Setophaga magnolia
Setophaga palmarum
Setophaga pensylvanica
Setophaga tigrina
Setophaga virens

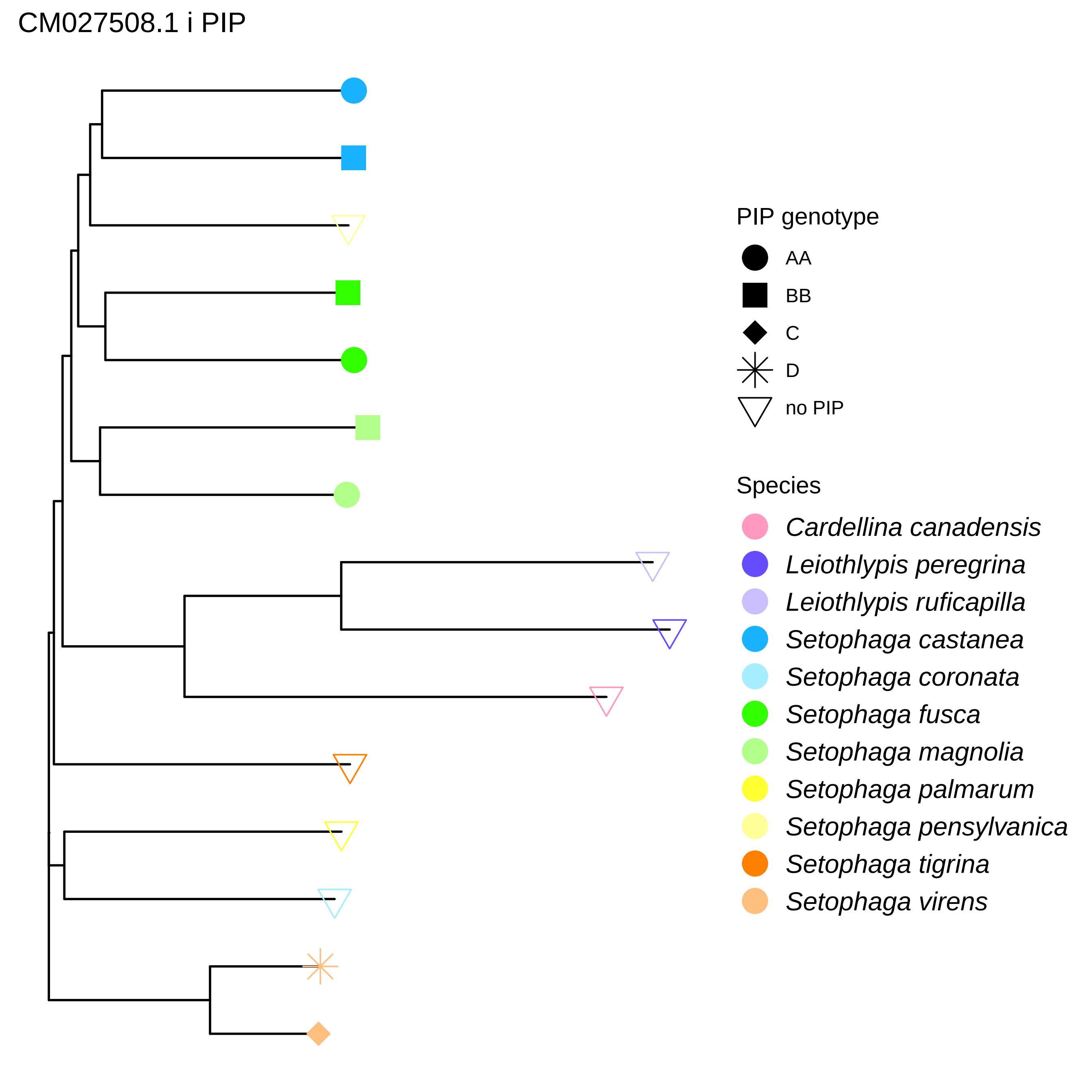
CM027508.1 i PIP
PIP genotype
AA
BB
C
D
no PIP
Species
Cardellina canadensis
Leiothlypis peregrina
Leiothlypis ruficapilla
Setophaga castanea
Setophaga coronata
Setophaga fusca
Setophaga magnolia
Setophaga palmarum
Setophaga pensylvanica
Setophaga tigrina
Setophaga virens

CM027508.1 ii PIP

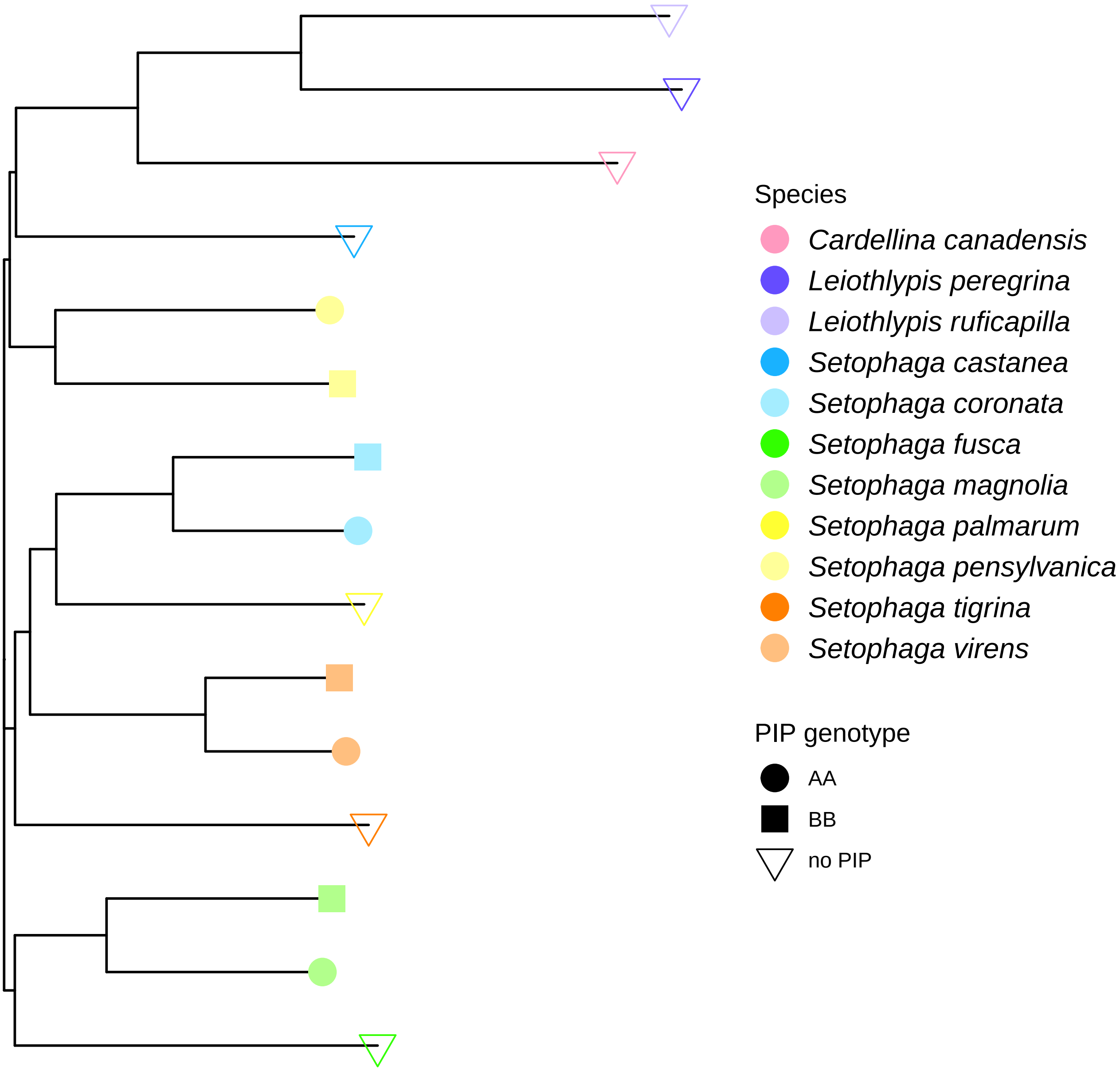

CM027510.1 COMP

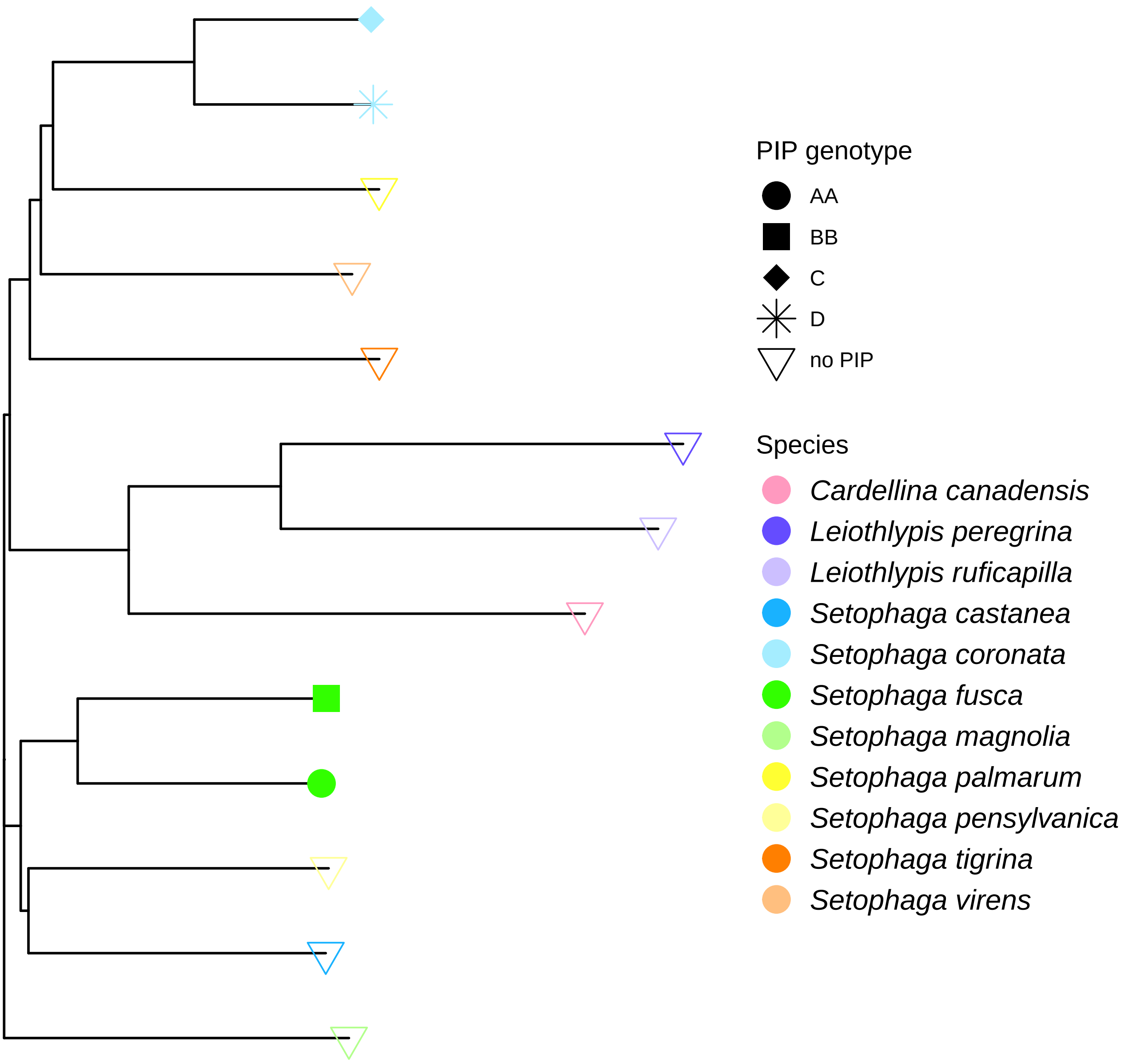

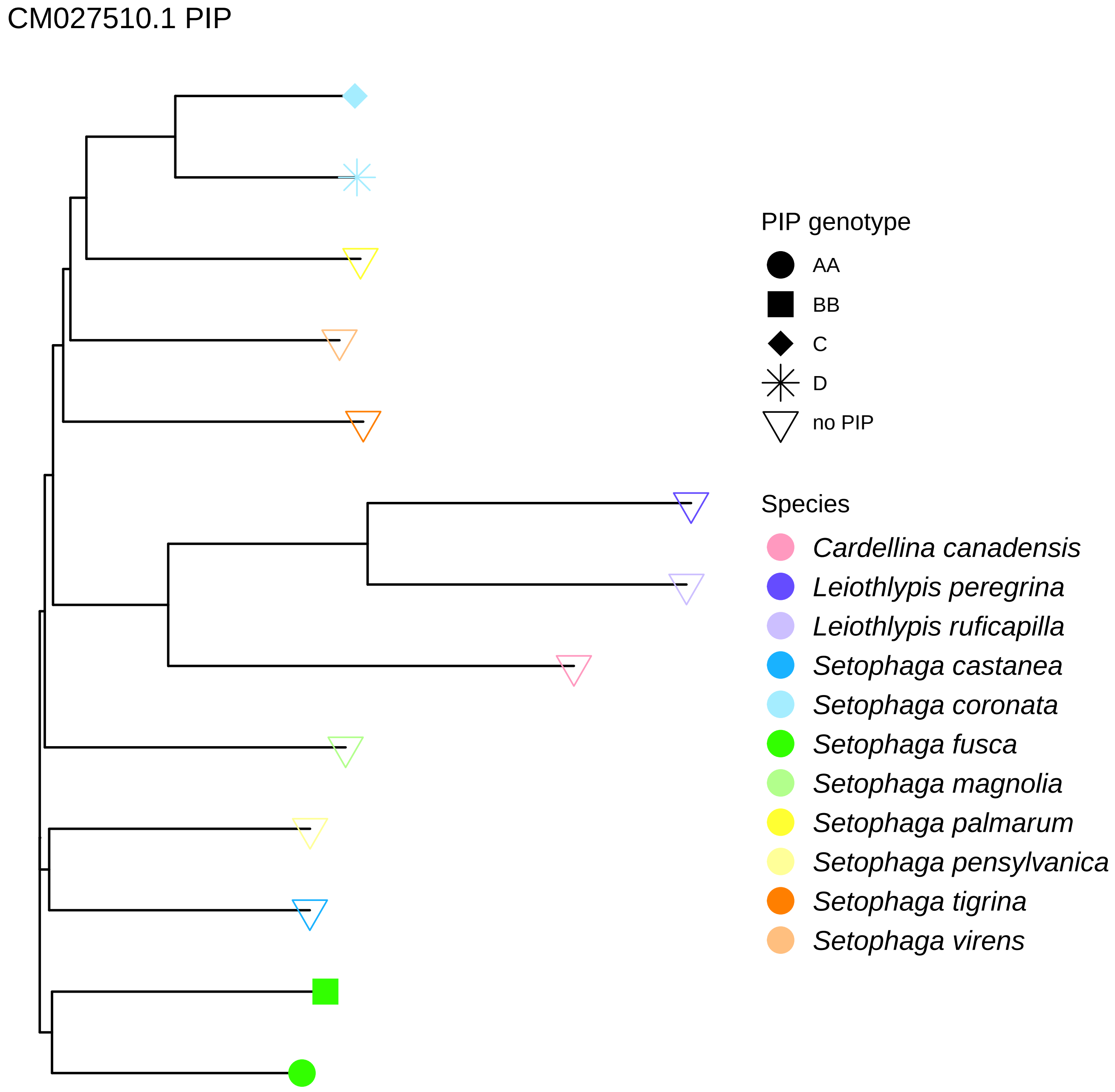
CM027510.1 PIP
PIP genotype
AA
BB
C
D
no PIP
Species
Cardellina canadensis
Leiothlypis peregrina
Leiothlypis ruficapilla
Setophaga castanea
Setophaga coronata
Setophaga fusca
Setophaga magnolia
Setophaga palmarum
Setophaga pensylvanica
Setophaga tigrina
Setophaga virens

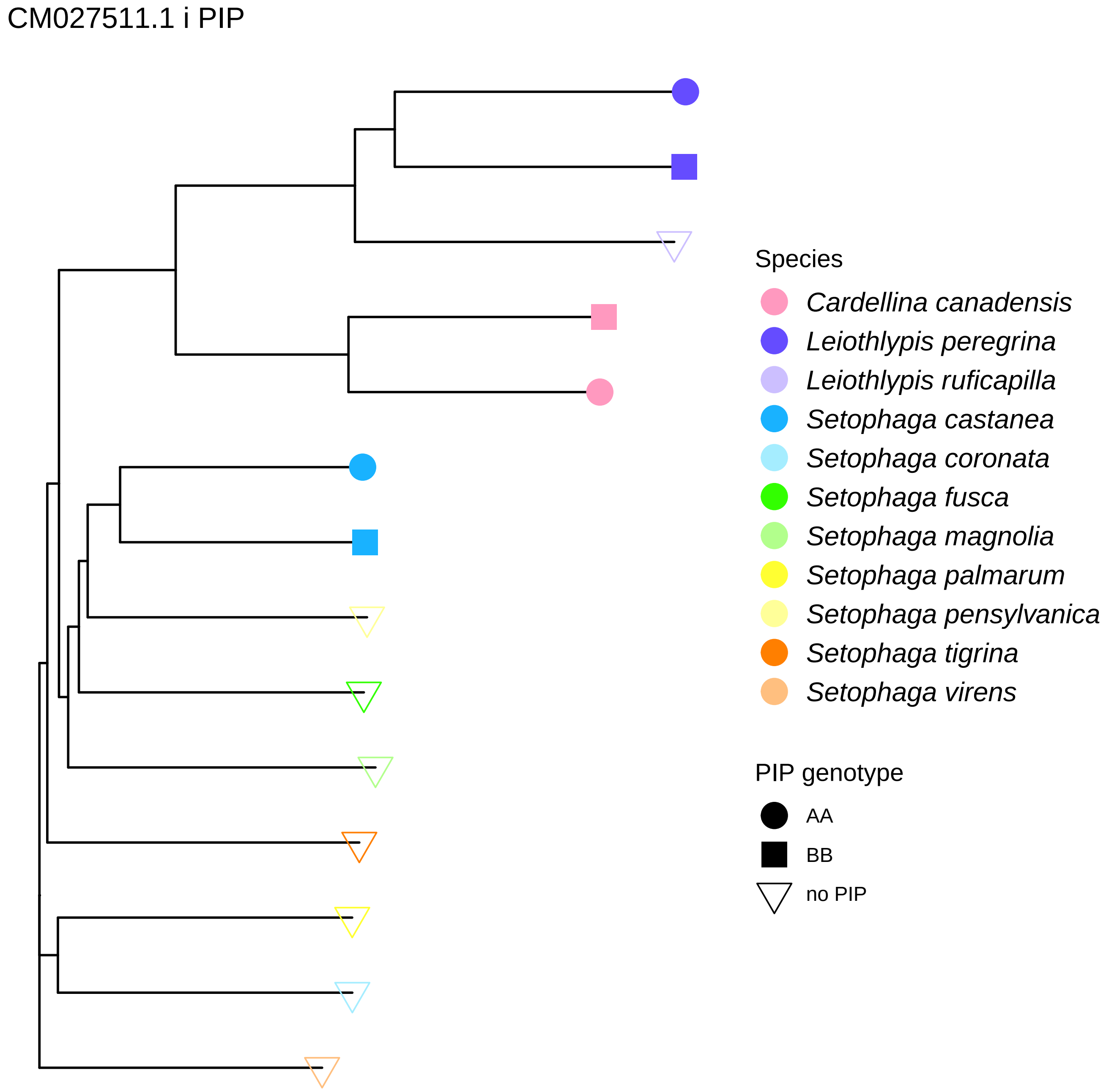
CM027511.1 i PIP
Species
Cardellina canadensis
Leiothlypis peregrina
Leiothlypis ruficapilla
Setophaga castanea
Setophaga coronata
Setophaga fusca
Setophaga magnolia
Setophaga palmarum
Setophaga pensylvanica
Setophaga tigrina
Setophaga virens
PIP genotype
AA
BB
no PIP

CM027511.1 ii PIP

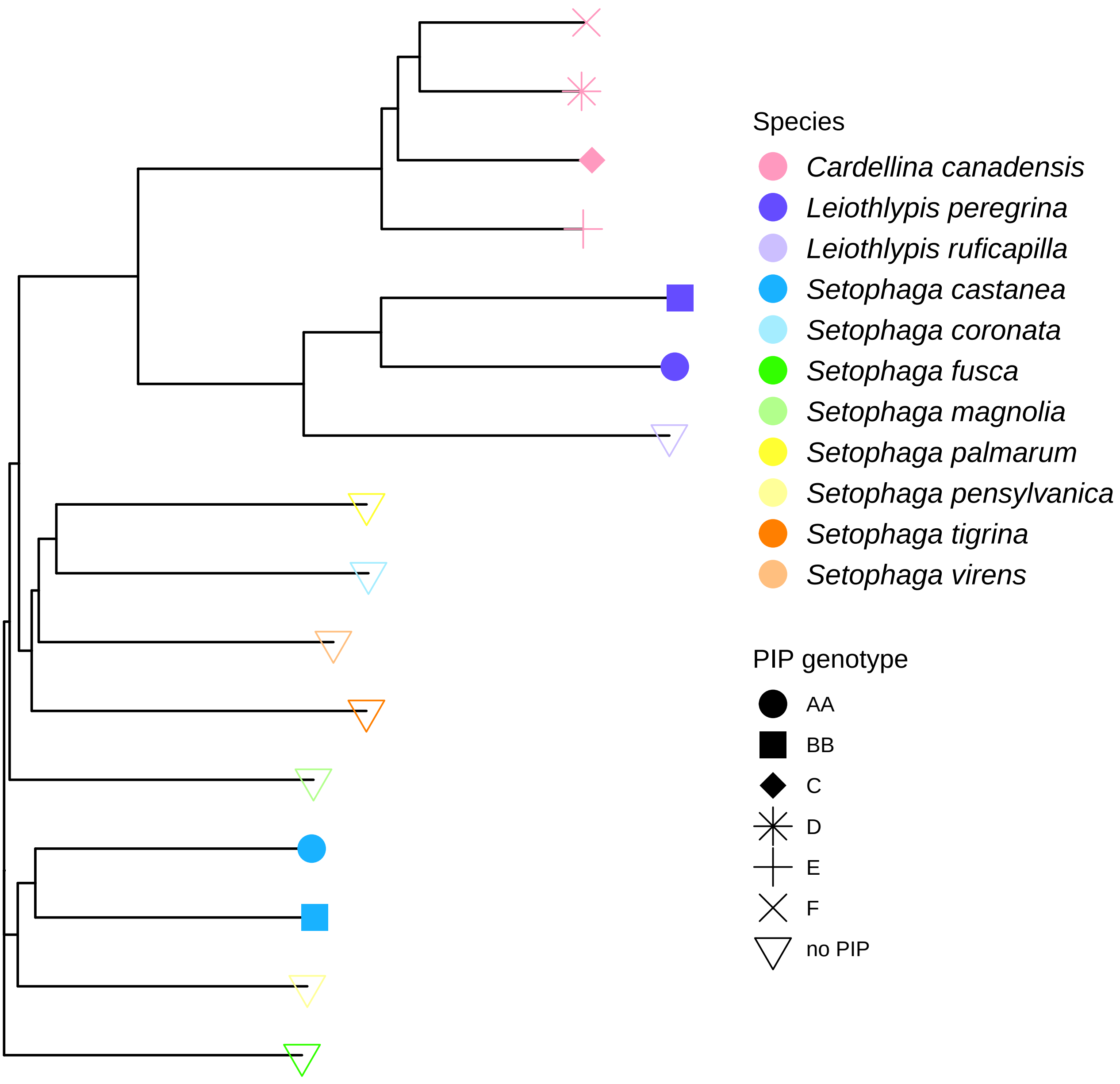

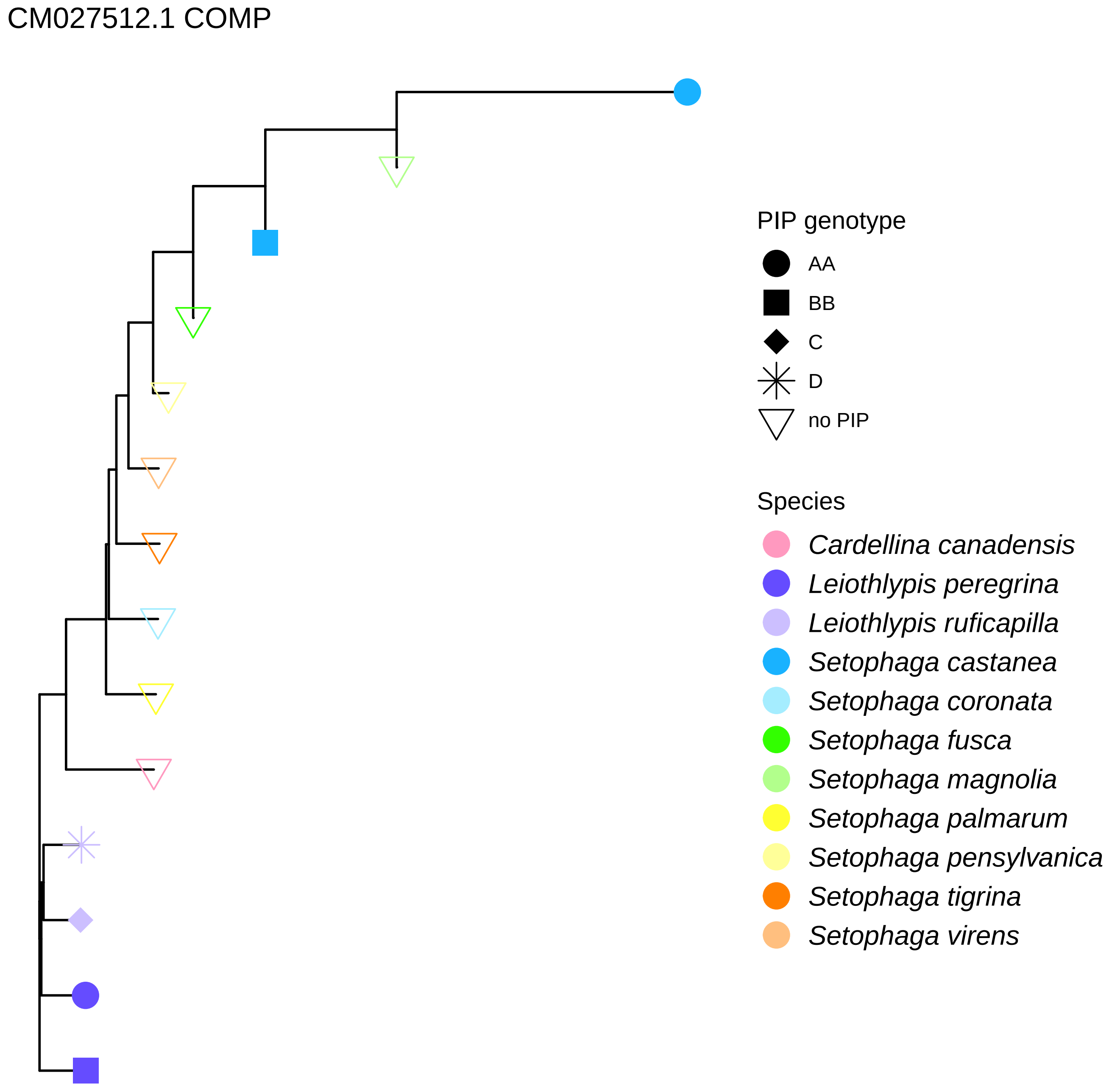
CM027512.1 COMP
PIP genotype
AA
BB
C
D
no PIP
Species
Cardellina canadensis
Leiothlypis peregrina
Leiothlypis ruficapilla
Setophaga castanea
Setophaga coronata
Setophaga fusca
Setophaga magnolia
Setophaga palmarum
Setophaga pensylvanica
Setophaga tigrina
Setophaga virens

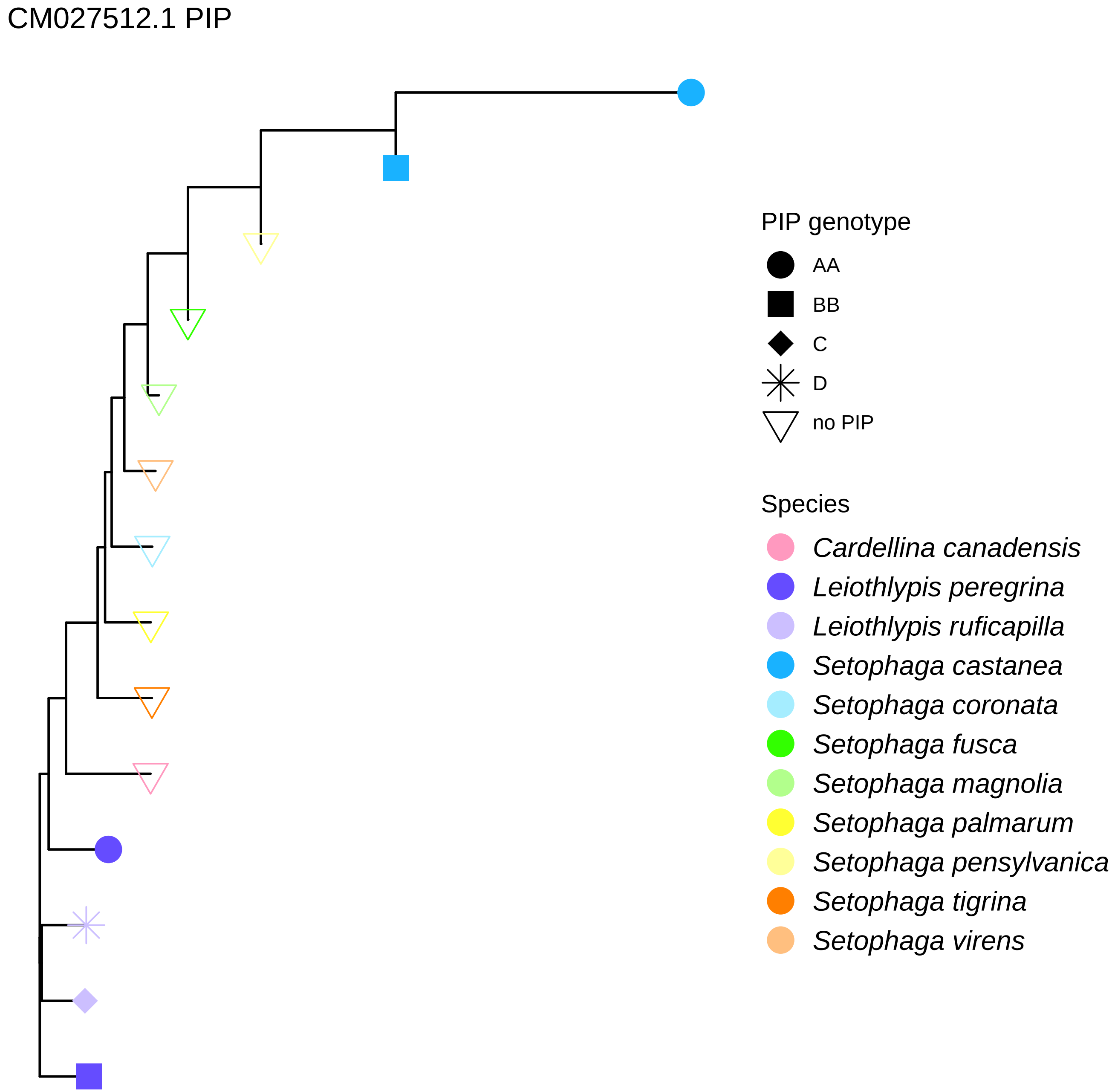

CM027512.1 PIP
PIP genotype
AA
BB
C
D
no PIP
Species
Cardellina canadensis
Leiothlypis peregrina
Leiothlypis ruficapilla
Setophaga castanea
Setophaga coronata
Setophaga fusca
Setophaga magnolia
Setophaga palmarum
Setophaga pensylvanica
Setophaga tigrina
Setophaga virens

CM027513.1 COMP

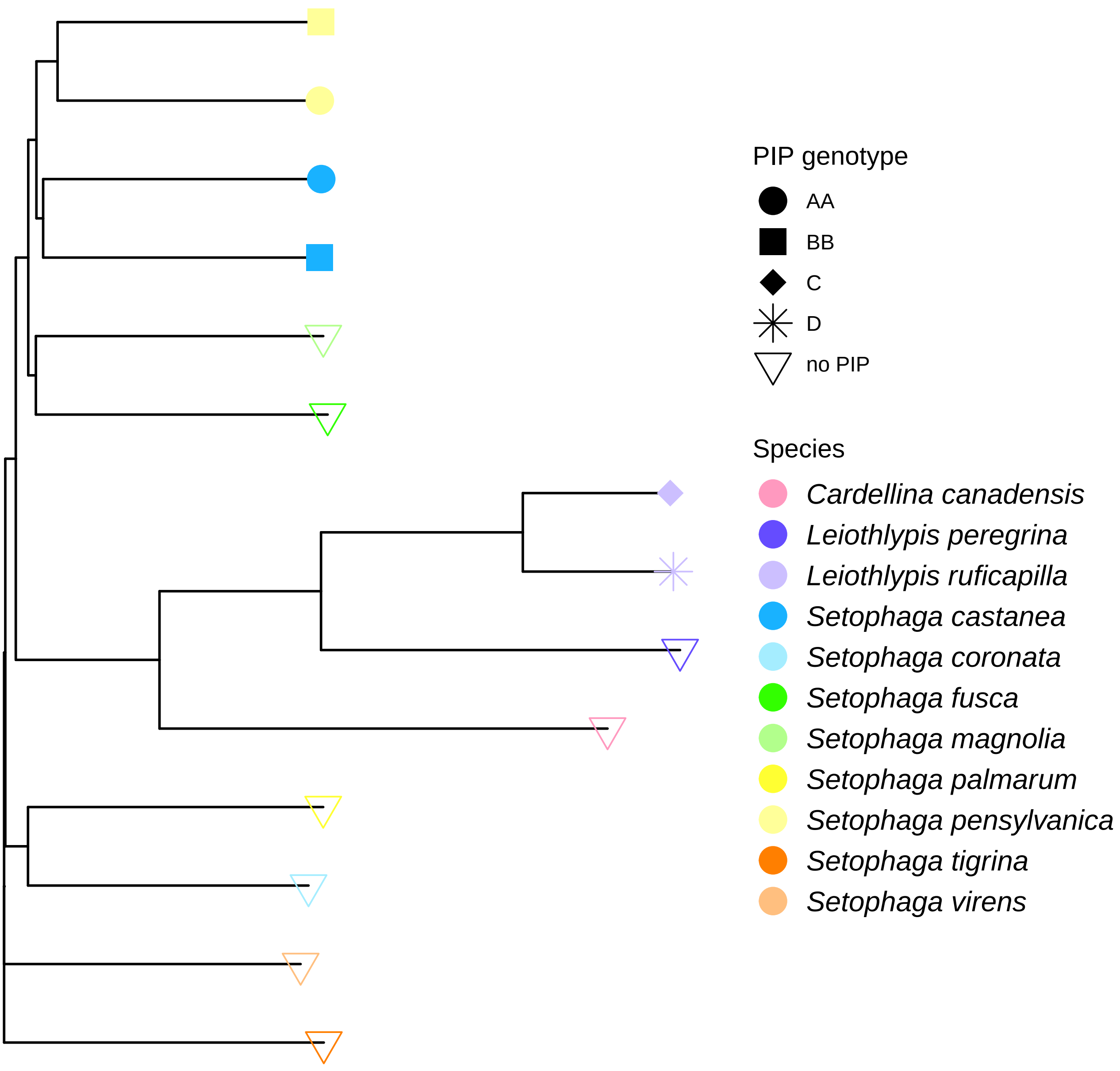

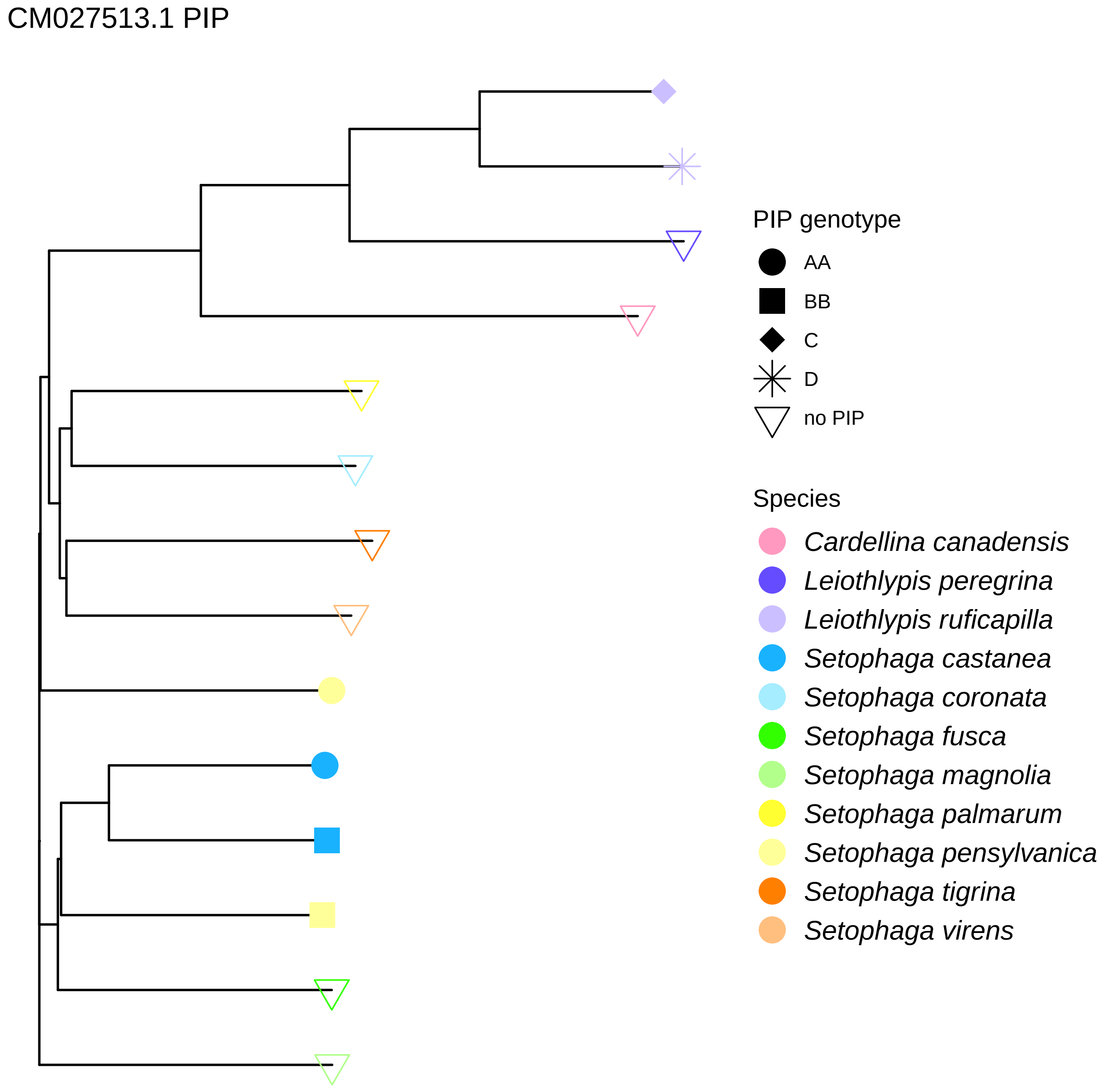
CM027513.1 PIP
PIP genotype
AA
BB
C
D
no PIP
Species
Cardellina canadensis
Leiothlypis peregrina
Leiothlypis ruficapilla
Setophaga castanea
Setophaga coronata
Setophaga fusca
Setophaga magnolia
Setophaga palmarum
Setophaga pensylvanica
Setophaga tigrina
Setophaga virens

### CM027518.1 COMP

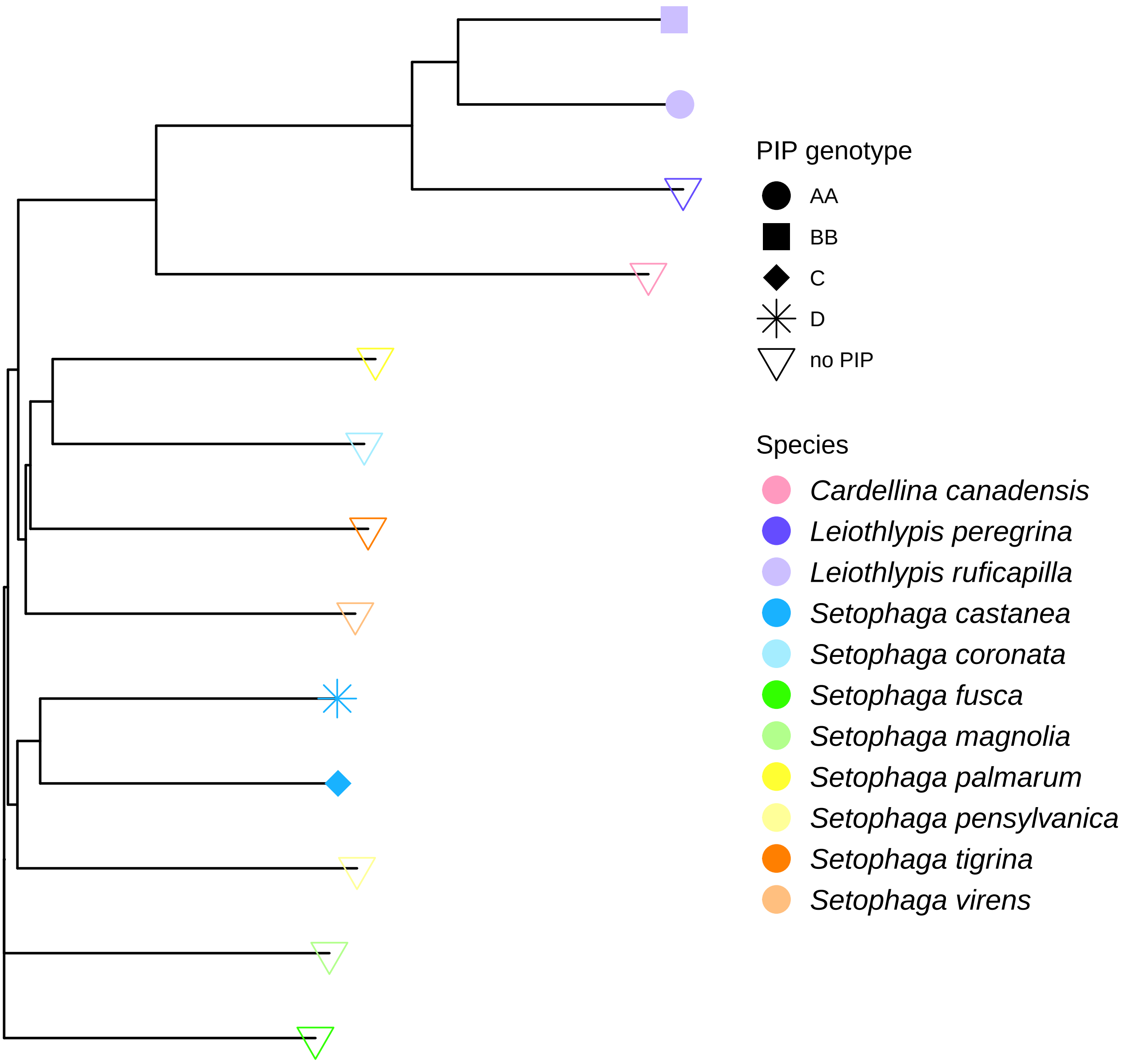

### CM027518.1 PIP

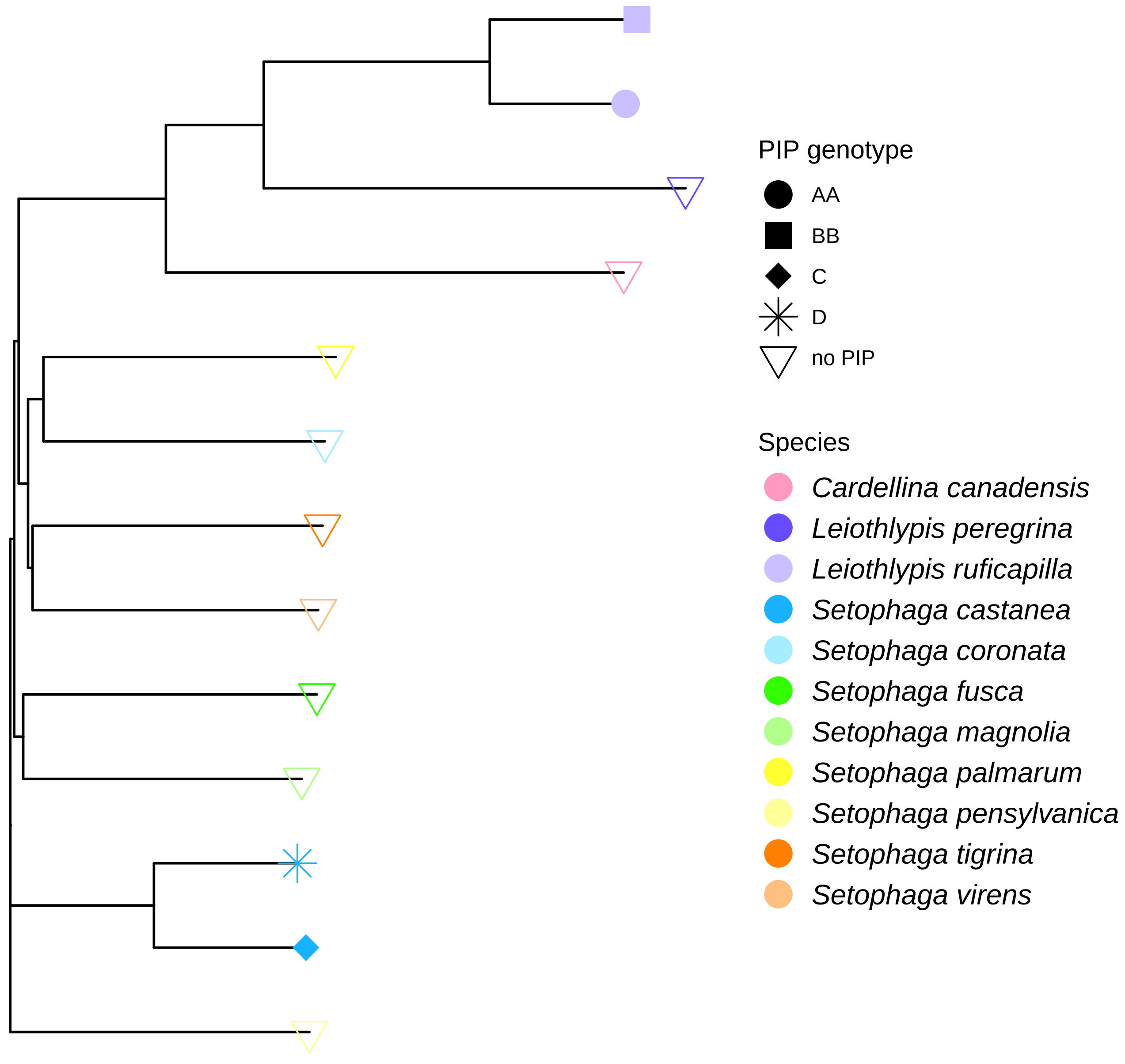

CM027530.1 COMP

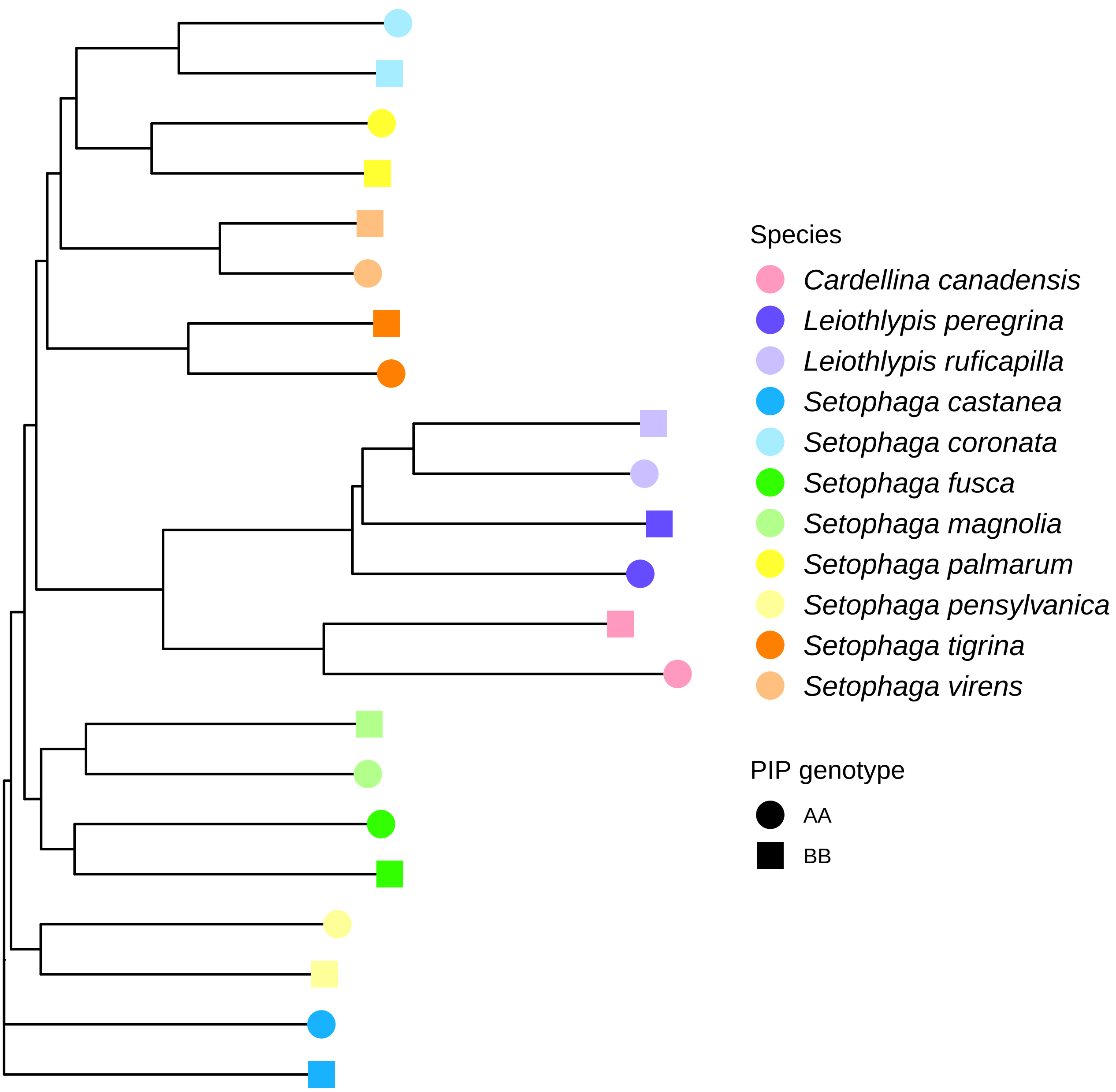

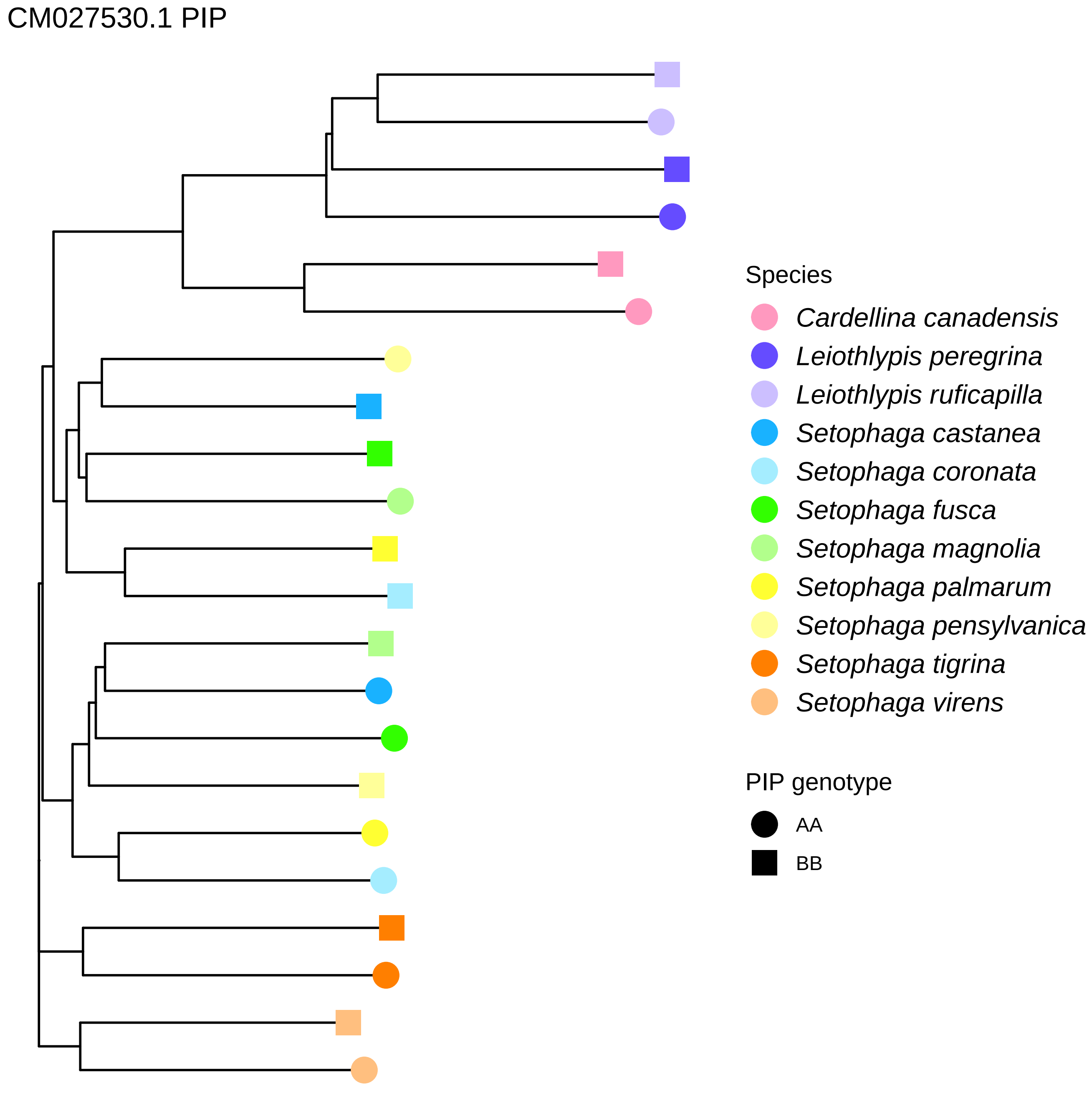
CM027530.1 PIP
Species
Cardellina canadensis
Leiothlypis peregrina
Leiothlypis ruficapilla
Setophaga castanea
Setophaga coronata
Setophaga fusca
Setophaga magnolia
Setophaga palmarum
Setophaga pensylvanica
Setophaga tigrina
Setophaga virens
PIP genotype
AA
BB

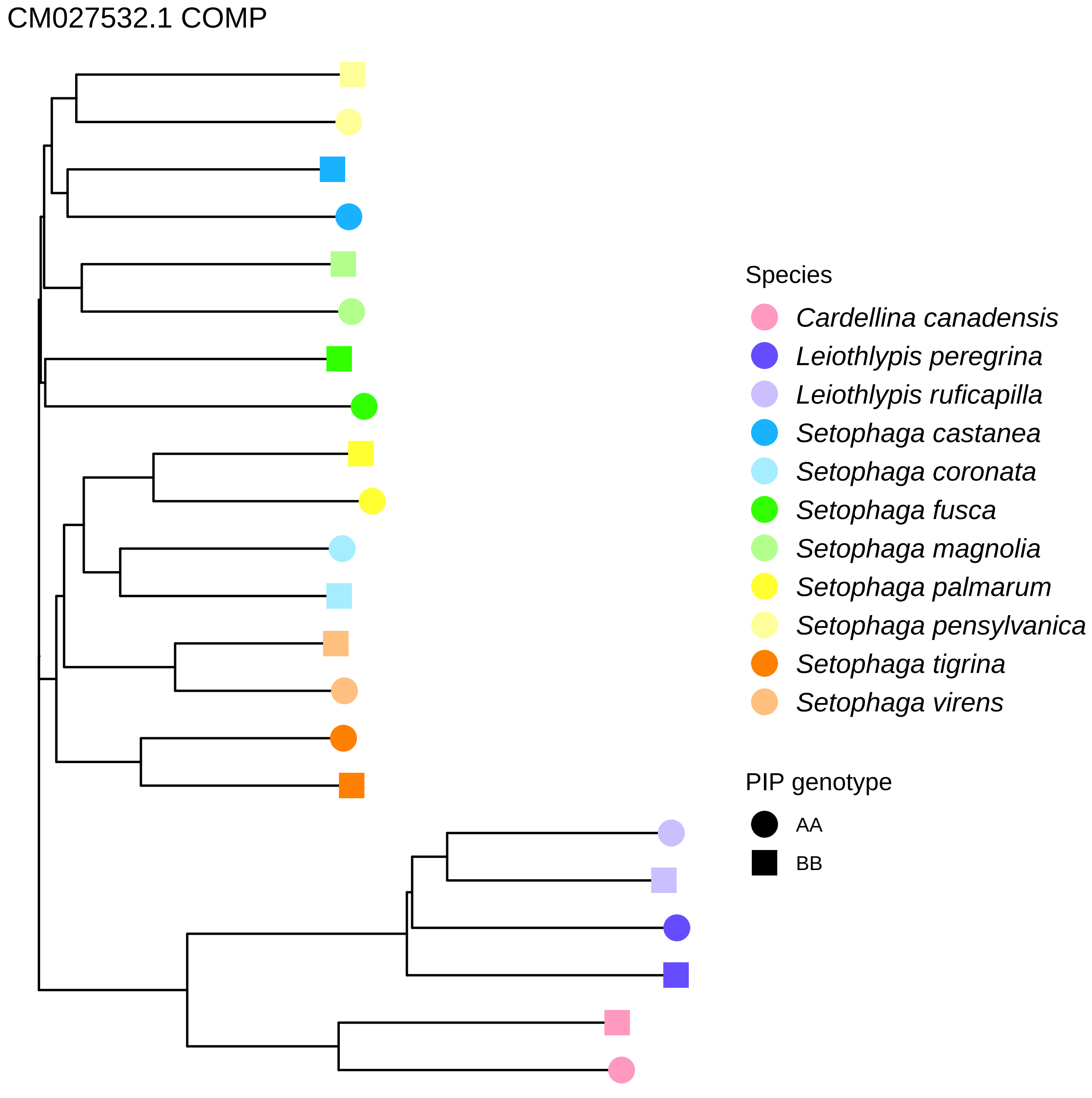
CM027532.1 COMP
Species
Cardellina canadensis
Leiothlypis peregrina
Leiothlypis ruficapilla
Setophaga castanea
Setophaga coronata
Setophaga fusca
Setophaga magnolia
Setophaga palmarum
Setophaga pensylvanica
Setophaga tigrina
Setophaga virens
PIP genotype
AA
BB

CM027532.1 PIP

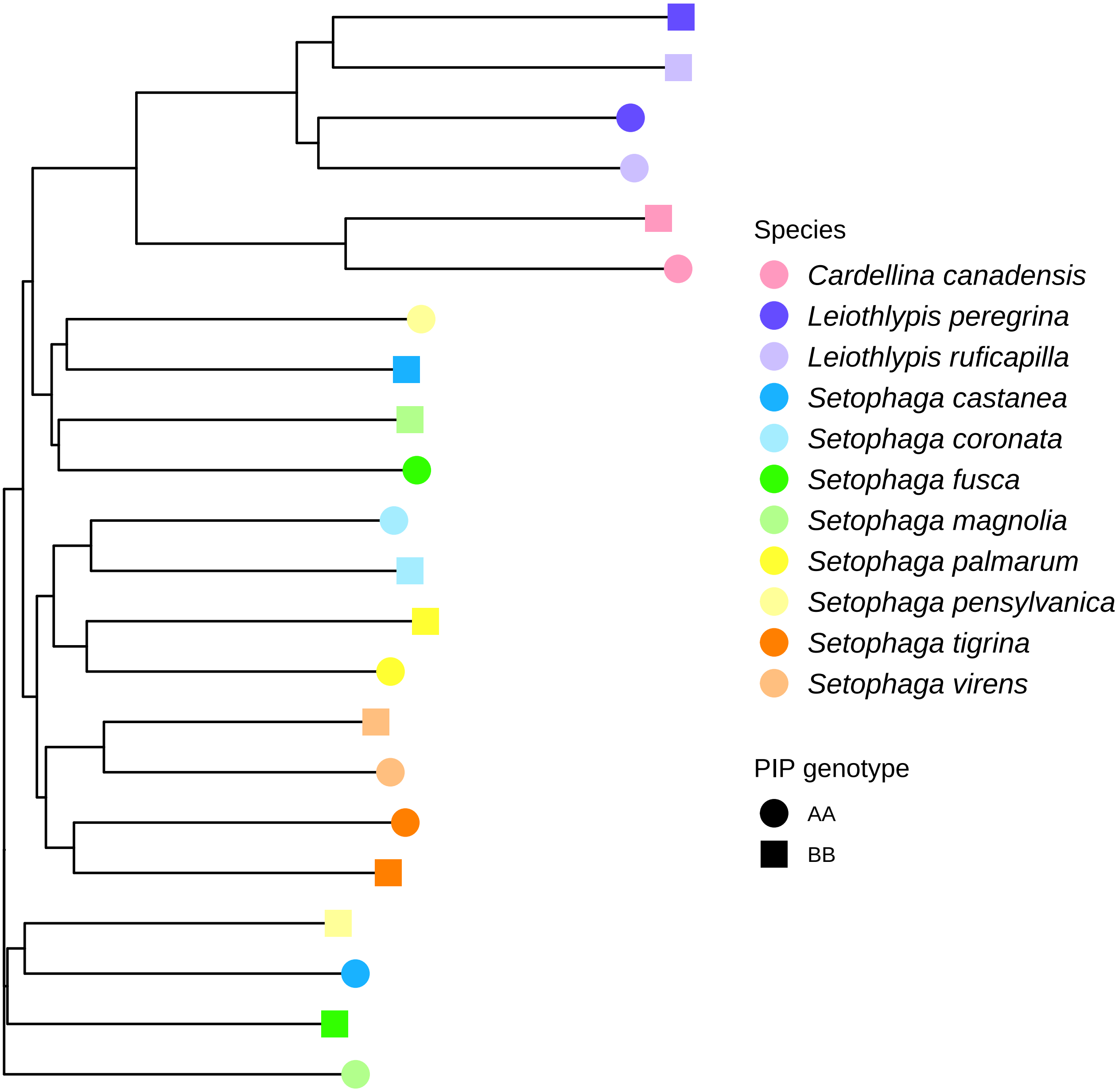

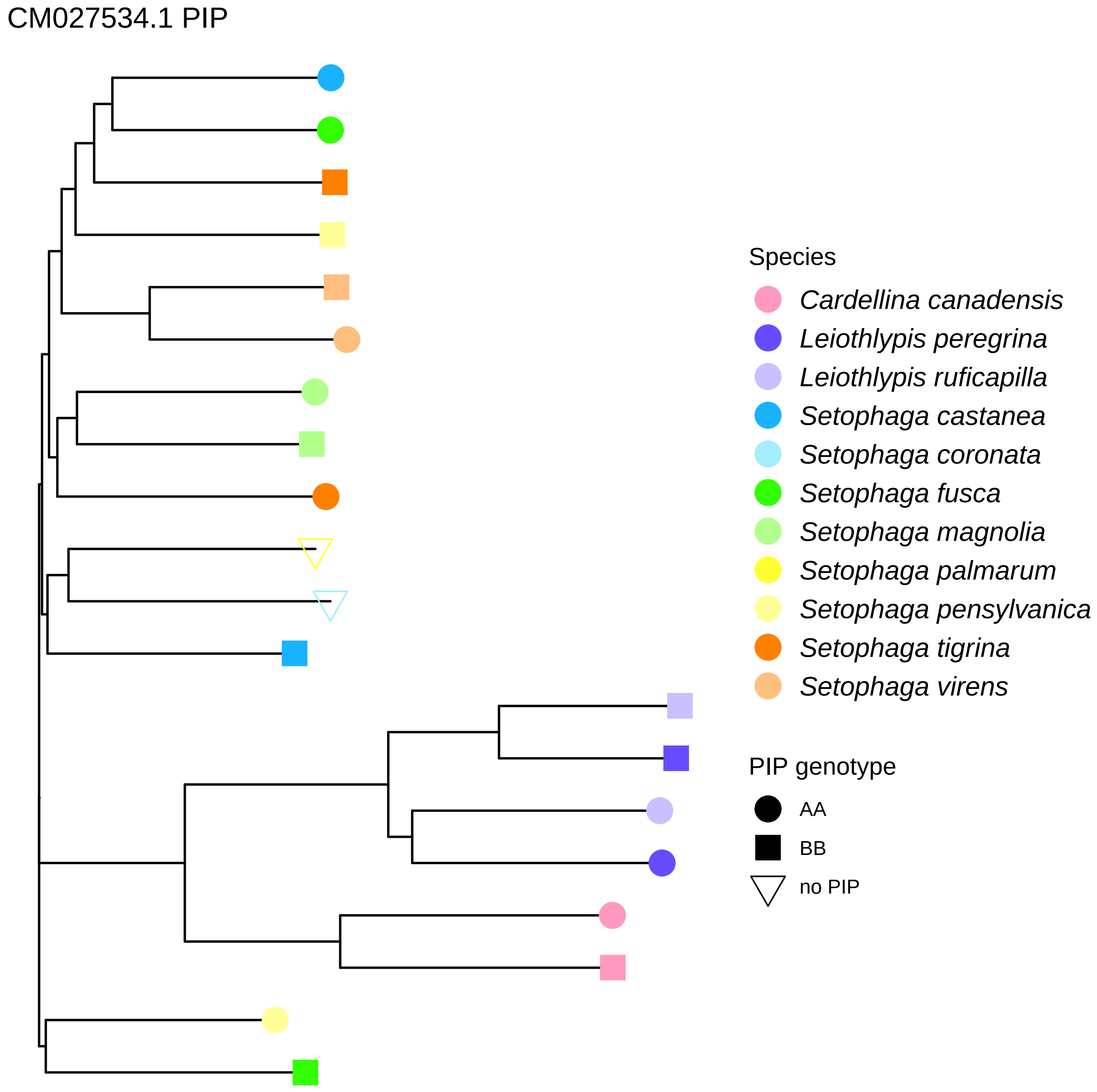
CM027534.1 PIP
Species
Cardellina canadensis
Leiothlypis peregrina
Leiothlypis ruficapilla
Setophaga castanea
Setophaga coronata
Setophaga fusca
Setophaga magnolia
Setophaga palmarum
Setophaga pensylvanica
Setophaga tigrina
Setophaga virens
PIP genotype
AA
BB
no PIP

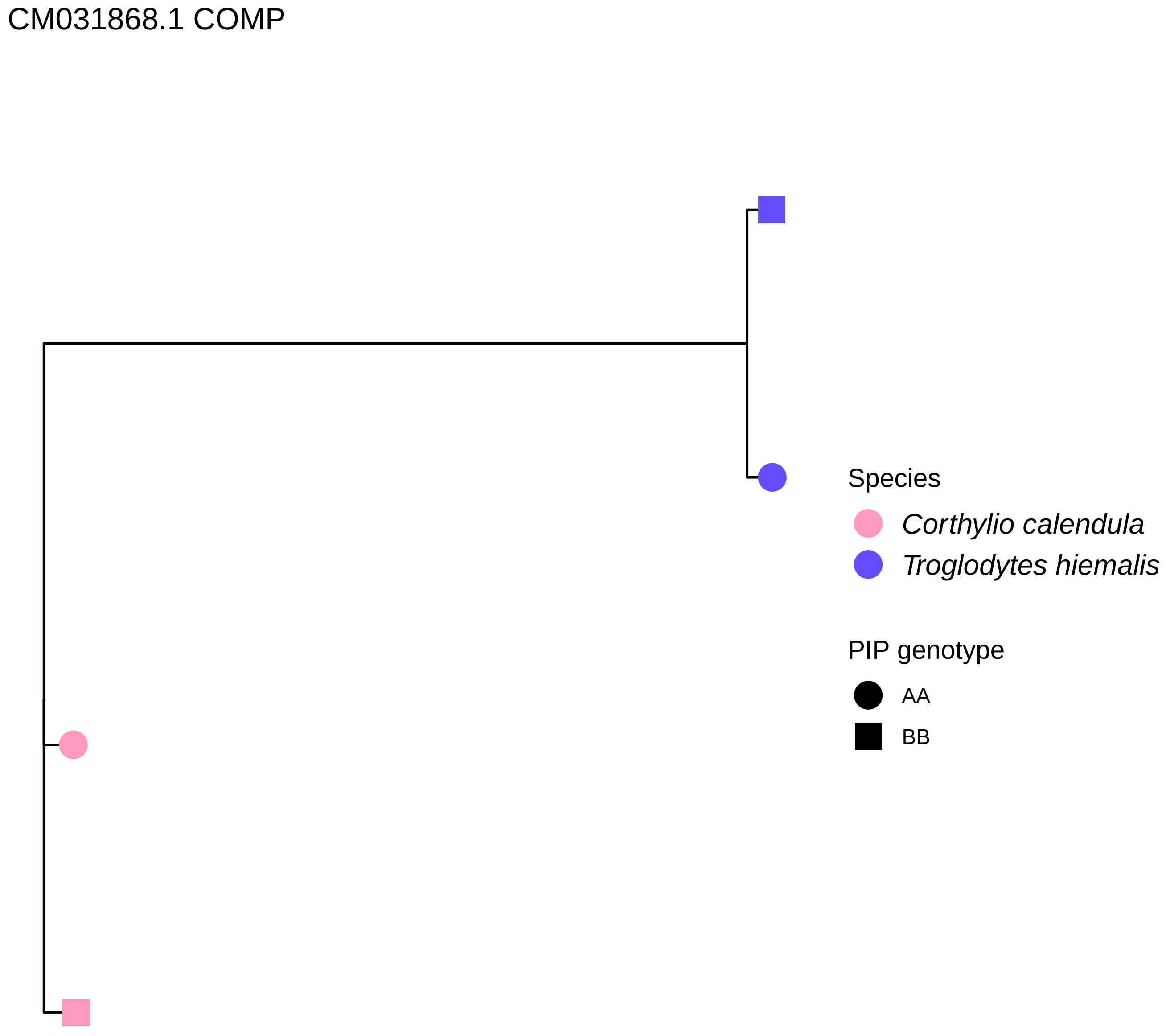
CM031868.1 COMP
Species
Corthylio calendula
Troglodytes hiemalis
PIP genotype
AA
BB

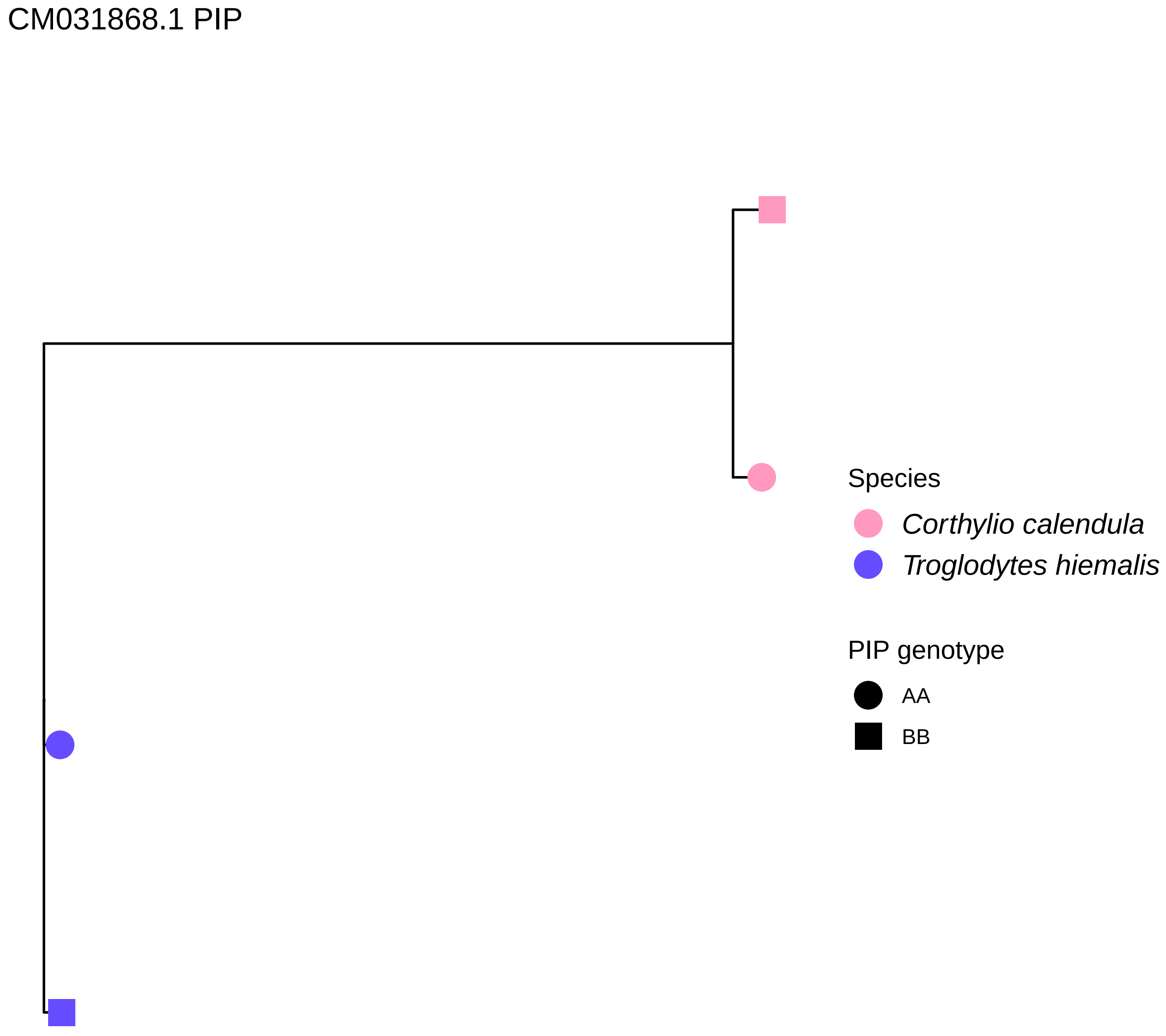
CM031868.1 PIP
Species
Corthylio calendula
Troglodytes hiemalis
PIP genotype
AA
BB

CM036369.1 COMP

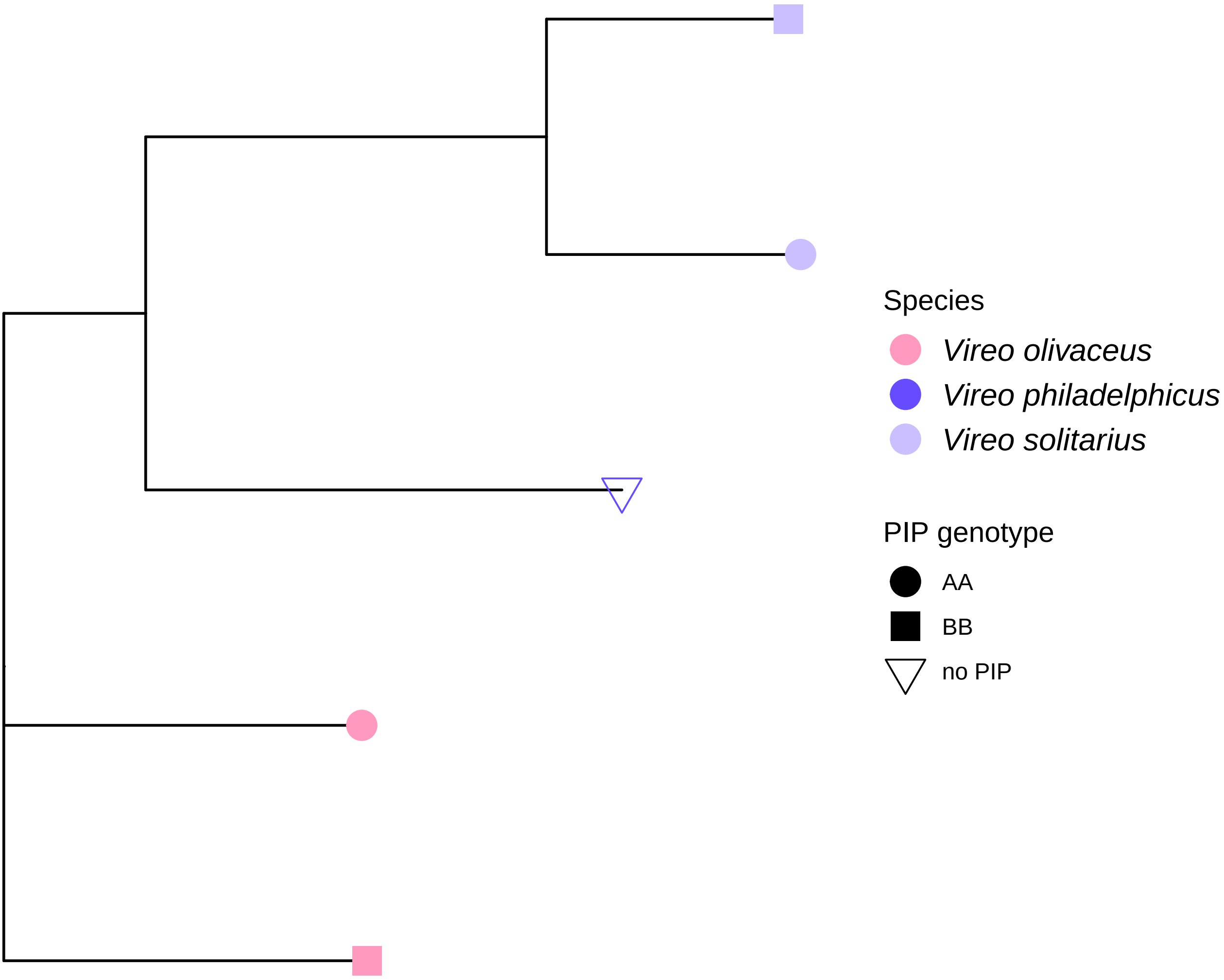

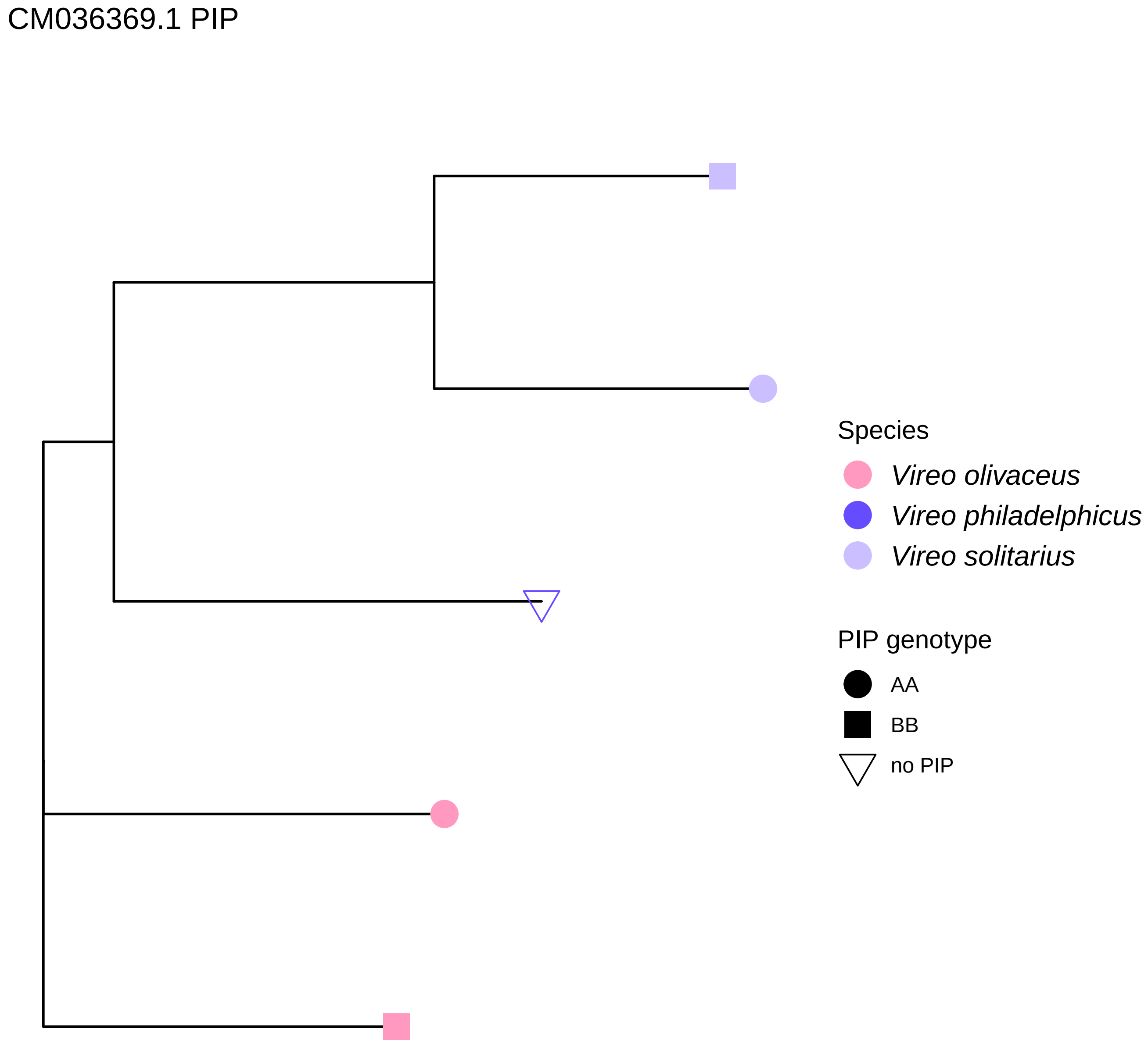
CM036369.1 PIP
Species
Vireo olivaceus
Vireo philadelphicus
Vireo solitarius
PIP genotype
AA
BB
no PIP

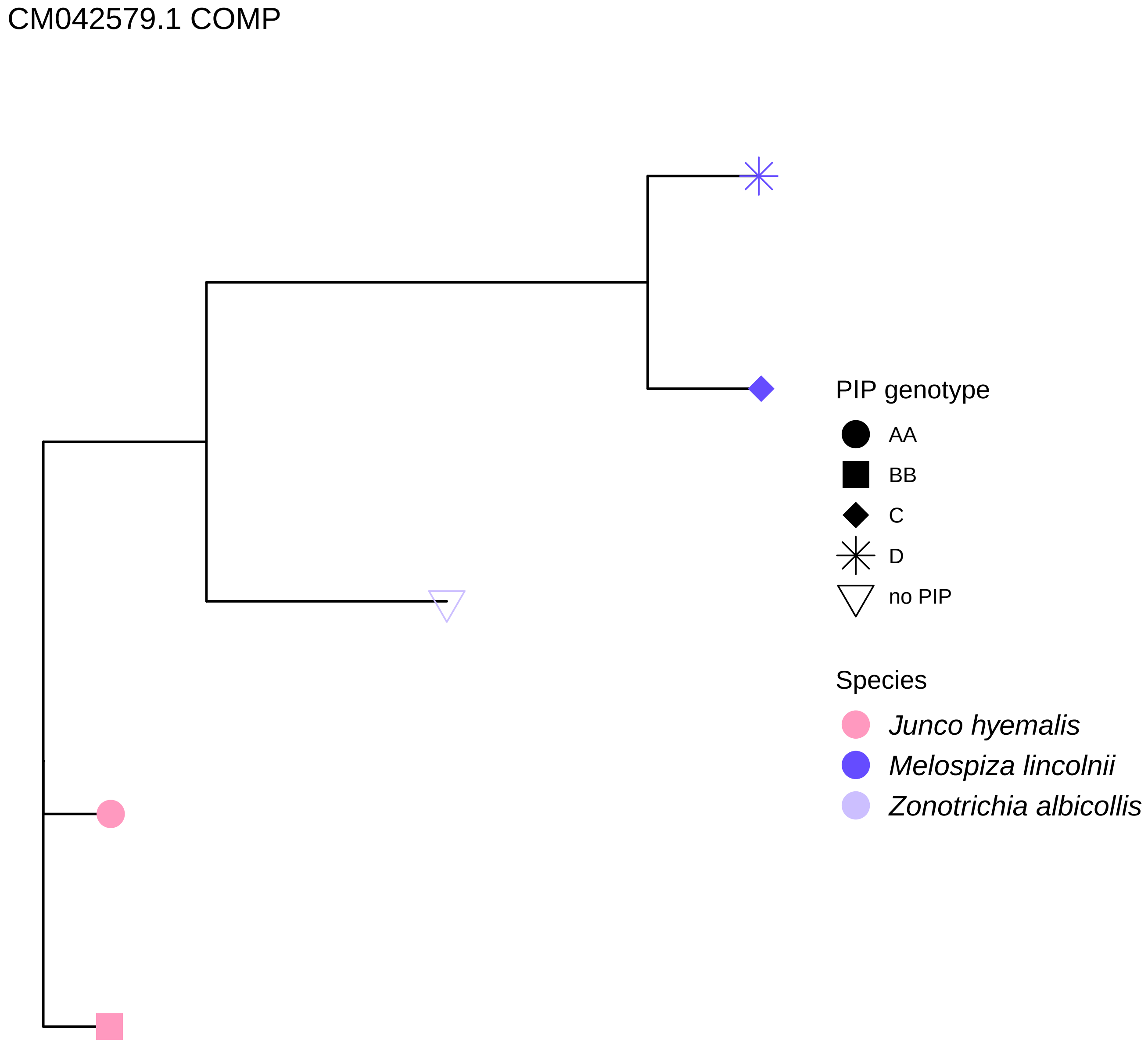
CM042579.1 COMP
PIP genotype
AA
BB
C
D
no PIP
Species
Junco hyemalis
Melospiza lincolnii
Zonotrichia albicollis

### CM042579.1 PIP

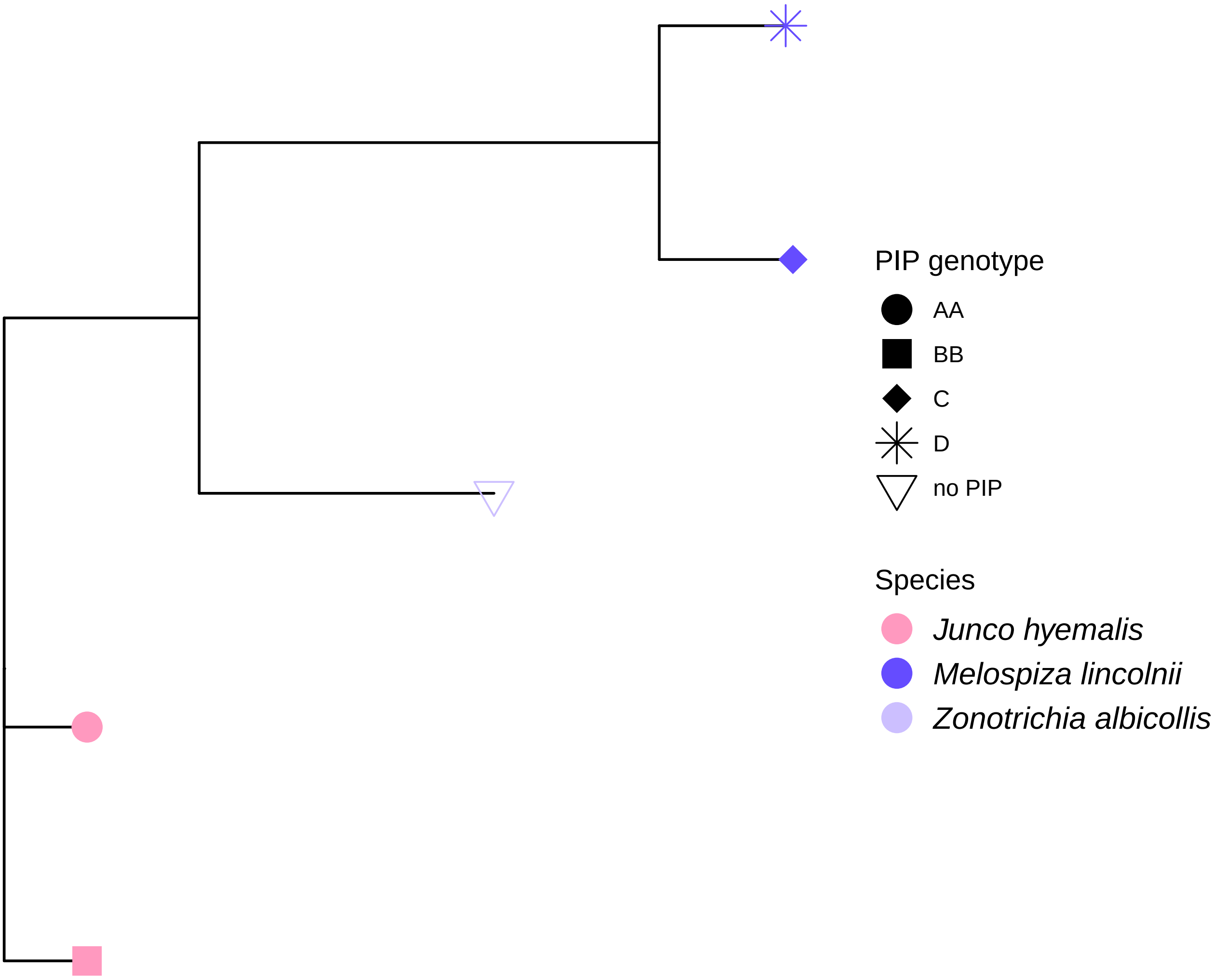

CM042598.1 COMP

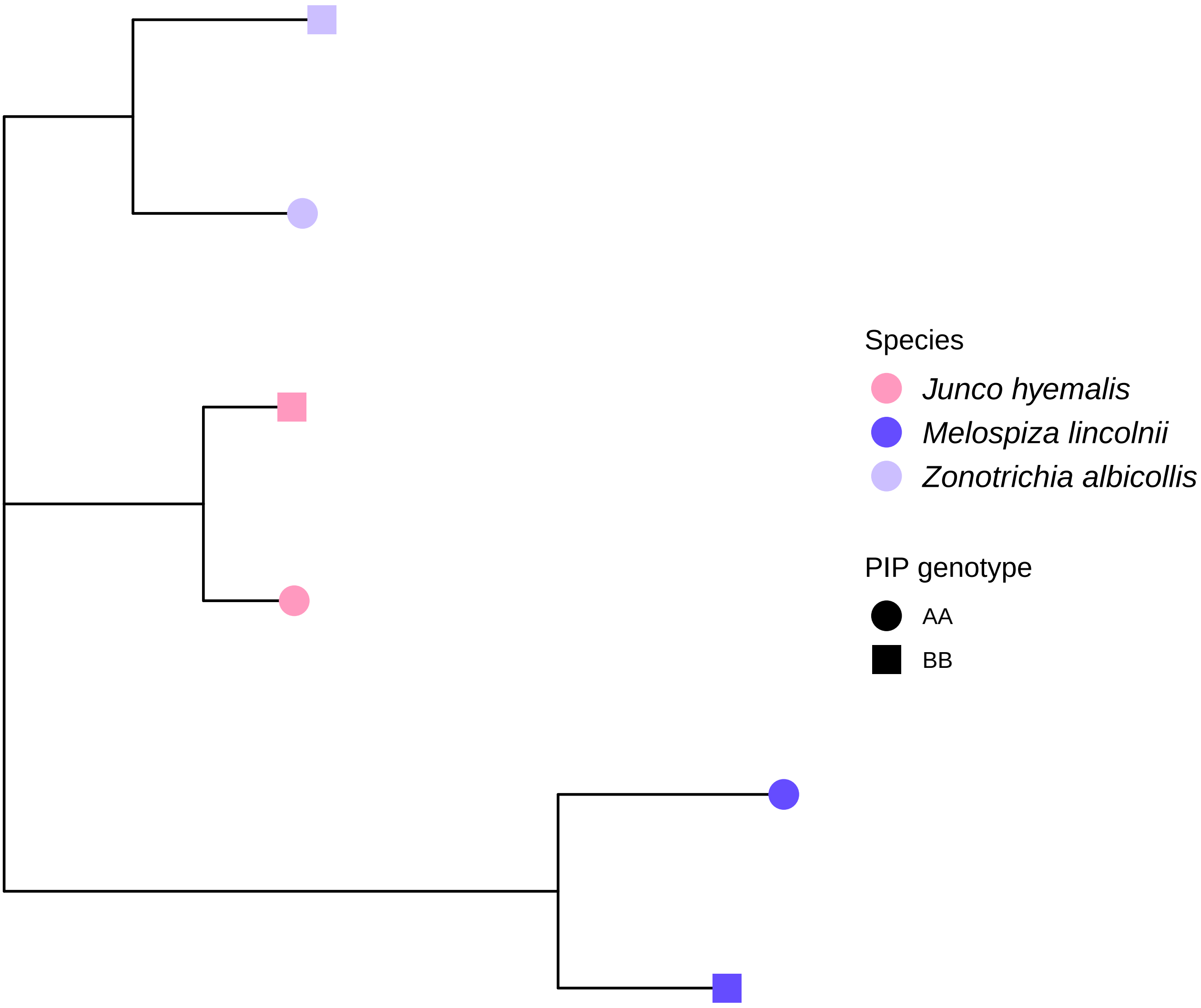

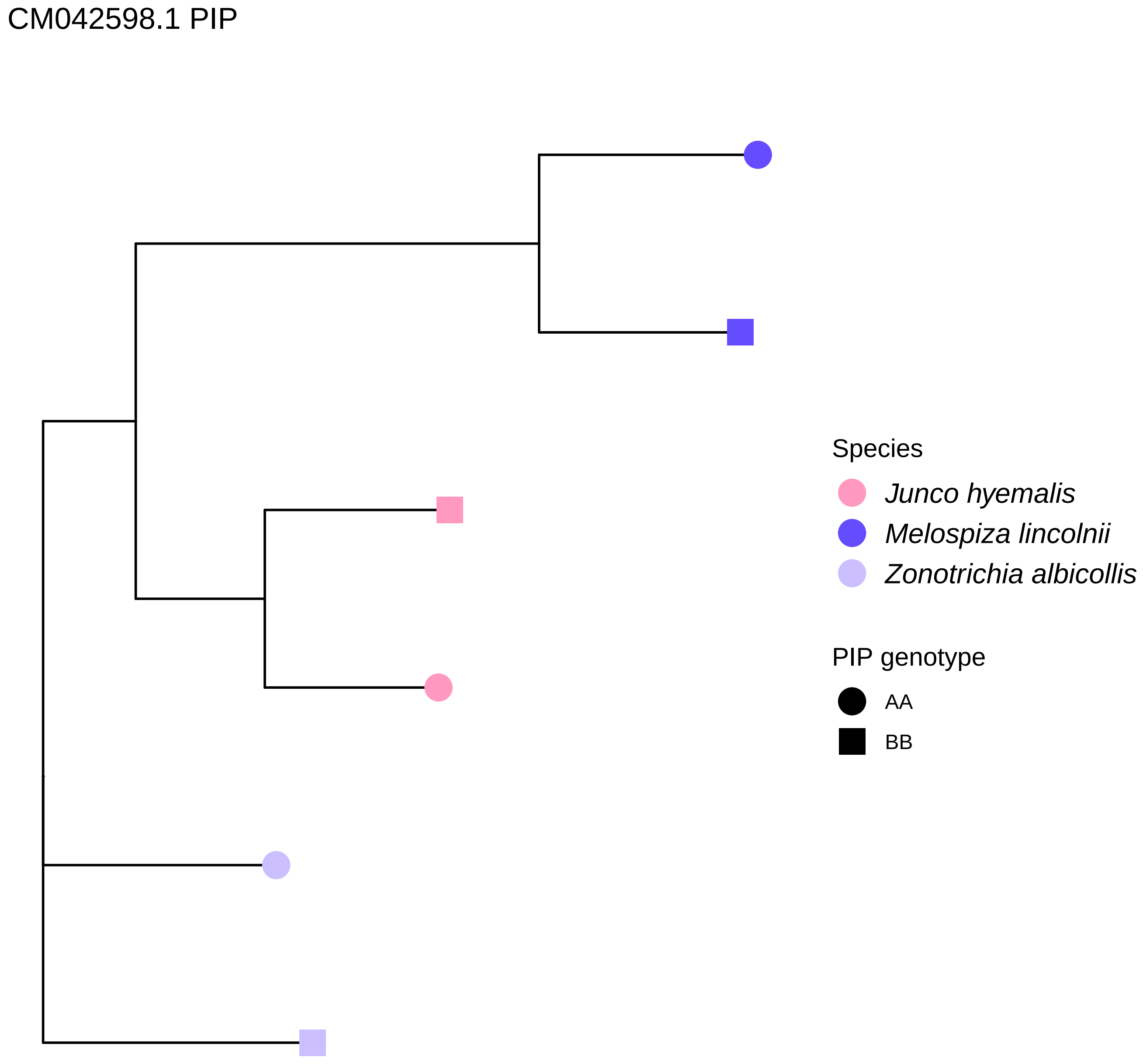
CM042598.1 PIP
Species
Junco hyemalis
Melospiza lincolnii
Zonotrichia albicollis
PIP genotype
AA
BB
