## Supplementary Information for "Large inversion polymorphisms are widespread in North American songbirds"

**This PDF file includes:**

Supporting text
Figures S1 to S5
Tables S1 to S2
Legends for Datasets S1 to S5
SI References

Other supporting materials for this manuscript include the following:

Datasets S1 to S5

### Supporting Information Text

*Genetic distance to the reference genome does not correlate with the number of PIPs discovered.*

Mapping reads to reference genomes can introduce biases that potentially affect our ability to detect inversion polymorphisms, and these biases may vary with the genetic distance between the reference genome species and the focal mapped species (Prasad et al. 2022; Thorburn et al. 2023). To facilitate discovery of possible inversions, we mapped our study species to chromosome-assembled reference genomes from the most closely related species available on GenBank at the time of analysis, but some species' closest chromosome-assembled relatives were in different genera or even different families than the study species. Therefore, we used linear models to assess the relationship across species between the number of putative inversion polymorphisms (PIPs) we discovered and a proxy for genetic distance between reference genome species and mapped species (hereafter, "reference D").

Our proxy for reference D is the genetic distance between the mitochondrial ND2 haplotype from the reference genome species compared with that of the mapped species. We used a mitochondrial proxy because it was possible for us to assemble accurate, reference-free high-coverage mitochondrial genes even from low-coverage whole genome sequencing data (Pegan et al. 2025; Dierckxsens et al. 2016). Further, mitochondrial genetic distances are sufficient to capture the broad range of evolutionary distances involved in our reference mapping, from conspecific reference genomes to reference genomes from a different taxonomic family than the mapped species. For our study species, we used one example of an ND2 haplotype from our dataset for each species, which we assembled as described in Pegan et al. 2024. We chose random samples from within the central portion of the boreal region (Michigan or Minnesota, USA, or Manitoba, Canada when no samples were available from the central USA). We downloaded a copy of ND2 from GenBank for each reference genome species not included in our study species. One exception is that no mitochondrial genes were available for the reference species *Chiroxiphia lanceolata*, to which we mapped our Empidonax study species, so we used a copy of ND2 from its congener *Chiroxiphia pareola*. We estimated the ND2 genetic distance between each study species and its reference genome species (i.e., reference D) using the function 'dist.dna()' in the R package 'ape'9. For four species that were mapped to conspecific reference genomes (*Certhia americana, Junco hyemalis, Setophaga coronata, Catharus ustulatus*), we assigned reference D to be 0.

We created a linear model in R evaluating the effect of the reference D predictor on the number of PIPs discovered (Table 1, main text). We scaled the D predictor to have a mean of 0 and a standard deviation of 1 prior to modeling. The linear model indicates that the relationship between D and number of PIPs discovered is not significant (effect = -0.79, std. err. = 0.67, p = 0.25; Fig. S5). Although the specific effects of reference genome choice on PIP discovery remain to be investigated in depth, our results demonstrate that it is possible to identify large numbers of PIPs (>10) even in species mapped to the reference genome of a relatively distantly related species (Fig. S5).

*Confirming the correspondence between PIPs and known inversions in Zonotrichia albicollis.*

Among our 35 study species, a discrete phenotypic polymorphism is only known in 1 species, the sparrow *Zonotrichia albicollis*, which displays a range-wide plumage polymorphism (white-striped vs tan-striped crown feathers) associated with known inversion polymorphisms (Tuttle et al. 2016). Given that the *Z. albicollis* plumage polymorphism is already known to be associated with inversions, we use this species as an opportunity to ensure that our methods could detect these inversion polymorphisms and that they conformed to expectations based on past studies.

At the time of analysis, a chromosome-assembled reference genome for *Zonotrichia albicollis* was not available, so we mapped this species' sequence data to the chromosome-assembled reference genome of *Junco hyemalis*. The inversion polymorphism called *2/2$^{m}$*, typically considered to lie on chromosome 2 of *Z. albicollis* (Tuttle et al. 2016; Thorneycroft 1975), instead mapped to chromosome 3 of the *J. hyemalis* reference (GenBank accession number CM042576.1). Similarly, the polymorphism called *3/3$^{a}$* by (Thorneycroft 1975) is presumably the large polymorphism in *Z. albicollis* that mapped to chromosome 1 (GenBank accession number CM042573.1) of the *J. hyemalis* reference. We confirmed the identity of *2/2$^{m}$* as mapped to CM042576.1 by comparing the genotypes and phenotypes of 22 individuals with voucher skin specimens at the University of Michigan Museum of Zoology (Table S1). We found that the color phenotype of specimens (tan-striped vs white-striped; (Thorneycroft 1975)) perfectly aligned with assignment to different PCA clusters on CM042576.1 (Table S1). This polymorphism produced only two clusters in PCA, consistent with the observation that individuals homozygous for the *2$^{m}$* "white" inversion haplotype are extremely rare (Thorneycroft 1975; Tuttle et al. 2016). The two clusters in our PCA therefore represent individuals homozygous for the "tan" haplotype (tan-striped individuals) and heterozygotes (white-striped individuals), and we did not observe any "super-white" individuals with a homozygous *2$^{m}$* genotype.

*Annotation of high-order repeats on reference chromosomes.*

High-order repeats (HORs) are repetitive regions that tend to be associated with centromeres (Qi et al. 2025). We annotated HORs onto our reference chromosomes to explore the relationship between centromeric repeats and PIPs and to see if we could infer whether PIPs were pericentric or paracentric.

Our ability to annotate HORs is significantly impacted by the quality of each reference genome and the methods by which it was created and assembled. Repetitive regions cause problems with genome assembly and even the most advanced assembly methods face challenges accurately assembling centromeres (Logsdon et al. 2024). The previously published reference genomes we used in this study were assembled using a variety of methods, including methods that are not suitable for accurate resolution of repetitive elements. As such, the results of this analysis cannot be interpreted as a systematic investigation into the locations of HORs and centromeres on each chromosome and instead represent a preliminary exploration.

To annotate HORs, we used the program centroAnno (Qi et al. 2025) with its default settings and we plotted the results on plots of chromosomes in Dataset S2 (A and B panels). We found that centroAnno was only able to annotate HORs in about 55% of chromosomes with PIPs, which may reflect issues with missing/collapsed satellites in many of our reference genomes. Notably, no HORs were annotated on any chromosome from reference genomes we used for *Setophaga* and Paridae species (GCA_001746935.2 and GCA_001522545.3, respectively). By contrast, centroAnno found particularly extensive HOR content in the chromosomes of GCA_020740725.1—this is the sole reference genome in our study that was created long-read sequencing and assembled with hifiasm (Cheng et al. 2021), which are methods well suited to avoid collapsing repetitive regions (Dataset S2 pages 28-33, on plots showing *Vireo* species). On chromosomes with HOR annotations, we were unable to consistently and confidently infer the locations of centromeres because HORs were frequently annotated in multiple locations along the chromosome. This pattern may reflect the fact that centromeric repeats can also exist outside of centromeres (Gozashti et al. 2025). Based on these limitations, we refrain from drawing conclusions about the status of PIPs as paracentric or pericentric.

When we examined the results of HOR annotation, we observed many instances where HORs were annotated close to PIP breakpoints (e.g., Dataset S2 pages 3, 4, 6, 15, 27, 30). The presence of HORs near PIP breakpoints is consistent with the observation that centromeric repeats can be moved around by inversions, and may even cause inversions to form (Gozashti et al. 2025). In other cases where HORs lie within or surrounding PIPs (e.g., Dataset S2 pages 3, 31). Some PIPs could represent polymorphic centromeres segregating in the population as two highly differentiated haplotypes that do not experience recombination (Irwin et al. 2025). These possibilities will require investigation in future studies based on long-read sequencing methods that can better resolve complex genomic regions like HORs.

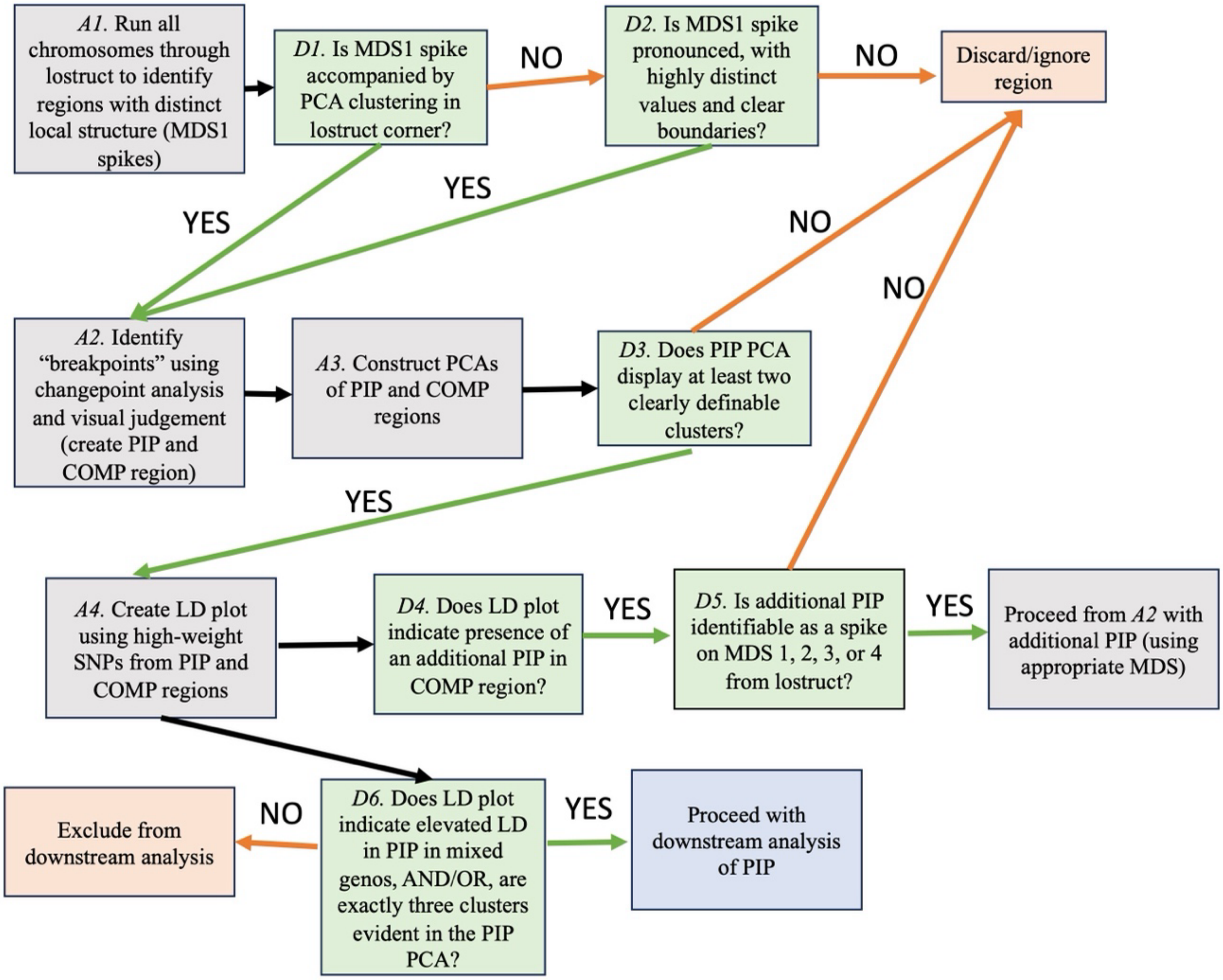


**Fig. S1**. A flow chart demonstrates how we identified putative inversion polymorphisms (PIPs) using a series of analyses (*A1-A4*) and decisions (*D1-D6*). For a detailed overview of each element in the flow chart, including definitions of terms and descriptions of the programs used for each analysis, see *Materials and Methods*. Dataset S2 shows plots for each PIP, including those corresponding to analyses *A1-A4*.

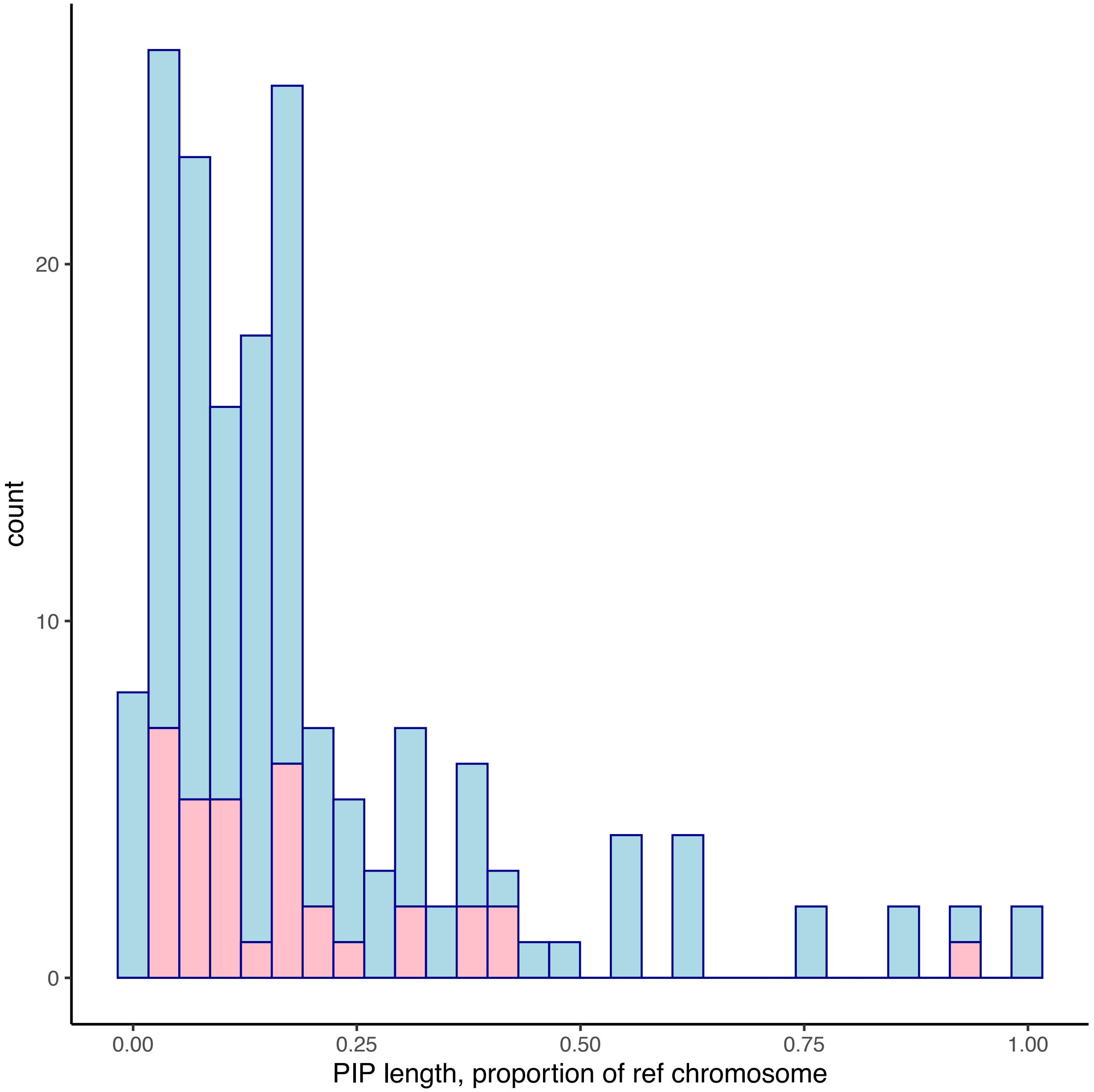


**Fig. S2.** The distribution of PIP lengths as a proportion of reference chromosome length. Many PIPs take up a substantial proportion of the chromosomes they are mapped to. In one microchromosome, CM027534.1 (from the GCA_001746935.2 *Setophaga coronata* reference genome), a 3-cluster PCA pattern was present across the entire chromosome in most of our parulid study species (see also Table 2 in the main text, Dataset S2). In this case the PIP length is approximately equal to the chromosome length (the bar at proportion = 1). It is not clear whether this pattern involves a large inversion whose actual breakpoints are undetectable with our methods, or if recombination on this microchromosome is suppressed in general for other reasons.

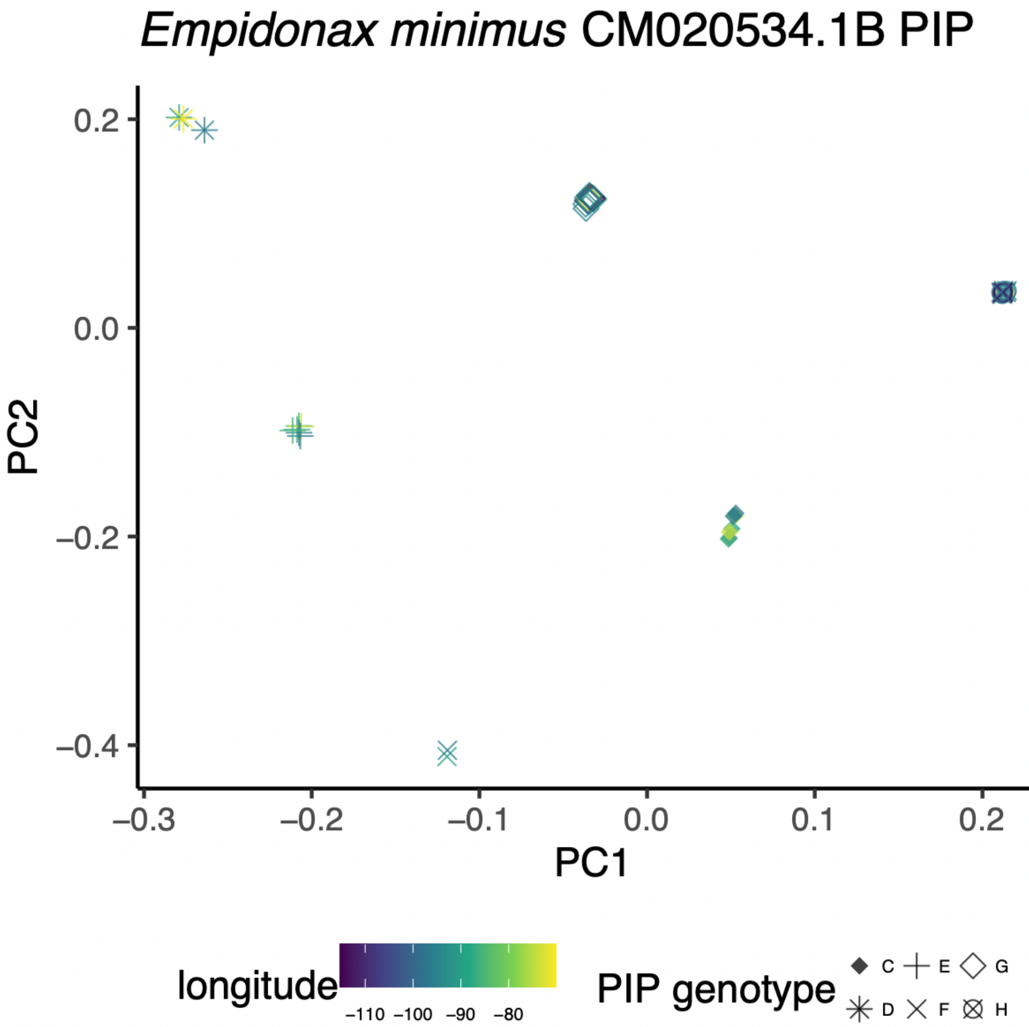


**Fig. S3.** A PCA of putative inversion polymorphism (PIP) in *Empidonax minimus* produces 6 clusters of individuals. The underlying polymorphism may have 3 alleles, corresponding to six possible genotypes, all of which are present in the population. Genotypes are labeled C though H arbitrarily. Points are colored by longitude of sampling, but no spatial patterns are evident in the PCA.

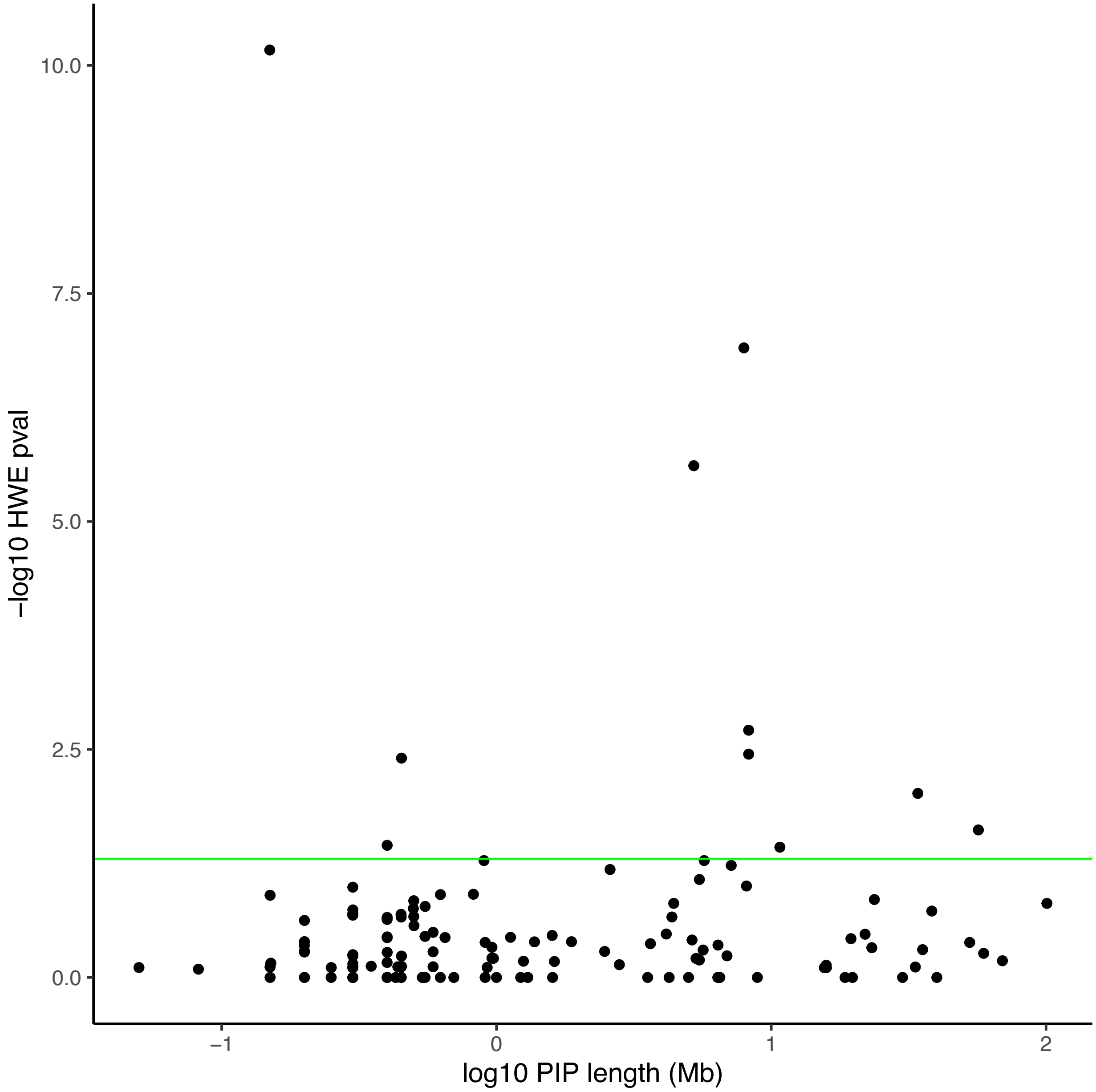


**Fig. S4.** The relationship between PIP length and the p-value of tests for deviation from Hardy-Weinberg Equilibrium (see also Table 2 in the main text). The green line is plotted at $p = 0.05$, the alpha-level for statistical significance. There is no clear tendency for large inversions to be more likely to deviate from HWE.

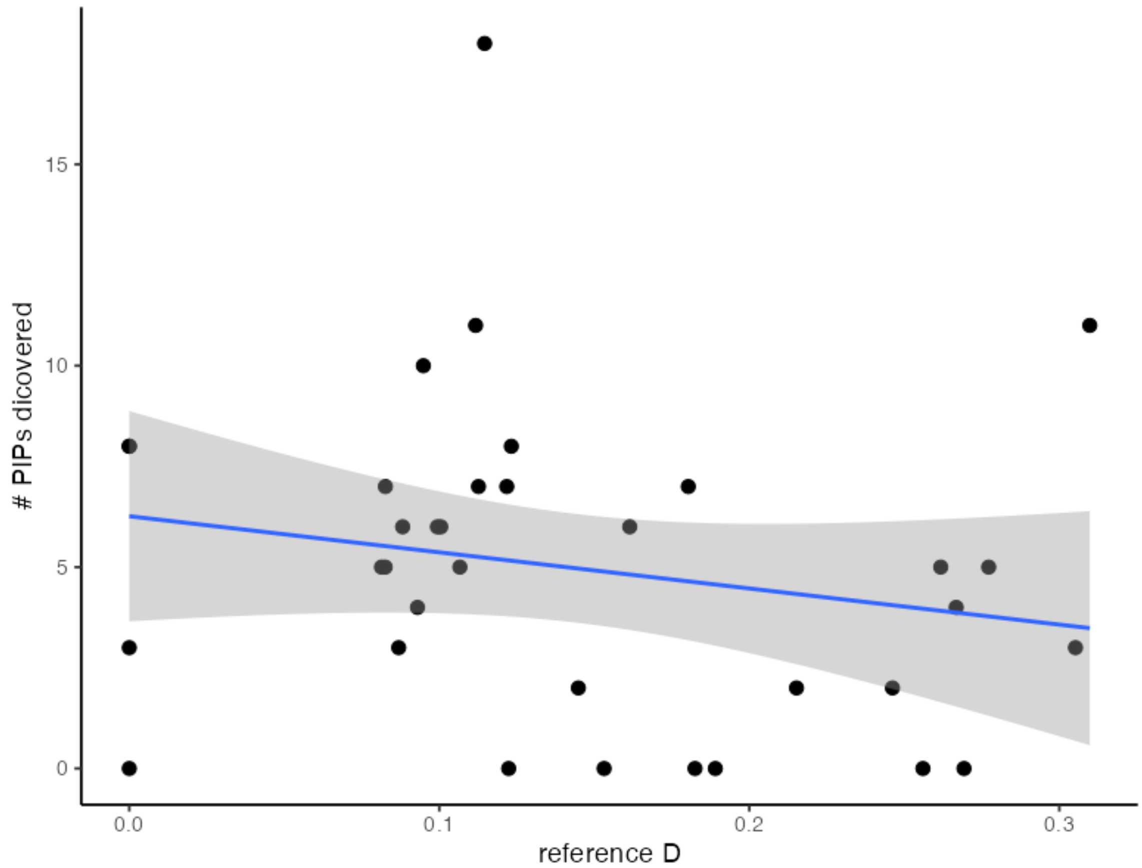


**Fig. S5**. We found putative inversion polymorphisms (PIPs) in species mapped to a wide range of reference genomes, including those mapped to reference genomes of relatively distantly related species (i.e., species with high “reference D”, representing genetic distance in the mitochondrial ND2 gene). The number of PIPs discovered does not show a significant relationship with reference D.

**Table S1**. Samples from *Zonotrichia albicollis* with voucher skin specimens at the University of Michigan Museum of Zoology (*n* = 22) demonstrate that the putative inversion polymorphism (PIP) we detected in *Z. albicollis* data mapped to the CM042576.1 reference chromosome matches phenotypic expectations of the well-studied *2/2^m^* inversion polymorphism in this species. We classified each skin specimen by its crown stripe color blindly (i.e., without consulting the PIP data), then compared its crown phenotype to its PCA-based genotype assignment for the CM042576.1 PIP. We found perfect correlation between crown phenotype and PIP genotype among these samples. The "C" and "D" genotype labels we assigned based on PCA clustering (see *Materials and Methods*) correspond to *2/2* homozygotes (tan-striped individuals) and $2/2^m$ heterozygotes (white-striped individuals) respectively (Tuttle et al. 2016).

| **Specimen number** | **Crown stripe phenotype** | **CM042576.1 PIP genotype** |
|---|---|---|
| UMMZ:Bird:248165 | Tan | C |
| UMMZ:Bird:244721 | Tan | C |
| UMMZ:Bird:244774 | Tan | C |
| UMMZ:Bird:247365 | Tan | C |
| UMMZ:Bird:245214 | Tan | C |
| UMMZ:Bird:245215 | Tan | C |
| UMMZ:Bird:245230 | Tan | C |
| UMMZ:Bird:245216 | Tan | C |
| UMMZ:Bird:244770 | Tan | C |
| UMMZ:Bird:244776 | White | D |
| UMMZ:Bird:247370 | White | D |
| UMMZ:Bird:247364 | White | D |
| UMMZ:Bird:247372 | White | D |
| UMMZ:Bird:247366 | White | D |
| UMMZ:Bird:247367 | White | D |
| UMMZ:Bird:245225 | White | D |
| UMMZ:Bird:245231 | White | D |
| UMMZ:Bird:245250 | White | D |
| UMMZ:Bird:245232 | White | D |
| UMMZ:Bird:248161 | White | D |
| UMMZ:Bird:248164 | White | D |
| UMMZ:Bird:245355 | White | D |

**Table S2**. Putative inversion polymorphisms (PIPs) found to be overlapping on the same reference chromosome in multiple species, which we used to create the genetic distance based trees shown in Dataset S3. The second column lists the species mapped to the relevant chromosome which had overlapping PIPs identified on that chromosome. The breadth of the "shared PIP" (spanned by values shown in shared PIP start and end columns) represents the largest possible region that fully overlaps all species' PIPs, and therefore does not exactly match that of the individual PIPs identified in each overlapping species. Similarly, the "Shared COMP" region was chosen based on all species mapped to the relevant chromosome and exactly matches the size of the shared PIP and therefore is different from COMP regions chosen for each species individually. Note that in some cases, we were unable to identify a suitable shared COMP region from the same chromosome as a shared PIP due to the size and/or complexity of the PIP. The final column indicates whether each tree shows evidence of trans-specific clustering, i.e. pairs of conspecifics that cluster according to PIP genotype instead of species (Dataset S3). In one case (CM027512.1), the COMP and PIP region share an unexpected clustering pattern.

| **Shared PIP name** | **Species with PIP** | **Shared PIP start** | **Shared PIP end** | **Shared COMP start** | **Shared COMP end** | **Trans-specific clustering?** |
|---|---|---|---|---|---|---|
| CM019929.1 | *Oporornis agilis, Geothlypis philadelphia* | 1375000 | 1774820 | 500000 | 899820 | T |
| CM020365.1 | *Catharus fuscescens, Catharus guttatus* | 1275000 | 1575156 | 3000000 | 3300156 | F |
| CM020367.1_i | *Catharus fuscescens, Catharus guttatus, Catharus ustulatus* | 275044 | 724997 | 1000000 | 1449953 | T |
| CM020367.1_ii | *Catharus fuscescens, Catharus guttatus, Catharus ustulatus* | 1975032 | 2275004 | NA | NA | T |
| CM020368.1 | *Catharus fuscescens, Catharus guttatus, Catharus ustulatus* | 1375003 | 2284362 | 90641 | 1000000 | T |
| CM020371.1_i | *Catharus fuscescens, Catharus guttatus, Catharus ustulatus* | 275003 | 374995 | NA | NA | T |
| CM020371.1_ii | *Catharus fuscescens, Catharus guttatus, Catharus ustulatus* | 725199 | 924952 | NA | NA | T |
| CM020371.1_iii | *Catharus guttatus, Catharus ustulatus* | 1374999 | 1424991 | NA | NA | T |
| CM020534.1_i | *Empidonax flaviventris, Empidonax minimus* | 35275020 | 70824606 | 1 | 25000000 | F |
| CM020534.1_ii | *Empidonax alnorum, Empidonax flaviventris, Empidonax minimus* | 82025000 | 120403895 | NA | NA | T |
| CM020543.1 | *Empidonax flaviventris, Empidonax minimus* | 18124367 | 22063292 | 9000000 | 12938925 | F |
| CM027508.1_i | *Setophaga castanea, Setophaga fusca, Setophaga magnolia, Setophaga virens* | 43175065 | 48124955 | 100000000 | 104949890 | F |
| CM027508.1_ii | *Setophaga coronata, Setophaga magnolia, Setophaga pensylvanica Setophaga virens* | 48124954 | 49624979 | NA | NA | F |

| | | | | | | |
|---|---|---|---|---|---|---|
| CM027510.1 | *Setophaga coronata, Setophaga fusca* | 5324999 | 12174998 | 40000000 | 46849999 | F |
| CM027511.1_i | *Cardellina canadensis, Leiothlypis peregrina, Setophaga castanea* | 4124873 | 23224972 | NA | NA | F |
| CM027511.1_ii | *Cardellina canadensis, Leiothlypis peregrina, Setophaga castanea* | 26074951 | 51375065 | NA | NA | F |
| CM027512.1 | *Leiothlypis peregrina, Leiothlypis ruficapilla, Setophaga castanea* | 2724053 | 3225008 | 30000000 | 30500955 | NA |
| CM027513.1 | *Leiothlypis ruficapilla, Setophaga castanea, Setophaga pensylvanica* | 35674998 | 38264695 | 5000000 | 7589697 | T |
| CM027518.1 | *Leiothlypis ruficapilla, Setophaga castanea* | 17324986 | 18024642 | 5000000 | 5699656 | F |
| CM027530.1 | All 11 species aligned to this chromosome (see Dataset S1) | 574805 | 874807 | 1300000 | 1600002 | T |
| CM027532.1 | All 11 species aligned to this chromosome (see Dataset S1) | 2374079 | 2774194 | 1500000 | 1900115 | T |
| CM027534.1 | All species aligned to this chromosome except *Setophaga pensylvanica, Setophaga coronata* (see Dataset S1) | 525259 | 775051 | NA | NA | T |
| CM031868.1 | *Corthylio calendula, Troglodytes hiemalis* | 1 | 5324946 | 10000000 | 15324945 | F |
| CM036369.1 | *Vireo olivaceus, Vireo solitarius* | 1 | 925008 | 6000000 | 6925007 | F |
| CM042579.1 | *Junco hyemalis, Melospiza lincolnii* | 6425278 | 26175546 | 40000000 | 59750268 | F |
| CM042598.1 | *Junco hyemalis, Melospiza lincolnii, Zonotrichia albicollis* | 774997 | 1174898 | 2000000 | 2399901 | F |

**Dataset S1** (separate file). Data from each of the putative inversion polymorphisms (PIPs) we identified, including the species, the GenBank accession number of the reference genome we mapped the species' data to, the species of the reference genome, and the GenBank accession number of the relevant chromosome. The name of each PIP is usually the reference chromosome accession number, except in cases where more than one PIP was detected on a chromosome, in which case a subscript (A, B, or C) is added to the accession number. The column "PIP unique status" indicates whether a PIP was identified on the corresponding reference chromosome in more than one species or not. The PIP start and PIP end columns provide the approximate breakpoints of the PIP as mapped to the reference chromosome (see Methods), which we used to estimate PIP length. "LD supported" indicates whether the PIP's status as an inversion was corroborated by linkage disequilibrium analysis or not (see *Materials and Methods*). "PCA cluster number" indicates how many clusters the PIP produces in PCA, which is assumed to correspond to how many genotypes are present in the population. "IBD slope" gives a slope of isolation by distance for the PIP, indicating whether there is spatial structure in the distribution of the PIP haplotypes (see *Materials and Methods*). IBD slope is only estimated for PIPs with exactly 3 PCA clusters (i.e. those represented by exactly 3 genotypes, as expected for biallelic variants with all possible genotypes present in the population). "comp_break_right" and "comp_break_left" columns indicate where comparison ("COMP") regions are located on chromosomes—COMP regions are designated as all loci to the right of "comp_break_right" and/or all points to the left of "comp_break_left." Some  Plots associated with each PIP can be viewed in Dataset S2.

**Dataset S2** (separate file). For each putative inversion polymorphism (PIP), we show 8 plots generated by the analyses we used to identify PIPs (Fig. S1) and to evaluate some of their characteristics. The species and name of the PIP is given at the top of each 8-plot page. **A.** Analysis with lostruct (Fig. S1, *A1*) indicates which segments of chromosomes have distinct population structure, as identified by multidimensional scaling (MDS) (Li & Ralph 2019). The highlighted red region is identified as a PIP (Fig. S1, *A2*). Panel A shows MDS1 values along each chromosome. **B.** MDS2 values from lostruct along each chromosome. In some cases, PIPs may be highlighted on MDS2 and not MDS1, particularly when more than one PIP is present on a chromosome. **C.** PCA plots from each PIP region (Fig. S1, *A3)* show distinct clustering patterns consistent with inversion (Mérot 2020). Genotypes are assigned to individuals based on these clusters. In cases where PCA produces 3 clusters, individuals in the central cluster are assumed to be heterozygotes for the polymorphism (AB genotype), while those in the right and left clusters are assumed to be homozygotes (AA and BB genotypes; (Mérot 2020)). Otherwise, individuals are assigned arbitrary genotype codes (C-H, depending on the number of clusters). **D.** The cluster pattern is not present in the comparison (COMP) region of the chromosome, where no inversion polymorphisms are thought to be present. **E.** Boxplots show individual heterozygosity values from inside the PIP region and inside a comparison (COMP) region in the same chromosome. **F.** A waffle plot shows PIP genotype frequencies of the sampled population. Genotypes are assigned based on PCA (see panel C). **G.** Linkage disequilibrium (LD) is plotted from a subsample of SNPs in the PIP and COMP regions (Fig. S1, *A4*). Inversions are expected to produce a pattern where LD in a sample of individuals with both classes of homozygous genotypes (upper triangle) is elevated in the PIP region compared with the COMP region (Mérot 2020). This pattern should not exist in sample of individuals that all share the same genotype (lower triangle). PIPs that meet these expectations are considered "LD-supported" PIPs. **H.** Individuals are shown in geographic space, colored by their PIP genotype, illustrating the spatial distributions of the genotypes. Individual locations are jittered to allow all points to be seen across the range (actual sampling locations can be seen in Fig. 1A). Note that in some cases, we were unable to identify a suitable COMP region from the same chromosome as a PIP due to the size and/or complexity of the PIP, and COMP data are therefore not shown in panels D and E in these cases. Chromosomes that have more than one PIP on them have only a single COMP region (if any).

**Dataset S3** (separate file). In 26 cases, PIPs were identified in multiple species that were mapped to the same reference genome. Here we provide neighbor-joining genetic distance trees for each shared PIP, along with trees from comparison (COMP) regions of the same chromosome. Note that it was not always possible to identify an appropriate COMP region, and that PIPs from the same chromosome share a single COMP region. Each tree includes all species mapped to a given reference chromosome, including one randomly-chosen representative of each identified PIP genotype for each species. We excluded putative heterozygotes (genotype=AB) from this analysis. In cases where species did not have an identified PIP on the relevant chromosome, we chose one individual randomly. The shape at each tip on the tree indicates the PIP genotype of that individual, while color represents species. In some cases, branching patterns show individuals clustering by inversion genotype rather than by species, suggesting the existence of trans-specific polymorphisms. Note that AA and BB genotypes are named arbitrarily based on haplotype frequencies within their respective species, not homology—i.e., in cases of trans-specific polymorphisms, the A haplotype of one species may be apparently homologous with the B haplotype of another species.

**Dataset S4** (separate file). Individual-level data for each putative inversion polymorphism (PIP) we identified, including metadata about the sample (catalog number and the institution where the sample came from, latitude and longitude of the sampling locality), the species and PIP name, the genotype assigned to the individual for the corresponding PIP, and the heterozygosity of the individual within the PIP region and an associated comparison (COMP) region from the same chromosome. The NCBI SRA accession number for each individual is provided. Plots associated with each PIP can be viewed in Dataset S2. We do not include metadata from samples of the 7 species for which no PIPs were found (*Picoides arcticus, Dryobates villosus, Perisoreus canadensis, Vireo philadelphicus, Poecile hudsonicus, Regulus satrapa, Certhia americana*); metadata from the samples we examined in these species can be found in the supplementary materials of (Pegan et al. 2025).

Institution abbreviations:
AMNH = American Museum of Natural History
CMNH = Cleveland Museum of Natural History
CUMV = Cornell University Museum of Vertebrates
MMNH = Bell Museum of Natural History
MVZ = UC Berkeley Museum of Vertebrate Zoology
NYSM = New York State Museum
RAM = Royal Alberta Museum
ROM = Royal Ontario Museum
UAM = University of Alaska Museum
UMMZ = University of Michigan Museum of Zoology

**Dataset S5** (separate file). Individual-level data from samples used to create genetic distance based trees of shared PIPs. Samples are listed with their species, the name of the shared PIP used in the analyses, the name of the species-level PIP that overlapped the shared region (i.e. the PIP name used in most other parts of the study, including Dataset S1-S3), and the genotype assigned to the individual at that PIP. Further metadata for each individual can be found in Dataset S2, with one exception: a sample from the species *Vireo philadelphicus* is included here because we used it to make a tree for an overlapping PIP shared by other Vireo species, but no PIPs were identified in *Vireo philadelphicus*. Further metadata for that sample is available in (Pegan et al. 2025). The NCBI SRA accession number for each individual is provided.

### Supporting Information References

Cheng, H., G. T. Concepcion, X. Feng, H. Zhang, and H. Li. 2021. Haplotype-resolved de novo assembly using phased assembly graphs with hifiasm. Nat. Methods 18:170–175. Springer US.

Dierckxsens N, Mardulyn P, Smits G. 2016. NOVOPlasty : de novo assembly of organelle genomes from whole genome data. Nucleic Acids Res. 45:10.1093/nar/gkw955. doi: 10.1093/nar/gkw955.

Gozashti, L., O. S. Harringmeyer, L. Gozashti, O. S. Harringmeyer, and H. E. Hoekstra. 2025. How repeats rearrange chromosomes: The molecular basis of chromosomal inversions in deer mice. Cell Rep. 44:115644.

Irwin, D., S. Bensch, C. Charlebois, G. David, A. Geraldes, S. Kumar, G. Bettina, H. Paul, H. Jessica, H. I. Vladimir, and V. I. Irina. 2025. The distribution and dispersal of large haploblocks in a superspecies. Mol. Ecol. 0:17731.

Li H, Ralph P. 2019. Local PCA shows how the effect of population structure differs along the genome. Genetics. 211:289–304. doi: 10.1534/genetics.118.301747.

Logsdon, G. A., A. N. Rozanski, F. Ryabov, T. Potapova, V. A. Shepelev, C. R. Catacchio, D. Porubsky, Y. Mao, D. Yoo, M. Rautiainen, S. Koren, S. Nurk, J. K. Lucas, and K. Hoekzema. 2024. The variation and evolution of complete human centromeres. Nature 629:136–145. Springer US.

Mérot C. 2020. Making the most of population genomic data to understand the importance of chromosomal inversions for adaptation and speciation. Mol Ecol. 29:2513–2516. doi: 10.1111/mec.15500.

Pegan, T. M., Berv, J. S., Gulson-Castillo, E. R., Kimmitt, A. A. & Winger, B. M. The pace of mitochondrial molecular evolution varies with seasonal migration distance. Evolution (N Y) 78, 160–173 (2024).

Pegan, T. M., A. A. Kimmitt, B. W. Benz, B. C. Weeks, Y. Aubry, T. M. Burg, J. Hudon, A. W. Jones, J. J. Kirchman, K. C. Ruegg, and B. M. Winger. 2025. Long-distance seasonal migration to the tropics promotes genetic diversity but not gene flow in boreal birds. Nat. Ecol. Evol. 9:957–969. Springer US.

Prasad A, Lorenzen ED, Westbury M V. 2022. Evaluating the role of reference-genome phylogenetic distance on evolutionary inference. Mol Ecol Resour. 22:45–55. doi: 10.1111/1755-0998.13457.

Qi, J., J. Ma, R. Han, Z. Han, T. Yu, and G. Li. 2025. De novo annotation of centromere with centroAnno. bioRxiv.

Thorburn DMJ et al. 2023. Origin matters: Using a local reference genome improves measures in population genomics. Mol Ecol Resour. 23:1706–1723. doi: 10.1111/1755-0998.13838.

Thorneycroft HB. 1975. A cytogenetic study of the White-throated Sparrow, *Zonotrichia albicollis* (Gmelin). Evolution (N Y). 29:611–621.

Tuttle EM et al. 2016. Divergence and functional degradation of a sex chromosome-like supergene. Current Biology. 26:344–350. doi: 10.1016/j.cub.2015.11.069.
